# Retinotopic organization of feedback projections in primate early visual cortex: implications for active vision

**DOI:** 10.1101/2022.04.27.489651

**Authors:** Mingli Wang, Yujie Hou, Loïc Magrou, Joonas A. Autio, Pierre Misery, Tim Coalson, Erin Reid, Yuanfang Xu, Camille Lamy, Arnauld Falchier, Qi Zhang, Mu-Ming Poo, Colette Dehay, Matthew F. Glasser, Takuya Hayashi, Kenneth Knoblauch, David Van Essen, Zhiming Shen, Henry Kennedy

## Abstract

Feedback connections play a major role in many theories of brain function. Previous studies of feedback connections to early visual areas have mainly concentrated on the representation of central visual fields. Here, injections of tracers at different eccentricities in areas V1 and V2 revealed retinotopically organized feedback. Peripheral injections revealed projections from 15 areas that are not labeled by central injections. Connection strengths for the majority of projections vary significantly with eccentricity in a systematic fashion with respect to distance and origin; whereas projections to central and upper visual fields are significantly stronger from ventral stream areas, peripheral and lower field projections are stronger from the dorsal stream. Non-invasive functional connectivity suggests a similar anatomical organization in humans. These features are discussed with respect to the cognitive and perceptual roles of these feedback pathways.

## Introduction

The macaque visual cortex is an intensively studied system for investigating information processing in the cerebral cortex. The primary visual cortex (area 17 or V1) receives the major sensory input from the retina via the thalamus. Projections from area V1 target higher-order areas in the cortical hierarchy, and these ascending, feedforward (FF) pathways are typically reciprocated by feedback (FB) projections. FF and FB pathways are distinguished by their laminar origins and terminations (Rockland and Pandya, 1979) and collectively form a cortical hierarchy that mediate distributed processing across the connectome (Felleman and Van Essen, 1991). Early models of cortical function were largely based on FF connectivity, which underpin the elaboration of receptive field properties presumed to reflect the extraction of information from the ascending flow of activity (Hubel and Wiesel, 1962; Zeki, 1978). However, in the last 20 years top-down FB pathways, which are twice as numerous as the FF pathways (Markov et al., 2014b), have become particularly intriguing to neuroscientists (Bullier, 2006). While there is evidence that FB pathways play a modulatory role (Klink et al., 2017), these pathways are increasingly considered to play an important role in assigning meaning to the inherent uncertainty of the sensory input to the brain (Gregory, 1997), formalized as Bayesian inferences of the causes of sensation (Lee and Mumford, 2003). This has led to predictive processing models, where the brain is hypothesized to use sensory inputs to continuously update a model of the world whose representation is distributed across the cortical hierarchy. According to predictive processing, prior information at individual levels of the cortical hierarchy generates descending predictions that interact with earlier levels to generate prediction errors that ascend the hierarchy and are in turn updated with new descending predictions, etc. until an interpretation is assigned to the input (Clark, 2013; Friston, 2010; Keller and Mrsic-Flogel, 2018; Markov and Kennedy, 2013; Rao and Ballard, 1999).

Multisensory integration has been well documented in higher-order, association areas across parietal, temporal and frontal regions (Macaluso and Driver, 2005; van Atteveldt et al., 2014). The use of anatomical tracers with increased sensitivity has led to a large increase in the observed connectivity of cortical areas in rodents (Gamanut et al., 2018) and non-human primates (Markov et al., 2014b; Theodoni et al., 2021). Increases in the estimated density and node degree of the cortical inter-areal connectome suggest greater likelihood of individual cortical areas being responsive to multiple sensory modalities. Indeed, recently interest has turned to multisensory integration in primary areas. Tracer injections in macaque area V1 revealed eccentricity-dependent projections from auditory areas preferentially targeting cortex representing the peripheral visual field (Falchier et al., 2002) subsequently confirmed in marmoset (Majka et al., 2019). These findings have spurred a reappraisal of multisensory integration in lower-level cortices, for review see (Ghazanfar and Schroeder, 2006; van Atteveldt et al., 2014). In addition to elucidating the perceptual and cognitive roles of primary areas, multisensory integration in primary areas is particularly interesting for investigating predictive processes. In human cortex, natural sounds can be decoded from complex fMRI activity in early visual cortex (Vetter et al., 2014), and with a higher accuracy in peripheral compared to central representations (Vetter et al., 2020) in conformity with the underlying anatomy (Eckert et al., 2008). Much of the work in this domain is increasing our understanding of hierarchical processing. For instance, a recent study of the circuit mechanism that enables long-range cortical cross modal phenomena shows that auditory inputs to visual cortex can suppress responses to predictable input and is therefore highly relevant to predictive processing theory (Garner and Keller, 2022).

How does eccentricity-dependent multisensory integration in macaque area V1 relate to the known physiology of area V1? The highly contrasting effects of loss of foveal versus peripheral vision on human behavior (Faye, 1984) led physiologists and psychologists to examine the effects of retinal eccentricity information processing in the visual cortex (Yu et al., 2015). These studies show that while there are major differences with eccentricity in the spatial frequency tuning of individual neurons in area V1, the shift to higher speed sensitivities with increased eccentricity (Orban et al., 1986) is largely explained by magnification scaling so that overall spatiotemporal computations appear remarkably constant across the visual field (Virsu et al., 1982; Yu et al., 2015). One possibility is that the apparent uniformity suggested by investigations of spatiotemporal tuning across eccentricity in area V1 using non-natural stimuli in anesthetized preparations (Yu et al., 2010) will essentially reflect FF processes and will not be sensitive to the contextual influences of top-down FB effects.

It is often incorrectly assumed that FB to area V1 is overwhelmingly from visual cortical areas. In fact, retrograde tracer injections in the central representation of area V1 back label neurons in 34 cortical areas (in an atlas of 91 areas) across occipital, parietal, temporal and prefrontal regions (Markov et al., 2014a). In the high-density cortical connectome, however, binary connectivity has little predictive power of the properties of the network (Markov et al., 2013). Features such as global and local communication efficiencies modeled as network conduction efficiencies and the heterogeneity of the network emerge from weighted measures of connectivity (Ercsey-Ravasz et al., 2013). The role of weight-distance relations is such that the network is spatially embedded, where a determinate feature is an exponential decline in connection weight with distance, meaning that the probability of two areas being connected can be expressed in terms of an Exponential Distance Rule (EDR) (Knoblauch et al., 2016). Additionally, two major functional streams dominate the visual cortical landscape, an occipital-parietal dorsal stream implicated in spatial vision, and an occipital-temporal ventral stream for object recognition (Mishkin et al., 1983; Ungerleider and Mishkin, 1982). This where/what dichotomy was later reformulated in terms of action/perception (Goodale and Milner, 1992; Goodale et al., 1991); for a more nuanced view see (Rossetti et al., 2017; Sheth and Young, 2016).

Here we address whether FB from the dorsal and ventral streams to visual cortex is shaped by the EDR. This is plausible because the foveal and upper field representations are expected to be closer to the ventral stream, whereas the representation of the lower visual field to be closer to the dorsal stream (Kravitz et al., 2011; Kravitz et al., 2013). The EDR would suggest that these spatial relations of the functional streams would correlate with increased weight of FB projections from the dorsal stream to the lower field representation in dorsal V1 and V2 compared to the upper field representation in ventral V1 and V2. Similarly, the spatial location of the central representation in V1 and V2 would increase the FB frequency from the ventral stream areas. Such findings could provide a structural basis for Previc’s ecological theory of vision, which proposes a functional specialization where the lower visual field is preferentially engaged in peripersonal space and the upper visual field in extrapersonal space (Previc, 1990).

In this manner, there are two factors that we anticipate may correlate with regional variation of FB connectivity to areas V1 and V2. Expression of the EDR rule would predict *quantitative* changes in the sets of FB projections from individual extrastriate areas to specific retinotopic locations, and *qualitative* differences in the origin of cortical FB to dorsal and ventral peripheral subdivisions of V1 and V2, representing upper and lower visual fields. Numerous authors have addressed the influence of eccentricity on the cortical input to areas V1 and V2 (Borra and Rockland, 2011; Falchier et al., 2002; Majka et al., 2019; Rockland and Ojima, 2003). However, none of these studies provided an extensive quantitative estimate of cortical afferents over a wide range of eccentricities combined with an exploration of dorsal and ventral subdivisions. Tracer injections in the central representation of area V1 reveal the distribution of connectivity strengths following a lognormal profile, reflecting the projections from 34 cortical areas (Kennedy et al., 2013; Markov et al., 2014a). Here, building on this knowledge we provide an extended, comprehensive and detailed analysis that allows comparing connectivity profiles of central representations with paracentral and far periphery representations in the upper and lower visual fields. Exploiting previously published data (Glasser et al., 2018; Glasser et al., 2016) we show that partial correlation of functional connectivity data in humans replicate the biases seen in visuotopic connectivity to the dorsal and ventral streams observed in macaques. Our findings reveal numerous long-distance FB projections to peripheral representations from sources including frontal regions not previously described.

Perception can be conjectured as a closed loop convergence which needs to be addressed in the freely moving individual (Ahissar and Assa, 2016; Gallant et al., 1998; Schroeder et al., 2010). During active visual sensing there is a systematic exploration of the world via patterns of saccades and fixations that accompany a contextual modulation of the activity in area V1 (Barczak et al., 2019). While the observed rhythmic changes in activity could be of subcortical origin, their laminar patterns conform to known top-down influences of visual cortex (Ito et al., 2022). The changing patterns of connectivity with eccentricity that we report could play a role in the dynamic coupling of foveal and extra-foveal processing of information that is required in active vision.

## Results

In a first instance we show that surface mapping of connectivity following injections in different subdivisions leads to broadly overlapping projection zones containing back-labeled neurons; however, the density of labeling shows important variation according to the location of the target subdivision in the upper, lower or central visual field. We analyze these variations by means of weight index (FLN see methods) the values of which are related to distances between target and source areas. Numerous studies have argued that the connectivity profile at a cortical location is an important characteristic related to function (Bressler and Menon, 2010; Markov et al., 2014a; Passingham et al., 2002). In this part of the Results Section we examine the connectivity profile, which bears out our prediction that the central, upper and lower visual fields are characterized by distinct connectivity profiles. Examination of these profiles reveals numerous exceptions to the EDR, reminding us that the EDR is an average statistical property of the cortex. Finally, the results reveal numerous instances, particularly to far peripheral subdivisions, of FB projections from non-visual areas in retrosplenial, cingulate, frontal and prefrontal regions.

The full set of 21 injection sites in Table S1 have been mapped to the atlas surface and are shown in **Figure 1**. The extent of each injection and its relation to the borders of the subregion injected are shown in line drawings in Supplementary Figure S1. Examples of connectivity following injections in different subdivisions are shown in Figure S2.

**Figure 1.**
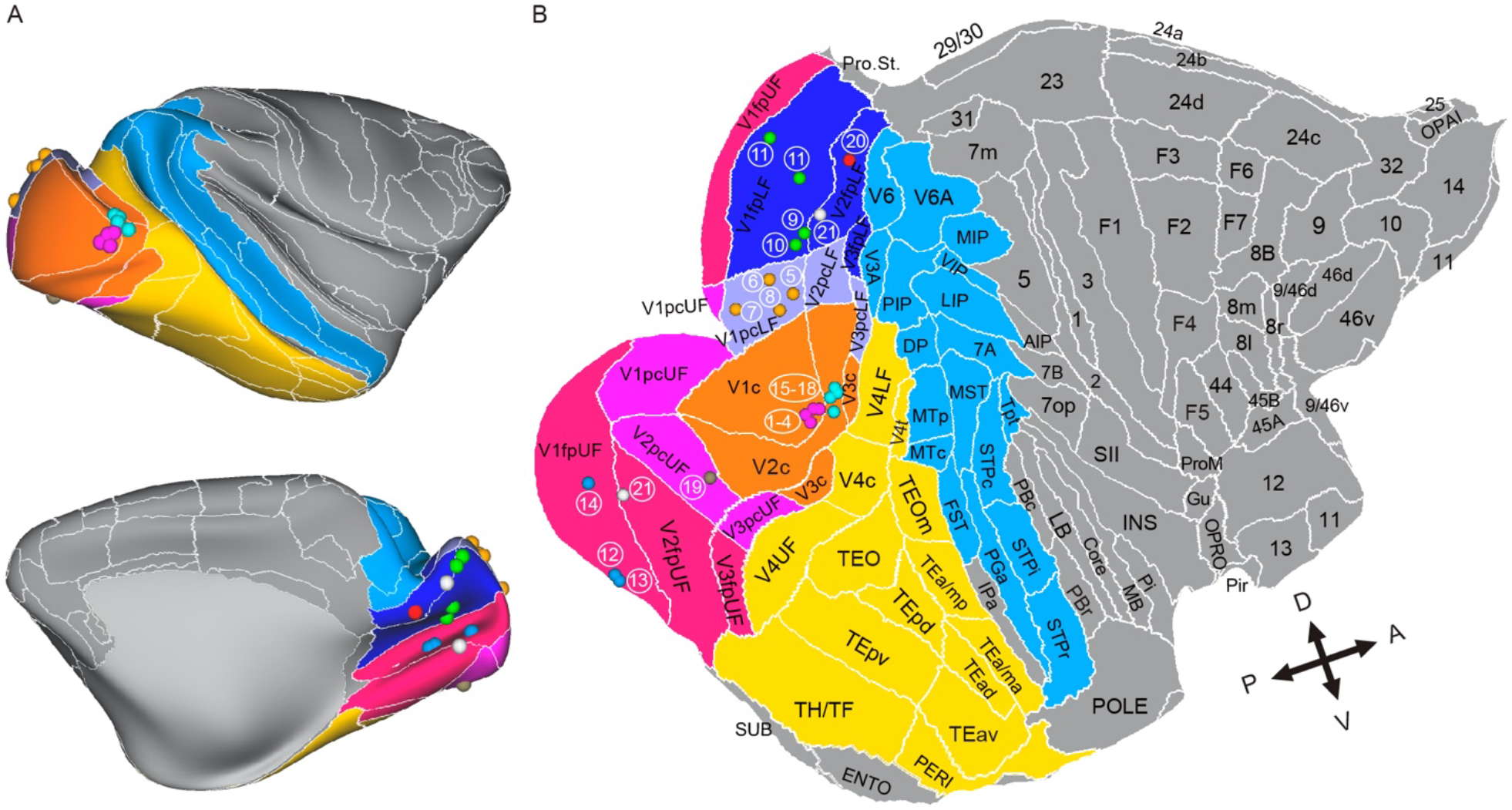
Retinotopic atlas and injection sites in area V1 and V2 retinotopic subdivisions. Injections used in this study in different eccentricity subdivisions are illustrated on the 106-area atlas surface on the lateral and medial surfaces (A) and the flat map (B). Injections in different subdivisions are indicated in different colors. Blue, dorsal stream areas; yellow, ventral stream areas. White numbers in B refer to case numbers reported in Table S1.

Inspection of injection sites (see methods) showed that overall injections were successfully limited to the targeted retinotopic subregion. However, in case 20 directed at the far periphery dorsal V2 (i.e lower visual field), the injection showed a small involvement of the supragranular layers of ventral V2 (i.e. upper visual field; **Figure S1**). We compared the back labeling in dorsal V1 against ventral V1, which showed labeling in dorsal V1 corresponding to 2.7% that in ventral V1 suggesting that the contamination was minimal. In case 21 (V2fpUF/LF) the injection was targeted at upper-field V2 but there was a considerable involvement of the LF and this injection is considered UF/LF injection of the far periphery.

### Non-parcellated mapping of FB connections to retinotopic subdivisions of early visual cortex

A realistic representation of the spatial distribution of labeled neurons following injections in retinotopic subdivisions can be obtained by visualizing the distribution of labeled neurons on inflated and flatmap views of the cortical surface **(Figure 2)**. This shows that while there is extensive colocalization following injections in different subdivisions there are clear regional differences in labeling density.

**Figure 2.**
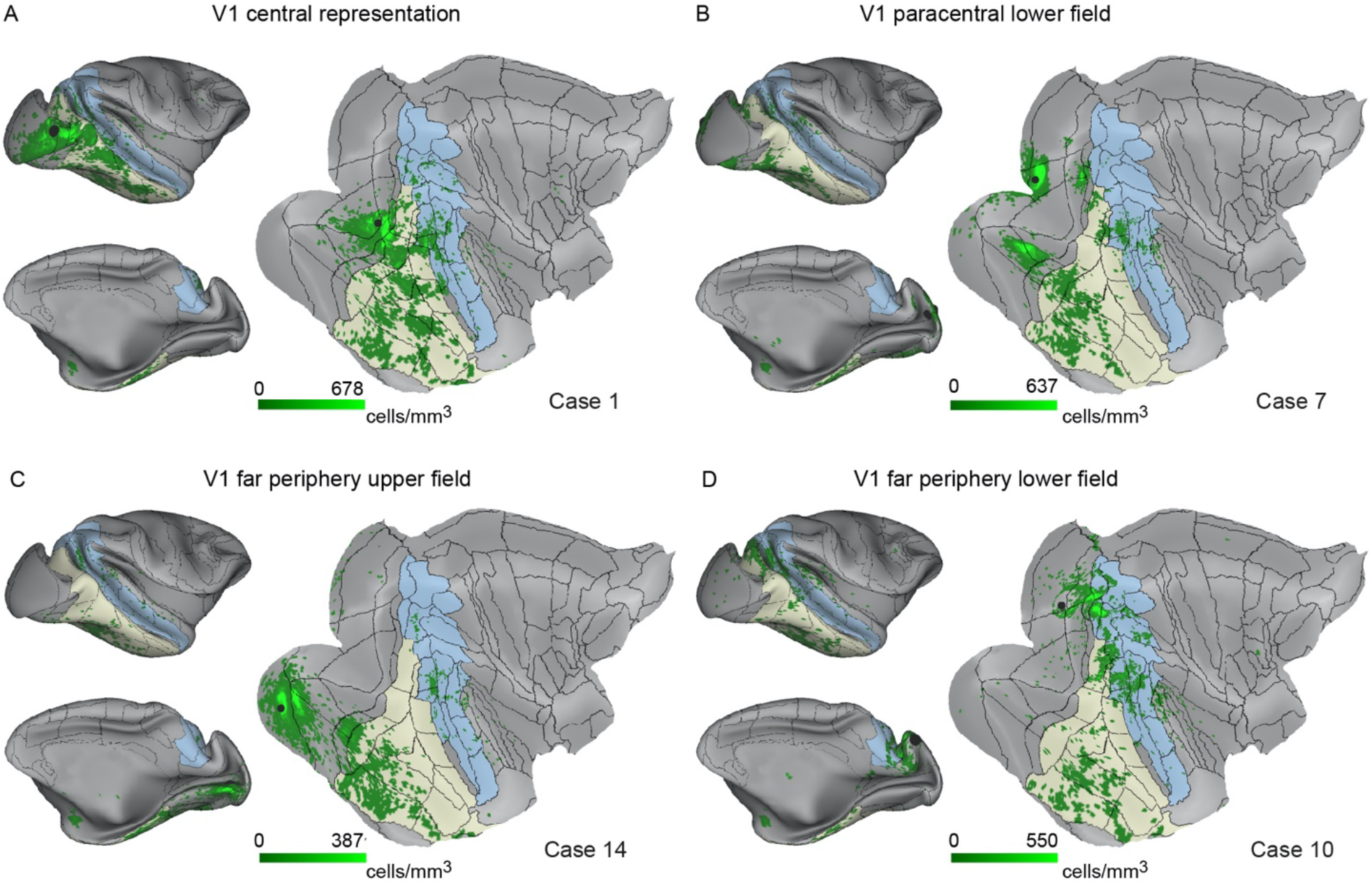
Non-parcellated dense surface mapping of feedback projections to area V1 retinotopic subdivisions. The distribution of the labeled cells of the representative injections in (A) V1c, (B) V1pcLF, (C) V1fpUF and (D) V1fpLF are illustrated on the flat map, the lateral and medial view of the mid-thickness surface of the individual brain. The black dots indicate the injection sites. The injection IDs and case names are indicated.

### Spatial distribution of areal FLN values projecting to retinotopic subdivisions

Tracer injections in the cortex lead to a wide range of connection weights, with FLNe values spanning 5 to 6 orders of magnitude (Markov et al., 2014a; Markov et al., 2011). As in previous studies (Markov et al., 2014a; Markov et al., 2011), connection weights generally conform to a lognormal distribution, and there was in general good consistency across repeat injections (see **Figure 5** below for details). **Figure 3** shows flat maps of the cortex with each row showing results for different retinotopic subdivisions of V1 and V2. The left-hand image of each panel indicates FLNe on a log scale and shows the pattern of labeling with respect to the dorsal and ventral stream areas, outlined in blue and yellow. The middle column indicates the mean distance of each area from the injected subregion. The far-right column indicates those ‘Projections that were Unique to Peripheral eccentricities’ (PUP), i.e., were not detected following injections in the central representation (panel A for area V1 and panel E for area V2). To facilitate aligning distance and FLNe, the PUPs are outlined in fine red lines in the first and second columns.

**Figure 3.**
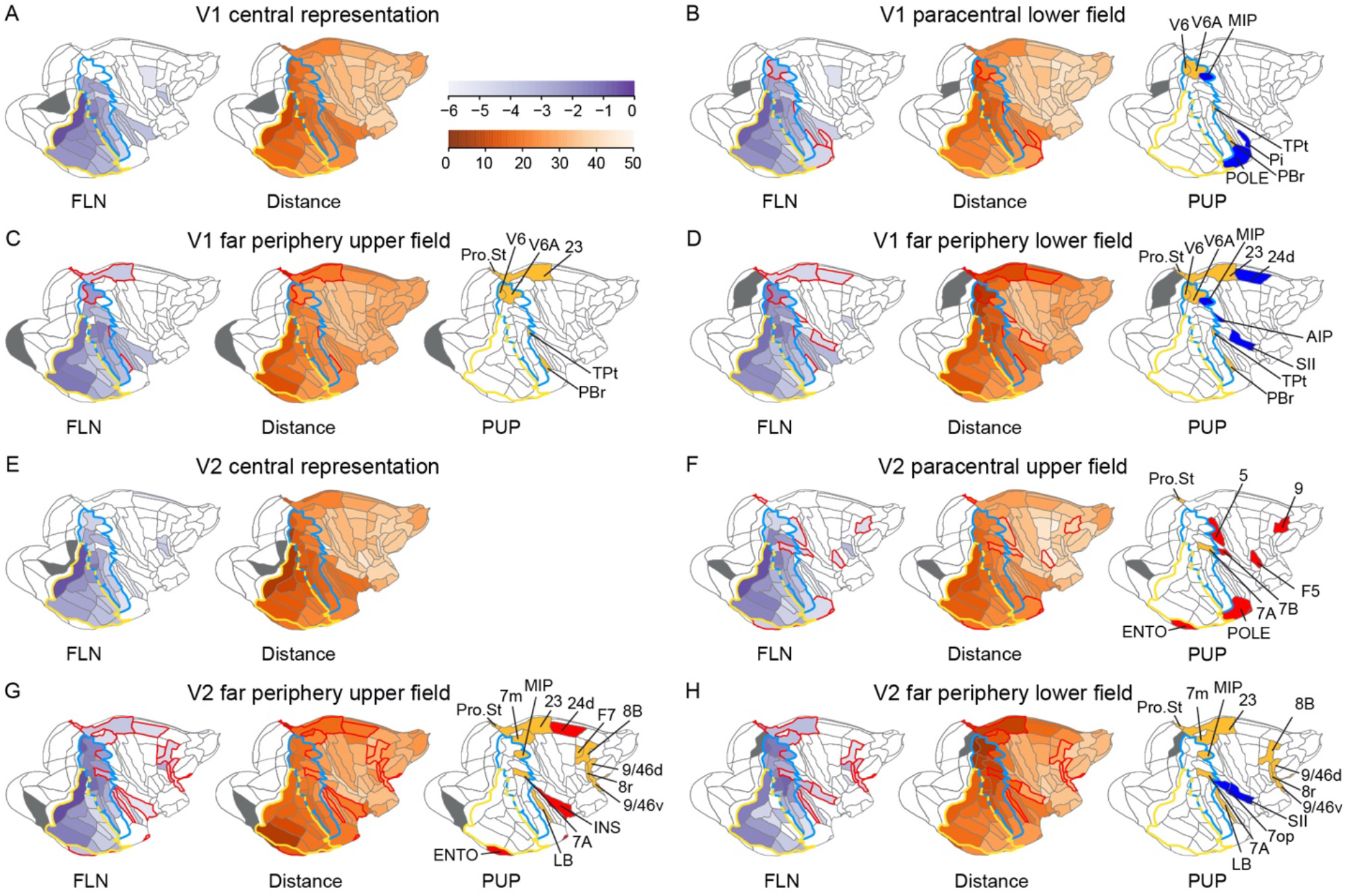
Surface maps of projections to area V1 (panels A-D) and V2 (panels E-H). Left-hand, log FLNe values; middle distances (mm); right-hand, projections unique to periphery (PUP), indicated in red outlines for distance and weight maps. PUPs are color coded in right-hand panel, blue projecting to lower field, red to upper field, yellow to both upper and lower. Areas of ventral and dorsal streams outlined in yellow and blue, respectively. Dorsal stream areas: 7A, STPc, STPr, STPi, FST, TPt, MST, MT, LIP, PIP, MIP, VIP, PGa, V3A, DP, V6 and V6A; ventral stream areas: TEO, TEOm, TEpv, TEpd, TEav, TEad, TEa/ma, TEa/mp, TH/TF, PERI, V4 and V4t. Subdivisions injected indicated in black.

Injections of tracers in the central representation of area V1 (V1c) (**Figure 3A**) label a wide expanse of occipital, temporal, parietal cortex and the frontal eye field in prefrontal cortex and include projections from auditory areas, which are known to connect to visual cortex in macaque (Markov et al., 2014a). Projections to V1c emanate from all areas of the ventral stream but fail to label the dorsomedial stream areas V6, V6A MIP as well as TPt.

In contrast to V1c, injection into the paracentral lower field of V1 (V1pcLF) (**Figure 3B**) labels the full complement of dorsal and ventral stream areas. These injections lead to 7 PUPs including the mediodorsal stream areas (V6, V6A, MIP) as well as TPt, Pi, PBr and POLE. Injection in the far periphery lower field of V1 (V1fpLF) (**Figure 3D**) generates a similar trend in observed spatial patterns observed going from V1c to pcLF; a reduction of FLNe in the ventral stream particularly in the dorsal subdivisions (TEO, TEpd) and the rostral extremity of the ventral stream (TEa/am, TEad, TEav and PERI), accompanied by a frank increase of FLNe in the dorsal stream particularly in the mediodorsal subdivisions. Five of the PUPs labeled by the pc injection (V6, V6A, MIP, TPt, PBr) are also labeled by the fp injection which in addition labels areas 23, 24d, AIP and SII. Compared to injection in V1fpLF, injections in the far periphery of the UF (V1fpUF) (**Figure 3C**) show stronger labeling in the ventral stream particularly in ventral subdivisions (TH/TF, TEpd), and notably weaker labeling in the dorsal stream including an absence of labeled cells in V3A and DP. Note an absence of labeling in V4t. Importantly, PUPs following injections in the fp of both UF and LF are somewhat similar.

The different patterns of labeling observed after injections of the different retinotopic subdivisions partially reflect changes in the distances of the source areas from the injected subdivisions as shown in the middle column of **Figure 2**. Both injections in the fpLF and fpUF revealed area 23 as a PUP and 24d after injection in fpLF, note areas 23 and 24d have similar distances from the two injected subdivisions and significantly shorter than the distances from V1c and V1pc to these areas.

Likewise, the mediodorsal dorsal stream areas (V6, V6A and MIP) showed a similar distance to V1pcLF and V1fpUF (**Figure 3B, C**) and similar densities of labeling. This contrasts with V1fpLF (**Figure 3D**) where compared to V1pcLF (**Figure 3B**) and V1fpUF (**Figure 3C**), the distance from injections site to mediodorsal dorsal stream areas decreased and this too was accompanied by an increase in labeling density.

Like injections in area V1c, injections in area V2c (**Figure 3E-H**), label all ventral stream areas and area 8_L_ and 8m in prefrontal cortex, as well as all dorsal stream areas with the exception of MIP and 7A. However, whereas V1c injections label auditory areas CORE, MB, LB, PBc and PBr, injections in V2c only label MB and PBc. Injections in V2pcUF (**Figure 3F**) label all dorsal stream areas with the exception of TPt. Compared to injections in peripheral V1, peripheral V2 injections label more PUPs and in more rostral locations. V2pcUF labels numerous PUPs, including frontal (area F5) and prefrontal cortex (area 9), parietal cortex (5, 7A, 7B) and temporal cortex (ENTO) as well as the PUPs observed following V1pcLF injection (POLE) and V1fpUF, V1fpLF injections (Pro.St).

The injection in V2fpUF/LF (**Figure 3G**) contaminated the lower field means that here variations of dorsal versus ventral stream areas are of doubtful value. Nevertheless, we observe increases of intensity of labeling in dorsal stream areas compared to V2pcUF particularly in V6, V6A, MIP, V3A, MT, DP and 7A. The distribution of PUPs after V2fpUF/LF and V2fpLF includes areas 23 and 24d in the medial wall as found following injections in V1fpLF and V1fpUF/LF. Injection in V2fpUF/LF (**Figure 3G**) also labels 2 PUPs (ENTO, 7A) also found after V2pcUF.

The injection in V2fpLF (**Figure 3H**) labeled all dorsal stream areas except area IPa and all ventral stream areas except V4t, TEOm, TEa/ma and TEad. Injection in V2fpLF leads to increased levels of labeling in mediodorsal dorsal stream areas (V6, V6A, MIP) compared with injections at all other eccentricities and revealed 12 PUPs: Pro.St, 23, 7m, MIP, 7A, 7op, SII, LB, 8B, 8r, 9/46d and 9/46v. The majority of PUPs are common to both injections and could be the consequence of the involvement of LF and UF in the injection reported in **Figure 3G**, alternatively the injection of the fp in the LF and UF leads to similar PUPs, as was the case in the V1 injections. In any case, there are only 2 PUPs appearing to be specific to the V2fpUF/LF (ENTO, INS) and to V2fpLF (7op, SII) suggesting that there might not be a PUP signature of V2UF and V2LF.

Again, we note that variation of FLNe can be related to variation of distances. Hence, as in area V1, the high level of labeling of the mediodorsal areas following injections in V2fpLF compared to more central injections correlates with shorter distances, and once again we observe PUPs in the cingulate cortex (areas 23 and 24d) following the fp injections.

### Spatial embedding of FB projections to different retinotopic subdivisions

Here we explore how the physical distances of the retinotopic subdivisions to areas in the dorsal and ventral streams shape the collective FLNe values of FB connections from the two functional streams. In **Figure 4A** we calculated the ratio of dorsal vs ventral stream neurons projecting to the different retinotopic subdivisions of V1 and V2. This shows that in both areas, ventral stream projections dominate the central representation of V1 and V2 and all subdivisions of the upper visual field as well as the paracentral divisions of lower field. This contrasts with the domination of dorsal stream projections to the fpLF subdivisions of both areas V1and V2. Similar results are indicated by considering the cumulative input (i.e. FLNe) from the functional streams to the retinotopic subdivisions (**Figure 4B**). To what extent do these results reflect the distances of the subdivisions from the functional streams? In **Figure 4C**, we show that in area V1 the average distance of the upper field injection sites to the ventral stream is significantly shorter than to dorsal stream areas. The inverse is found in the lower visual field where dorsal stream distances tended to be shorter reaching significance in V1fpLF. Similar findings were found in area V2 (**Figure 4D**); upper field distances were significantly shorter to ventral stream areas, while in the lower visual field distances were significantly shorter to dorsal areas and more markedly so than in area V1. Interestingly, both in V1 and V2, the central representation was equidistant to both streams.

**Figure 4.**
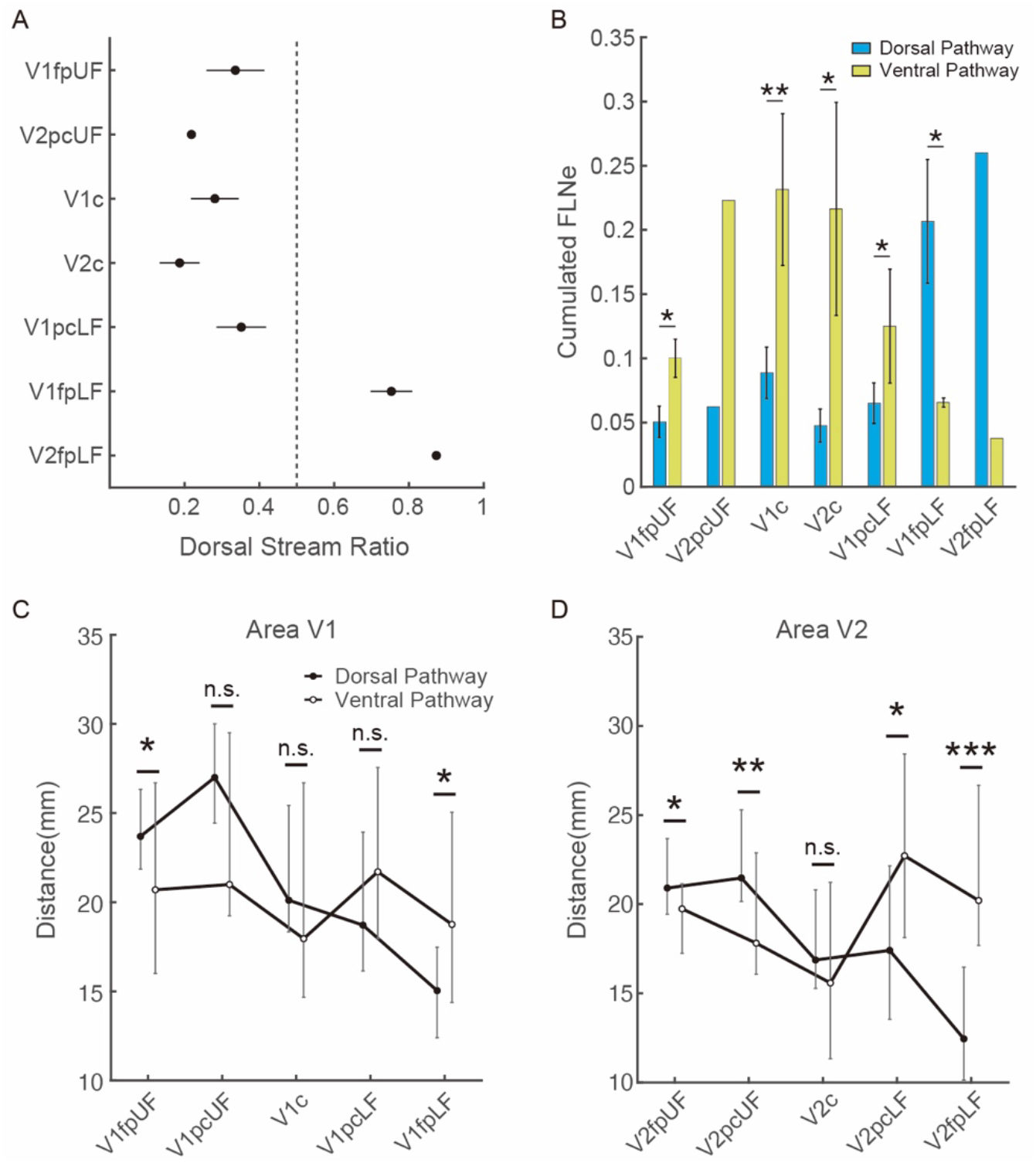
Spatial embedding: Differential strength of projections to retinotopic subdivisions of areas V1 and V2. (A) Dorsal Stream Ratio for the injected subdivisions, as values approaches 0, the connectivity is dominated by the ventral stream. (B) Cumulated FLNe value from dorsal and ventral areas to retinotopic subdivisions of areas V1 and V2. C; D) tractographic distances from dorsal and ventral pathway areas to V1 (C) and V2 (D). The error bars indicate 25% and 75% quantiles (interquartile range) of total distance. Significance, between the dorsal and ventral streams estimated via permutation tests (n = 10000) which randomly re-assigned the distances to different streams under the null hypothesis that the average distance to the injection site was the same for dorsal and ventral areas. *, P < 0.05; **, P < 0.01; ***, P < 0.005; n.s., not significant.

### Influence of eccentricity on connectivity profiles

Tracer injections lead to a stereotypic broad distribution of weights ranging that exhibit a lognormal distribution (**Figure 5A**) (Markov et al., 2014a; Markov et al., 2011). Here we anticipate that repeat injection in retinotopic subdivisions will likewise generate specific connectivity profiles.

**Figure 5.**
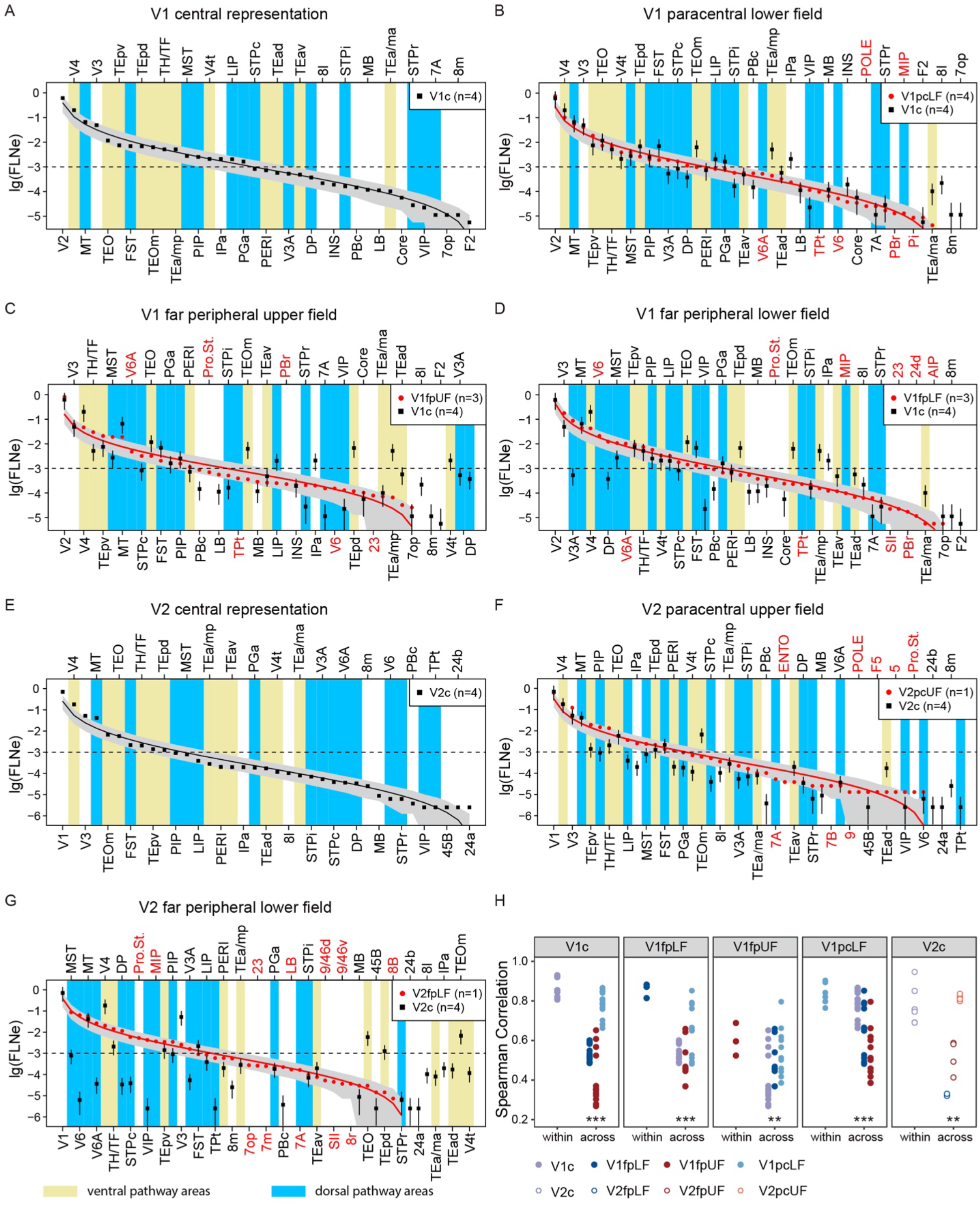
Connectivity profiles of retinotopic subdivisions in area V1 (panels A-D) and area V2 (panels E-G). (A) black squares, means log(FLNe) and SD of V1c values. (B-G) In each subdivisions, areas are ordered by mean log(FLNe) in red; The grey envelope, 95% confidence band indicated by a negative binomial distribution, dispersion=7. Blue, dorsal stream areas; yellow, ventral stream areas. H, Spearman correlation of connectivity profiles of injections within and across V1 subdivisions, including connections projecting to only center and only periphery. Each panel illustrates the result for each subdivision. Within: the correlation of connectivity profiles within the subdivision; across: correlation of connectivity profiles across subdivisions. Wilcoxon test, ***, P < 0.001, **, P < 0.01.

Figure 5. compares the lognormal connectivity profiles in area V1 and V2 obtained at different eccentricities. Comparison of the weights of projections resulting from peripheral and central injections shows that they differentially involve areas in dorsal (blue bars) and ventral stream areas (yellow bars). Hence going to -3 log (FLNe) there are 7 blue bars in V1c, 8 in V1pcLF and the maximum number is 12 in V1fpLF (compared to 9 in V1fpUF). With this change in order there is a change in the average Spearman rank coefficient (see later).

Here we address the extent to which changes in eccentricity differentially impact FLNe of FB projections from individual areas. In V1pcLF (**Figure 5B**) there are two instances (TEOm, TEa/mp) of a significant FLNe difference in FB of ventral stream areas to the central and peripheral targets. In both cases the peripheral injections results in lower FLNe values. For dorsal areas there are two cases of significant differences (V3A and VIP) where the peripheral injections result in higher values. In the V1fpLF (**Figure 5D**) there are six instances of significant differences for the ventral stream areas (V4, TEO, TEpd, TEOm, TEa/mp), and in all cases the peripheral injection resulted in lower values. At this eccentricity there is a significant increase in FLNe of three of the dorsal stream areas (V3A, DP, VIP). In aggregate, these findings support our hypothesis that the LF of area V1 is preferentially connected to the dorsal stream.

In V1fpUF (**Figure 5C**) there are 6 instances of eccentricity dependence of FLNe in ventral stream areas and in all cases the peripheral injection resulted in a significant decrease (V4, TEOm, TEpd, TEa/mp and TEad). At this eccentricity in the dorsal stream there are 7 instances of significant difference where two are lower following the peripheral injections (MT, LIP) and 5 are higher (area 7A, MST, STPc, STPi and STPr). The ventral dominance for V1fpUF shown in **Figure 5C** can be explained by the fact that the decrease of ventral area FLNe with respect to the central representation is compensated by connectivity increases in the dorsal stream areas, thus resulting in a similar DSR value.

In **Figure 5**, names of PUP areas are in red. This figure shows that there are notable examples of PUPs with FLNe values comparable to the projections to the central representation, *i*.*e*. PUPs do not constitute a category of exceptionally weak connections. This is particularly the case for V6A and Pro.St following injection in V1fpUF and V6, V6A and Pro.St. following injection in V1fpLF. There are between 2 to 5 areas, shown on the far right of the central distribution, that were labeled by the central but not the peripheral injections. These central-only projections were generally weak except for V4t, V3A and DP, which were not labeled after injecting V1fpUF.

Injections in area V2 likewise show important variations in the FLNe values of the FB pathways from dorsal and ventral stream areas to different retinotopic subdivisions (**Figure 5E-G**). As in area V1, there is only a systematic bias in the fpLF injections of V2 (**Figure 5G**), where there are 8 instances (V6, V6A, DP, STPc, VIP, PIP, TPt, 8m) of a significant difference between peripheral and central FLNe values in dorsal stream areas, and in these cases the FLNe is higher following peripheral compared to central injections. In V2fpLF there are significant differences in 4 ventral stream areas compared to central injections, where 3 return lower FLNe values (V4, TEO and TEpd) and one instance a higher FLNe value (TH/TF). In the case of V2fpLF there are 6 instances of center-only projections having relatively high FLNe values (4 ventral stream areas (TEa/ma, TEad, TEOm, V4t) and two dorsal stream areas (8_L_, IPa). Again, for the two eccentricities examined in V2 and as in area V1, PUPs are observed to have FLNe values comparable to those of areas found across all eccentricities.

In this analysis we excluded the injection of V2fpUF/LF shown in **Figure 2G** because of the involvement of fpLF. This injection revealed very similar PUPs with both the LF and UF injections, and the weights of these PUPs are comparable to those shown in V2pcUF and V2fpLF (data not shown).

A Spearman correlation analysis showed that with the exception of V1fpUF injections in each retinotopic subdivision result in connectivity profiles that are highly consistent, similar to the consistency of connectivity profiles of an entire cortical area across individuals (**Figure 5H**) (Markov et al., 2014a). This contrasts with across subdivision analyses which, in all cases, showed that connectivity profiles differed significantly.

The examination of weights for injections at different eccentricities shows that there is a trend for the fpLF to be more strongly connected to dorsal stream areas than to ventral stream areas. The other eccentricities conversely have a globally stronger input from ventral compared to dorsal stream areas. A further clarification is to look at the relative dominance of FB projections to different retinotopic domains. We define a dominant domain as one for which the log(FLNe) value of the projection from a given area to a particular subdivision is higher than the other two subdivisions to a specified extent. A probabilistic map of domination was made by adjusting the threshold from 0.3 to 1. The results in **Figure 6** show that FB projections to the central representation of V1 dominate in the ventral stream (notably V4, TEO, TEOm, FST, TEa/mp, TEpd, TEad and IPa), in frontal F2 and prefrontal 8m. This contrasts with the FB projections to the far periphery of area V1 where the dorsal stream dominates (notably in areas V6, V3A, DP, MIP, VIP, AIP), frontal SII and area cingulate 24d. The FB projections to the far periphery of the upper-field dominate the ventral component of the ventral stream (TH/TF), the auditory associative area PBr, the dorsal stream area STPr and the retrosplenial area 23.

**Figure 6.**
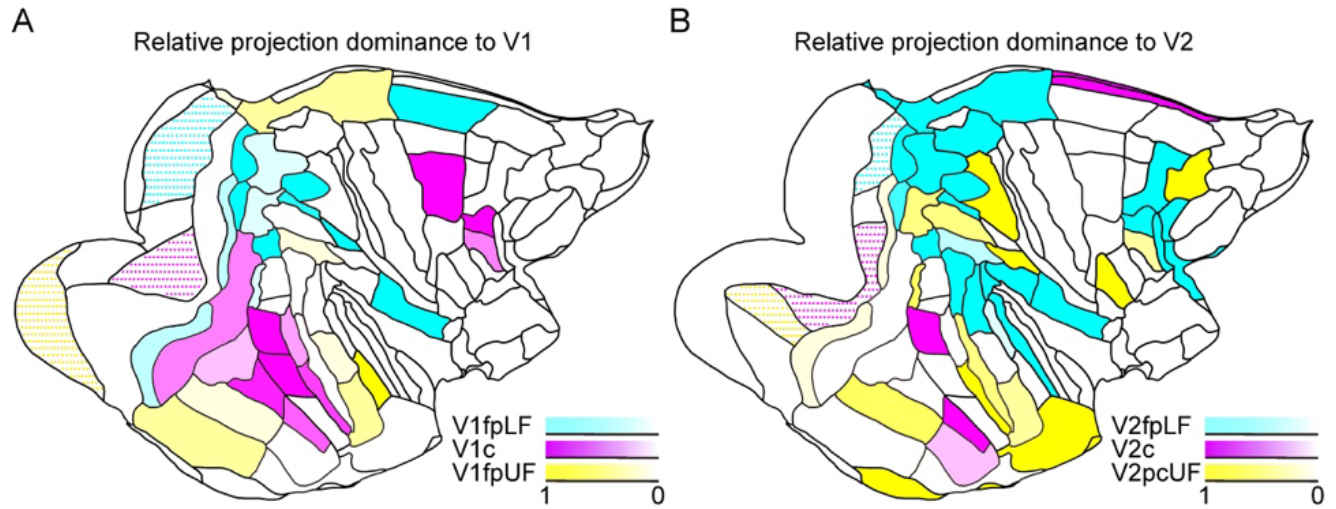
Probabilistic map of relative projection dominance to retinotopic subdivisions of area V1 (A) and V2 (B). Areas projecting dominantly to V1 (A) and V2 (B) V2c. Central dominance magenta, fpLF dominance blue, fpUF Yellow.

The dominance of FB projections to area V2 compared to area V1 are more dispersed and somewhat more rostral (**Figure 6B**). There is a marked contraction of dominance to the central representation (TEO, TEad, 24a and 24b). This contrasts with dominance to pcUF which extends across the temporal, parietal and prefrontal regions. Projections to the fpLF of area V2 stretch far dorsally dominating in area Pro.St, 23, VIP, MST, STPc, TPt, PBc, LB, 45B, 9/46d, 9/46v.

### Influence of eccentricity on non-visual projections to visual cortex (Figure 7)

In the present study, we find strong evidence of projections to area V1 from auditory areas targeting central and peripheral subdivisions and UF and LF (**Figure 7A**). Projections to the central representation of area V1 originate from the primary auditory area CORE and the auditory associative areas PBc, MB and LB. Injections in the far periphery lead to labeling of a wider range of associative areas included PBr and TPt and are significantly stronger to V1fpUF and V1fpLF (top left and top right panels in **Figure 7A**). The case V1fpLF, which shows stronger projections compared to central representations the injection is offset from the representation of the vertical meridian, whereas one of the two injections (case 11) at this eccentricity that showed more modest elevations is located near to the representation of the vertical meridian. This effect has been previously reported by Rosa’s group (Majka et al., 2019). The auditory projection to V1pcLF appears intermediate in robustness to projections to central and far periphery of the upper and lower visual fields. The projections from auditory cortex to area V2 are significantly less robust that that to area V1 in so far as it does not include projections from CORE and only involves 3 associative areas to the central representation (MB, PBc, TPt) (**Figure 7A bottom row**). There is an effect of eccentricity that might be more marked in fpLF, which in addition to having projections from MB, PBc and TPt also has projections from LB.

**Figure 7.**
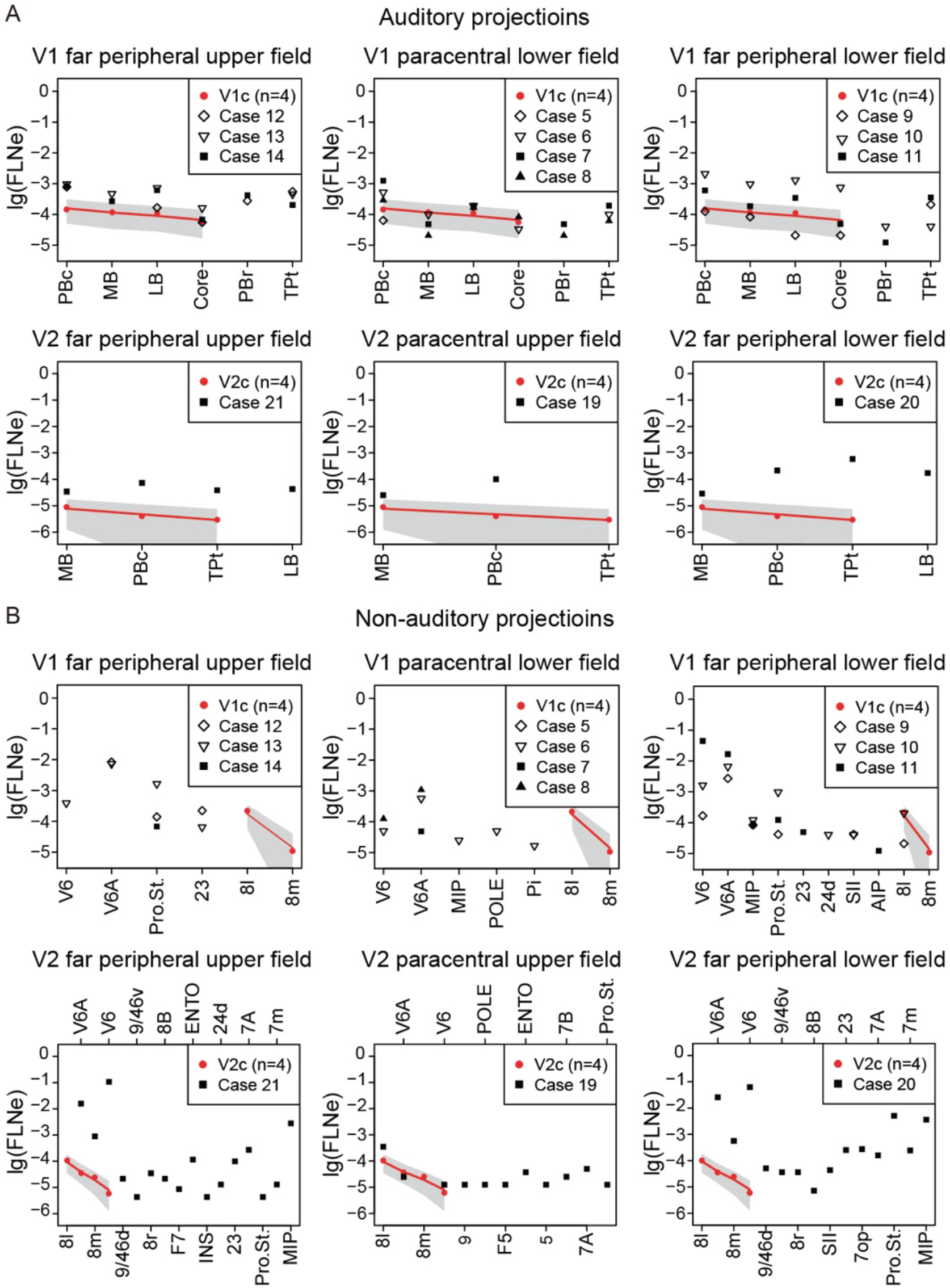
Influence of eccentricity on projections to peripheral subdivisions of area V1 and V2. (A) Auditory projections (B) Non-auditory projections. Grey envelope, 95% confidence band indicated by a negative binomial distribution, dispersion=7; red dots, central projections

### Similar results in humans using non-invasive functional connectivity

We examined human resting-state fMRI data from the Human Connectome Project (HCP) to demonstrate retinotopic relationships similar to those shown above (**Figures 2 and 3**) using invasive anatomical connectivity in the macaque. Glasser et al. (Glasser et al., 2016) previously used a V1 spatial regression model to map the visuotopic organization of extrastriate visual areas based on multiple linear regressions (see Methods section *‘Analysis of human functional connectivity’* and also Supplementary Methods Figure 9) (Glasser et al., 2016). **Figure 8A** illustrates results from a lower field versus upper field regressor. Positive (yellow and red) regions have stronger functional connectivity with the lower field; this includes nearly all dorsal stream areas (up on the cortical flatmap). Negative (blue) regions, including nearly all ventral stream areas (down on the flatmap) have stronger functional connectivity with the upper visual field. Similarly, **Figure 8B** shows central versus peripheral regressor results, where positive regions have stronger functional connectivity with central visual fields and negative regions have stronger functional connectivity with peripheral representations. Although dorsal and ventral stream areas do have both central and peripheral representations in many cases, the dorsal stream areas tend to have stronger and larger peripheral representations and the ventral stream areas to have stronger and larger central representations. Finally, **Figure 8C** illustrates a full-rank partial correlation functional connectivity analysis based on the 180 area-per-hemisphere HCP multimodal parcellation. (correlation between a pair of areas after controlling for all other areas) seeded from the right auditory core. This reveals specific functional connectivity between the auditory core and V1 (but not V2 or V3) and also to primary somatosensory cortex area 3b (Glasser et al., 2018). These findings are highly consistent with the present invasive tract-tracing results shown above in the macaque and suggest that humans likely have similar biases in anatomical connectivity.

**Figure 8.**
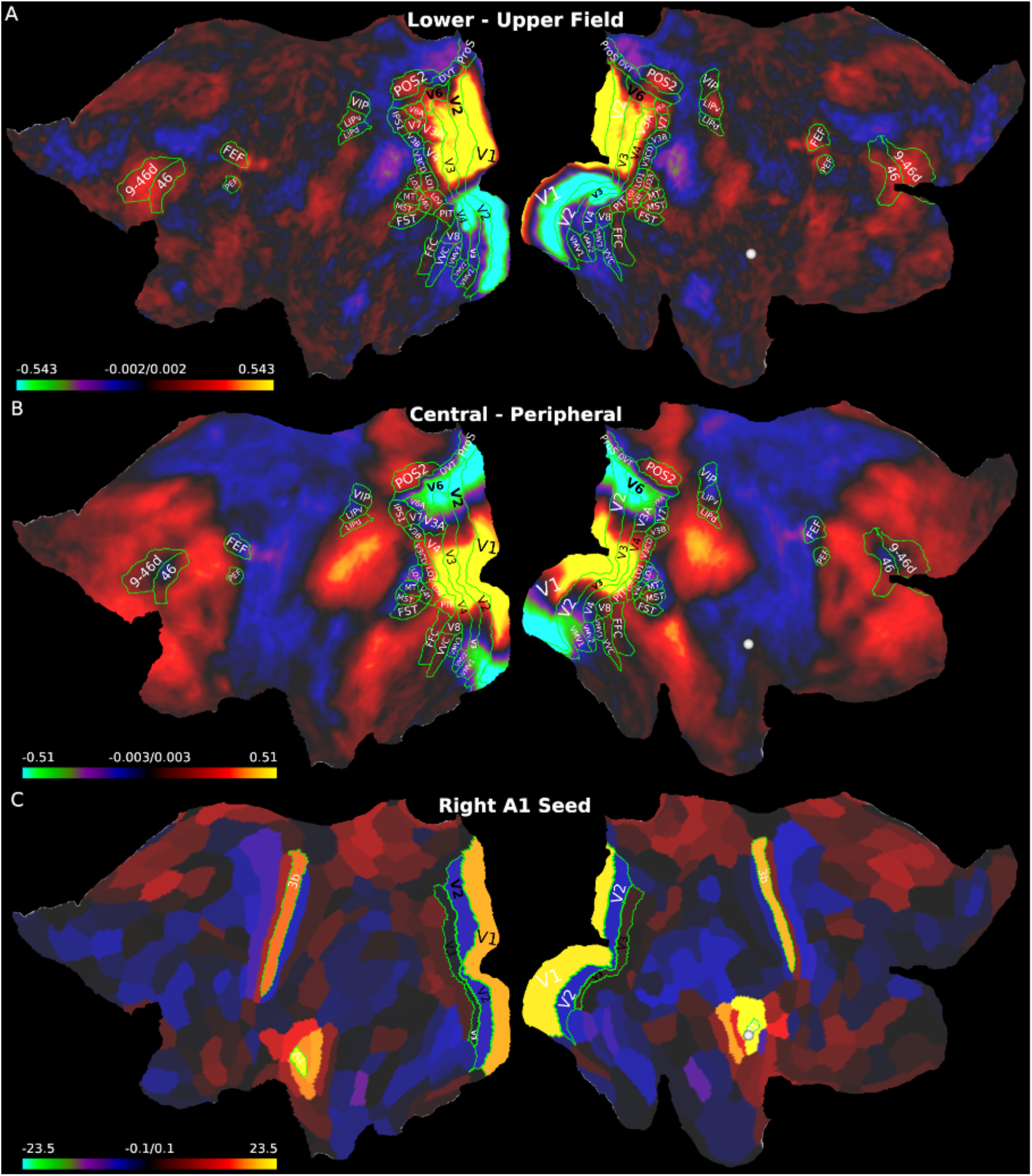
Human partial correlation functional connectivity contrasts and results displayed on left and right hemisphere cortical flatmaps. A. The contrast in resting-state functional connectivity between lower field (positive) and upper field (negative) while controlling for other visuotopic signals within V1. B. The contrast in functional connectivity between central (positive) and peripheral (negative) in the same model. C. Full-rank partial connectivity results seeded in right A1 (white sphere). The maps in A and B are from the same HCP 210P dataset shown in Supplementary Methods Fig. 8B, D, respectively of Glasser et al. (Glasser et al., 2016) but displayed on flatmaps instead of inflated surfaces and using a different palette range. The maps in C are from the 449 HCP-YA subjects that were previously published in (Glasser et al., 2018).

## Discussion

In the high-density inter-areal cortical network where over two-thirds of possible connections actually exist, it is the strength of existing connections, the so-called connectivity profile, that determines the properties of the network. A major defining feature of cortical connectivity is the exponential decline of connectivity with distance, the so-called EDR. In the present study we examined how the EDR shapes the variation of connectivity profiles in different retinotopic subdivisions. However, the EDR is a statistical property of the cortex. Paradoxically while it is highly predictive of network properties it predicts a much narrower distribution of connection weights with distance than what is observed meaning that it is an average, statistical property of cortical geometry (Horvat et al., 2016). Hence, while the predicted bias of dorsal stream FB connections to the representation of the lower visual field is supported by the present findings, the projections of individual areas nevertheless provide numerous counterexamples as can be seen in the comparisons of the connectivity profiles across different eccentricities.

Previc’s ecological theory of vision proposes that far vision is dominant in the upper visual field while near vision is emphasized in the dorsal visual stream and the peripheral lower visual field (Previc, 1990). It accounts for numerous observations of psychophysical and behavioral differences in the lower and upper visual field (Bjoertomt et al., 2002; Lane et al., 2013; Levine and McAnany, 2005; Weiss et al., 2000; Zito et al., 2016). We do not find evidence of a fingerprint of the upper, lower and central visual fields in terms of non-visual inputs. But we do find evidence supporting differential connectivity of dorsal and ventral streams to retinotopic subdivisions, namely following an economy of wiring principal the dorsal visual stream is preferentially connected to the far periphery lower visual field, whereas the upper and central visual fields are preferentially connected to ventral visual areas. We report similar findings using non-invasive functional connectivity in humans, supporting the hypothesis that humans have similar underlying visuotopic biases in anatomical connectivity to V1.

The present results report FB projections to areas V1 and V2 from 37 to 45 areas including extrastriate visual areas as well as a number of non-visual areas in retrosplenial, cingulate, parietal, frontal, prefrontal and temporal regions. Here we identify four major channels, each potentially conveying distinct contextual influences to the retinotopic subdivisions of the early visual cortex (**Figure 7**).

### An auditory channel: primary auditory area CORE and auditory associative areas MB, LB, PBr and TPt (Figure 7A)

In recent years advances in signal analysis have led to a revision of the notion of unimodal primary areas suggesting that the neocortex might essentially be multisensory in function (Cappe and Barone, 2005; Fu et al., 2003; Gau et al., 2020; Ghazanfar and Schroeder, 2006; Hasson et al., 2016; Henschke et al., 2015; Liang et al., 2013; Macaluso and Driver, 2005; Murray et al., 2016). The major focus on the projection of non-visual areas to early visual cortex in the primate brain has been on those originating from auditory cortex which in addition to multisensory integration and their impact on perception (Bedny, 2017; Deen et al., 2015; Kanjlia et al., 2021; Kupers and Ptito, 2014; van Wassenhove et al., 2005), have been implicated in cortical plasticity and cross modal reorganization (Gau et al., 2020; Vetter et al., 2020) and as a model system for investigating predictive processing in the cortex (Beierholm et al., 2020; Gau et al., 2020; Rohe et al., 2019; Rohe and Noppeney, 2015; Vetter et al., 2020).

A number of studies have reported projections in macaque from auditory to early visual areas (Falchier et al., 2002; Majka et al., 2019; Markov et al., 2014b; Rockland and Ojima, 2003). In macaque retrograde tracers revealed projections from primary auditory cortex (CORE) and auditory associative areas in a low to high gradient across the central, paracentral to far-periphery retinal representations of area V1 (Falchier et al., 2002). Projections from associative auditory area to the far periphery V1 and V2 were subsequently reported following injections of anterograde tracers in LB (Rockland and Ojima, 2003). More recently, a gradient of auditory input to area V1 was reported in marmoset; LB was found to project to the central representation of area V1, with a stronger projection to peripheral representations from CORE, MB, LB and PBc (Majka et al., 2019). This study found no projections in the marmoset from auditory cortex to area V2. The present study confirms previous reports that projections of CORE, MB, LB and PBc to the central representation of V1 are robust (Markov et al., 2014a), and somewhat stronger to paracentral and far periphery representations of area V1, at least to representations offset from the vertical meridian (**Figure 7A**). Projections from auditory cortex to area V2 did not include projections from CORE were found to be globally less robust than to area V1 (**Figure 7A**). Similarly in humans with non-invasive functional connectivity, we found selective connectivity between A1/CORE to V1, but not V2 (or V3).

The influence of auditory inputs to area V1 is extremely rapid (Giard and Peronnet, 1999), and the termination of auditory connections in areas V1 and V2 have a FB signature (Rockland and Ojima, 2003) as do the laminar locations of the projections as revealed in the present study and elsewhere (Falchier et al., 2002; Majka et al., 2019; Markov et al., 2014a). The auditory influence relayed by FF projection from V1 to V2 and the feedback from auditory cortex to V1 and V2 could generate the mixed FF and FB effects that have been observed (Foxe and Schroeder, 2005) and which can be further elucidated using high resolution fMRI (Vetter et al., 2020).

### A retrosplenial channel: motion areas prostriata and area 23 (Figure 7B)

Our findings show that the limbic area prostriata has a relatively robust projection to the far periphery of the upper and lower visual field of areas V1 and V2, whereas projections to the paracentral representation are considerably weaker and completely absent to the central representations. Prostriata has a limbic cytoarchitecture (Sanides, 1972), and recent connectivity and physiology studies suggest that it may constitute a primitive visual motion region (Smith, 2021; Yu et al., 2012). Electrophysiological recordings and fMRI reveal an extensive representation of the peripheral visual field with very large receptive fields, little eccentricity dependence, low stimulus selectivity, and short latency responses to very fast stimuli in human (Mikellidou et al., 2017) and marmoset (Yu et al., 2012). Tracer studies reveal connections with the peripheral representation of the early visual cortex in macaque (Sousa et al., 1991) and rodent (Lu et al., 2020). In rodents and perhaps in humans, prostriata receives a direct input from the dorsal lateral geniculate (Chen et al., 2021; Kurzawski et al., 2020). In addition to the projection from prostriata we found a considerably weaker projection from the retrosplenial area 23, which has also been proposed to constitute a primate visual motion area (Smith, 2021).

The large size of the receptive fields of prostriata and its extensive connectivity with limbic, hippocampal, cingulate, parietal, medial prefrontal, orbitofrontal and frontopolar cortices suggests an anatomical bypass across the cortical hierarchy thereby allowing peripheral vision to exert a rapid and widespread control over behavior and cognition (Yu et al., 2012). These features suggest that prostriata is a member of a network that is distinct from the ventral and dorsal streams (Mikellidou et al., 2017; Saleem, 2020; Tamietto and Leopold, 2018; Yu et al., 2012). There have been several suggestions as to how prostriata contributes to behavior including monitoring the periphery for sudden motion, participating in postural control and spatial navigation given recent work in rodent (Mikellidou et al., 2017; Tamietto and Leopold, 2018). Our findings showing that prostriata has very strong projections uniquely to the periphery of areas V1 and V2 suggests that it is relaying a unique mix of contextual information to the early visual areas. In humans, prostriata also has strong peripheral connectivity.

### A dorsomedial dorsal stream channel: the V6 motion area and the V6A and MIP sensory-motor areas (Figure 7B)

Area V1c lacked any projections from V6, V6A and MIP, while area V2c received only weak projections from V6 and V6A, and none from MIP (**Figure 7B**). Areas V6, V6A and MIP had strong projections and in that order to the far periphery of the lower field, which were significantly stronger than to V1pcLF, where values were intermediate to the center and far periphery lower field. In ventral stream dominated subregions of V1 and V2 (*i*.*e*. all regions including the central representation and excepting the far periphery lower field) the strongest projections were from V6A followed by V6 and then MIP. This contrasted with the far periphery lower field representations, where the strongest projection was from V6 followed by V6A and then MIP.

There is a clear distinction between V6 which is considered to be a visual motion area and V6A and MIP which are visuomotor areas involved in online control of prehension (Bakola et al., 2017; Galletti and Fattori, 2018). Information from V6 takes a dorsomedial path to V6A and MIP and a dorsolateral path to MT, V3A and LIP. Our findings show that V6 and V6A projections to areas V1 and V2 vary with eccentricity in keeping with their emphasis on the periphery representation, contrasting with MT where our results show that strength of connections are relatively constant across eccentricity. These observations with respect to the two motion areas V6 and MT are consistent with their different functional roles in motion analysis (Pitzalis et al., 2012). In humans (**Figure 8**), V6 and V6A have strong peripheral and lower field functional connectivity.

### A frontal and prefrontal channel: frontal eye fields (areas 8_L_, 8M) and frontal areas insula, F7, 8r, 8B, 9/46d and 9/46v (Figure 7B)

This wide range of areas projecting to the far periphery of the lower field of area V2 might also project to the far periphery of the upper field given that a similar pattern of projections is revealed following the V2fpUF/LF injection that was contaminated by leakage into the fpLF. There is extensive literature showing that in the congenitally blind the “visual” cortex is activated by linguistic stimuli and shows enhanced functional connectivity with language response frontal regions (Amedi et al., 2003; Bedny et al., 2011). More recently, a resting state and task fMRI investigation exploiting a go/no-go paradigm revealed interactions of visual cortex with frontal executive function networks (Kanjlia et al., 2021). The authors of the study argue that the influence of the prefrontal cortex is not the consequence of large-scale anatomical changes in blindness, but rather the efficacy of known connectivity patterns between fronto-parietal cortex and the visual system. The direct projections of the prefrontal cortex to the far periphery of area V2 that we report here would be ideally suited for supporting the reported top-down executive functions. In humans (**Figure 8**), areas 9/46d and 46 have stronger lower field and peripheral functional connectivity relative to other prefrontal areas.

### Active sensing and multimodal integration in the peripheral representation of the early visual cortex

The principal finding of the present study is that compared to foveal representation, the peripheral representation of the early visual areas in the macaque has a more extensive connectivity with numerous non-visual areas. A window to the functional potential of these connections can be gleaned from work in the congenitally blind where higher-cognitive functions are associated with “visual” cortex. However, such considerations do not further our understanding of the function of these connections in the normal brain, which raises the difficult issue of the role of area V1 in higher cognitive functions including visual consciousness (Gilbert and Li, 2013; Linton, 2021; Tong, 2003). Progress in this direction has been made in rodents, and one way of approaching these findings is to consider the mouse primate differences. The fovea and its cortical representation is largely a primate-specific feature, while the peripheral representation in primate retina is similar to the entire retina in mouse (Huberman and Niell, 2011). Non-visual connections to rodent early visual cortex have been known for many years and recently have been shown increasingly to support higher-cognitive functions (Poort et al., 2015). In this way, the peripheral representation of the primate cortex possesses a connectivity that is more rodent-like and where non-visual inputs may be enriching the cognitive capacity of these early visual areas. In the present study, we find inputs from the retrosplenial, frontal and prefrontal cortex to the peripheral representation of the early visual cortex. The input from the retrosplenial and motor cortex that we report could be homologous to motor signals to mouse visual cortex, which allows locomotion based predictions (Niell and Stryker, 2010; Pakan et al., 2018).

A complete account of perception requires an active agent. There is considerable evidence that motor signals are fed back into the sensory processing (Merriam and Colby, 2005). An equivalent sensorimotor coupling reported in mouse visual cortex in the primate could involve the coupling of foveal and extrafoveal processing via saccadic eye movements, allowing natural vision to be conceived as an active sensing of the environment (Ahissar and Assa, 2016; Gallant et al., 1998; Huber-Huber et al., 2021; Ito et al., 2022; Schroeder et al., 2010). In the framework of active sensing, top-down influences on the peripheral representation of the early visual areas can be related to modulation of excitability and stimulus evoked activity to a peripheral target (Barczak et al., 2019). In this respect, in addition to motor inputs, our results reveal inputs to peripheral representation from frontal working memory circuits in area 9/46, which is in conformity with recent electrophysiological findings in awake behaving monkey showing attentional and working memory effects on processing in area V1 (van Kerkoerle et al., 2017).

## Conclusion

A large body of work shows the importance of quantitative, weighted data in the elaboration of large-scale models of the cerebral cortex and its spatially embedded nature (Bassett and Sporns, 2017; Beul et al., 2017; Burt et al., 2018; Chaudhuri et al., 2015; Ercsey-Ravasz et al., 2013; Froudist-Walsh et al., 2021; Goulas et al., 2019; Markov et al., 2013; Roberts et al., 2017; Theodoni et al., 2021). The present study provides evidence that connection weights of cortical projections to areas V1 and V2 are profoundly influenced by representation of the visual field. We anticipate that this data, fully available in the Supplementary Information, will allow the elaboration of improved computational models that will significantly enhance our understanding of the network properties of the primate cortical connectome.

## Author contributions

Surgery MW, PM, CL, CD, ZS, HK; Data acquisition, MW, LM, PM, YX, CL, AF, QZ, ZS, HK; Data curation MW, YH, PM, CL, ZS; Data analysis MW, YH, LM, JAA, PM, TC, ER, CL, MFG, TH, DVE, KK, ZS, HK; software MW, SH, TC, TH, ZS; Visualization MW, YH; Supervision TH, DVE, KK, ZS, HK; Proposed the project MMP, CD, ZS, HK; Experimental design MFG, TH, KK, DVE, ZS, HK; wrote the first draft HK, with assistance from MFG, TH, DVE, KK; writing-review editing, all authors.

## Acknowledgments

HK thanks Martin Vinck for early discussion. This study is supported by a grant DUAL_STREAMS ANR-19-CE37-0025 (K.K.); LABEX CORTEX ANR-11-LABX-0042, Université de Lyon ANR-11-IDEX-0007 (H.K.); CORTICITY ANR-17-HBPR-0003 (H.K.); INSERM IRP CORTICAL_CONNECTOME (H.K., C.D.); NIH R01 MH60974 (DCVE, MFG); Shanghai Municipal Science and Technology Major Project 2018SHZDZX05 (M.-M.P); International Partnership Program of Chinese Academy of Sciences 153D31KYSB20170059 (M.-M.P); the Scientific Instrument Developing Project of CAS YJKYYQ20190052 (M.-M.P); Lingang Lab LG202104-02-04 (M.-M.P); Strategic Priority Research Program of Chinese Academy of Science, XDB32070100; Brain/MINDS-beyond from Japan Agency of Medical Research and Development (AMED) JP22dm0307006 (T.H., J.A.A.).

## STAR Method

The full set of 21 injection sites listed in Table S1 have been mapped to the atlas surface and are shown in Figure 1. The extent of each injection and its relation to the borders of the subregion injected are shown in line drawings in Supplementary Figure S1. Examples of connectivity following injections in different subdivisions are shown in Figure S2.

### Surgery and histology

Surgical procedures were in accordance with European requirements 86/609/EEC and approved by the appropriate veterinary and ethical bodies. All the experiments carried out in IoN were approved by the Ethics Committee of the Institute of Neuroscience, Chinese Academy of Science. Experiments were carried out in cynomolgus (Macaca fascicularis). A total of 21 injections were performed in 15 monkeys (Table S1).

Following premedication with atropine (1.25 mg, i.m.) and dexamethasone (4 mg, i.m.), monkeys were prepared for surgery under ketamine hydrochloride (20 mg/kg, i.m.) and chlorpromazine (2 mg/kg, i.m.). Anesthesia was continued with halothane or 0.5 - 2% isoflurane in N2O/O2 (70/30).

### Tracer leakage with deep injections

Characterizing the effect of retinotopic representation on connectivity profiles via tracer injections in defined subregions of areas V1 and V2 is challenging. It involves deep injections where contamination resulting from leak of tracer can obscure the predicted dependency of topography on eccentricity. Single injections of Diamidino Yellow (DY) (2 - 3% in water), Fast Blue (FsB) (2 - 3% in water) or Cholera toxin subunit B conjugated with Alexa Fluor (Life Technologies) (1-3% in PBS) were made using Hamilton syringes. In cases C027, C054 and C060, tracers were injected with glass pipettes (100 - 200 um diameter). In brains M121, M122, M101, M103, M146, M148, M096 and M097, stereotypic elongated injections (1 to 4mm) were made in the cortical gray matter; tracer was delivered at regular intervals as the needle was retracted thereby aiming to leave a bolus of tracer in the cortex. For the other cases, the injections were performed in an angle that is perpendicular to the cortical surface. The volume of tracer that has been injected in each case was listed in Table S1. Injection sites are shown in Figures S1. Injection of areas in non-exposed cortical regions was aided by an image-guided stereotaxic system (Brainsight® Frameless, Rogue Research Inc.) (Frey et al., 2004). The target area was identified on TI-weighted (T1w) MRI 1.5-Tesla scans or MRI 3-Tesla (Prisma Siemens) acquired at 0.5 mm isotropic resolution. Visualization of sulcal landmarks in a 3D reconstruction of the monkey brain allowed optimization of injection trajectory.

After a survival period of 11-13 days, animals were deeply anesthetized before being perfused with 4-8% paraformaldehyde / 0.05% glutaraldehyde in phosphate buffer (0.1 M, pH=7.4). Cryoprotection was ensured by sucrose (Kennedy and Bullier, 1985) or glycerol perfusion gradients (Rosene et al., 1986). Brains were removed, cut coronally into anterior and posterior halves, kept in the cryoprotecting liquid overnight or until sinking. Coronal 40µm thick sections were cut on a freezing microtome. Every sixth section was mounted on gelatinized glass slides and used to explore projection pathways, and subsequently stained with cresyl violet after the neurons were mapped.

### Charting labeled neurons and reconstruction of injection sites

Injection sites and their relationship to areal borders and underlying white matter are shown in line drawings of multiple sections through each injection site (Figure S1).

Sections with fluorescent labeling were digitized with Axioscan Z1 (Zeiss) using a 10x objective (Plan-Apochromat 10×0.45) and a fluorescence camera (Carera set Orca Flash 4.0) exploiting a LED light source Colibri 7 (channels -R[G/Y]B-UV) and filter set 43 HE (CTB-555), filter set 50 HE (CTB-647) and a customized long pass filter lp465 nm / dichroic mirror dc455 nm (FB/DY). Connectivity was examined throughout the whole hemisphere injected. Neurons were plotted with Mercator (ExploraNova) or customized software PlotFast.

### Quantification of connection weights and laminar distribution

When exploring for labeled neurons, 1 section in 2 (i.e. at 240 μm intervals) was examined throughout cortical gray matter, which enabled the identification of many new-found projections. For each injection, the number of labeled neurons in a given source area relative to the total number of labeled neurons in the brain (including those in the injected area) defines the fraction of labeled neurons (FLN) of the source area (Falchier et al., 2002; Markov et al., 2014a; Markov et al., 2011). The extrinsic FLN (FLNe) of an area is estimated from the number of labeled neurons in that area relative to the total number of labeled neurons less the neurons intrinsic to the injected area. We quantified the laminar distribution of labeled neurons with respect to the counts of labeled neurons through-out the projection zone i.e. the region containing the full complement of labeled neurons in a given source area following the injection to the target area. This allows defining the percentage of supragranular layer neurons (SLN) which corresponds to the total number of supragranular layer neurons divided by the sum of supra-plus infragranular layer neurons (Markov et al., 2014a). A SLN of 100% indicates all parent neurons are located in the supragranular layers and a SLN of 0 indicates that all parent neurons are located in the infragranular layers. SLN values above 50% characterize FF projections and SLN values below 50% FB projections (Markov et al., 2014a).

### Generation of the M150 106-parcel macaque atlas

We generated an updated macaque surface-based cortical atlas using a new anatomical substrate (the M150 right hemisphere) and a significantly revised parcellation that is based on the (Markov et al., 2014a) M132 91-area parcellation but includes a finer-grained parcellation of subdivisions of early retinotopic areas into multiple sub-parcels.

### Reconstruction of M150 surface

This was generated from a female M. fascicularis (5.2 kg). An in vivo T1w structural MRI scan was obtained using a flexible coil (Flex-Small, Siemens) and a simple coil “L11”and a 3-Tesla MRI scanner (MAGNETOME Prisma, Siemens Healthineers, Erlangen) at 0.5mm isotropic resolution using an MPRAGE sequence (0.5 mm isotropic resolution, TR=3500ms, TI=1050ms, TE=3.8ms, FA=8 degree, bandwidth=130Hz/pixel, number of average=4). An NHP_HCP pipeline was used to generate a cortical surface reconstruction (Autio et al., 2021; Donahue et al., 2016). Since the M150 MRI scan was a legacy style protocol (before optimization for HCP-style analysis), the NHP-HCP pipeline was customized to allow surface reconstruction using the T1w volume alone, instead of using both T1w and T2w volumes in the recommended protocol (Autio et al., 2020). The biasfield of the T1w volume was corrected by presuming a strong bias in ‘fsl_anat’ in FSL (version 6.0.4) before reconstructing the cortical grey/white matter surface (white surface) and cortical outer surface (pial surface) using FreeSurfer version 5.3-HCP plus a midthickness surface midway between pial and white surfaces. Cortical surfaces were registered to a ‘sulc’ (sulcal depth) template of cynomolgus using the MSMSulc algorithm (Robinson et al., 2014). Prior to the surface registration, the cynomolgus sulc template was created by averaging sulc maps from multiple subjects (N=24), using data collected in an HCP-style study, NHP_NNP (Hayashi et al., 2021). Surfaces were reconstructed and registered with MSMSulc to the Yerke19_v1.2 template as described previously (Autio et al., 2020). Finally, surfaces were resampled using Connectome Workbench (‘Workbench’; version 1.5.0) to the 164k and 32k fs_LR standard meshes for the macaque (Donahue et al., 2016) with mean spacing of 0.26 and 0.57 mm in average midthickness surfaces, respectively.

### Retinotopic subdivisions

The retinotopic maps of V1 from Figures 4, 10, and 11 of Van Essen et al. (Van Essen et al., 1984) were used to delineate contours indicating the estimated horizontal meridian and isoeccentricity contours for 3 and 8 degrees on the Yerkes19_v1.2 template surface. V1 was subdivided into 5 sectors: V1c includes the central 3 degrees but does not include separate upper vs lower field sectors because these were not accurately mapped in the original study; V1pcLF and V1pcUF include the paracentral representations from 3 to 8 degrees for the lower field and upper field, respectively; V1fpLF and V1fpUF include the peripheral representations exceeding 8 degrees for the lower field and upper field, respectively. These were transferred to the M150 surface using the MSMSulc registration to the SpecMac24cyno template, which is also registered to the Yerkes19_v1.2 template as described above.

Corresponding maps of V2c, V2pcLF, V2pcUF, V2fpLF, and V2fpUF and V3c, V3pcLF, V3pcUF, V3fpLF, and V3fpUF were delineated on the M150 flatmap surface using folding-based landmarks and the retinotopic maps and other information in Figs. 2, 5 in (Zhu and Vanduffel, 2019), and Figs 4 and 5 in (Burkhalter et al., 1986), and (Olavarria and VanEssen, 1997). We also used tracer injections in V2pcUF and V2fpUF from case M148 that were registered to Yerkes19 (and thus to M150) as additional constraints on the borders between V2 and V3. For area V4 we identified a central representation, V4c and separate lower-field (V4LF) and upper-field (V4UF) subdivisions based on (Gattass et al., 1988; Zhu and Vanduffel, 2019). For area MT we identified separate central and peripheral representations based on (Gattass et al., 1988; Kolster et al., 2009).

### Calculation of interareal distances

The interareal distance of cortico-cortical connectivity cannot be directly measured by retrograde neuronal tracers since the routes in the white matter tracts are rarely stained. Previous work estimated the interareal distances by measuring the shortest physical distance between geometric centers (of areas) through the white matter, thereby approximating axonal trajectories between areas (Ercsey-Ravasz et al., 2013). In the current study, we refined our estimates of interareal distances by applying the current M150 cortical areal parcellations of 106-area including visual eccentricity to high-resolution, cortical surface-based diffusion tractography in a single cynomolgus monkey. The diffusion data was collected in an NHP study (Hayashi et al., 2021) with an HCP-style protocol using a 24-channel macaque head coil and an 3T MRI scanner (Autio et al., 2020). Probabilistic diffusion tractography was performed using a surface-based approach (Autio et al., 2020; Donahue et al., 2016) seeding from the greyordinates (matrix 1 in FSL’s probtrackx), allowing calculation of the counts and distances for the tracts between all the cortical surface vertices in both hemispheres with 32k meshes and the medial wall excluded (a 54k by 54k dense connectome). The dense distance map was parcellated by the 106-parcel-per-hemisphere M150 cortical areal labels (a total of 212 areas) using weighted average of the distances across vertices per parcel using the tract count as a weight (Donahue et al., 2016).

### Parcellation-free mapping of tracer density to atlas surface

In 2 monkeys (4 injections) we used the native brain as an intermediate step to map retrograde tracer density to the atlas surface. For these 2 monkeys, a T1w scan was available, a cortical surface reconstruction was generated and resampled to the Yerkes19_v1.2 template mesh (similar to above section ‘M150 surface’). We used the following approach to accurately match slices of the MRI scan to the corresponding histological slices and the associated pial/white surface contours drawn for each section. A Workbench scene file was generated that included side-by-side panels displaying section contours, histology section images (when available), and a coronal MRI slice (with pial, layer 4, and white matter surface contours displayed), starting with a slice close to the anterior commissure. The MRI coronal slice number was adjusted until the MRI slice and histological contours/image were reasonably well aligned. If needed, a slightly oblique MRI slice orientation was chosen to improve the alignment. The scene was saved, and the process was repeated for additional section/MRI-slice pairs 2 - 3 mm apart, with further fine tuning of the slice obliqueness until all regions were well matched for a fixed obliqueness.

Using this alignment between histology section contours and individual-subject surface contours, a custom Matlab-based GUI was used to manually match corresponding segments of the contours. The cortical ribbon plus closed subcortical contours (putamen, claustrum, and amygdala, when available) were flood-filled, and each section (plus associated labeled neurons) was initially brought into approximate alignment with the MRI-based surface contours and atlas-based subcortical structures using non-shearing translation and rotation. For sections containing a physically separate portion (e.g., the temporal pole), the separate pieces were aligned independently. The elastix registration tool (Klein, 2010; Shamonin et al., 2013) was then used to align the histology contours to the surface contours and thereby to transform all retrogradely labeled neurons to their corresponding locations in the MRI slice. Neurons lying between the pial and white surfaces were categorized as cortical, and those below the white surface were categorized as subcortical. A custom script then computed labeled cortical and subcortical cell density (cells/mm^3) and mapped the cortical cell density to the individual’s surface using the 164k standard mesh.

### Statistical Analysis

All statistical analyses were performed using R (R Development Core Team, 2022). Comparisons of cumulated FLNe values between dorsal and ventral streams were performed with paired t-tests. To compare distances of dorsal and ventral areas to different visual field representations, a permutation test was performed in which distances were randomly permuted 10,000 times across visual field regions and the dorsal/ventral differences were compared with the distribution of permuted differences.

To obtain connectivity profiles, FLNe values were ordered from largest to smallest. Our previous studies (Markov et al., 2014a; Markov et al., 2011) demonstrated that the ordered FLNe values followed the order statistics for a lognormal distribution. We fit the predicted reversed cumulative distribution function to the data based on the mean and standard deviation of the log10 FLNe values. Confidence intervals and confidence bands were calculated based on a negative binomial distribution of the raw counts with dispersion parameter set to 7, that has been previously found to describe the data well (Markov et al., 2014a; Markov et al., 2011) and then normalized these values by the total counts to put them on the scale of the FLNe.

The Wilcoxon test was used to test the significance of differences between Spearman correlations of connectivity profiles of injections within and across V1 subdivisions.

### Analysis of human functional connectivity

We used resting-state fMRI data from the Human Connectome Project (HCP) to characterize functional connectivity patterns involving retinotopic subdivisions of V1 and their relation to dorsal and ventral stream areas in extrastriate cortex. Previously in humans, a V1 spatial regression model was used to map human visual areas outside of V1 with resting state functional connectivity (see Supplementary Methods Section 4.4 in (Glasser et al., 2016) applied to the 210 ‘parcellation’ (210P) subjects. The spatial regressors were defined solely within the confines of V1 (whose boundaries were mapped by a separate process) and then used to identify visuotopically organized higher visual areas based on multiple linear regressions (first a spatial multiple regression of the regressors into V1 resting state fMRI data to generate V1 timeseries for each regressor and then a temporal regression of these V1 timeseries into all of the resting state fMRI data, thereby producing spatial maps with information outside the confines of V1 - see Supplementary Methods Figure 9 in (Glasser et al., 2016)). The results of each spatial regressor represent the partial functional connectivity of that regressor to the rest of the brain while controlling for the other regressors (partial correlation functional connectivity is thought to better represent direct anatomical connectivity than does full correlation functional connectivity (Smith, 2021). For a parcellated analyses of full-rank partial correlation functional connectivity between auditory cortex and all other areas in the HCP_MMP1.0 180-area-per-hemisphere parcellation (Glasser et al., 2016) we used data from the 449 HCP-YA subjects that were previously published in (Glasser et al., 2018).

## Data availability

The datasets associated with the surface-based analyses shown in are available at: https://balsa.wustl.edu. All the FLN and SLN data of the injections is available at: https://core-nets.org

## Supplementary Information

Figure S1: Injection sites

Figure S2: Exemplar connection of representative injection of the subdivisions injected

Figure S3: The 106-area atlas (pdf)

Table S1: Animal and injections used in the present study. c: central representation; fpUF: far periphery upper field; fpLF: far periphery lower field; pcUF: paracentral upper field; pcLF: paracentral lower field; CTB-555/647: cholera toxin subunit B, conjugated with Alexa 555/647; F: female; M: male.

Table S2: FLN and SLN of all cases used in the study

Table S3: Inter-areal distances

Table S4: Index of abbreviations

## Notes

### Competing Interest Statement

The authors have declared no competing interest.

## References

Ahissar, E., and Assa, E. (2016). Perception as a closed-loop convergence process. Elife 5.

Amedi, A., Raz, N., Pianka, P., Malach, R., and Zohary, E. (2003). Early ‘visual’ cortex activation correlates with superior verbal memory performance in the blind. Nat Neurosci 6, 758–766.

Autio, J.A., Glasser, M.F., Ose, T., Donahue, C.J., Bastiani, M., Ohno, M., Kawabata, Y., Urushibata, Y., Murata, K., Nishigori, K., Yamaguchi, M., Hori, Y., Yoshida, A., Go, Y., Coalson, T.S., Jbabdi, S., Sotiropoulos, S.N., Kennedy, H., Smith, S., Van Essen, D.C., and Hayashi, T. (2020). Towards HCP-Style macaque connectomes: 24-Channel 3T multi-array coil, MRI sequences and preprocessing. Neuroimage 215, 116800.

Autio, J.A., Zhu, Q., Li, X., Glasser, M.F., Schwiedrzik, C.M., Fair, D.A., Zimmermann, J., Yacoub, E., Menon, R.S., Van Essen, D.C., Hayashi, T., Russ, B., and Vanduffel, W. (2021). Minimal specifications for non-human primate MRI: Challenges in standardizing and harmonizing data collection. NeuroImage 236, 118082.

Bakola, S., Passarelli, L., Huynh, T., Impieri, D., Worthy, K.H., Fattori, P., Galletti, C., Burman, K.J., and Rosa, M.G.P. (2017). Cortical Afferents and Myeloarchitecture Distinguish the Medial Intraparietal Area (MIP) from Neighboring Subdivisions of the Macaque Cortex. eNeuro 4.

Barczak, A., Haegens, S., Ross, D.A., McGinnis, T., Lakatos, P., and Schroeder, C.E. (2019). Dynamic Modulation of Cortical Excitability during Visual Active Sensing. Cell Rep 27, 3447–3459 e3443.

Bassett, D.S., and Sporns, O. (2017). Network neuroscience. Nat Neurosci 20, 353–364.

Bedny, M. (2017). Evidence from Blindness for a Cognitively Pluripotent Cortex. Trends Cogn Sci 21, 637–648.

Bedny, M., Pascual-Leone, A., Dodell-Feder, D., Fedorenko, E., and Saxe, R. (2011). Language processing in the occipital cortex of congenitally blind adults. Proc Natl Acad Sci U S A 108, 4429–4434.

Beierholm, U., Rohe, T., Ferrari, A., Stegle, O., and Noppeney, U. (2020). Using the past to estimate sensory uncertainty. Elife 9.

Beul, S.F., Barbas, H., and Hilgetag, C.C. (2017). A Predictive Structural Model of the Primate Connectome. Sci Rep 7, 43176.

Bjoertomt, O., Cowey, A., and Walsh, V. (2002). Spatial neglect in near and far space investigated by repetitive transcranial magnetic stimulation. Brain 125, 2012–2022.

Borra, E., and Rockland, K.S. (2011). Projections to early visual areas v1 and v2 in the calcarine fissure from parietal association areas in the macaque. Front Neuroanat 5, 35.

Bressler, S.L., and Menon, V. (2010). Large-scale brain networks in cognition: emerging methods and principles. Trends Cogn Sci 14, 277–290.

Bullier, J. (2006). What is Fed Back? In 23 Problems in Systems Neuroscience, J.L. van Hemmen, and T.J. Sejnowski, eds. (Oxford University Press USA), pp. 103–132.

Burkhalter, A., Felleman, D.J., Newsome, W.T., and Van Essen, D.C. (1986). Anatomical and physiological asymmetries related to visual areas V3 and VP in macaque extrastriate cortex. Vision Res 26, 63–80.

Burt, J.B., Demirtaş, M., Eckner, W.J., Navejar, N.M., Ji, J.L., Martin, W.J., Bernacchia, A., Anticevic, A., and Murray, J.D. (2018). Hierarchy of transcriptomic specialization across human cortex captured by structural neuroimaging topography. Nat Neurosci 21, 1251–1259.

Cappe, C., and Barone, P. (2005). Heteromodal connections supporting multisensory integration at low levels of cortical processing in the monkey. Eur J Neurosci 22, 2886–2902.

Chaudhuri, R., Knoblauch, K., Gariel, M.A., Kennedy, H., and Wang, X.J. (2015). A Large-Scale Circuit Mechanism for Hierarchical Dynamical Processing in the Primate Cortex. Neuron 88, 419–431.

Chen, C.H., Hu, J.M., Zhang, S.Y., Xiang, X.J., Chen, S.Q., and Ding, S.L. (2021). Rodent Area Prostriata Converges Multimodal Hierarchical Inputs and Projects to the Structures Important for Visuomotor Behaviors. Front Neurosci 15, 772016.

Clark, A. (2013). Whatever next? Predictive brains, situated agents, and the future of cognitive science. Behav Brain Sci 36, 181–204.

Deen, B., Saxe, R., and Bedny, M. (2015). Occipital cortex of blind individuals is functionally coupled with executive control areas of frontal cortex. J Cogn Neurosci 27, 1633–1647.

Donahue, C.J., Sotiropoulos, S., Jbabdi, S., Herandez-Ferandez, M., Behrens, T., Kennedy, H., Knoblauch, K., Coalson, T., Glasser, M., and Van Essen, D. (2016). Using diffusion tractography to predict cortical connection strength and distance: a quantitative comparison with tracers in the monkey. J Neurosci 36, 6758–6770.

Eckert, M.A., Kamdar, N.V., Chang, C.E., Beckmann, C.F., Greicius, M.D., and Menon, V. (2008). A cross-modal system linking primary auditory and visual cortices: evidence from intrinsic fMRI connectivity analysis. Hum Brain Mapp 29, 848–857.

Ercsey-Ravasz, M., Markov, N.T., Lamy, C., Van Essen, D.C., Knoblauch, K., Toroczkai, Z., and Kennedy, H. (2013). A predictive network model of cerebral cortical connectivity based on a distance rule. Neuron 80, 184–197.

Falchier, A., Clavagnier, S., Barone, P., and Kennedy, H. (2002). Anatomical evidence of multimodal integration in primate striate cortex. J Neurosci 22, 5749–5759.

Faye, E.E. (1984). Clinical low vision Vol 50 (Boston: Little, Brown).

Felleman, D.J., and Van Essen, D.C. (1991). Distributed hierarchical processing in the primate cerebral cortex. Cereb Cortex 1, 1–47.

Foxe, J.J., and Schroeder, C.E. (2005). The case for feedforward multisensory convergence during early cortical processing. Neuroreport 16, 419–423.

Frey, S., Comeau, R., Hynes, B., Mackey, S., and Petrides, M. (2004). Frameless stereotaxy in the nonhuman primate. Neuroimage 23, 1226–1234.

Friston, K. (2010). The free-energy principle: a unified brain theory? Nat Rev Neurosci 11, 127–138.

Froudist-Walsh, S., Bliss, D.P., Ding, X., Rapan, L., Niu, M., Knoblauch, K., Zilles, K., Kennedy, H., Palomero-Gallagher, N., and Wang, X.J. (2021). A dopamine gradient controls access to distributed working memory in the large-scale monkey cortex. Neuron.

Fu, K.M., Johnston, T.A., Shah, A.S., Arnold, L., Smiley, J., Hackett, T.A., Garraghty, P.E., and Schroeder, C.E. (2003). Auditory cortical neurons respond to somatosensory stimulation. J Neurosci 23, 7510–7515.

Gallant, J.L., Connor, C.E., and Van Essen, D.C. (1998). Neural activity in areas V1, V2 and V4 during free viewing of natural scenes compared to controlled viewing. Neuroreport 9, 85–90.

Galletti, C., and Fattori, P. (2018). The dorsal visual stream revisited: Stable circuits or dynamic pathways? Cortex 98, 203–217

Gamanut, R., Kennedy, H., Toroczkai, Z., Ercsey-Ravasz, M., Van Essen, D.C., Knoblauch, K., and Burkhalter, A. (2018). The Mouse Cortical Connectome, Characterized by an Ultra-Dense Cortical Graph, Maintains Specificity by Distinct Connectivity Profiles. Neuron 97, 698–715 e610.

Garner, A.R., and Keller, G.B. (2022). A cortical circuit for audio-visual predictions. Nat Neurosci 25, 98–105.

Gattass, R., Sousa, A.P.B., and Gross, C.G. (1988). Visuotopic organization and extend of V3 and V4 of the macaque. J Neurosci 8, 1831–1845.

Gau, R., Bazin, P.L., Trampel, R., Turner, R., and Noppeney, U. (2020). Resolving multisensory and attentional influences across cortical depth in sensory cortices. Elife 9.

Ghazanfar, A.A., and Schroeder, C.E. (2006). Is neocortex essentially multisensory? Trends Cogn Sci 10, 278–285.

Giard, M.H., and Peronnet, F. (1999). Auditory-visual integration during multimodal object recognition in humans: a behavioral and electrophysiological study. J Cogn Neurosci 11, 473–490.

Gilbert, C.D., and Li, W. (2013). Top-down influences on visual processing. Nat Rev Neurosci 14, 350–363.

Glasser, M.F., Coalson, T.S., Bijsterbosch, J.D., Harrison, S.J., Harms, M.P., Anticevic, A., Van Essen, D.C., and Smith, S.M. (2018). Using temporal ICA to selectively remove global noise while preserving global signal in functional MRI data. Neuroimage 181, 692–717.

Glasser, M.F., Coalson, T.S., Robinson, E.C., Hacker, C.D., Harwell, J., Yacoub, E., Ugurbil, K., Andersson, J., Beckmann, C.F., Jenkinson, M., Smith, S.M., and Van Essen, D.C. (2016). A multi-modal parcellation of human cerebral cortex. Nature 536, 171–178.

Goodale, M.A., and Milner, A.D. (1992). Separate visual pathways for perception and action. Trends Neurosci 15, 20–25.

Goodale, M.A., Milner, A.D., Jakobson, L.S., and Carey, D.P. (1991). A neurological dissociation between perceiving objects and grasping them. Nature 349, 154–156.

Goulas, A., Majka, P., Rosa, M.G.P., and Hilgetag, C.C. (2019). A blueprint of mammalian cortical connectomes. PLoS Biol 17, e2005346.

Gregory, R.L. (1997). Knowledge in perception and illusion. Philos Trans R Soc Lond B Biol Sci 352, 1121–1127.

Hasson, U., Andric, M., Atilgan, H., and Collignon, O. (2016). Congenital blindness is associated with large-scale reorganization of anatomical networks. Neuroimage 128, 362–372.

Hayashi, T., Hou, Y., Glasser, M.F., Autio, J.A., Knoblauch, K., Inoue-Murayama, M., Coalson, T., Yacoub, E., Smith, S., Kennedy, H., and Van Essen, D.C. (2021). The nonhuman primate neuroimaging and neuroanatomy project. Neuroimage 229, 117726.

Henschke, J.U., Noesselt, T., Scheich, H., and Budinger, E. (2015). Possible anatomical pathways for short-latency multisensory integration processes in primary sensory cortices. Brain Struct Funct 220, 955–977.

Horvat, S., Gamanut, R., Ercsey-Ravasz, M., Magrou, L., Gamanut, B., Van Essen, D.C., Burkhalter, A., Knoblauch, K., Toroczkai, Z., and Kennedy, H. (2016). Spatial Embedding and Wiring Cost Constrain the Functional Layout of the Cortical Network of Rodents and Primates. PLoS Biol 14, e1002512.

Hubel, D.H., and Wiesel, T.N. (1962). Receptive fields binocular interaction and functional architecture in the cat visual cortex. J Physiol 160, 106–154.

Huber-Huber, C., Buonocore, A., and Melcher, D. (2021). The extrafoveal preview paradigm as a measure of predictive, active sampling in visual perception. J Vis 21, 12.

Huberman, A.D., and Niell, C.M. (2011). What can mice tell us about how vision works? Trends Neurosci 34, 464–473.

Ito, J., Joana, C., Yamane, Y., Fujita, I., Tamura, H., Maldonado, P.E., and Grun, S. (2022). Latency shortening with enhanced sparseness and responsiveness in V1 during active visual sensing. Sci Rep 12, 6021.

Kanjlia, S., Loiotile, R.E., Harhen, N., and Bedny, M. (2021). ‘Visual’ cortices of congenitally blind adults are sensitive to response selection demands in a go/no-go task. Neuroimage 236, 118023.

Keller, G.B., and Mrsic-Flogel, T.D. (2018). Predictive Processing: A Canonical Cortical Computation. Neuron 100, 424–435.

Kennedy, H., and Bullier, J. (1985). A double-labeling investigation of the afferent connectivity to cortical areas V1 and V2 of the macaque monkey. J Neurosci 5, 2815–2830.

Kennedy, H., Knoblauch, K., and Toroczkai, Z. (2013). Why data coherence and quality is critical for understanding interareal cortical networks. Neuroimage 80, 37–45.

Klein, S., Staring, M., Murphy, K., Viergever, M. A., Pluim, J.P.W. (2010). Elastix: a toolbox for intensity based medical image registration. IEEE Transactions on medical imaging 29, 196–205.

Klink, P.C., Dagnino, B., Gariel-Mathis, M.A., and Roelfsema, P.R. (2017). Distinct Feedforward and Feedback Effects of Microstimulation in Visual Cortex Reveal Neural Mechanisms of Texture Segregation. Neuron 95, 209–220 e203.

Knoblauch, K., Ercsey-Ravasz, M., Kennedy, H., and Toroczkai, Z. (2016). The brain in space. In Micro-, meso-and macro-connectomics of the brain, H. Kennedy, D. Van Essen, and Y. Christen, eds. (Springer, Heidelberg), pp. 45–74.

Kolster, H., Mandeville, J.B., Arsenault, J.T., Ekstrom, L.B., Wald, L.L., and Vanduffel, W. (2009). Visual field map clusters in macaque extrastriate visual cortex. J Neurosci 29, 7031–7039.

Kravitz, D.J., Saleem, K.S., Baker, C.I., and Mishkin, M. (2011). A new neural framework for visuospatial processing. Nat Rev Neurosci 12, 217–230.

Kravitz, D.J., Saleem, K.S., Baker, C.I., Ungerleider, L.G., and Mishkin, M. (2013). The ventral visual pathway: an expanded neural framework for the processing of object quality. Trends Cogn Sci 17, 26–49.

Kupers, R., and Ptito, M. (2014). Compensatory plasticity and cross-modal reorganization following early visual deprivation. Neurosci Biobehav Rev 41, 36–52.

Kurzawski, J.W., Mikellidou, K., Morrone, M.C., and Pestilli, F. (2020). The visual white matter connecting human area prostriata and the thalamus is retinotopically organized. Brain Struct Funct 225, 1839–1853.

Lane, A.R., Ball, K., Smith, D.T., Schenk, T., and Ellison, A. (2013). Near and far space: Understanding the neural mechanisms of spatial attention. Hum Brain Mapp 34, 356–366.

Lee, T.S., and Mumford, D. (2003). Hierarchical Bayesian inference in the visual cortex. J Opt Soc Am A Opt Image Sci Vis 20, 1434–1448.

Levine, M.W., and McAnany, J.J. (2005). The relative capabilities of the upper and lower visual hemifields. Vision Res 45, 2820–2830.

Liang, M., Mouraux, A., Hu, L., and Iannetti, G.D. (2013). Primary sensory cortices contain distinguishable spatial patterns of activity for each sense. Nat Commun 4, 1979.

Linton, P. (2021). V1 as an egocentric cognitive map. Neurosci Conscious 2021, iab017.

Lu, W., Chen, S., Chen, X., Hu, J., Xuan, A., and Ding, S.L. (2020). Localization of area prostriata and its connections with primary visual cortex in rodent. J Comp Neurol 528, 389–406.

Macaluso, E., and Driver, J. (2005). Multisensory spatial interactions: a window onto functional integration in the human brain. Trends Neurosci 28, 264–271.

Majka, P., Rosa, M.G.P., Bai, S., Chan, J.M., Huo, B.X., Jermakow, N., Lin, M.K., Takahashi, Y.S., Wolkowicz, I.H., Worthy, K.H., Rajan, R., Reser, D.H., Wojcik, D.K., Okano, H., and Mitra, P.P. (2019). Unidirectional monosynaptic connections from auditory areas to the primary visual cortex in the marmoset monkey. Brain Struct Funct 224, 111–131.

Markov, N.T., Ercsey-Ravasz, M., Van Essen, D.C., Knoblauch, K., Toroczkai, Z., and Kennedy, H. (2013). Cortical high-density counter-stream architectures. Science 342, 1238406.

Markov, N.T., Ercsey-Ravasz, M.M., Ribeiro Gomes, A.R., Lamy, C., Magrou, L., Vezoli, J., Misery, P., Falchier, A., Quilodran, R., Gariel, M.A., Sallet, J., Gamanut, R., Huissoud, C., Clavagnier, S., Giroud, P., Sappey-Marinier, D., Barone, P., Dehay, C., Toroczkai, Z., Knoblauch, K., Van Essen, D.C., and Kennedy, H. (2014a). A weighted and directed interareal connectivity matrix for macaque cerebral cortex. Cereb Cortex 24, 17–36.

Markov, N.T., and Kennedy, H. (2013). The importance of being hierarchical. Curr Opin Neurobiol 23, 187–194.

Markov, N.T., Misery, P., Falchier, A., Lamy, C., Vezoli, J., Quilodran, R., Gariel, M.A., Giroud, P., Ercsey-Ravasz, M., Pilaz, L.J., Huissoud, C., Barone, P., Dehay, C., Toroczkai, Z., Van Essen, D.C., Kennedy, H., and Knoblauch, K. (2011). Weight Consistency Specifies Regularities of Macaque Cortical Networks. Cereb Cortex 21, 1254–1272.

Markov, N.T., Vezoli, J., Chameau, P., Falchier, A., Quilodran, R., Huissoud, C., Lamy, C., Misery, P., Giroud, P., Barone, P., Dehay, C., Ullman, S., Knoblauch, K., and Kennedy, H. (2014b). The Anatomy of Hierarchy: Feedforward and feedback pathways in macaque visual cortex. J Comp Neurol 522, 225–259.

Merriam, E.P., and Colby, C.L. (2005). Active vision in parietal and extrastriate cortex. Neuroscientist 11, 484–493.

Mikellidou, K., Kurzawski, J.W., Frijia, F., Montanaro, D., Greco, V., Burr, D.C., and Morrone, M.C. (2017). Area Prostriata in the Human Brain. Curr Biol 27, 3056–3060 e3053.

Mishkin, M., Ungerleider, L.G., and Macko, K.A. (1983). Object vision and spatial vision: two cortical pathways. Trends Neurosci 6, 414–417.

Murray, M.M., Thelen, A., Thut, G., Romei, V., Martuzzi, R., and Matusz, P.J. (2016). The multisensory function of the human primary visual cortex. Neuropsychologia 83, 161–169.

Niell, C.M., and Stryker, M.P. (2010). Modulation of visual responses by behavioral state in mouse visual cortex. Neuron 65, 472–479.

Olavarria, J.F., and VanEssen, D.C. (1997). The global pattern of cytochrome oxidase stripes in visual area V2 of the macaque monkey. Cerebral Cortex 7, 395–404.

Orban, G.A., Kennedy, H., and Bullier, J. (1986). Velocity sensitivity and direction selectivity of neurons in areas V1 and V2 of the monkey: influence of eccentricity. J Neurophysiol 56, 462–480.

Pakan, J.M., Francioni, V., and Rochefort, N.L. (2018). Action and learning shape the activity of neuronal circuits in the visual cortex. Curr Opin Neurobiol 52, 88–97.

Passingham, R.E., Stephan, K.E., and Kotter, R. (2002). The anatomical basis of functional localization in the cortex. Nat Rev Neurosci 3, 606–616.

Pitzalis, S., Fattori, P., and Galletti, C. (2012). The functional role of the medial motion area V6. Front Behav Neurosci 6, 91.

Poort, J., Khan, A.G., Pachitariu, M., Nemri, A., Orsolic, I., Krupic, J., Bauza, M., Sahani, M., Keller, G.B., Mrsic-Flogel, T.D., and Hofer, S.B. (2015). Learning Enhances Sensory and Multiple Non-sensory Representations in Primary Visual Cortex. Neuron 86, 1478–1490.

Previc, F.H. (1990). Functional specialization in the lower and upper visual fields in humans: Its ecological origins ans neurophysiological implications. Behav Brain Sci 13, 519–575.

R Development Core Team (2022). R: A language and environment for statistical computing. (Vienna, Austria., R Foundation for statistical computing. http://www.R-project.org).

Rao, R.P., and Ballard, D.H. (1999). Predictive coding in the visual cortex: a functional interpretation of some extra-classical receptive-field effects. Nat Neurosci 2, 79–87.

Roberts, J.A., Perry, A., Roberts, G., Mitchell, P.B., and Breakspear, M. (2017). Consistency-based thresholding of the human connectome. Neuroimage 145, 118–129.

Rockland, K.S., and Ojima, H. (2003). Multisensory convergence in calcarine visual areas in macaque monkey. Int J Psychophysiol 50, 19–26.

Rockland, K.S., and Pandya, D.N. (1979). Laminar origins and terminations of cortical connections of the occipital lobe in the rhesus monkey. Brain Res 179, 3–20.

Rohe, T., Ehlis, A.C., and Noppeney, U. (2019). The neural dynamics of hierarchical Bayesian causal inference in multisensory perception. Nat Commun 10, 1907.

Rohe, T., and Noppeney, U. (2015). Cortical hierarchies perform Bayesian causal inference in multisensory perception. PLoS Biol 13, e1002073.

Rosene, D.L., Roy, N.J., and Davis, B.J. (1986). A cryoprotection method that facilitates cutting frozen sections of whole monkey brains for histological and histochemical processing without freezing artifact. J Histochem Cytochem 34, 1301–1315.

Rossetti, Y., Pisella, L., and McIntosh, R.D. (2017). Rise and fall of the two visual systems theory. Ann Phys Rehabil Med 60, 130–140.

Saleem, A.B. (2020). Two stream hypothesis of visual processing for navigation in mouse. Curr Opin Neurobiol 64, 70–78.

Sanides, F. (1972). Representation in the cerebral cortex and its areal lamination patterns. In The structure and function of the nervous tissue, G.F. Bourne, ed. (New York and London: Academic Press), pp. 329–453.

Schroeder, C.E., Wilson, D.A., Radman, T., Scharfman, H., and Lakatos, P. (2010). Dynamics of Active Sensing and perceptual selection. Curr Opin Neurobiol 20, 172–176.

Shamonin, D.P., Bron, E.E., Lelieveldt, B.P., Smits, M., Klein, S., Staring, M., and Alzheimer’s Disease Neuroimaging, I. (2013). Fast parallel image registration on CPU and GPU for diagnostic classification of Alzheimer’s disease. Front Neuroinform 7, 50.

Sheth, B.R., and Young, R. (2016). Two Visual Pathways in Primates Based on Sampling of Space: Exploitation and Exploration of Visual Information. Front Integr Neurosci 10, 37.

Smith, A.T. (2021). Cortical visual area CVs as a cingulate motor area: a sensorimotor interfacefor the control of locomotion. Brain Struct Funct 226, 2931–2950.

Sousa, A.P., Pinon, M.C., Gattass, R., and Rosa, M.G. (1991). Topographic organization of cortical input to striate cortex in the Cebus monkey: a fluorescent tracer study. J Comp Neurol 308, 665–682.

Tamietto, M., and Leopold, D.A. (2018). Visual Cortex: The Eccentric Area Prostriata in the Human Brain. Curr Biol 28, R17–R19.

Theodoni, P., Majka, P., Reser, D.H., Wojcik, D.K., Rosa, M.G.P., and Wang, X.J. (2021). Structural Attributes and Principles of the Neocortical Connectome in the Marmoset Monkey. Cereb Cortex 32, 15–28.

Tong, F. (2003). Primary visual cortex and visual awareness. Nat Rev Neurosci 4, 219–229.

Ungerleider, L.G., and Mishkin, M. (1982). Two cortical visual systems. In Analysis of visual behavior, D.J. Ingle, M.A. Goodale, and R.J.R. Mansfield, eds. (Cambridge, Massachusetts: MIT Press), pp. 549–586.

van Atteveldt, N., Murray, M.M., Thut, G., and Schroeder, C.E. (2014). Multisensory integration: flexible use of general operations. Neuron 81, 1240–1253.

Van Essen, D.C., Newsome, W.T., and Maunsell, J.H. (1984). The visual field representation in striate cortex of the macaque monkey: asymmetries, anisotropies, and individual variability. Vision Res 24, 429–448.

van Kerkoerle, T., Self, M.W., and Roelfsema, P.R. (2017). Layer-specificity in the effects of attention and working memory on activity in primary visual cortex. Nat Commun 8, 13804.

van Wassenhove, V., Grant, K.W., and Poeppel, D. (2005). Visual speech speeds up the neural processing of auditory speech. Proc Natl Acad Sci U S A 102, 1181–1186.

Vetter, P., Bola, L., Reich, L., Bennett, M., Muckli, L., and Amedi, A. (2020). Decoding Natural Sounds in Early “Visual” Cortex of Congenitally Blind Individuals. Curr Biol 30, 3039–3044 e3032.

Vetter, P., Smith, F.W., and Muckli, L. (2014). Decoding sound and imagery content in early visual cortex. Curr Biol 24, 1256–1262.

Virsu, V., Rovamo, J., Laurinen, P., and Nasanen, R. (1982). Temporal contrast sensitivity and cortical magnification. Vision Res 22, 1211–1217.

Weiss, P.H., Marshall, J.C., Wunderlich, G., Tellmann, L., Halligan, P.W., Freund, H.J., Zilles, K., and Fink, G.R. (2000). Neural consequences of acting in near versus far space: a physiological basis for clinical dissociations. Brain 123, 2531–2541.

Yu, H.H., Chaplin, T.A., Davies, A.J., Verma, R., and Rosa, M.G. (2012). A specialized area in limbic cortex for fast analysis of peripheral vision. Curr Biol 22, 1351–1357.

Yu, H.H., Chaplin, T.A., and Rosa, M.G. (2015). Representation of central and peripheral vision in the primate cerebral cortex: Insights from studies of the marmoset brain. Neurosci Res 93, 47–61.

Yu, H.H., Verma, R., Yang, Y., Tibballs, H.A., Lui, L.L., Reser, D.H., and Rosa, M.G. (2010). Spatial and temporal frequency tuning in striate cortex: functional uniformity and specializations related to receptive field eccentricity. Eur J Neurosci 31, 1043–1062.

Zeki, S.M. (1978). Functional specialisation in the visual cortex of the rhesus monkey. Nature 274, 423–428.

Zhu, Q., and Vanduffel, W. (2019). Submillimeter fMRI reveals a layout of dorsal visual cortex in macaques, remarkably similar to New World monkeys. Proc Natl Acad Sci U S A 116, 2306–2311.

Zito, G.A., Cazzoli, D., Muri, R.M., Mosimann, U.P., and Nef, T. (2016). Behavioral Differences in the Upper and Lower Visual Hemifields in Shape and Motion Perception. Front Behav Neurosci 10, 128.

