## Supplementary material for "Retinotopic organization of feedback projections in primate early visual cortex: implications for active vision": Figure S1

#### Case 1: C054RH (V1c)

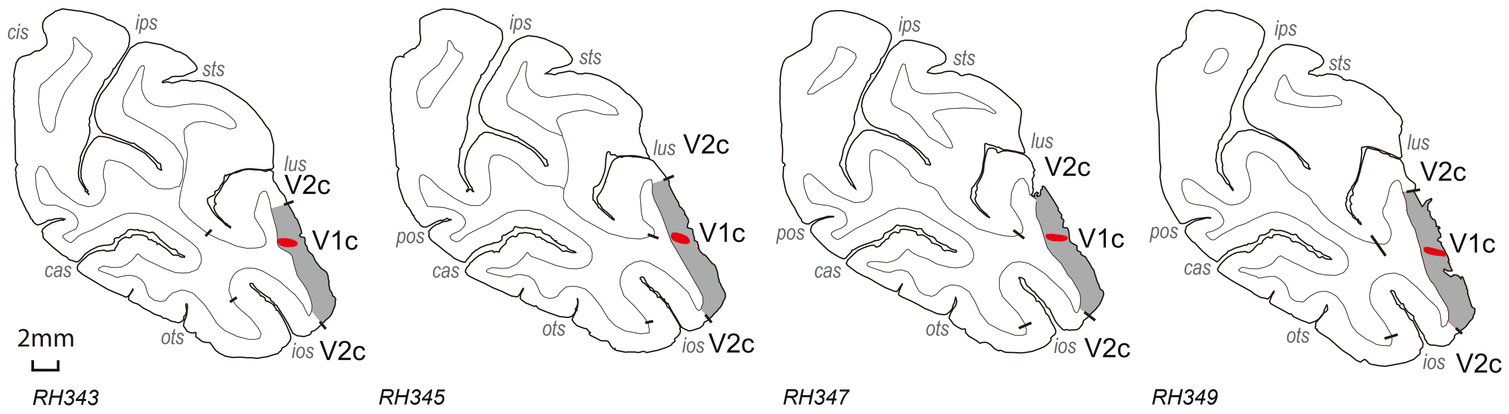


#### Case 2: C054RH (V1c)

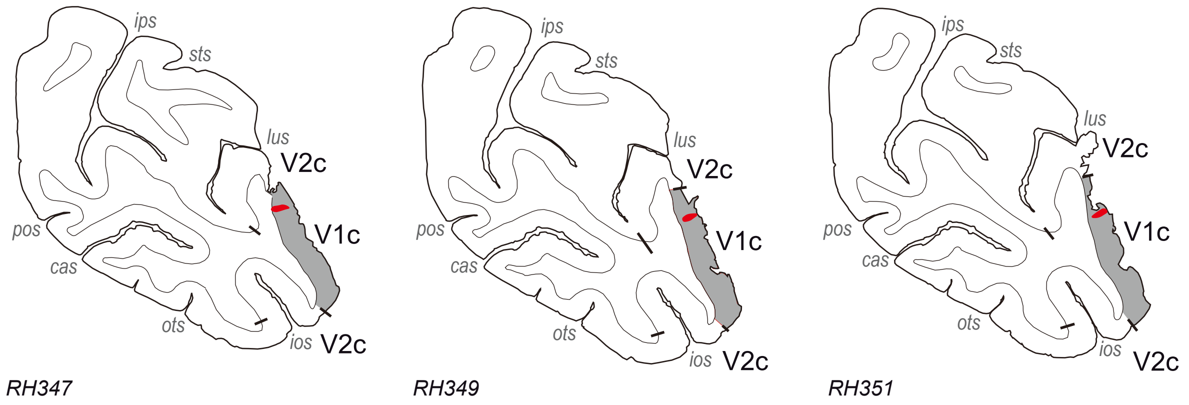


#### Case 3: C081RH (V1c)

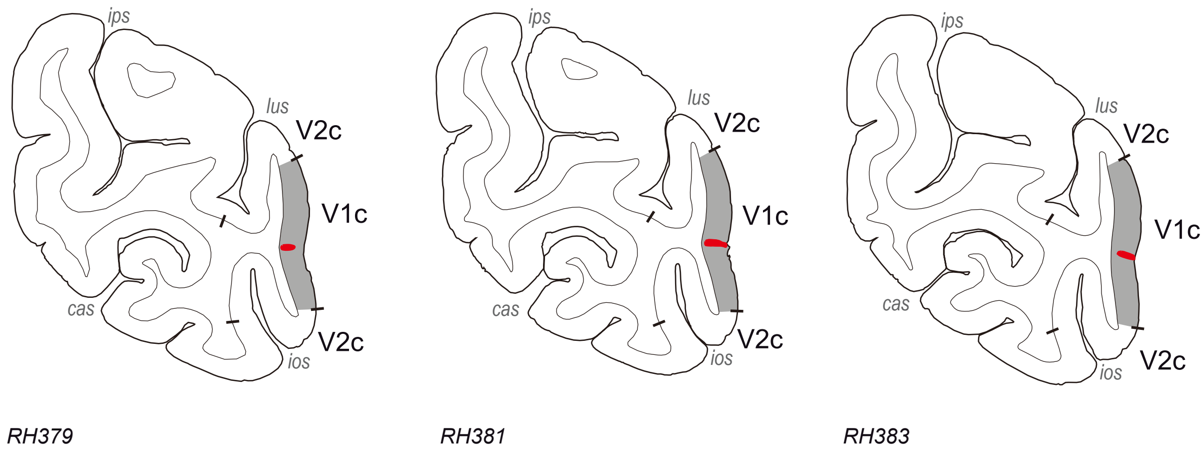


#### Case 4: M121RH (V1c)

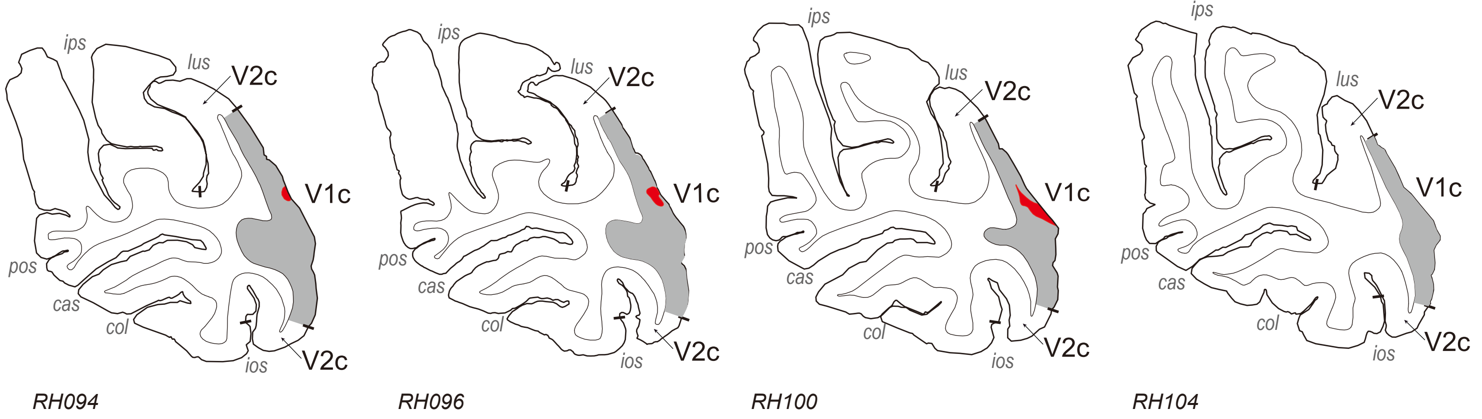

#### Case 5: C062RH (V1pcLF)

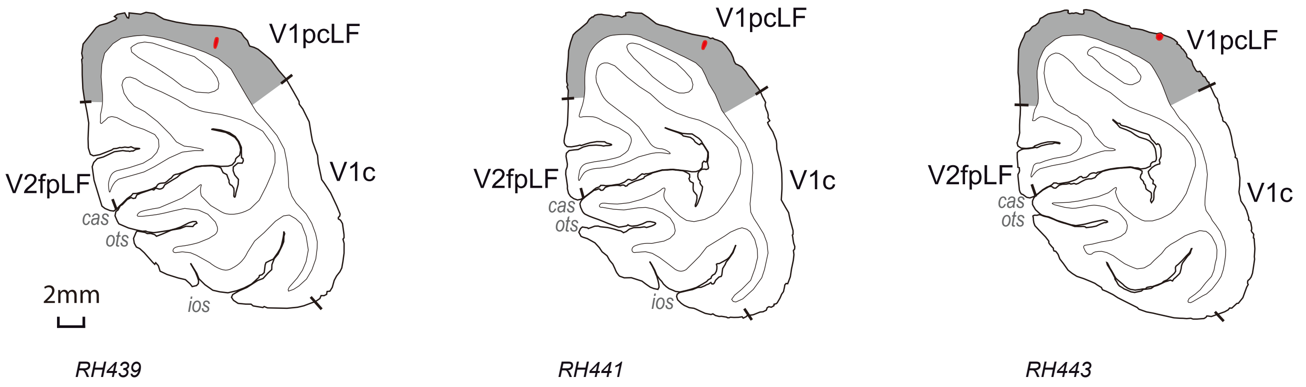


#### Case 6: M122RH (V1pcLF)

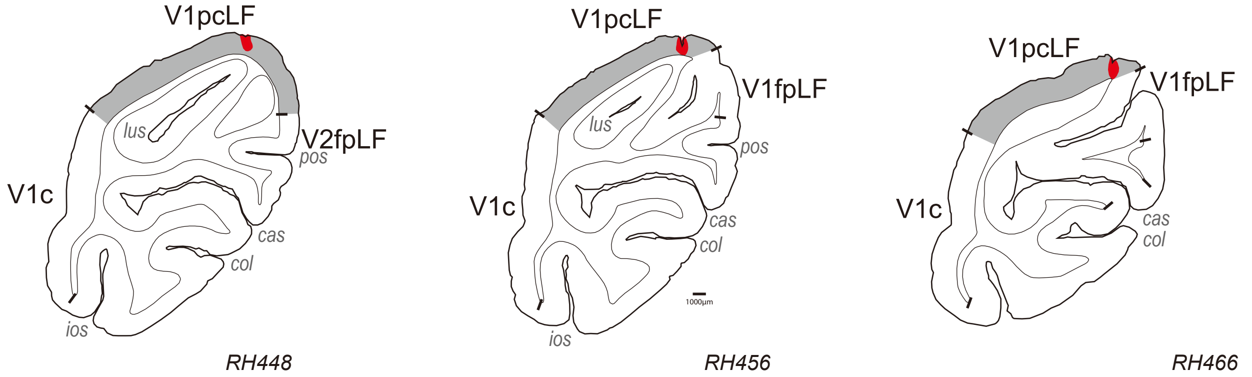


#### Case 7: C060RH (V1pcLF)

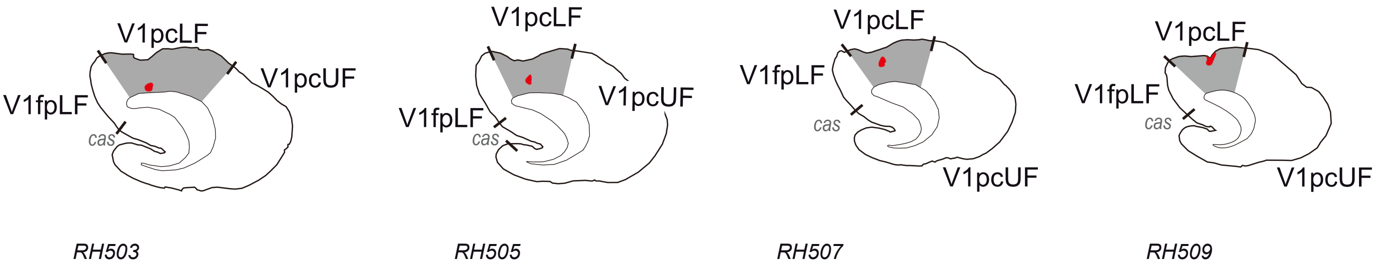


#### Case 8: C081RH (V1pcLF)

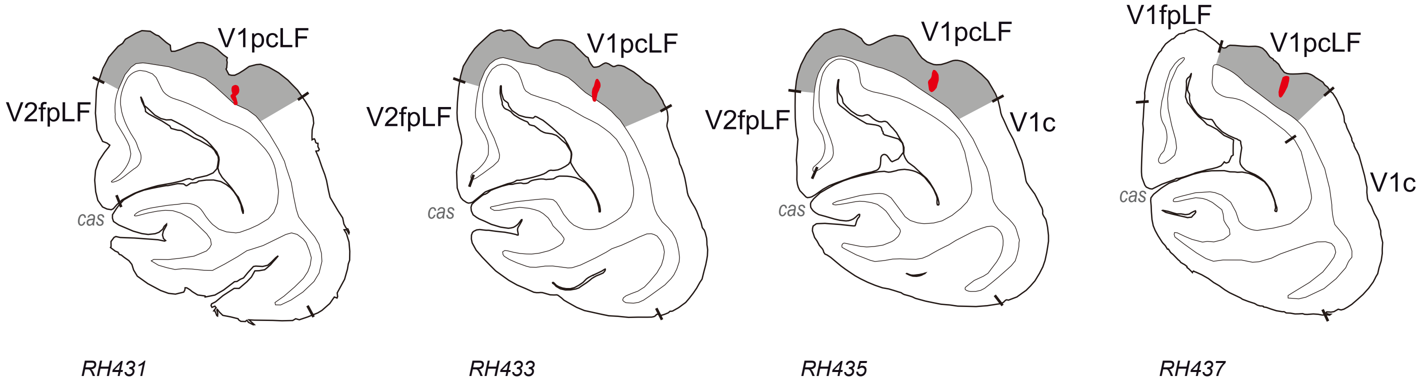

### Case 9: C027RH (V1fpLF)

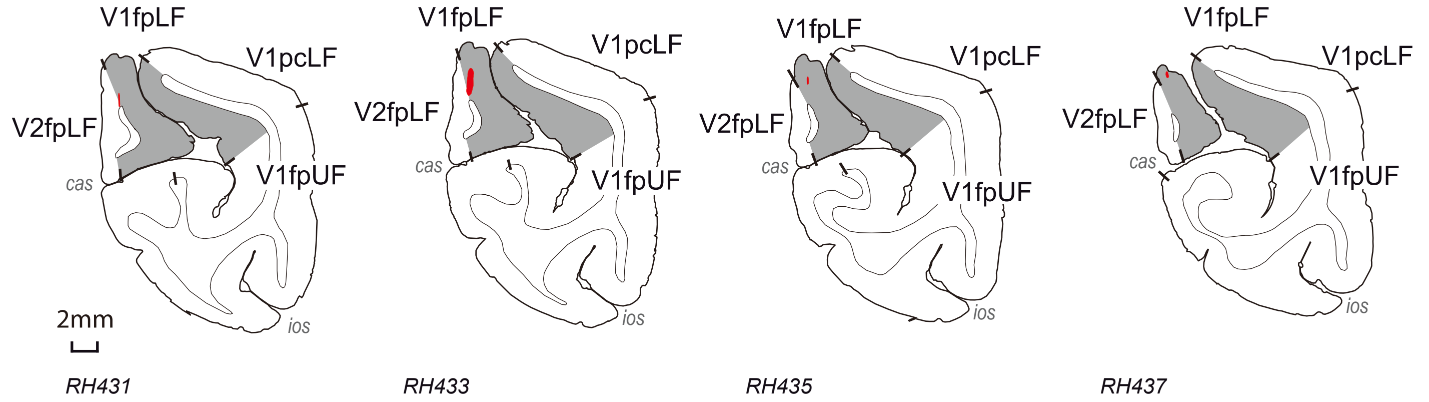


### Case 10: C054RH (V1fpLF)

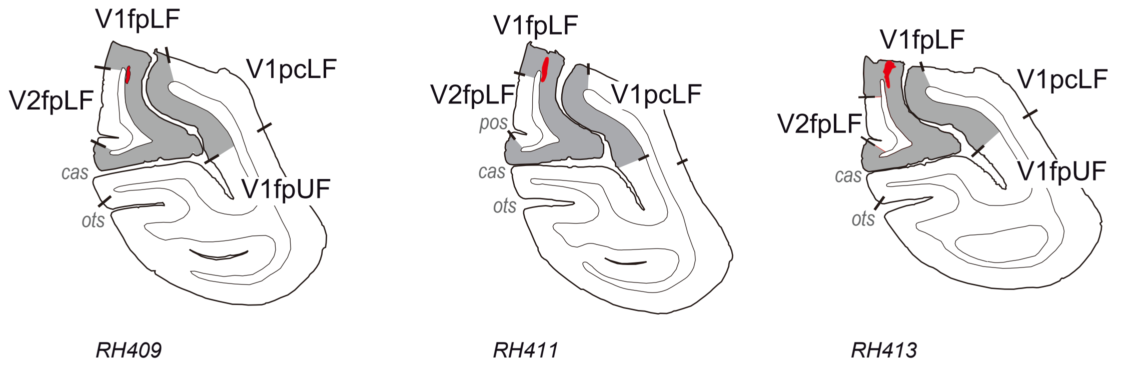


### Case 11: C081RH (V1fpLF)

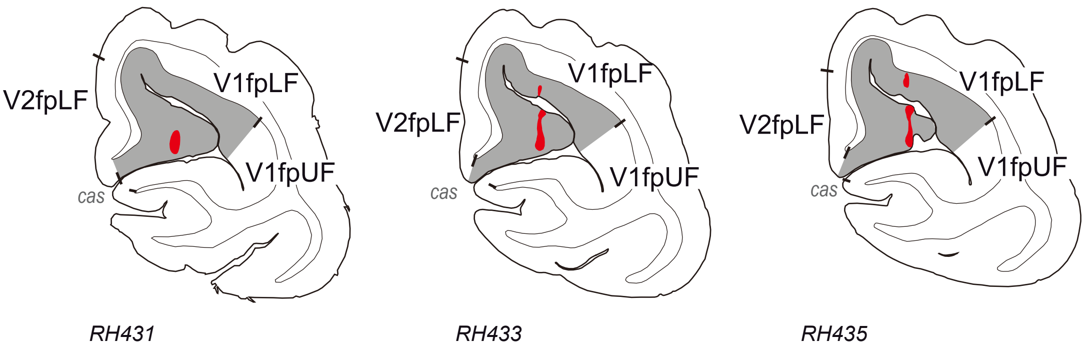


### Case 12: C057RH (V1fpUF)

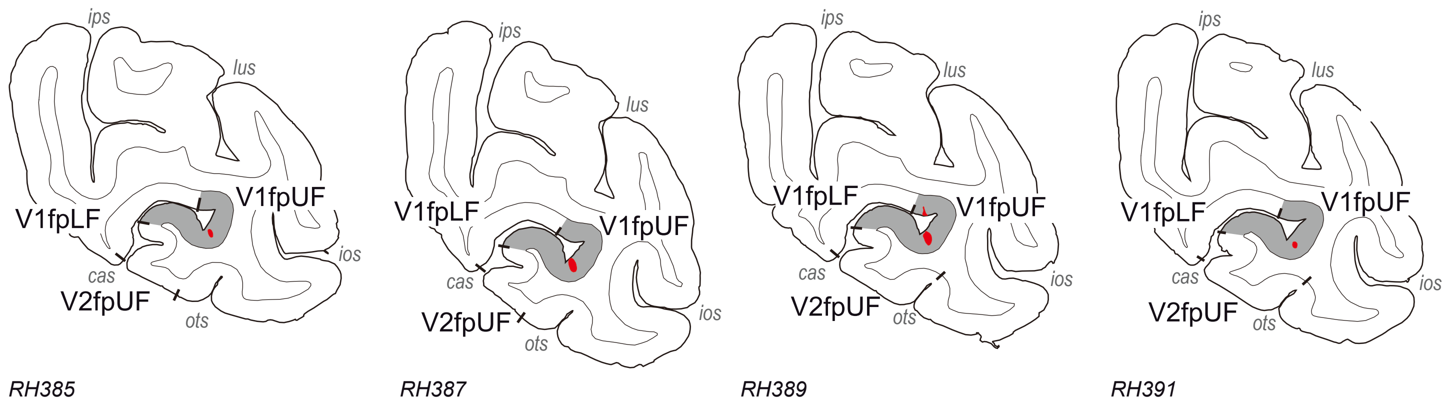

#### Case 13: C083RH (V1fpUF)

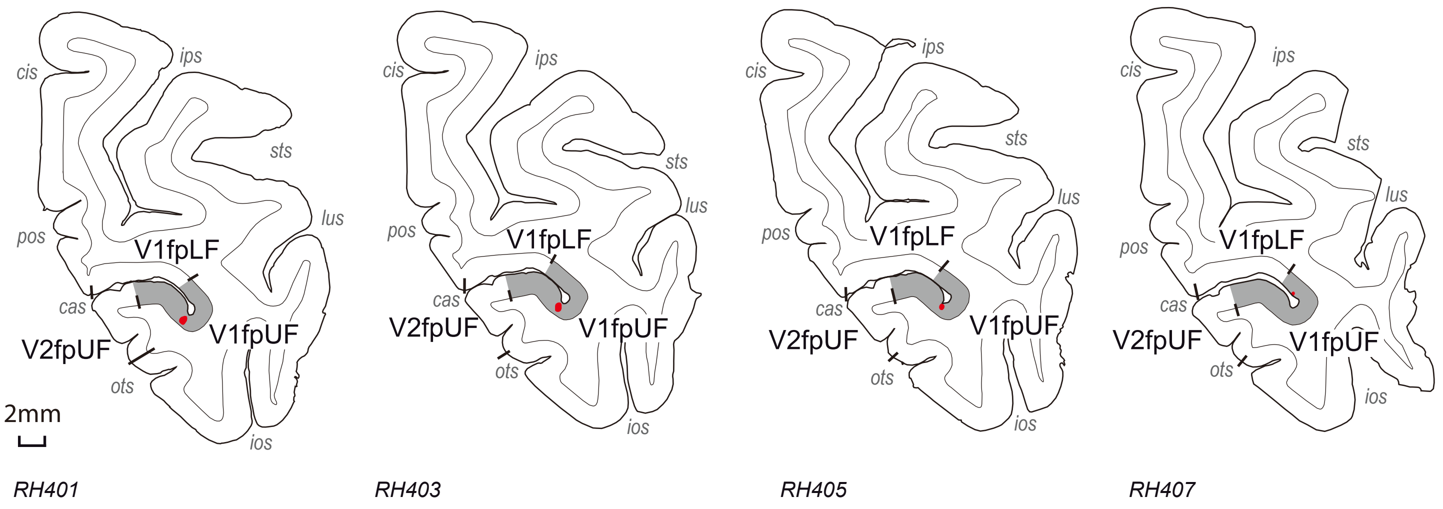


#### Case 14: C060RH (V1fpUF)

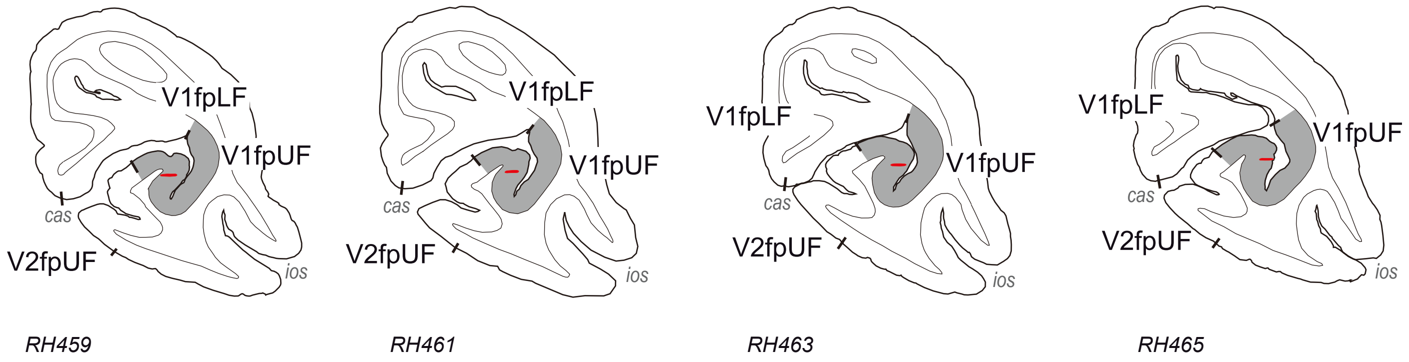


#### Case 15: M101LH (V2c)

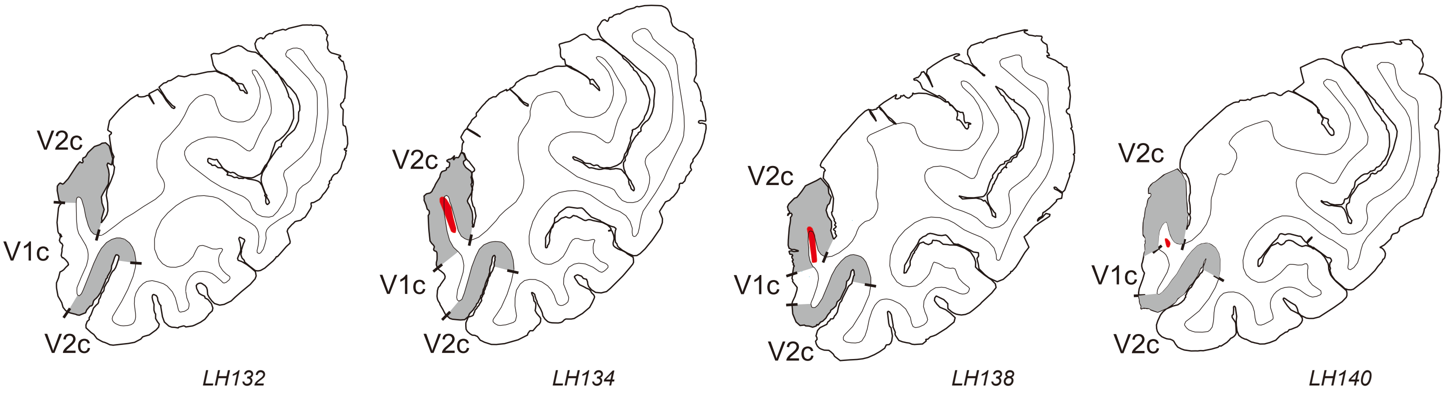


#### Case 16: M101RH (V2c)

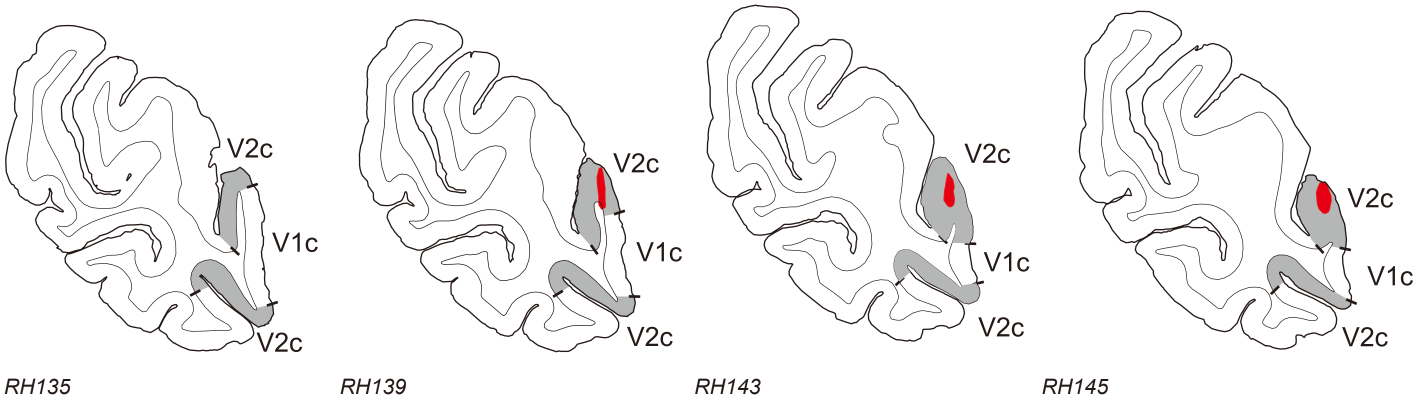

#### Case 17: M103LH (V2c)

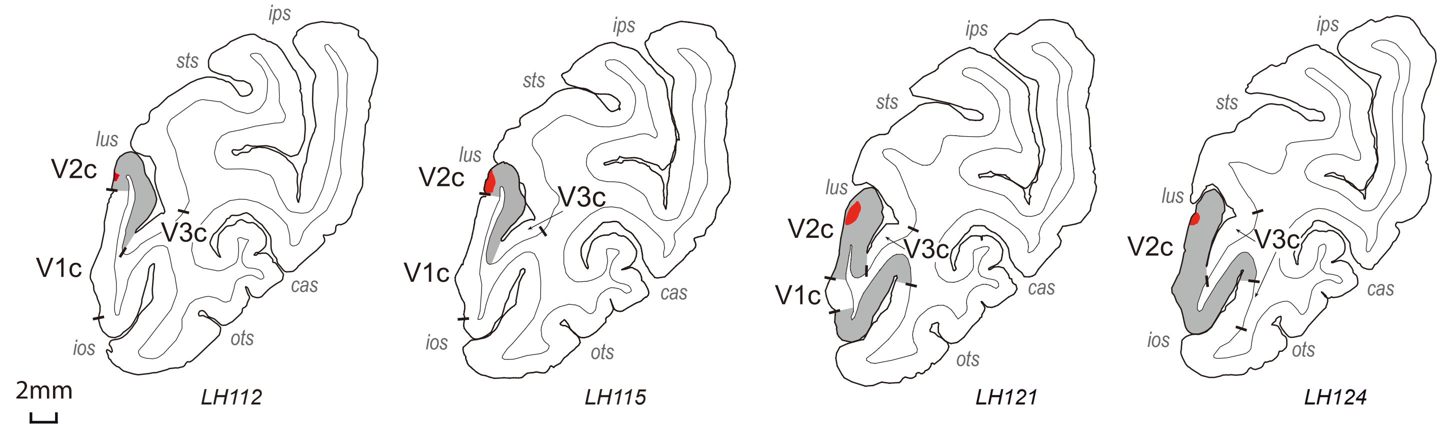


#### Case 18: M146LH (V2c)

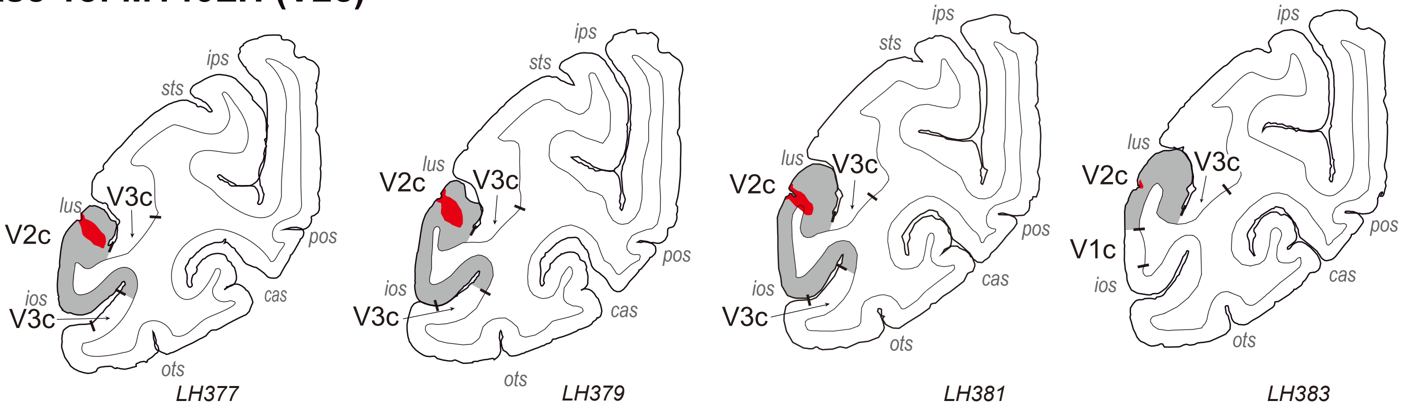


#### Case 19: M148RH (V2pcUF)

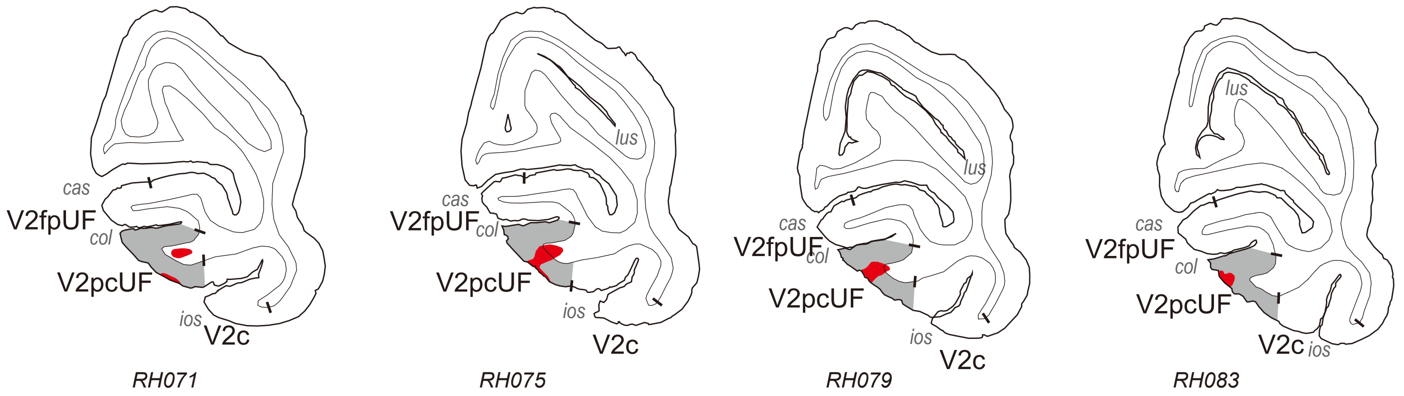


#### Case 20: M097LH (V2fpLF)

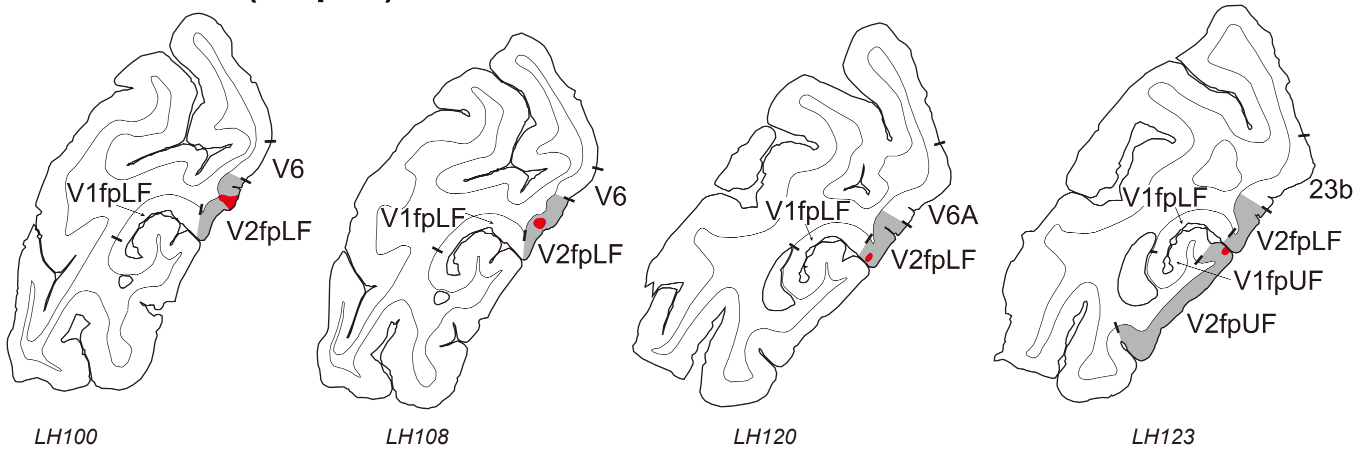

#### Case 21: M096LH (V2fpUF/LF)

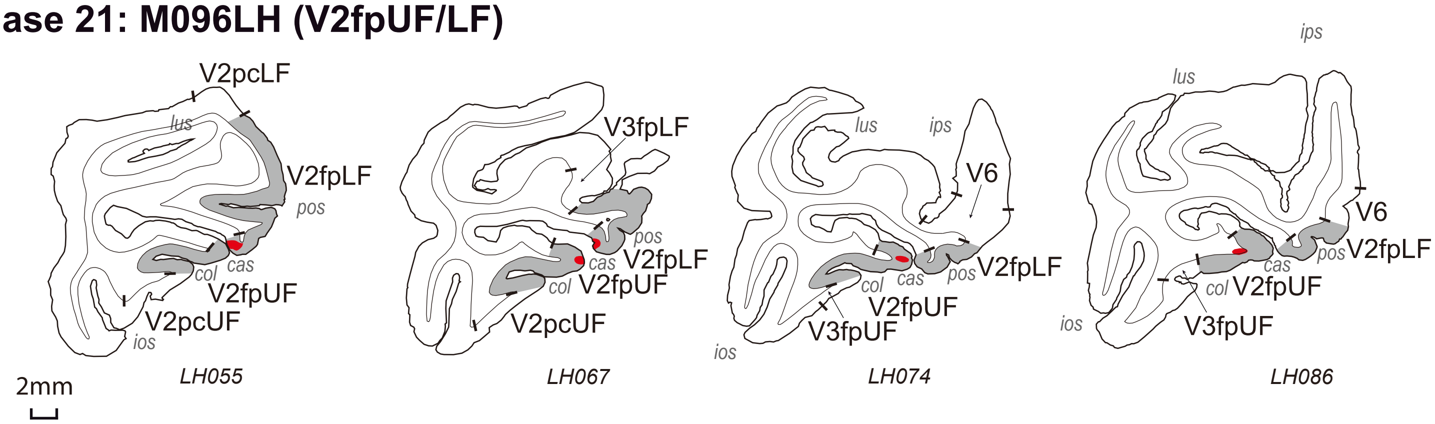
