## Supplementary material for "Retinotopic organization of feedback projections in primate early visual cortex: implications for active vision": Figure S2

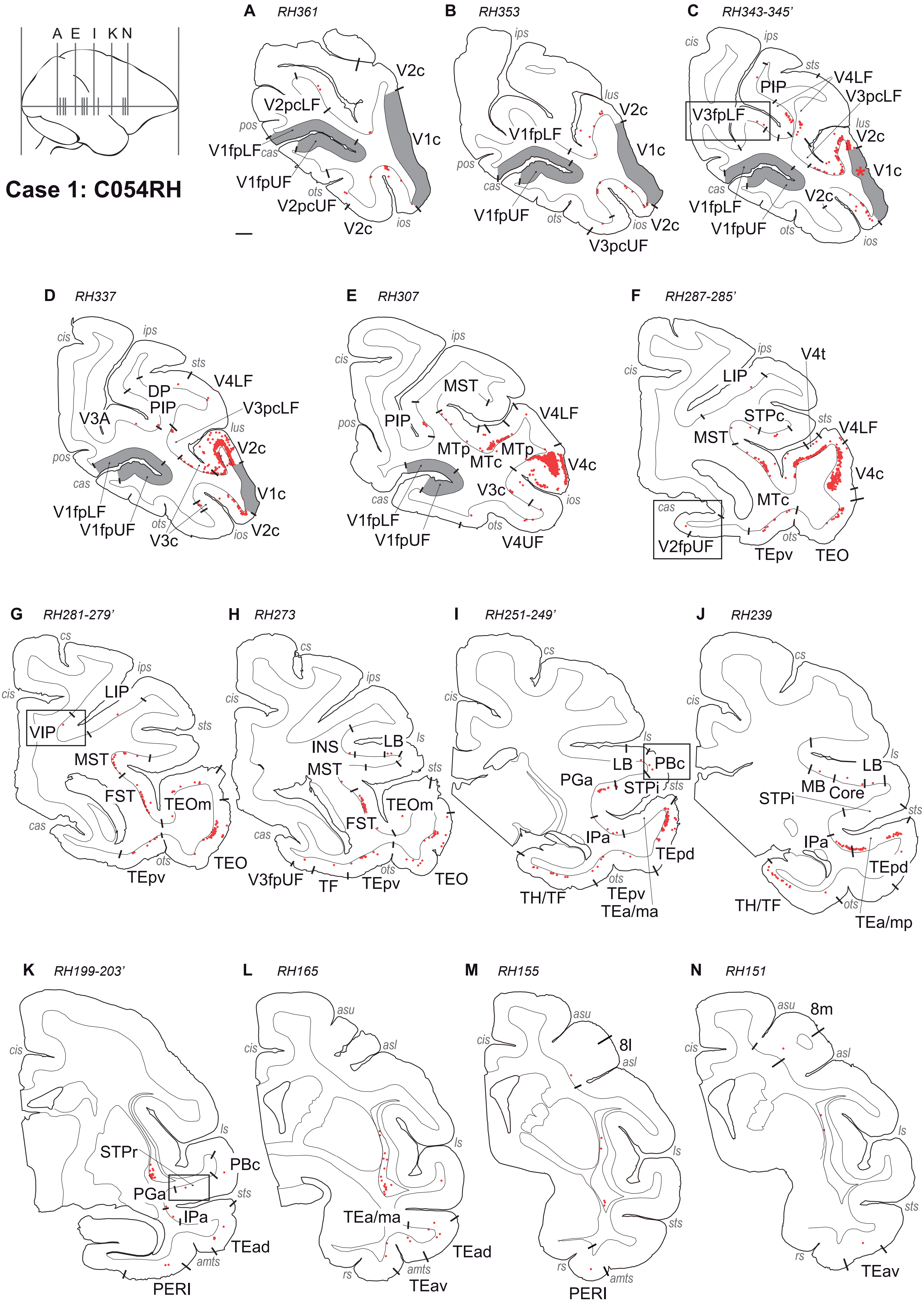
A E I K N
Case 1: C054RH
A RH361
V2c
V2pcLF
pos
V1c
V1fpLF
cas
V1fpUF
ots
V2pcUF
ios
V2c
B RH353
ips
lus
V2c
V1fpLF
V1c
pos
cas
V1fpUF
ots
ios
V2c
V3pcUF
C RH343-345'
cis
ips
sts
V4LF
PIP
V3pcLF
V3fpLF
lus
V2c
V1c
cas
V2c
V1fpLF
V1fpUF
ots
ios
D RH337
cis
ips
sts
V4LF
DP
PIP
V3pcLF
V3A
lus
V2c
pos
cas
V1c
V1fpLF
V1fpUF
ots
V3c
ios
V2c
E RH307
ips
cis
MST
pos
PIP
V4LF
MTp
MTp
MTc
V4c
V3c
ios
cas
V1fpLF
ots
V1fpUF
V4UF
F RH287-285'
ips
V4t
cis
LIP
STPc
sts
V4LF
MST
V4c
MTc
cas
ots
V2fpUF
TEpv
TEO
G RH281-279'
cs
ips
cis
LIP
VIP
sts
MST
FST
TEOm
cas
ots
TEO
TEpv
H RH273
cs
ips
cis
INS
LB
ls
MST
sts
TEOm
FST
cas
ots
V3fpUF
TF
TEpv
TEO
I RH251-249'
cs
cis
ls
LB
PBc
PGa
STPi
sts
IPa
TEpd
ots
TH/TF
TEpv
TEa/ma
J RH239
cs
cis
ls
LB
MB
Core
STPi
sts
IPa
TEpd
ots
TH/TF
TEa/mp
K RH199-203'
cis
ls
STPr
PBc
PGa
sts
IPa
TEad
amts
PERI
L RH165
asu
asl
cis
ls
TEa/ma
TEad
rs
amts
TEav
M RH155
asu
8l
asl
cis
ls
sts
rs
amts
PERI
N RH151
asu
8m
asl
cis
ls
sts
rs
TEav

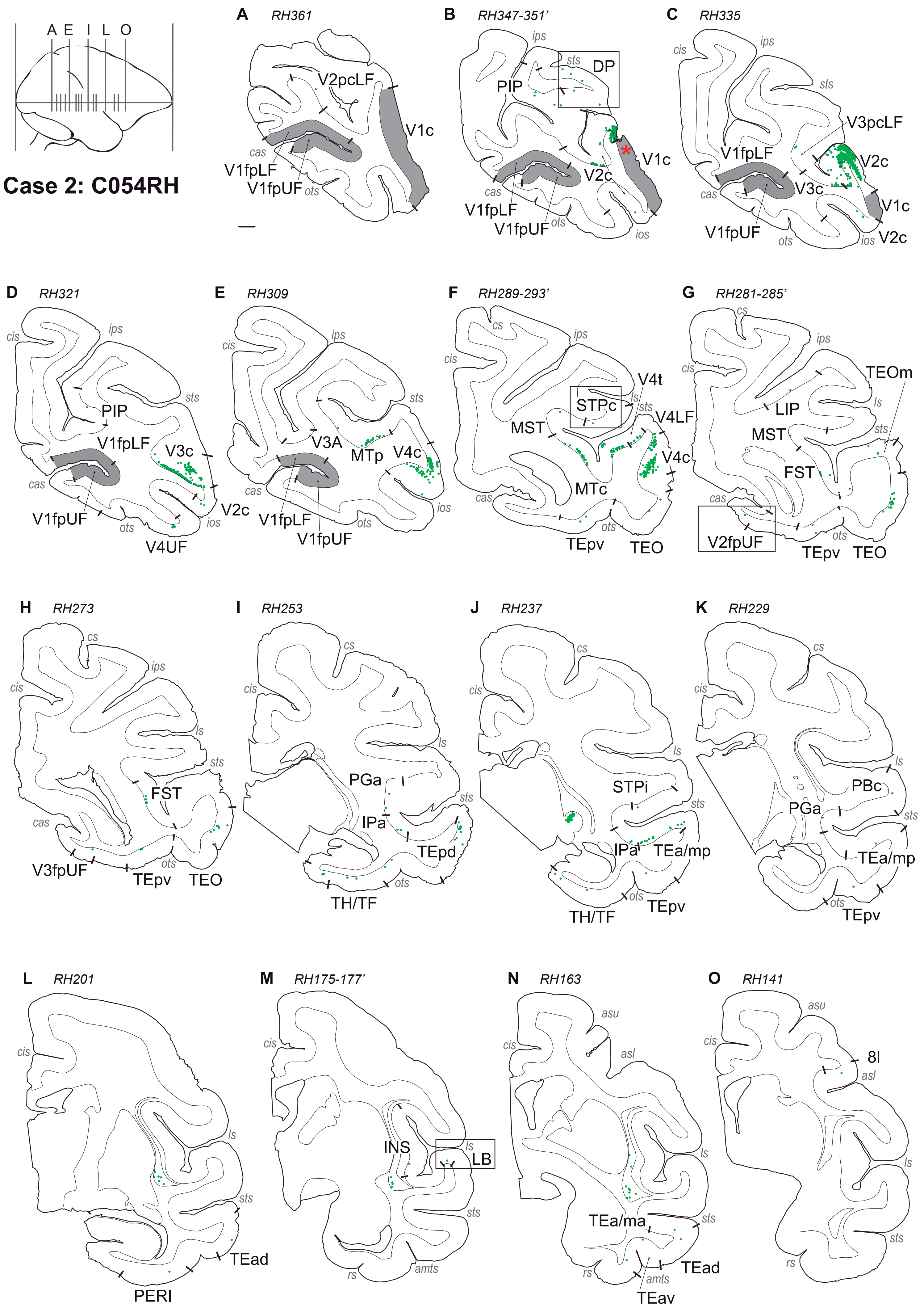

Case 2: C054RH
A E I L O
A RH361
V2pcLF
V1c
cas
V1fpLF
V1fpUF
ots
B RH347-351'
ips
sts
DP
PIP
V1c
V2c
cas
V1fpLF
V1fpUF
ots
ios
C RH335
cis
ips
sts
V3pcLF
V1fpLF
V2c
cas
V3c
V1c
V1fpUF
ots
ios
V2c
D RH321
cis
ips
sts
PIP
V1fpLF
V3c
cas
V2c
V1fpUF
ots
ios
V4UF
E RH309
cis
ips
sts
V3A
MTp
V4c
cas
ios
V1fpLF
ots
V1fpUF
F RH289-293'
cs
ips
cis
V4t
STPc
ls
sts
V4LF
MST
V4c
MTc
cas
ots
TEpv
TEO
G RH281-285'
cs
ips
cis
TEOm
ls
LIP
MST
sts
FST
cas
V2fpUF
ots
TEpv
TEO
H RH273
cs
ips
cis
ls
sts
FST
cas
V3fpUF
ots
TEpv
TEO
I RH253
cs
cis
ls
PGa
sts
IPa
TEpd
ots
TH/TF
J RH237
cs
cis
ls
STPi
sts
IPa
TEa/mp
ots
TEpv
TH/TF
K RH229
cs
cis
ls
PBc
PGa
sts
TEa/mp
ots
TEpv
L RH201
cis
ls
sts
TEad
PERI
M RH175-177'
cis
INS
ls
LB
sts
rs
amts
N RH163
asu
cis
asl
ls
TEa/ma
sts
rs
amts
TEad
TEav
O RH141
asu
cis
8l
asl
ls
sts
rs

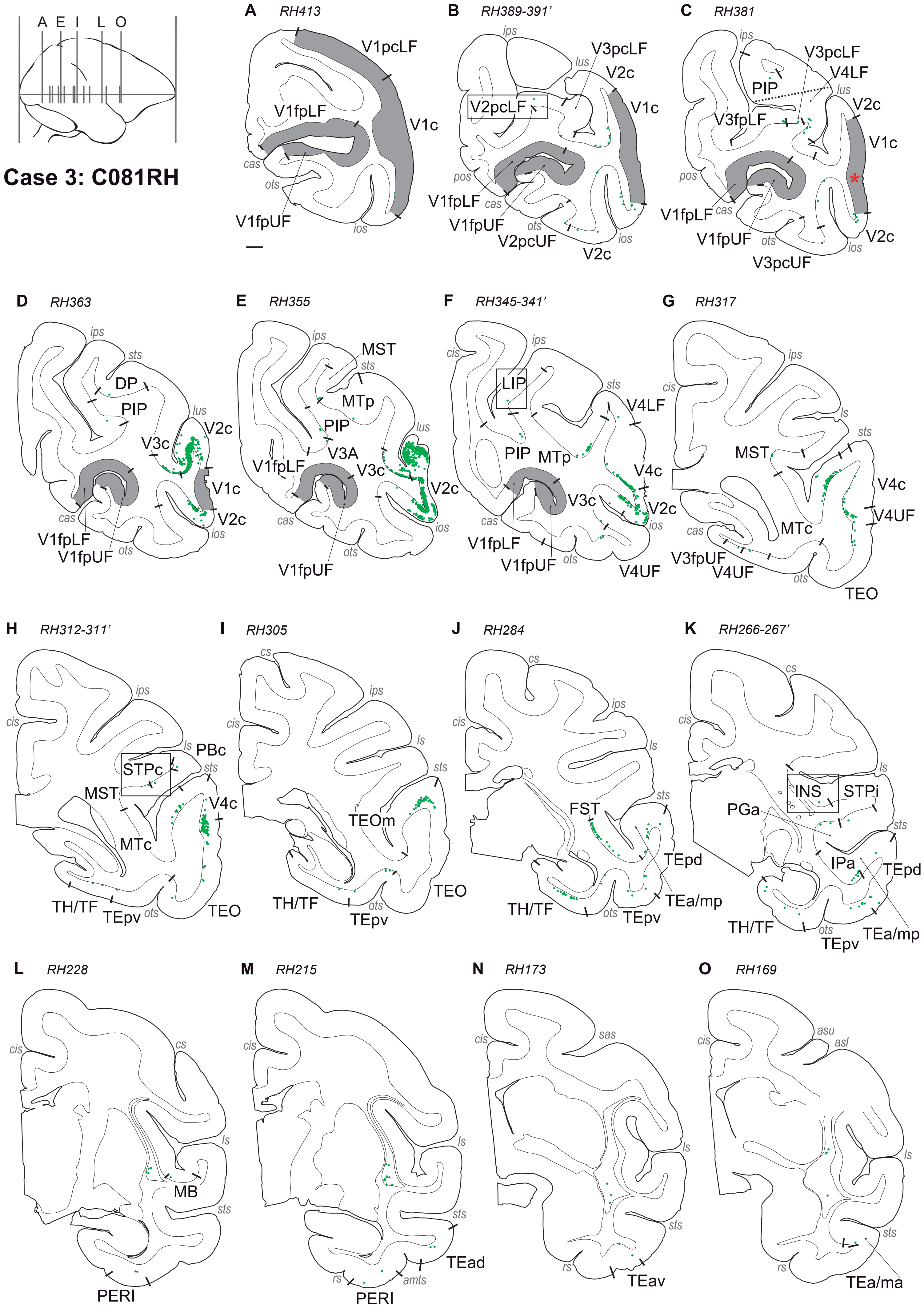

A E I L O
Case 3: C081RH
A RH413
V1pcLF
V1fpLF
V1c
cas
ots
V1fpUF
ios
B RH389-391'
ips
lus
V3pcLF
V2c
V2pcLF
V1c
pos
V1fpLF
cas
V1fpUF
ots
V2pcUF
ios
V2c
C RH381
ips
V3pcLF
V4LF
PIP
lus
V2c
V3fpLF
V1c
pos
V1fpLF
cas
V1fpUF
ots
ios
V2c
V3pcUF
D RH363
ips
sts
DP
PIP
lus
V2c
V3c
V1c
cas
V2c
ios
V1fpLF
V1fpUF
ots
E RH355
ips
MST
sts
MTp
PIP
lus
V3A
V1fpLF
V3c
V2c
cas
ios
ots
V1fpUF
F RH345-341'
cis
ips
LIP
sts
V4LF
PIP
MTp
V4c
V3c
V2c
cas
ios
V1fpLF
V1fpUF
ots
V4UF
G RH317
ips
cis
ls
sts
MST
V4c
V4UF
MTc
cas
V3fpUF
V4UF
ots
TEO
H RH312-311'
ips
cis
ls
PBc
STPc
sts
MST
V4c
MTc
TH/TF
TEpv
ots
TEO
I RH305
cs
ips
cis
ls
sts
TEOm
TEO
TH/TF
ots
TEpv
J RH284
cs
ips
cis
ls
sts
FST
TEpd
TH/TF
ots
TEpv
TEa/mp
K RH266-267'
cs
cis
ls
INS
STPi
PGa
sts
IPa
TEpd
TH/TF
ots
TEpv
TEa/mp
L RH228
cis
cs
ls
MB
sts
PERI
M RH215
cis
ls
sts
TEad
rs
amts
PERI
N RH173
cis
sas
ls
sts
rs
TEav
O RH169
asu
cis
asl
ls
sts
rs
TEa/ma

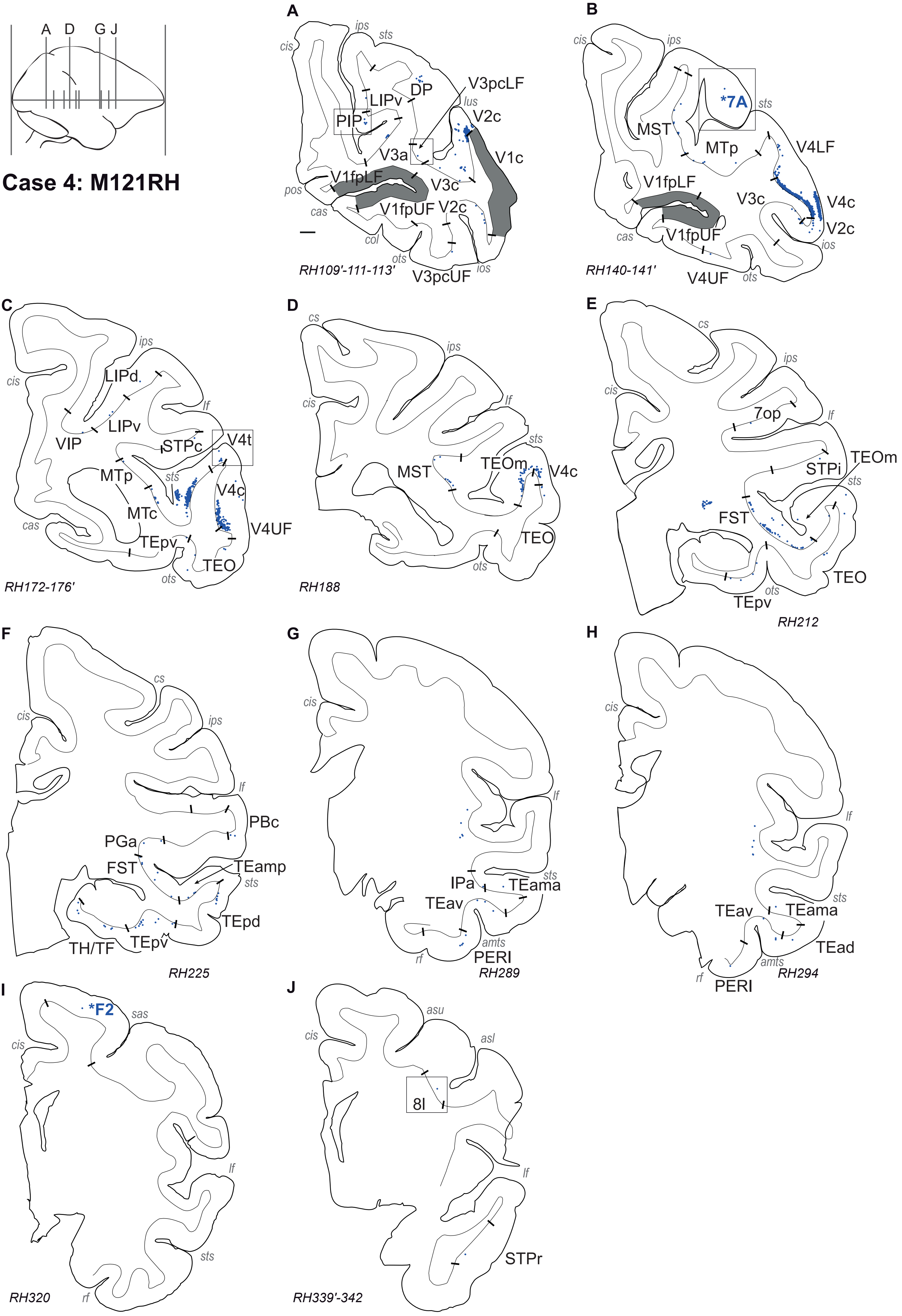
A D G J
Case 4: M121RH
A
ips sts cis
DP V3pcLF
LIPv lus
PIP V2c
V3a V1c
pos V1fpLF V3c
cas V1fpUF V2c
col ots ios
RH109'-111-113' V3pcUF
B
cis ips
*7A sts
MST MTp V4LF
V1fpLF V3c V4c
cas V1fpUF V2c ios
RH140-141' V4UF ots
C
ips cis LIPd
LIPv lf
VIP STPc V4t
MTp sts V4c
MTc V4UF
cas TEpv TEO
RH172-176' ots
D
cs ips cis lf
sts MST TEOm V4c
TEO ots
RH188
E
cs ips cis
7op lf
TEOm STPi sts
FST TEO
TEpv ots
RH212
F
cs cis ips
lf PBc
PGa FST TEamp sts
TEpd TH/TF TEpv
RH225
G
cis lf
IPa sts TEama TEav
amts rf PERI
RH289
H
cis lf
sts TEav TEama TEad
amts rf PERI
RH294
I
*F2 sas cis
lf sts
RH320 rf
J
asu cis asl
8l lf
STPr
RH339'-342

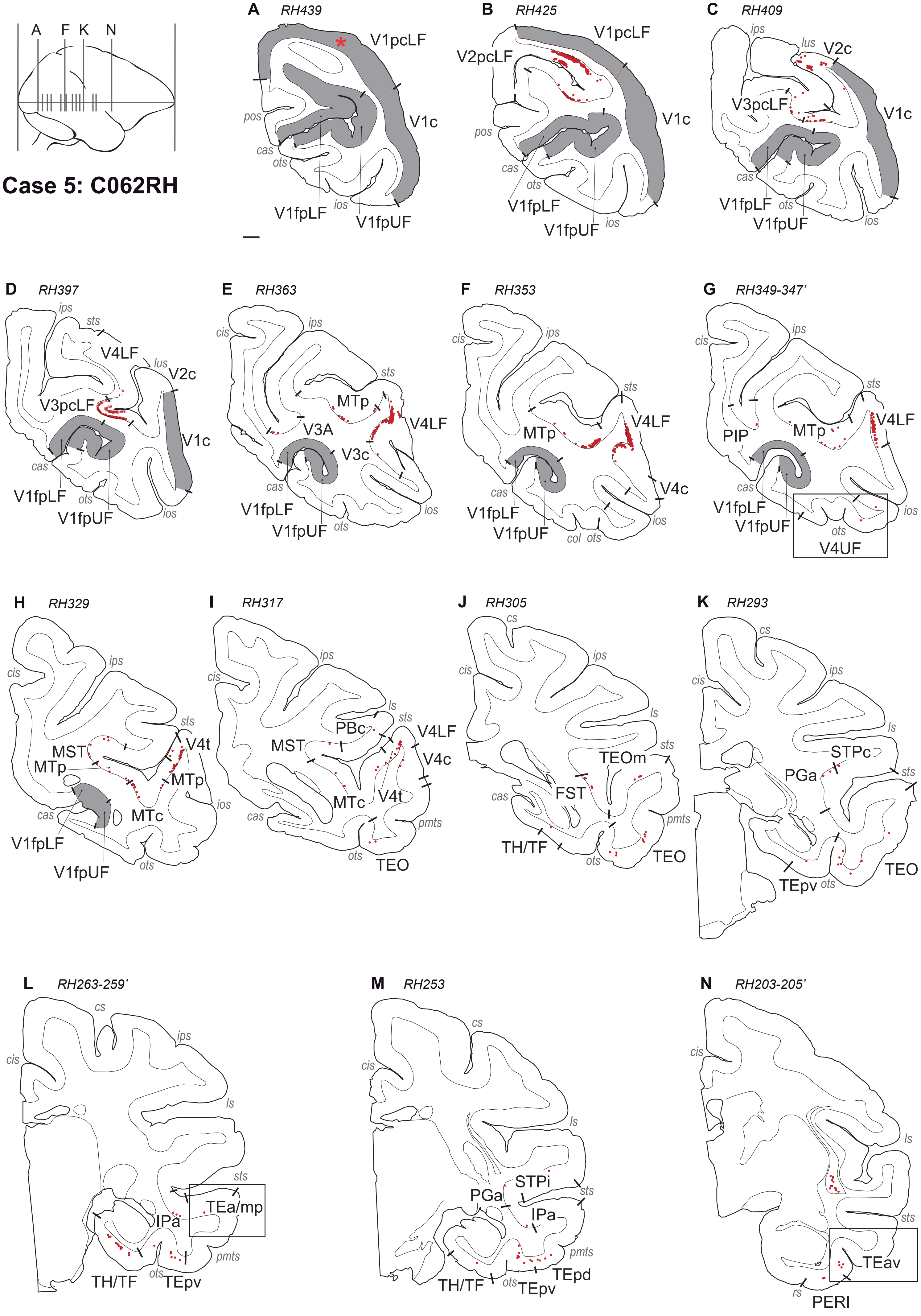

Case 5: C062RH
A F K N
A RH439
V1pcLF
V1c
pos
cas
ots
ios
V1fpLF
V1fpUF
B RH425
V2pcLF
V1pcLF
V1c
pos
cas
ots
V1fpLF
V1fpUF
ios
C RH409
ips
lus
V2c
V3pcLF
V1c
cas
V1fpLF
ots
V1fpUF
ios
D RH397
ips
sts
V4LF
lus
V2c
V3pcLF
V1c
cas
V1fpLF
ots
V1fpUF
ios
E RH363
cis
ips
sts
MTp
V4LF
V3A
V3c
cas
V1fpLF
V1fpUF
ots
ios
F RH353
cis
ips
sts
V4LF
MTp
V4c
cas
V1fpLF
V1fpUF
col
ots
ios
G RH349-347'
cis
ips
sts
V4LF
PIP
MTp
cas
V1fpLF
V1fpUF
ots
V4UF
ios
H RH329
cis
ips
sts
V4t
MST
MTp
MTp
MTc
cas
V1fpLF
V1fpUF
ots
ios
I RH317
cis
ips
ls
sts
PBc
V4LF
MST
V4c
MTc
V4t
cas
pmts
ots
TEO
J RH305
cs
cis
ips
ls
sts
TEOm
FST
cas
pmts
TH/TF
ots
TEO
K RH293
cs
cis
ips
ls
STPc
PGa
sts
TEO
TEpv
ots
L RH263-259'
cs
ips
cis
ls
sts
TEa/mp
IPa
pmts
TH/TF
ots
TEpv
M RH253
cs
cis
ls
STPi
PGa
sts
IPa
pmts
TEpd
TH/TF
ots
TEpv
N RH203-205'
cis
ls
sts
TEav
rs
PERI

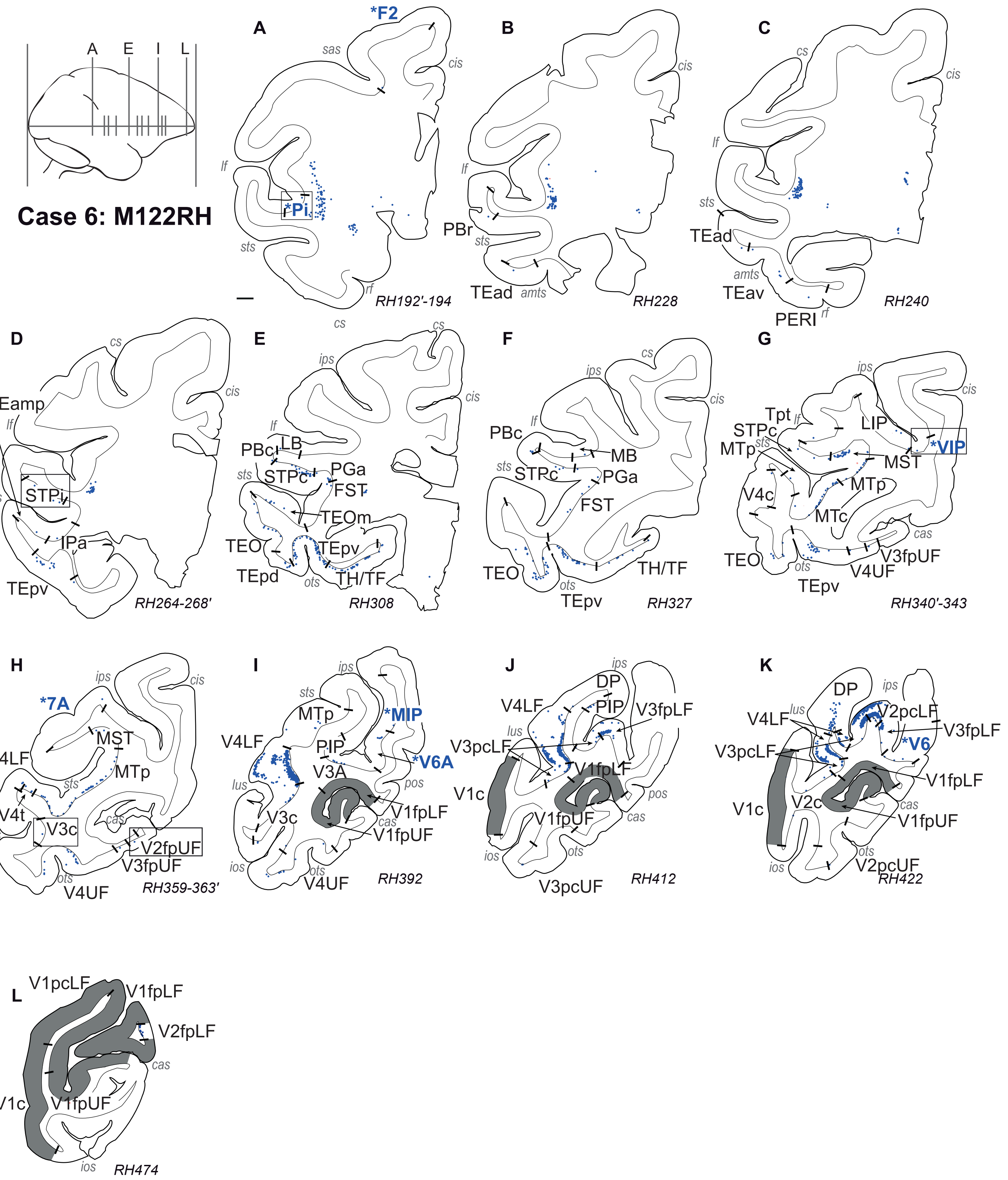

Case 6: M122RH
A
E
I
L
A
*F2
sas
cis
lf
*Pi
sts
rf
RH192'-194
B
cis
lf
PBr
sts
TEad
amts
RH228
C
cs
cis
lf
sts
TEad
amts
TEav
PERI
rf
RH240
D
cs
cis
TEamp
lf
sts
STPi
IPa
TEpv
RH264-268'
E
cs
ips
cis
lf
PBcl
LB
sts
STPc
PGa
FST
TEOm
TEO
TEpv
TEpd
ots
TH/TF
RH308
F
cs
ips
cis
PBc
lf
MB
sts
STPc
PGa
FST
TEO
ots
TEpv
TH/TF
RH327
G
ips
cis
Tpt
lf
STPc
LIP
MTp
sts
*VIP
MST
V4c
MTp
MTc
cas
TEO
ots
V3fpUF
V4UF
TEpv
RH340'-343
H
ips
cis
*7A
MST
V4LF
MTp
sts
V4t
V3c
cas
ios
V2fpUF
V3fpUF
ots
V4UF
RH359-363'
I
ips
sts
MTp
*MIP
V4LF
PIP
*V6A
V3A
lus
pos
V3c
V1fpLF
cas
V1fpUF
ios
ots
V4UF
RH392
J
ips
DP
V4LF
PIP
V3fpLF
lus
V3pcLF
V1fpLF
V1c
pos
V1fpUF
cas
ios
ots
V3pcUF
RH412
K
DP
ips
V4LF
lus
V2pcLF
V3fpLF
V3pcLF
*V6
V1fpLF
V1c
V2c
cas
V1fpUF
ios
ots
V2pcUF
RH422
L
V1pcLF
V1fpLF
V2fpLF
cas
V1c
V1fpUF
ios
RH474

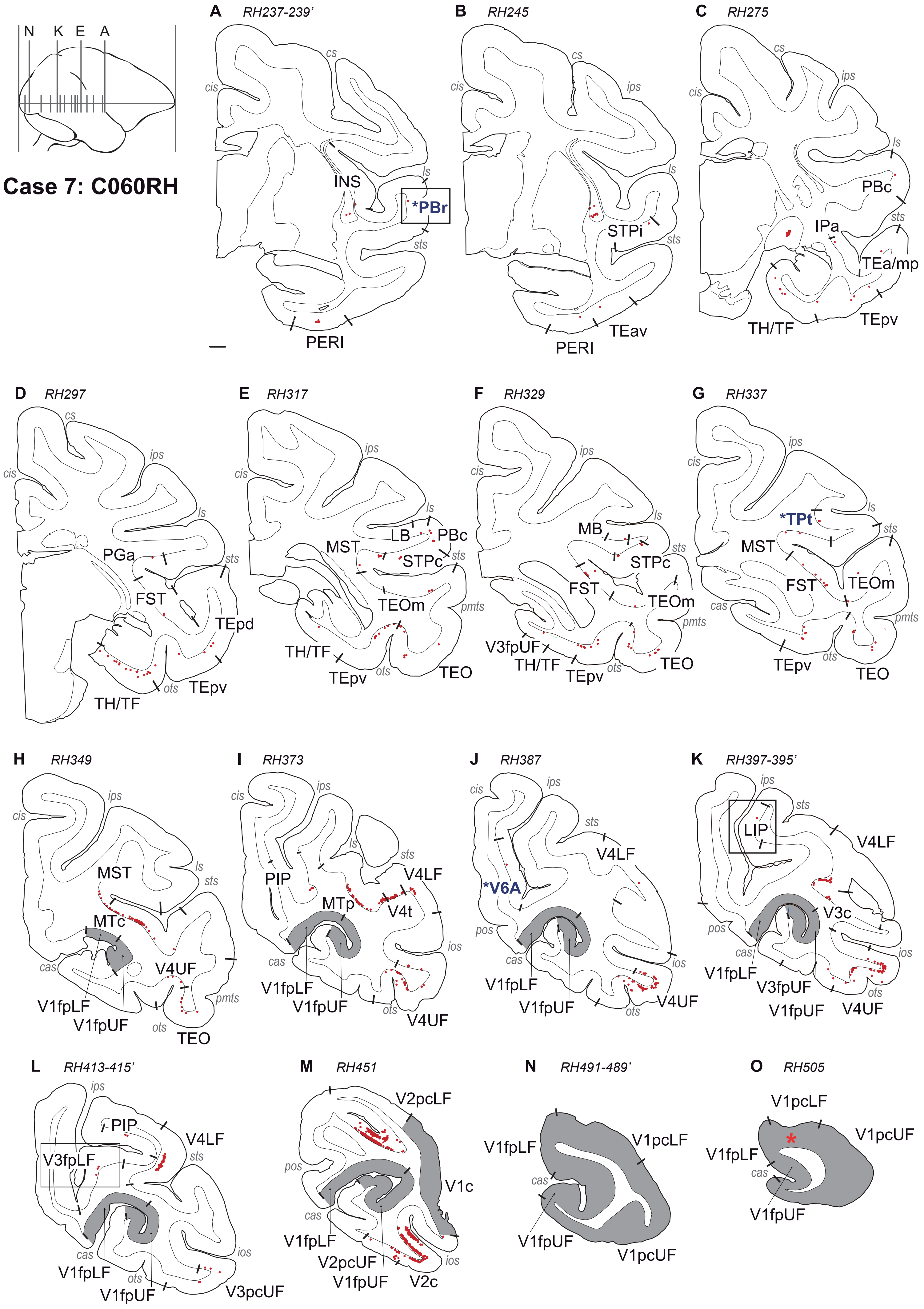
N K E A
Case 7: C060RH
A RH237-239'
cs
cis
INS
ls
*PBr
sts
PERI
B RH245
cs
cis
ips
ls
STPi
sts
PERI
TEav
C RH275
cs
ips
cis
ls
PBc
sts
IPa
TEa/mp
TH/TF
TEpv
D RH297
cs
cis
ips
ls
PGa
sts
FST
TEpd
TH/TF
ots
TEpv
E RH317
ips
cis
ls
LB
PBc
MST
STPc
sts
TEOm
pmts
TH/TF
TEpv
ots
TEO
F RH329
ips
cis
ls
MB
STPc
sts
FST
TEOm
pmts
V3fpUF
TH/TF
TEpv
ots
TEO
G RH337
ips
cis
*TPt
ls
sts
MST
FST
TEOm
cas
pmts
TEpv
ots
TEO
H RH349
ips
cis
MST
ls
sts
MTc
cas
V4UF
V1fpLF
V1fpUF
pmts
ots
TEO
I RH373
ips
cis
ls
sts
PIP
V4LF
MTp
V4t
cas
ios
V1fpLF
V1fpUF
ots
V4UF
J RH387
cis
ips
sts
V4LF
*V6A
pos
cas
ios
V1fpLF
V1fpUF
ots
V4UF
K RH397-395'
ips
sts
LIP
V4LF
V3c
pos
cas
ios
V1fpLF
V3fpUF
V1fpUF
ots
V4UF
L RH413-415'
ips
PIP
V4LF
sts
V3fpLF
cas
ios
V1fpLF
ots
V1fpUF
V3pcUF
M RH451
V2pcLF
pos
V1c
cas
V1fpLF
V2pcUF
V1fpUF
ios
V2c
N RH491-489'
V1fpLF
V1pcLF
cas
V1fpUF
V1pcUF
O RH505
V1pcLF
V1pcUF
V1fpLF
cas
V1fpUF

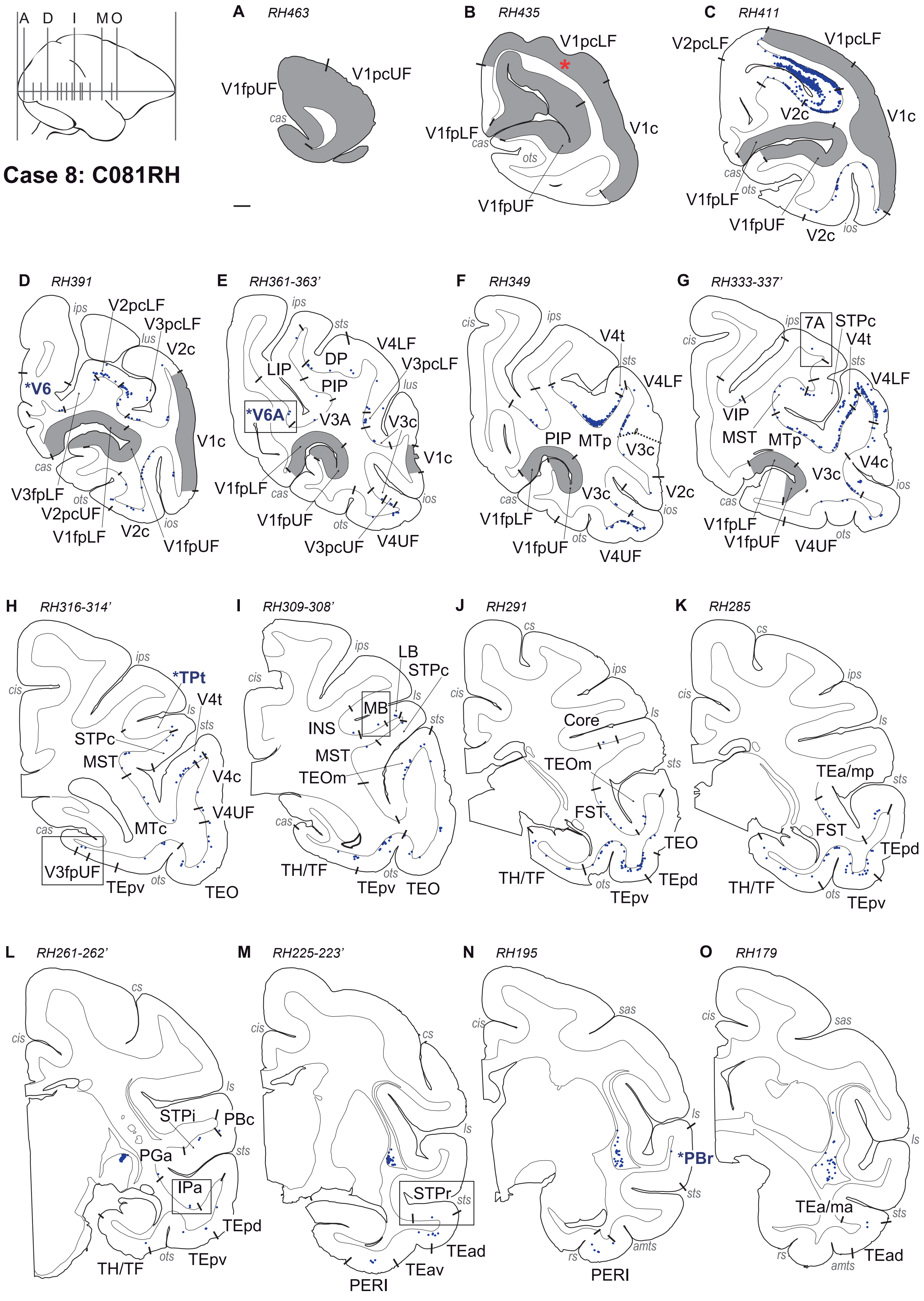

Case 8: C081RH
A D I M O
A RH463
V1pcUF
V1fpUF
cas
B RH435
V1pcLF
V1fpLF
V1c
cas
ots
V1fpUF
C RH411
V2pcLF
V1pcLF
V2c
V1c
cas
ots
V1fpLF
V1fpUF
V2c
ios
D RH391
ips
V2pcLF
V3pcLF
lus
V2c
*V6
V1c
cas
V3fpLF
ots
V2pcUF
V1fpLF
V2c
ios
V1fpUF
E RH361-363'
ips
sts
V4LF
DP
V3pcLF
LIP
PIP
lus
*V6A
V3A
V3c
V1c
V1fpLF
cas
V1fpUF
ots
V3pcUF
V4UF
ios
F RH349
cis
ips
V4t
sts
V4LF
PIP
MTp
V3c
V2c
cas
V3c
ios
V1fpLF
V1fpUF
ots
V4UF
G RH333-337'
cis
ips
7A
STPc
V4t
sts
V4LF
VIP
MST
MTp
V4c
V3c
ios
cas
V1fpLF
V1fpUF
V4UF
ots
H RH316-314'
ips
*TPt
cis
V4t
ls
sts
STPc
MST
V4c
V4UF
cas
MTc
V3fpUF
TEpv
ots
TEO
I RH309-308'
ips
LB
cis
STPc
ls
MB
INS
sts
MST
TEOm
cas
TH/TF
ots
TEpv
TEO
J RH291
cs
ips
cis
ls
Core
TEOm
sts
FST
TEO
TH/TF
ots
TEpd
TEpv
K RH285
cs
ips
cis
ls
TEa/mp
sts
FST
TEpd
TH/TF
ots
TEpv
L RH261-262'
cs
cis
ls
STPi
PBc
PGa
sts
IPa
TEpd
TH/TF
ots
TEpv
M RH225-223'
cis
cs
ls
STPr
sts
TEad
TEav
PERI
N RH195
sas
cis
ls
*PBr
sts
amts
rs
PERI
O RH179
cis
sas
ls
TEa/ma
sts
rs
amts
TEad

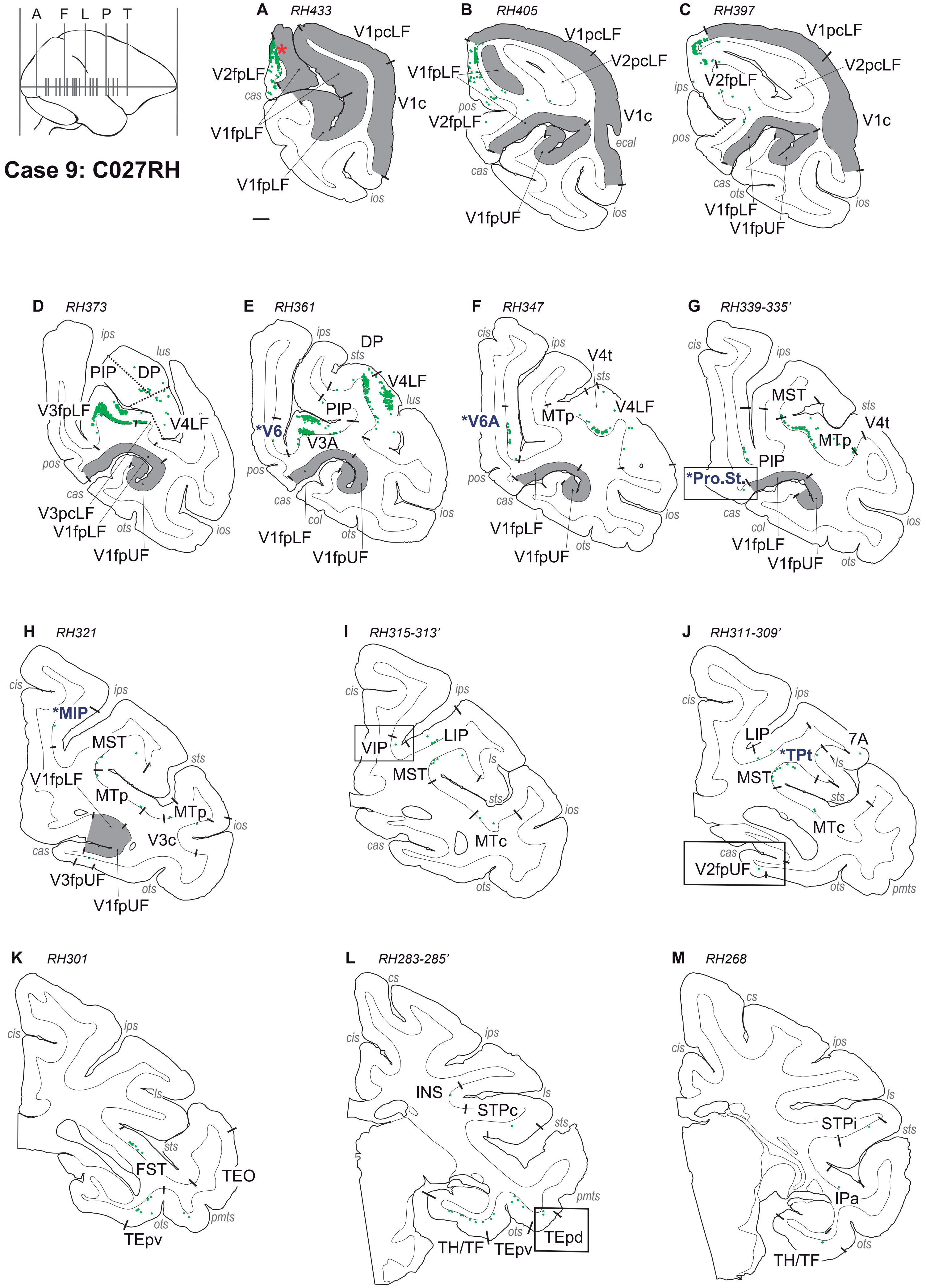

A F L P T
Case 9: C027RH
A RH433
V1pcLF
V2fpLF
cas
V1fpLF
V1c
V1fpLF
ios
B RH405
V1pcLF
V1fpLF
V2pcLF
pos
V2fpLF
V1c
ecal
cas
V1fpUF
ios
C RH397
V1pcLF
V2pcLF
ips
V2fpLF
pos
V1c
cas
ots
V1fpLF
V1fpUF
ios
D RH373
ips
lus
PIP
DP
V3fpLF
V4LF
pos
cas
V3pcLF
V1fpLF
ots
ios
V1fpUF
E RH361
ips
DP
sts
V4LF
lus
PIP
*V6
V3A
pos
cas
col
ots
ios
V1fpLF
V1fpUF
F RH347
cis
ips
V4t
sts
V4LF
*V6A
MTp
pos
cas
ios
V1fpLF
V1fpUF
ots
G RH339-335'
cis
ips
MST
sts
V4t
MTp
PIP
*Pro.St.
cas
col
V1fpLF
V1fpUF
ots
ios
H RH321
cis
ips
*MIP
MST
sts
V1fpLF
MTp
MTp
ios
V3c
cas
V3fpUF
ots
V1fpUF
I RH315-313'
cis
ips
VIP
LIP
ls
MST
sts
ios
cas
MTc
ots
J RH311-309'
cis
ips
LIP
7A
*TPt
ls
MST
sts
MTc
cas
V2fpUF
ots
pmts
K RH301
cis
ips
ls
sts
FST
TEO
pmts
ots
TEpv
L RH283-285'
cs
cis
ips
INS
ls
STPc
sts
pmts
ots
TH/TF
TEpv
TEpd
M RH268
cs
cis
ips
ls
STPi
sts
IPa
ots
TH/TF

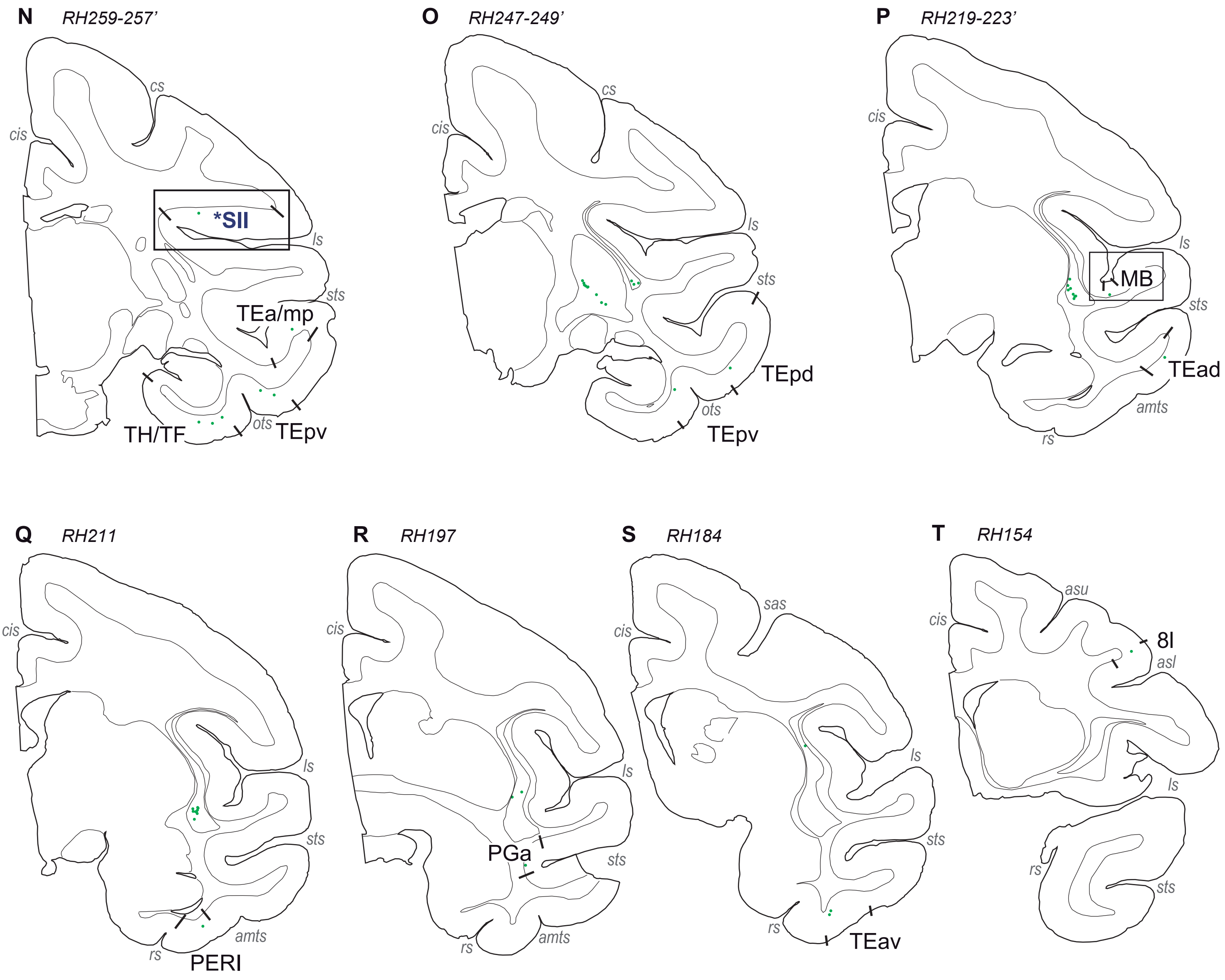
N
RH259-257'
cs
cis
*SII
ls
sts
TEa/mp
TH/TF
ots
TEpv
O
RH247-249'
cs
cis
ls
sts
TEpd
ots
TEpv
P
RH219-223'
cis
ls
MB
sts
TEad
amts
rs
Q
RH211
cis
ls
sts
amts
rs
PERI
R
RH197
cis
ls
PGa
sts
rs
amts
S
RH184
sas
cis
ls
sts
rs
TEav
T
RH154
asu
cis
8l
asl
ls
rs
sts

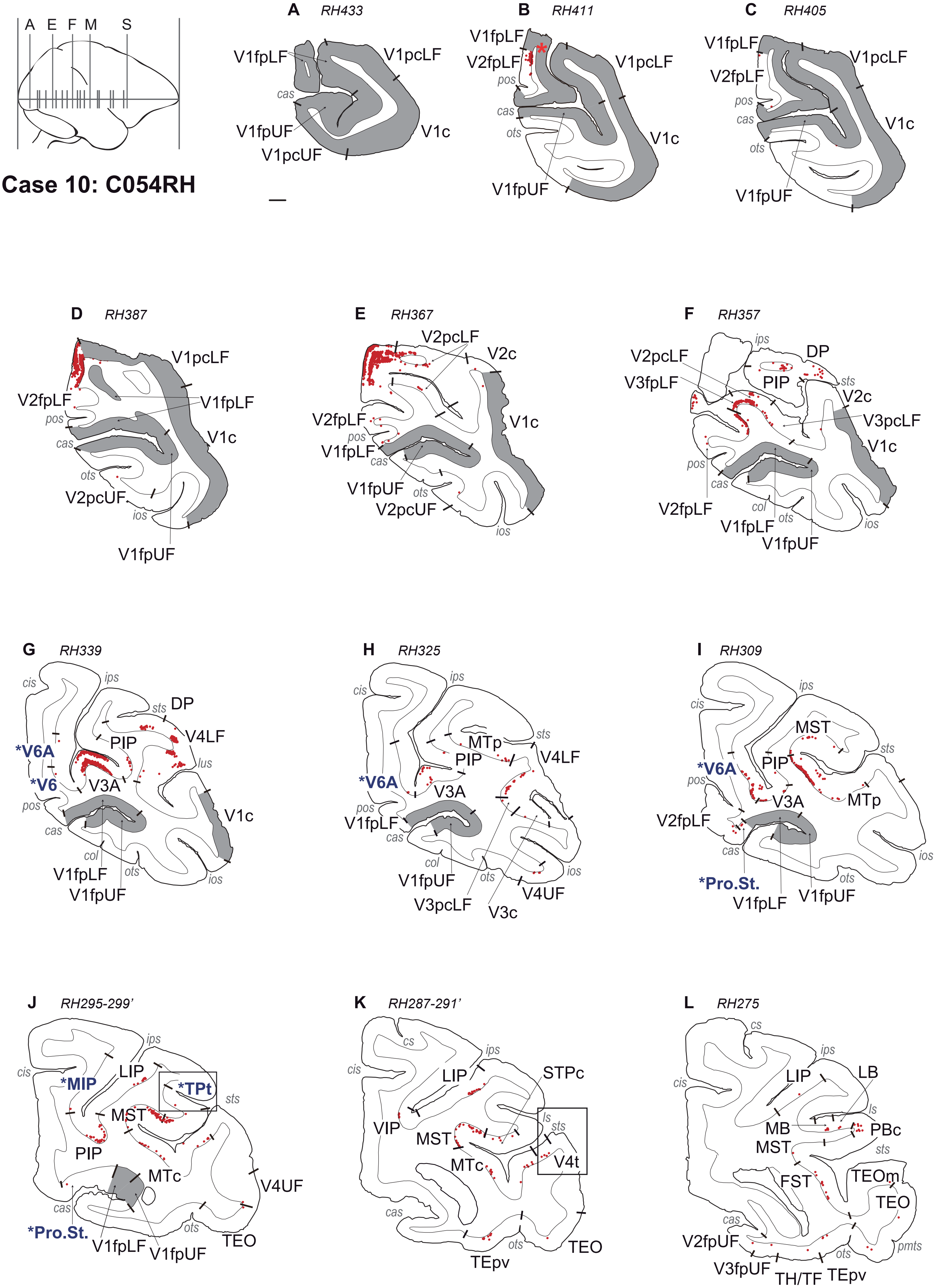

Case 10: C054RH
A E F M S
A RH433
V1fpLF V1pcLF cas V1fpUF V1c V1pcUF
B RH411
V1fpLF V2fpLF pos cas ots V1pcLF V1c V1fpUF
C RH405
V1fpLF V2fpLF pos cas ots V1pcLF V1c V1fpUF
D RH387
V1pcLF V2fpLF pos V1fpLF cas V1c ots V2pcUF ios V1fpUF
E RH367
V2pcLF V2c V2fpLF pos V1c V1fpLF cas V1fpUF ots V2pcUF ios
F RH357
ips DP V2pcLF V3fpLF PIP sts V2c V3pcLF V1c pos cas V2fpLF col ots ios V1fpLF V1fpUF
G RH339
cis ips sts DP V4LF PIP *V6A lus *V6 V3A pos V1c cas col V1fpLF ots ios V1fpUF
H RH325
ips cis sts MTp V4LF PIP *V6A V3A pos V1fpLF cas col ios V1fpUF ots V4UF V3pcLF V3c
I RH309
ips cis MST sts *V6A PIP pos V3A MTp V2fpLF ios cas ots *Pro.St. V1fpLF V1fpUF
J RH295-299'
ips cis *MIP LIP *TPt sts MST PIP MTc V4UF cas ots *Pro.St. V1fpLF V1fpUF TEO
K RH287-291'
cs ips cis LIP STPc VIP ls sts MST MTc V4t cas ots TEpv TEO
L RH275
cs ips cis LIP LB ls MB PBc MST sts FST TEOm TEO cas V2fpUF ots pmts V3fpUF TH/TF TEpv

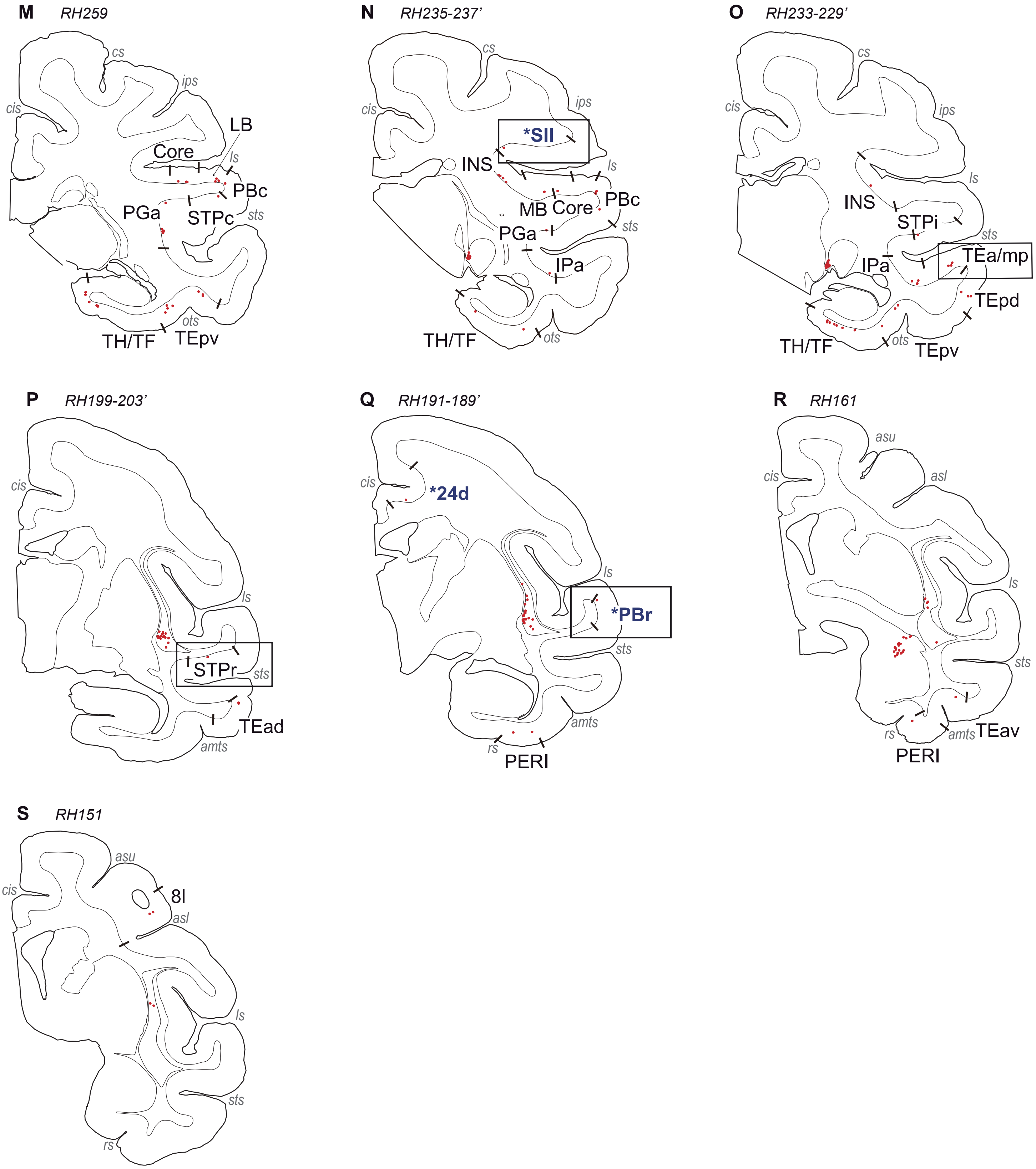

M
RH259
cs
ips
cis
LB
Core
ls
PBc
PGa
STPc
sts
ots
TH/TF
TEpv
N
RH235-237'
cs
ips
cis
*SII
INS
ls
PBc
MB
Core
PGa
sts
IPa
TH/TF
ots
O
RH233-229'
cs
ips
cis
ls
INS
STPi
sts
TEa/mp
IPa
TEpd
TH/TF
ots
TEpv
P
RH199-203'
cis
ls
STPr
sts
TEad
amts
Q
RH191-189'
cis
*24d
ls
*PBr
sts
amts
rs
PERI
R
RH161
asu
cis
asl
ls
sts
rs
amts
TEav
PERI
S
RH151
asu
cis
8l
asl
ls
sts
rs

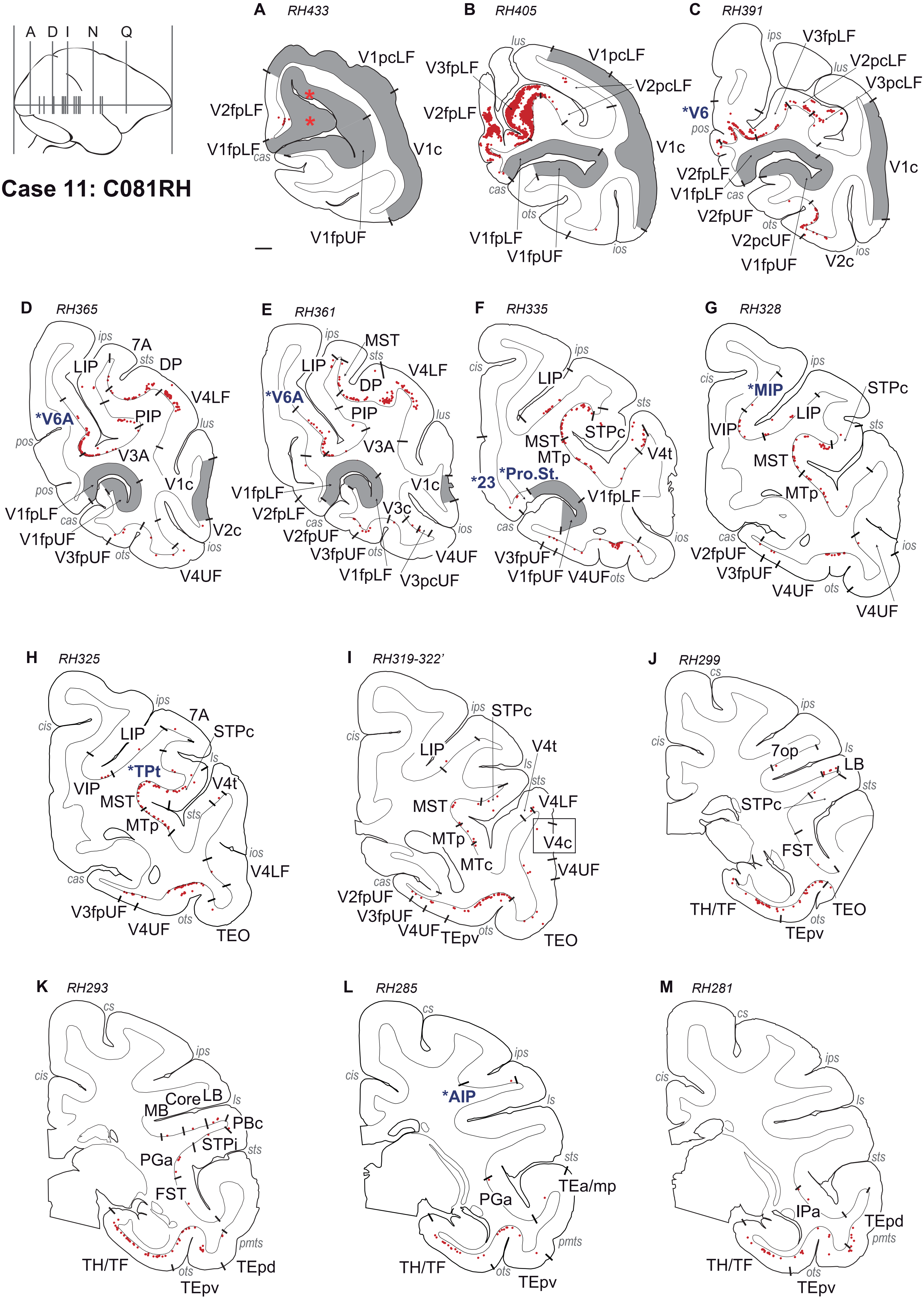

Case 11: C081RH
A D I N Q
A RH433
V1pcLF
V2fpLF
V1fpLF
cas
V1c
V1fpUF
B RH405
lus
V1pcLF
V3fpLF
V2pcLF
V2fpLF
V1c
cas
ots
V1fpLF
V1fpUF
ios
C RH391
ips
V3fpLF
lus
V2pcLF
V3pcLF
*V6
pos
V2fpLF
V1fpLF
cas
V1c
V2fpUF
ots
V2pcUF
V1fpUF
V2c
ios
D RH365
ips
7A
sts
DP
LIP
V4LF
*V6A
PIP
lus
pos
V3A
pos
V1c
V1fpLF
cas
V1fpUF
V2c
V3fpUF
ots
ios
V4UF
E RH361
ips
MST
sts
LIP
V4LF
DP
*V6A
PIP
lus
V3A
V1fpLF
V1c
V2fpLF
V3c
cas
V2fpUF
ios
V3fpUF
ots
V4UF
V1fpLF
V3pcUF
F RH335
ips
cis
LIP
sts
MST
STPc
MTp
V4t
*23
*Pro.St.
V1fpLF
cas
ios
V3fpUF
V1fpUF
V4UF
ots
G RH328
ips
cis
*MIP
STPc
LIP
VIP
sts
MST
MTp
cas
ios
V2fpUF
V3fpUF
V4UF
ots
V4UF
H RH325
ips
cis
7A
LIP
STPc
*TPt
ls
VIP
V4t
MST
sts
MTp
ios
V4LF
cas
V3fpUF
V4UF
ots
TEO
I RH319-322'
ips
STPc
cis
V4t
LIP
ls
sts
MST
V4LF
MTp
V4c
MTc
V4UF
cas
V2fpUF
V3fpUF
V4UF
TEpv
ots
TEO
J RH299
cs
ips
cis
ls
7op
LB
sts
STPc
FST
TH/TF
TEO
ots
TEpv
K RH293
cs
ips
cis
LB
Core
ls
MB
PBc
STPi
sts
PGa
FST
pmts
TH/TF
ots
TEpd
TEpv
L RH285
cs
ips
cis
*AIP
ls
sts
TEa/mp
PGa
pmts
TH/TF
ots
TEpv
M RH281
cs
ips
cis
ls
sts
IPa
TEpd
pmts
TH/TF
ots
TEpv

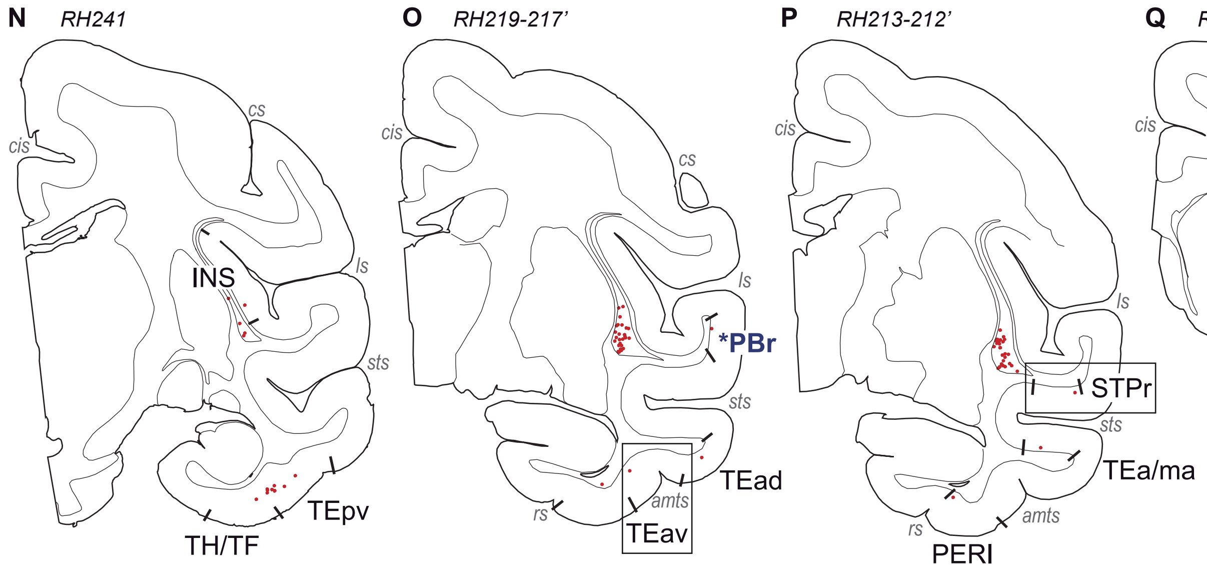
N
RH241
cs
cis
INS
ls
sts
TEpv
TH/TF
O
RH219-217'
cis
cs
ls
*PBr
sts
TEad
amts
rs
TEav
P
RH213-212'
cis
ls
STPr
sts
TEa/ma
amts
rs
PERI
Q
RH141
asu
cis
asl
rs
sts
PERI

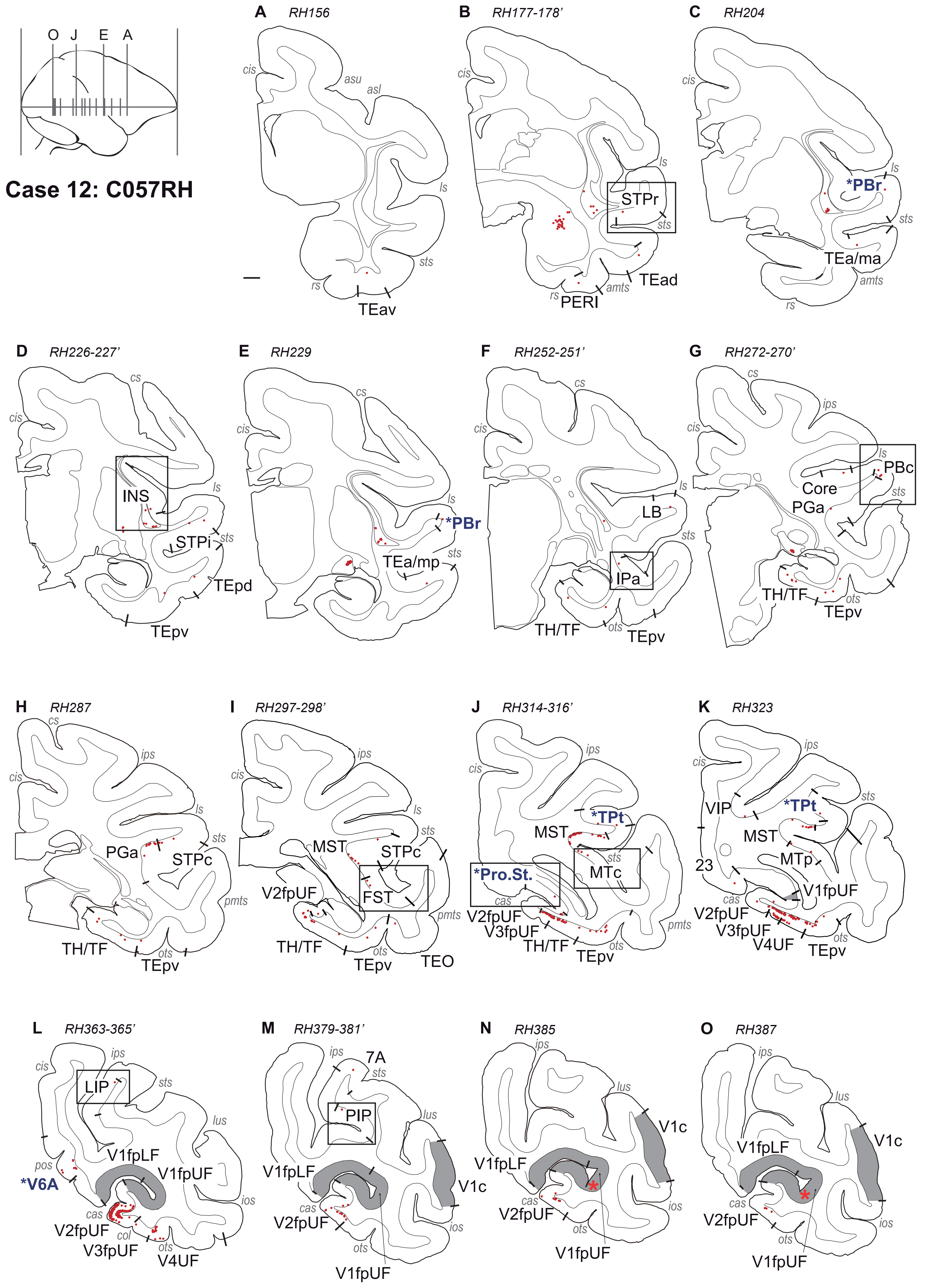

O J E A
Case 12: C057RH
A RH156
B RH177-178'
C RH204
D RH226-227'
E RH229
F RH252-251'
G RH272-270'
H RH287
I RH297-298'
J RH314-316'
K RH323
L RH363-365'
M RH379-381'
N RH385
O RH387
cis
asu
asl
ls
sts
rs
amts
cs
ips
pmts
ots
cas
pos
col
lus
ios
TEav
STPr
TEad
PERI
*PBr
TEa/ma
INS
STPi
TEpd
TEpv
TEa/mp
LB
IPa
TH/TF
PBc
Core
PGa
STPc
MST
FST
V2fpUF
TEO
*TPt
*Pro.St.
MTc
V3fpUF
VIP
23
MTp
V1fpUF
V4UF
LIP
*V6A
V1fpLF
7A
PIP
V1c

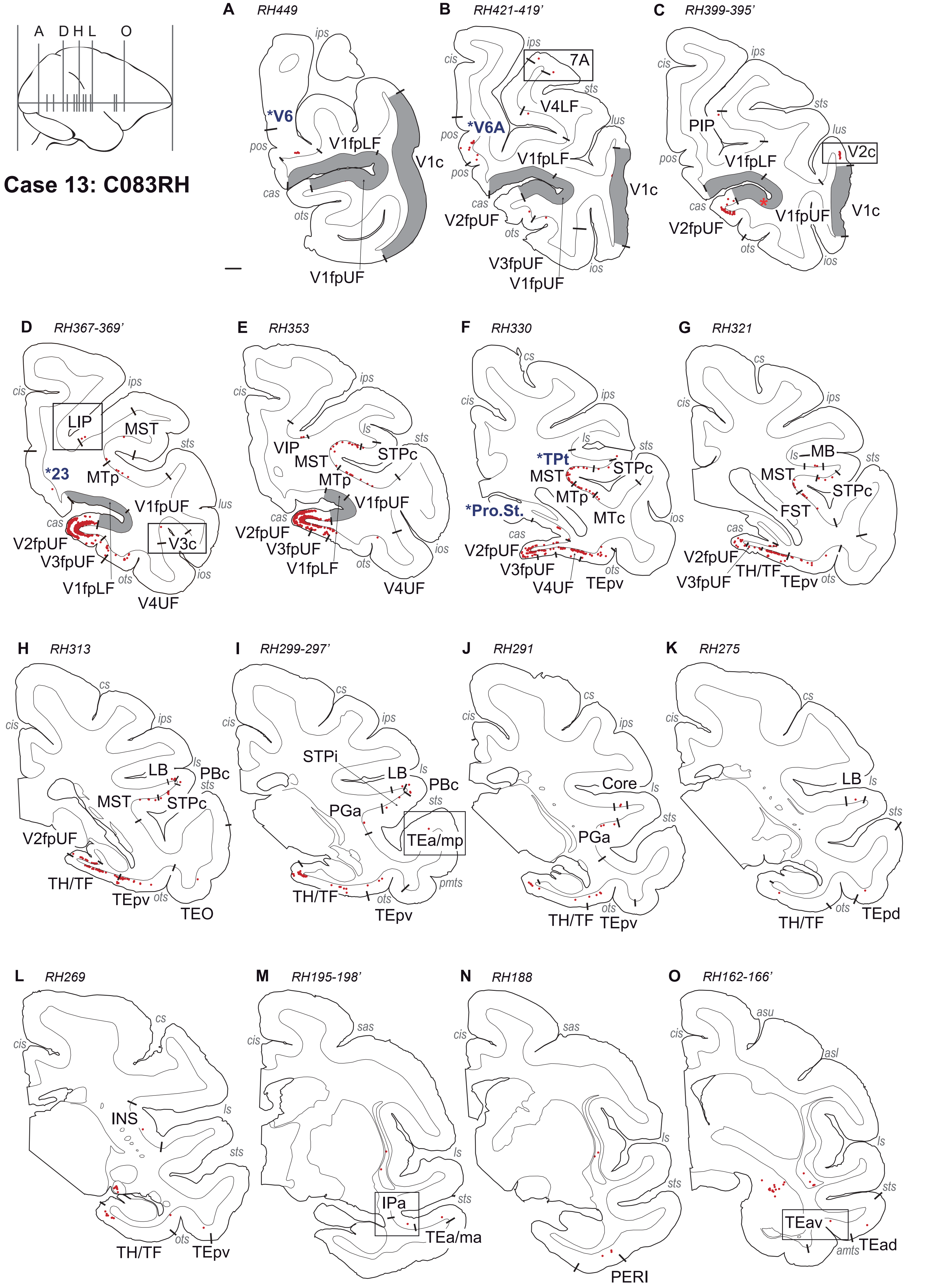

A D H L O
Case 13: C083RH
A RH449
ips
*V6
pos
cas
ots
V1fpLF
V1c
V1fpUF
B RH421-419'
ips
cis
7A
sts
V4LF
lus
pos
*V6A
pos
V1fpLF
cas
V1c
V2fpUF
ots
V3fpUF
V1fpUF
ios
C RH399-395'
ips
cis
sts
PIP
lus
pos
V1fpLF
V2c
cas
V1fpUF
V1c
V2fpUF
ots
ios
D RH367-369'
cis
ips
LIP
MST
sts
*23
MTp
V1fpUF
lus
cas
V2fpUF
V3c
V3fpUF
V1fpLF
ots
ios
V4UF
E RH353
cis
ips
ls
sts
VIP
MST
STPc
MTp
V1fpUF
cas
V2fpUF
V3fpUF
V1fpLF
ots
ios
V4UF
F RH330
cs
cis
ips
ls
sts
*TPt
STPc
MST
MTp
*Pro.St.
MTc
ios
cas
V2fpUF
V3fpUF
ots
V4UF
TEpv
G RH321
cs
cis
ips
ls
MB
sts
MST
STPc
FST
cas
V2fpUF
V3fpUF
TH/TF
TEpv
ots
H RH313
cs
cis
ips
ls
LB
PBc
sts
MST
STPc
V2fpUF
TH/TF
TEpv
ots
TEO
I RH299-297'
cs
cis
ips
STPi
LB
ls
PBc
sts
PGa
TEa/mp
pmts
TH/TF
ots
TEpv
J RH291
cs
cis
ips
Core
ls
sts
PGa
ots
TH/TF
TEpv
K RH275
cs
cis
LB
ls
sts
ots
TEpd
TH/TF
L RH269
cs
cis
INS
ls
sts
ots
TH/TF
TEpv
M RH195-198'
sas
cis
ls
sts
IPa
TEa/ma
N RH188
sas
cis
ls
sts
PERI
O RH162-166'
asu
cis
asl
ls
sts
TEav
amts
TEad

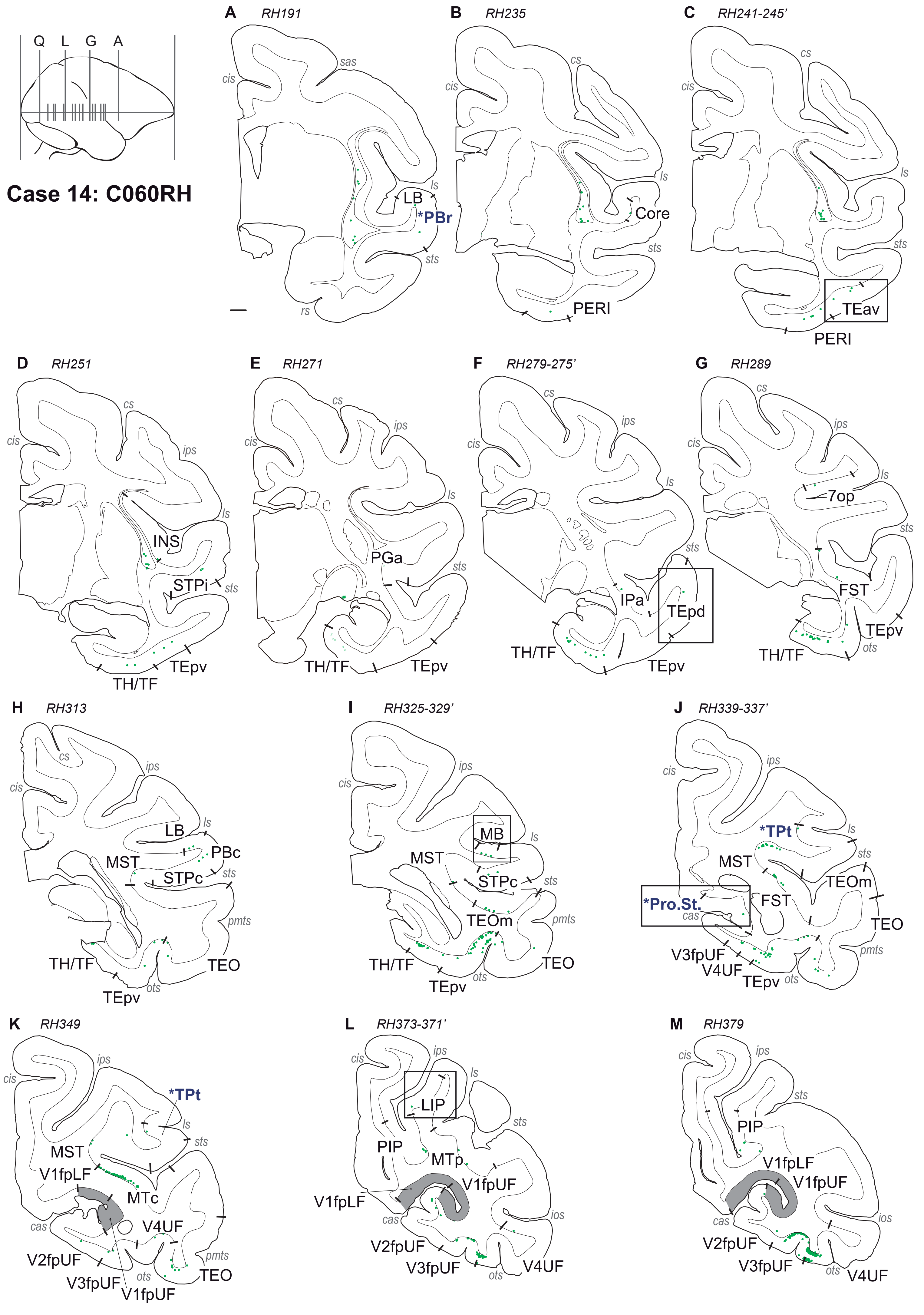
Case 14: C060RH
Q L G A
A RH191
sas
cis
ls
LB
*PBr
sts
rs
B RH235
cs
cis
ls
Core
sts
PERI
C RH241-245'
cs
cis
ls
sts
TEav
PERI
D RH251
cs
cis
ips
ls
INS
STPi
sts
TEpv
TH/TF
E RH271
cs
ips
cis
ls
PGa
sts
TH/TF
TEpv
F RH279-275'
cs
ips
cis
ls
sts
IPa
TEpd
TH/TF
TEpv
G RH289
cs
ips
cis
ls
7op
sts
FST
TEpv
ots
TH/TF
H RH313
cs
ips
cis
ls
LB
PBc
MST
STPc
sts
pmts
TH/TF
TEO
TEpv
ots
I RH325-329'
ips
cis
MB
ls
MST
STPc
sts
TEOm
pmts
TH/TF
TEO
TEpv
ots
J RH339-337'
ips
cis
*TPt
ls
sts
MST
TEOm
*Pro.St.
cas
FST
TEO
pmts
V3fpUF
V4UF
TEpv
ots
K RH349
ips
cis
*TPt
ls
sts
MST
V1fpLF
MTc
cas
V4UF
pmts
V2fpUF
V3fpUF
ots
TEO
V1fpUF
L RH373-371'
cis
ips
ls
LIP
sts
PIP
MTp
V1fpUF
V1fpLF
cas
ios
V2fpUF
V3fpUF
ots
V4UF
M RH379
cis
ips
sts
PIP
V1fpLF
V1fpUF
cas
ios
V2fpUF
V3fpUF
ots
V4UF

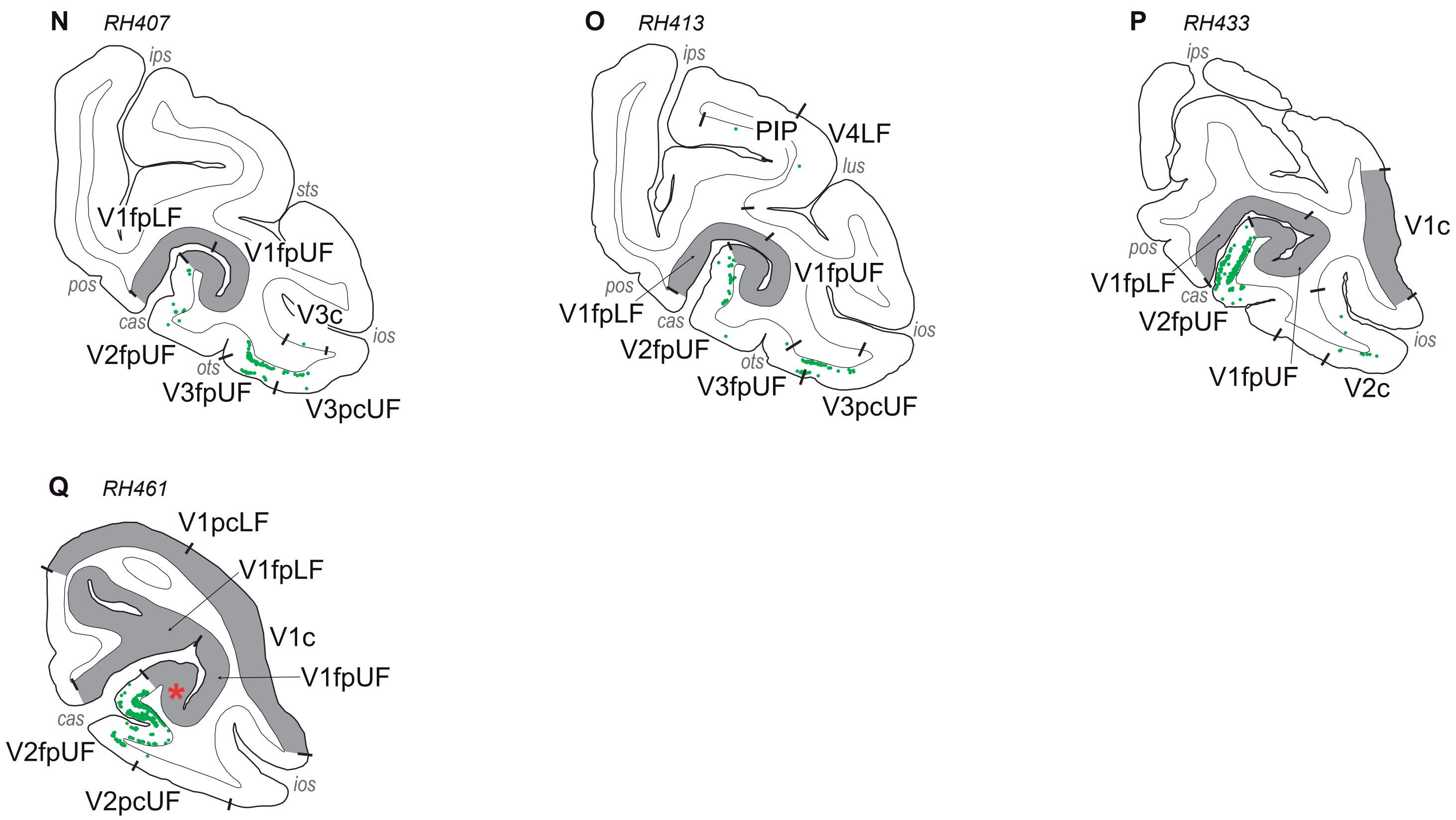

N
RH407
ips
sts
V1fpLF
V1fpUF
pos
V3c
cas
ios
V2fpUF
ots
V3fpUF
V3pcUF
O
RH413
ips
PIP
V4LF
lus
V1fpUF
pos
V1fpLF
cas
ios
V2fpUF
ots
V3fpUF
V3pcUF
P
RH433
ips
V1c
pos
V1fpLF
cas
V2fpUF
ios
V1fpUF
V2c
Q
RH461
V1pcLF
V1fpLF
V1c
V1fpUF
cas
V2fpUF
ios
V2pcUF

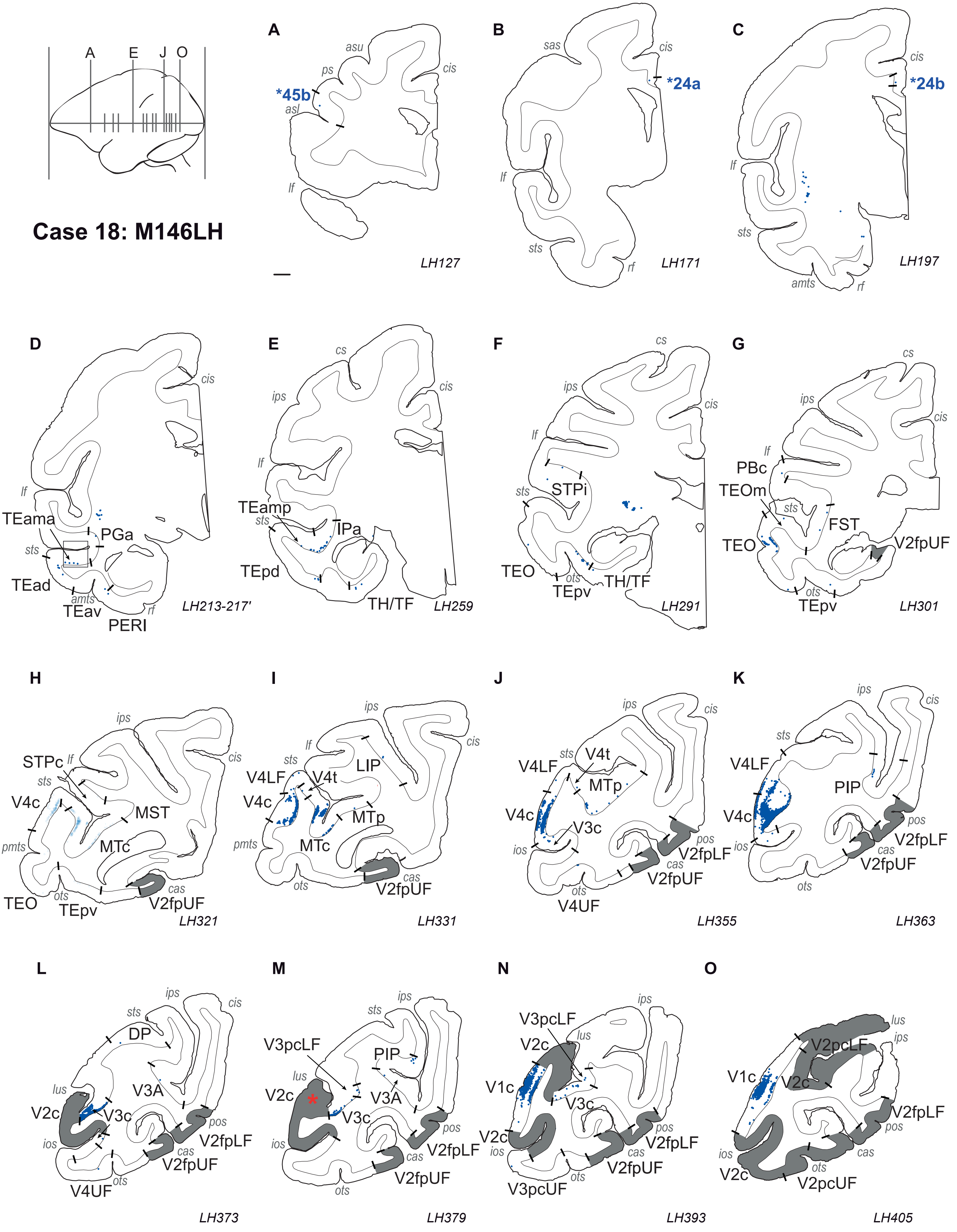
Case 18: M146LH
A
E
J
O
A
asu
ps
*45b
asl
cis
lf
LH127
B
sas
cis
*24a
lf
sts
rf
LH171
C
cis
*24b
lf
sts
amts
rf
LH197
D
cis
lf
TEama
PGa
sts
TEad
amts
TEav
rf
PERI
LH213-217'
E
cs
ips
cis
lf
TEamp
sts
IPa
TEpd
TH/TF
LH259
F
cs
ips
cis
lf
STPi
sts
TEO
ots
TEpv
TH/TF
LH291
G
cs
ips
cis
lf
PBc
TEOm
sts
FST
TEO
V2fpUF
ots
TEpv
LH301
H
ips
STPc
lf
cis
sts
V4c
MST
pmts
MTc
TEO
ots
TEpv
cas
V2fpUF
LH321
I
ips
lf
cis
sts
V4LF
V4t
LIP
V4c
MTp
pmts
MTc
ots
cas
V2fpUF
LH331
J
ips
cis
sts
V4t
V4LF
MTp
V4c
V3c
ios
pos
V2fpLF
cas
V2fpUF
ots
V4UF
LH355
K
ips
cis
sts
V4LF
PIP
V4c
pos
ios
V2fpLF
cas
V2fpUF
ots
LH363
L
ips
sts
cis
DP
lus
V3A
V2c
V3c
pos
ios
V2fpLF
cas
V2fpUF
ots
V4UF
LH373
M
ips
sts
V3pcLF
PIP
lus
V2c
V3A
V3c
pos
ios
V2fpLF
cas
V2fpUF
ots
LH379
N
ips
V3pcLF
lus
V2c
V1c
V3c
V2c
pos
ios
V2fpLF
cas
V2fpUF
ots
V3pcUF
LH393
O
lus
ips
V2pcLF
V1c
V2c
V2fpLF
pos
ios
cas
V2c
ots
V2pcUF
LH405

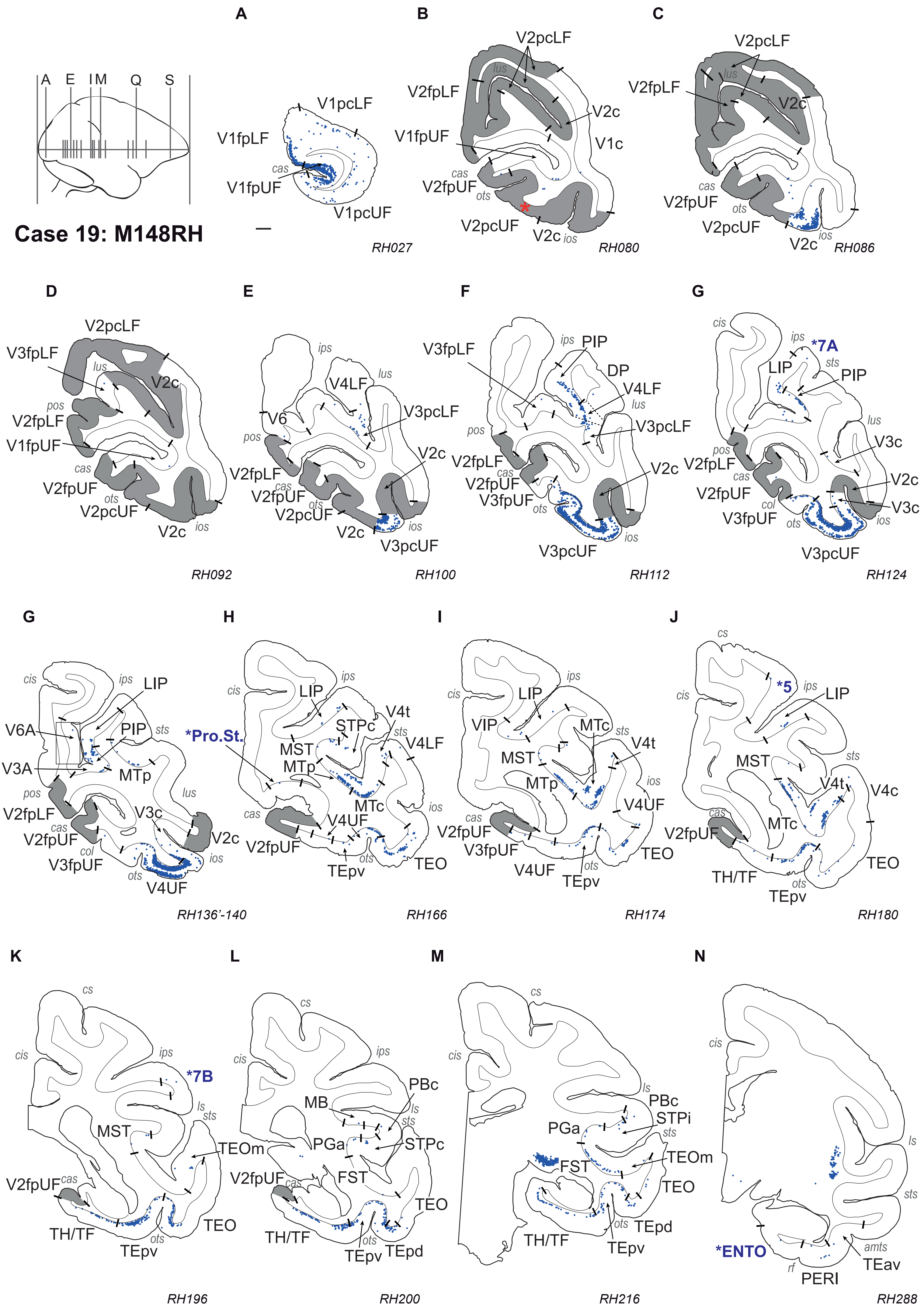
Case 19: M148RH
A
E
I M
Q
S
V1pcLF
V1fpLF
V1fpUF
V1pcUF
cas
RH027
B
V2pcLF
V2fpLF
V1fpUF
V2c
V1c
V2fpUF
V2pcUF
lus
cas
ots
ios
RH080
C
RH086
D
V3fpLF
pos
RH092
E
V6
V4LF
V3pcLF
V3pcUF
ips
RH100
F
PIP
DP
RH112
G
*7A
LIP
V3c
V3fpUF
col
sts
cis
RH124
V6A
V3A
MTp
V4UF
RH136'-140
H
*Pro.St.
STPc
V4t
MST
MTc
TEpv
TEO
RH166
I
VIP
RH174
J
*5
V4c
TH/TF
cs
RH180
K
*7B
TEOm
ls
RH196
L
PBc
MB
PGa
FST
TEpd
RH200
M
STPi
RH216
N
*ENTO
PERI
TEav
amts
rf
RH288

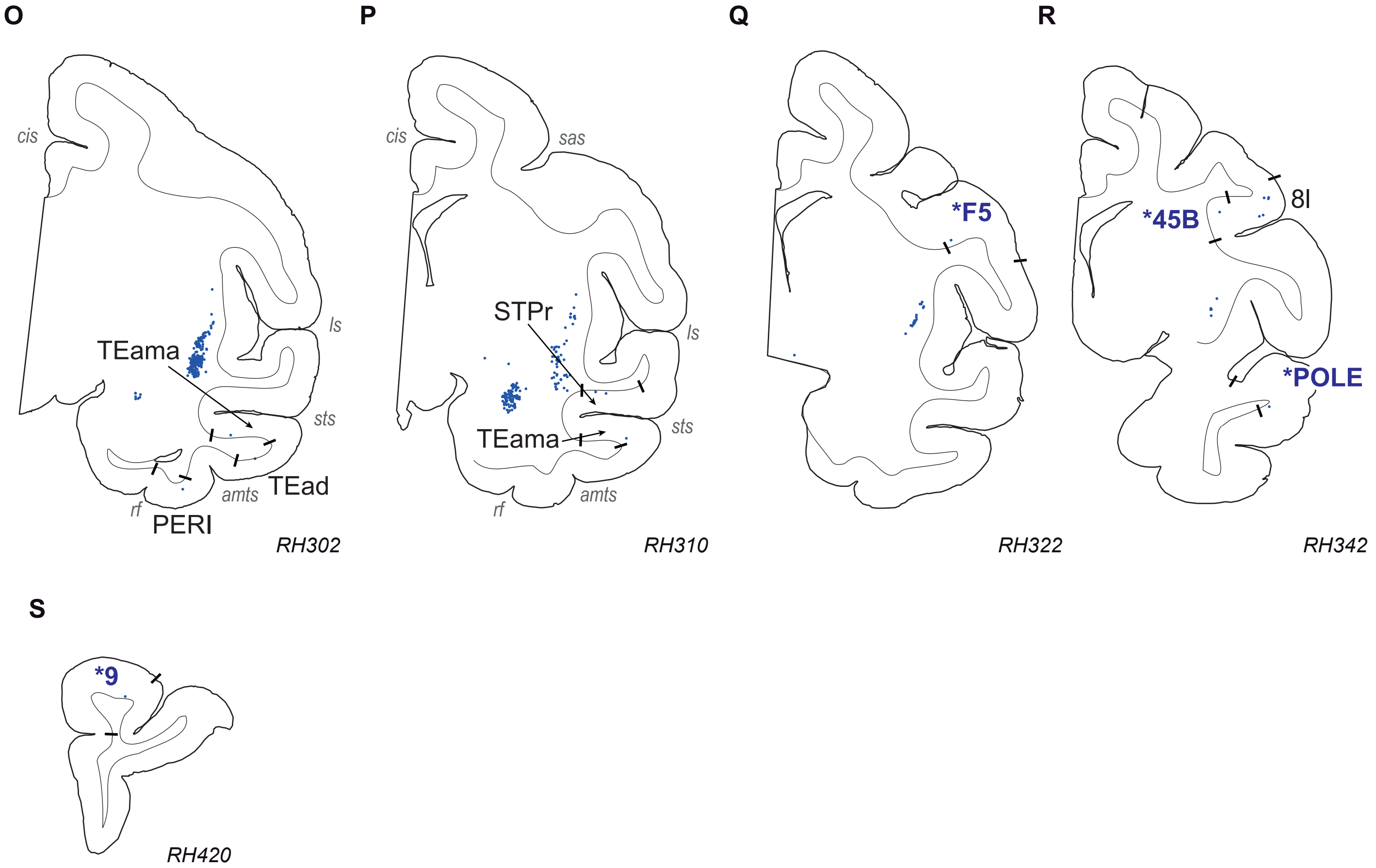

O
cis
ls
TEama
sts
TEad
amts
rf
PERI
RH302
P
cis
sas
STPr
ls
sts
TEama
amts
rf
RH310
Q
*F5
RH322
R
*45B
8l
*POLE
RH342
S
*9
RH420

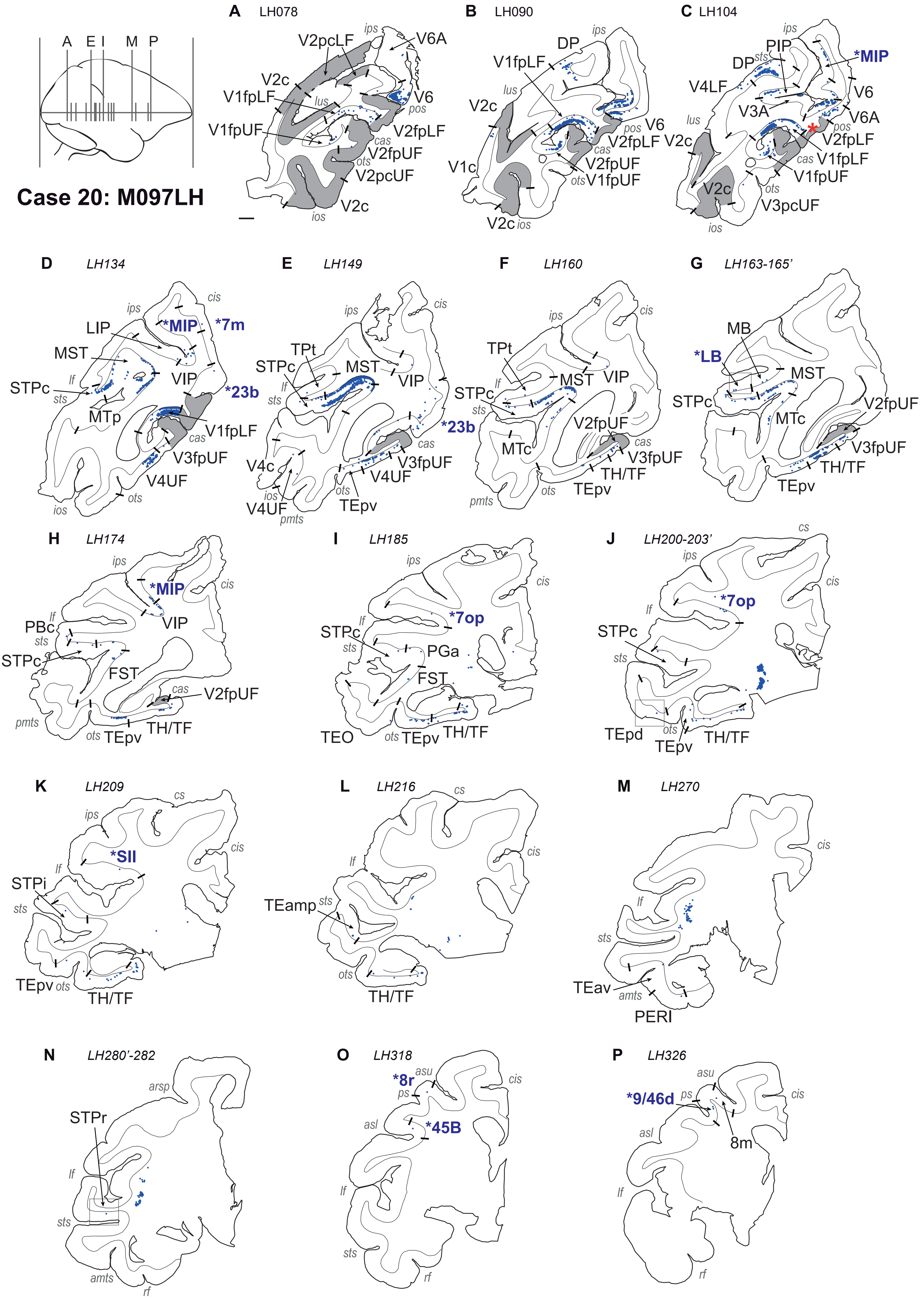
Case 20: M097LH
A LH078
B LH090
C LH104
D LH134
E LH149
F LH160
G LH163-165'
H LH174
I LH185
J LH200-203'
K LH209
L LH216
M LH270
N LH280'-282
O LH318
P LH326
*MIP
*7m
*23b
*LB
*7op
*SII
*8r
*45B
*9/46d
V2pcLF
V6A
V2c
V1fpLF
V1fpUF
V6
V2fpLF
V2fpUF
V2pcUF
DP
PIP
V4LF
V3A
V3pcUF
V1c
LIP
MST
STPc
VIP
MTp
V3fpUF
V4UF
TPt
V4c
TEpv
MTc
TH/TF
MB
PBc
FST
PGa
TEO
TEpd
STPi
TEamp
TEav
PERI
STPr
8m

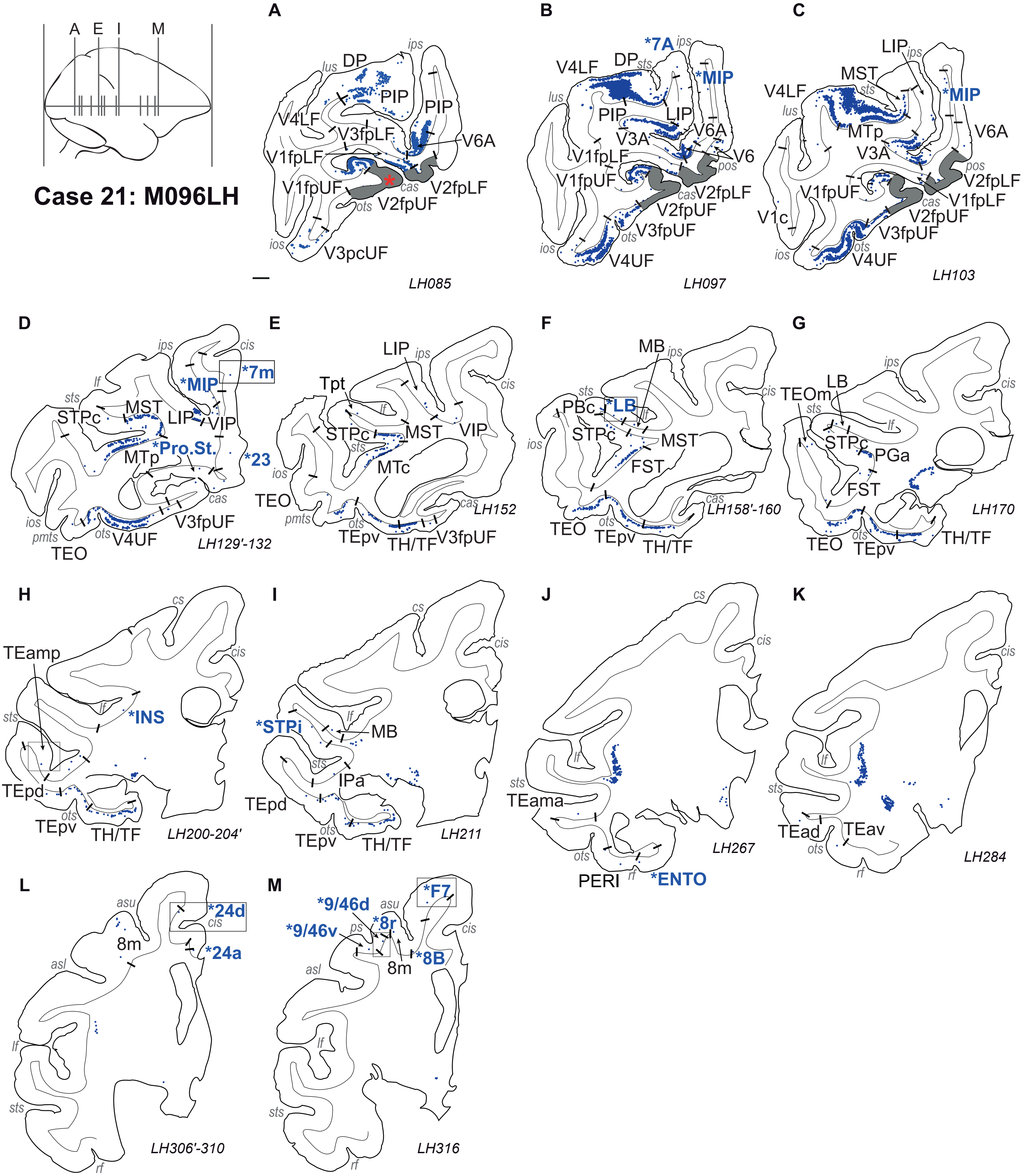
A E I M
Case 21: M096LH
A
B
C
D
E
F
G
H
I
J
K
L
M
LH085
LH097
LH103
LH129'-132
LH152
LH158'-160
LH170
LH200-204'
LH211
LH267
LH284
LH306'-310
LH316
*7A
*MIP
*7m
*Pro.St.
*23
*LB
*INS
*STPi
*ENTO
*24d
*24a
*F7
*9/46d
*9/46v
*8r
*8B
DP
PIP
V4LF
V3fpLF
V6A
V1fpLF
V1fpUF
V2fpLF
V2fpUF
V3pcUF
LIP
V3A
V6
V3fpUF
V4UF
MST
MTp
V1c
STPc
VIP
TEO
Tpt
MTc
TEpv
TH/TF
PBc
MB
FST
TEOm
LB
PGa
TEamp
TEpd
IPa
TEama
PERI
TEad
TEav
8m
