## Supplementary material for "Retinotopic organization of feedback projections in primate early visual cortex: implications for active vision": Figure S3

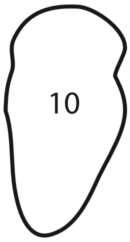
10

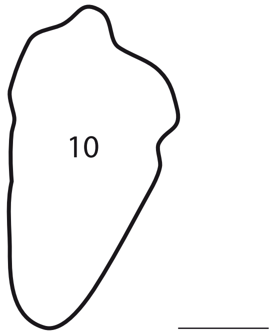

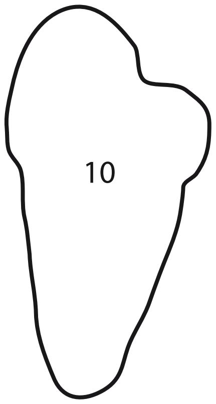
10

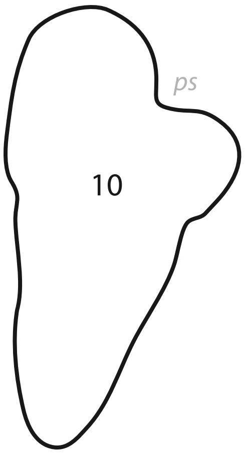
ps
10

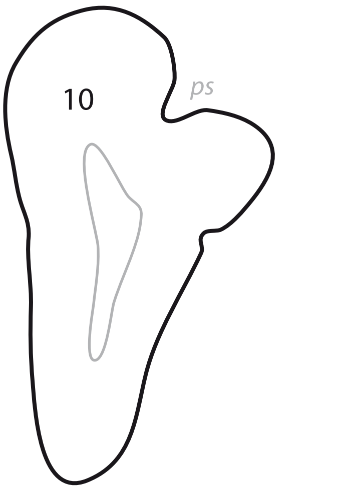
10
ps

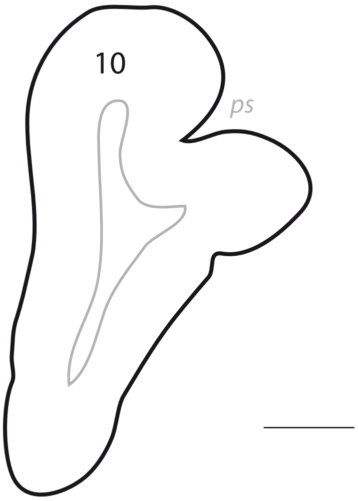
10
ps

# p. 007

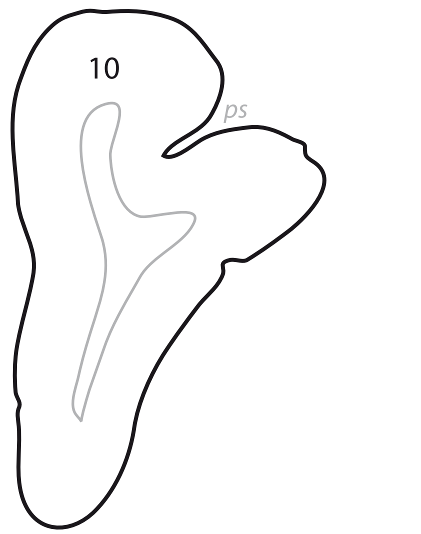

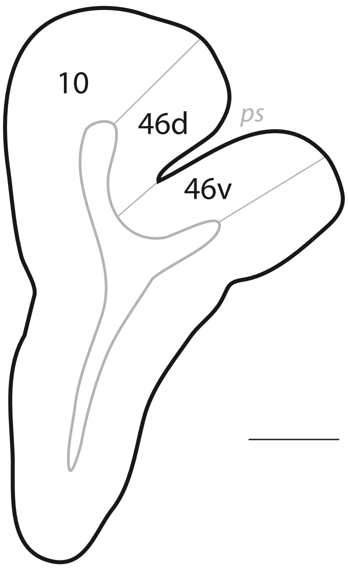
10
46d
ps
46v

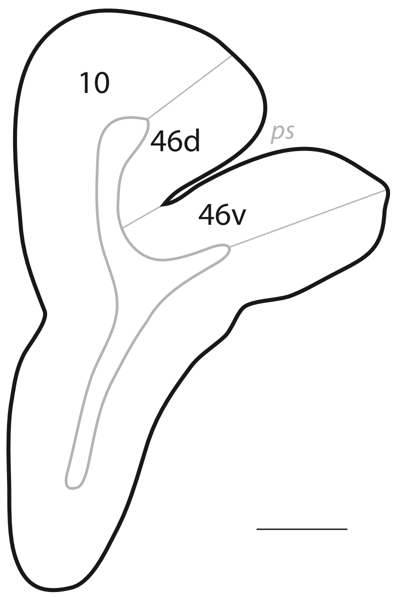
10
46d
ps
46v

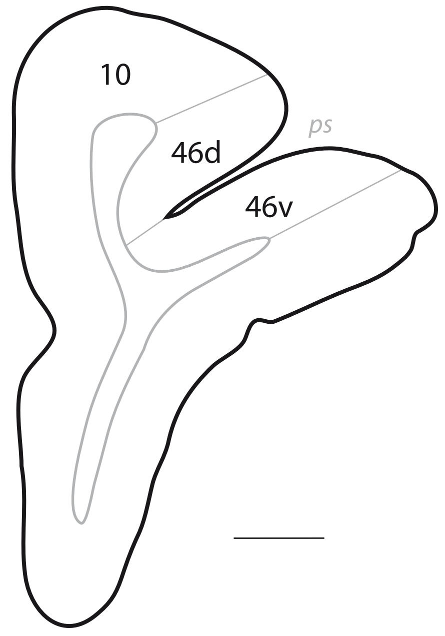
10
46d
ps
46v

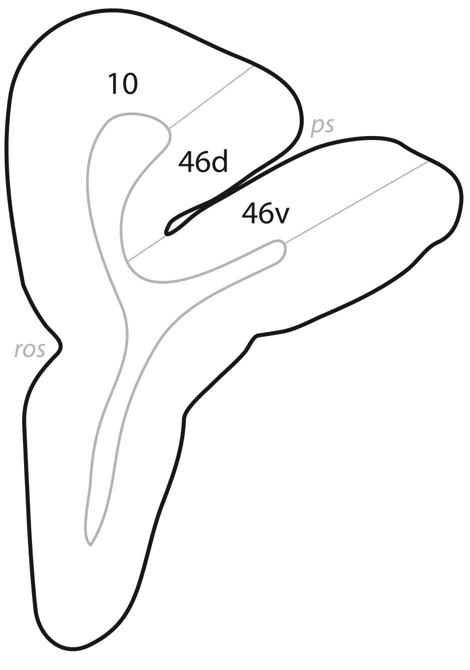
10
ps
46d
46v
ros

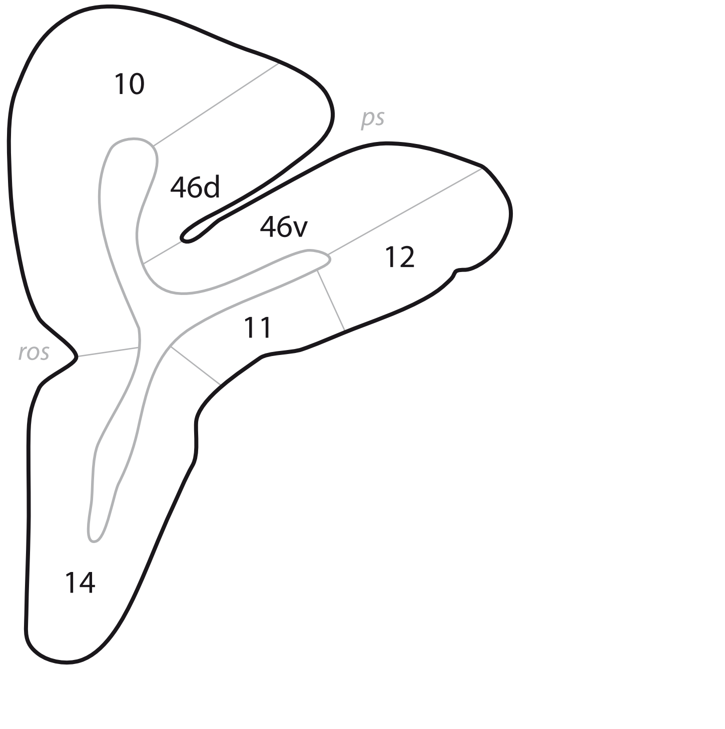
10
ps
46d
46v
12
11
ros
14

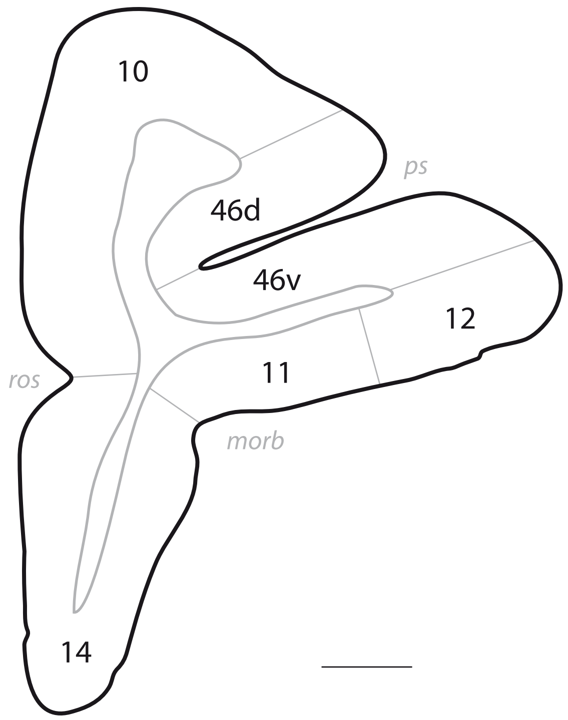
10
ps
46d
46v
12
ros
11
morb
14

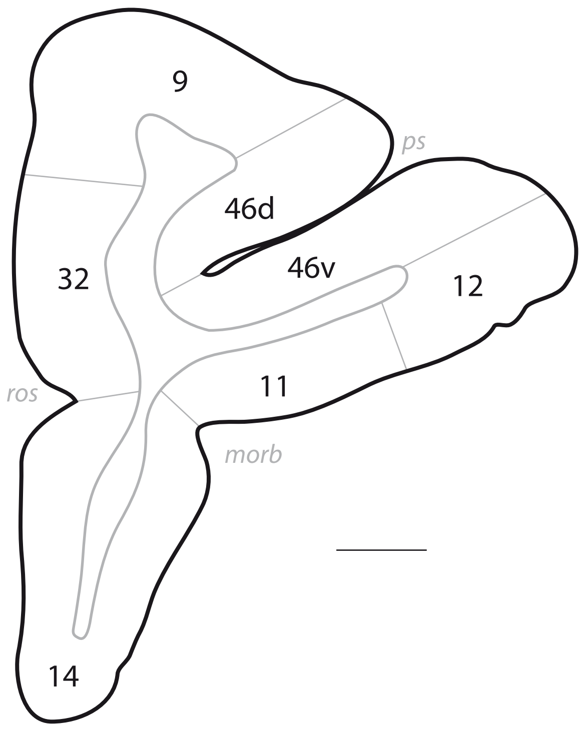
9
ps
46d
46v
32
12
ros
11
morb
14

# p. 015

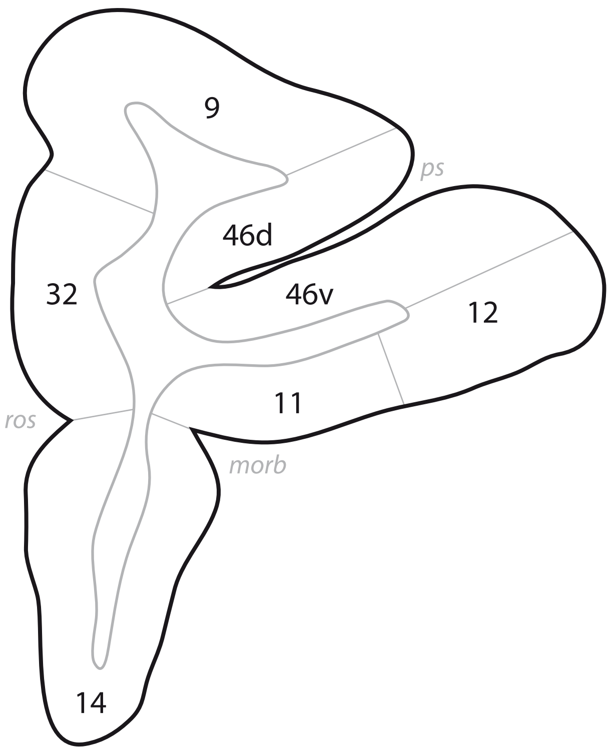

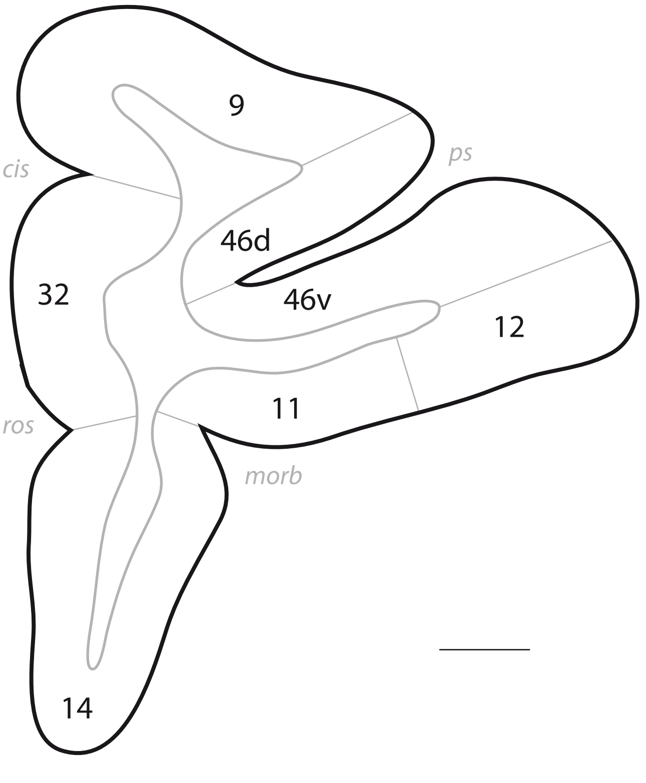
9
cis
ps
46d
32
46v
12
11
ros
morb
14

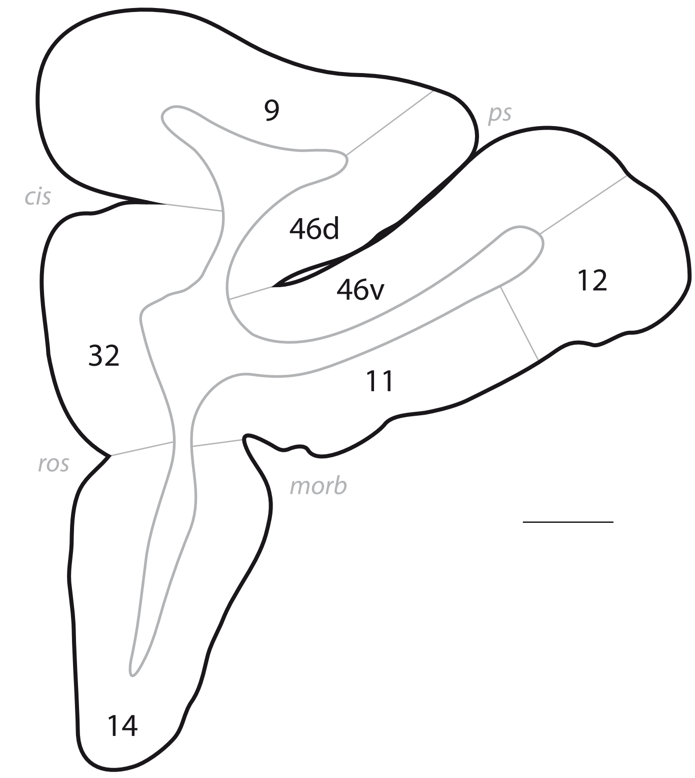
9
ps
cis
46d
46v
12
32
11
ros
morb
14

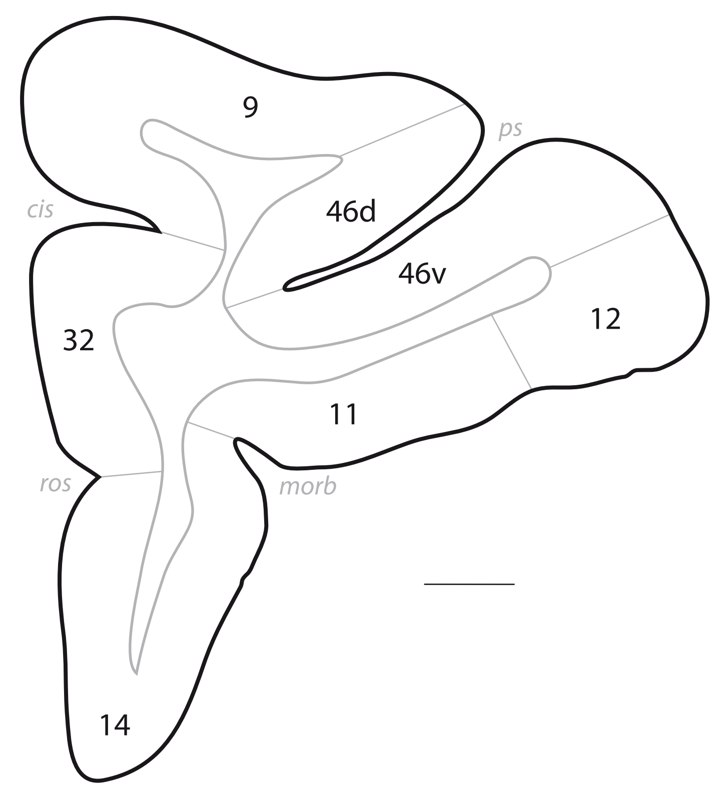
9
ps
cis
46d
46v
12
32
11
ros
morb
14

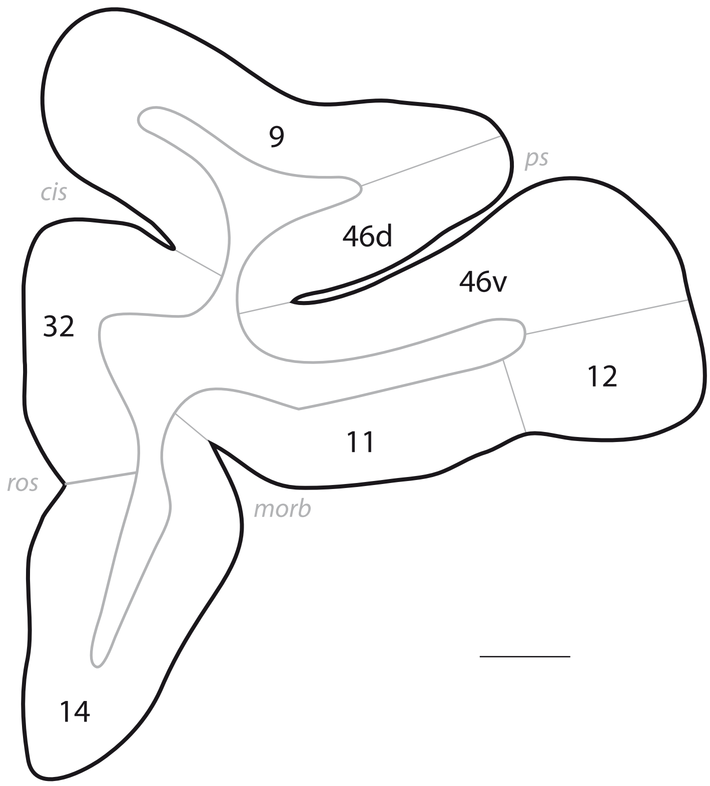
9
ps
cis
46d
46v
32
12
11
ros
morb
14

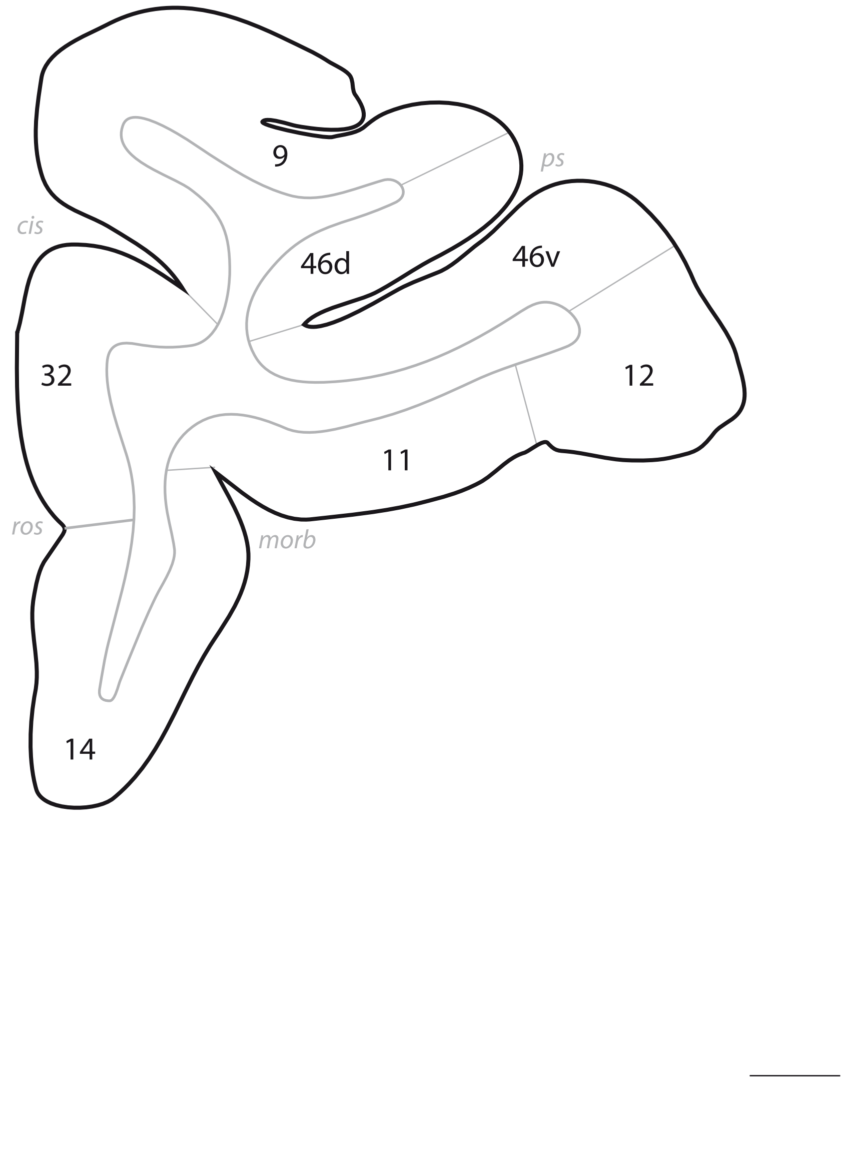
9
ps
cis
46d
46v
32
12
11
ros
morb
14

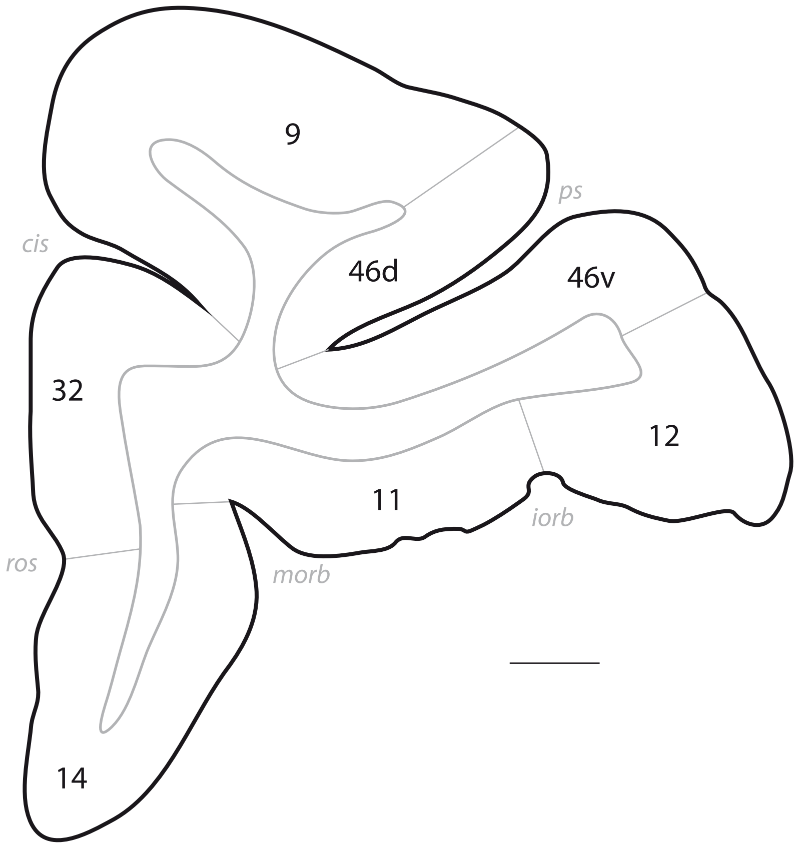
9
ps
cis
46d
46v
32
12
11
iorb
ros
morb
14

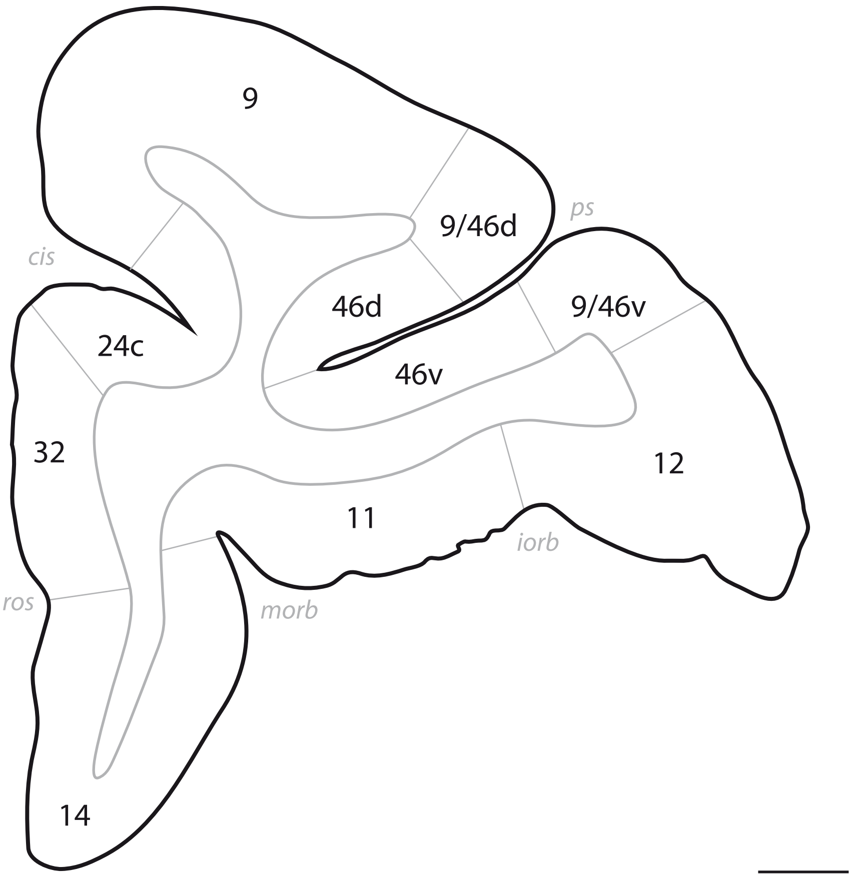
9
9/46d
ps
cis
46d
9/46v
24c
46v
32
12
11
iorb
ros
morb
14

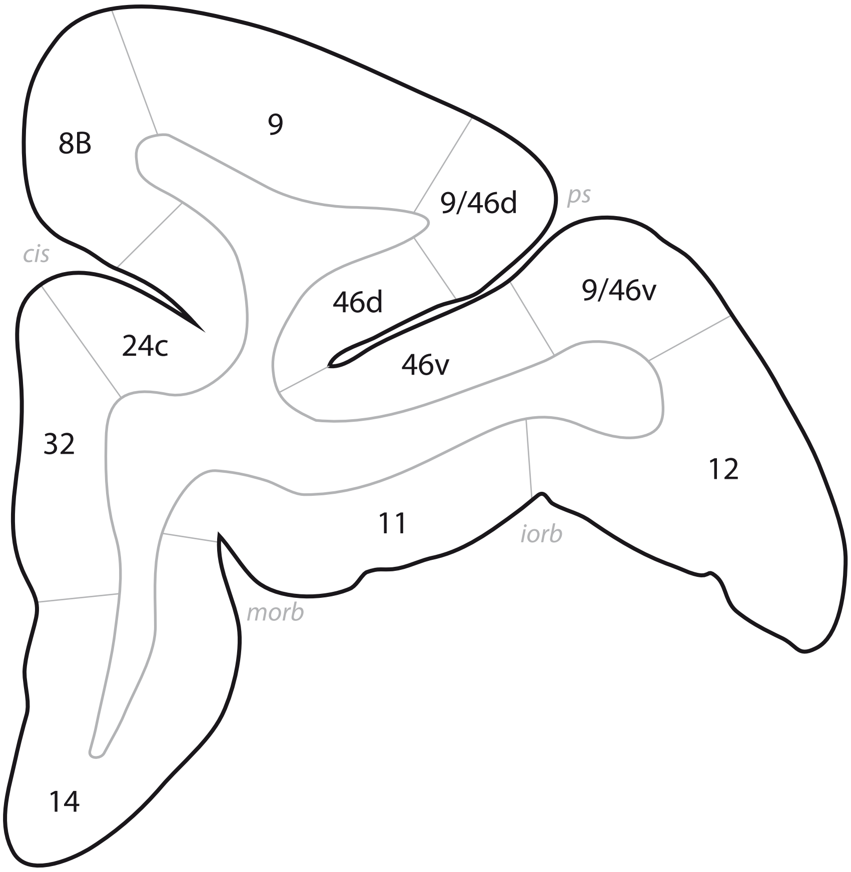
8B
9
9/46d
ps
cis
46d
9/46v
24c
46v
32
12
11
iorb
morb
14

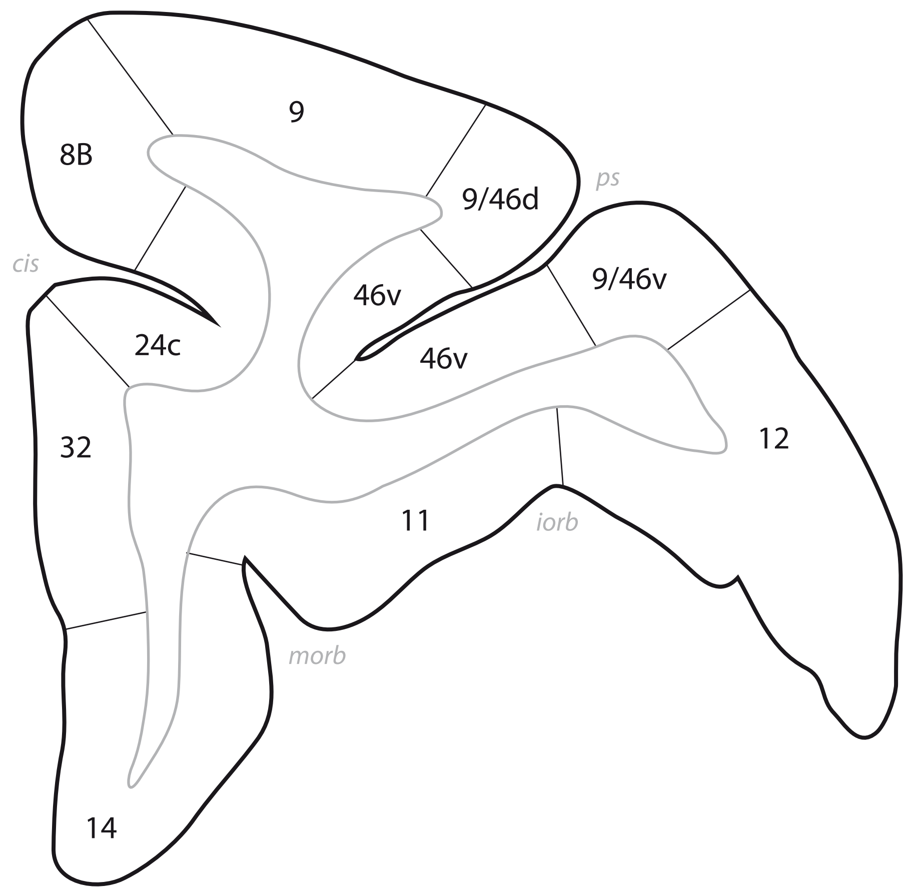
9
8B
ps
9/46d
cis
9/46v
46v
24c
46v
12
32
11
iorb
morb
14

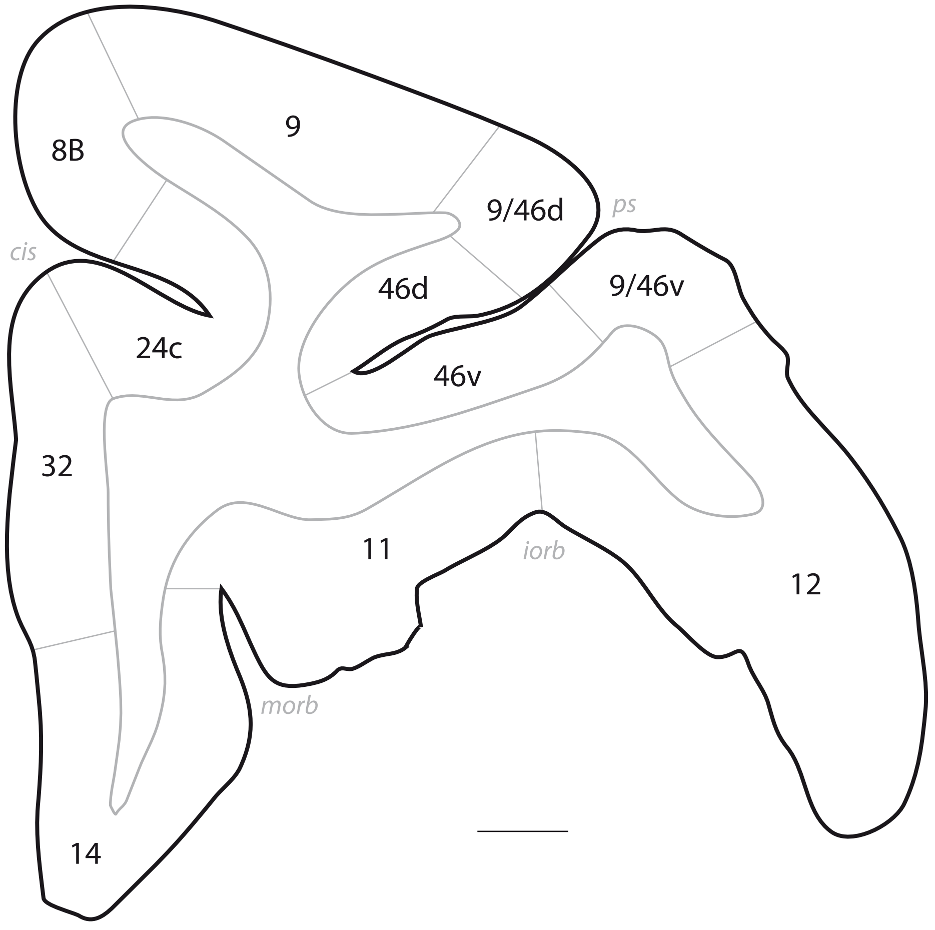
9
8B
9/46d
ps
cis
46d
9/46v
24c
46v
32
11
iorb
12
morb
14

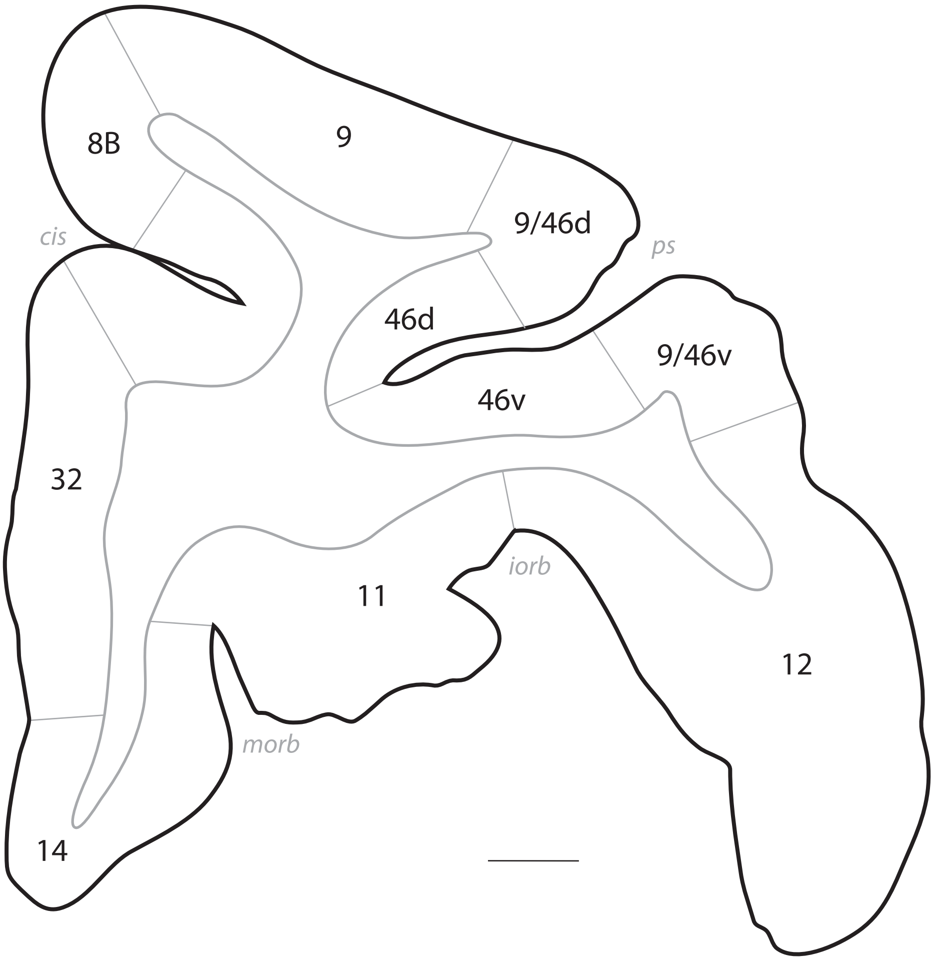
8B
9
9/46d
cis
ps
46d
9/46v
46v
32
11
iorb
12
morb
14

# p. 027

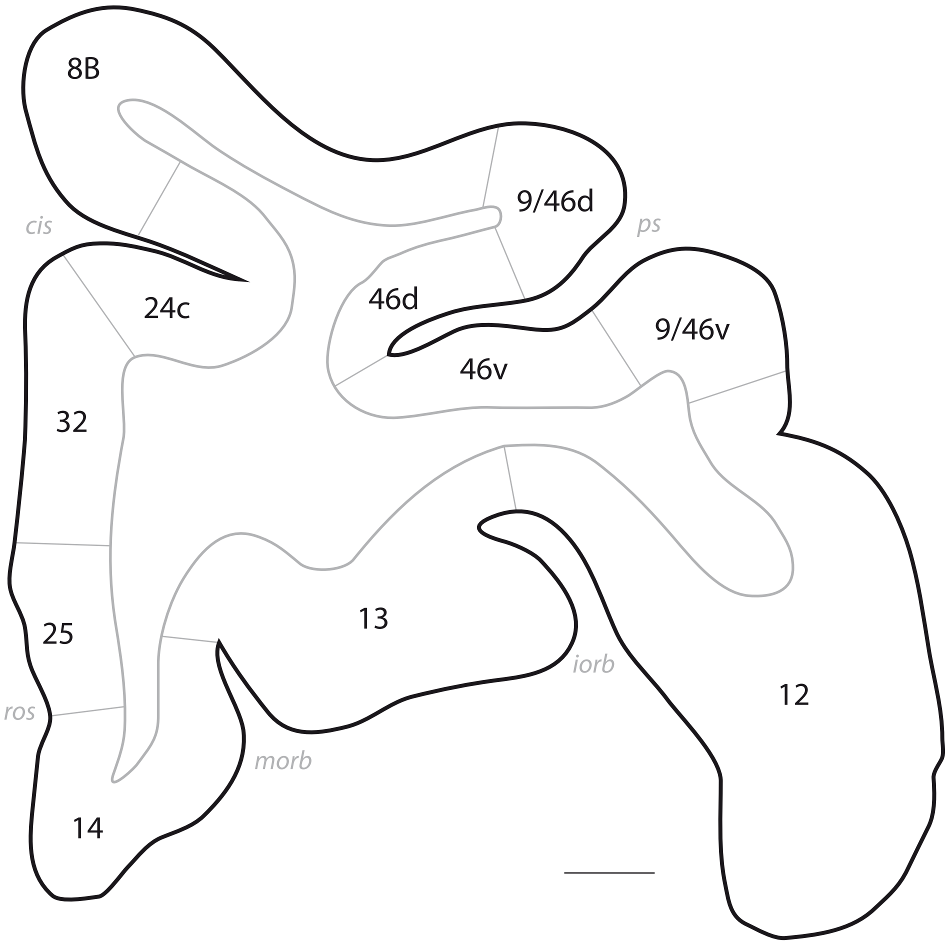

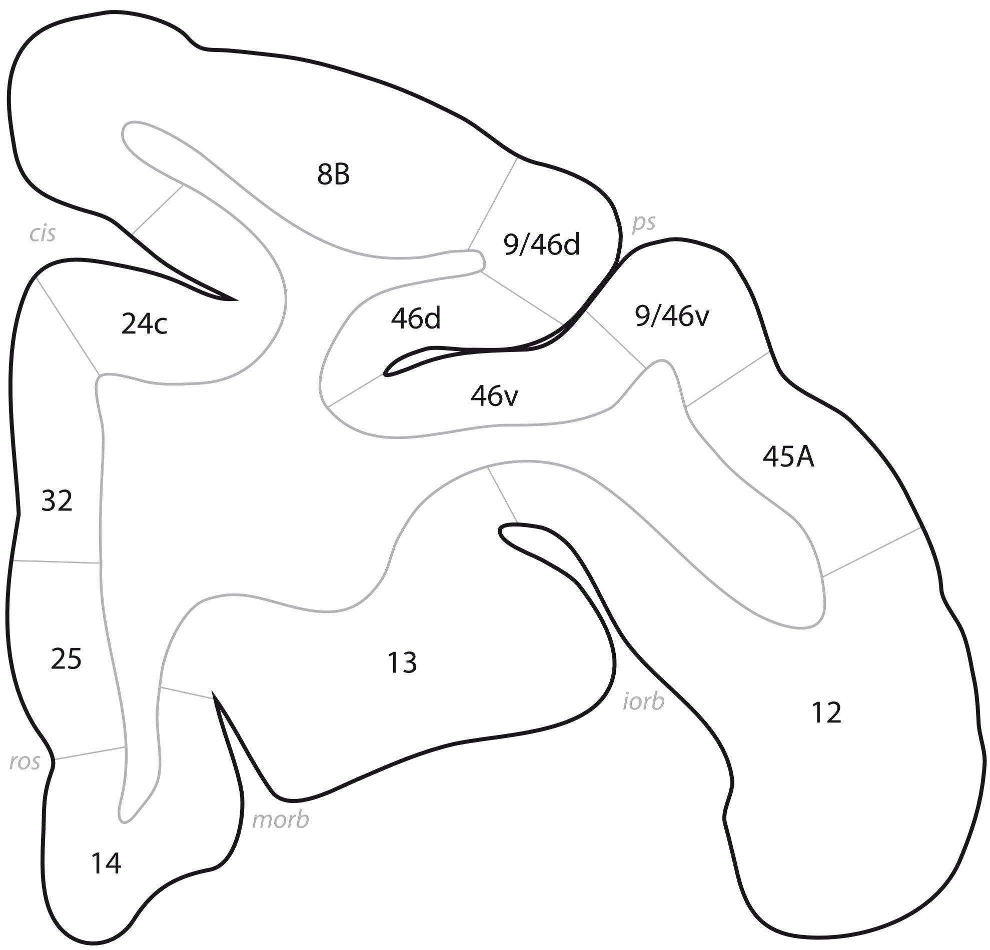
8B
9/46d
ps
cis
24c
46d
9/46v
46v
45A
32
25
13
iorb
12
ros
morb
14

# p. 029

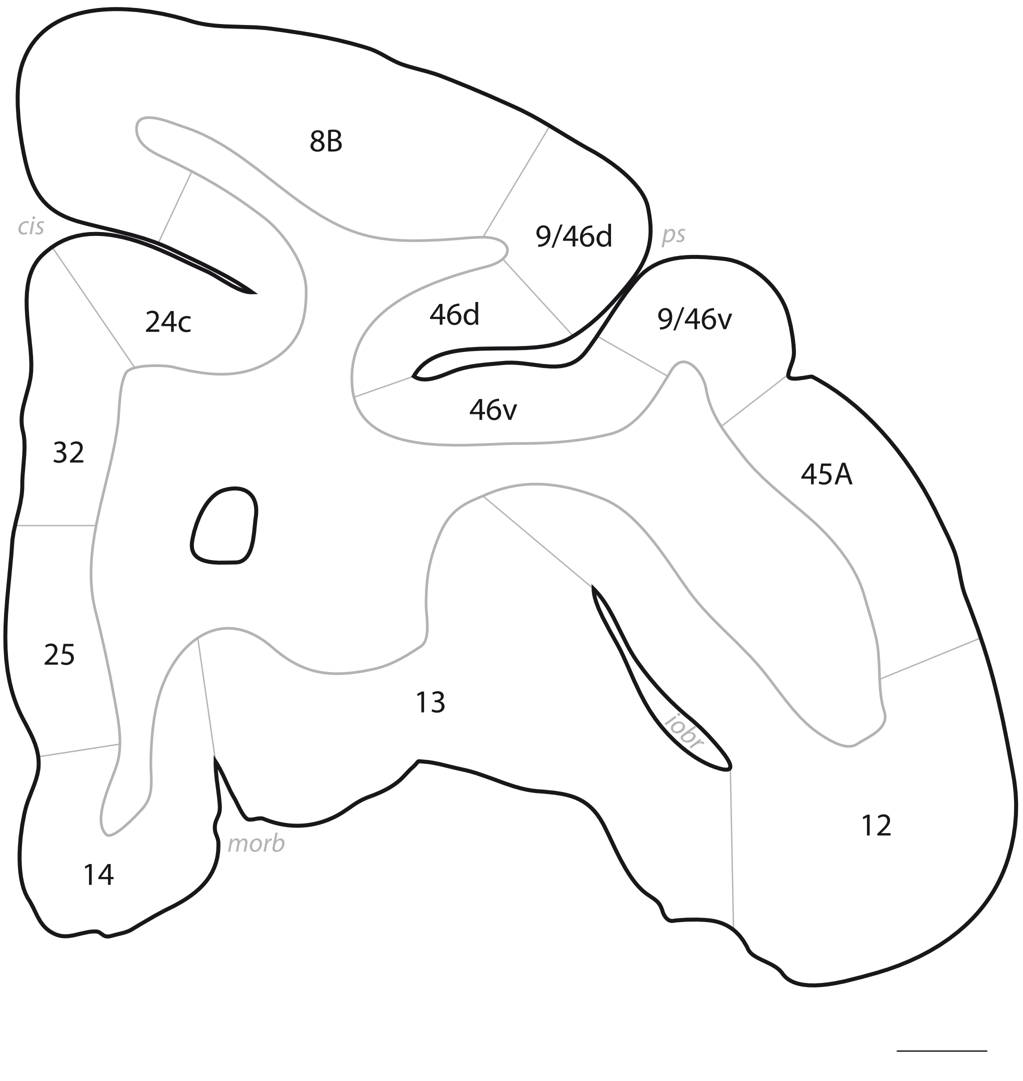

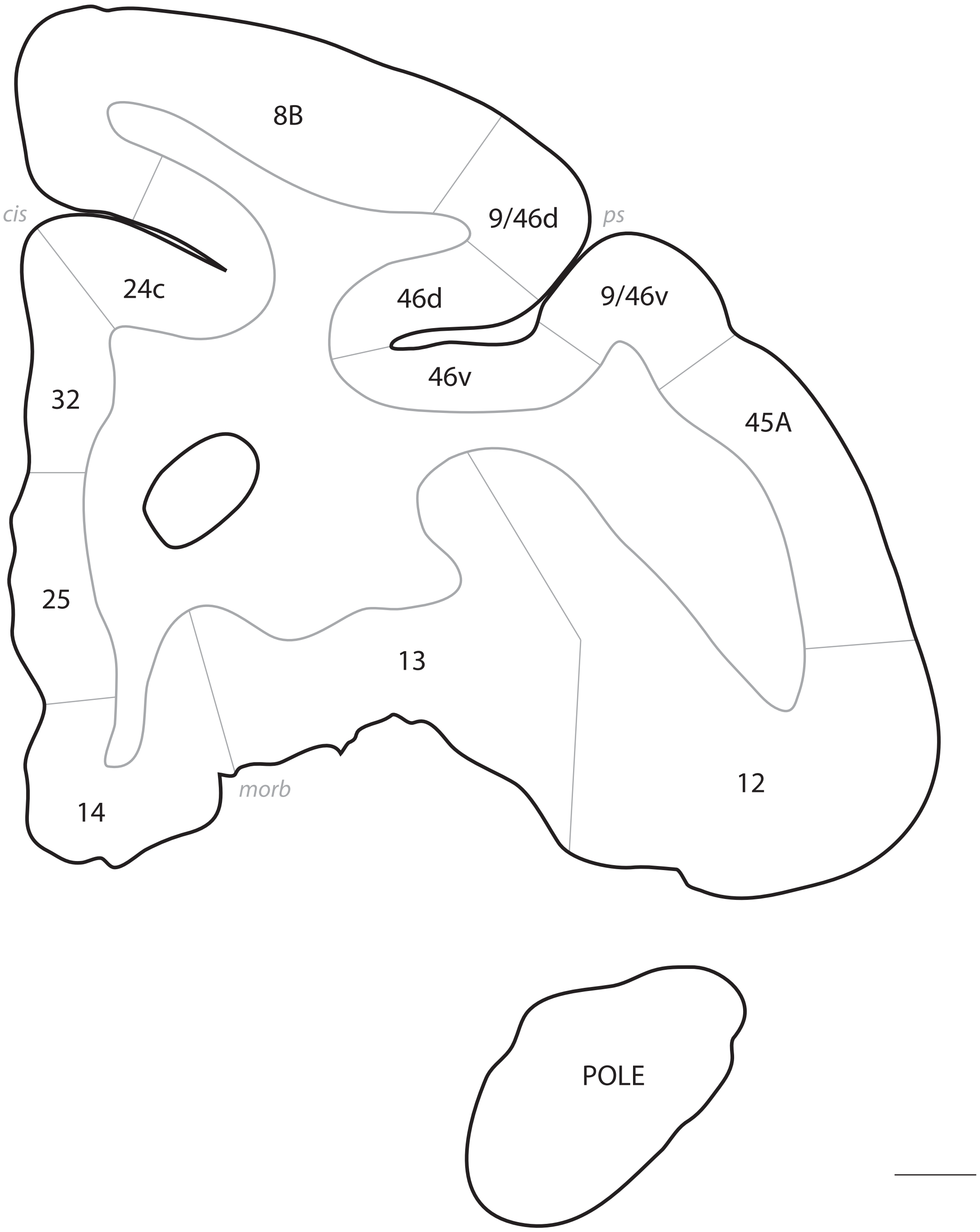
8B
cis
9/46d
ps
24c
46d
9/46v
46v
32
45A
25
13
12
14
morb
POLE

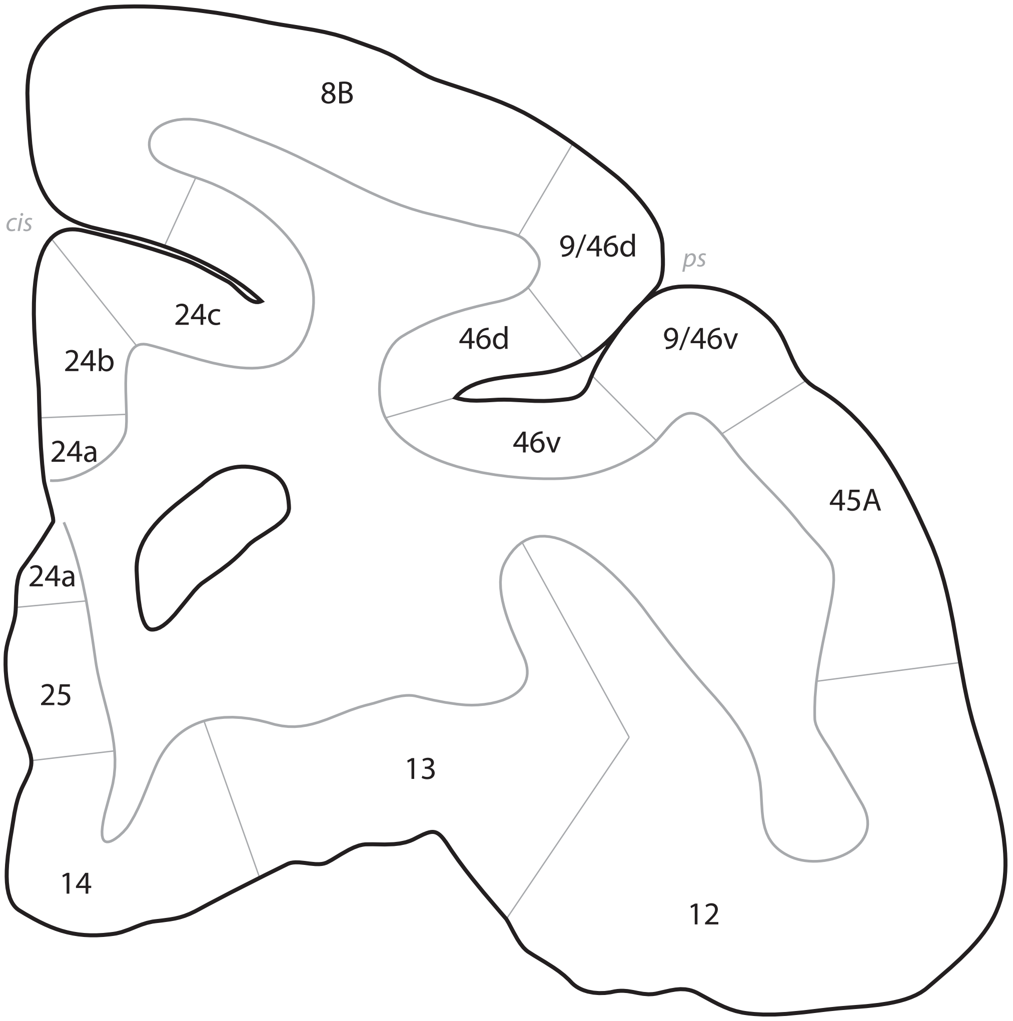
8B
cis
9/46d
ps
24c
46d
9/46v
24b
46v
24a
45A
24a
25
13
14
12

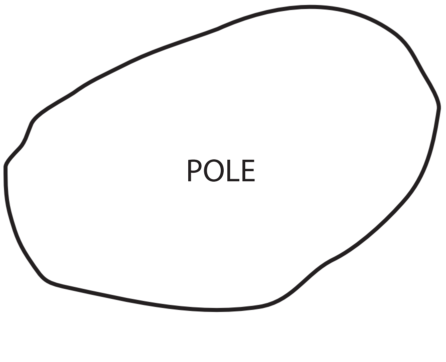
POLE

# p. 032

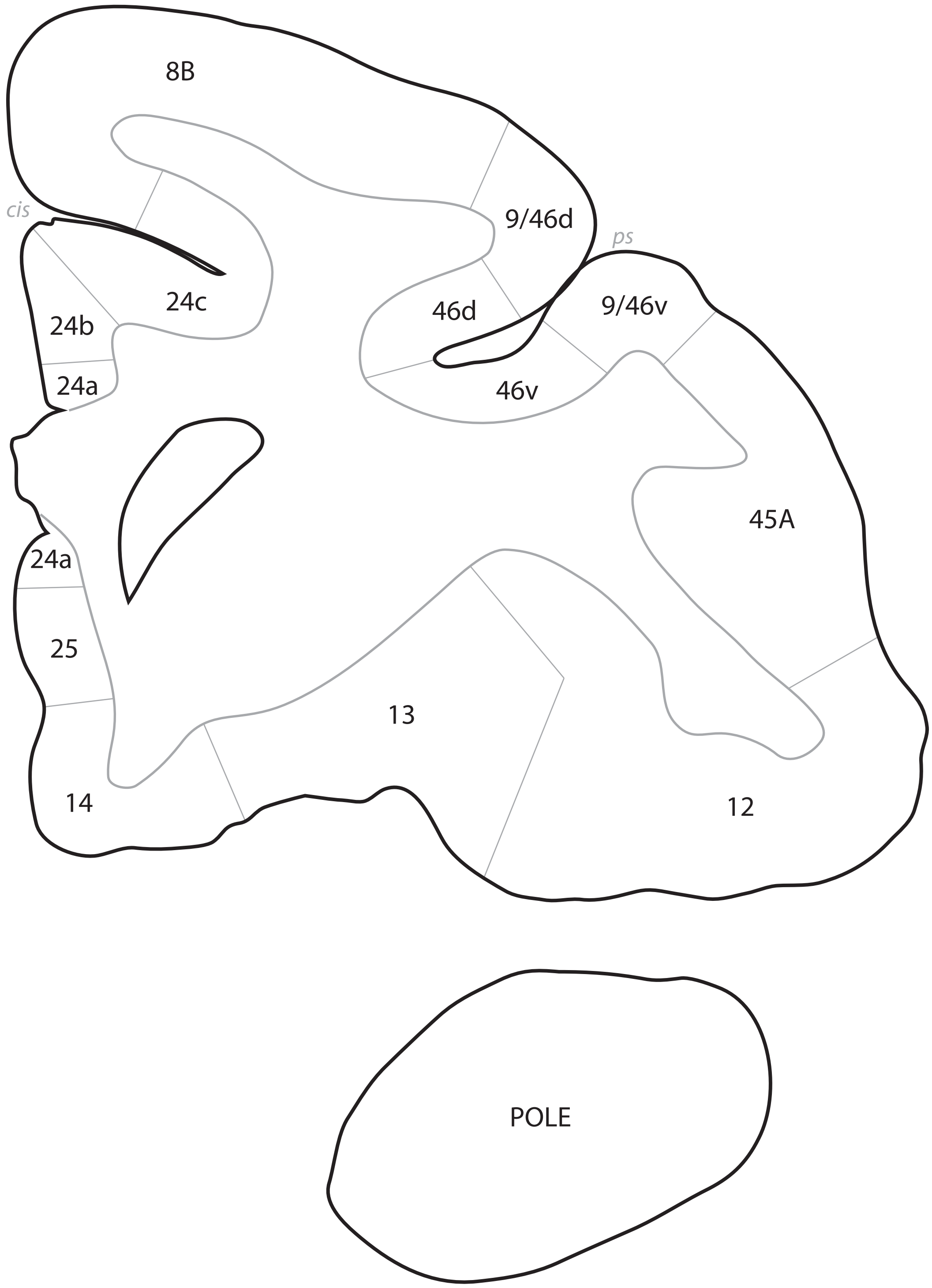

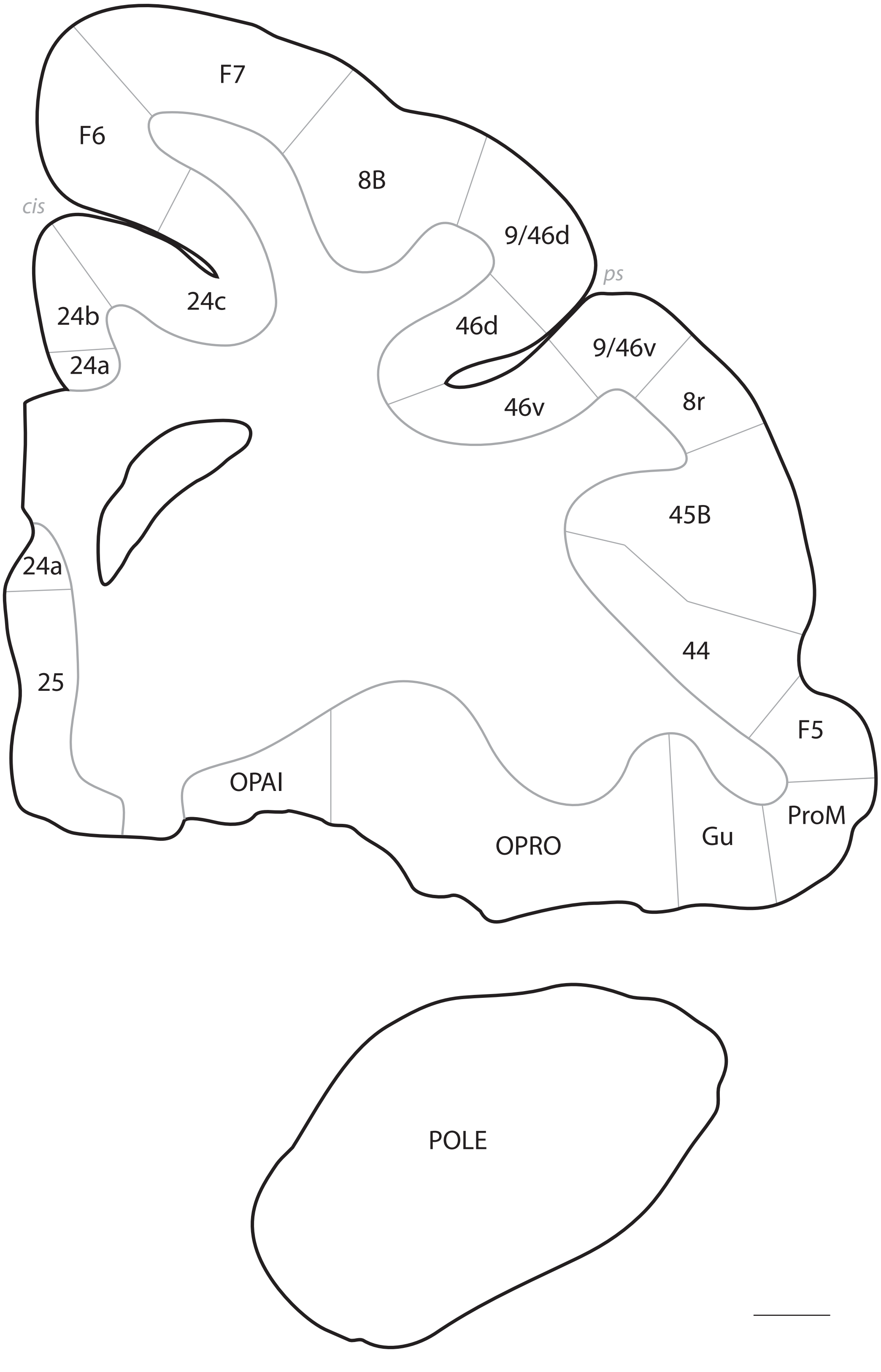
F7
F6
8B
cis
9/46d
ps
24c
24b
24a
46d
9/46v
8r
46v
45B
24a
44
25
F5
OPAI
ProM
Gu
OPRO
POLE

# p. 034

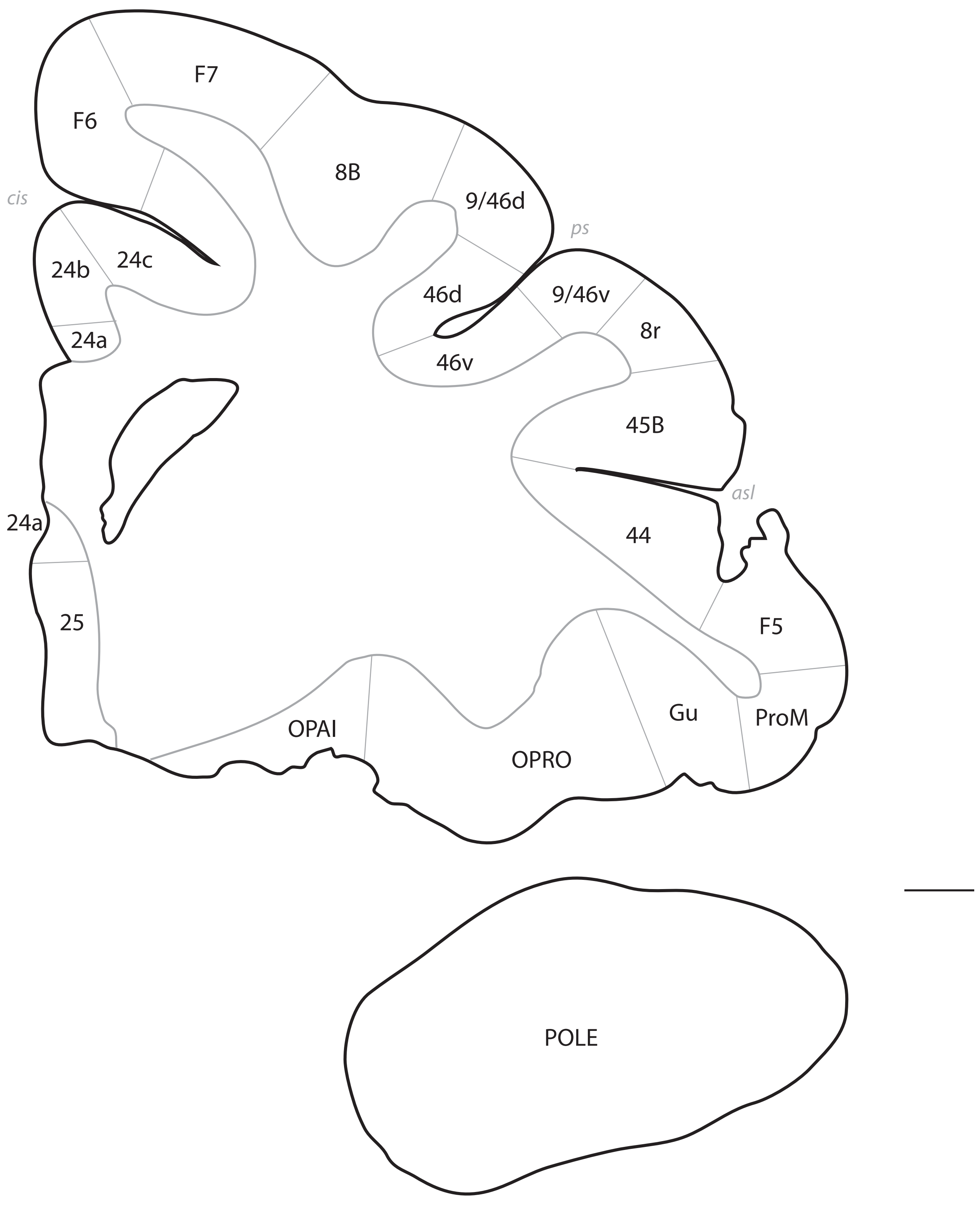

F7
F6
8B
9/46d
cis
ps
24b
24c
46d
9/46v
24a
8r
46v
45B
asl
24a
44
25
F5
Gu
ProM
OPAI
OPRO
POLE

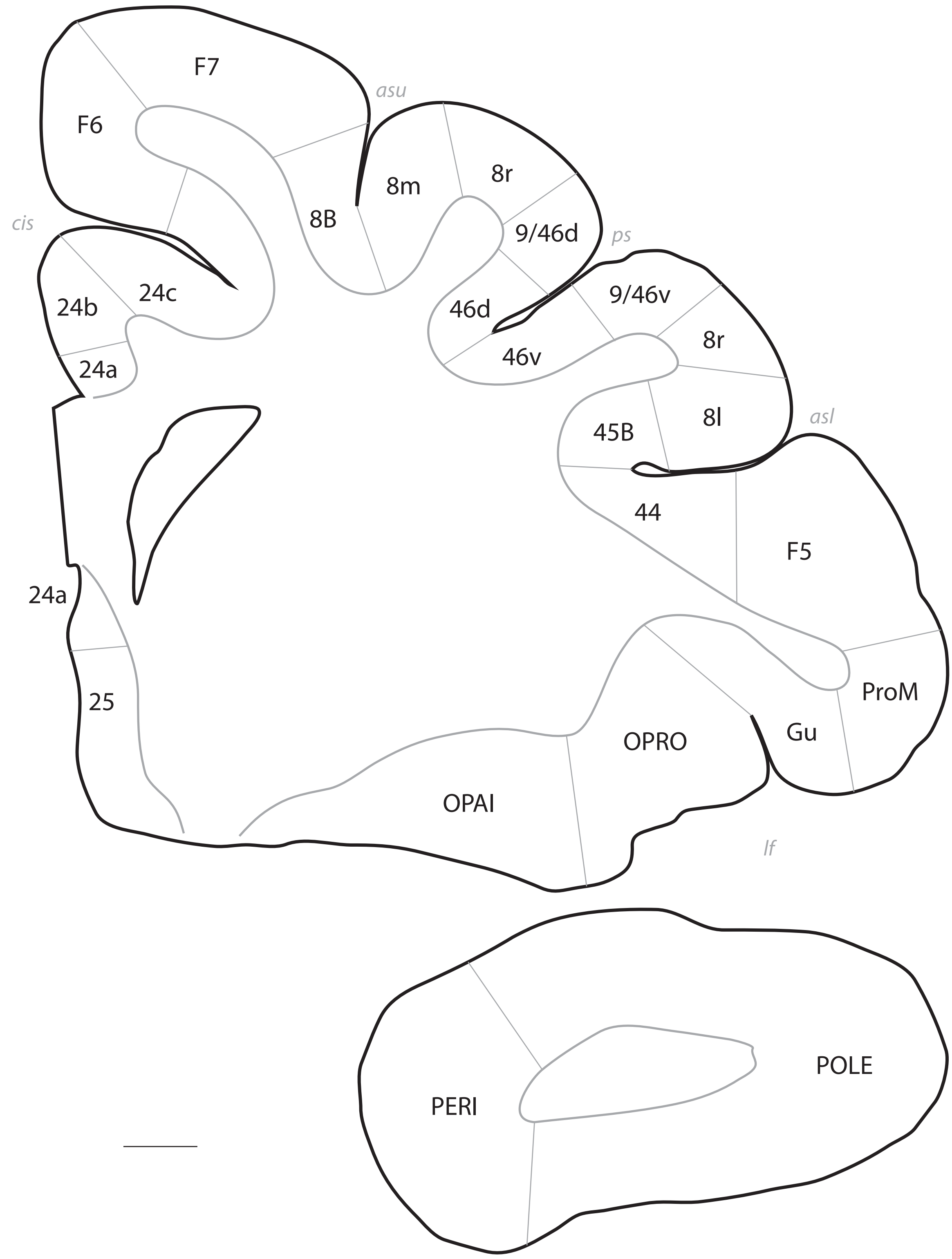
F7
asu
F6
8m
8r
8B
cis
9/46d
ps
24c
24b
9/46v
46d
8r
24a
46v
8l
asl
45B
44
F5
24a
ProM
25
Gu
OPRO
OPAI
lf
POLE
PERI

# p. 036

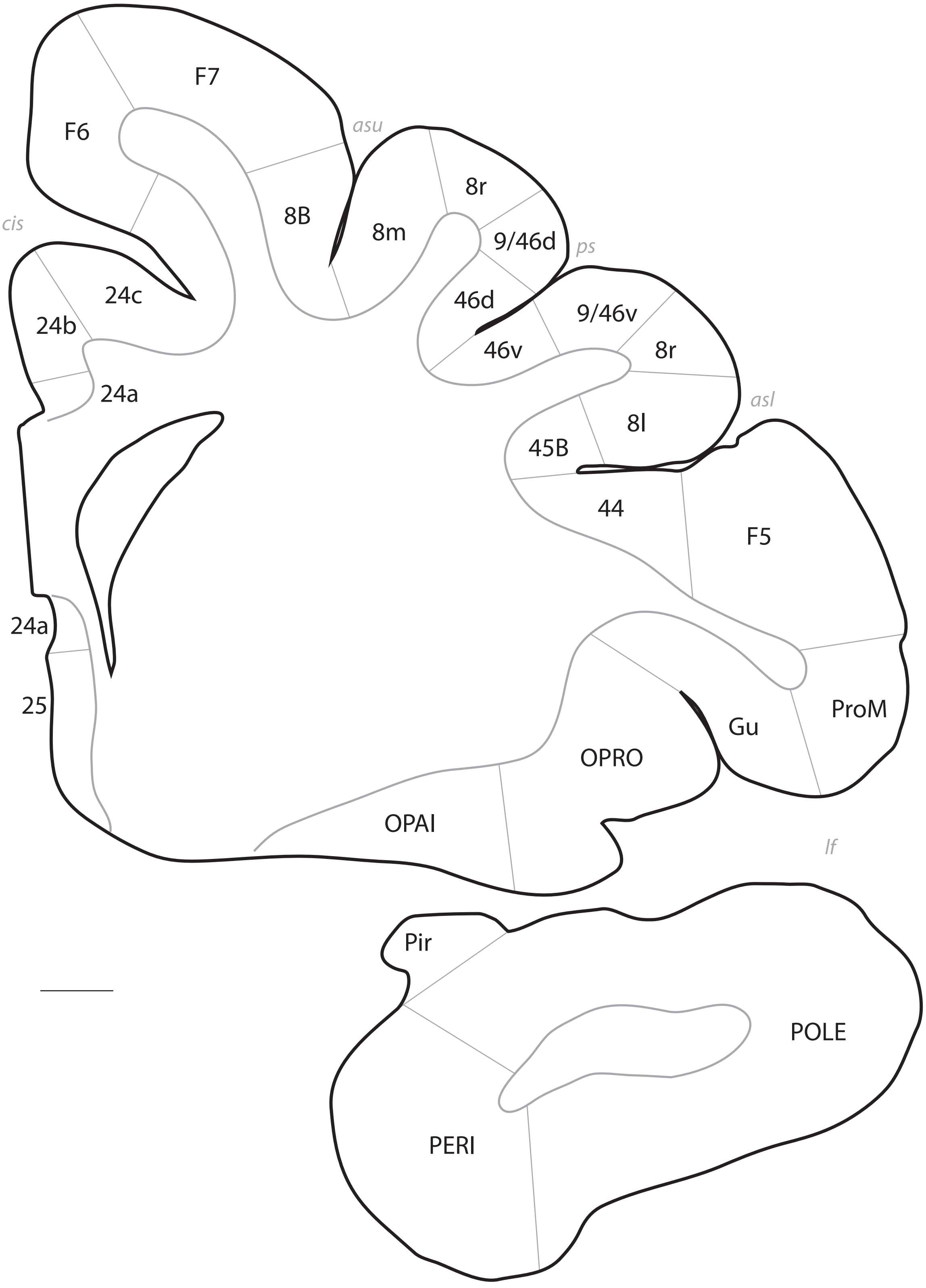
F7
asu
F6
8B
8r
cis
8m
9/46d
ps
24c
46d
9/46v
24b
46v
8r
24a
asl
8l
45B
44
F5
24a
ProM
25
Gu
OPRO
OPAI
lf
Pir
POLE
PERI

# p. 037

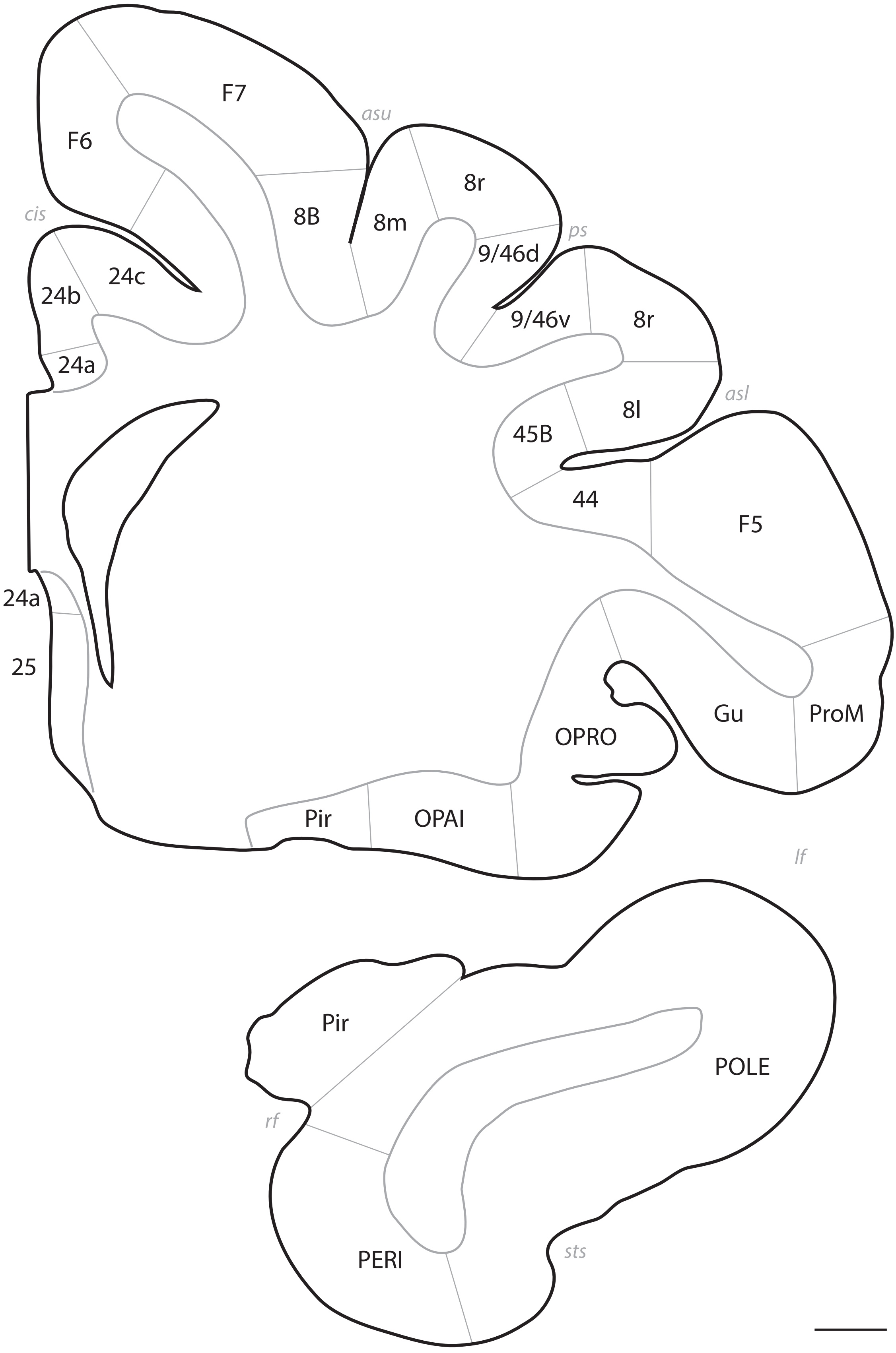
F7
asu
F6
8r
8B
8m
cis
ps
9/46d
24c
24b
9/46v
8r
24a
asl
8l
45B
44
F5
24a
25
ProM
Gu
OPRO
Pir
OPAI
lf
Pir
POLE
rf
PERI
sts

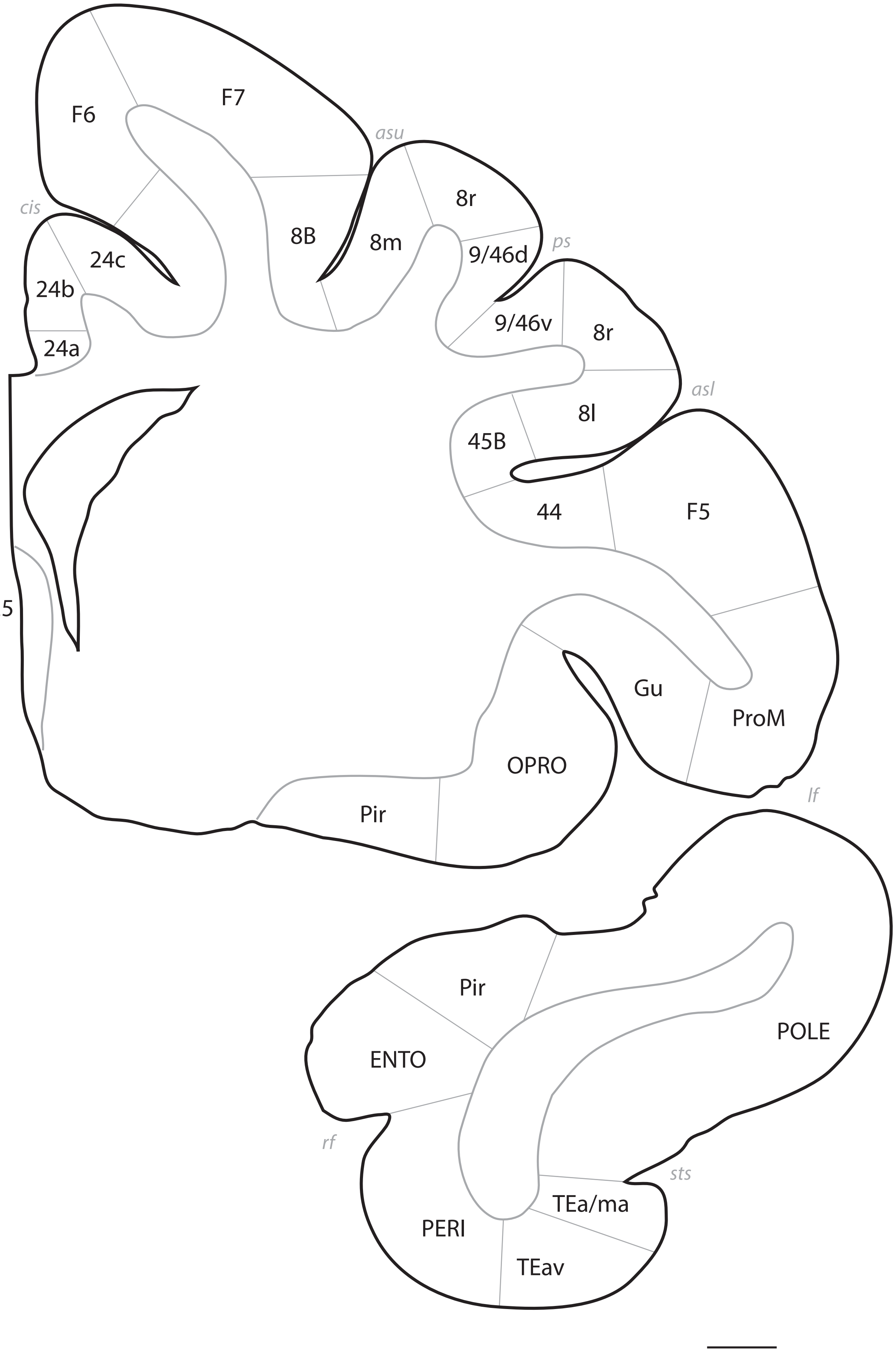
F6
F7
asu
cis
24c
24b
24a
8B
8m
8r
9/46d
ps
9/46v
8r
asl
8l
45B
44
F5
25
Gu
ProM
OPRO
Pir
lf
Pir
ENTO
POLE
rf
sts
TEa/ma
PERI
TEav

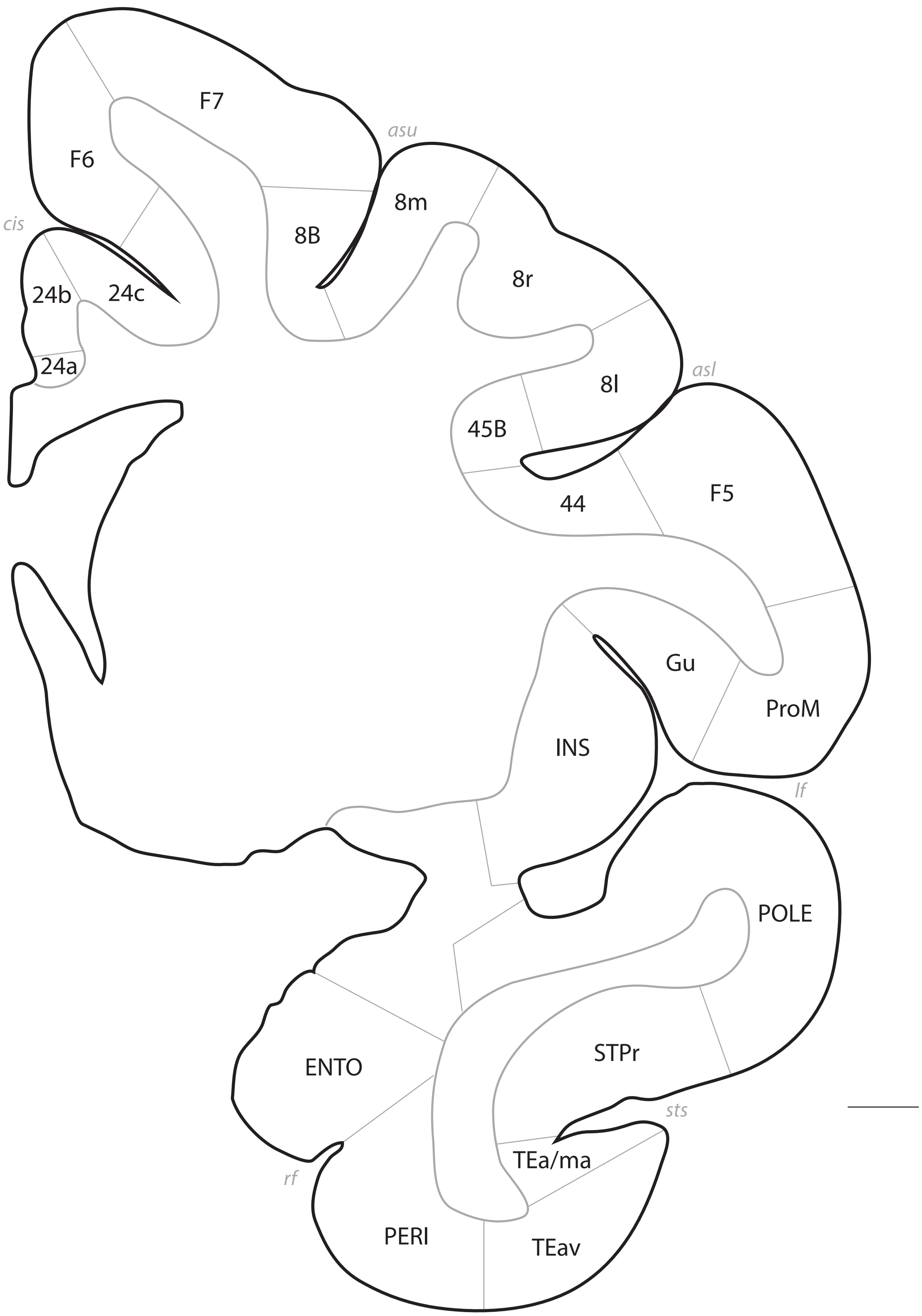
F7
asu
F6
8m
cis
8B
8r
24b
24c
24a
8l
asl
45B
44
F5
Gu
ProM
INS
lf
POLE
ENTO
STPr
sts
TEa/ma
rf
PERI
TEav

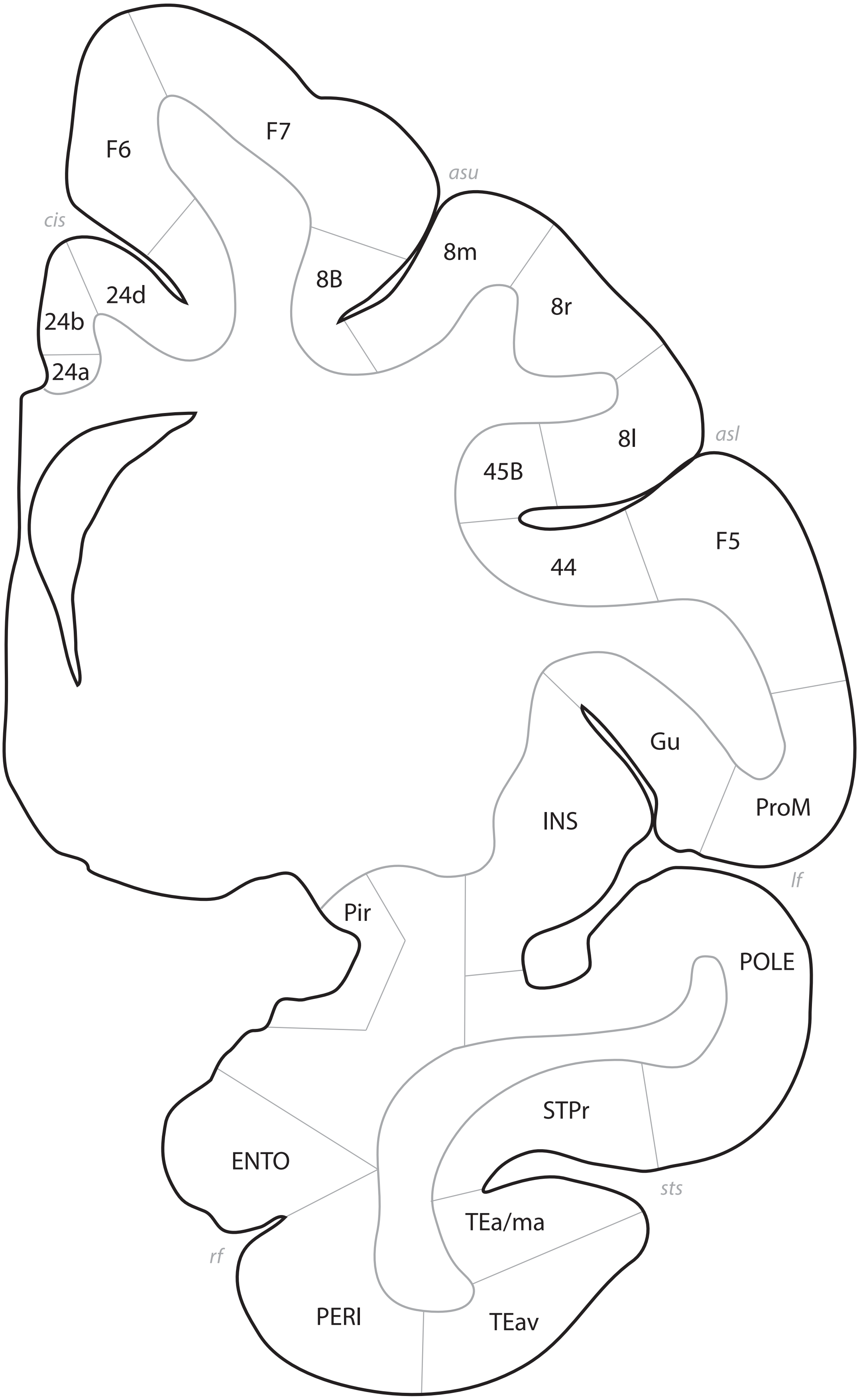
F6
F7
asu
cis
24d
24b
24a
8B
8m
8r
8l
asl
45B
44
F5
Gu
INS
ProM
lf
Pir
POLE
STPr
ENTO
sts
TEa/ma
rf
PERI
TEav

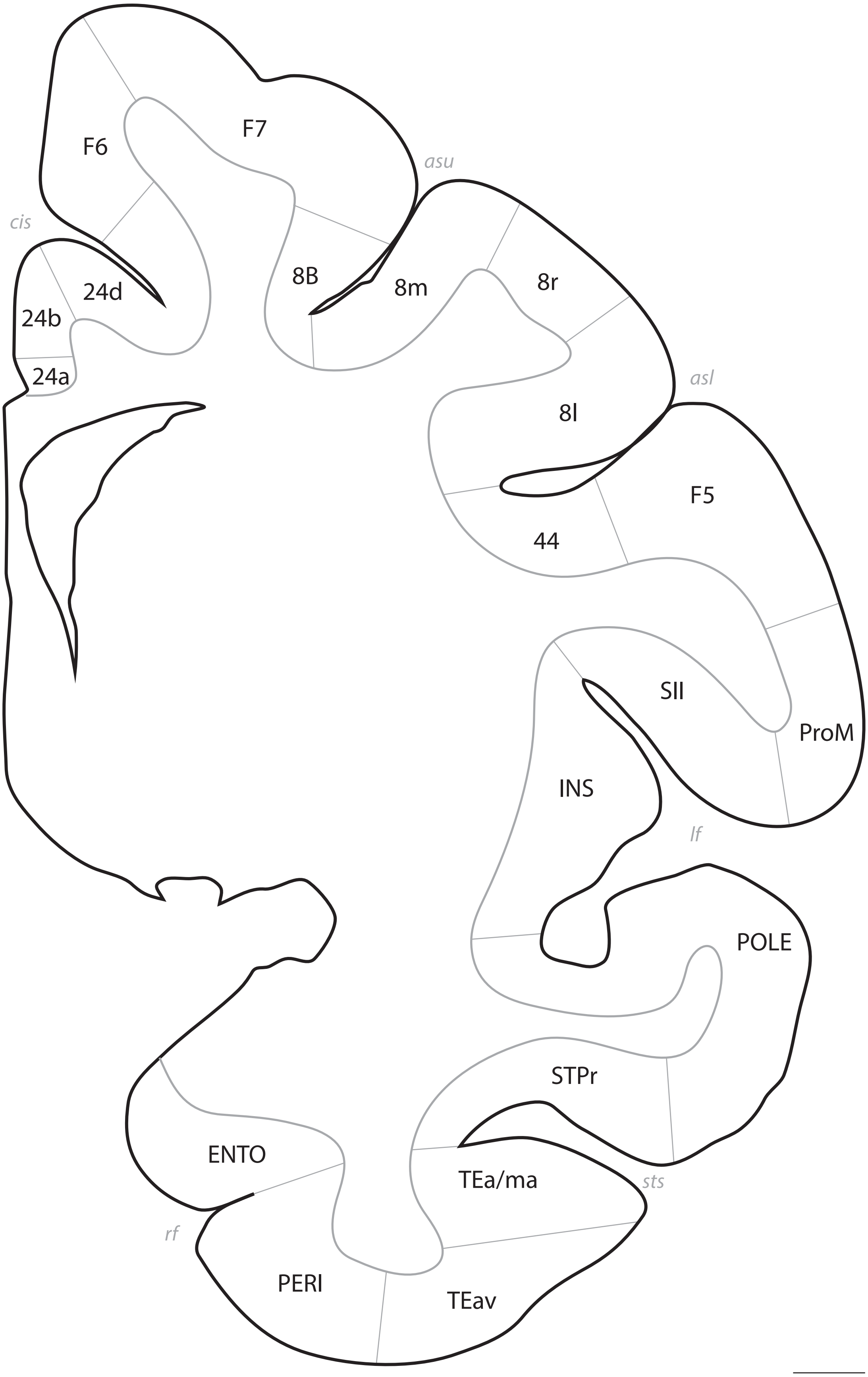
F7
F6
asu
cis
8B
24d
8m
8r
24b
24a
asl
8l
F5
44
SII
ProM
INS
lf
POLE
STPr
TEa/ma
sts
ENTO
rf
PERI
TEav

# p. 042

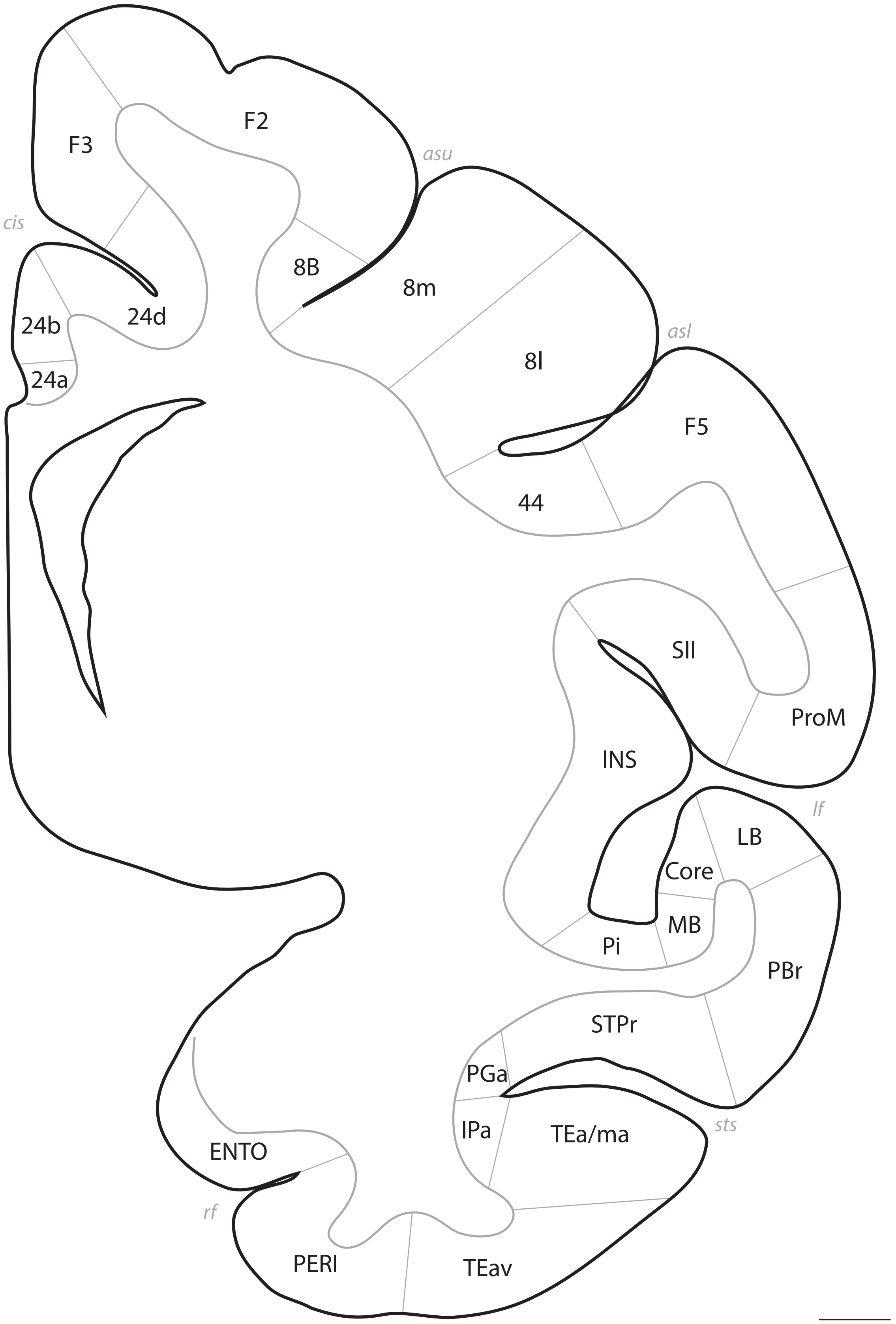
F3
F2
asu
cis
8B
8m
24d
24b
24a
asl
8l
F5
44
SII
ProM
INS
lf
LB
Core
MB
Pi
PBr
STPr
PGa
IPa
TEa/ma
sts
ENTO
rf
PERI
TEav

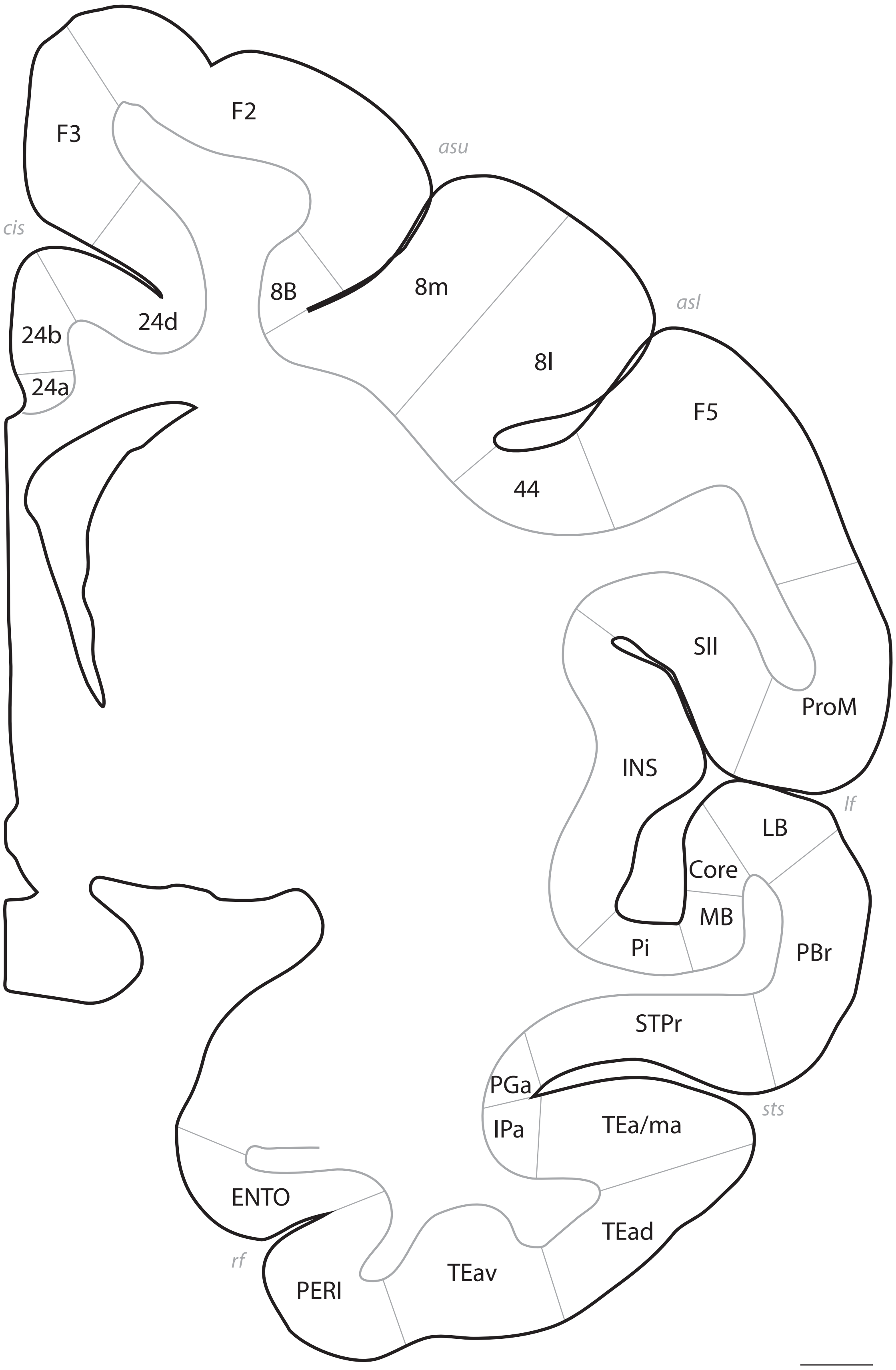
F3
F2
asu
cis
8B
8m
24d
24b
24a
asl
8l
F5
44
SII
ProM
INS
LB
lf
Core
MB
Pi
PBr
STPr
PGa
IPa
TEa/ma
sts
ENTO
TEad
rf
PERI
TEav

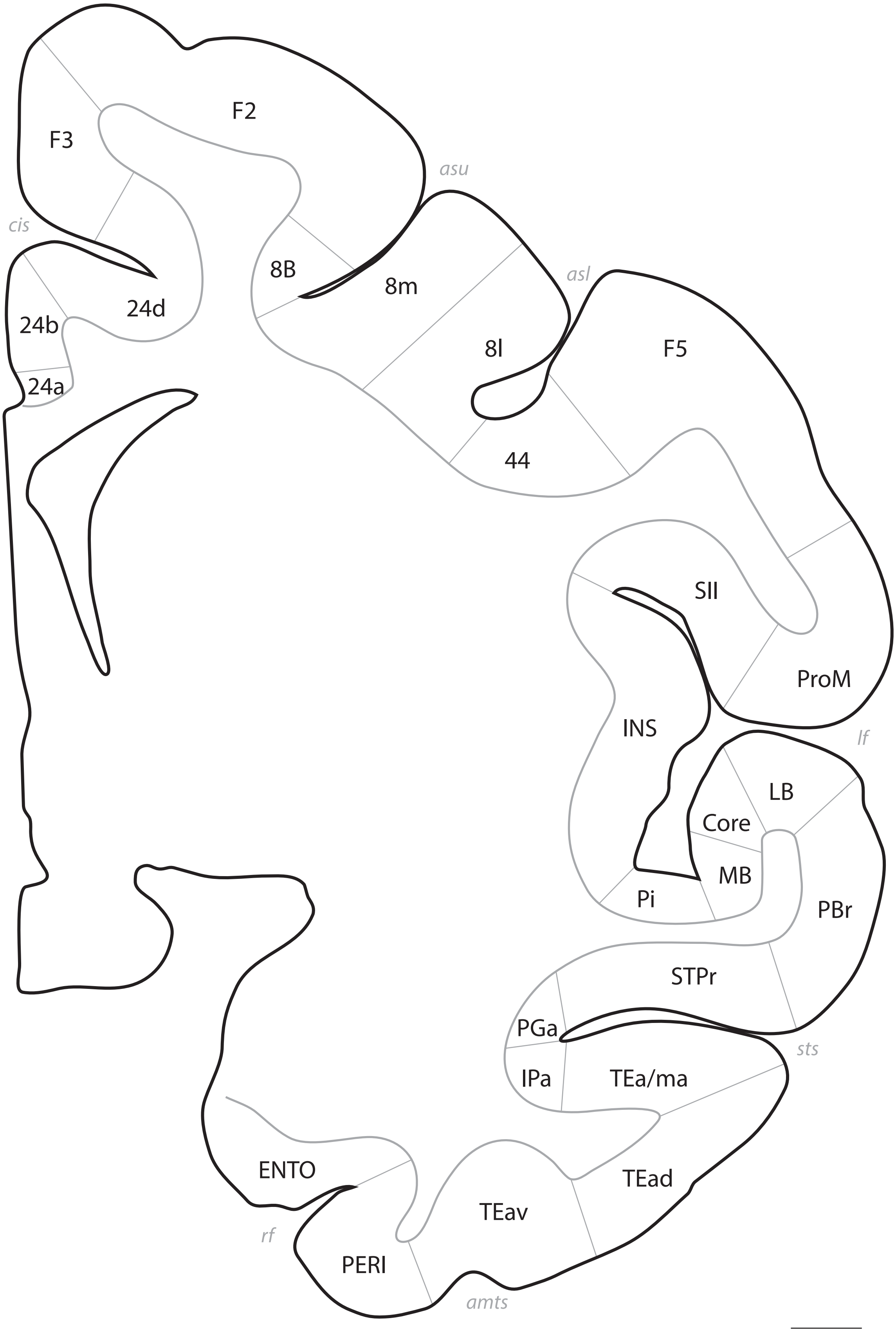
F2
F3
asu
cis
8B
8m
24d
24b
asl
8l
F5
24a
44
SII
ProM
INS
lf
LB
Core
MB
Pi
PBr
STPr
PGa
sts
IPa
TEa/ma
ENTO
TEad
TEav
rf
PERI
amts

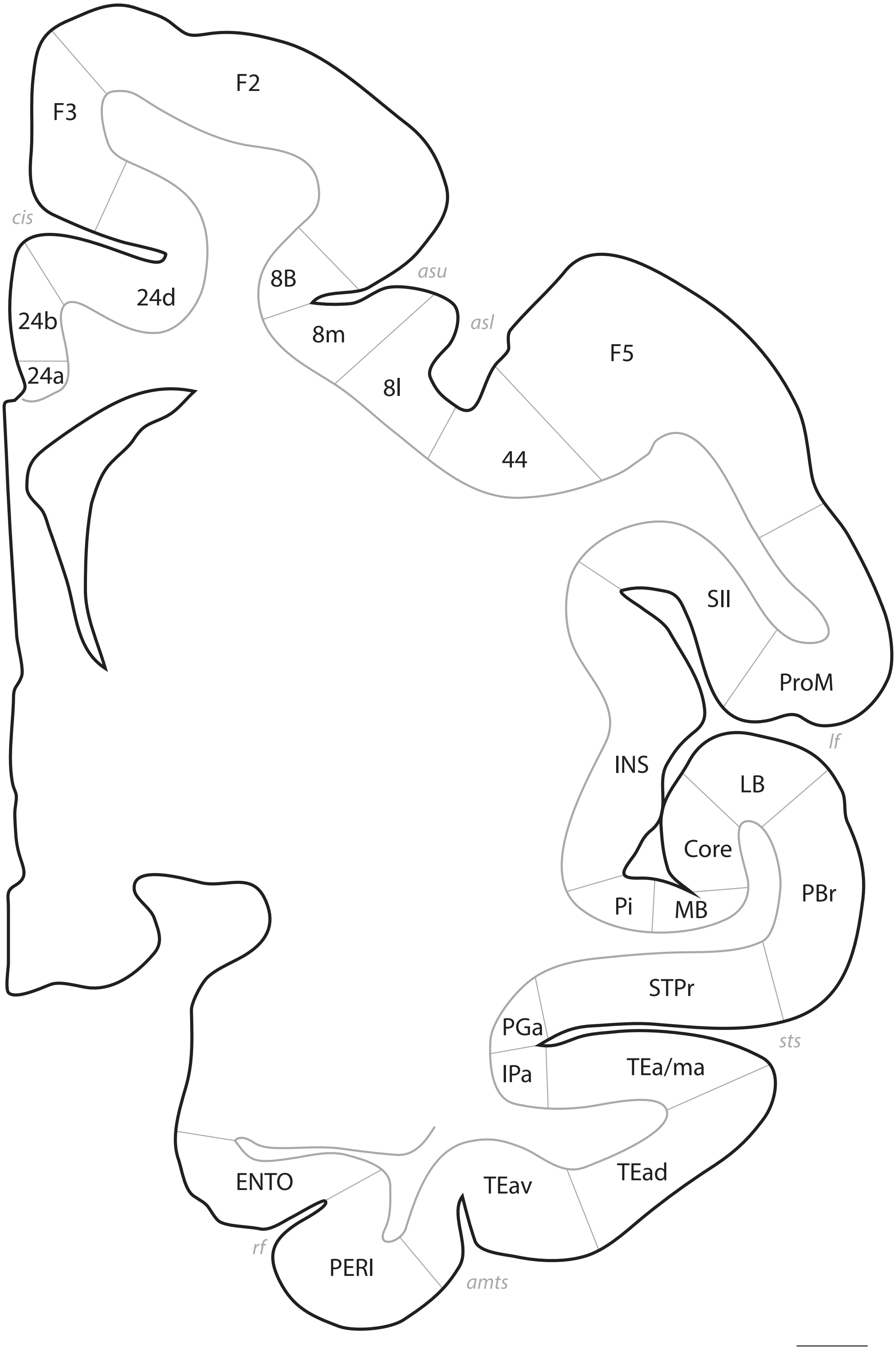
F3
F2
cis
24d
8B
asu
24b
8m
asl
F5
24a
8l
44
SII
ProM
INS
lf
LB
Core
PBr
Pi
MB
STPr
PGa
sts
IPa
TEa/ma
TEad
ENTO
TEav
rf
PERI
amts

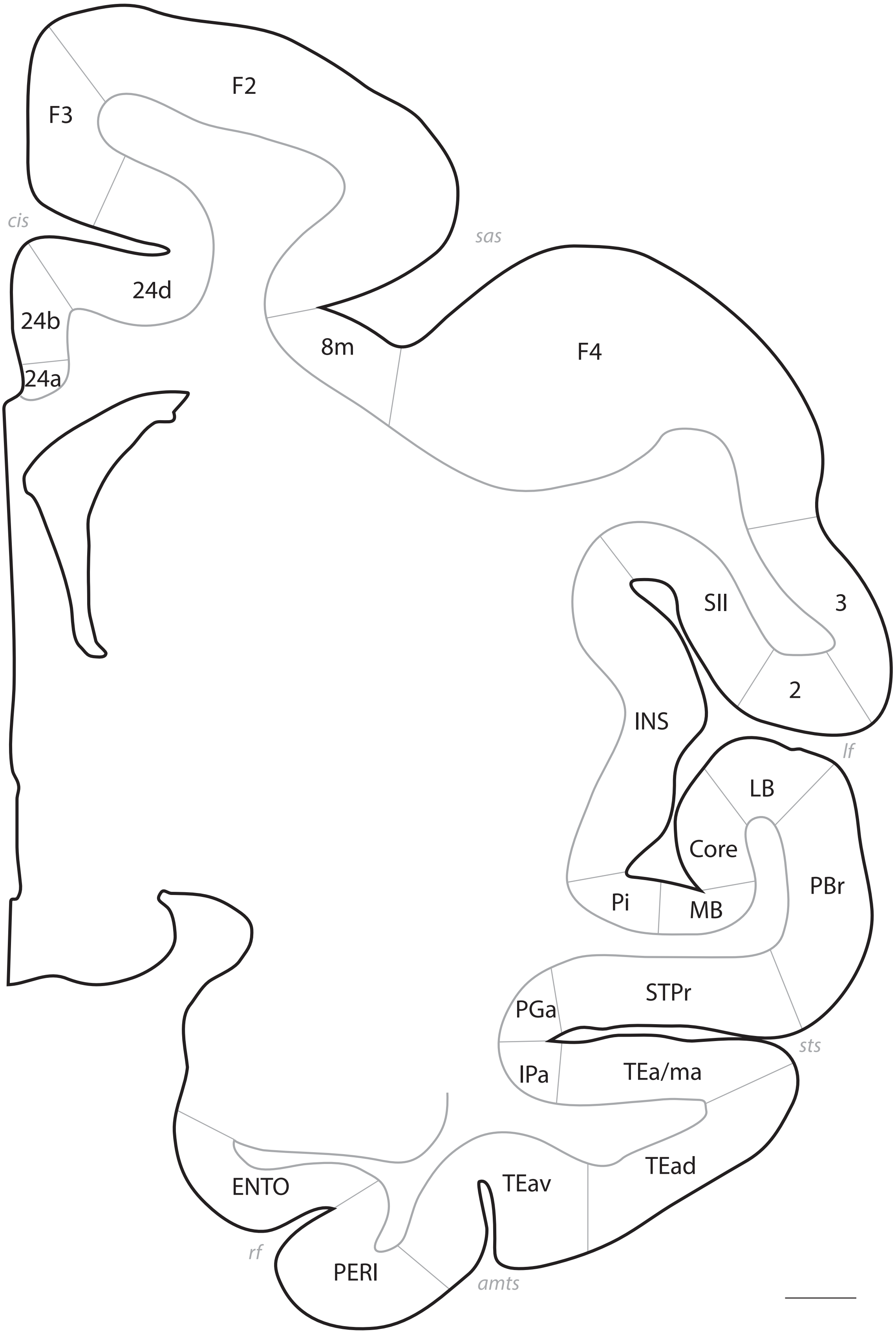
F2
F3
cis
sas
24d
24b
24a
8m
F4
SII
3
INS
2
lf
LB
Core
PBr
Pi
MB
STPr
PGa
IPa
TEa/ma
sts
TEad
ENTO
TEav
rf
PERI
amts

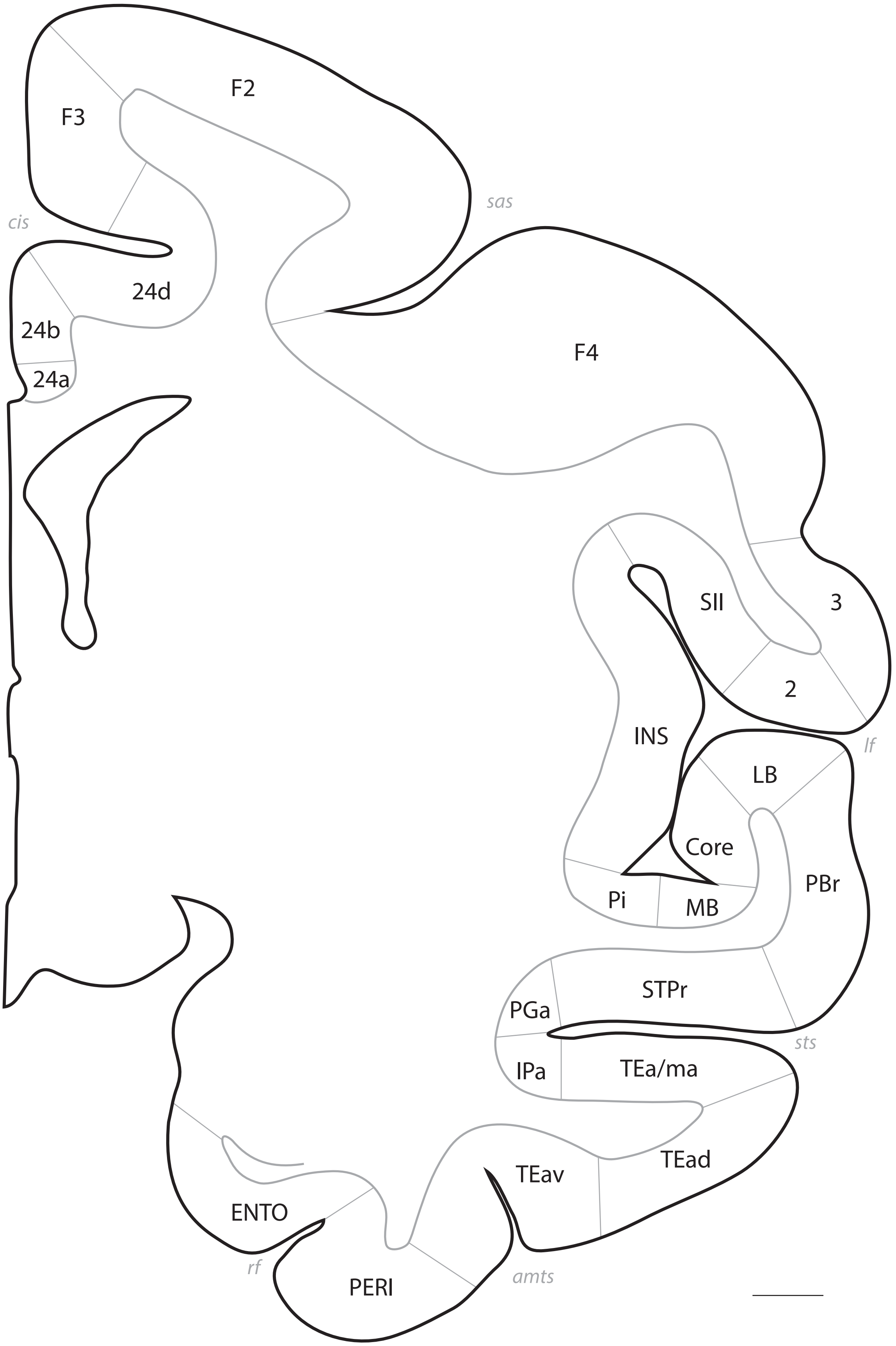
F2
F3
sas
cis
24d
24b
24a
F4
SII
3
2
INS
lf
LB
Core
PBr
Pi
MB
STPr
PGa
sts
IPa
TEa/ma
TEad
TEav
ENTO
rf
PERI
amts

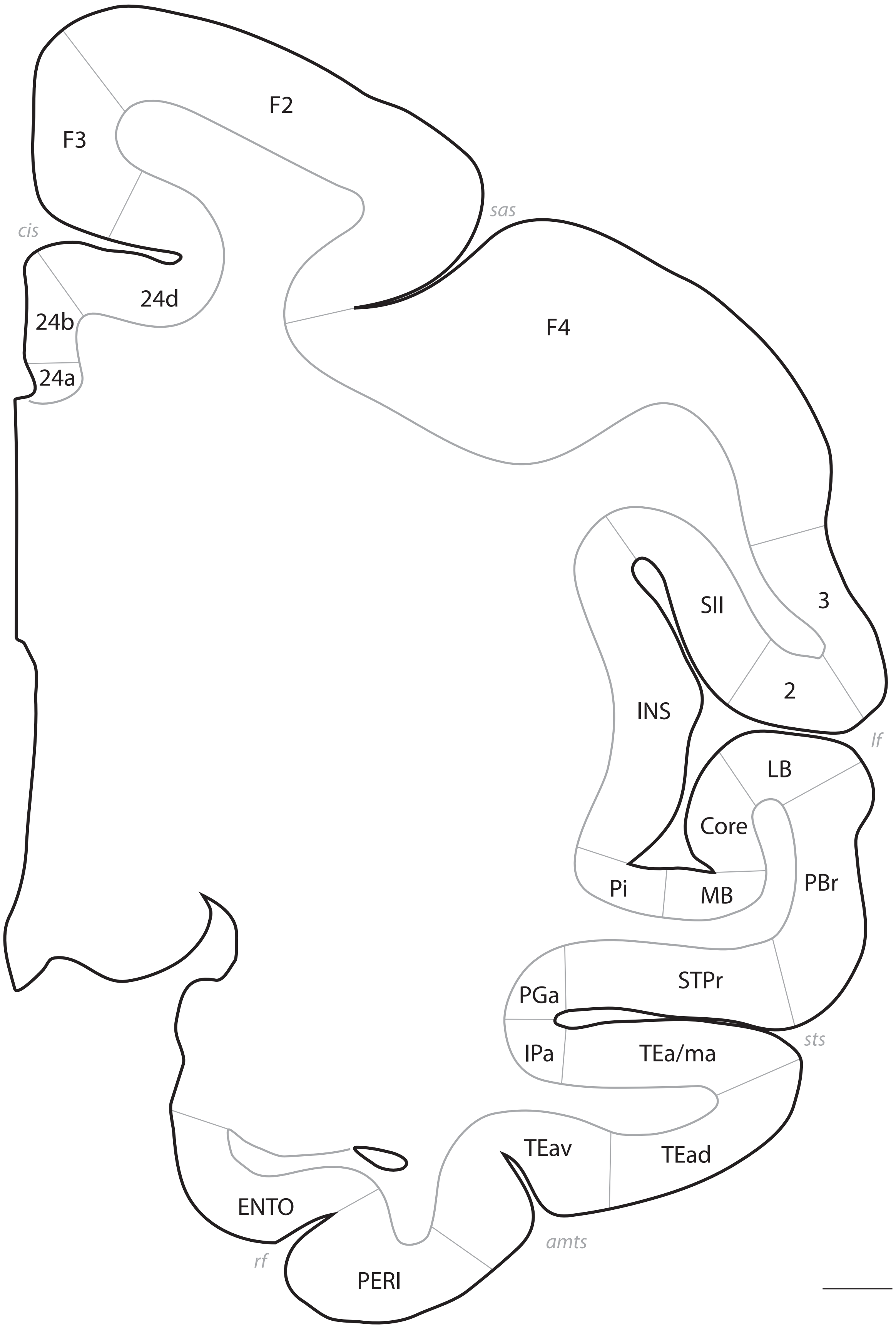
F2
F3
sas
cis
24d
24b
24a
F4
3
SII
2
INS
lf
LB
Core
PBr
Pi
MB
STPr
PGa
sts
IPa
TEa/ma
TEav
TEad
ENTO
amts
rf
PERI

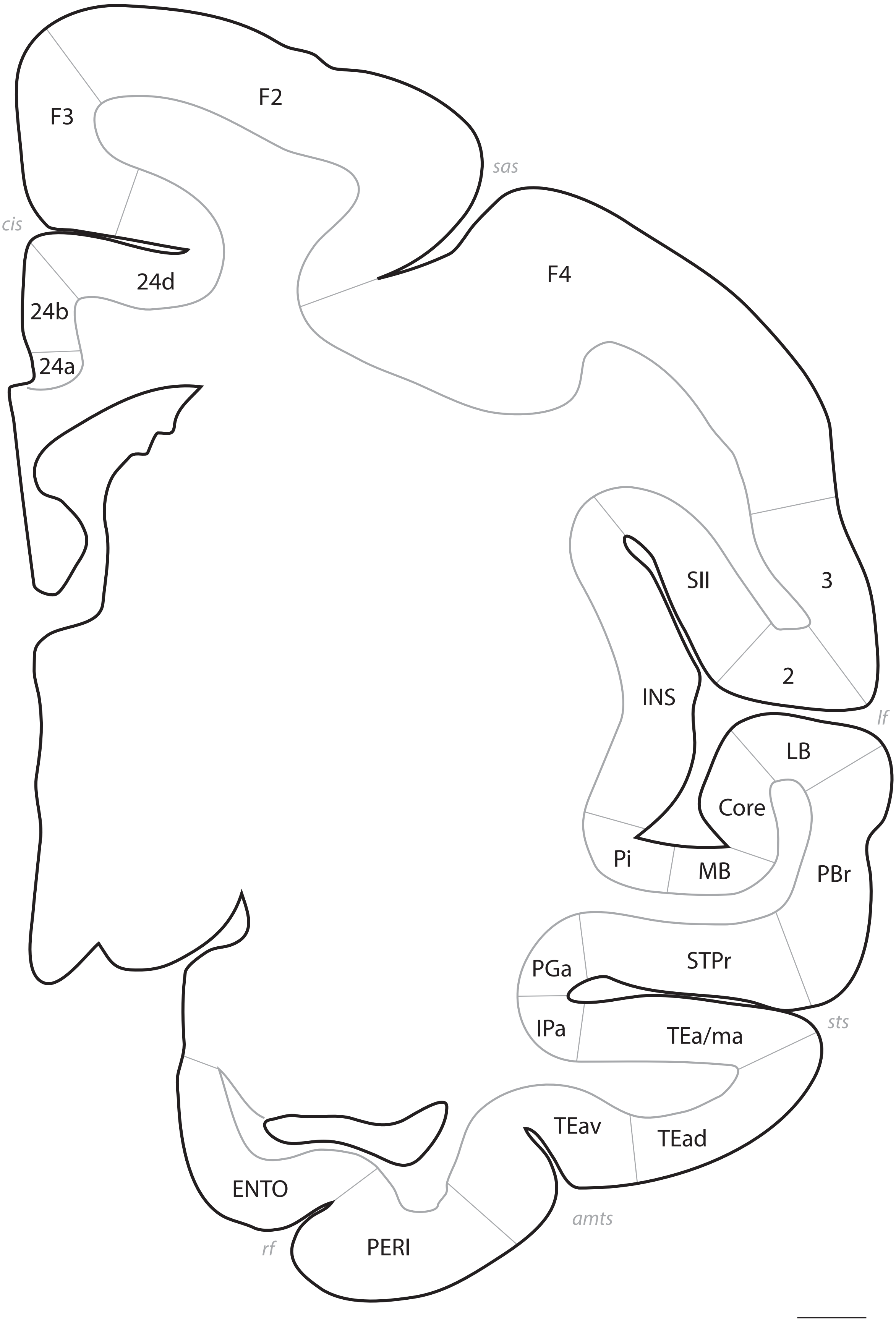
F3
F2
sas
cis
24d
F4
24b
24a
SII
3
2
INS
lf
LB
Core
Pi
MB
PBr
STPr
PGa
IPa
TEa/ma
sts
TEav
TEad
ENTO
amts
rf
PERI

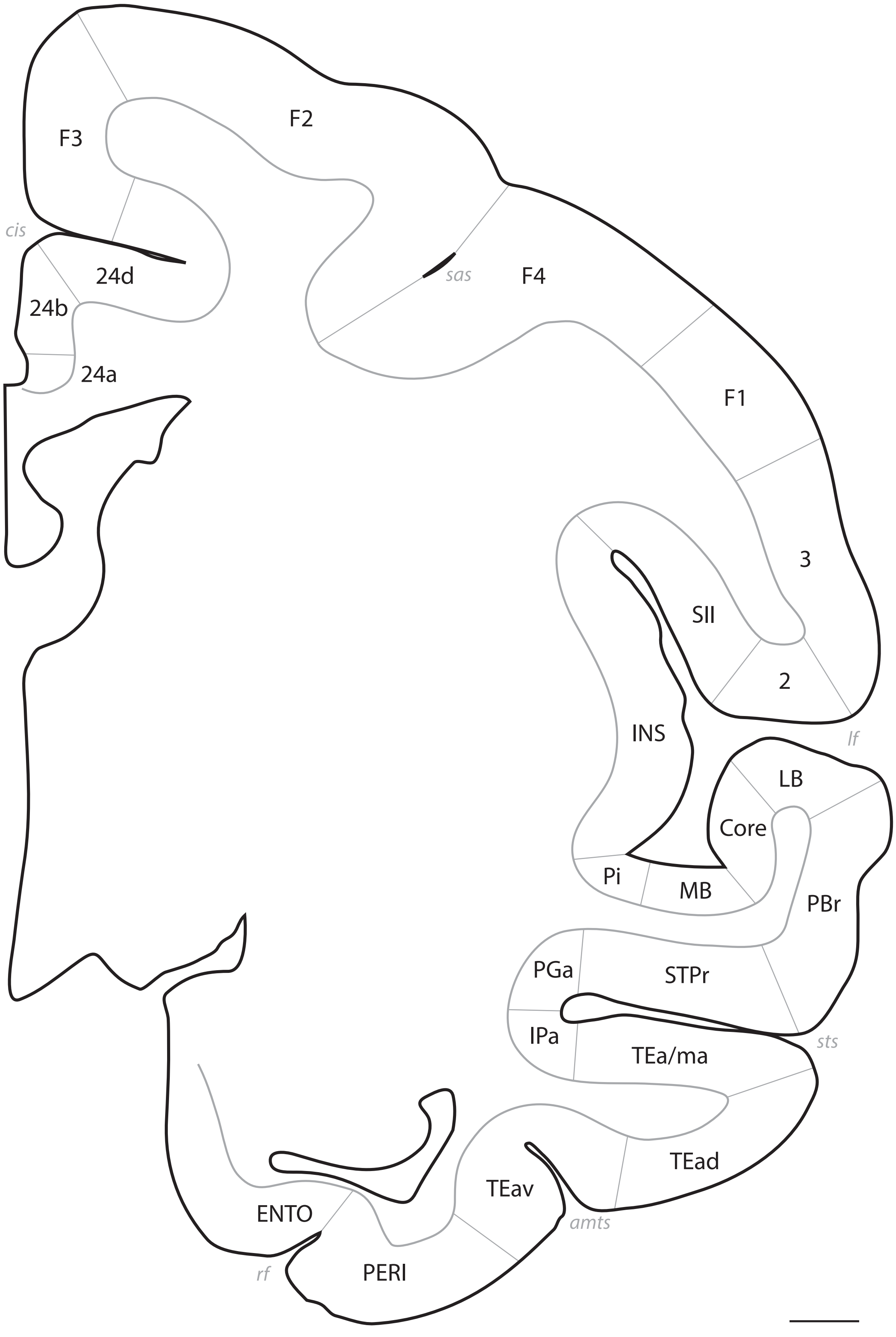
F3
F2
cis
24d
sas
F4
24b
24a
F1
3
SII
2
INS
lf
LB
Core
Pi
MB
PBr
PGa
STPr
IPa
TEa/ma
sts
TEad
TEav
ENTO
amts
rf
PERI

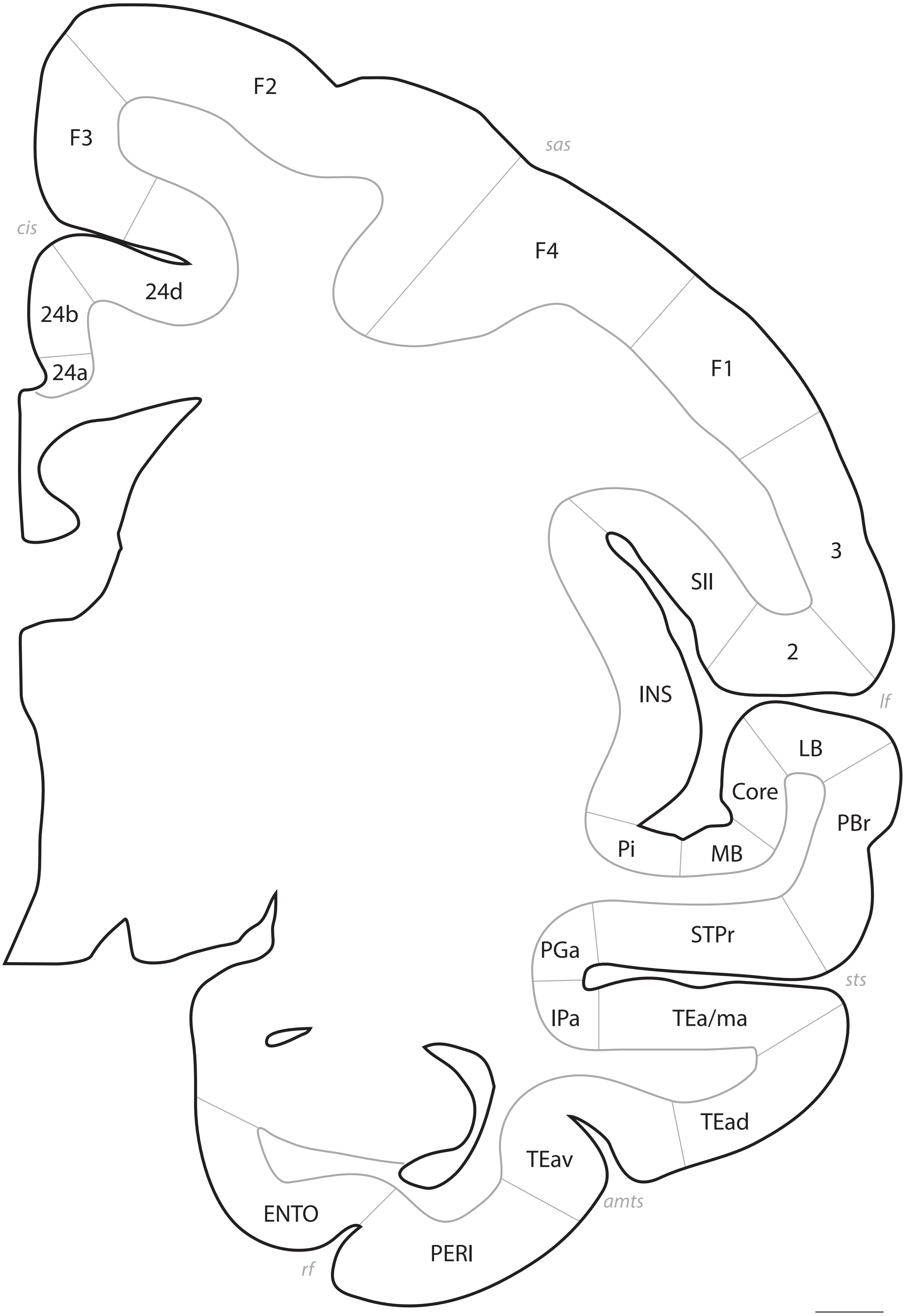
F2
F3
sas
F4
cis
24d
24b
F1
24a
3
SII
2
INS
lf
LB
Core
PBr
Pi
MB
STPr
PGa
sts
IPa
TEa/ma
TEad
TEav
ENTO
amts
PERI
rf

# p. 052

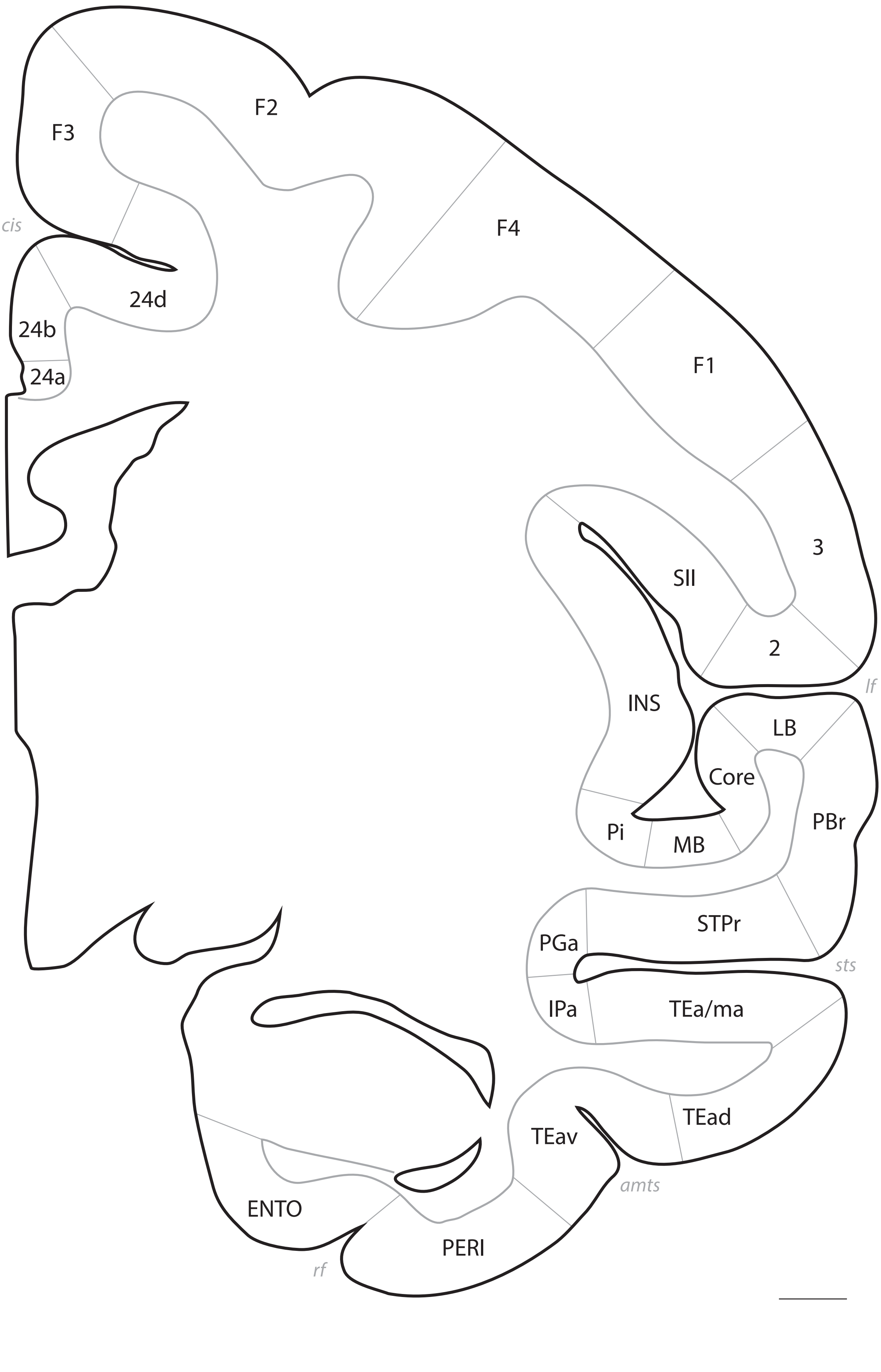
F2
F3
F4
cis
24d
24b
24a
F1
3
SII
2
INS
lf
LB
Core
PBr
Pi
MB
STPr
PGa
sts
IPa
TEa/ma
TEad
TEav
amts
ENTO
PERI
rf

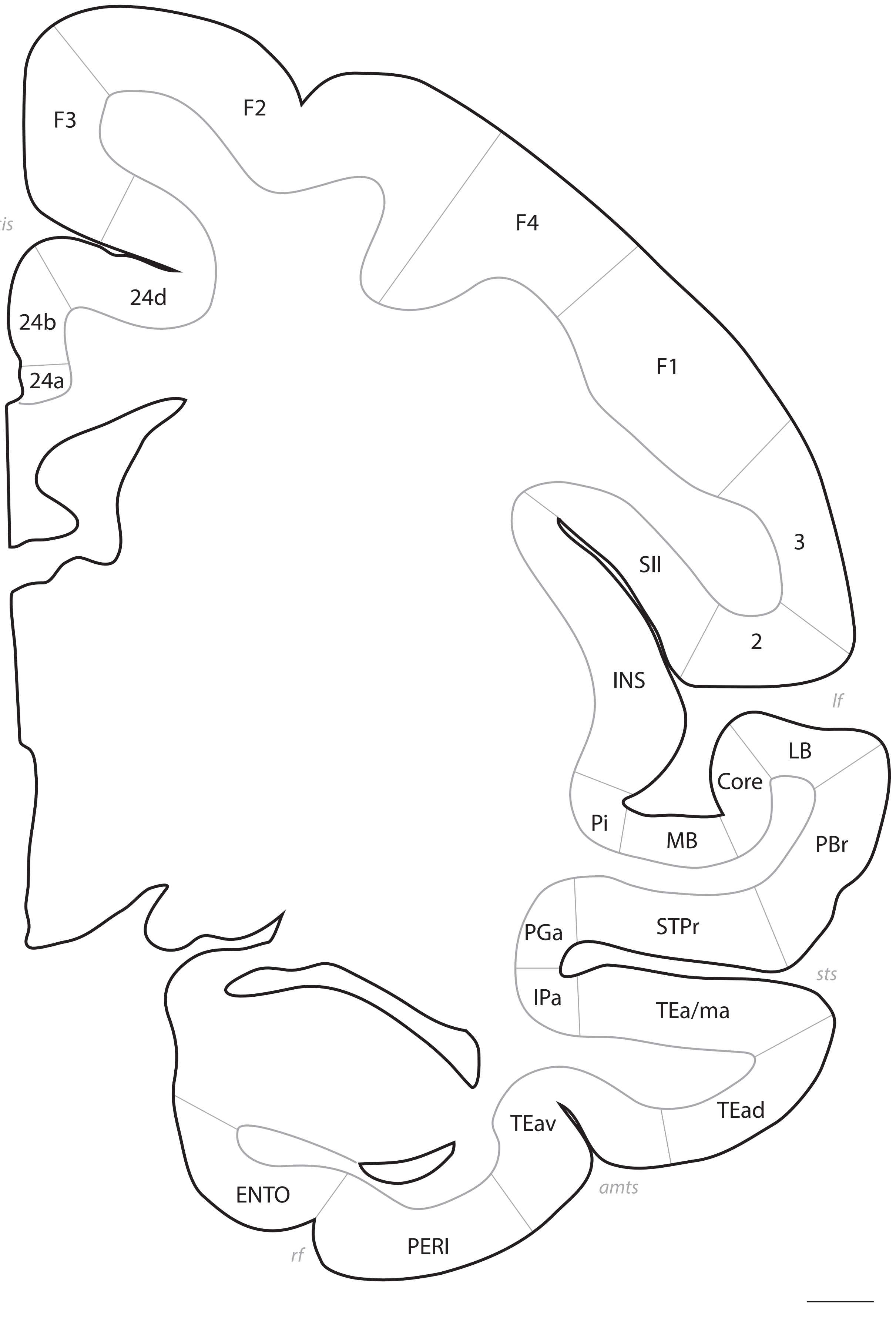
F3
F2
F4
cis
24d
24b
24a
F1
3
SII
2
INS
lf
LB
Core
Pi
MB
PBr
PGa
STPr
sts
IPa
TEa/ma
TEad
TEav
ENTO
amts
PERI
rf

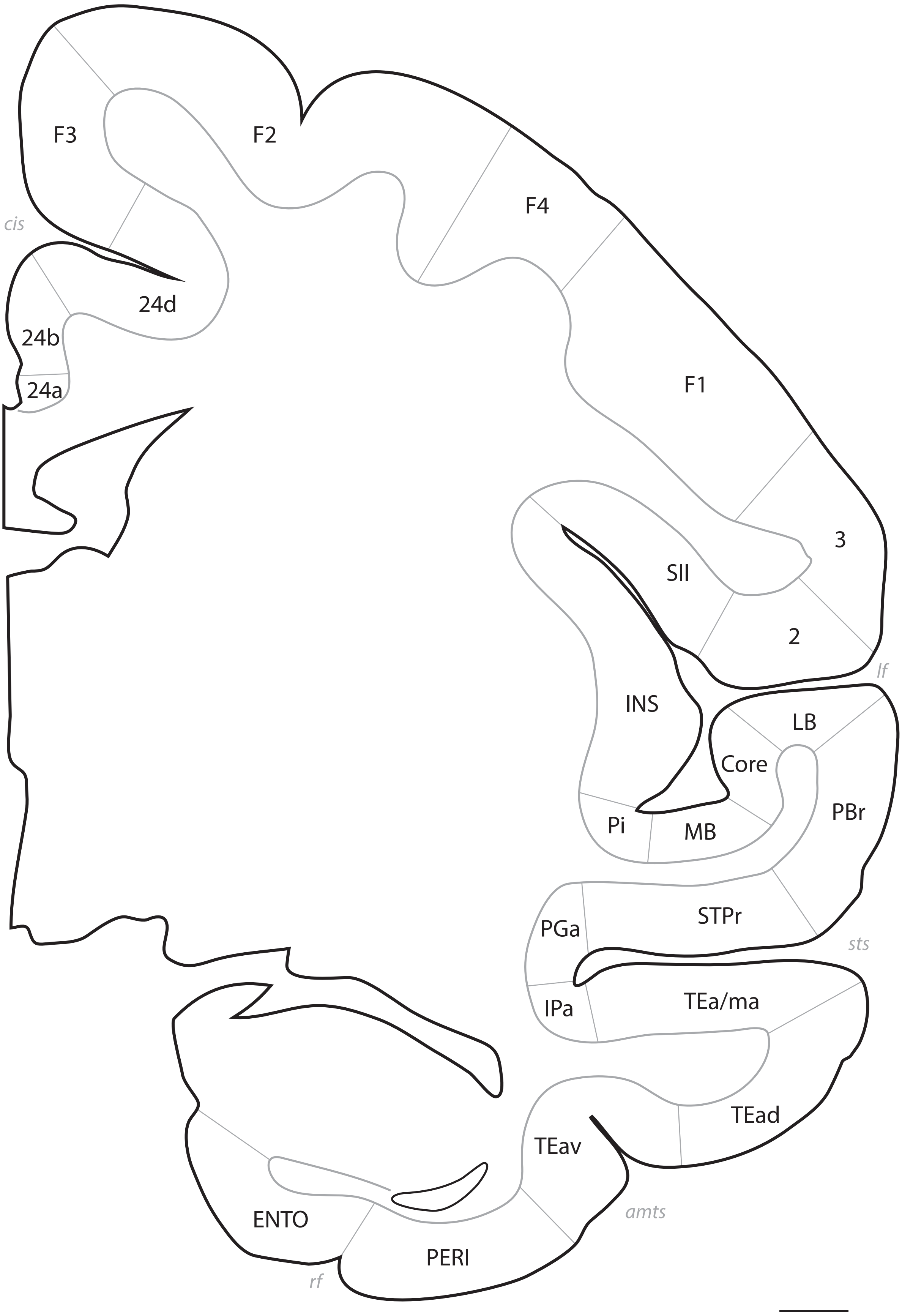
F3
F2
F4
cis
24d
24b
24a
F1
3
SII
2
lf
INS
LB
Core
PBr
Pi
MB
STPr
PGa
sts
TEa/ma
IPa
TEad
TEav
amts
ENTO
PERI
rf

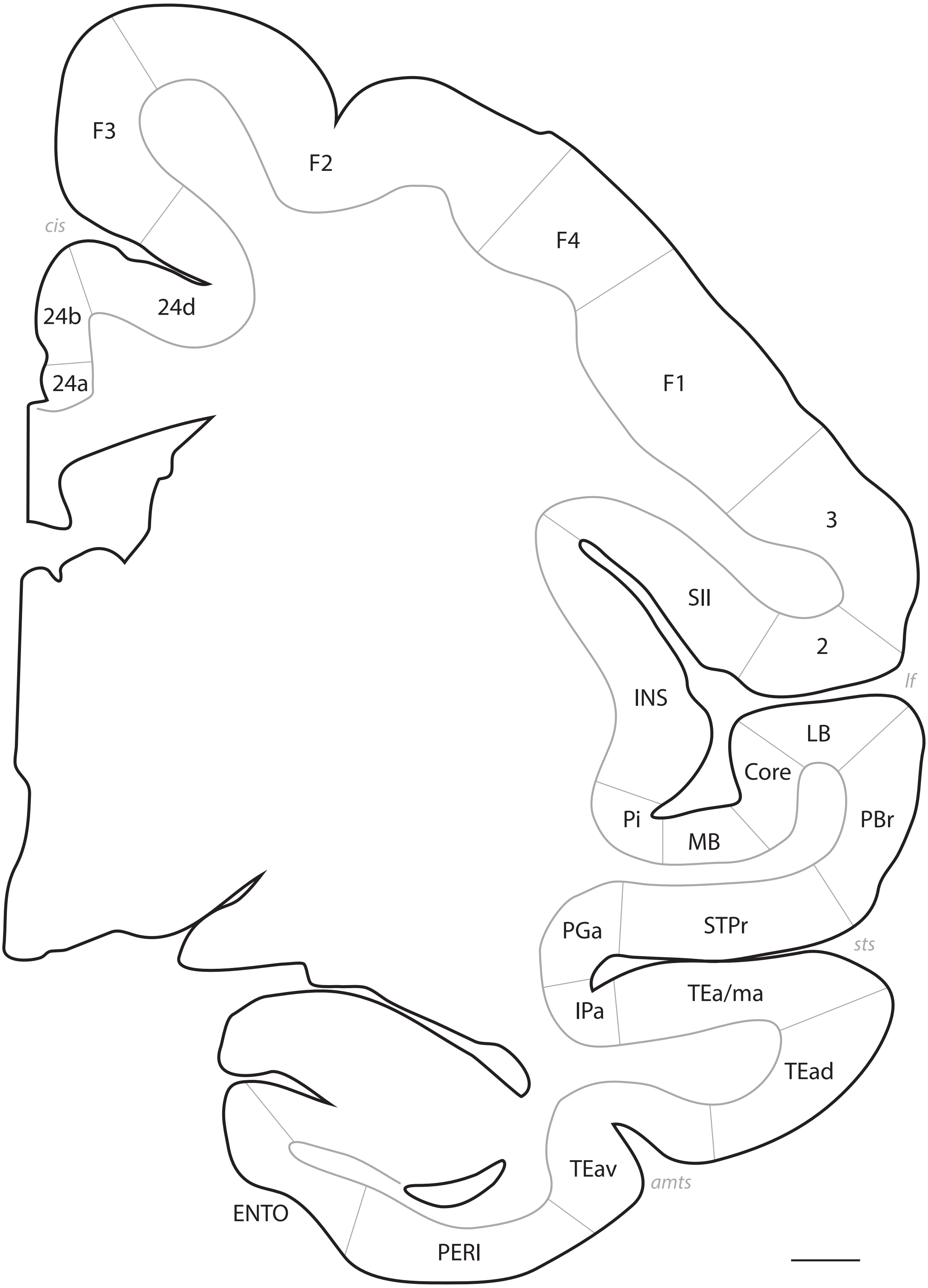
F3
F2
cis
F4
24b
24d
24a
F1
3
SII
2
lf
INS
LB
Core
PBr
Pi
MB
PGa
STPr
sts
TEa/ma
IPa
TEad
TEav
amts
ENTO
PERI

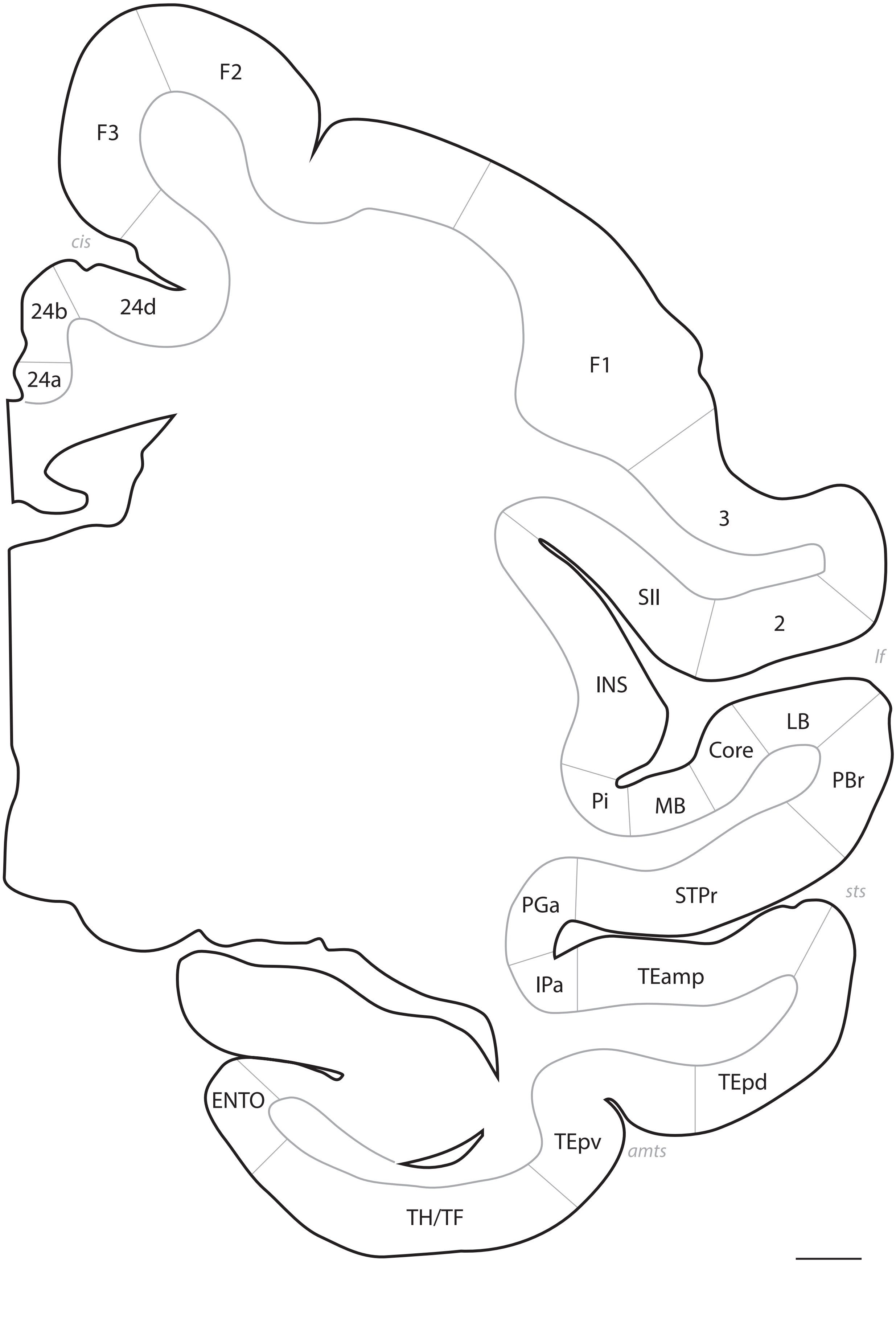
F2
F3
cis
24b
24d
24a
F1
3
SII
2
lf
INS
LB
Core
PBr
Pi
MB
STPr
sts
PGa
TEamp
IPa
TEpd
ENTO
TEpv
amts
TH/TF

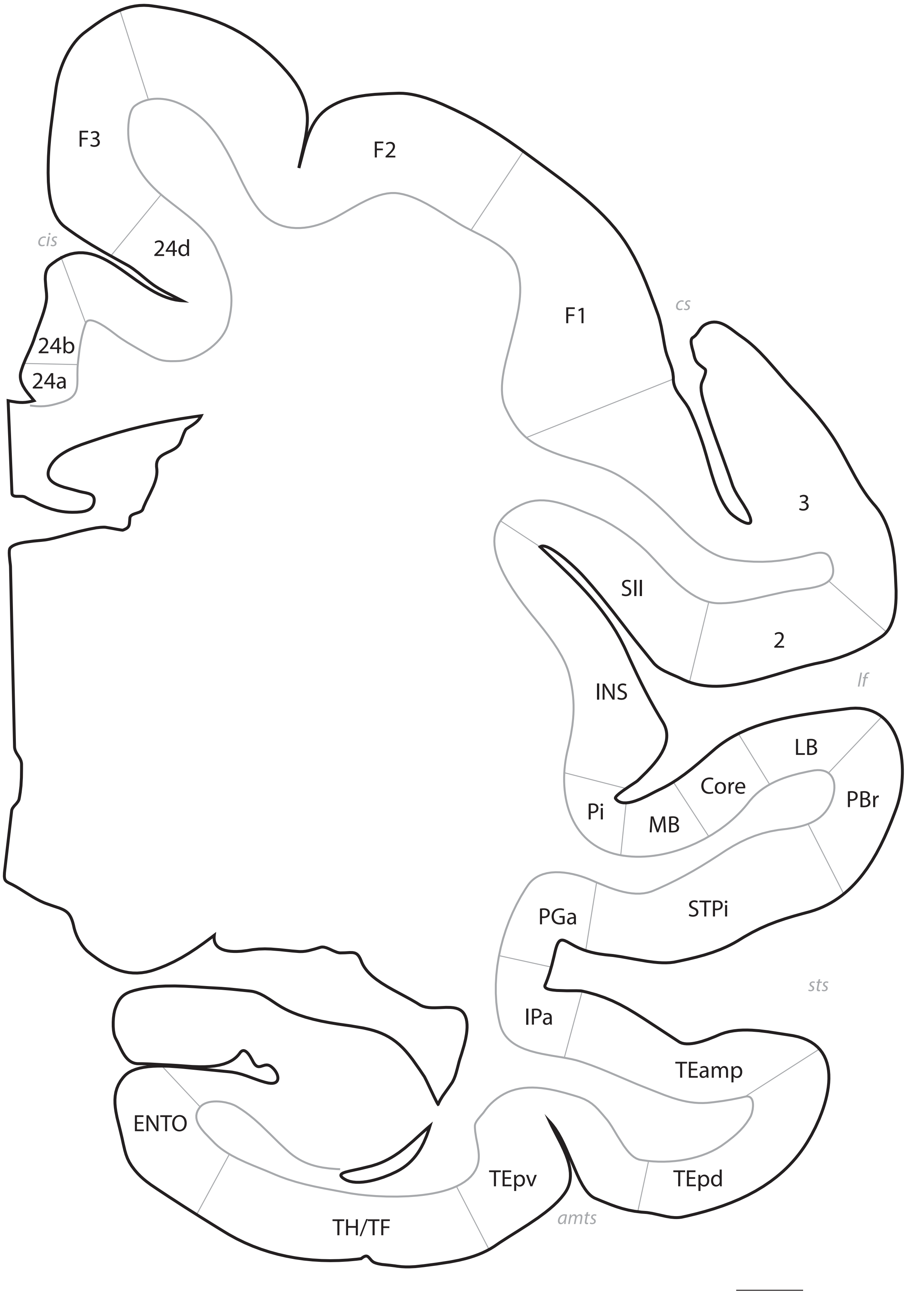
F3
F2
cis
24d
F1
cs
24b
24a
3
SII
2
lf
INS
LB
Core
PBr
Pi
MB
PGa
STPi
sts
IPa
TEamp
ENTO
TEpv
TEpd
amts
TH/TF

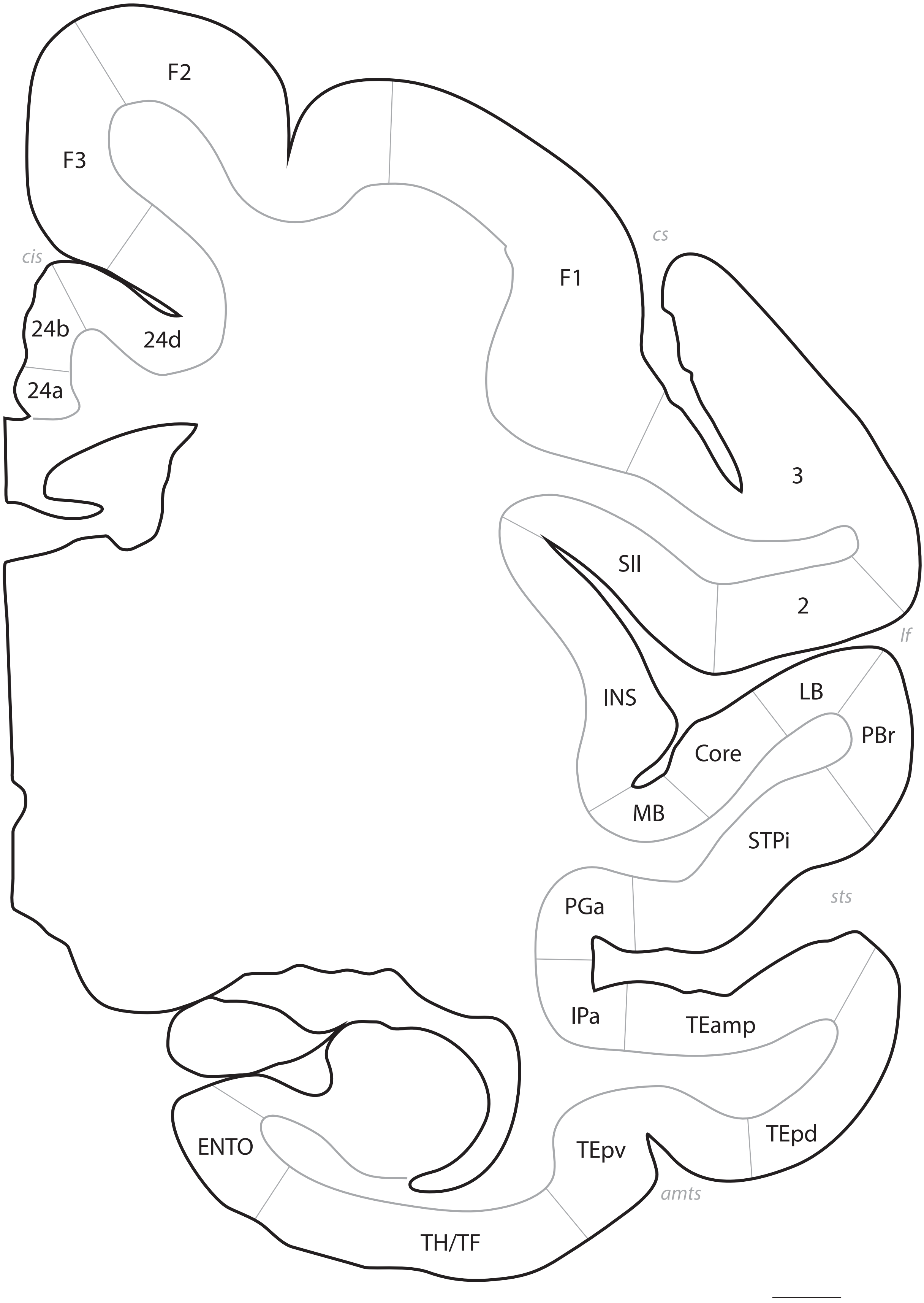
F2
F3
cis
24b
24d
24a
cs
F1
3
SII
2
lf
INS
LB
PBr
Core
MB
STPi
sts
PGa
IPa
TEamp
TEpd
ENTO
TEpv
amts
TH/TF

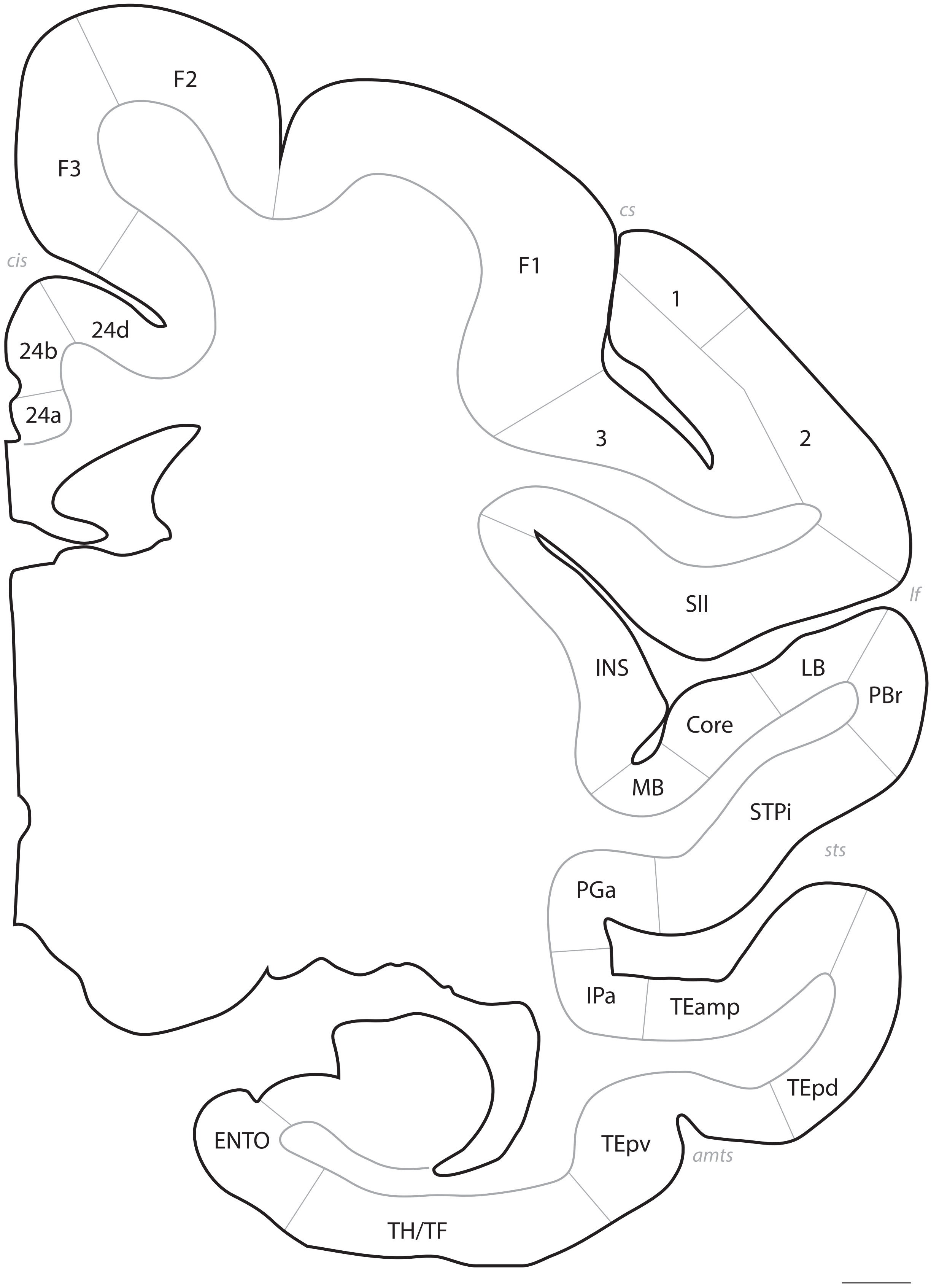
F2
F3
cis
24d
24b
24a
cs
F1
1
3
2
SII
lf
INS
LB
PBr
Core
MB
STPi
sts
PGa
IPa
TEamp
TEpd
ENTO
TEpv
amts
TH/TF

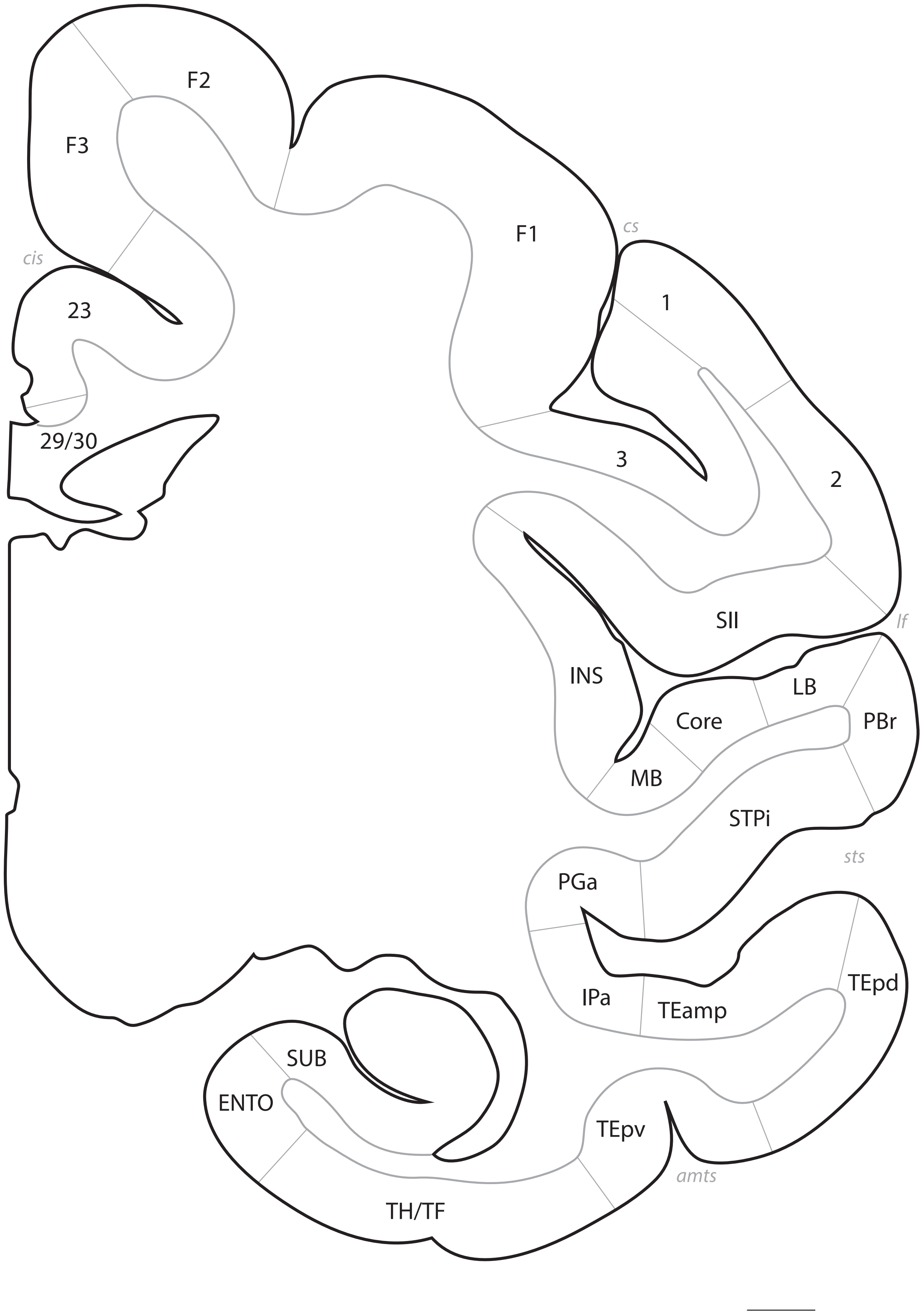
F2
F3
F1
cs
cis
23
1
29/30
3
2
SII
lf
INS
LB
Core
PBr
MB
STPi
sts
PGa
TEpd
IPa
TEamp
SUB
ENTO
TEpv
amts
TH/TF

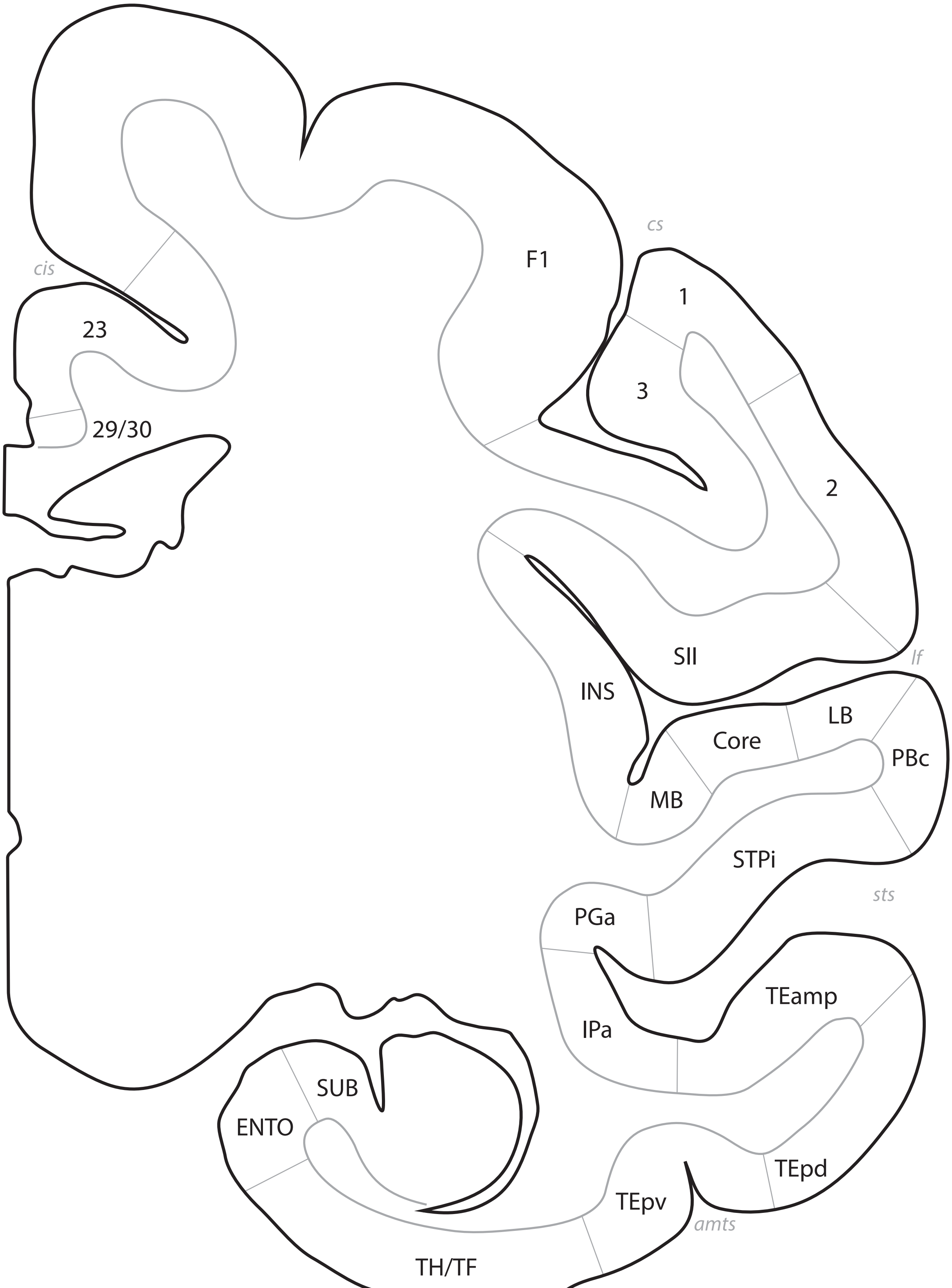
cs
F1
cis
23
29/30
1
3
2
SII
lf
INS
LB
Core
PBc
MB
STPi
sts
PGa
TEamp
IPa
SUB
ENTO
TEpd
TEpv
amts
TH/TF

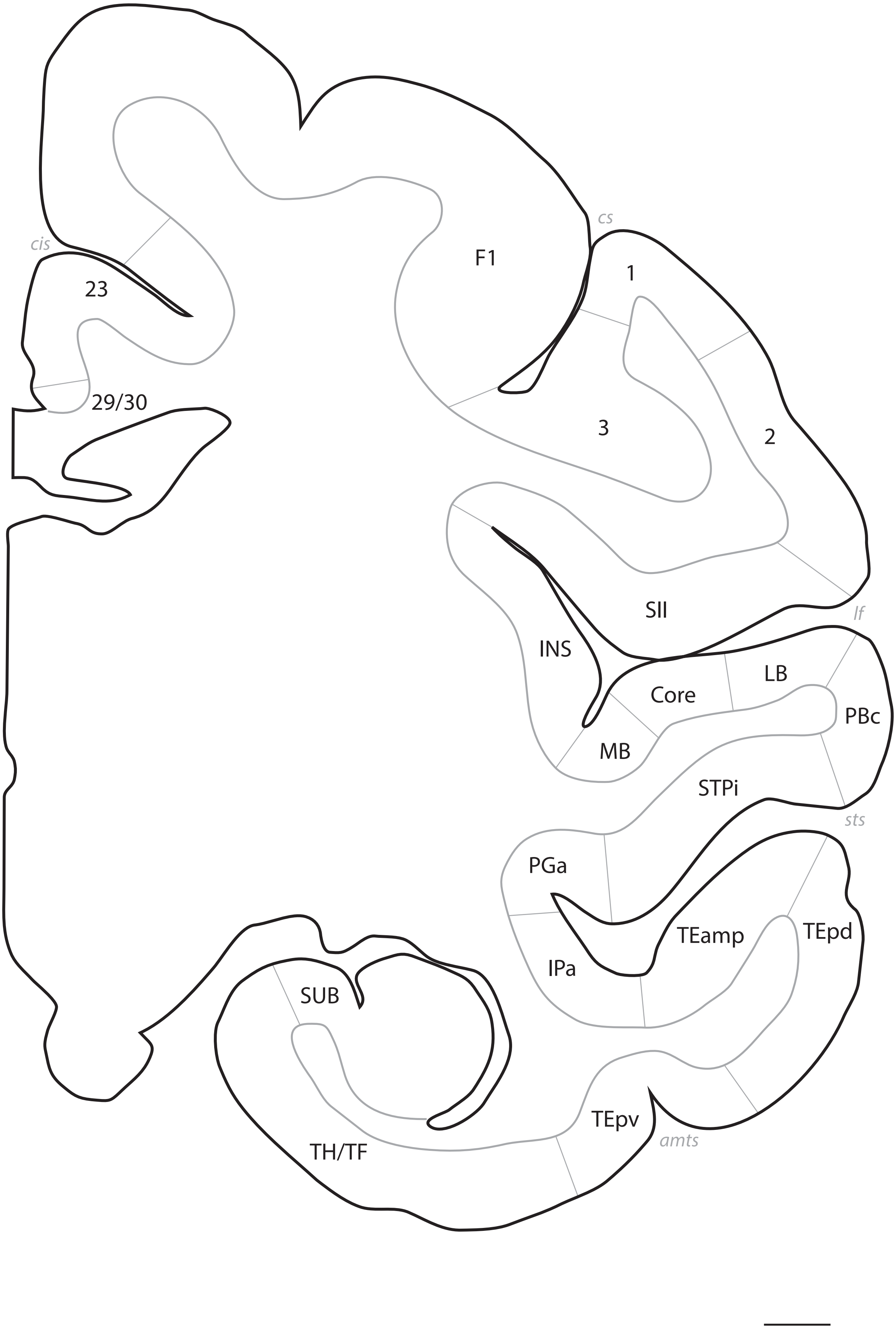
cs
cis
F1
1
23
3
2
29/30
SII
lf
INS
LB
Core
PBc
MB
STPi
sts
PGa
TEamp
TEpd
IPa
SUB
TEpv
amts
TH/TF

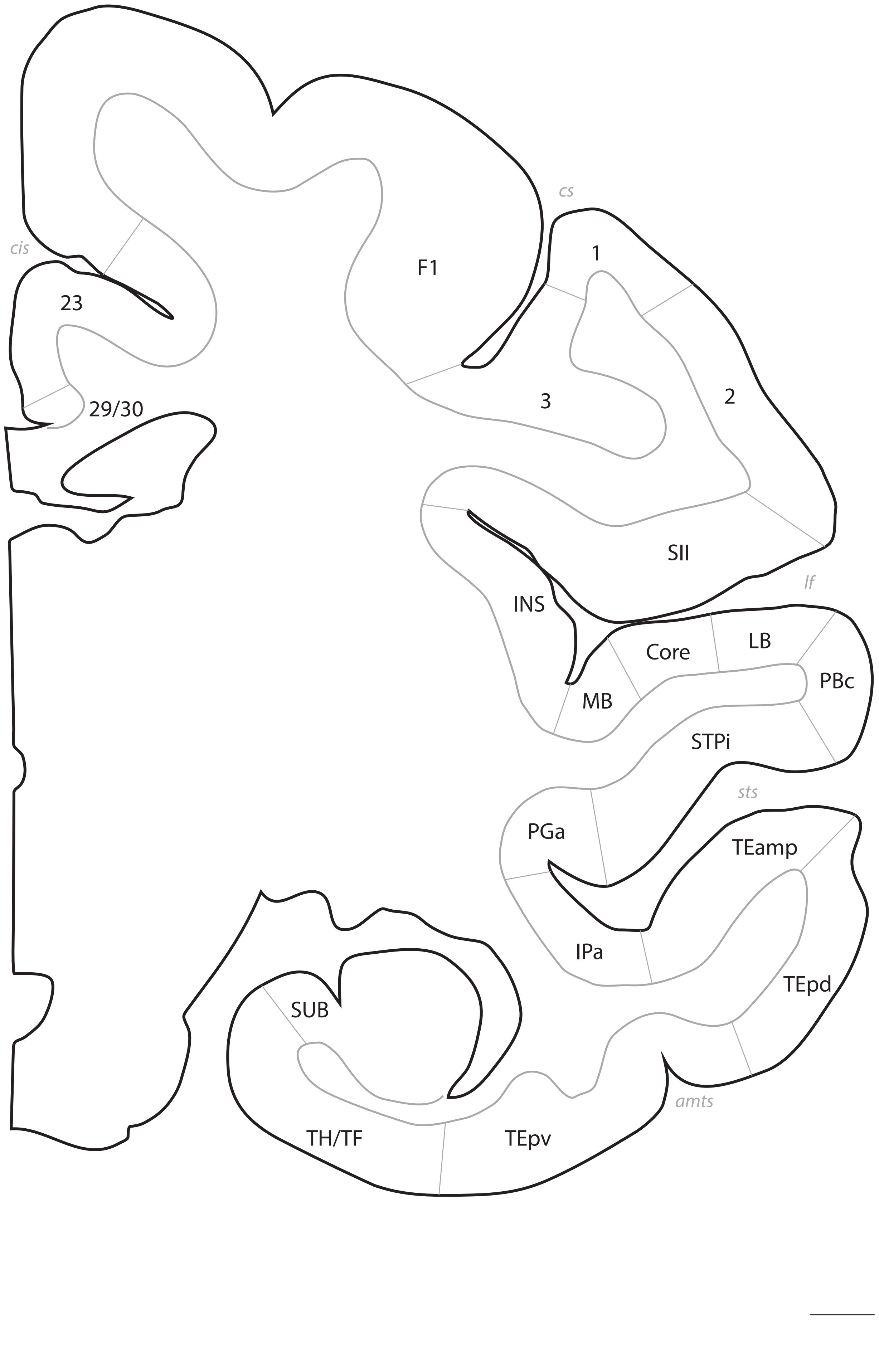
cs
cis
1
F1
23
3
2
29/30
SII
lf
INS
LB
Core
PBc
MB
STPi
sts
PGa
TEamp
IPa
TEpd
SUB
amts
TH/TF
TEpv

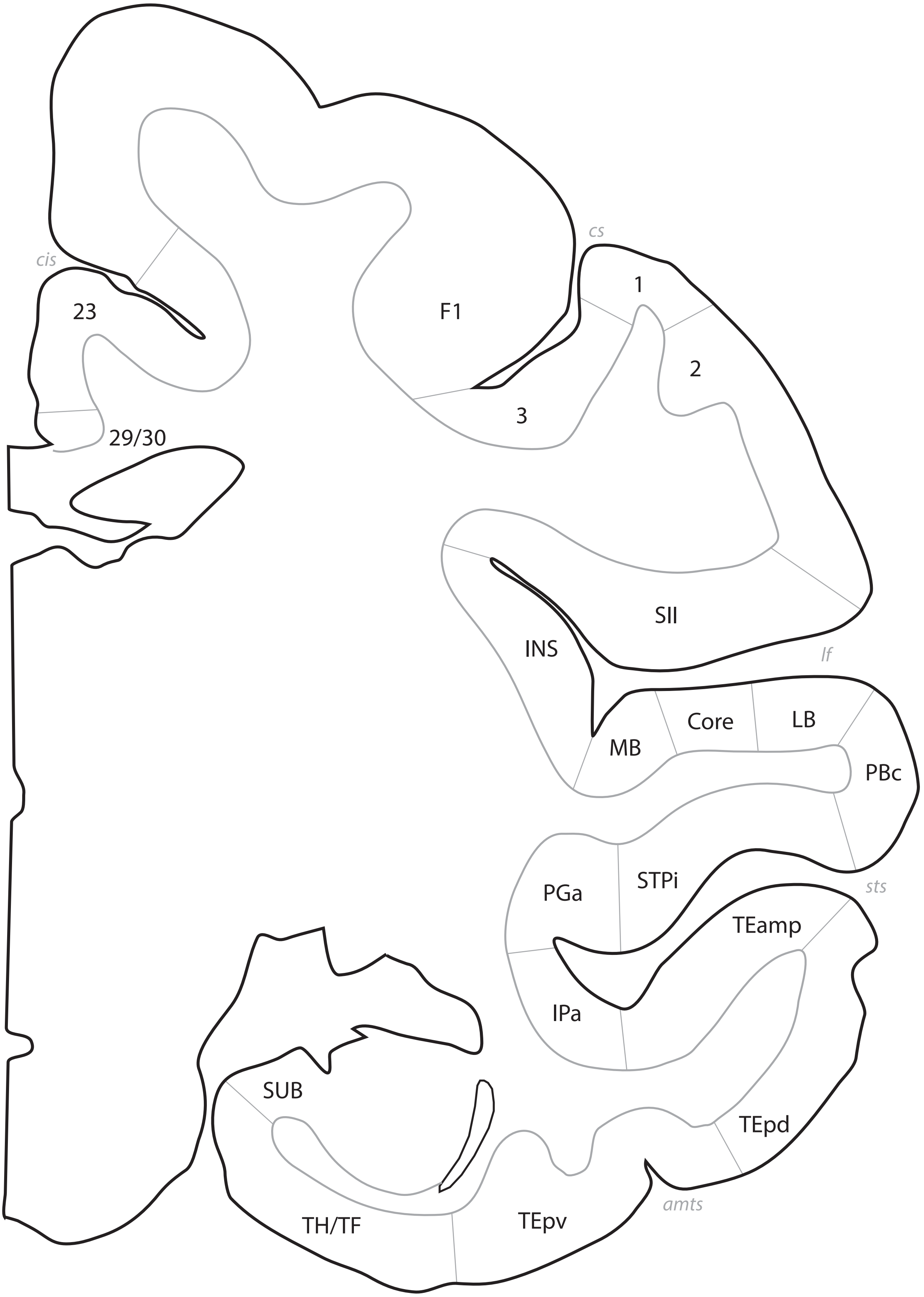
cs
cis
1
23
F1
2
3
29/30
SII
INS
lf
Core
LB
MB
PBc
STPi
PGa
sts
TEamp
IPa
SUB
TEpd
TH/TF
TEpv
amts

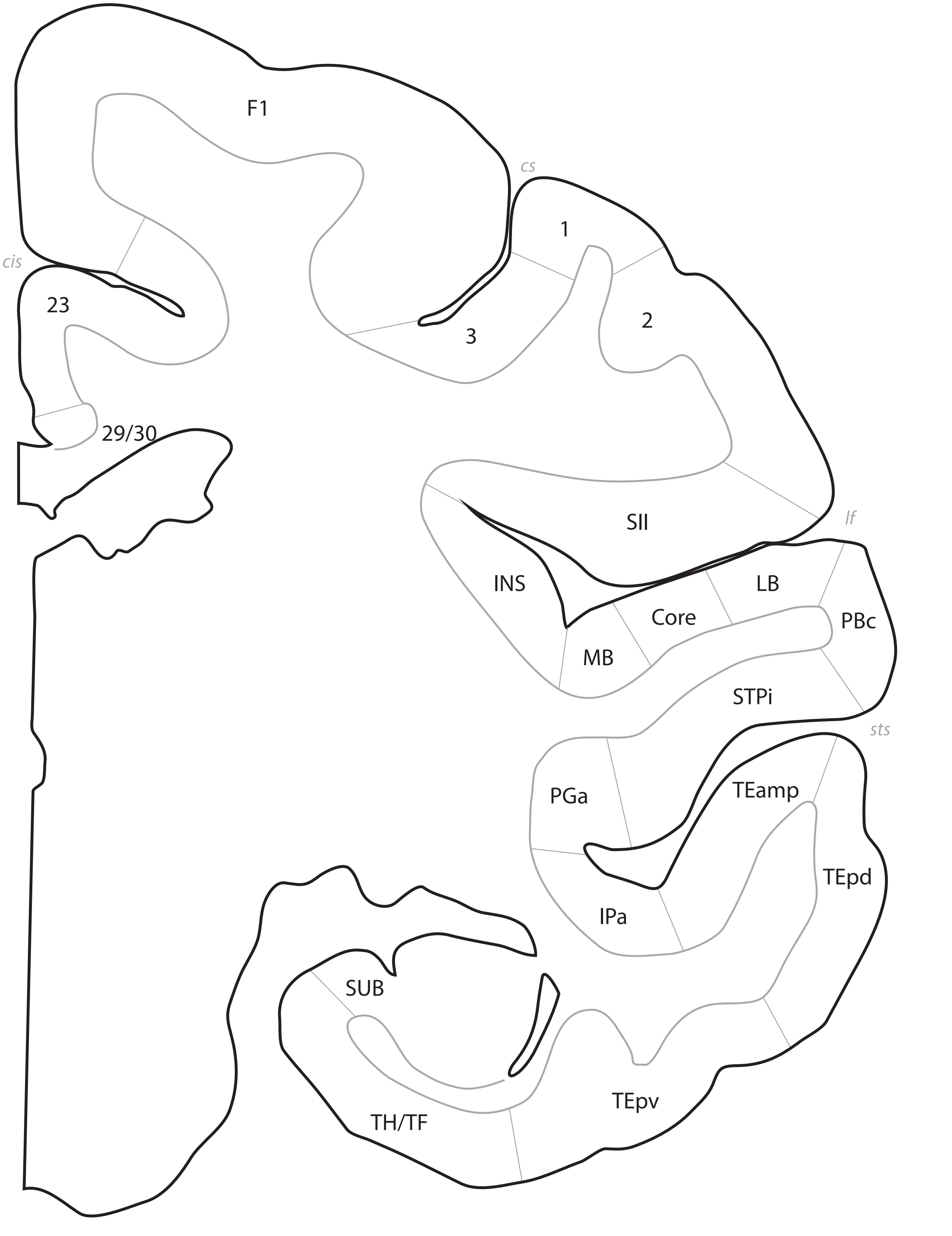
F1
cs
1
cis
23
3
2
29/30
SII
lf
INS
LB
Core
PBc
MB
STPi
sts
PGa
TEamp
TEpd
IPa
SUB
TEpv
TH/TF

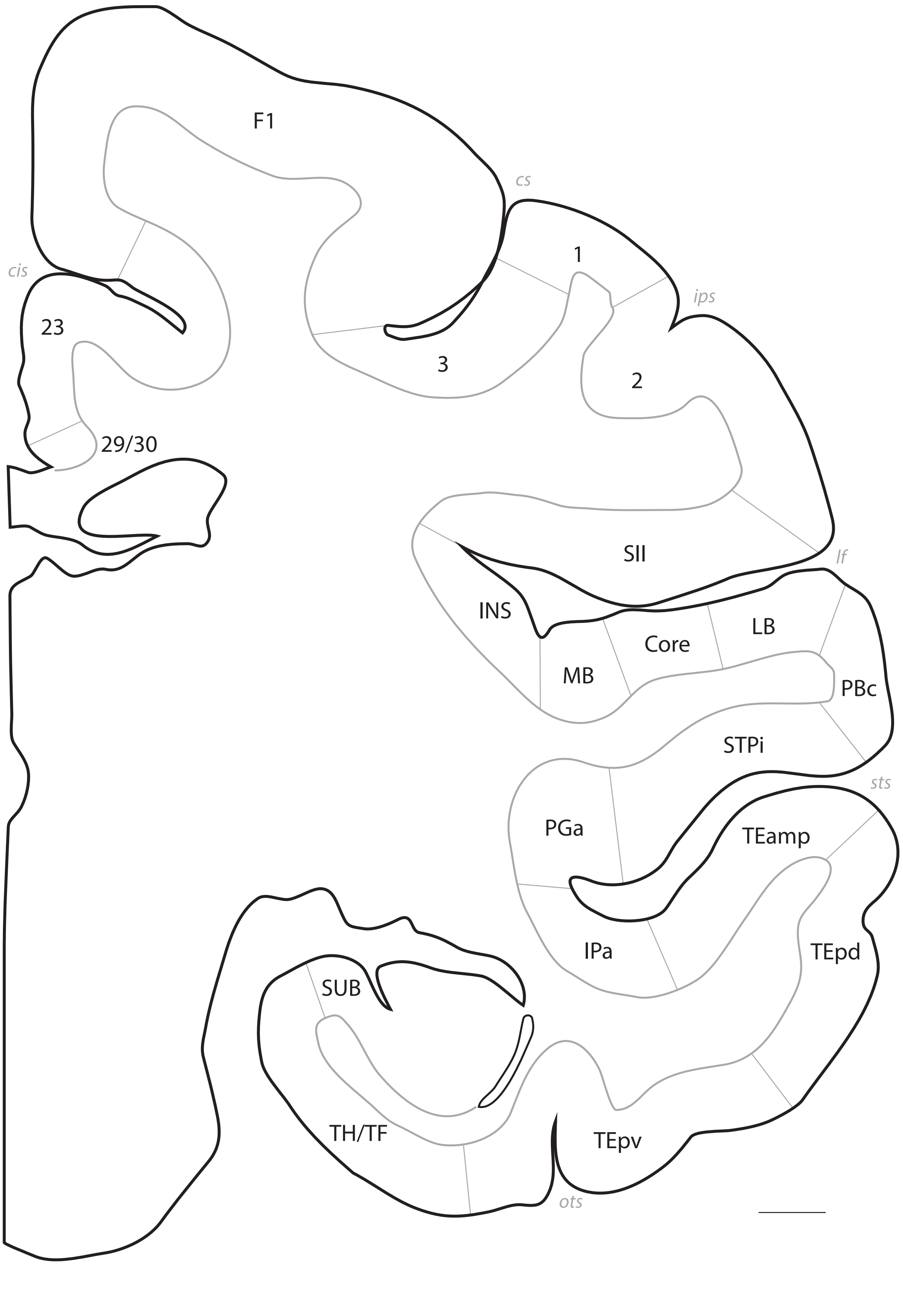
F1
cs
cis
1
ips
23
3
2
29/30
SII
lf
INS
LB
Core
MB
PBc
STPi
sts
PGa
TEamp
IPa
TEpd
SUB
TH/TF
TEpv
ots

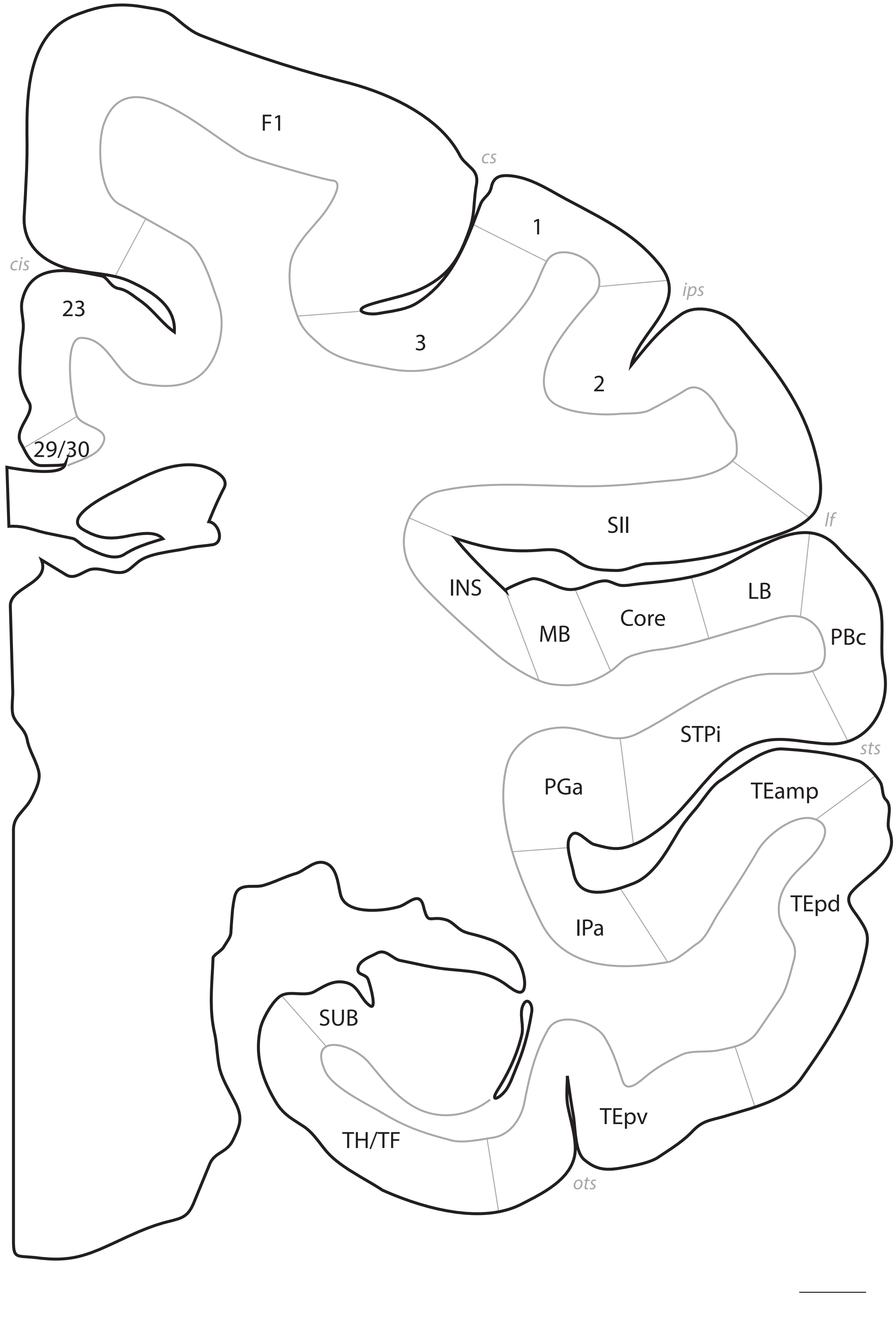
F1
cs
1
cis
ips
23
3
2
29/30
SII
lf
INS
LB
Core
MB
PBc
STPi
sts
PGa
TEamp
IPa
TEpd
SUB
TEpv
TH/TF
ots

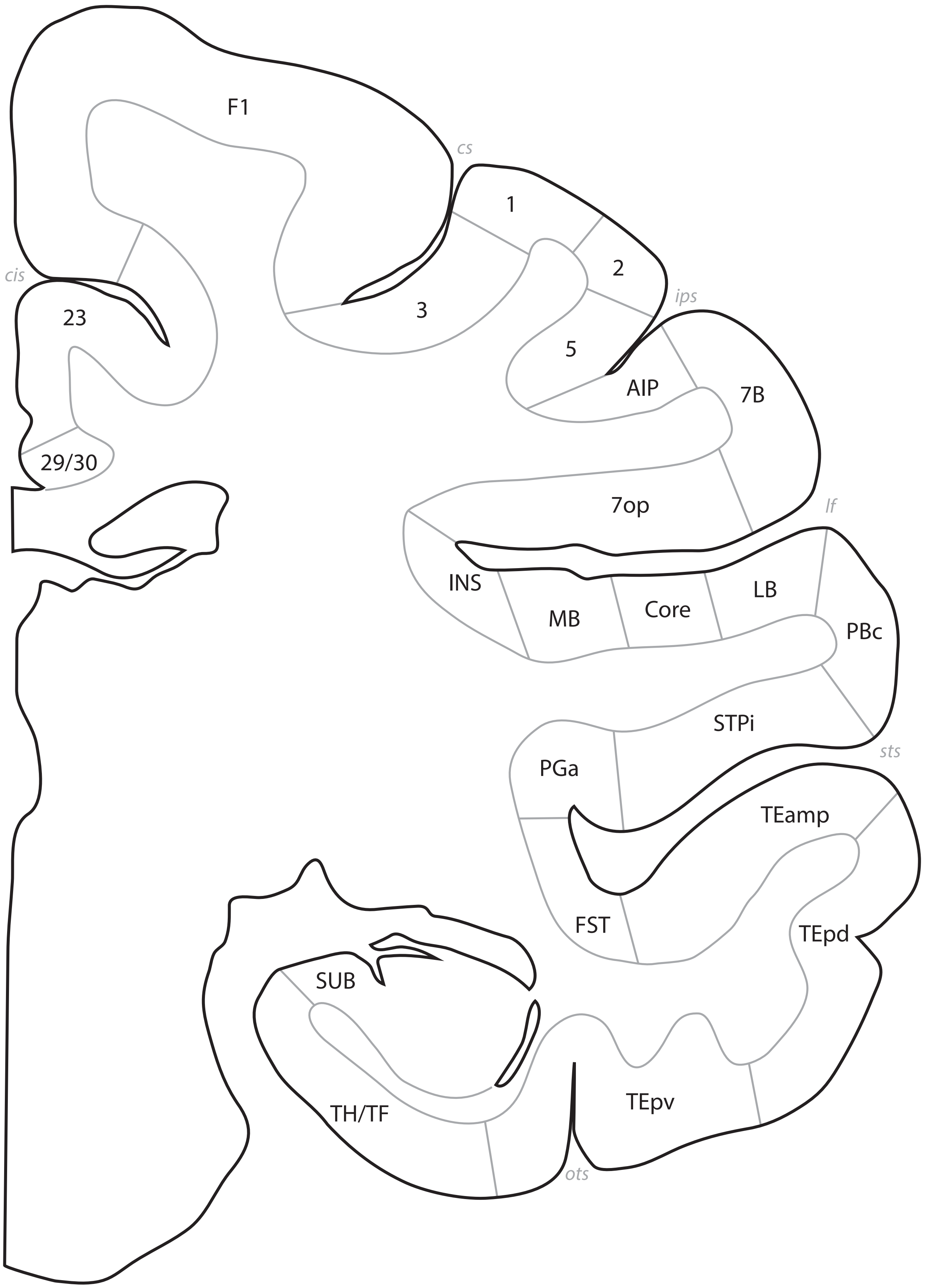
F1
cs
1
2
3
cis
23
5
ips
AIP
7B
29/30
7op
lf
INS
MB
Core
LB
PBc
STPi
PGa
sts
TEamp
FST
TEpd
SUB
TH/TF
TEpv
ots

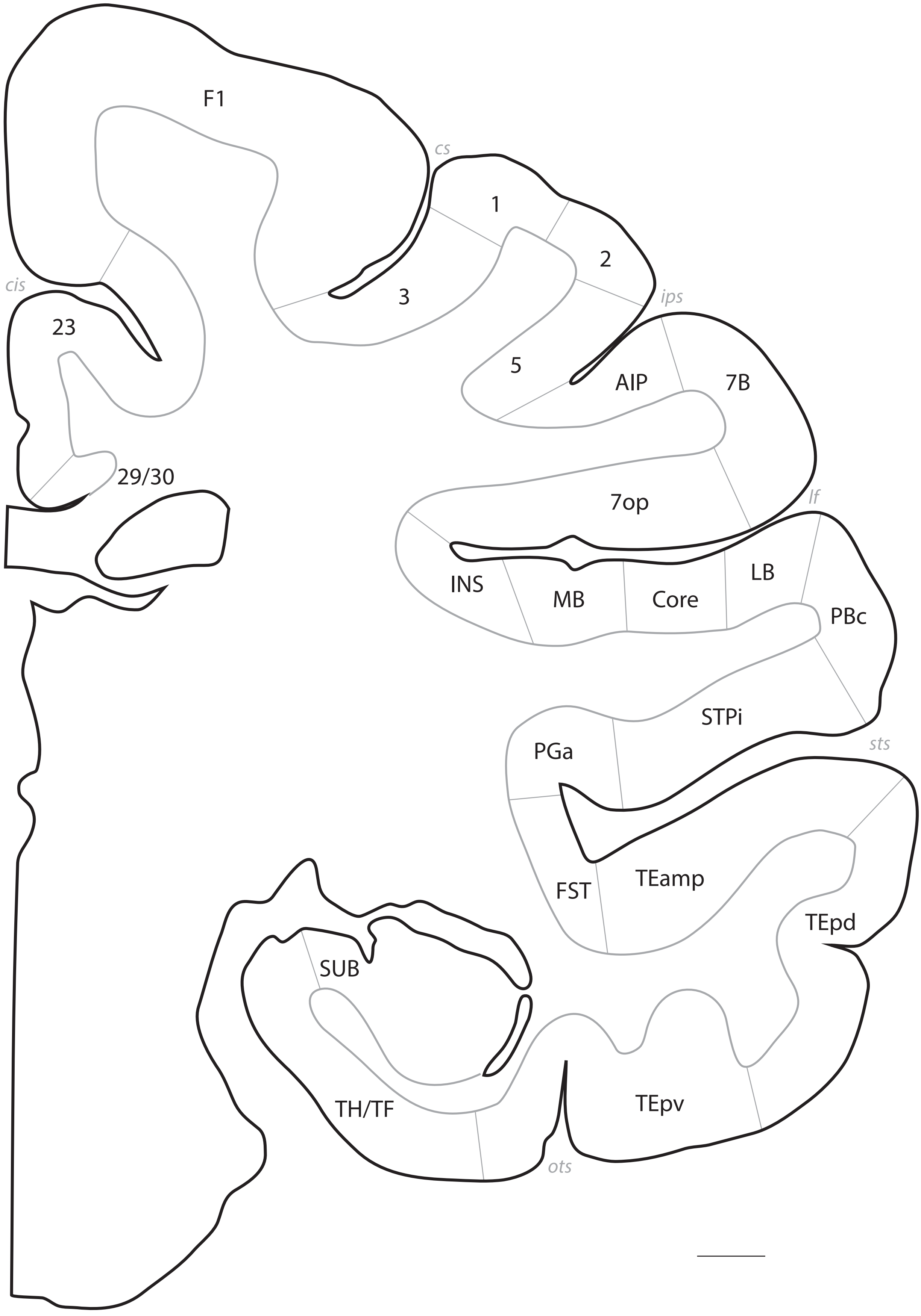
F1
cs
1
2
3
cis
23
ips
5
AIP
7B
29/30
7op
lf
INS
MB
Core
LB
PBc
STPi
sts
PGa
FST
TEamp
TEpd
SUB
TH/TF
TEpv
ots

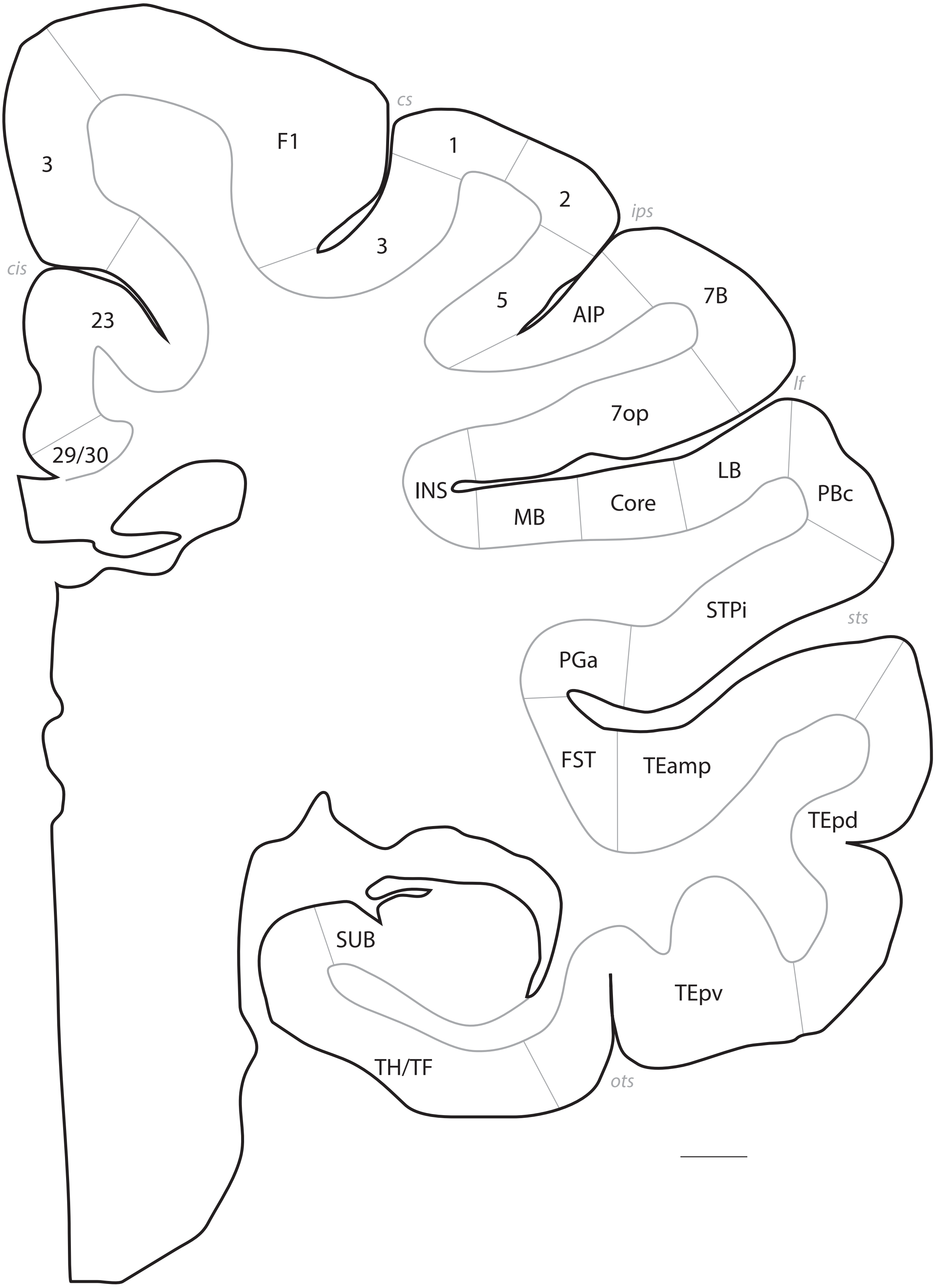
cs
F1
3
1
2
ips
3
cis
23
5
AIP
7B
29/30
7op
lf
INS
MB
Core
LB
PBc
STPi
sts
PGa
FST
TEamp
TEpd
SUB
TEpv
TH/TF
ots

# p. 071

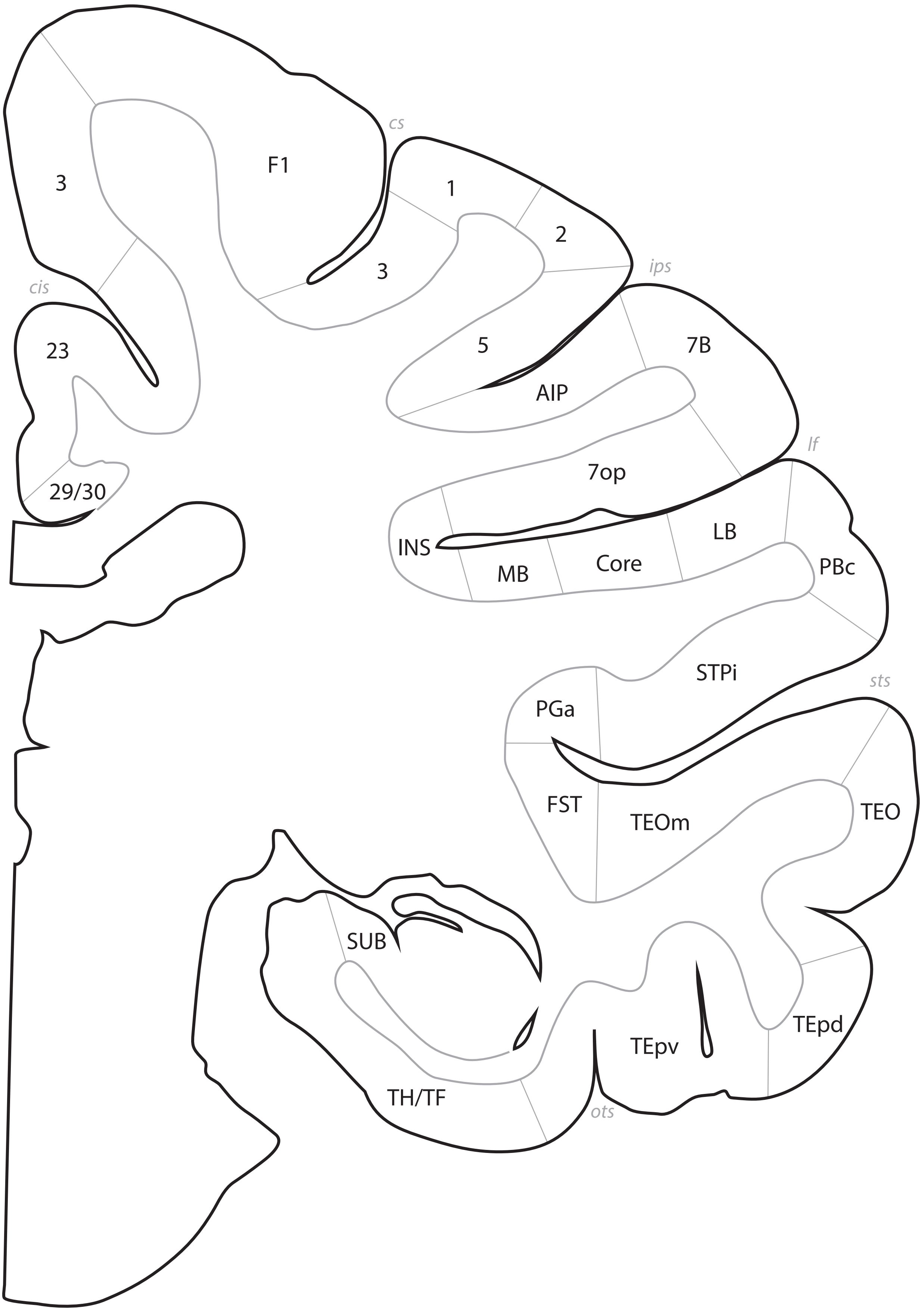

cs
3
F1
1
3
2
ips
cis
23
5
7B
AIP
29/30
lf
7op
INS
LB
MB
Core
PBc
STPi
sts
PGa
FST
TEOm
TEO
SUB
TEpd
TEpv
TH/TF
ots

# p. 072

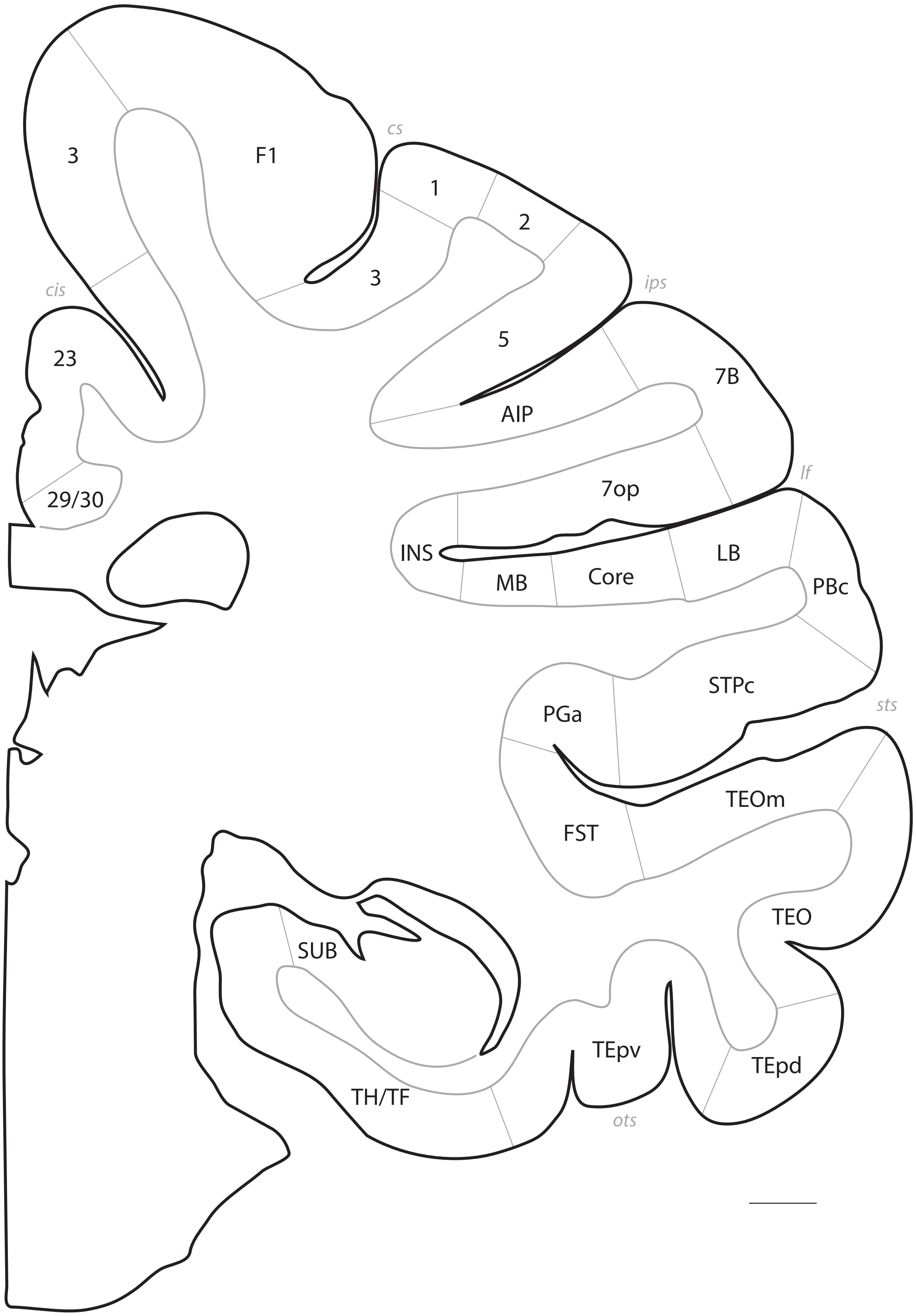
cs
3
F1
1
2
3
cis
ips
23
5
7B
AIP
29/30
7op
lf
INS
LB
MB
Core
PBc
STPc
sts
PGa
TEOm
FST
TEO
SUB
TEpv
TEpd
TH/TF
ots

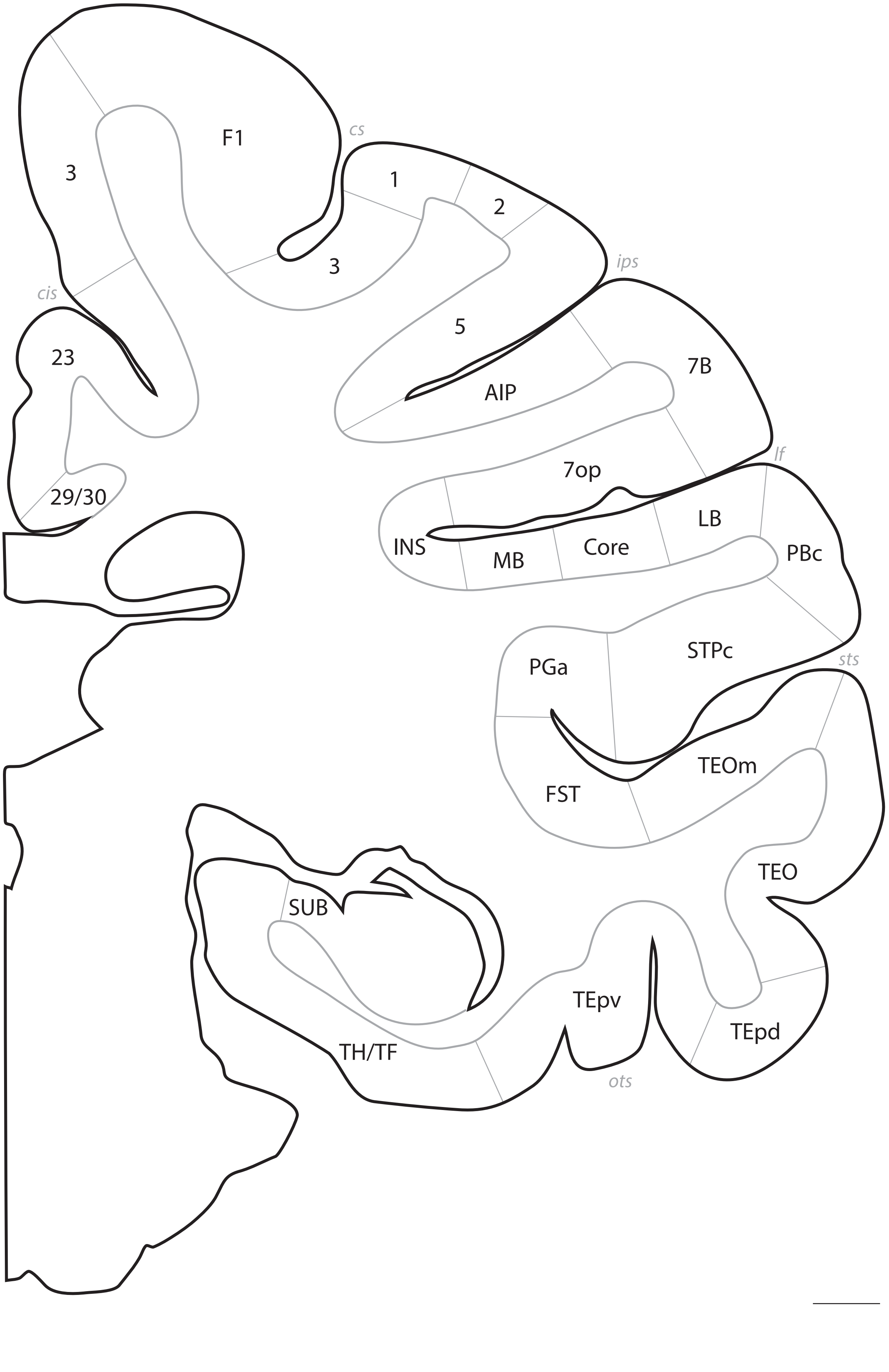
cs
F1
3
1
2
3
ips
5
cis
23
AIP
7B
29/30
7op
lf
LB
INS
Core
MB
PBc
STPc
sts
PGa
TEOm
FST
TEO
SUB
TEpv
TEpd
TH/TF
ots

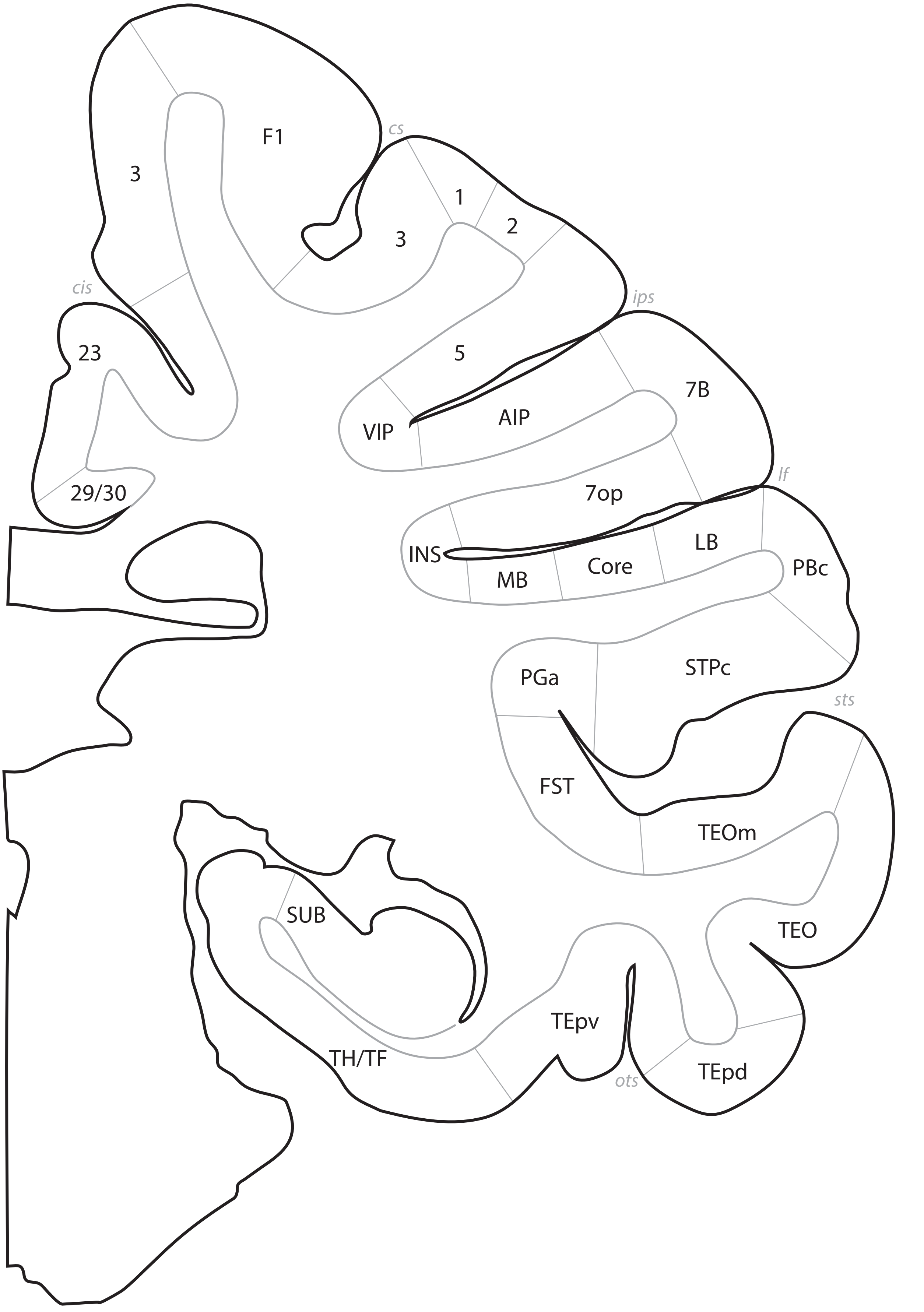
F1
3
cs
1
2
3
cis
23
5
ips
VIP
AIP
7B
29/30
7op
lf
INS
LB
Core
MB
PBc
PGa
STPc
sts
FST
TEOm
SUB
TEO
TEpv
TH/TF
TEpd
ots

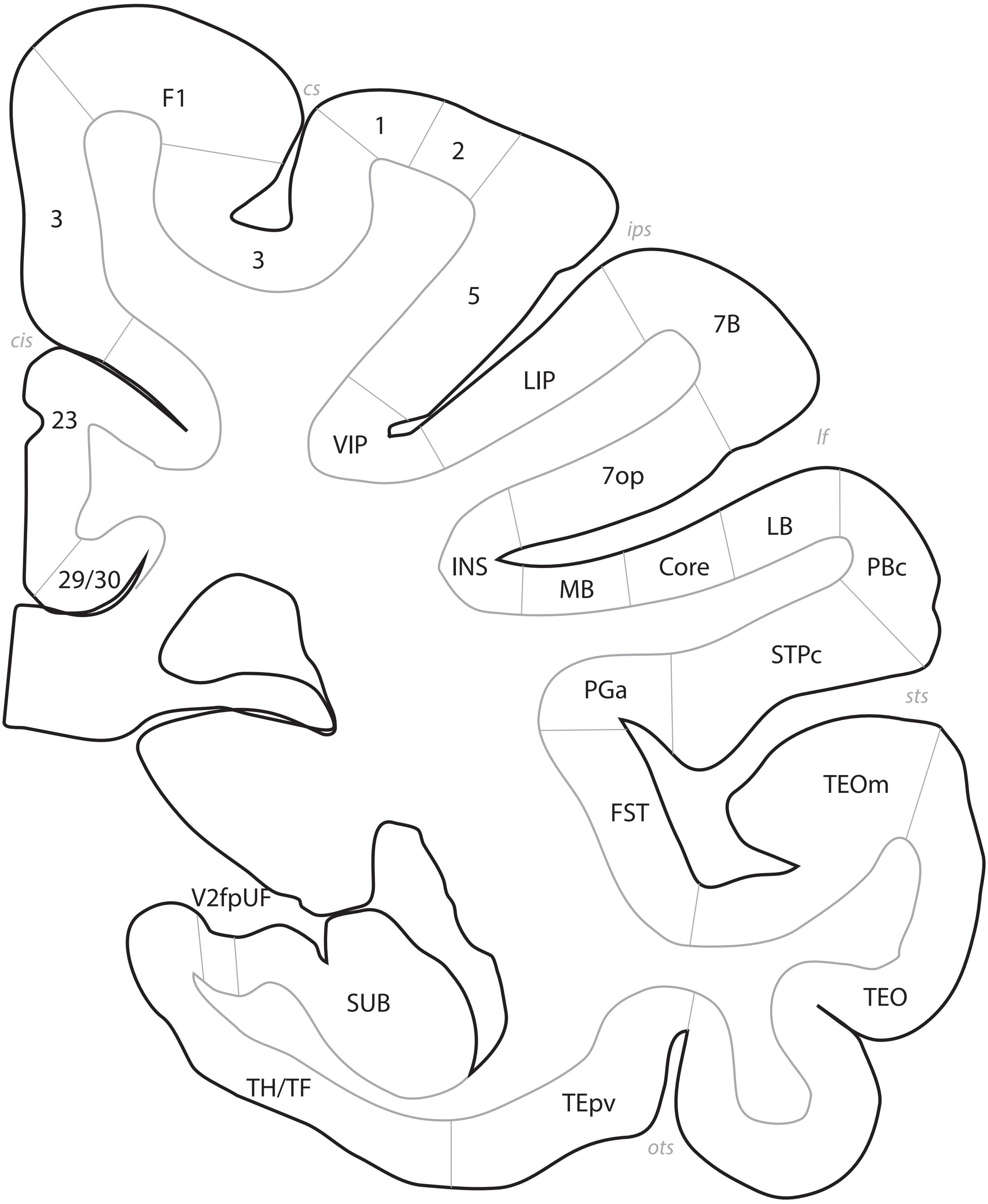

F1
cs
1
2
3
3
ips
5
7B
cis
LIP
23
VIP
lf
7op
LB
INS
Core
PBc
MB
29/30
STPc
PGa
sts
TEOm
FST
V2fpUF
SUB
TEO
TH/TF
TEpv
ots

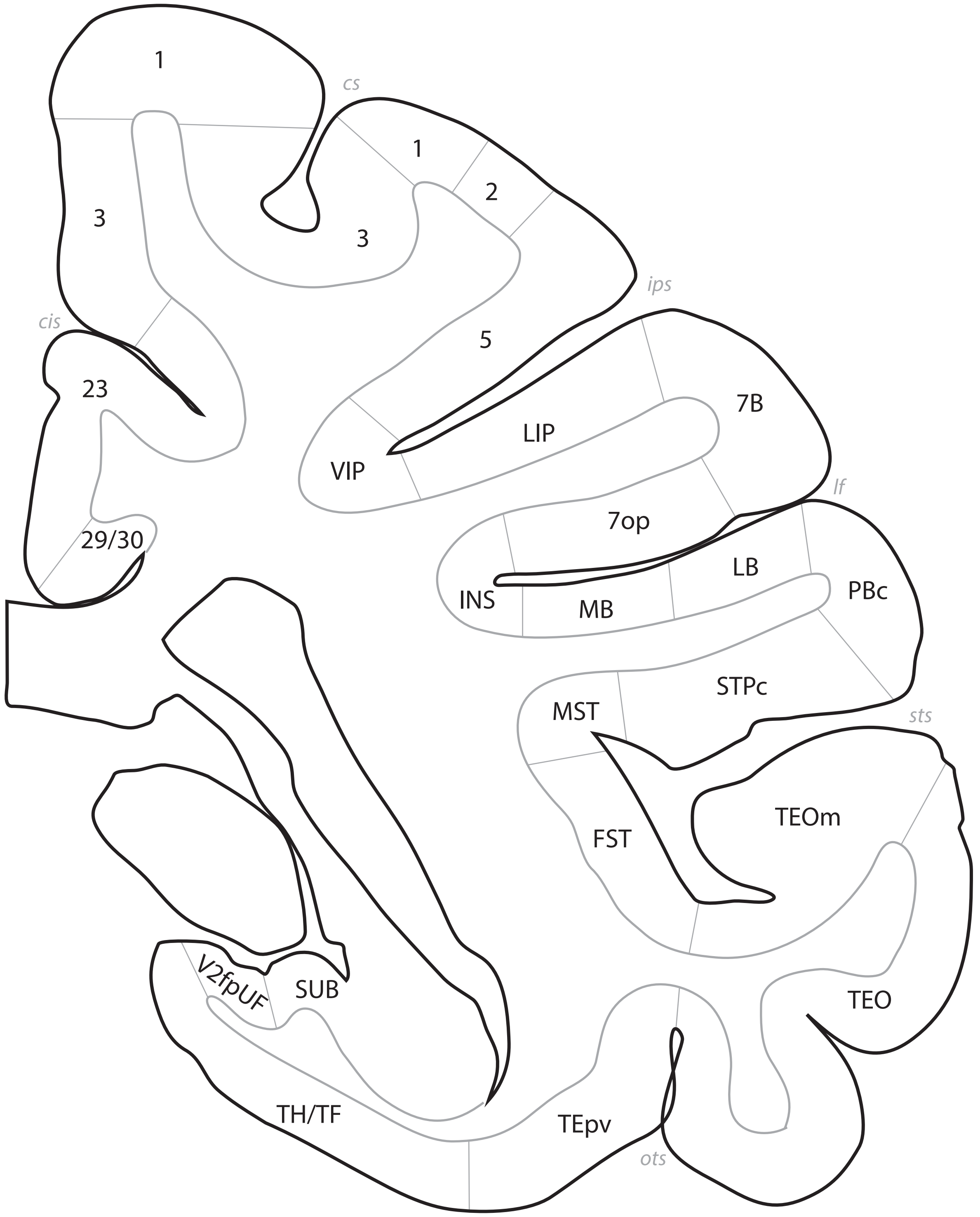

cs
1
1
2
3
3
3
cis
ips
5
23
7B
LIP
VIP
29/30
lf
7op
LB
INS
MB
PBc
STPc
MST
sts
TEOm
FST
V2fpUF
SUB
TEO
TH/TF
TEpv
ots

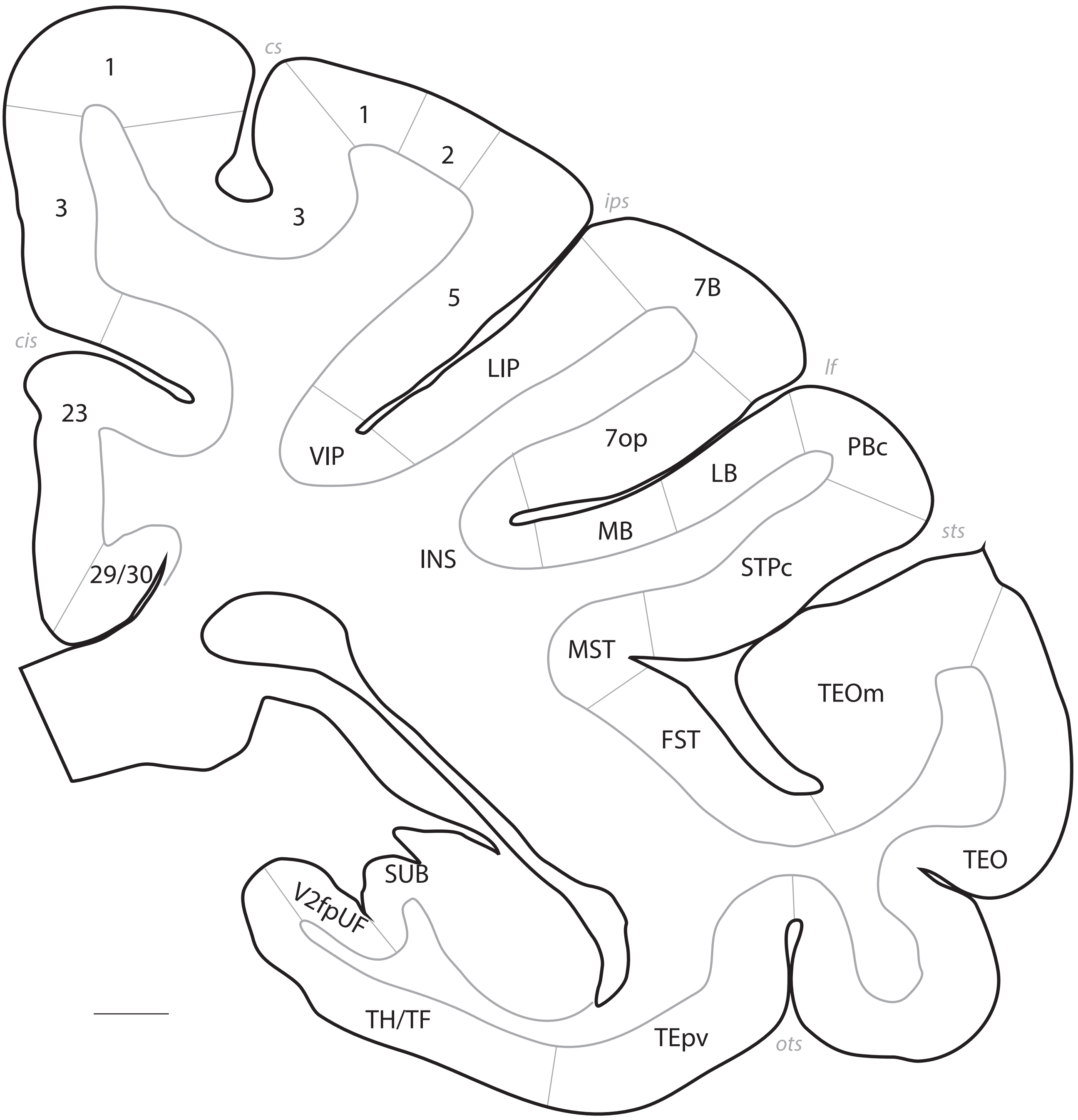

cs
1
1
2
3
3
ips
5
7B
cis
LIP
lf
23
7op
PBc
VIP
LB
MB
sts
INS
STPc
29/30
MST
TEOm
FST
SUB
TEO
V2fpUF
TH/TF
TEpv
ots

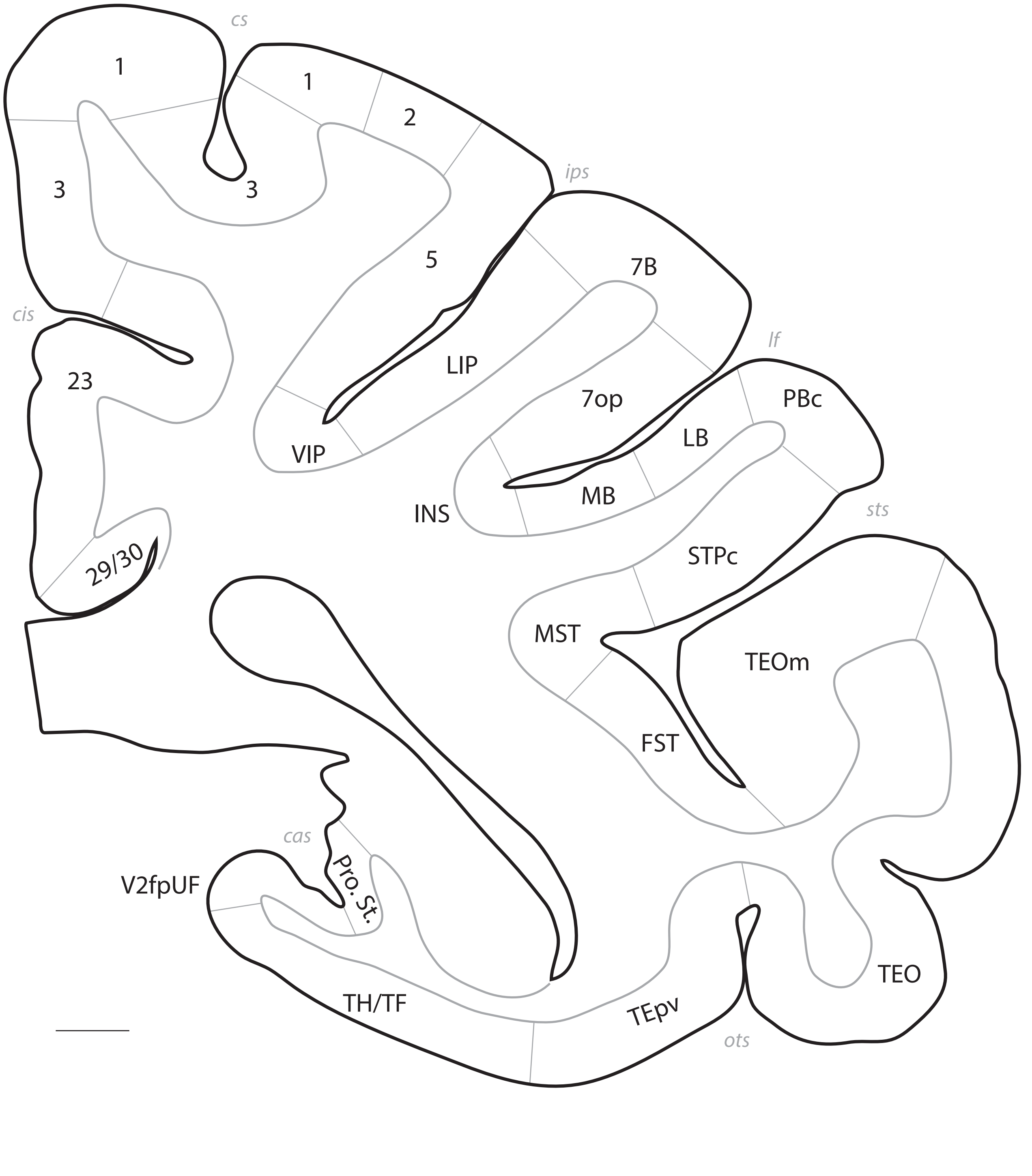

cs
1
1
2
3
3
ips
5
7B
cis
23
LIP
lf
7op
PBc
VIP
LB
INS
MB
sts
STPc
29/30
MST
TEOm
FST
cas
Pro. St.
V2fpUF
TEO
TH/TF
TEpv
ots

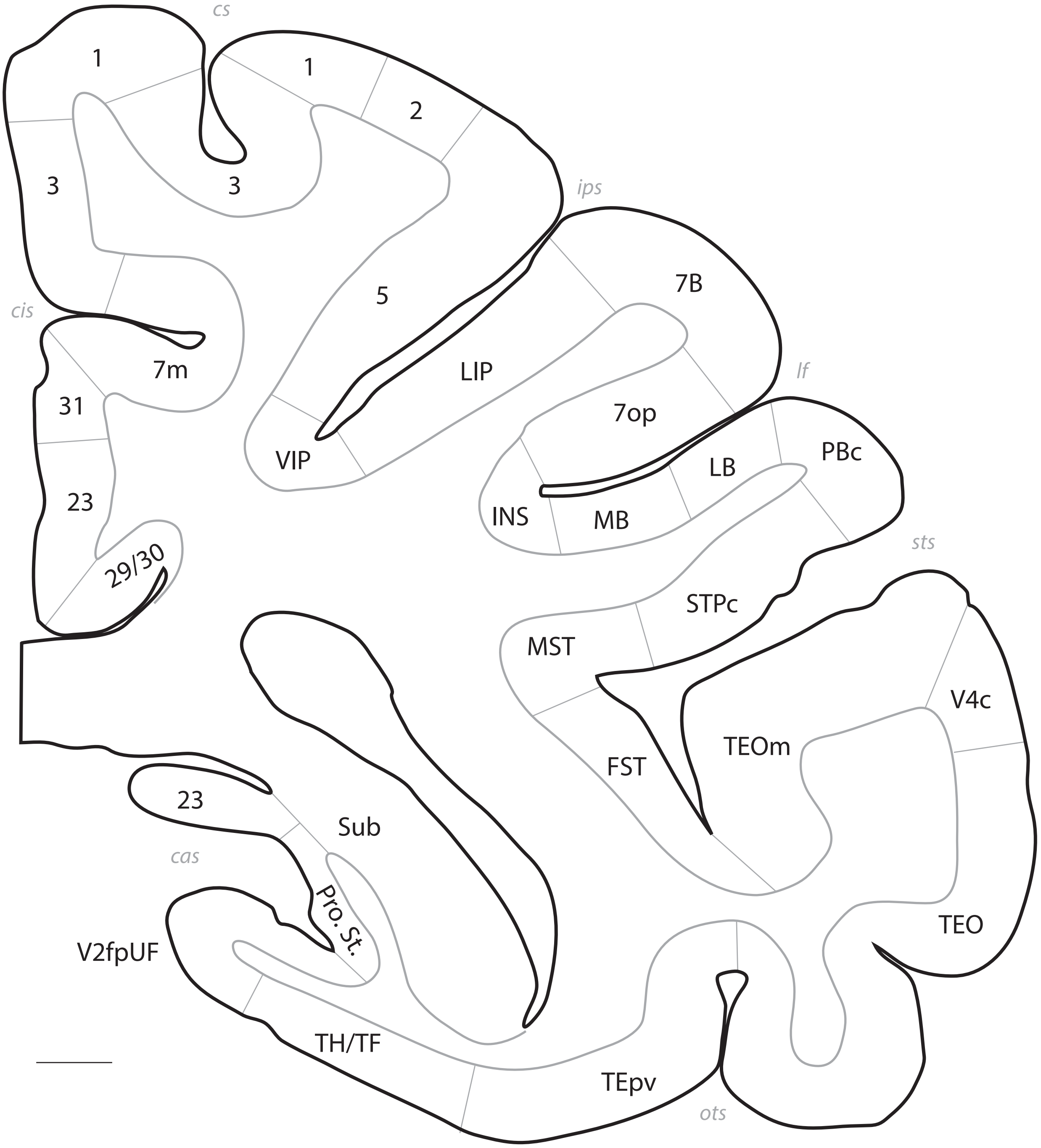
cs
1
1
2
3
3
ips
5
7B
cis
7m
LIP
lf
31
7op
VIP
LB
PBc
23
INS
MB
sts
29/30
STPc
MST
V4c
TEOm
FST
23
Sub
cas
Pro. St.
TEO
V2fpUF
TH/TF
TEpv
ots

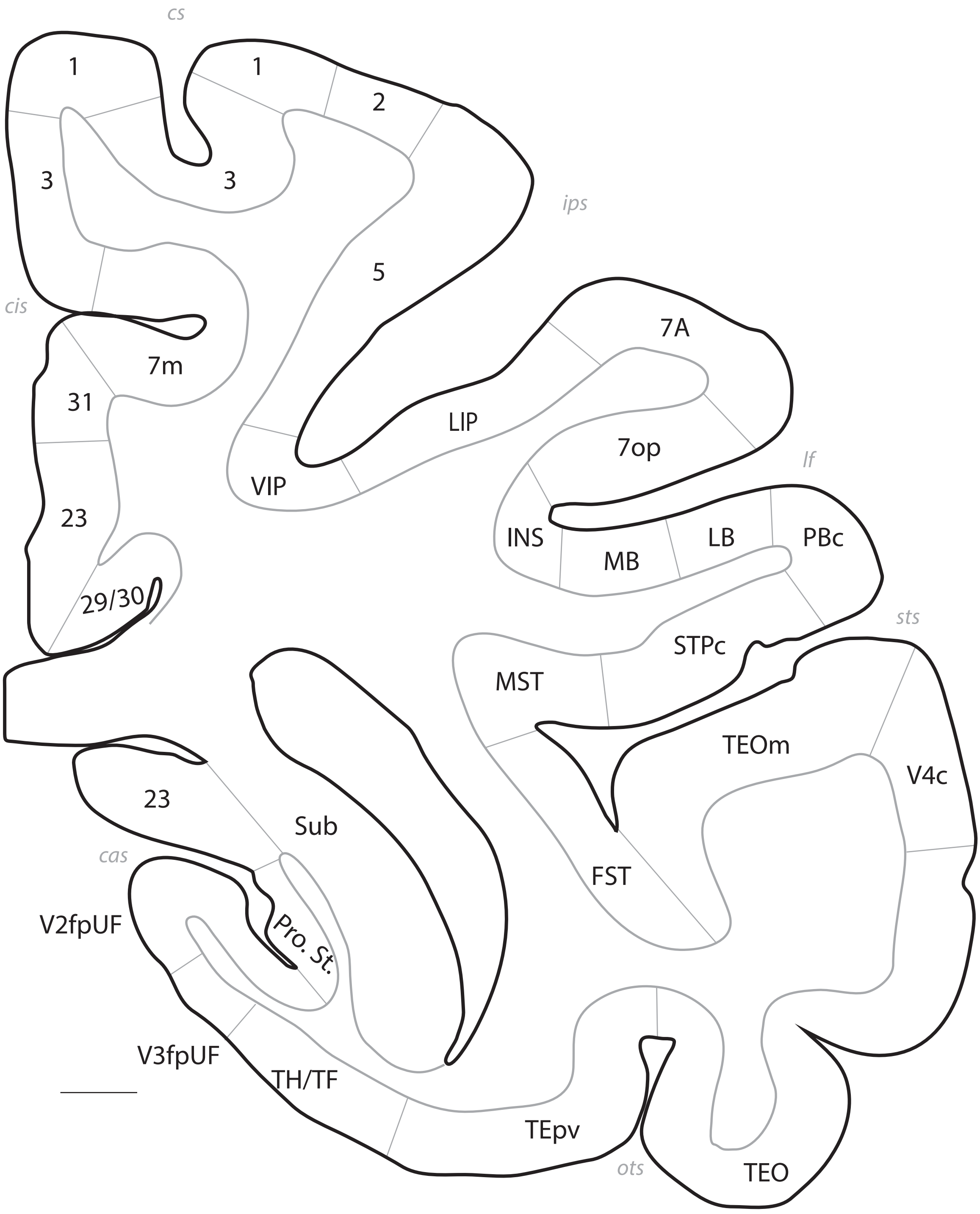
cs
1
1
2
3
3
ips
5
cis
7m
7A
31
LIP
7op
lf
VIP
23
INS
MB
LB
PBc
29/30
sts
STPc
MST
TEOm
V4c
23
Sub
cas
FST
V2fpUF
Pro. St.
V3fpUF
TH/TF
TEpv
ots
TEO

# p. 081

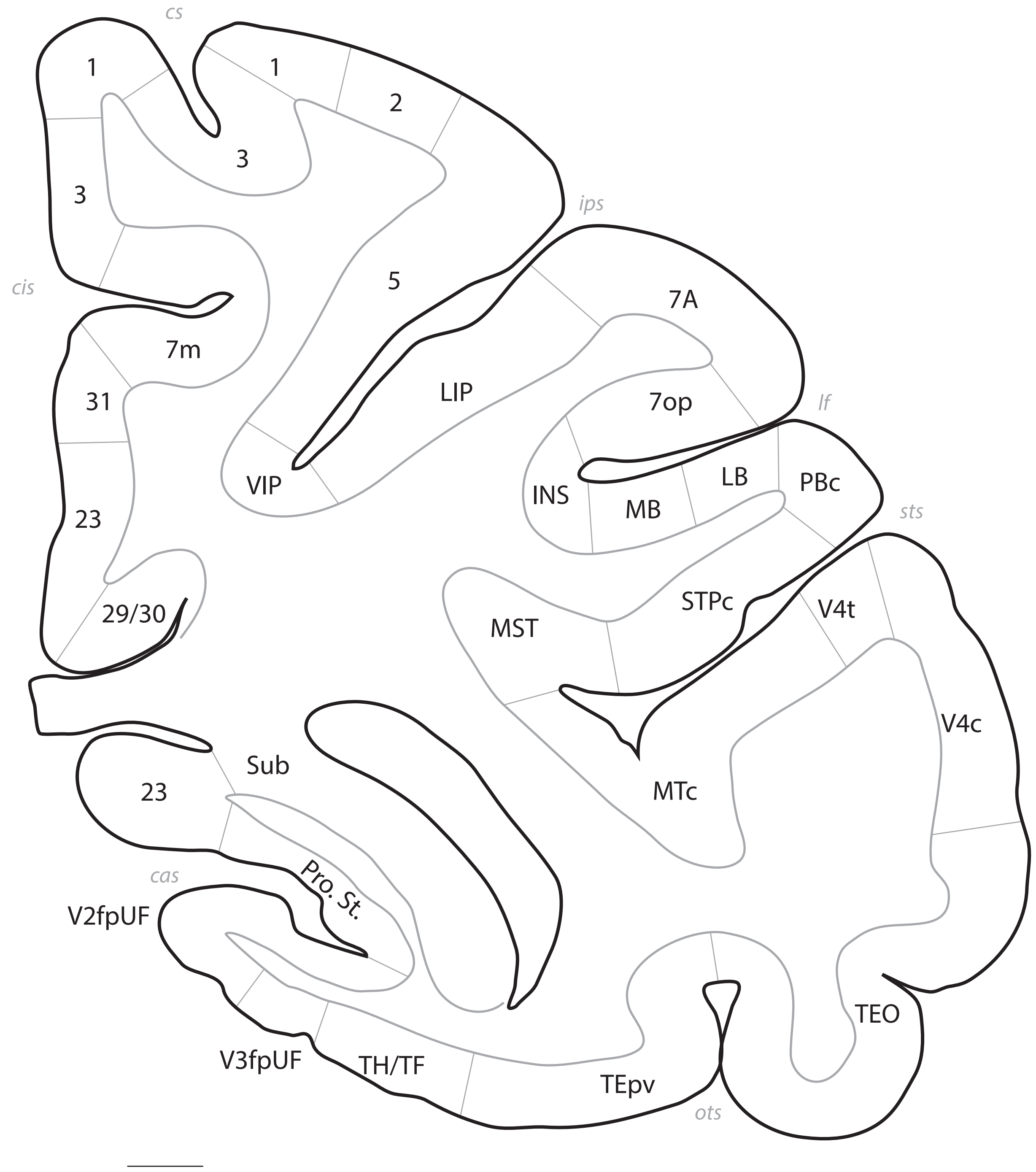
cs
1
1
2
3
3
ips
5
cis
7A
7m
31
LIP
7op
lf
VIP
INS
LB
PBc
MB
sts
23
STPc
MST
V4t
29/30
V4c
Sub
MTc
23
cas
Pro. St.
V2fpUF
TEO
V3fpUF
TH/TF
TEpv
ots

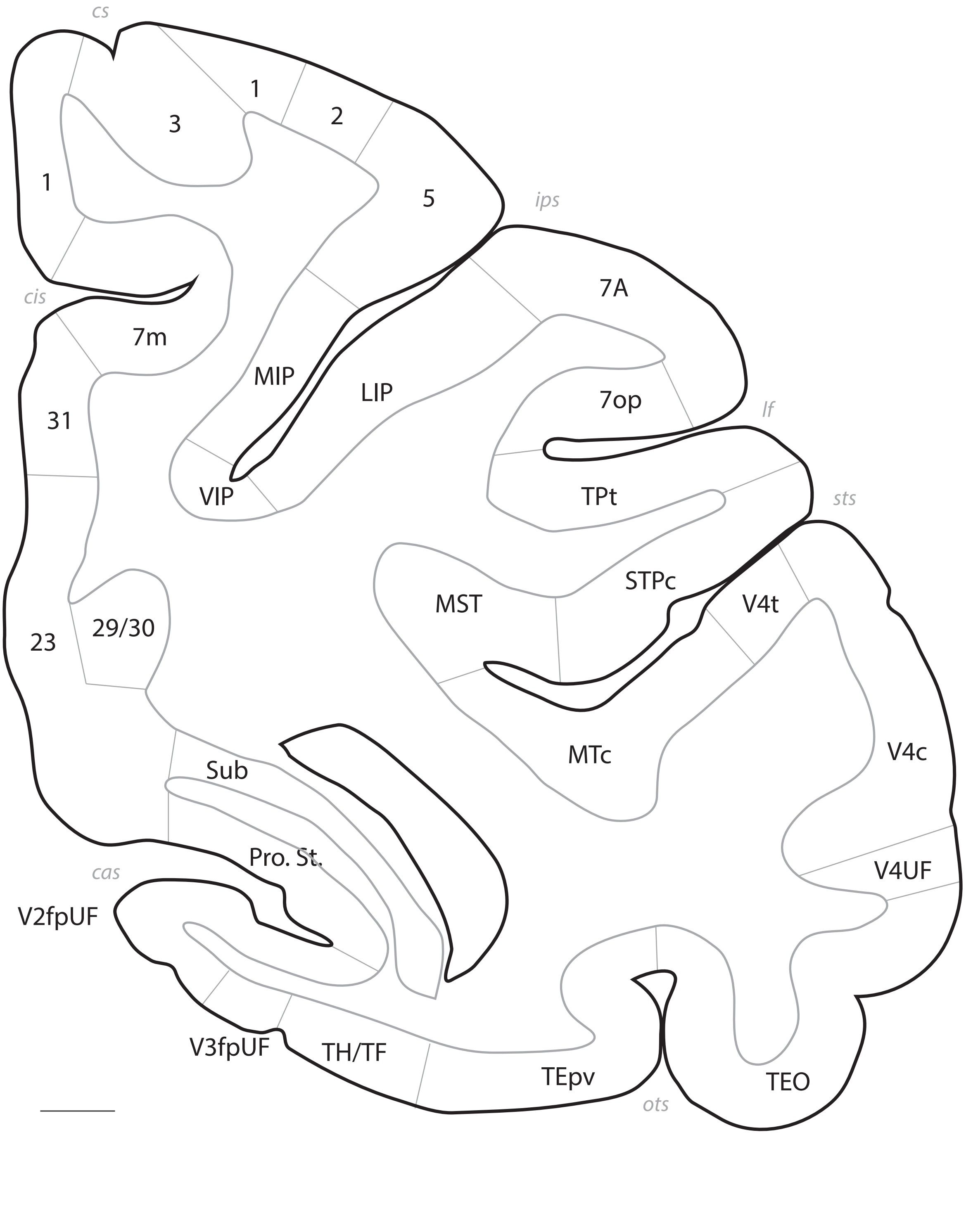
cs
1
3
1
2
5
ips
cis
7m
7A
MIP
LIP
7op
31
lf
VIP
TPt
sts
STPc
MST
V4t
23
29/30
MTc
V4c
Sub
Pro. St.
V4UF
cas
V2fpUF
V3fpUF
TH/TF
TEpv
ots
TEO

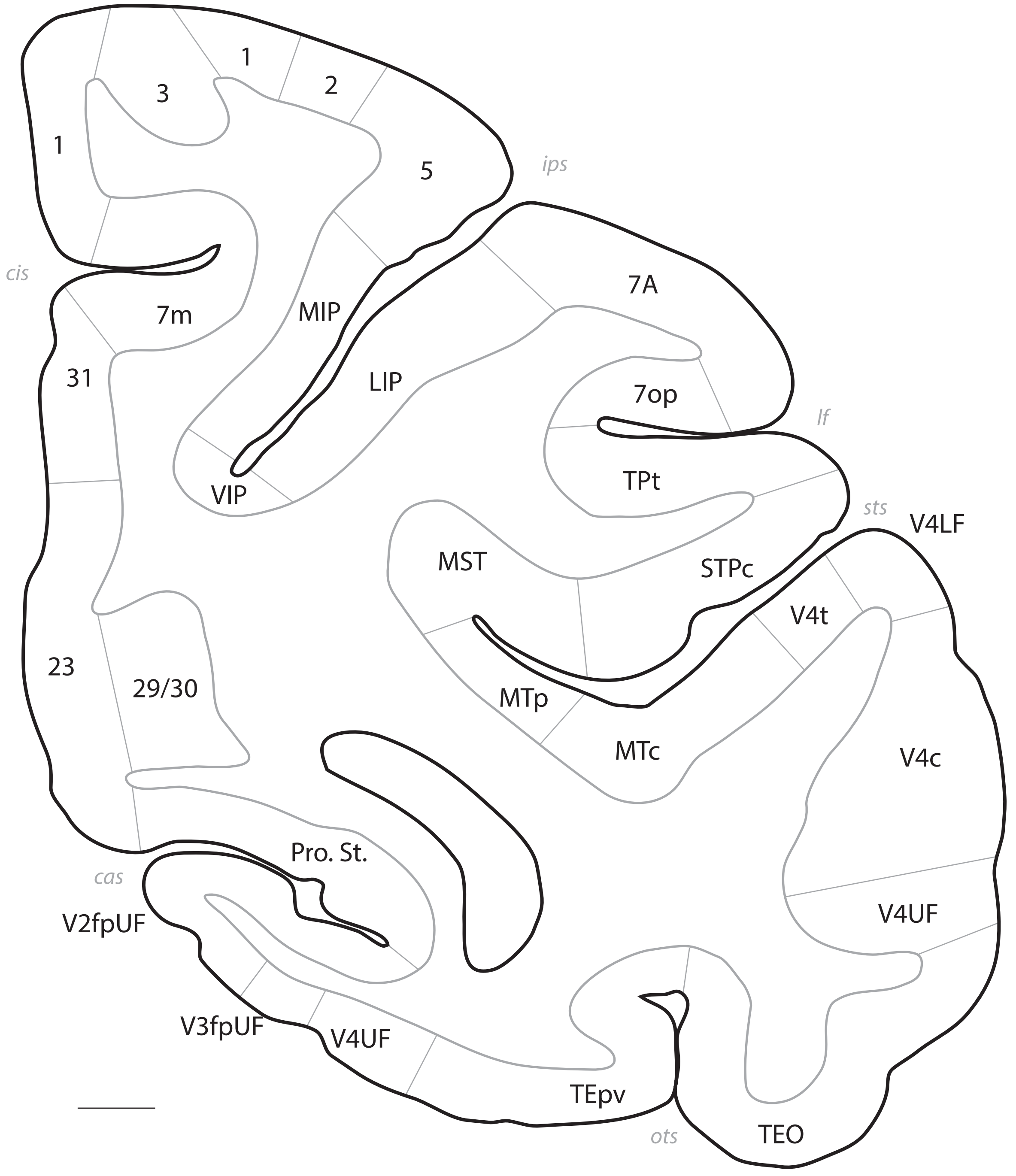

1
3
1
2
5
ips
cis
7m
MIP
7A
31
LIP
7op
lf
VIP
TPt
sts
V4LF
MST
STPc
V4t
23
29/30
MTp
MTc
V4c
Pro. St.
cas
V4UF
V2fpUF
V3fpUF
V4UF
TEpv
ots
TEO

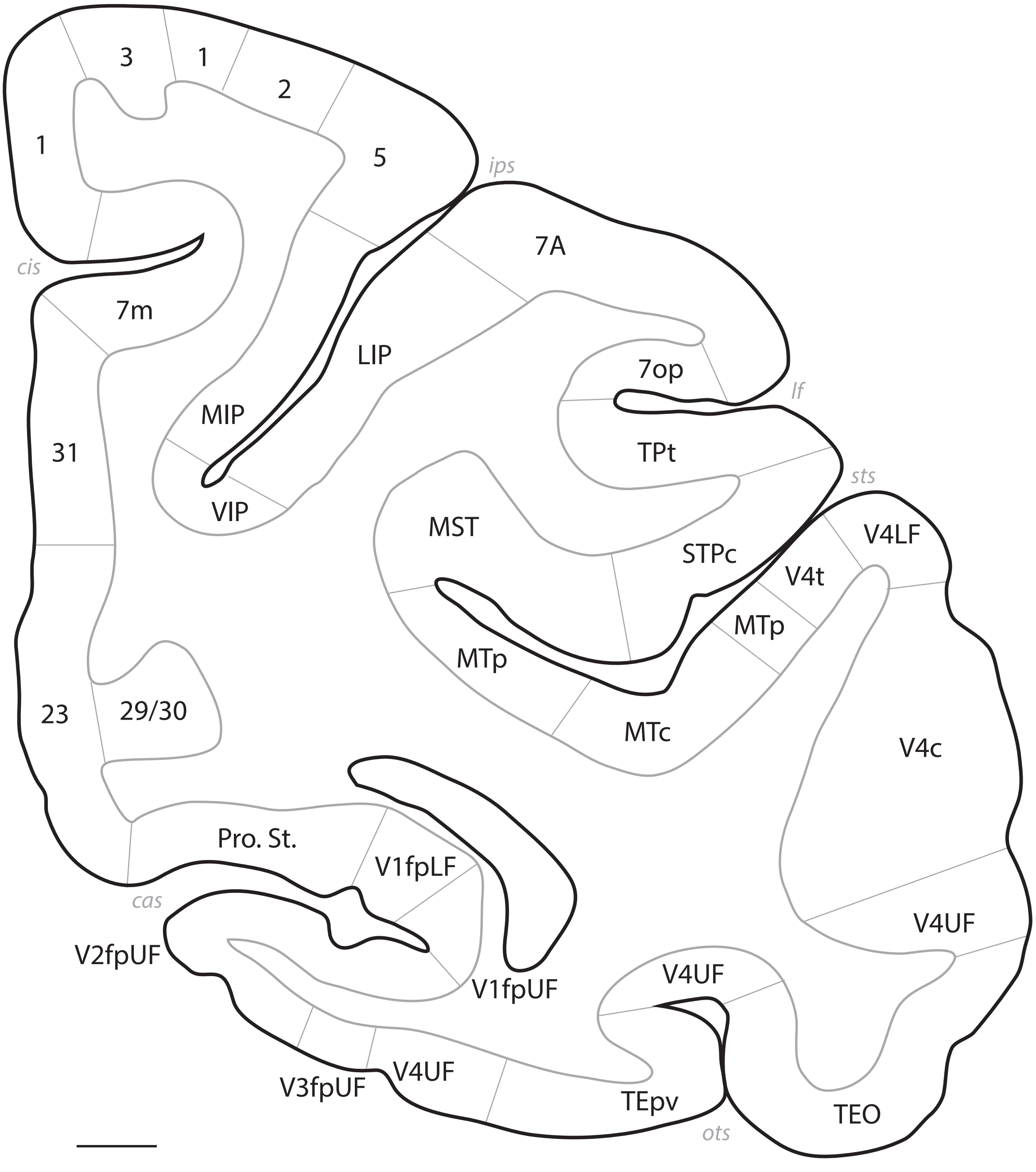
3
1
2
1
5
ips
7A
cis
7m
LIP
7op
lf
31
MIP
TPt
VIP
sts
MST
V4LF
STPc
V4t
MTp
MTp
23
29/30
MTc
V4c
Pro. St.
V1fpLF
V4UF
cas
V2fpUF
V4UF
V1fpUF
V4UF
V3fpUF
TEpv
TEO
ots

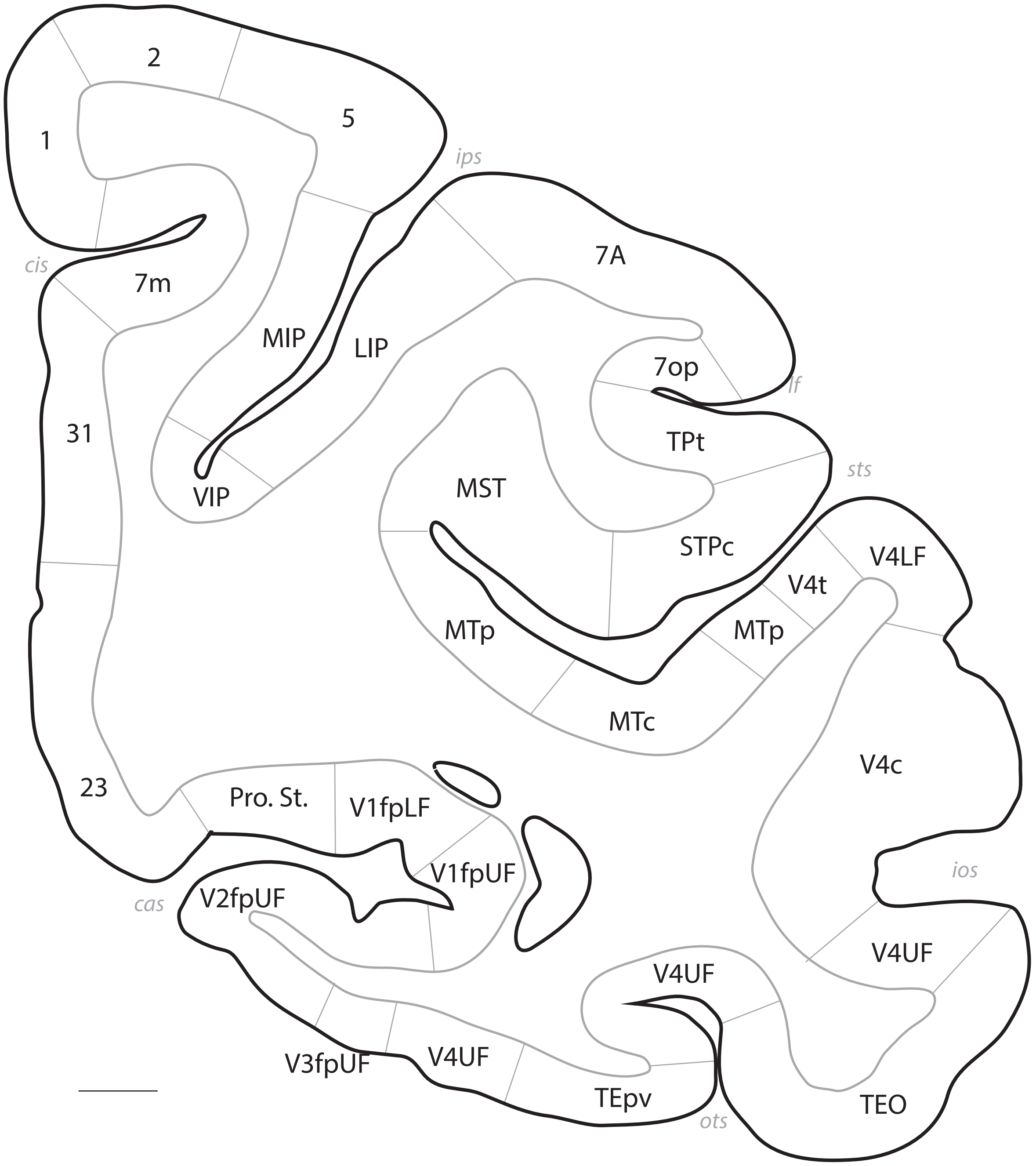

2
5
1
ips
7A
cis
7m
MIP
LIP
7op
lf
31
TPt
sts
MST
VIP
STPc
V4LF
V4t
MTp
MTp
MTc
V4c
23
Pro. St.
V1fpLF
V1fpUF
ios
cas
V2fpUF
V4UF
V4UF
V4UF
V3fpUF
TEpv
ots
TEO

# p. 086

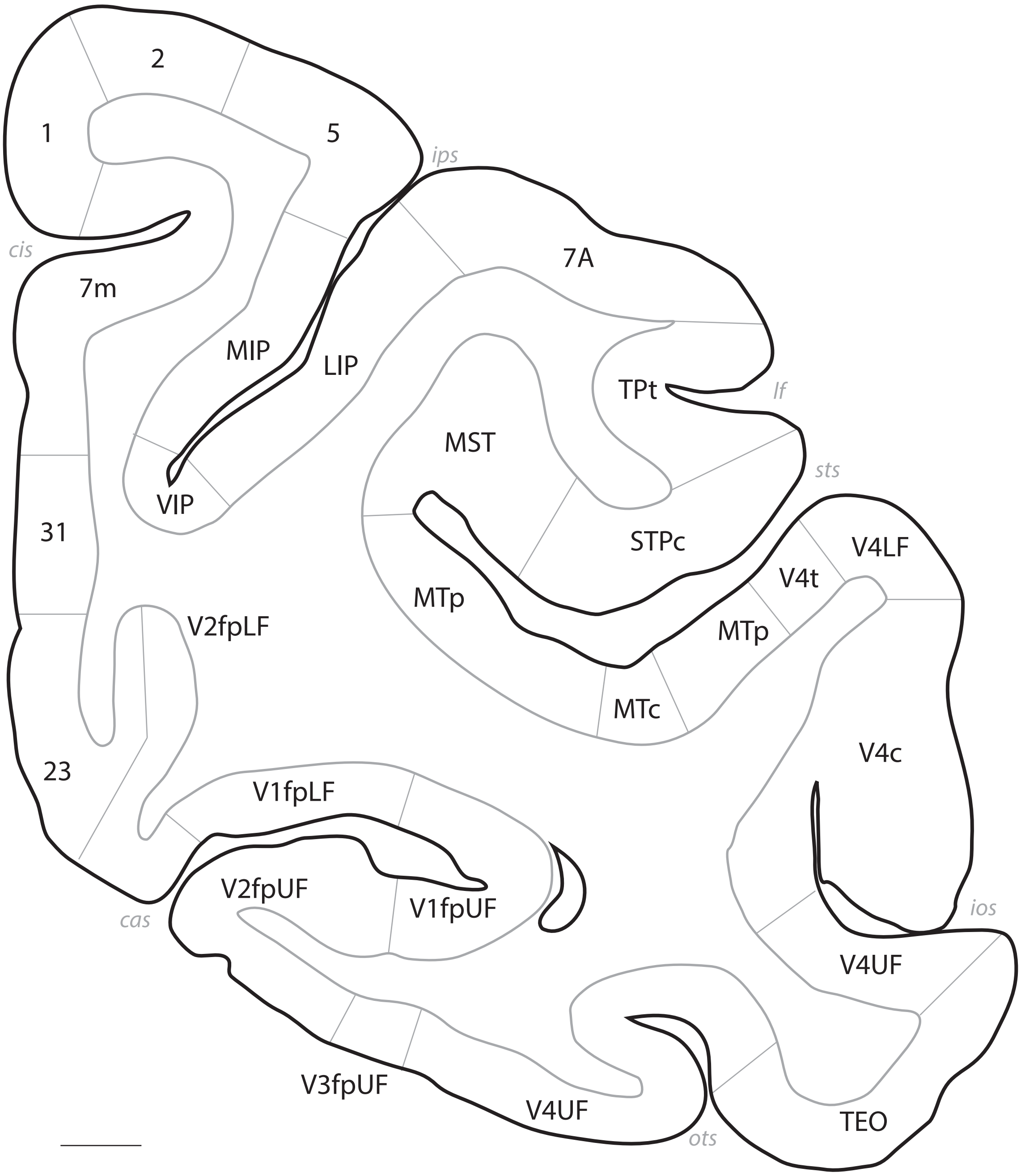

2
1
5
ips
cis
7m
7A
MIP
LIP
TPt
lf
MST
sts
VIP
31
STPc
V4LF
V4t
MTp
V2fpLF
MTp
MTc
V4c
23
V1fpLF
V2fpUF
V1fpUF
cas
ios
V4UF
V3fpUF
V4UF
TEO
ots

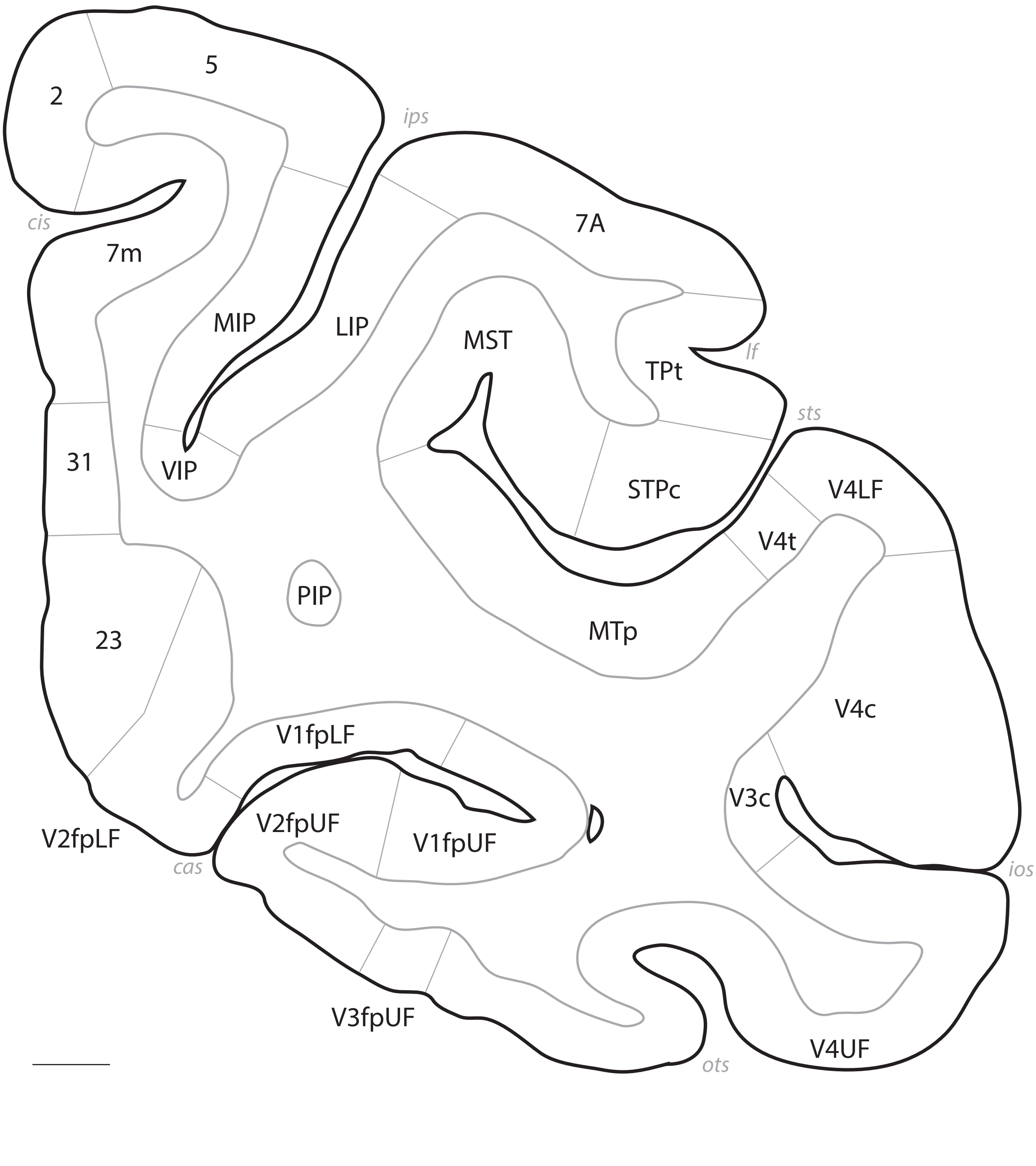

5
2
ips
cis
7m
7A
MIP
LIP
MST
TPt
lf
sts
VIP
31
STPc
V4LF
V4t
PIP
23
MTp
V4c
V1fpLF
V3c
V2fpUF
V2fpLF
V1fpUF
cas
ios
V3fpUF
V4UF
ots

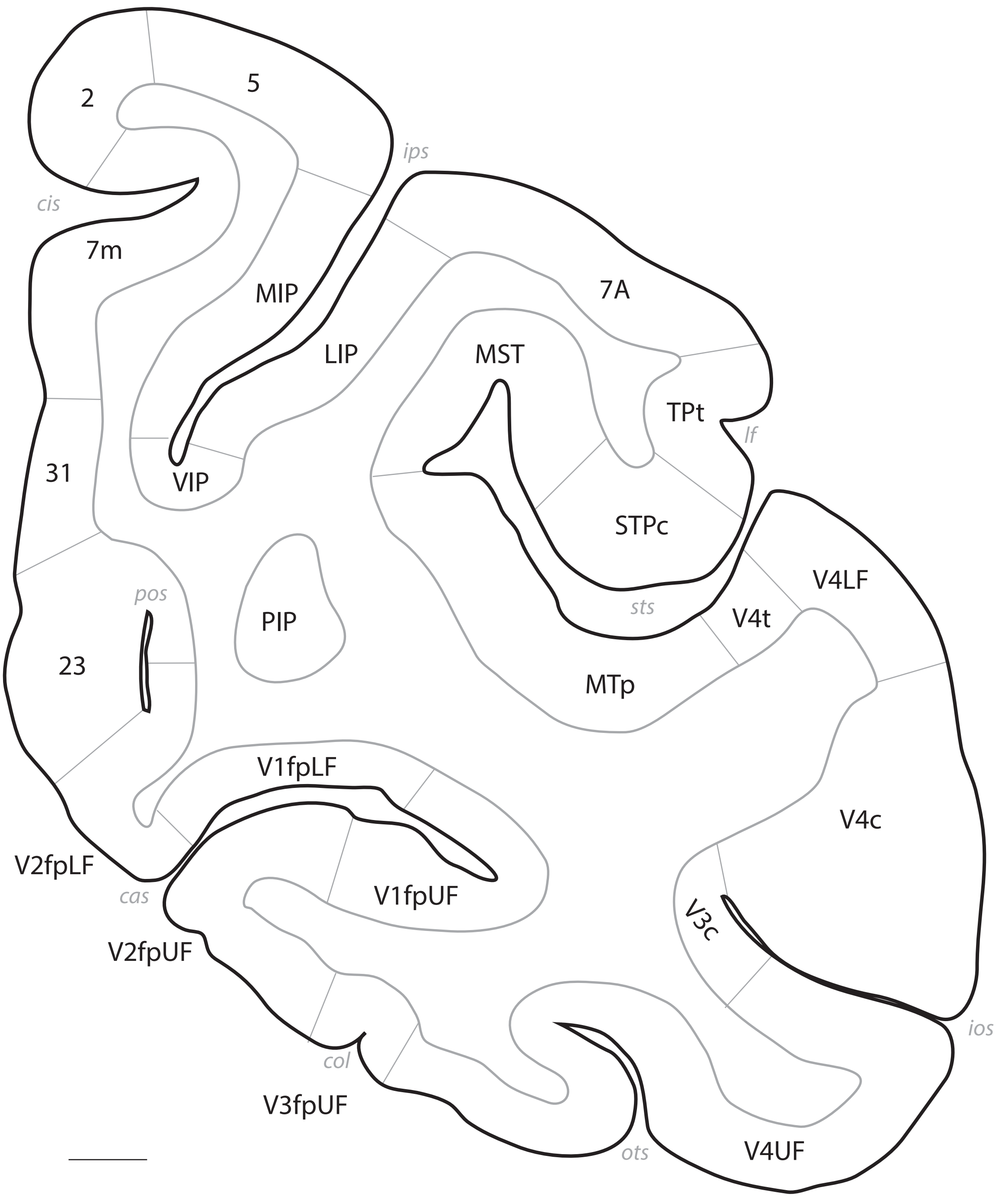
2
5
ips
cis
7m
MIP
LIP
7A
MST
TPt
lf
VIP
31
STPc
pos
PIP
sts
V4t
V4LF
23
MTp
V1fpLF
V4c
V2fpLF
cas
V1fpUF
V3c
V2fpUF
ios
col
V3fpUF
ots
V4UF

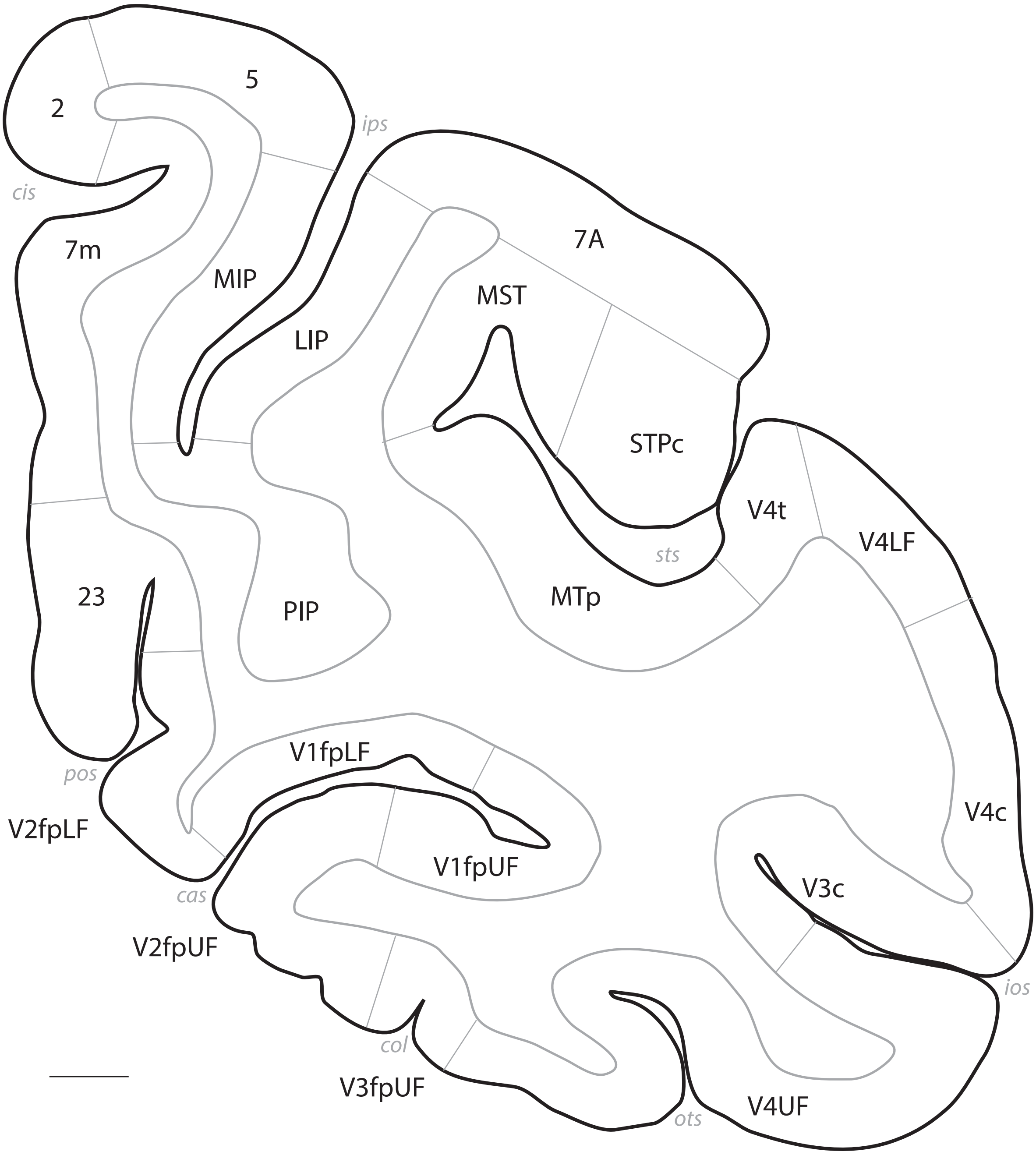
2
5
ips
cis
7m
MIP
7A
MST
LIP
STPc
V4t
V4LF
sts
23
PIP
MTp
V1fpLF
pos
V2fpLF
V4c
V1fpUF
V3c
cas
V2fpUF
ios
col
V3fpUF
ots
V4UF

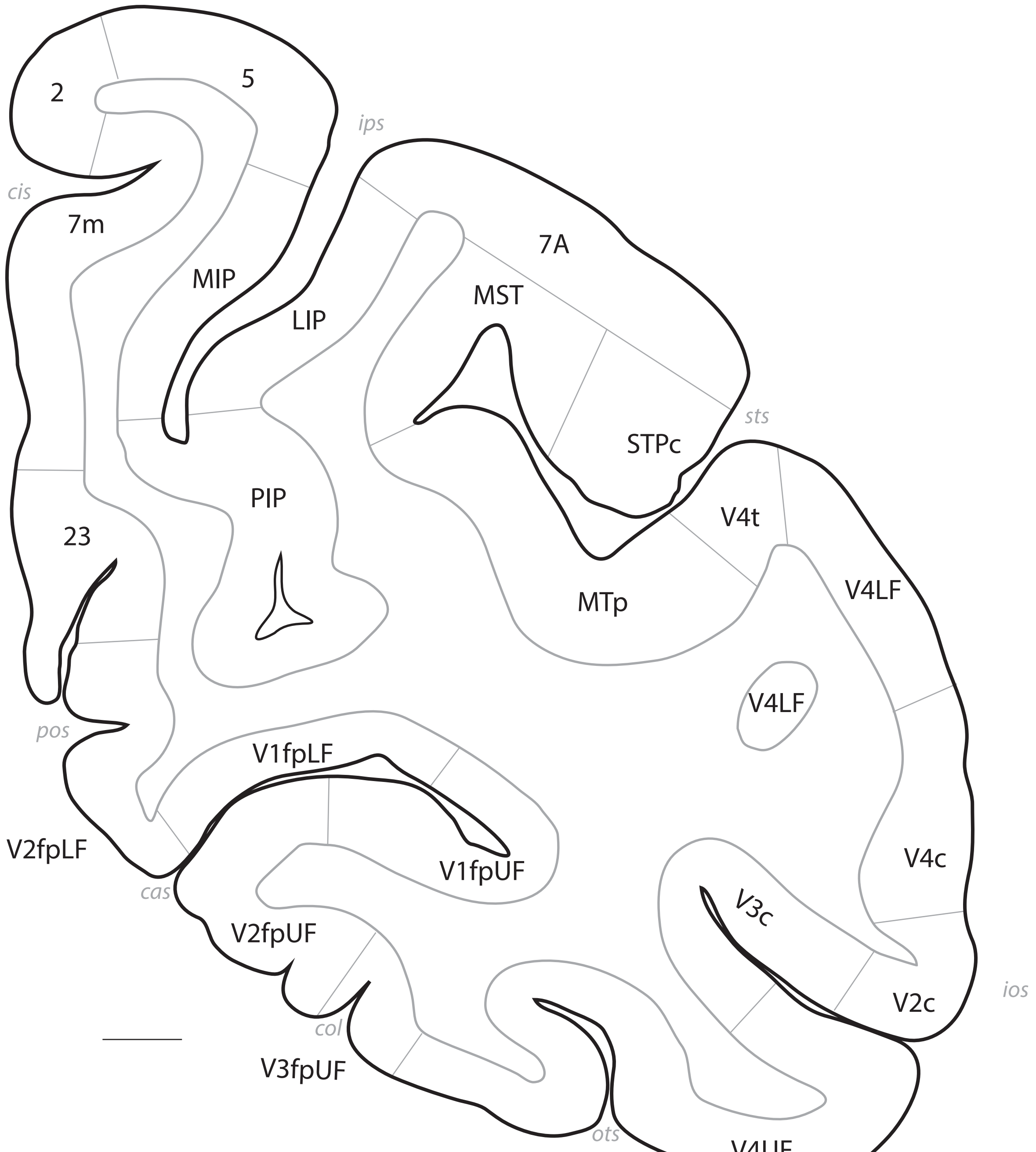
2
5
ips
cis
7m
MIP
LIP
7A
MST
PIP
sts
STPc
V4t
23
MTp
V4LF
V4LF
pos
V1fpLF
V2fpLF
V1fpUF
cas
V2fpUF
V3c
V4c
ios
V2c
col
V3fpUF
ots
V4UF

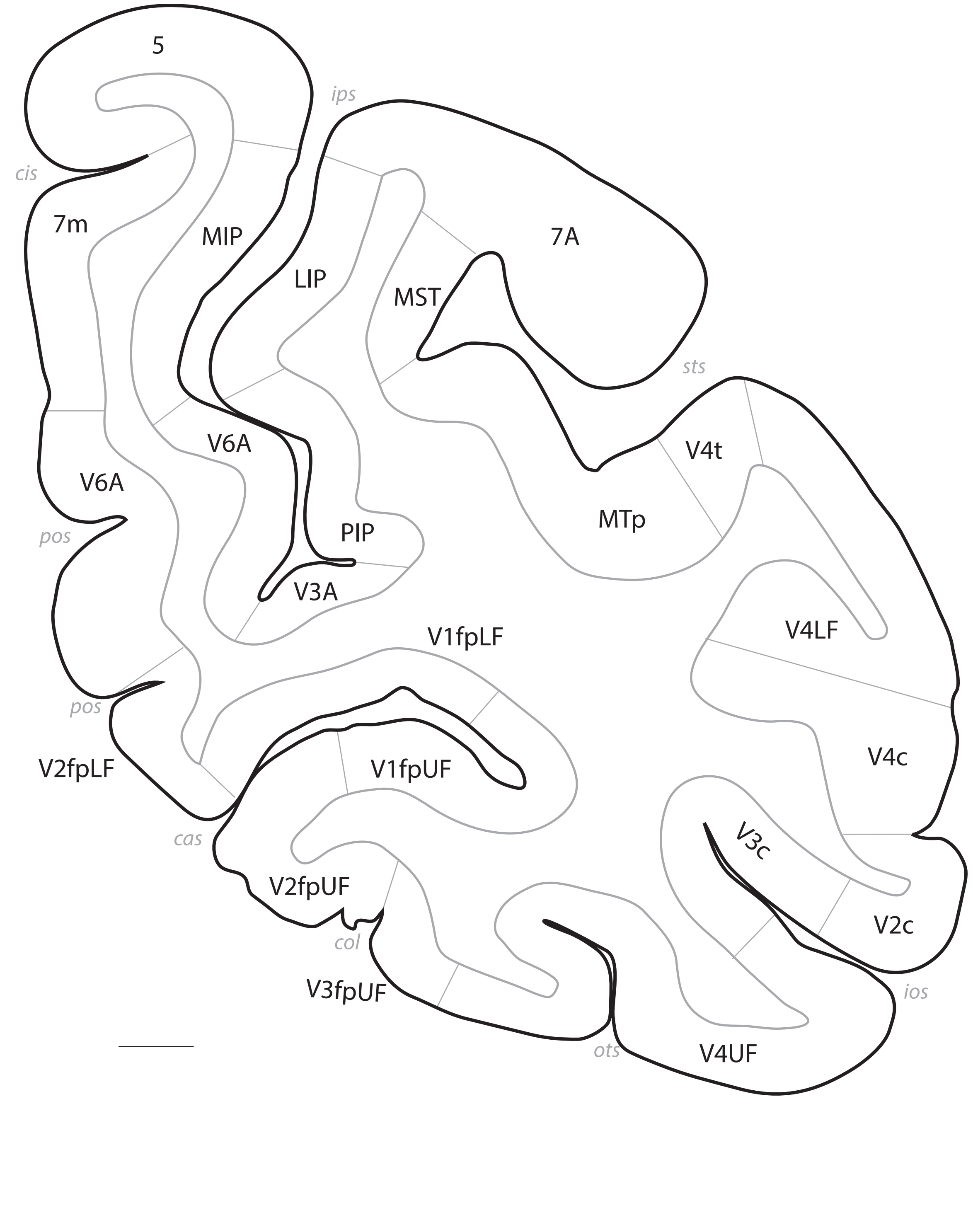
5
ips
cis
7m
MIP
LIP
MST
7A
sts
V6A
V6A
V4t
MTp
PIP
pos
V3A
V1fpLF
V4LF
pos
V2fpLF
V1fpUF
V4c
cas
V3c
V2fpUF
V2c
col
ios
V3fpUF
ots
V4UF

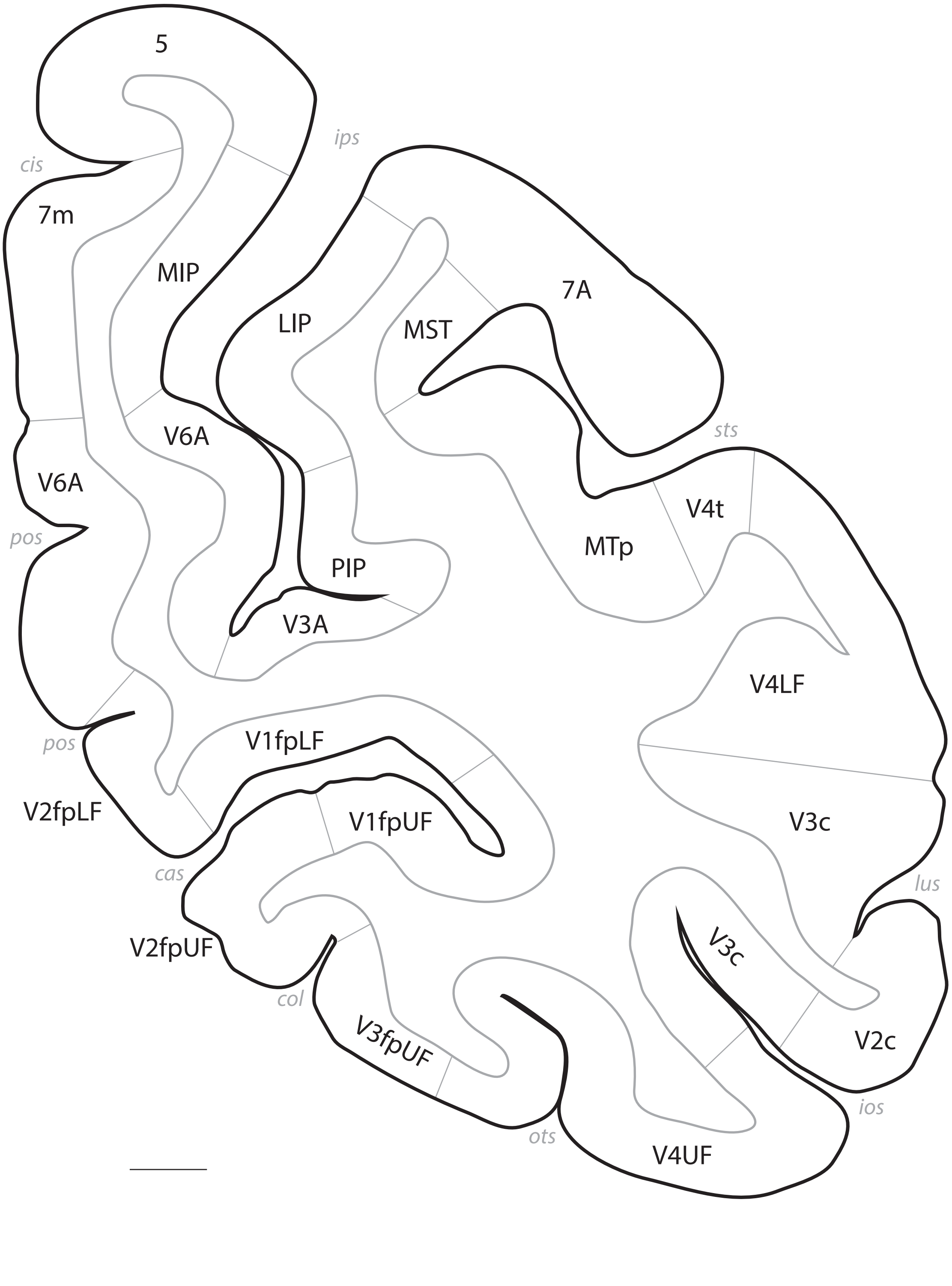

5
ips
cis
7m
MIP
LIP
MST
7A
V6A
V6A
sts
pos
V4t
MTp
PIP
V3A
V4LF
pos
V1fpLF
V2fpLF
V1fpUF
V3c
lus
cas
V2fpUF
V3c
col
V3fpUF
V2c
ios
ots
V4UF

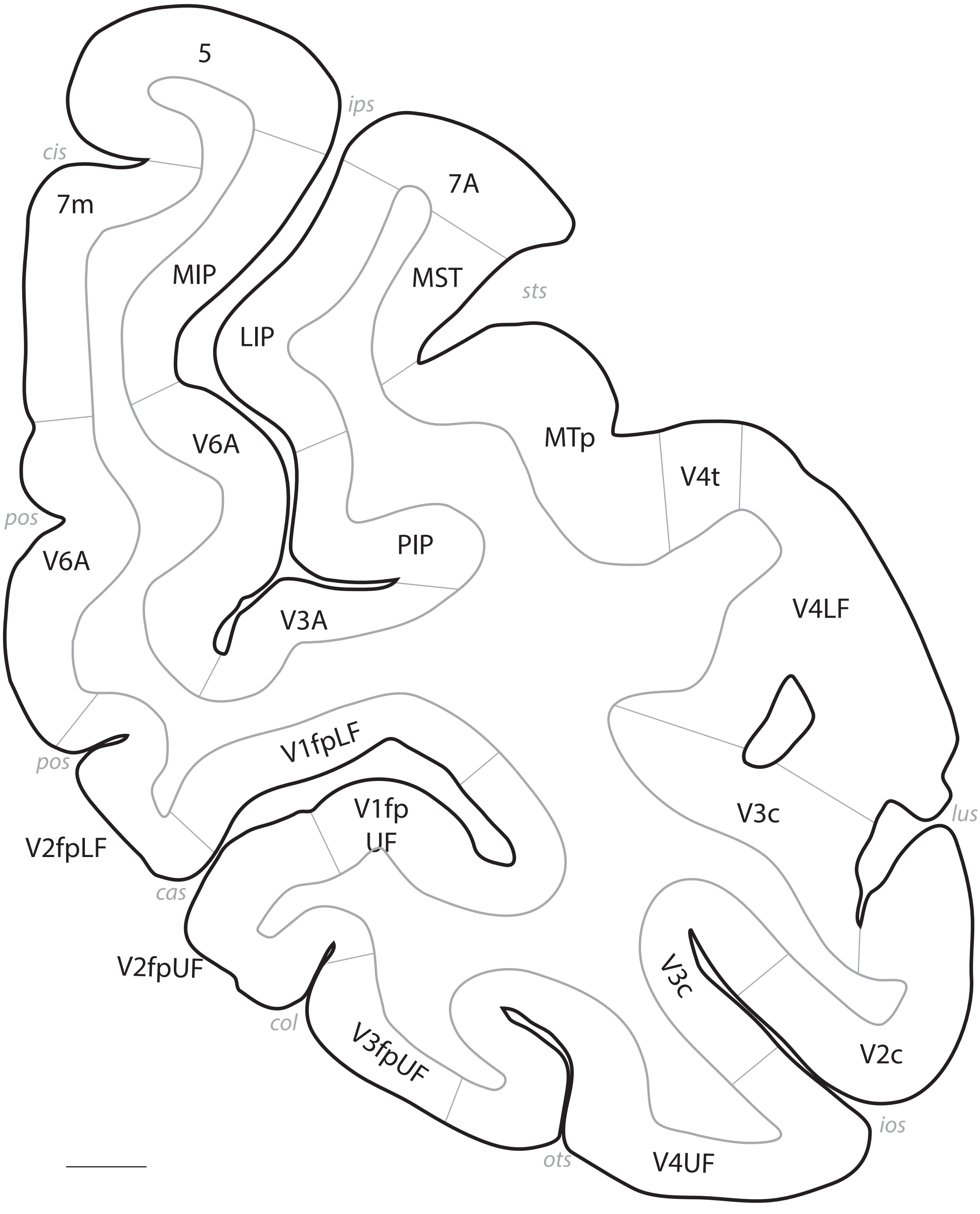
5
ips
cis
7A
7m
MIP
MST
sts
LIP
V6A
MTp
V4t
pos
V6A
PIP
V4LF
V3A
V1fpLF
pos
V1fp
UF
V3c
lus
V2fpLF
cas
V2fpUF
V3c
col
V3fpUF
V2c
ios
ots
V4UF

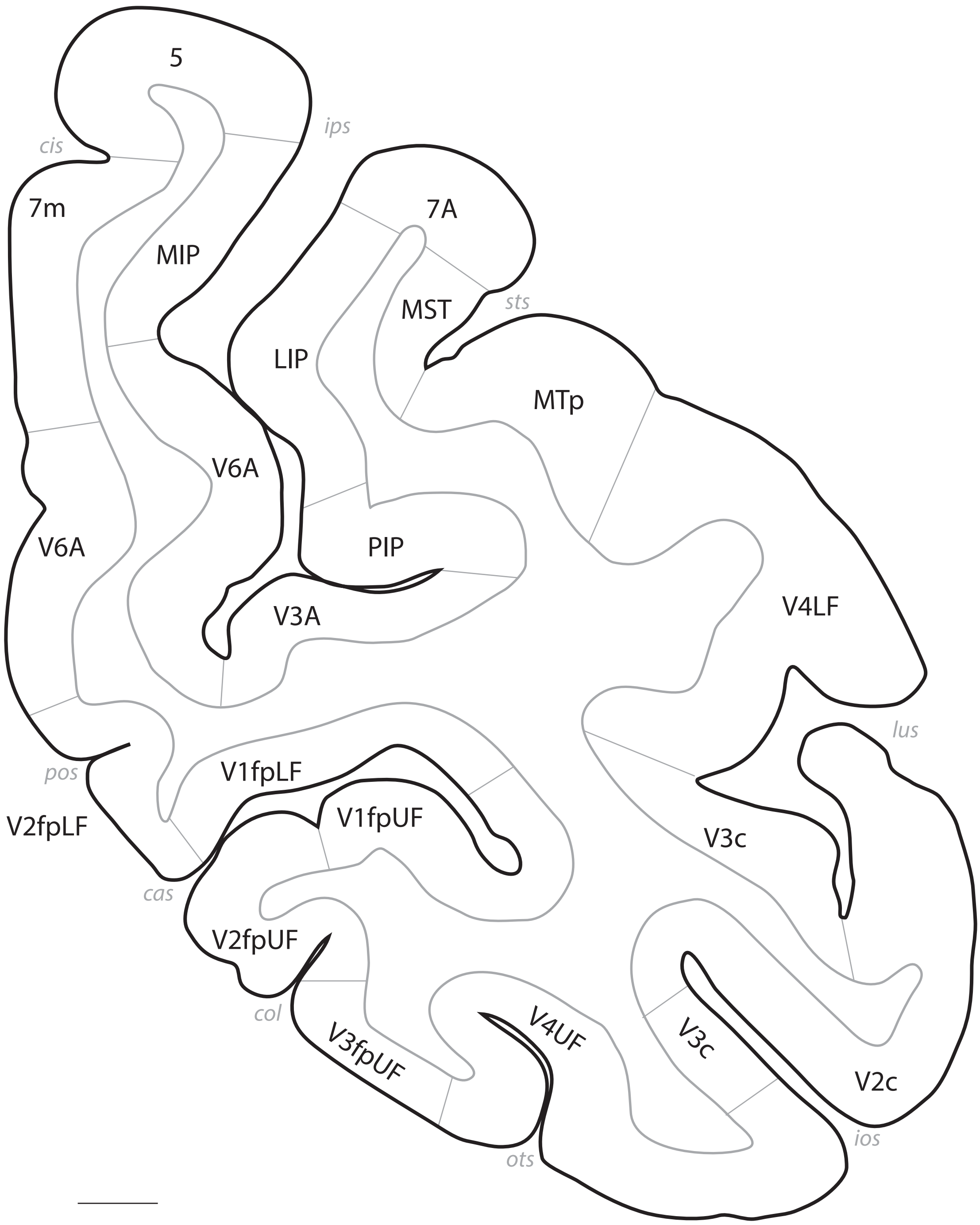
5
ips
cis
7m
MIP
7A
sts
MST
LIP
MTp
V6A
V6A
PIP
V3A
V4LF
lus
pos
V1fpLF
V2fpLF
V1fpUF
V3c
cas
V2fpUF
col
V4UF
V3fpUF
V3c
V2c
ios
ots

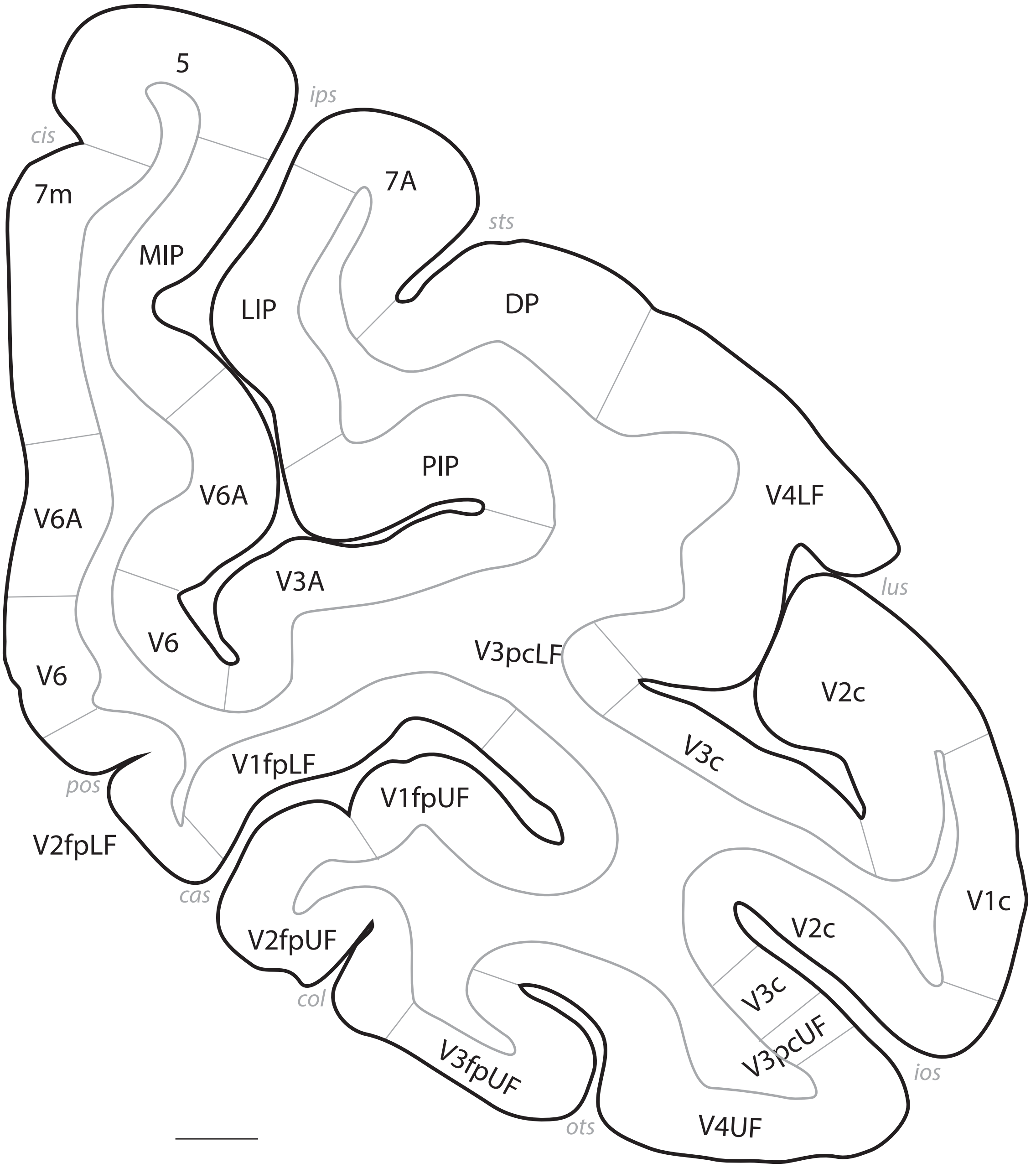
5
ips
cis
7A
7m
sts
MIP
LIP
DP
PIP
V6A
V4LF
V6A
V3A
lus
V6
V3pcLF
V2c
V6
V3c
V1fpLF
pos
V1fpUF
V2fpLF
V1c
cas
V2c
V2fpUF
V3c
col
V3pcUF
V3fpUF
ios
ots
V4UF

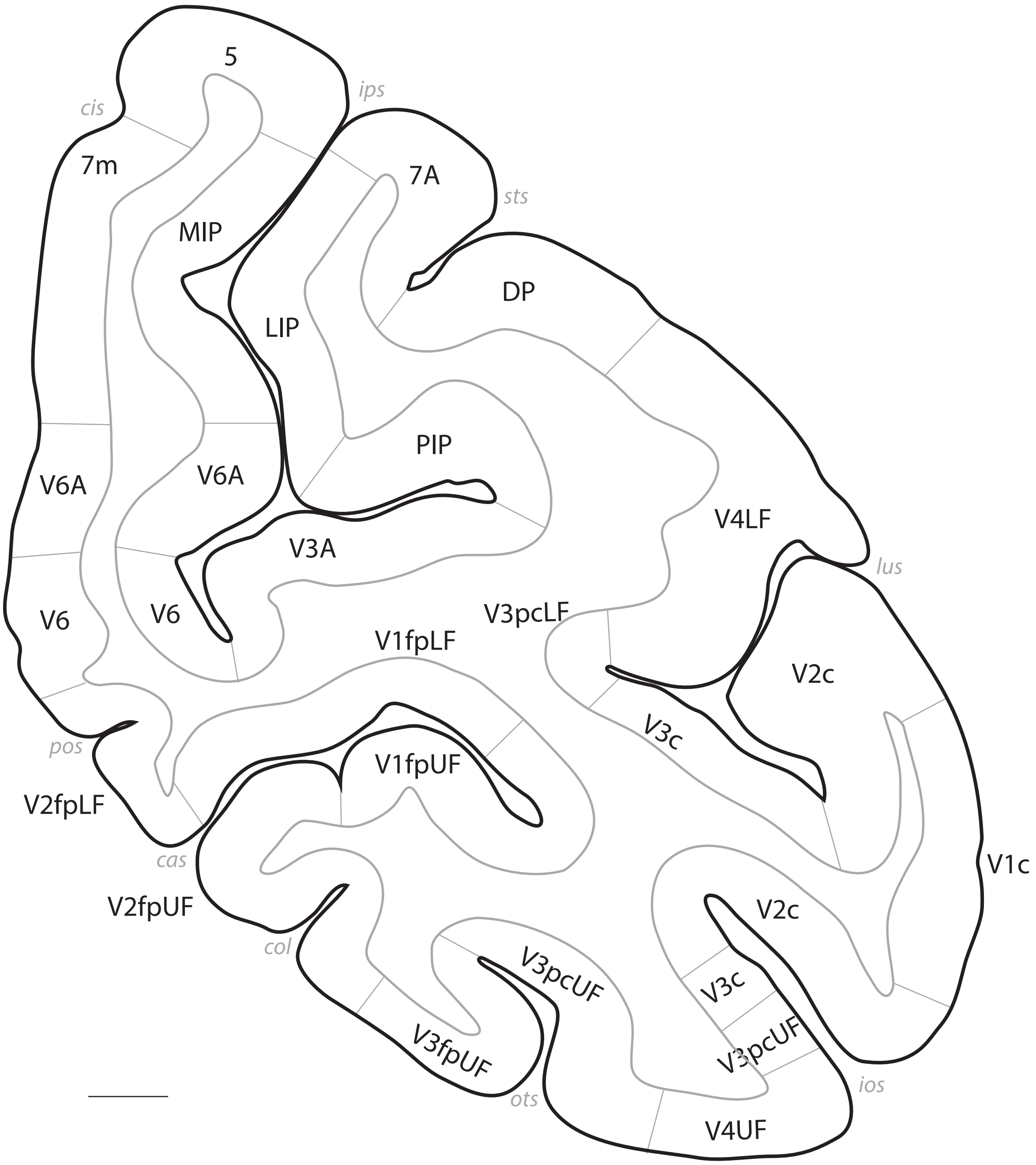
5
cis
ips
7m
7A
sts
MIP
DP
LIP
PIP
V6A
V6A
V4LF
V3A
lus
V6
V6
V3pcLF
V1fpLF
V2c
V3c
pos
V1fpUF
V2fpLF
cas
V1c
V2fpUF
V2c
col
V3pcUF
V3c
V3pcUF
V3fpUF
ios
ots
V4UF

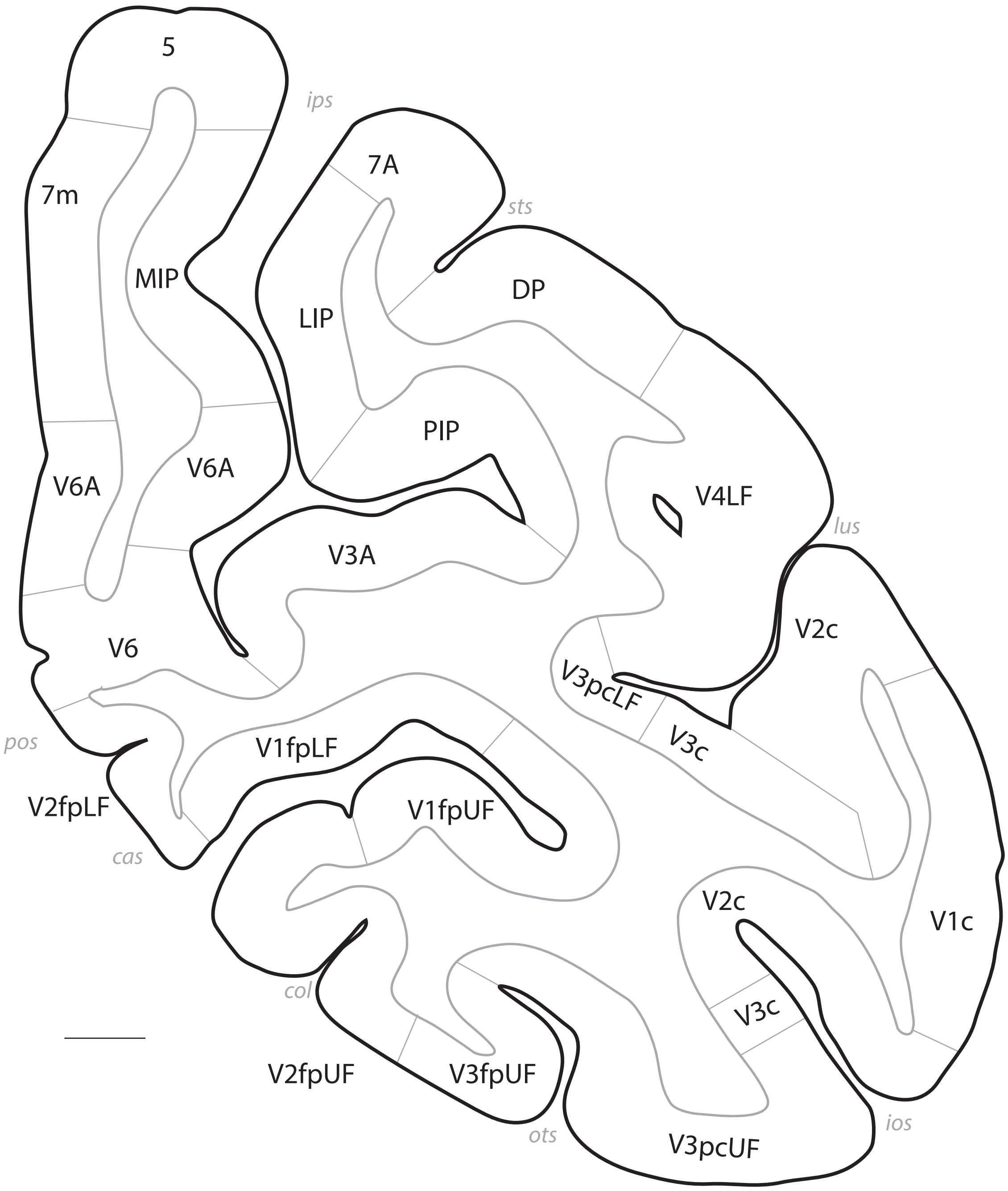
5
ips
7m
MIP
7A
sts
LIP
DP
PIP
V6A
V6A
V4LF
lus
V3A
V6
V2c
V3pcLF
V3c
pos
V1fpLF
V2fpLF
V1fpUF
cas
V2c
V1c
col
V3c
V2fpUF
V3fpUF
ots
V3pcUF
ios

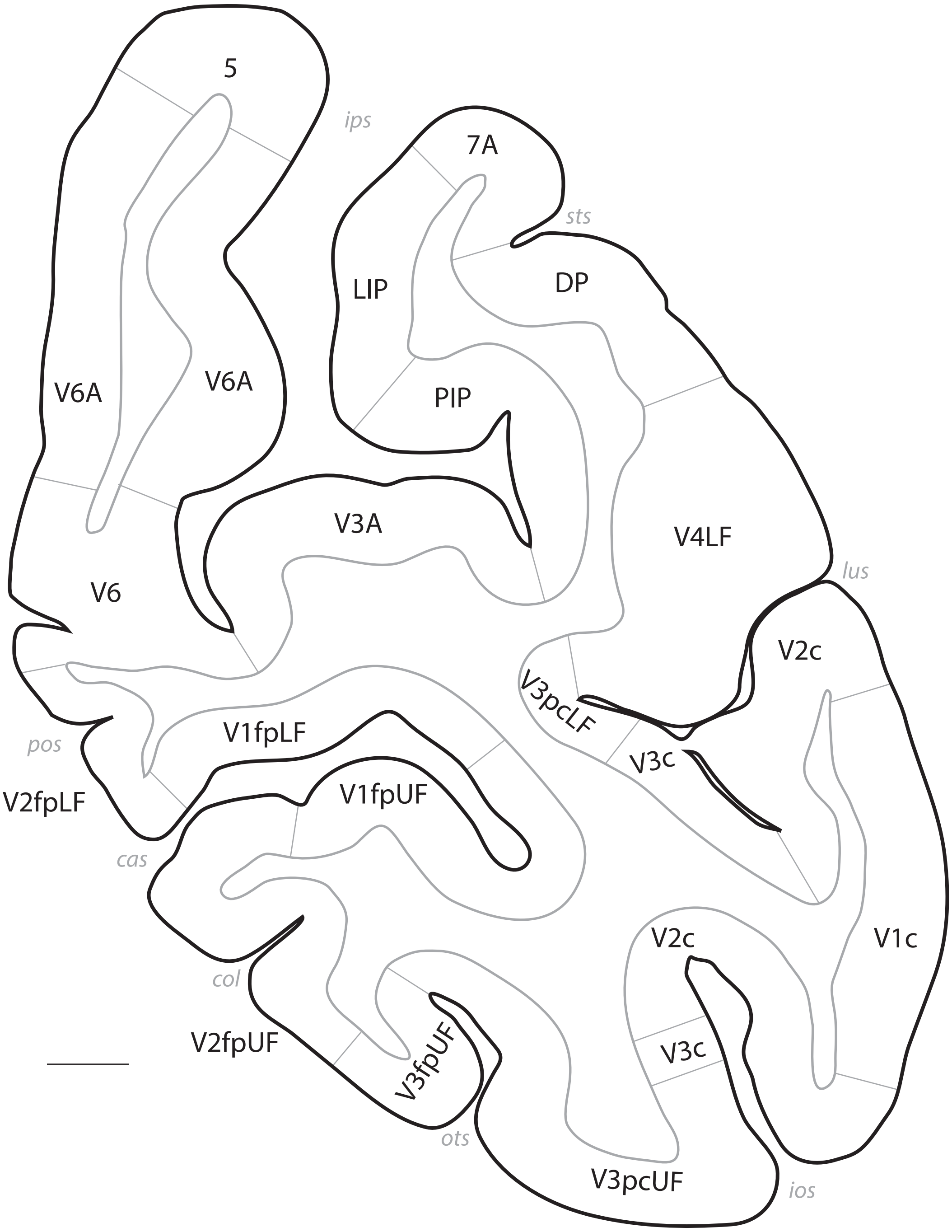
5
ips
7A
sts
LIP
DP
V6A
V6A
PIP
V3A
V4LF
lus
V6
V2c
V3pcLF
V1fpLF
V3c
pos
V2fpLF
V1fpUF
cas
V2c
V1c
col
V2fpUF
V3fpUF
V3c
ots
V3pcUF
ios

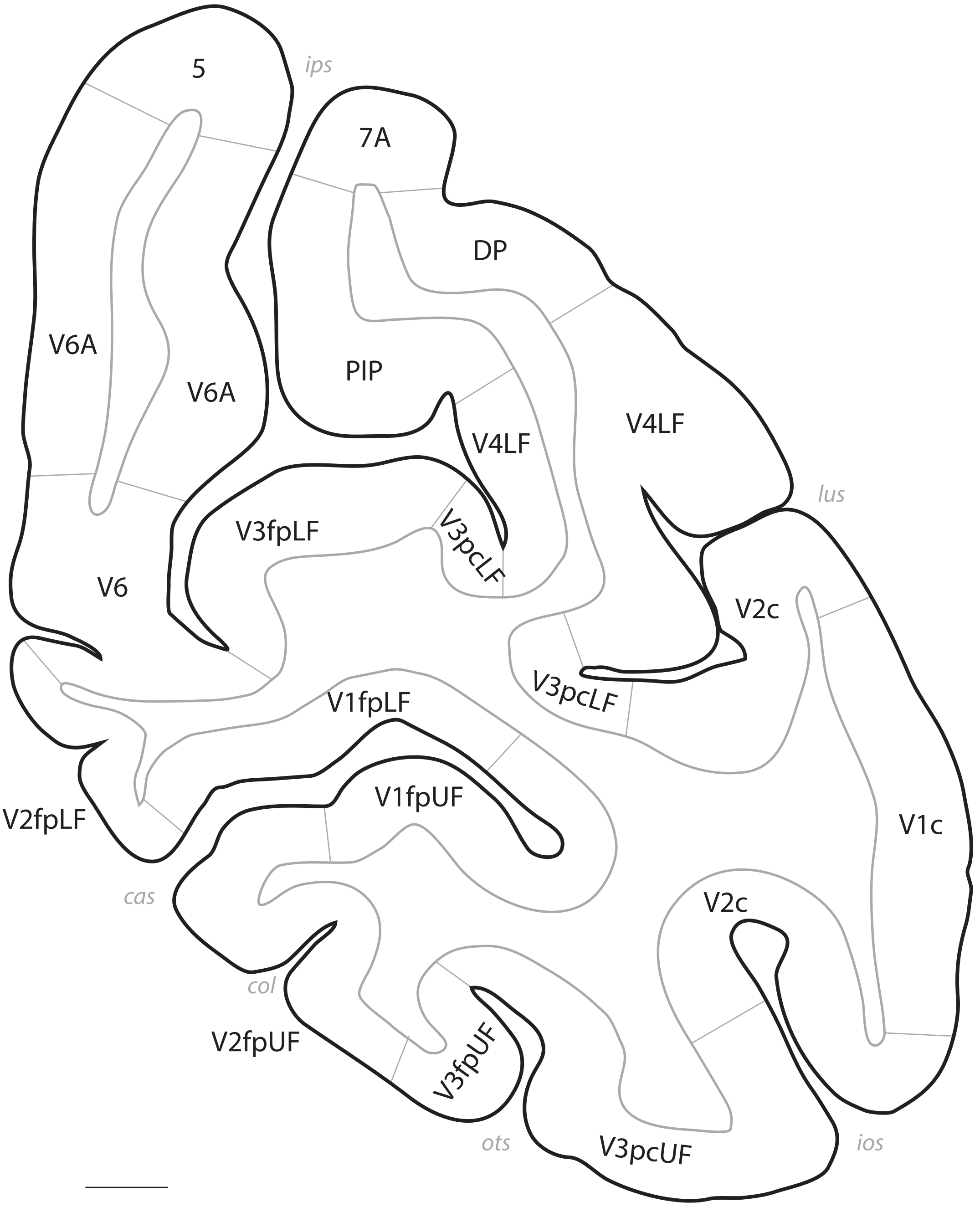
5
ips
7A
DP
PIP
V6A
V6A
V4LF
V4LF
lus
V3fpLF
V3pcLF
V6
V2c
V3pcLF
V1fpLF
V1fpUF
V2fpLF
V1c
cas
V2c
col
V2fpUF
V3fpUF
ots
V3pcUF
ios

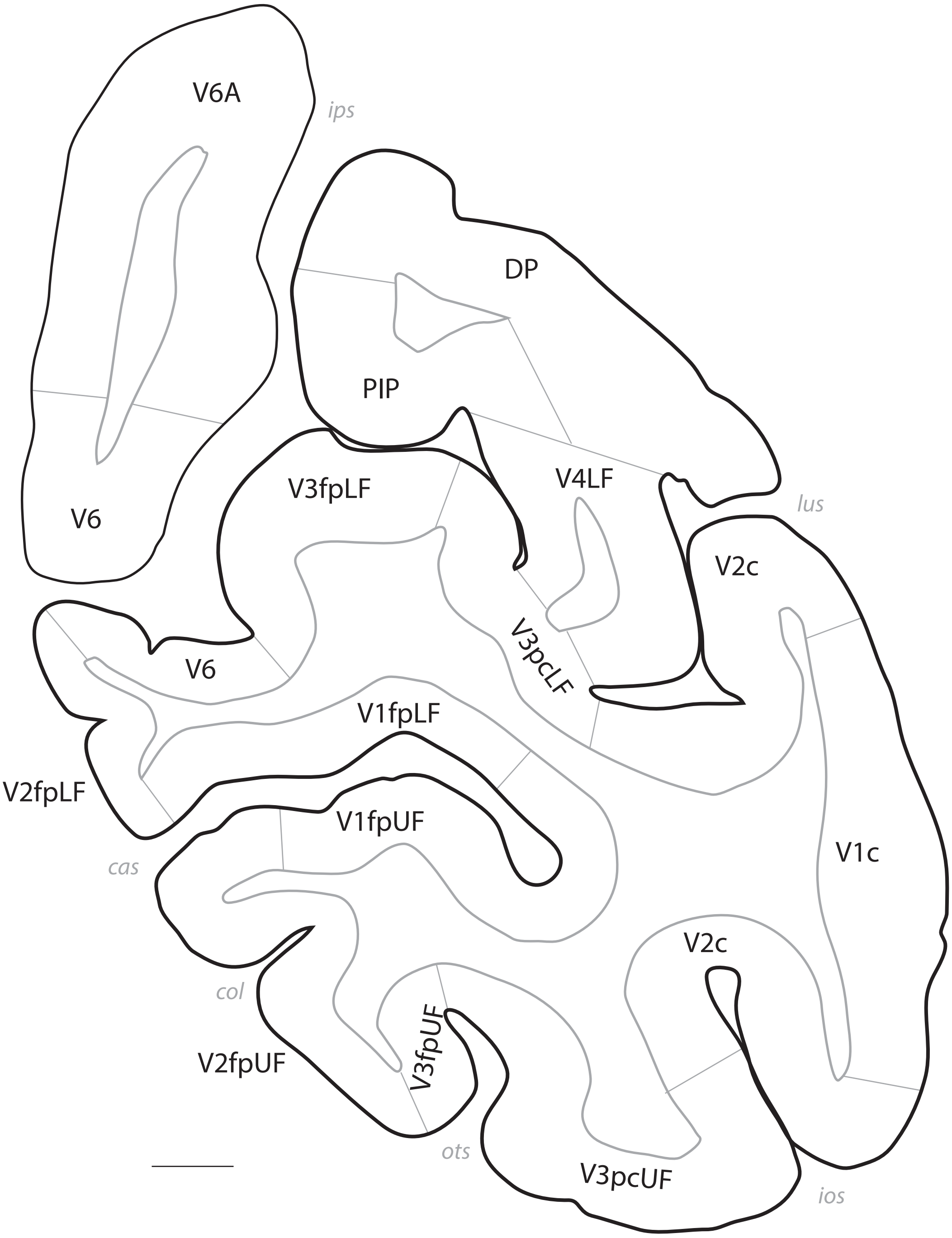
V6A
ips
DP
PIP
V6
V3fpLF
V4LF
lus
V2c
V6
V3pcLF
V1fpLF
V2fpLF
V1fpUF
cas
V1c
V2c
col
V3fpUF
V2fpUF
ots
V3pcUF
ios

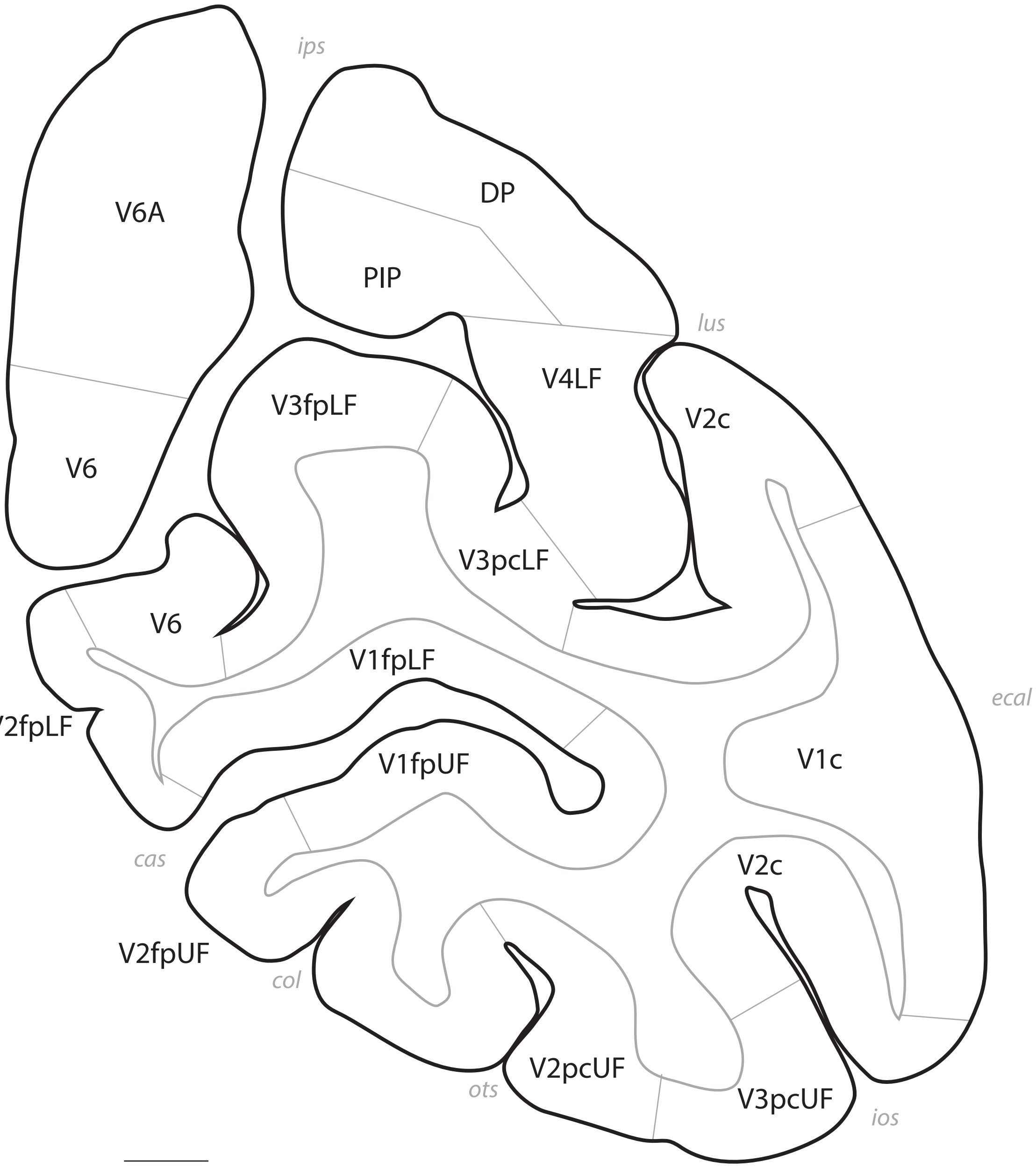
ips
V6A
DP
PIP
lus
V4LF
V3fpLF
V2c
V6
V3pcLF
V6
V1fpLF
ecal
V2fpLF
V1fpUF
V1c
V2c
cas
V2fpUF
col
V2pcUF
ots
V3pcUF
ios

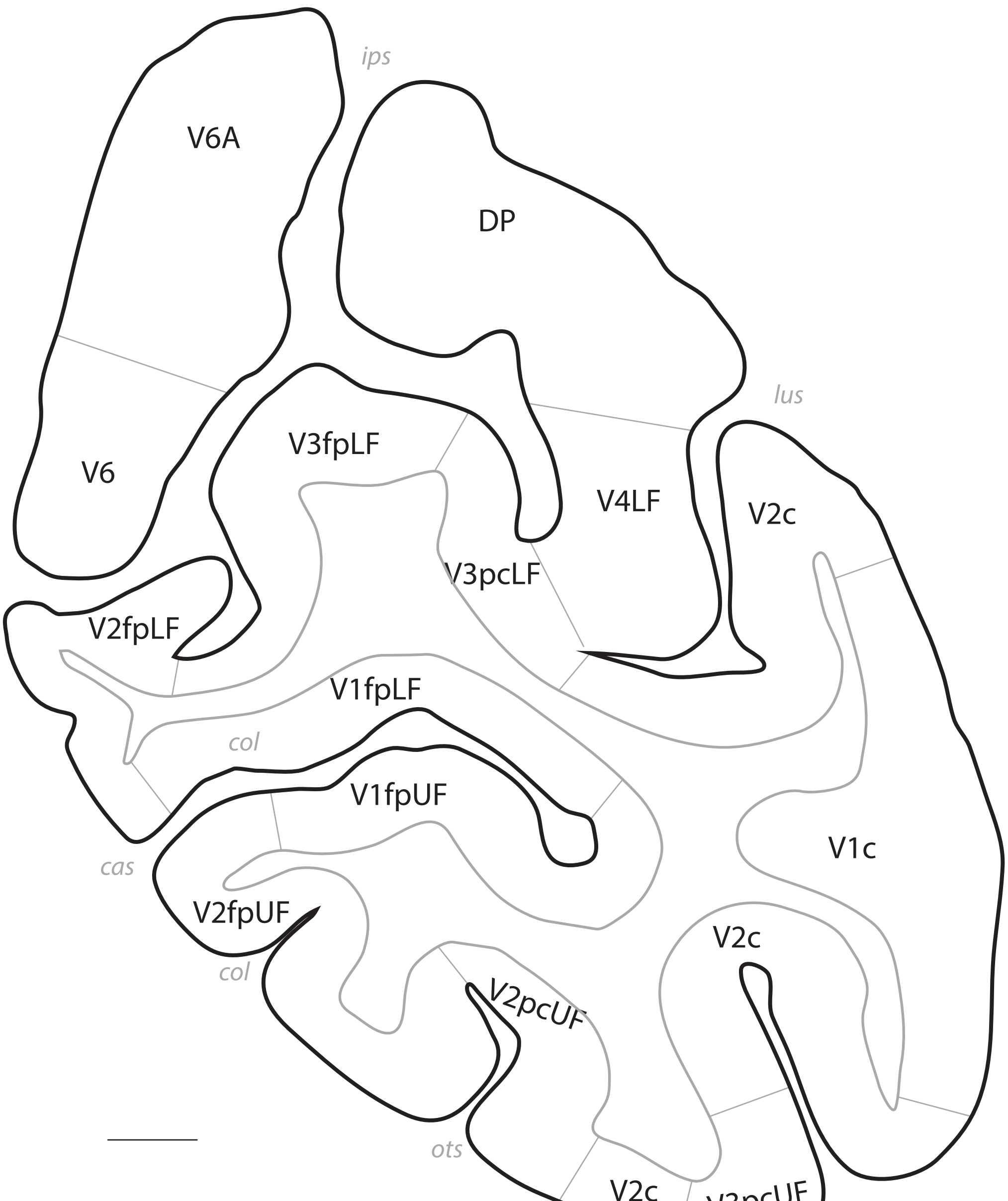
ips
V6A
DP
lus
V6
V3fpLF
V4LF
V2c
V3pcLF
V2fpLF
V1fpLF
col
V1fpUF
ecal
V1c
cas
V2fpUF
V2c
col
V2pcUF
ots
V2c
V3pcUF
ios

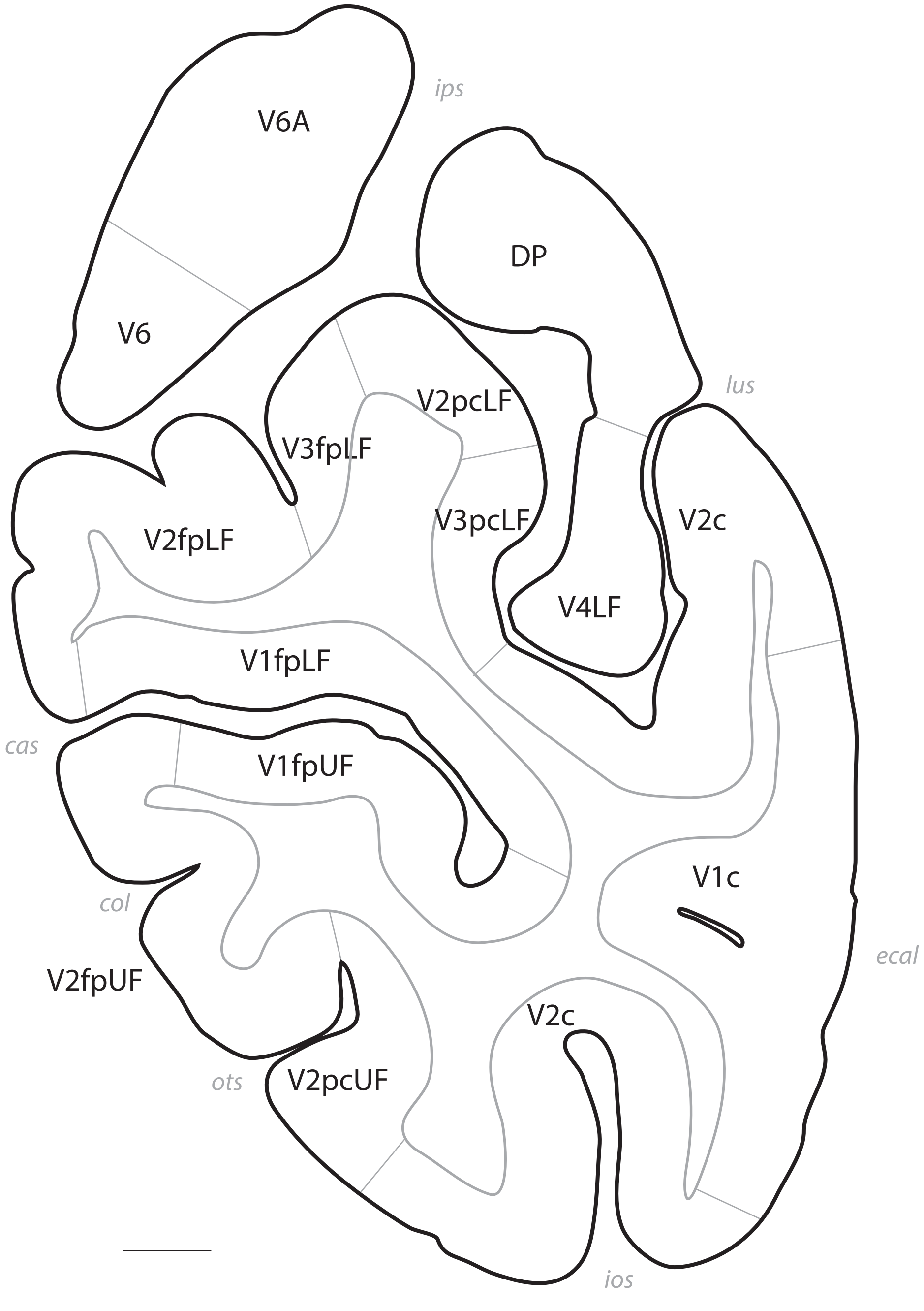
ips
V6A
DP
V6
lus
V2pcLF
V3fpLF
V3pcLF
V2c
V2fpLF
V4LF
V1fpLF
cas
V1fpUF
V1c
col
ecal
V2fpUF
V2c
ots
V2pcUF
ios

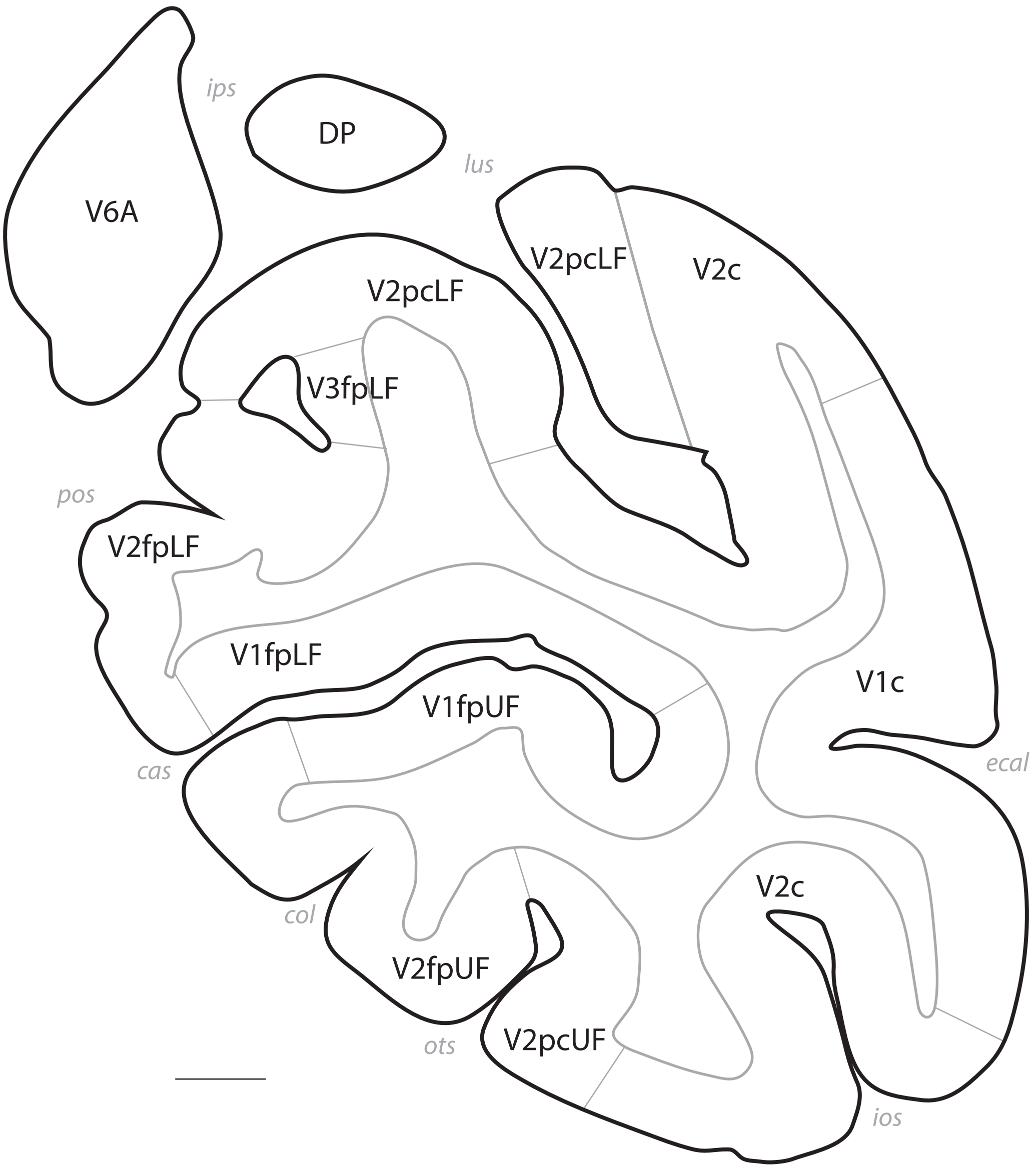
ips
DP
lus
V6A
V2pcLF
V2c
V2pcLF
V3fpLF
pos
V2fpLF
V1fpLF
V1fpUF
V1c
ecal
cas
V2c
col
V2fpUF
ots
V2pcUF
ios

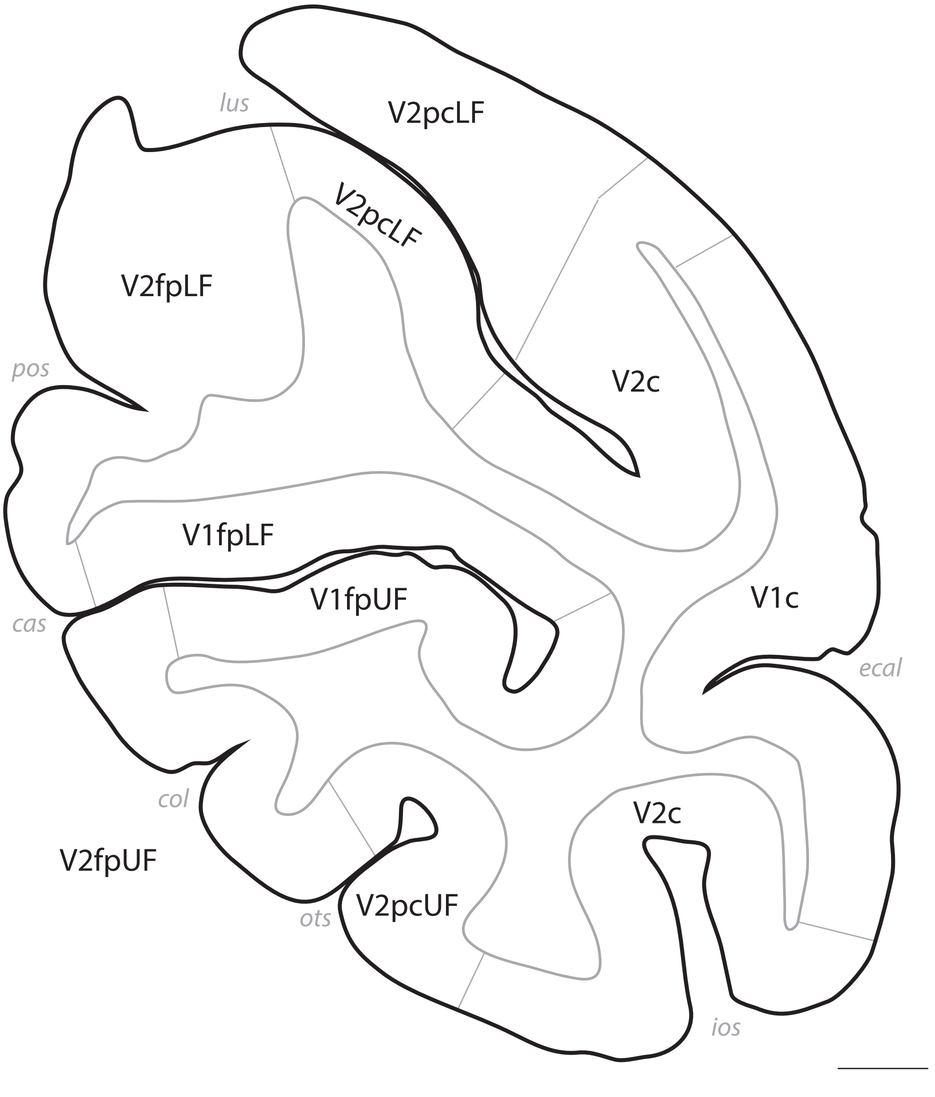
lus
V2pcLF
V2pcLF
V2fpLF
pos
V2c
V1fpLF
V1fpUF
V1c
cas
ecal
col
V2c
V2fpUF
ots
V2pcUF
ios

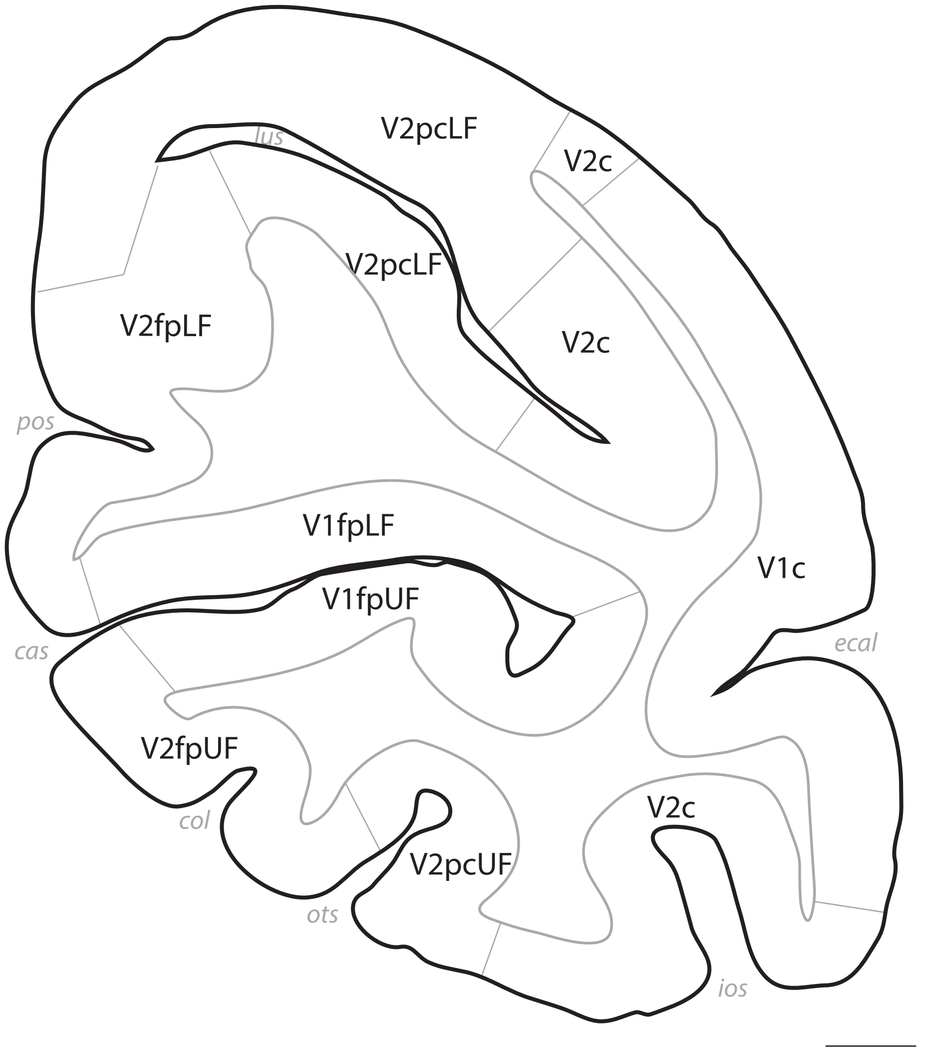

V2pcLF
lus
V2c
V2pcLF
V2fpLF
V2c
pos
V1fpLF
V1c
V1fpUF
cas
ecal
V2fpUF
V2c
col
V2pcUF
ots
ios

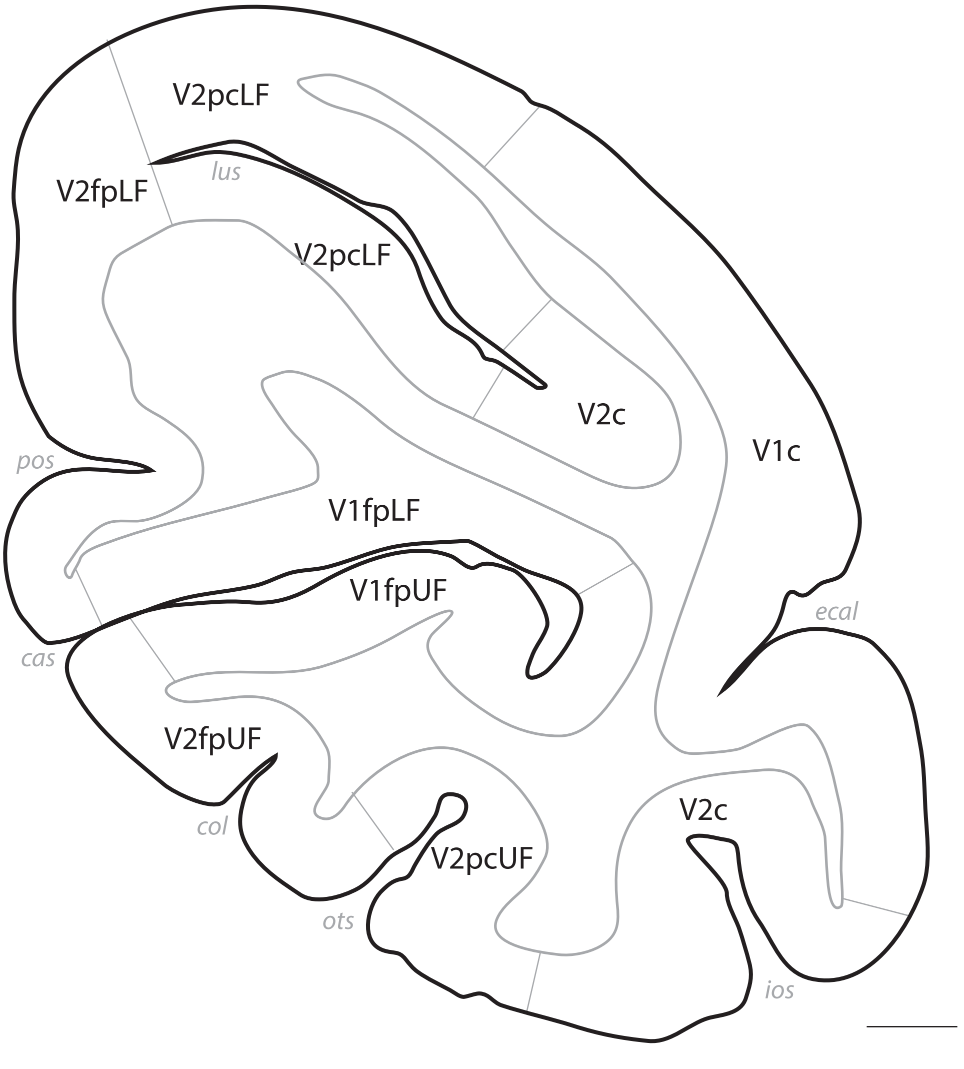
V2pcLF
V2fpLF
lus
V2pcLF
V2c
V1c
pos
V1fpLF
V1fpUF
ecal
cas
V2fpUF
col
V2c
V2pcUF
ots
ios

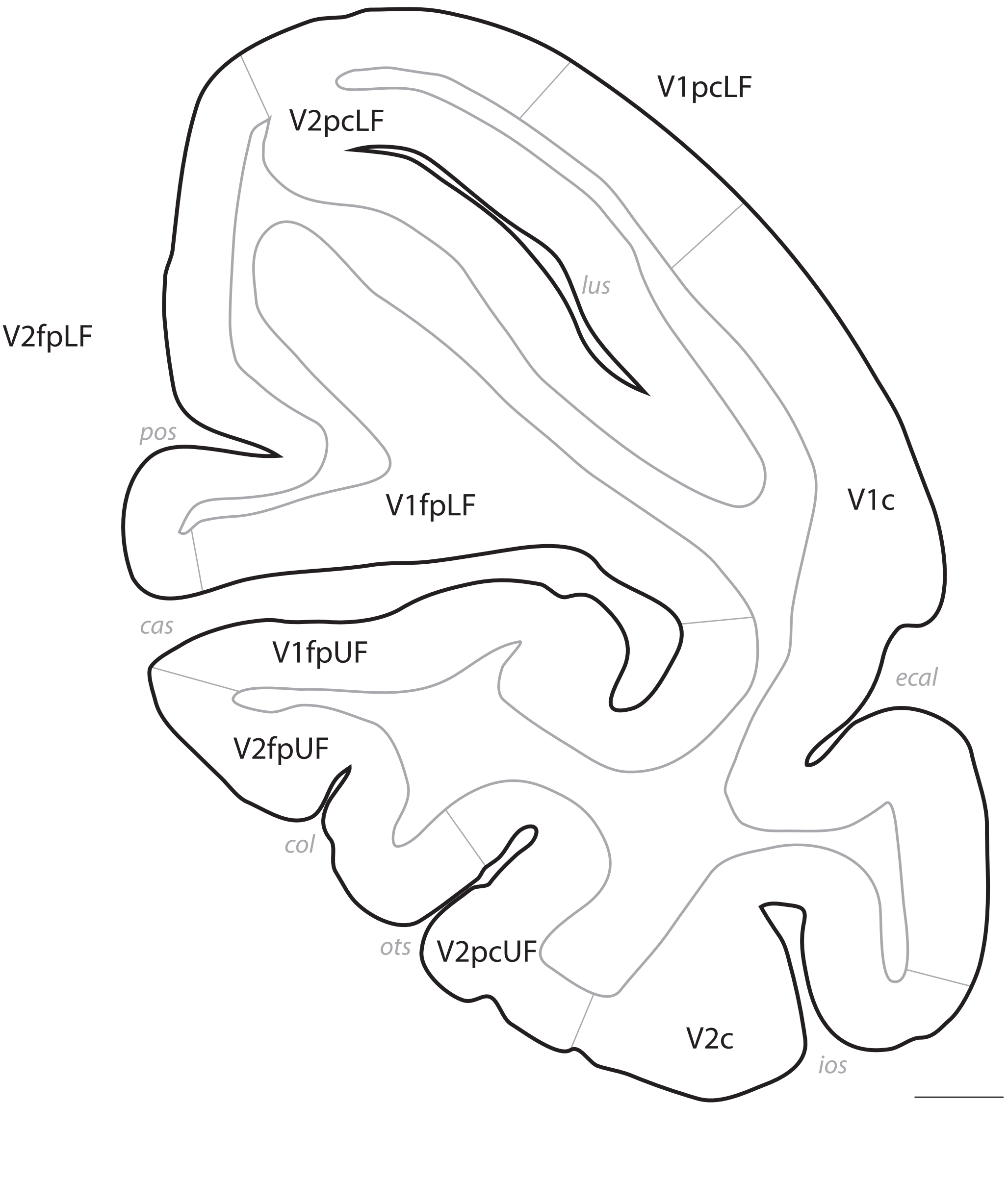
V1pcLF
V2pcLF
lus
V2fpLF
pos
V1fpLF
V1c
cas
V1fpUF
ecal
V2fpUF
col
ots
V2pcUF
V2c
ios

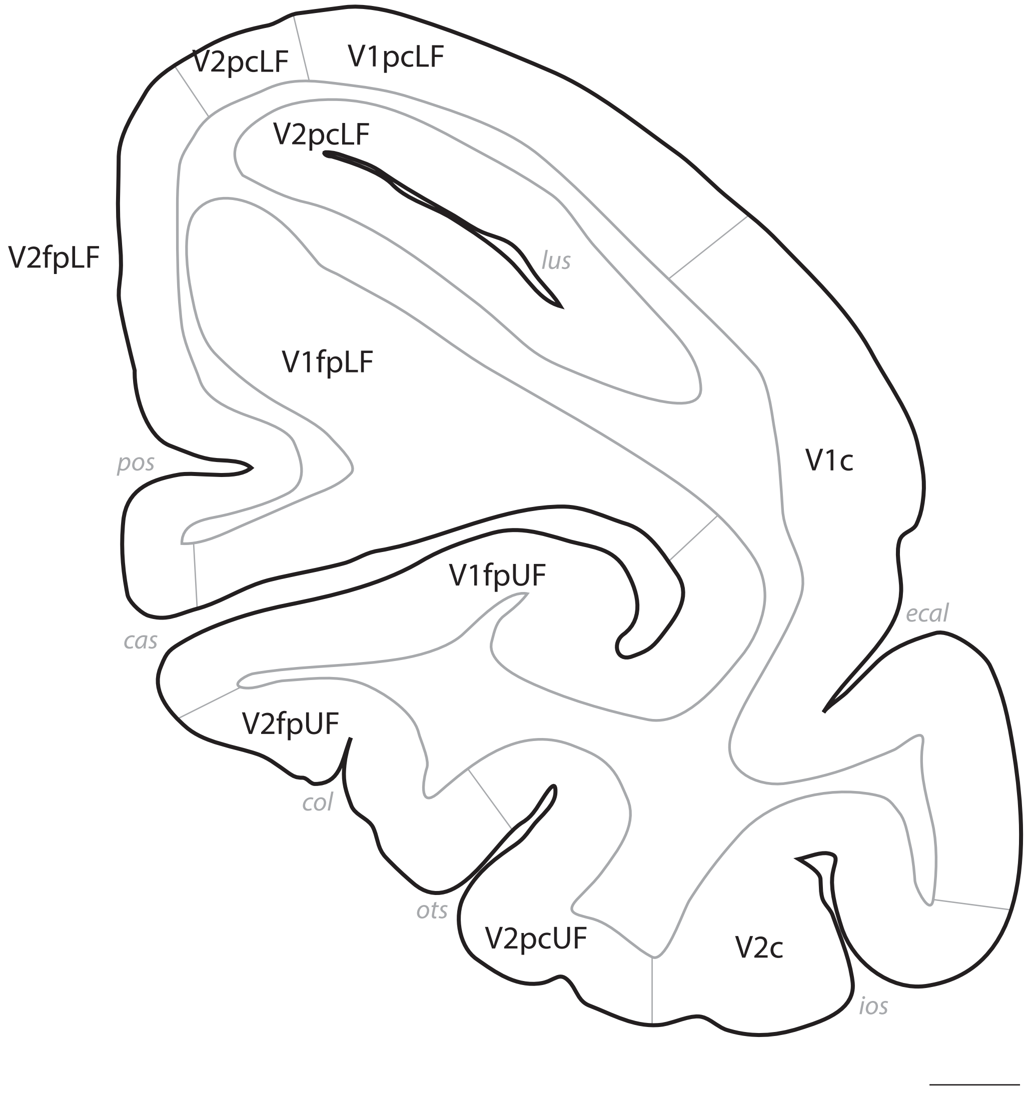
V2pcLF
V1pcLF
V2pcLF
lus
V2fpLF
V1fpLF
V1c
pos
V1fpUF
ecal
cas
V2fpUF
col
ots
V2pcUF
V2c
ios

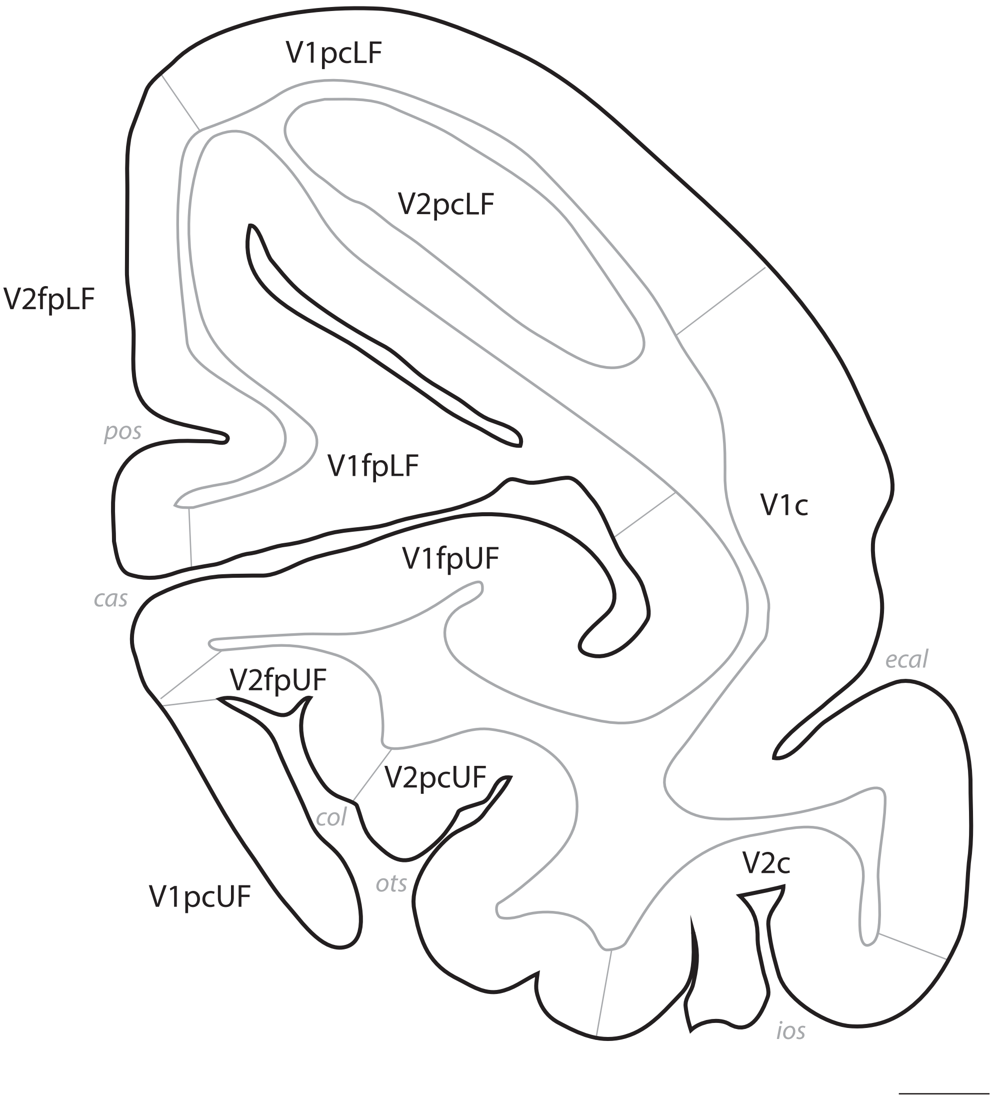
V1pcLF
V2pcLF
V2fpLF
pos
V1fpLF
V1c
V1fpUF
cas
ecal
V2fpUF
V2pcUF
col
ots
V1pcUF
V2c
ios

p. 111

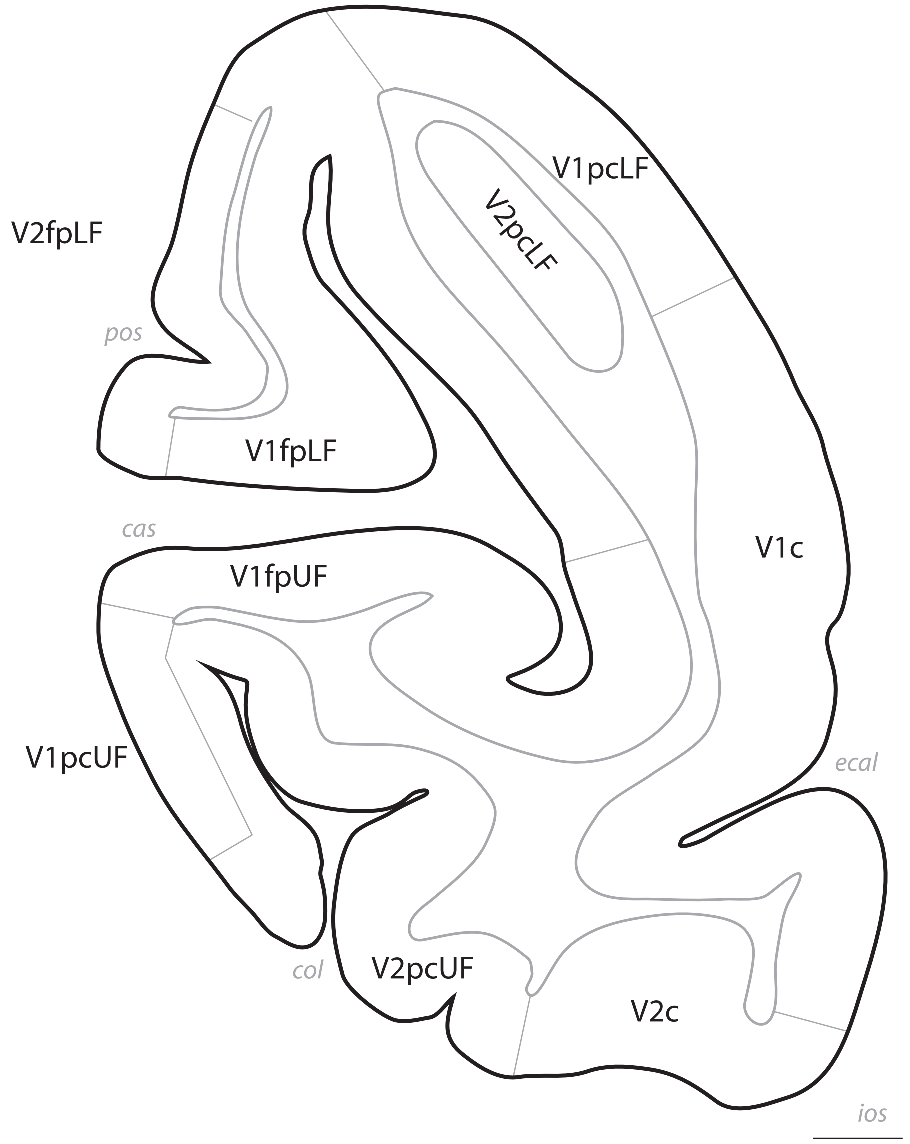
V2fpLF
V1pcLF
V2pcLF
pos
V1fpLF
cas
V1c
V1fpUF
V1pcUF
ecal
col
V2pcUF
V2c
ios

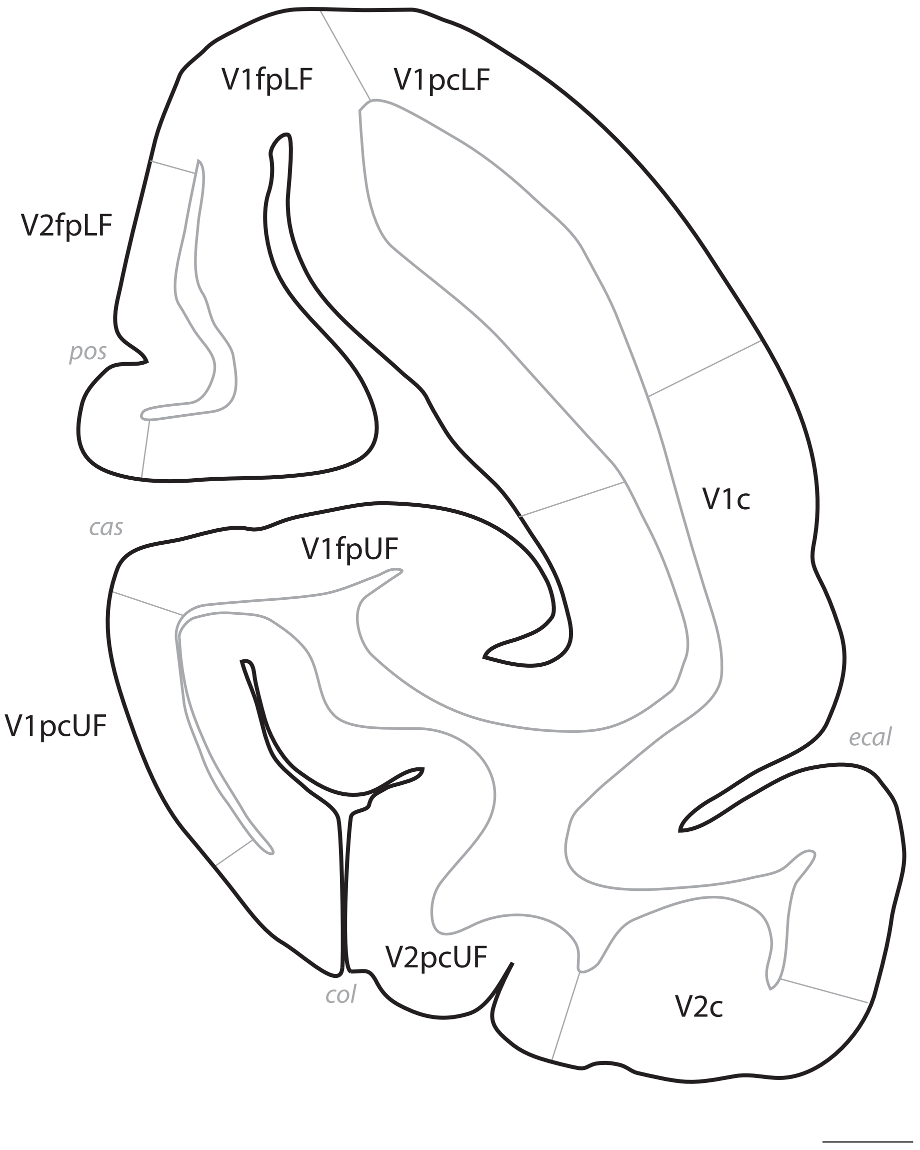
V1fpLF
V1pcLF
V2fpLF
pos
V1c
cas
V1fpUF
V1pcUF
ecal
V2pcUF
col
V2c

# p. 113

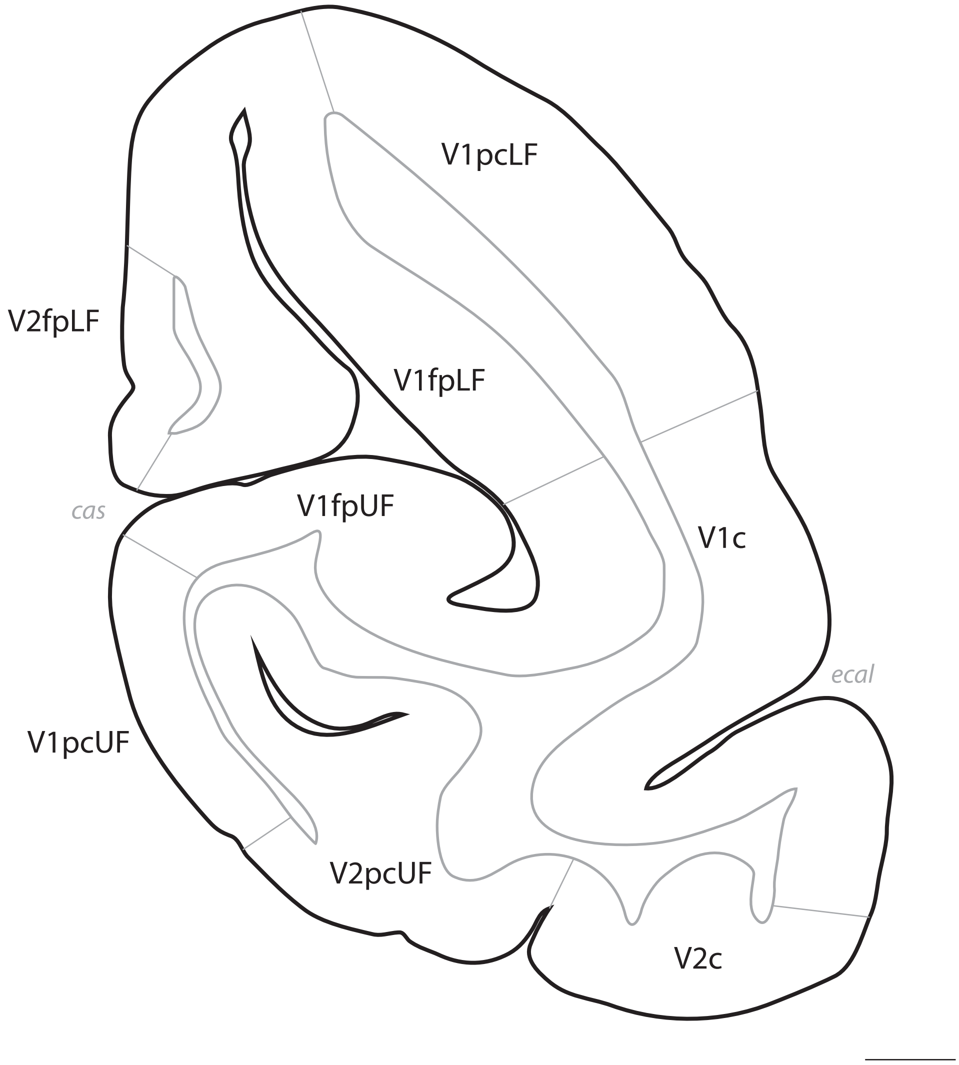

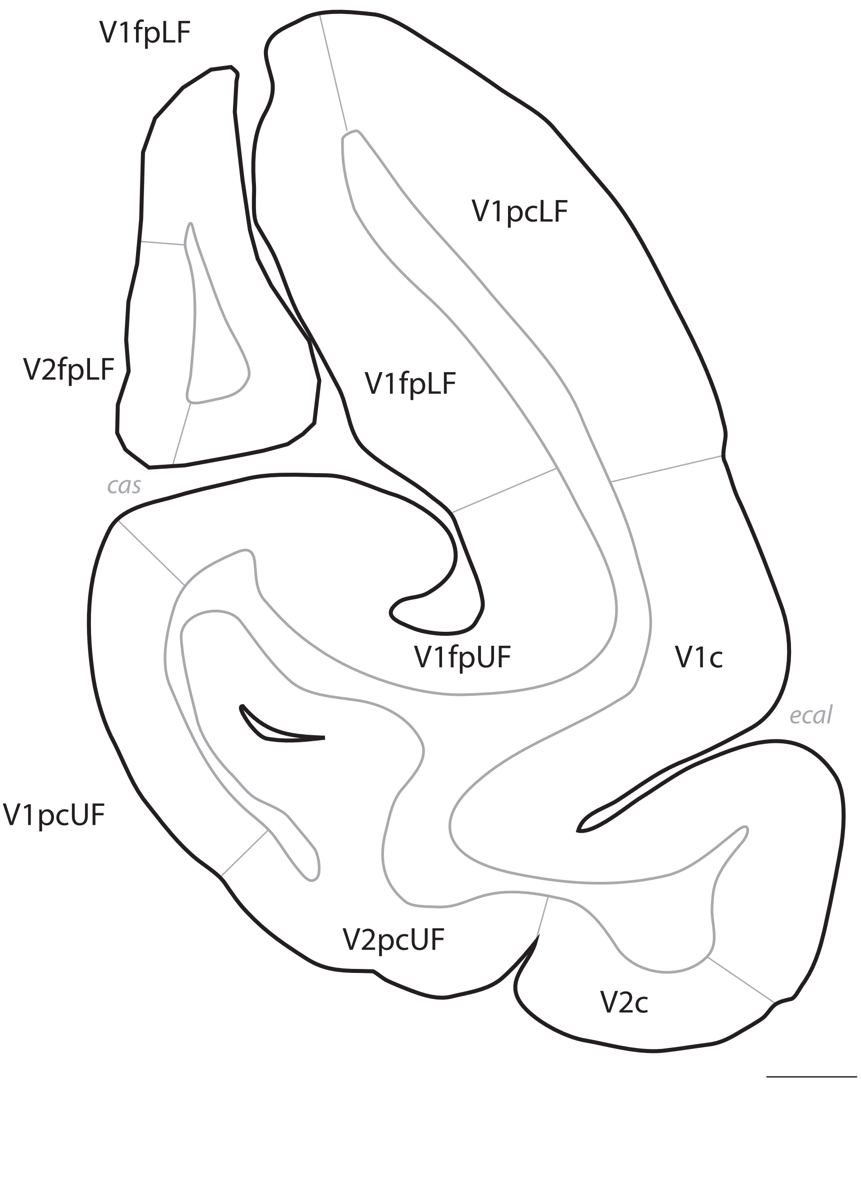
V1fpLF
V1pcLF
V2fpLF
V1fpLF
cas
V1fpUF
V1c
ecal
V1pcUF
V2pcUF
V2c

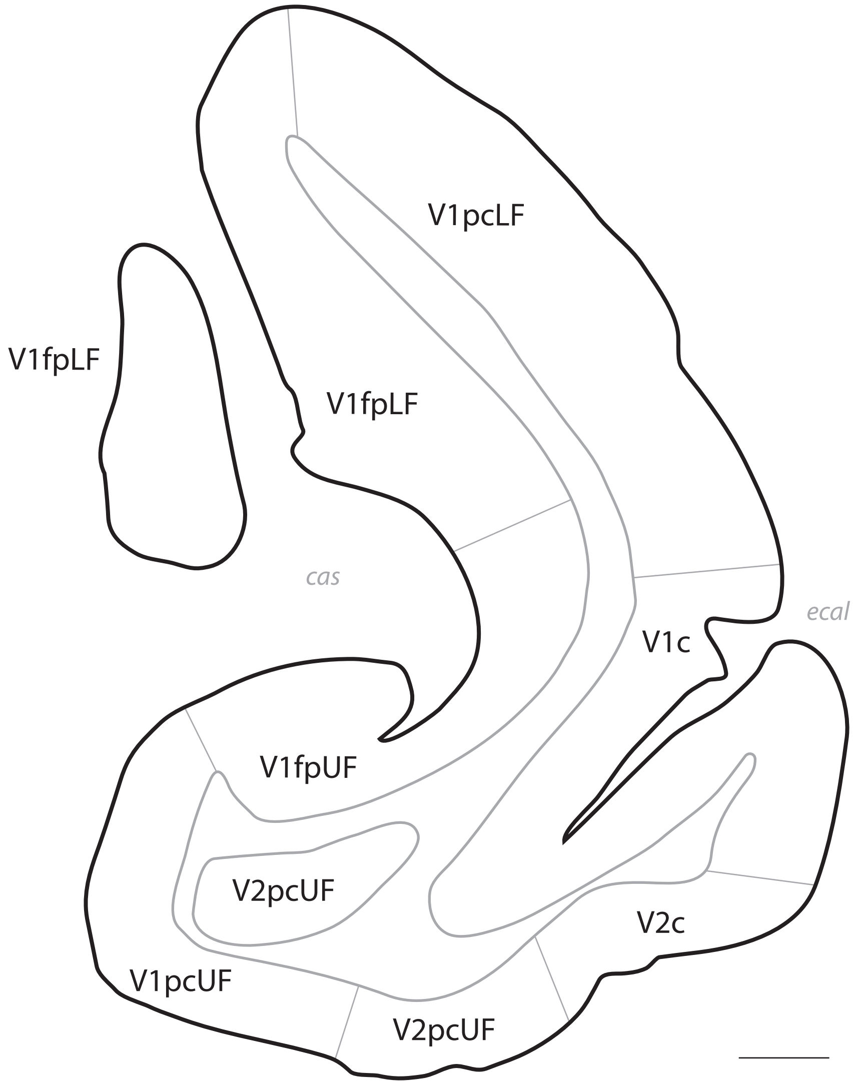
V1pcLF
V1fpLF
V1fpLF
cas
ecal
V1c
V1fpUF
V2pcUF
V2c
V1pcUF
V2pcUF

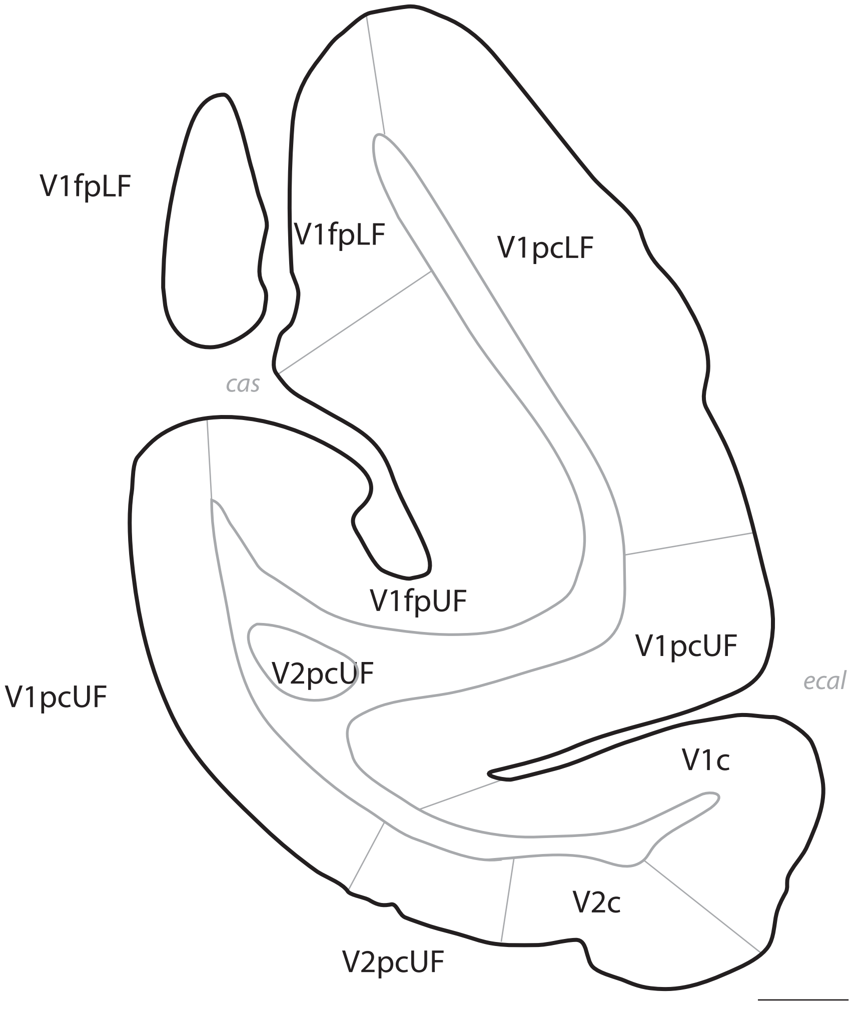
V1fpLF
V1fpLF
V1pcLF
cas
V1fpUF
V1pcUF
V2pcUF
V1pcUF
ecal
V1c
V2c
V2pcUF

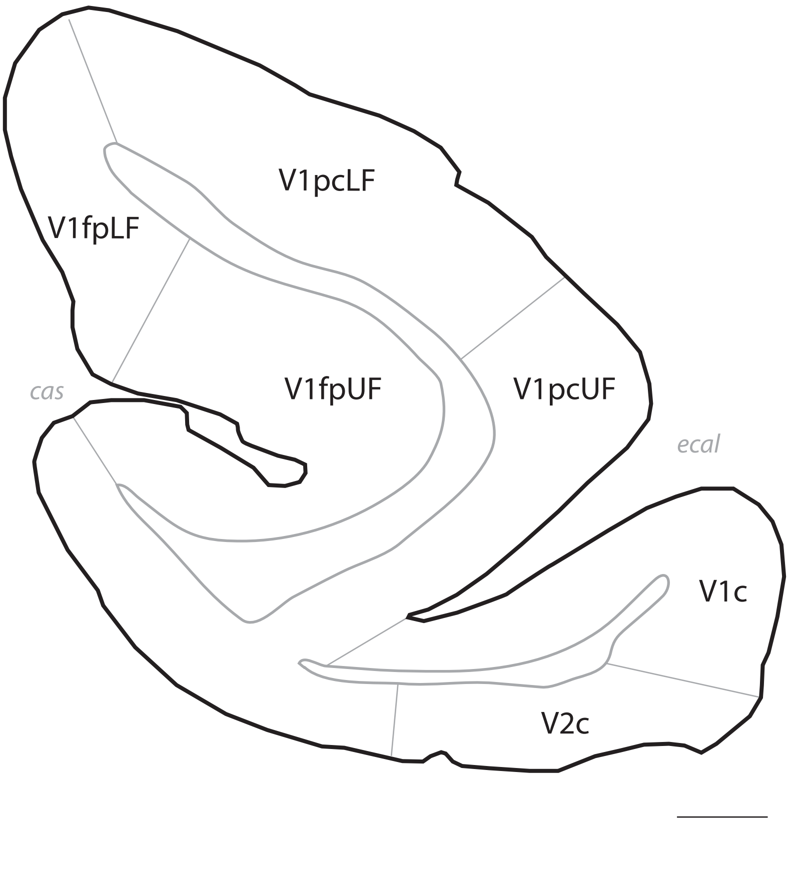
V1pcLF
V1fpLF
cas
V1fpUF
V1pcUF
ecal
V1c
V2c

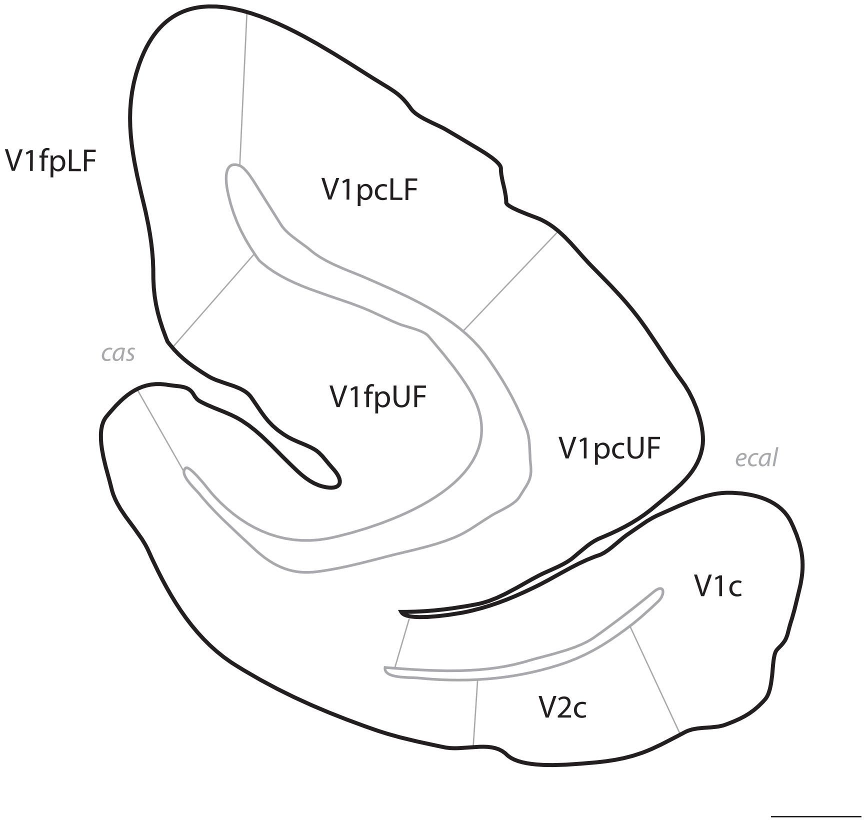
V1fpLF
V1pcLF
cas
V1fpUF
V1pcUF
ecal
V1c
V2c

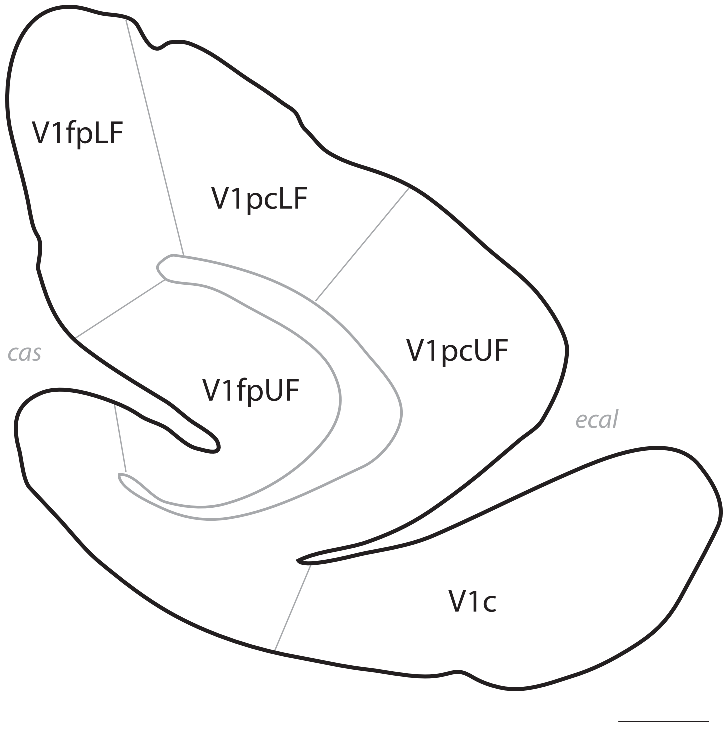
V1fpLF
V1pcLF
cas
V1fpUF
V1pcUF
ecal
V1c

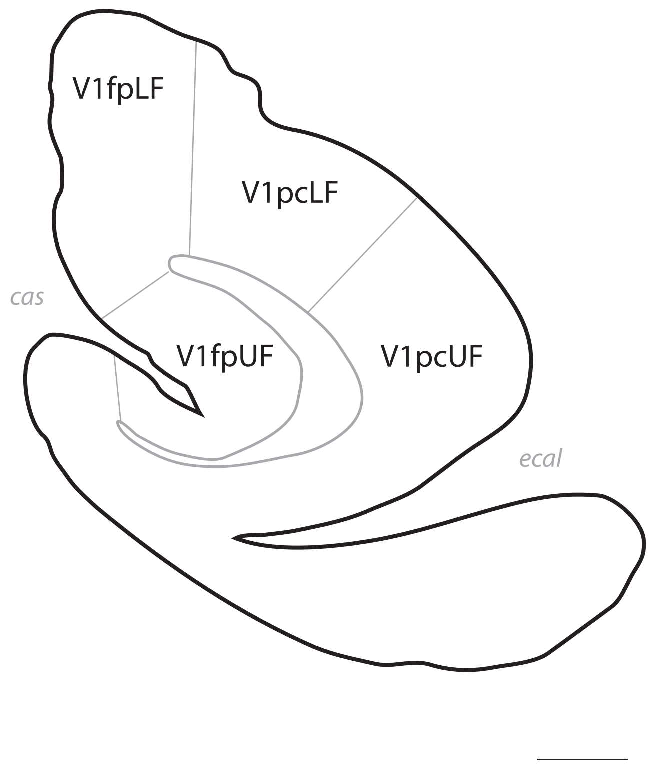
V1fpLF
V1pcLF
cas
V1fpUF
V1pcUF
ecal

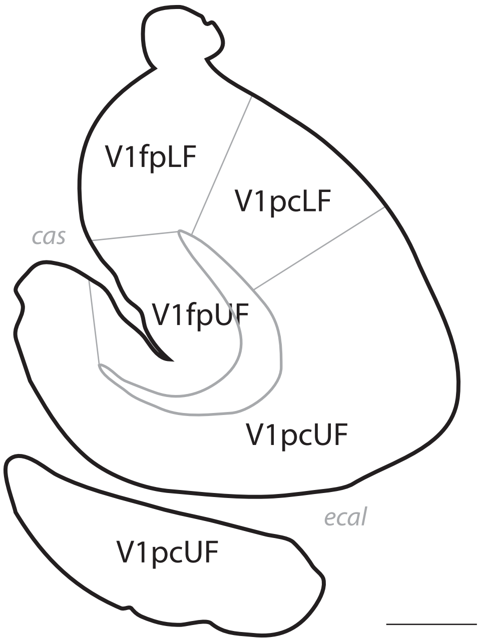
V1fpLF
V1pcLF
cas
V1fpUF
V1pcUF
ecal
V1pcUF

# p. 122

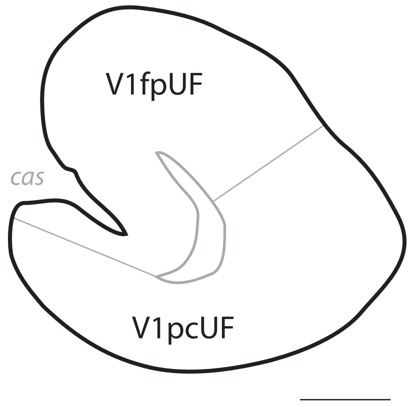

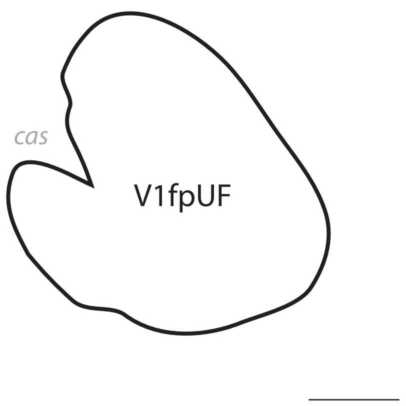
cas
V1fpUF

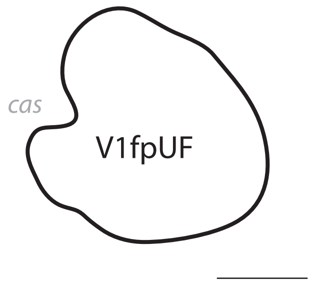
cas
V1fpUF
