## Supplementary material for "Retinotopic organization of feedback projections in primate early visual cortex: implications for active vision": Table S1

| Injection ID | Retinotopic Location | Monkey/ Hemisphere | Tracer | Age | Sex | Tracer Volume Injected | Layer Injection span |
|---|---|---|---|---|---|---|---|
| Case 1 | V1c | C054RH | CTB-555 | 11 | F | 2ul (1%) | 1-6 |
| Case 2 | V1c | C054RH | CTB-647 | 11 | F | 2ul (1%) | 1-6 |
| Case 3 | V1c | C081RH | CTB-647 | 12 | F | 2ul (1%) | 1-6 |
| Case 4 | V1c | M121RH | DY | 13 | F | 0.3µl (2%) | 1-6 |
| Case 5 | V1pcLF | C062RH | FB | 14 | F | 0.2ul (3%) | 1-6 |
| Case 6 | V1pcLF | M122RH | FB | / | M | 0.3µl (3%) | 1-6 |
| Case 7 | V1pcLF | C060RH | FB | 7 | F | 0.2ul (3%) | 1-6 |
| Case 8 | V1pcLF | C081RH | CTB-555 | 12 | F | 2ul (1%) | 1-6 |
| Case 9 | V1fpLF | C027RH | CTB-647 | 10 | F | 2ul (1%) | 1-6 |
| Case 10 | V1fpLF | C054RH | FB | 11 | F | 0.2ul (3%) | 1-6 |
| Case 11 | V1fpLF | C081RH | FB | 12 | F | 0.1ul (3%) | 1-6 |
| Case 12 | V1fpUF | C057RH | FB | 10 | F | 0.09ul (3%) | 1-4 |
| Case 13 | V1fpUF | C083RH | FB | 11 | F | 0.08ul (3%) | 1-6 |
| Case 14 | V1fpUF | C060RH | CTB-647 | 7 | F | 0.3ul (3%) | 1-6 |
| Case 15 | V2c | M101LH | DY | / | M | 0,65µl (3%) | 1-6 |
| Case 16 | V2c | M101RH | FB | / | M | 0.4µl (3%) | 1-6 |
| Case 17 | V2c | M103LH | DY | 4 | F | 0.2µl (3%) | 1-6 |
| Case 18 | V2c | M146LH | DY | 16 | F | 0.35µl (2%) | 1-6 |
| Case 19 | V2pcUF | M148RH | FB | 13 | F | 0.4µl (3%) | 1-6 |
| Case 20* | V2fpLF | M097LH | FB | / | F | / | 1-6 |
| Case 21* | V2fpUF/ V2fpLF | M096LH | FB | / | M | 0.25µl (3%) | 1-6 |

Table 1 Animal and injections used in the present study. c: central representation; fpUF: far periphery upper field; fpLF: far periphery lower field; pcUF: paracentral upper field; pcLF: paracentral lower field; CTB-555/647: cholera toxin subunit B, conjugated with Alexa 555/647; F: female; M: male.
