## Supplementary material for "Retinotopic organization of feedback projections in primate early visual cortex: implications for active vision": Table S2

FLN and SLN of All Injections Used in The Study

| | Case | MONKEY | SOURCE | TARGET | TOT | SUP | SLN | FLN | lgFLN |
|---|---|---|---|---|---|---|---|---|---|
| 1 | Case1 | C054 | 1 | V1c | 0 | 0 | NA | 0 | -Inf |
| 2 | Case1 | C054 | 10 | V1c | 0 | 0 | NA | 0 | -Inf |
| 3 | Case1 | C054 | 11 | V1c | 0 | 0 | NA | 0 | -Inf |
| 4 | Case1 | C054 | 12 | V1c | 0 | 0 | NA | 0 | -Inf |
| 5 | Case1 | C054 | 13 | V1c | 0 | 0 | NA | 0 | -Inf |
| 6 | Case1 | C054 | 14 | V1c | 0 | 0 | NA | 0 | -Inf |
| 7 | Case1 | C054 | 2 | V1c | 0 | 0 | NA | 0 | -Inf |
| 8 | Case1 | C054 | 23 | V1c | 0 | 0 | NA | 0 | -Inf |
| 9 | Case1 | C054 | 24a | V1c | 0 | 0 | NA | 0 | -Inf |
| 10 | Case1 | C054 | 24b | V1c | 0 | 0 | NA | 0 | -Inf |
| 11 | Case1 | C054 | 24c | V1c | 0 | 0 | NA | 0 | -Inf |
| 12 | Case1 | C054 | 24d | V1c | 0 | 0 | NA | 0 | -Inf |
| 13 | Case1 | C054 | 25 | V1c | 0 | 0 | NA | 0 | -Inf |
| 14 | Case1 | C054 | 29/30 | V1c | 0 | 0 | NA | 0 | -Inf |
| 15 | Case1 | C054 | 3 | V1c | 0 | 0 | NA | 0 | -Inf |
| 16 | Case1 | C054 | 31 | V1c | 0 | 0 | NA | 0 | -Inf |
| 17 | Case1 | C054 | 32 | V1c | 0 | 0 | NA | 0 | -Inf |
| 18 | Case1 | C054 | 44 | V1c | 0 | 0 | NA | 0 | -Inf |
| 19 | Case1 | C054 | 45A | V1c | 0 | 0 | NA | 0 | -Inf |
| 20 | Case1 | C054 | 45B | V1c | 0 | 0 | NA | 0 | -Inf |
| 21 | Case1 | C054 | 46d | V1c | 0 | 0 | NA | 0 | -Inf |
| 22 | Case1 | C054 | 46v | V1c | 0 | 0 | NA | 0 | -Inf |
| 23 | Case1 | C054 | 5 | V1c | 0 | 0 | NA | 0 | -Inf |
| 24 | Case1 | C054 | 7A | V1c | 0 | 0 | NA | 0 | -Inf |
| 25 | Case1 | C054 | 7B | V1c | 0 | 0 | NA | 0 | -Inf |
| 26 | Case1 | C054 | 7m | V1c | 0 | 0 | NA | 0 | -Inf |
| 27 | Case1 | C054 | 7op | V1c | 0 | 0 | NA | 0 | -Inf |
| 28 | Case1 | C054 | 8B | V1c | 0 | 0 | NA | 0 | -Inf |
| 29 | Case1 | C054 | 8l | V1c | 6 | 2 | 0,333333333 | 3,37838788960555E-05 | -4,471290488 |
| 30 | Case1 | C054 | 8m | V1c | 2 | 0 | 0 | 1,12612929653518E-05 | -4,948411743 |
| 31 | Case1 | C054 | 8r | V1c | 0 | 0 | NA | 0 | -Inf |
| 32 | Case1 | C054 | 9 | V1c | 0 | 0 | NA | 0 | -Inf |
| 33 | Case1 | C054 | 9/46d | V1c | 0 | 0 | NA | 0 | -Inf |
| 34 | Case1 | C054 | 9/46v | V1c | 0 | 0 | NA | 0 | -Inf |
| 35 | Case1 | C054 | AIP | V1c | 0 | 0 | NA | 0 | -Inf |
| 36 | Case1 | C054 | CORE | V1c | 6 | 0 | 0 | 3,37838788960555E-05 | -4,471290488 |
| 37 | Case1 | C054 | DP | V1c | 12 | 0 | 0 | 6,75677577921109E-05 | -4,170260493 |
| 38 | Case1 | C054 | ENTO | V1c | 0 | 0 | NA | 0 | -Inf |
| 39 | Case1 | C054 | F1 | V1c | 0 | 0 | NA | 0 | -Inf |
| 40 | Case1 | C054 | F2 | V1c | 0 | 0 | NA | 0 | -Inf |
| 41 | Case1 | C054 | F3 | V1c | 0 | 0 | NA | 0 | -Inf |
| 42 | Case1 | C054 | F4 | V1c | 0 | 0 | NA | 0 | -Inf |
| 43 | Case1 | C054 | F5 | V1c | 0 | 0 | NA | 0 | -Inf |
| 44 | Case1 | C054 | F6 | V1c | 0 | 0 | NA | 0 | -Inf |
| 45 | Case1 | C054 | F7 | V1c | 0 | 0 | NA | 0 | -Inf |
| 46 | Case1 | C054 | FST | V1c | 390 | 6 | 0,015384615 | 0,002195952 | -2,658377132 |
| 47 | Case1 | C054 | Gu | V1c | 0 | 0 | NA | 0 | -Inf |
| 48 | Case1 | C054 | INSULA | V1c | 30 | 0 | 0 | 0,000168919 | -3,772320484 |
| 49 | Case1 | C054 | IPa | V1c | 236 | 10 | 0,042372881 | 0,001328833 | -2,876529736 |
| 50 | Case1 | C054 | LB | V1c | 14 | 0 | 0 | 7,88290507574627E-05 | -4,103313703 |
| 51 | Case1 | C054 | LIP | V1c | 54 | 0 | 0 | 0,000304055 | -3,517047979 |
| 52 | Case1 | C054 | MB | V1c | 20 | 0 | 0 | 0,000112613 | -3,948411743 |
| 53 | Case1 | C054 | MIP | V1c | 0 | 0 | NA | 0 | -Inf |
| 54 | Case1 | C054 | MST | V1c | 210 | 0 | 0 | 0,001182436 | -2,927222444 |
| 55 | Case1 | C054 | MTc | V1c | 3736 | 614 | 0,164346895 | 0,021036095 | -1,677034871 |
| 56 | Case1 | C054 | MTp | V1c | 506 | 2 | 0,003952569 | 0,002849107 | -2,545291222 |
| 57 | Case1 | C054 | OPAl | V1c | 0 | 0 | NA | 0 | -Inf |
| 58 | Case1 | C054 | OPro | V1c | 0 | 0 | NA | 0 | -Inf |
| 59 | Case1 | C054 | PBc | V1c | 16 | 0 | 0 | 9,00903437228145E-05 | -4,045321756 |
| 60 | Case1 | C054 | PBr | V1c | 0 | 0 | NA | 0 | -Inf |
| 61 | Case1 | C054 | PERI | V1c | 42 | 2 | 0,047619048 | 0,000236487 | -3,626192448 |
| 62 | Case1 | C054 | PGa | V1c | 128 | 0 | 0 | 0,000720723 | -3,142231769 |
| 63 | Case1 | C054 | Pi | V1c | 0 | 0 | NA | 0 | -Inf |
| 64 | Case1 | C054 | PIP | V1c | 194 | 0 | 0 | 0,001092345 | -2,961640009 |
| 65 | Case1 | C054 | Pir | V1c | 0 | 0 | NA | 0 | -Inf |
| 66 | Case1 | C054 | POLE | V1c | 0 | 0 | NA | 0 | -Inf |
| 67 | Case1 | C054 | Pro.St | V1c | 0 | 0 | NA | 0 | -Inf |
| 68 | Case1 | C054 | ProM | V1c | 0 | 0 | NA | 0 | -Inf |
| 69 | Case1 | C054 | SII | V1c | 0 | 0 | NA | 0 | -Inf |
| 70 | Case1 | C054 | STPc | V1c | 56 | 0 | 0 | 0,000315316 | -3,501253712 |
| 71 | Case1 | C054 | STPi | V1c | 14 | 0 | 0 | 7,88290507574627E-05 | -4,103313703 |
| 72 | Case1 | C054 | STPr | V1c | 4 | 0 | 0 | 2,25225859307036E-05 | -4,647381747 |
| 73 | Case1 | C054 | SUBI | V1c | 0 | 0 | NA | 0 | -Inf |

| | Case | MONKEY | SOURCE | TARGET | TOT | SUP | SLN | FLN | lgFLN |
|---|---|---|---|---|---|---|---|---|---|
| 74 | Case1 | C054 | TEa/ma | V1c | 10 | 0 | 0 | 5,63064648267591E-05 | -4,249441739 |
| 75 | Case1 | C054 | TEa/mp | V1c | 491 | 15 | 0,030549898 | 0,002764647 | -2,558360247 |
| 76 | Case1 | C054 | TEad | V1c | 68 | 0 | 0 | 0,000382884 | -3,416932826 |
| 77 | Case1 | C054 | TEav | V1c | 30 | 4 | 0,133333333 | 0,000168919 | -3,772320484 |
| 78 | Case1 | C054 | TEO | V1c | 1052 | 96 | 0,091254753 | 0,00592344 | -2,227425999 |
| 79 | Case1 | C054 | TEOm | V1c | 50 | 8 | 0,16 | 0,000281532 | -3,550471734 |
| 80 | Case1 | C054 | TEpd | V1c | 762 | 55 | 0,072178478 | 0,004290553 | -2,367486767 |
| 81 | Case1 | C054 | TEpv | V1c | 653 | 4 | 0,006125574 | 0,003676812 | -2,434528557 |
| 82 | Case1 | C054 | TH_TF | V1c | 465 | 0 | 0 | 0,002618251 | -2,581988786 |
| 83 | Case1 | C054 | TPt | V1c | 0 | 0 | NA | 0 | -Inf |
| 84 | Case1 | C054 | V1c | V1c | 0 | 0 | NA | 0 | -Inf |
| 85 | Case1 | C054 | V1fpLF | V1c | 0 | 0 | NA | 0 | -Inf |
| 86 | Case1 | C054 | V1fpUF | V1c | 0 | 0 | NA | 0 | -Inf |
| 87 | Case1 | C054 | V1pcLF | V1c | 6 | 0 | 0 | 3,37838788960555E-05 | -4,471290488 |
| 88 | Case1 | C054 | V1pcUF | V1c | 2 | 0 | 0 | 1,12612929653518E-05 | -4,948411743 |
| 89 | Case1 | C054 | V2c | V1c | 26758 | 7580 | 0,283279767 | 0,150664839 | -0,821988089 |
| 90 | Case1 | C054 | V2fpLF | V1c | 0 | 0 | NA | 0 | -Inf |
| 91 | Case1 | C054 | V2fpUF | V1c | 4 | 0 | 0 | 2,25225859307036E-05 | -4,647381747 |
| 92 | Case1 | C054 | V2pcLF | V1c | 12 | 0 | 0 | 6,75677577921109E-05 | -4,170260493 |
| 93 | Case1 | C054 | V2pcUF | V1c | 14 | 0 | 0 | 7,88290507574627E-05 | -4,103313703 |
| 94 | Case1 | C054 | V3A | V1c | 18 | 0 | 0 | 0,000101352 | -3,994169234 |
| 95 | Case1 | C054 | V3c | V1c | 4064 | 1622 | 0,399114173 | 0,022882947 | -1,640488039 |
| 96 | Case1 | C054 | V3fpLF | V1c | 6 | 0 | 0 | 3,37838788960555E-05 | -4,471290488 |
| 97 | Case1 | C054 | V3fpUF | V1c | 10 | 0 | 0 | 5,63064648267591E-05 | -4,249441739 |
| 98 | Case1 | C054 | V3pcLF | V1c | 64 | 2 | 0,03125 | 0,000360361 | -3,443261765 |
| 99 | Case1 | C054 | V3pcUF | V1c | 22 | 0 | 0 | 0,000123874 | -3,907019058 |
| 100 | Case1 | C054 | V4c | V1c | 12110 | 2642 | 0,218166804 | 0,068187129 | -1,166297596 |
| 101 | Case1 | C054 | V4LF | V1c | 560 | 264 | 0,471428571 | 0,003153162 | -2,501253712 |
| 102 | Case1 | C054 | V4t | V1c | 278 | 26 | 0,09352518 | 0,00156532 | -2,805396943 |
| 103 | Case1 | C054 | V4UF | V1c | 212 | 4 | 0,018867925 | 0,001193697 | -2,923105878 |
| 104 | Case1 | C054 | V6 | V1c | 0 | 0 | NA | 0 | -Inf |
| 105 | Case1 | C054 | V6A | V1c | 0 | 0 | NA | 0 | -Inf |
| 106 | Case1 | C054 | VIP | V1c | 2 | 0 | 0 | 1,12612929653518E-05 | -4,948411743 |
| 107 | Case2 | C054 | 1 | V1c | 0 | 0 | NA | 0 | -Inf |
| 108 | Case2 | C054 | 10 | V1c | 0 | 0 | NA | 0 | -Inf |
| 109 | Case2 | C054 | 11 | V1c | 0 | 0 | NA | 0 | -Inf |
| 110 | Case2 | C054 | 12 | V1c | 0 | 0 | NA | 0 | -Inf |
| 111 | Case2 | C054 | 13 | V1c | 0 | 0 | NA | 0 | -Inf |
| 112 | Case2 | C054 | 14 | V1c | 0 | 0 | NA | 0 | -Inf |
| 113 | Case2 | C054 | 2 | V1c | 0 | 0 | NA | 0 | -Inf |
| 114 | Case2 | C054 | 23 | V1c | 0 | 0 | NA | 0 | -Inf |
| 115 | Case2 | C054 | 24a | V1c | 0 | 0 | NA | 0 | -Inf |
| 116 | Case2 | C054 | 24b | V1c | 0 | 0 | NA | 0 | -Inf |
| 117 | Case2 | C054 | 24c | V1c | 0 | 0 | NA | 0 | -Inf |
| 118 | Case2 | C054 | 24d | V1c | 0 | 0 | NA | 0 | -Inf |
| 119 | Case2 | C054 | 25 | V1c | 0 | 0 | NA | 0 | -Inf |
| 120 | Case2 | C054 | 29/30 | V1c | 0 | 0 | NA | 0 | -Inf |
| 121 | Case2 | C054 | 3 | V1c | 0 | 0 | NA | 0 | -Inf |
| 122 | Case2 | C054 | 31 | V1c | 0 | 0 | NA | 0 | -Inf |
| 123 | Case2 | C054 | 32 | V1c | 0 | 0 | NA | 0 | -Inf |
| 124 | Case2 | C054 | 44 | V1c | 0 | 0 | NA | 0 | -Inf |
| 125 | Case2 | C054 | 45A | V1c | 0 | 0 | NA | 0 | -Inf |
| 126 | Case2 | C054 | 45B | V1c | 0 | 0 | NA | 0 | -Inf |
| 127 | Case2 | C054 | 46d | V1c | 0 | 0 | NA | 0 | -Inf |
| 128 | Case2 | C054 | 46v | V1c | 0 | 0 | NA | 0 | -Inf |
| 129 | Case2 | C054 | 5 | V1c | 0 | 0 | NA | 0 | -Inf |
| 130 | Case2 | C054 | 7A | V1c | 0 | 0 | NA | 0 | -Inf |
| 131 | Case2 | C054 | 7B | V1c | 0 | 0 | NA | 0 | -Inf |
| 132 | Case2 | C054 | 7m | V1c | 0 | 0 | NA | 0 | -Inf |
| 133 | Case2 | C054 | 7op | V1c | 0 | 0 | NA | 0 | -Inf |
| 134 | Case2 | C054 | 8B | V1c | 0 | 0 | NA | 0 | -Inf |
| 135 | Case2 | C054 | 8l | V1c | 2 | 0 | 0 | 1,12612929653518E-05 | -4,948411743 |
| 136 | Case2 | C054 | 8m | V1c | 0 | 0 | NA | 0 | -Inf |
| 137 | Case2 | C054 | 8r | V1c | 0 | 0 | NA | 0 | -Inf |
| 138 | Case2 | C054 | 9 | V1c | 0 | 0 | NA | 0 | -Inf |
| 139 | Case2 | C054 | 9/46d | V1c | 0 | 0 | NA | 0 | -Inf |
| 140 | Case2 | C054 | 9/46v | V1c | 0 | 0 | NA | 0 | -Inf |
| 141 | Case2 | C054 | AIP | V1c | 0 | 0 | NA | 0 | -Inf |
| 142 | Case2 | C054 | CORE | V1c | 4 | 0 | 0 | 2,25225859307036E-05 | -4,647381747 |
| 143 | Case2 | C054 | DP | V1c | 12 | 4 | 0,333333333 | 6,75677577921109E-05 | -4,170260493 |
| 144 | Case2 | C054 | ENTO | V1c | 0 | 0 | NA | 0 | -Inf |
| 145 | Case2 | C054 | F1 | V1c | 0 | 0 | NA | 0 | -Inf |
| 146 | Case2 | C054 | F2 | V1c | 0 | 0 | NA | 0 | -Inf |
| 147 | Case2 | C054 | F3 | V1c | 0 | 0 | NA | 0 | -Inf |
| 148 | Case2 | C054 | F4 | V1c | 0 | 0 | NA | 0 | -Inf |

| | Case | MONKEY | SOURCE | TARGET | TOT | SUP | SLN | FLN | lgFLN |
|---|---|---|---|---|---|---|---|---|---|
| 149 | Case2 | C054 | F5 | V1c | 0 | 0 | NA | 0 | -Inf |
| 150 | Case2 | C054 | F6 | V1c | 0 | 0 | NA | 0 | -Inf |
| 151 | Case2 | C054 | F7 | V1c | 0 | 0 | NA | 0 | -Inf |
| 152 | Case2 | C054 | FST | V1c | 96 | 4 | 0,041666667 | 0,000540542 | -3,267170506 |
| 153 | Case2 | C054 | Gu | V1c | 0 | 0 | NA | 0 | -Inf |
| 154 | Case2 | C054 | INSULA | V1c | 2 | 0 | 0 | 1,12612929653518E-05 | -4,948411743 |
| 155 | Case2 | C054 | IPa | V1c | 81 | 0 | 0 | 0,000456082 | -3,34095672 |
| 156 | Case2 | C054 | LB | V1c | 6 | 0 | 0 | 3,37838788960555E-05 | -4,471290488 |
| 157 | Case2 | C054 | LIP | V1c | 18 | 0 | 0 | 0,000101352 | -3,994169234 |
| 158 | Case2 | C054 | MB | V1c | 0 | 0 | NA | 0 | -Inf |
| 159 | Case2 | C054 | MIP | V1c | 0 | 0 | NA | 0 | -Inf |
| 160 | Case2 | C054 | MST | V1c | 34 | 0 | 0 | 0,000191442 | -3,717962822 |
| 161 | Case2 | C054 | MTc | V1c | 1090 | 126 | 0,11559633 | 0,006137405 | -2,212015241 |
| 162 | Case2 | C054 | MTp | V1c | 128 | 0 | 0 | 0,000720723 | -3,142231769 |
| 163 | Case2 | C054 | OPAl | V1c | 0 | 0 | NA | 0 | -Inf |
| 164 | Case2 | C054 | OPro | V1c | 0 | 0 | NA | 0 | -Inf |
| 165 | Case2 | C054 | PBc | V1c | 2 | 0 | 0 | 1,12612929653518E-05 | -4,948411743 |
| 166 | Case2 | C054 | PBr | V1c | 0 | 0 | NA | 0 | -Inf |
| 167 | Case2 | C054 | PERI | V1c | 4 | 0 | 0 | 2,25225859307036E-05 | -4,647381747 |
| 168 | Case2 | C054 | PGa | V1c | 26 | 0 | 0 | 0,000146397 | -3,834468391 |
| 169 | Case2 | C054 | Pi | V1c | 0 | 0 | NA | 0 | -Inf |
| 170 | Case2 | C054 | PIP | V1c | 30 | 0 | 0 | 0,000168919 | -3,772320484 |
| 171 | Case2 | C054 | Pir | V1c | 0 | 0 | NA | 0 | -Inf |
| 172 | Case2 | C054 | POLE | V1c | 0 | 0 | NA | 0 | -Inf |
| 173 | Case2 | C054 | Pro.St | V1c | 0 | 0 | NA | 0 | -Inf |
| 174 | Case2 | C054 | ProM | V1c | 0 | 0 | NA | 0 | -Inf |
| 175 | Case2 | C054 | SII | V1c | 0 | 0 | NA | 0 | -Inf |
| 176 | Case2 | C054 | STPc | V1c | 8 | 0 | 0 | 4,50451718614073E-05 | -4,346351752 |
| 177 | Case2 | C054 | STPi | V1c | 6 | 0 | 0 | 3,37838788960555E-05 | -4,471290488 |
| 178 | Case2 | C054 | STPr | V1c | 0 | 0 | NA | 0 | -Inf |
| 179 | Case2 | C054 | SUBI | V1c | 0 | 0 | NA | 0 | -Inf |
| 180 | Case2 | C054 | TEa/ma | V1c | 2 | 0 | 0 | 1,12612929653518E-05 | -4,948411743 |
| 181 | Case2 | C054 | TEa/mp | V1c | 114 | 4 | 0,035087719 | 0,000641894 | -3,192536887 |
| 182 | Case2 | C054 | TEad | V1c | 6 | 2 | 0,333333333 | 3,37838788960555E-05 | -4,471290488 |
| 183 | Case2 | C054 | TEav | V1c | 10 | 0 | 0 | 5,63064648267591E-05 | -4,249441739 |
| 184 | Case2 | C054 | TEO | V1c | 246 | 28 | 0,113821138 | 0,001385139 | -2,858506632 |
| 185 | Case2 | C054 | TEOm | V1c | 8 | 2 | 0,25 | 4,50451718614073E-05 | -4,346351752 |
| 186 | Case2 | C054 | TEpd | V1c | 156 | 10 | 0,064102564 | 0,000878381 | -3,05631714 |
| 187 | Case2 | C054 | TEpv | V1c | 93 | 2 | 0,021505376 | 0,00052365 | -3,28095879 |
| 188 | Case2 | C054 | TH_TF | V1c | 69 | 0 | 0 | 0,000388515 | -3,410592648 |
| 189 | Case2 | C054 | TPt | V1c | 0 | 0 | NA | 0 | -Inf |
| 190 | Case2 | C054 | V1c | V1c | 0 | 0 | NA | 0 | -Inf |
| 191 | Case2 | C054 | V1fpLF | V1c | 0 | 0 | NA | 0 | -Inf |
| 192 | Case2 | C054 | V1fpUF | V1c | 0 | 0 | NA | 0 | -Inf |
| 193 | Case2 | C054 | V1pcLF | V1c | 0 | 0 | NA | 0 | -Inf |
| 194 | Case2 | C054 | V1pcUF | V1c | 0 | 0 | NA | 0 | -Inf |
| 195 | Case2 | C054 | V2c | V1c | 11964 | 3596 | 0,300568372 | 0,067365055 | -1,171565334 |
| 196 | Case2 | C054 | V2fpLF | V1c | 0 | 0 | NA | 0 | -Inf |
| 197 | Case2 | C054 | V2fpUF | V1c | 2 | 0 | 0 | 1,12612929653518E-05 | -4,948411743 |
| 198 | Case2 | C054 | V2pcLF | V1c | 2 | 0 | 0 | 1,12612929653518E-05 | -4,948411743 |
| 199 | Case2 | C054 | V2pcUF | V1c | 0 | 0 | NA | 0 | -Inf |
| 200 | Case2 | C054 | V3A | V1c | 4 | 0 | 0 | 2,25225859307036E-05 | -4,647381747 |
| 201 | Case2 | C054 | V3c | V1c | 1790 | 722 | 0,403351955 | 0,010078857 | -1,996588708 |
| 202 | Case2 | C054 | V3fpLF | V1c | 0 | 0 | NA | 0 | -Inf |
| 203 | Case2 | C054 | V3fpUF | V1c | 2 | 0 | 0 | 1,12612929653518E-05 | -4,948411743 |
| 204 | Case2 | C054 | V3pcLF | V1c | 12 | 0 | 0 | 6,75677577921109E-05 | -4,170260493 |
| 205 | Case2 | C054 | V3pcUF | V1c | 0 | 0 | NA | 0 | -Inf |
| 206 | Case2 | C054 | V4c | V1c | 2538 | 268 | 0,105594957 | 0,014290581 | -1,844950121 |
| 207 | Case2 | C054 | V4LF | V1c | 142 | 86 | 0,605633803 | 0,000799552 | -3,097153394 |
| 208 | Case2 | C054 | V4t | V1c | 76 | 4 | 0,052631579 | 0,000427929 | -3,368628146 |
| 209 | Case2 | C054 | V4UF | V1c | 38 | 0 | 0 | 0,000213965 | -3,669658142 |
| 210 | Case2 | C054 | V6 | V1c | 0 | 0 | NA | 0 | -Inf |
| 211 | Case2 | C054 | V6A | V1c | 0 | 0 | NA | 0 | -Inf |
| 212 | Case2 | C054 | VIP | V1c | 0 | 0 | NA | 0 | -Inf |
| 213 | Case3 | C081 | 1 | V1c | 0 | 0 | NA | 0 | -Inf |
| 214 | Case3 | C081 | 10 | V1c | 0 | 0 | NA | 0 | -Inf |
| 215 | Case3 | C081 | 11 | V1c | 0 | 0 | NA | 0 | -Inf |
| 216 | Case3 | C081 | 12 | V1c | 0 | 0 | NA | 0 | -Inf |
| 217 | Case3 | C081 | 13 | V1c | 0 | 0 | NA | 0 | -Inf |
| 218 | Case3 | C081 | 14 | V1c | 0 | 0 | NA | 0 | -Inf |
| 219 | Case3 | C081 | 2 | V1c | 0 | 0 | NA | 0 | -Inf |
| 220 | Case3 | C081 | 23 | V1c | 0 | 0 | NA | 0 | -Inf |
| 221 | Case3 | C081 | 24a | V1c | 0 | 0 | NA | 0 | -Inf |
| 222 | Case3 | C081 | 24b | V1c | 0 | 0 | NA | 0 | -Inf |
| 223 | Case3 | C081 | 24c | V1c | 0 | 0 | NA | 0 | -Inf |

| | Case | MONKEY | SOURCE | TARGET | TOT | SUP | SLN | FLN | lgFLN |
|---|---|---|---|---|---|---|---|---|---|
| 224 | Case3 | C081 | 24d | V1c | 0 | 0 | NA | 0 | -Inf |
| 225 | Case3 | C081 | 25 | V1c | 0 | 0 | NA | 0 | -Inf |
| 226 | Case3 | C081 | 29/30 | V1c | 0 | 0 | NA | 0 | -Inf |
| 227 | Case3 | C081 | 3 | V1c | 0 | 0 | NA | 0 | -Inf |
| 228 | Case3 | C081 | 31 | V1c | 0 | 0 | NA | 0 | -Inf |
| 229 | Case3 | C081 | 32 | V1c | 0 | 0 | NA | 0 | -Inf |
| 230 | Case3 | C081 | 44 | V1c | 0 | 0 | NA | 0 | -Inf |
| 231 | Case3 | C081 | 45A | V1c | 0 | 0 | NA | 0 | -Inf |
| 232 | Case3 | C081 | 45B | V1c | 0 | 0 | NA | 0 | -Inf |
| 233 | Case3 | C081 | 46d | V1c | 0 | 0 | NA | 0 | -Inf |
| 234 | Case3 | C081 | 46v | V1c | 0 | 0 | NA | 0 | -Inf |
| 235 | Case3 | C081 | 5 | V1c | 0 | 0 | NA | 0 | -Inf |
| 236 | Case3 | C081 | 7A | V1c | 0 | 0 | NA | 0 | -Inf |
| 237 | Case3 | C081 | 7B | V1c | 0 | 0 | NA | 0 | -Inf |
| 238 | Case3 | C081 | 7m | V1c | 0 | 0 | NA | 0 | -Inf |
| 239 | Case3 | C081 | 7op | V1c | 0 | 0 | NA | 0 | -Inf |
| 240 | Case3 | C081 | 8B | V1c | 0 | 0 | NA | 0 | -Inf |
| 241 | Case3 | C081 | 8l | V1c | 0 | 0 | NA | 0 | -Inf |
| 242 | Case3 | C081 | 8m | V1c | 0 | 0 | NA | 0 | -Inf |
| 243 | Case3 | C081 | 8r | V1c | 0 | 0 | NA | 0 | -Inf |
| 244 | Case3 | C081 | 9 | V1c | 0 | 0 | NA | 0 | -Inf |
| 245 | Case3 | C081 | 9/46d | V1c | 0 | 0 | NA | 0 | -Inf |
| 246 | Case3 | C081 | 9/46v | V1c | 0 | 0 | NA | 0 | -Inf |
| 247 | Case3 | C081 | AIP | V1c | 0 | 0 | NA | 0 | -Inf |
| 248 | Case3 | C081 | CORE | V1c | 0 | 0 | NA | 0 | -Inf |
| 249 | Case3 | C081 | DP | V1c | 2 | 2 | 1 | 1,12612929653518E-05 | -4,948411743 |
| 250 | Case3 | C081 | ENTO | V1c | 0 | 0 | NA | 0 | -Inf |
| 251 | Case3 | C081 | F1 | V1c | 0 | 0 | NA | 0 | -Inf |
| 252 | Case3 | C081 | F2 | V1c | 0 | 0 | NA | 0 | -Inf |
| 253 | Case3 | C081 | F3 | V1c | 0 | 0 | NA | 0 | -Inf |
| 254 | Case3 | C081 | F4 | V1c | 0 | 0 | NA | 0 | -Inf |
| 255 | Case3 | C081 | F5 | V1c | 0 | 0 | NA | 0 | -Inf |
| 256 | Case3 | C081 | F6 | V1c | 0 | 0 | NA | 0 | -Inf |
| 257 | Case3 | C081 | F7 | V1c | 0 | 0 | NA | 0 | -Inf |
| 258 | Case3 | C081 | FST | V1c | 196 | 196 | 1 | 0,001103607 | -2,957185667 |
| 259 | Case3 | C081 | Gu | V1c | 0 | 0 | NA | 0 | -Inf |
| 260 | Case3 | C081 | INSULA | V1c | 2 | 2 | 1 | 1,12612929653518E-05 | -4,948411743 |
| 261 | Case3 | C081 | IPa | V1c | 49 | 49 | 1 | 0,000275902 | -3,559245659 |
| 262 | Case3 | C081 | LB | V1c | 0 | 0 | NA | 0 | -Inf |
| 263 | Case3 | C081 | LIP | V1c | 9 | 9 | 1 | 5,06758183440832E-05 | -4,295199229 |
| 264 | Case3 | C081 | MB | V1c | 1 | 1 | 1 | 5,63064648267591E-06 | -5,249441739 |
| 265 | Case3 | C081 | MIP | V1c | 0 | 0 | NA | 0 | -Inf |
| 266 | Case3 | C081 | MST | V1c | 44,5 | 44,5 | 1 | 0,000250564 | -3,601081728 |
| 267 | Case3 | C081 | MTc | V1c | 760 | 760 | 1 | 0,004279291 | -2,368628146 |
| 268 | Case3 | C081 | MTp | V1c | 1330,5 | 1330,5 | 1 | 0,007491575 | -2,12542686 |
| 269 | Case3 | C081 | OPAl | V1c | 0 | 0 | NA | 0 | -Inf |
| 270 | Case3 | C081 | OPro | V1c | 0 | 0 | NA | 0 | -Inf |
| 271 | Case3 | C081 | PBc | V1c | 4 | 4 | 1 | 2,25225859307036E-05 | -4,647381747 |
| 272 | Case3 | C081 | PBr | V1c | 0 | 0 | NA | 0 | -Inf |
| 273 | Case3 | C081 | PERI | V1c | 25 | 25 | 1 | 0,000140766 | -3,85150173 |
| 274 | Case3 | C081 | PGa | V1c | 23 | 23 | 1 | 0,000129505 | -3,887713903 |
| 275 | Case3 | C081 | Pi | V1c | 0 | 0 | NA | 0 | -Inf |
| 276 | Case3 | C081 | PIP | V1c | 62 | 62 | 1 | 0,0003491 | -3,457050049 |
| 277 | Case3 | C081 | Pir | V1c | 0 | 0 | NA | 0 | -Inf |
| 278 | Case3 | C081 | POLE | V1c | 0 | 0 | NA | 0 | -Inf |
| 279 | Case3 | C081 | Pro.St | V1c | 0 | 0 | NA | 0 | -Inf |
| 280 | Case3 | C081 | ProM | V1c | 0 | 0 | NA | 0 | -Inf |
| 281 | Case3 | C081 | SII | V1c | 0 | 0 | NA | 0 | -Inf |
| 282 | Case3 | C081 | STPc | V1c | 3 | 3 | 1 | 1,68919394480277E-05 | -4,772320484 |
| 283 | Case3 | C081 | STPi | V1c | 6 | 6 | 1 | 3,37838788960555E-05 | -4,471290488 |
| 284 | Case3 | C081 | STPr | V1c | 0 | 0 | NA | 0 | -Inf |
| 285 | Case3 | C081 | SUBI | V1c | 0 | 0 | NA | 0 | -Inf |
| 286 | Case3 | C081 | TEa/ma | V1c | 1 | 1 | 1 | 5,63064648267591E-06 | -5,249441739 |
| 287 | Case3 | C081 | TEa/mp | V1c | 43 | 43 | 1 | 0,000242118 | -3,615973283 |
| 288 | Case3 | C081 | TEad | V1c | 25 | 25 | 1 | 0,000140766 | -3,85150173 |
| 289 | Case3 | C081 | TEav | V1c | 19 | 19 | 1 | 0,000106982 | -3,970688138 |
| 290 | Case3 | C081 | TEO | V1c | 456 | 456 | 1 | 0,002567575 | -2,590476896 |
| 291 | Case3 | C081 | TEOm | V1c | 188 | 188 | 1 | 0,001058562 | -2,97528389 |
| 292 | Case3 | C081 | TEpd | V1c | 154 | 154 | 1 | 0,00086712 | -3,061921018 |
| 293 | Case3 | C081 | TEpv | V1c | 340 | 340 | 1 | 0,00191442 | -2,717962822 |
| 294 | Case3 | C081 | TH_TF | V1c | 240 | 240 | 1 | 0,001351355 | -2,869230497 |
| 295 | Case3 | C081 | TPt | V1c | 0 | 0 | NA | 0 | -Inf |
| 296 | Case3 | C081 | V1c | V1c | 0 | 0 | NA | 0 | -Inf |
| 297 | Case3 | C081 | V1fpLF | V1c | 0 | 0 | NA | 0 | -Inf |
| 298 | Case3 | C081 | V1fpUF | V1c | 0 | 0 | NA | 0 | -Inf |

| | Case | MONKEY | SOURCE | TARGET | TOT | SUP | SLN | FLN | lgFLN |
|---|---|---|---|---|---|---|---|---|---|
| 299 | Case3 | C081 | V1pcLF | V1c | 20 | 20 | 1 | 0,000112613 | -3,948411743 |
| 300 | Case3 | C081 | V1pcUF | V1c | 0 | 0 | NA | 0 | -Inf |
| 301 | Case3 | C081 | V2c | V1c | 13132 | 13132 | 1 | 0,07394165 | -1,131110865 |
| 302 | Case3 | C081 | V2fpLF | V1c | 0 | 0 | NA | 0 | -Inf |
| 303 | Case3 | C081 | V2fpUF | V1c | 0 | 0 | NA | 0 | -Inf |
| 304 | Case3 | C081 | V2pcLF | V1c | 14 | 14 | 1 | 7,88290507574627E-05 | -4,103313703 |
| 305 | Case3 | C081 | V2pcUF | V1c | 2 | 2 | 1 | 1,12612929653518E-05 | -4,948411743 |
| 306 | Case3 | C081 | V3A | V1c | 10 | 10 | 1 | 5,63064648267591E-05 | -4,249441739 |
| 307 | Case3 | C081 | V3c | V1c | 2114 | 2114 | 1 | 0,011903187 | -1,924336756 |
| 308 | Case3 | C081 | V3fpLF | V1c | 16 | 16 | 1 | 9,00903437228145E-05 | -4,045321756 |
| 309 | Case3 | C081 | V3fpUF | V1c | 1 | 1 | 1 | 5,63064648267591E-06 | -5,249441739 |
| 310 | Case3 | C081 | V3pcLF | V1c | 42 | 42 | 1 | 0,000236487 | -3,626192448 |
| 311 | Case3 | C081 | V3pcUF | V1c | 2 | 2 | 1 | 1,12612929653518E-05 | -4,948411743 |
| 312 | Case3 | C081 | V4c | V1c | 2306 | 2306 | 1 | 0,012984271 | -1,886582436 |
| 313 | Case3 | C081 | V4LF | V1c | 124,5 | 124,5 | 1 | 0,000701015 | -3,154272387 |
| 314 | Case3 | C081 | V4t | V1c | 5 | 5 | 1 | 2,81532324133795E-05 | -4,550471734 |
| 315 | Case3 | C081 | V4UF | V1c | 313 | 313 | 1 | 0,001762392 | -2,753897401 |
| 316 | Case3 | C081 | V6 | V1c | 0 | 0 | NA | 0 | -Inf |
| 317 | Case3 | C081 | V6A | V1c | 0 | 0 | NA | 0 | -Inf |
| 318 | Case3 | C081 | VIP | V1c | 0 | 0 | NA | 0 | -Inf |
| 1591 | Case4 | M121RH | 1 | V1c | 0 | 0 | NA | 0 | -Inf |
| 1592 | Case4 | M121RH | 10 | V1c | 0 | 0 | NA | 0 | -Inf |
| 1593 | Case4 | M121RH | 11 | V1c | 0 | 0 | NA | 0 | -Inf |
| 1594 | Case4 | M121RH | 12 | V1c | 0 | 0 | NA | 0 | -Inf |
| 1595 | Case4 | M121RH | 13 | V1c | 0 | 0 | NA | 0 | -Inf |
| 1596 | Case4 | M121RH | 14 | V1c | 0 | 0 | NA | 0 | -Inf |
| 1597 | Case4 | M121RH | 2 | V1c | 0 | 0 | NA | 0 | -Inf |
| 1598 | Case4 | M121RH | 23 | V1c | 0 | 0 | NA | 0 | -Inf |
| 1599 | Case4 | M121RH | 24a | V1c | 0 | 0 | NA | 0 | -Inf |
| 1600 | Case4 | M121RH | 24b | V1c | 0 | 0 | NA | 0 | -Inf |
| 1601 | Case4 | M121RH | 24c | V1c | 0 | 0 | NA | 0 | -Inf |
| 1602 | Case4 | M121RH | 24d | V1c | 0 | 0 | NA | 0 | -Inf |
| 1603 | Case4 | M121RH | 25 | V1c | 0 | 0 | NA | 0 | -Inf |
| 1604 | Case4 | M121RH | 29/30 | V1c | 0 | 0 | NA | 0 | -Inf |
| 1605 | Case4 | M121RH | 3 | V1c | 0 | 0 | NA | 0 | -Inf |
| 1606 | Case4 | M121RH | 31 | V1c | 0 | 0 | NA | 0 | -Inf |
| 1607 | Case4 | M121RH | 32 | V1c | 0 | 0 | NA | 0 | -Inf |
| 1608 | Case4 | M121RH | 44 | V1c | 0 | 0 | NA | 0 | -Inf |
| 1609 | Case4 | M121RH | 45A | V1c | 0 | 0 | NA | 0 | -Inf |
| 1610 | Case4 | M121RH | 45B | V1c | 0 | 0 | NA | 0 | -Inf |
| 1611 | Case4 | M121RH | 46d | V1c | 0 | 0 | NA | 0 | -Inf |
| 1612 | Case4 | M121RH | 46v | V1c | 0 | 0 | NA | 0 | -Inf |
| 1613 | Case4 | M121RH | 5 | V1c | 0 | 0 | NA | 0 | -Inf |
| 1614 | Case4 | M121RH | 7A | V1c | 2 | 2 | 1 | 1,12612929653518E-05 | -4,948411743 |
| 1615 | Case4 | M121RH | 7B | V1c | 0 | 0 | NA | 0 | -Inf |
| 1616 | Case4 | M121RH | 7m | V1c | 0 | 0 | NA | 0 | -Inf |
| 1617 | Case4 | M121RH | 7op | V1c | 2 | 0 | 0 | 1,12612929653518E-05 | -4,948411743 |
| 1618 | Case4 | M121RH | 8B | V1c | 0 | 0 | NA | 0 | -Inf |
| 1619 | Case4 | M121RH | 8l | V1c | 31 | 1 | 0,032258065 | 0,00017455 | -3,758080045 |
| 1620 | Case4 | M121RH | 8m | V1c | 0 | 0 | NA | 0 | -Inf |
| 1621 | Case4 | M121RH | 8r | V1c | 0 | 0 | NA | 0 | -Inf |
| 1622 | Case4 | M121RH | 9 | V1c | 0 | 0 | NA | 0 | -Inf |
| 1623 | Case4 | M121RH | 9/46d | V1c | 0 | 0 | NA | 0 | -Inf |
| 1624 | Case4 | M121RH | 9/46v | V1c | 0 | 0 | NA | 0 | -Inf |
| 1625 | Case4 | M121RH | AIP | V1c | 0 | 0 | NA | 0 | -Inf |
| 1626 | Case4 | M121RH | CORE | V1c | 0 | 0 | NA | 0 | -Inf |
| 1627 | Case4 | M121RH | DP | V1c | 40 | 16 | 0,4 | 0,000225226 | -3,647381747 |
| 1628 | Case4 | M121RH | ENTO | V1c | 0 | 0 | NA | 0 | -Inf |
| 1629 | Case4 | M121RH | F1 | V1c | 0 | 0 | NA | 0 | -Inf |
| 1630 | Case4 | M121RH | F2 | V1c | 1 | 0 | 0 | 5,63064648267591E-06 | -5,249441739 |
| 1631 | Case4 | M121RH | F3 | V1c | 0 | 0 | NA | 0 | -Inf |
| 1632 | Case4 | M121RH | F4 | V1c | 0 | 0 | NA | 0 | -Inf |
| 1633 | Case4 | M121RH | F5 | V1c | 0 | 0 | NA | 0 | -Inf |
| 1634 | Case4 | M121RH | F6 | V1c | 0 | 0 | NA | 0 | -Inf |
| 1635 | Case4 | M121RH | F7 | V1c | 0 | 0 | NA | 0 | -Inf |
| 1636 | Case4 | M121RH | FST | V1c | 546 | 27 | 0,049450549 | 0,003074333 | -2,512249096 |
| 1637 | Case4 | M121RH | Gu | V1c | 0 | 0 | NA | 0 | -Inf |
| 1638 | Case4 | M121RH | INSULA | V1c | 0 | 0 | NA | 0 | -Inf |
| 1639 | Case4 | M121RH | IPa | V1c | 6 | 1 | 0,166666667 | 3,37838788960555E-05 | -4,471290488 |
| 1640 | Case4 | M121RH | LB | V1c | 0 | 0 | NA | 0 | -Inf |
| 1641 | Case4 | M121RH | LIP | V1c | 283 | 6 | 0,021201413 | 0,001593473 | -2,797655303 |
| 1642 | Case4 | M121RH | MB | V1c | 0 | 0 | NA | 0 | -Inf |
| 1643 | Case4 | M121RH | MIP | V1c | 0 | 0 | NA | 0 | -Inf |
| 1644 | Case4 | M121RH | MST | V1c | 202 | 0 | 0 | 0,001137391 | -2,944090369 |
| 1645 | Case4 | M121RH | MTc | V1c | 3798 | 604 | 0,159031069 | 0,021385195 | -1,669886778 |

| | Case | MONKEY | SOURCE | TARGET | TOT | SUP | SLN | FLN | lgFLN |
|---|---|---|---|---|---|---|---|---|---|
| 1646 | Case4 | M121RH | MTp | V1c | 309 | 6 | 0,019417476 | 0,00173987 | -2,759483259 |
| 1647 | Case4 | M121RH | OPAl | V1c | 0 | 0 | NA | 0 | -Inf |
| 1648 | Case4 | M121RH | OPro | V1c | 0 | 0 | NA | 0 | -Inf |
| 1649 | Case4 | M121RH | PBc | V1c | 4 | 0 | 0 | 2,25225859307036E-05 | -4,647381747 |
| 1650 | Case4 | M121RH | PBr | V1c | 0 | 0 | NA | 0 | -Inf |
| 1651 | Case4 | M121RH | PERI | V1c | 59 | 1 | 0,016949153 | 0,000332208 | -3,478589727 |
| 1652 | Case4 | M121RH | PGa | V1c | 112 | 3 | 0,026785714 | 0,000630632 | -3,200223716 |
| 1653 | Case4 | M121RH | Pi | V1c | 0 | 0 | NA | 0 | -Inf |
| 1654 | Case4 | M121RH | PIP | V1c | 164 | 0 | 0 | 0,000923426 | -3,034597891 |
| 1655 | Case4 | M121RH | Pir | V1c | 0 | 0 | NA | 0 | -Inf |
| 1656 | Case4 | M121RH | POLE | V1c | 0 | 0 | NA | 0 | -Inf |
| 1657 | Case4 | M121RH | Pro.St | V1c | 0 | 0 | NA | 0 | -Inf |
| 1658 | Case4 | M121RH | ProM | V1c | 0 | 0 | NA | 0 | -Inf |
| 1659 | Case4 | M121RH | SII | V1c | 0 | 0 | NA | 0 | -Inf |
| 1660 | Case4 | M121RH | STPc | V1c | 80 | 4 | 0,05 | 0,000450452 | -3,346351752 |
| 1661 | Case4 | M121RH | STPi | V1c | 3 | 0 | 0 | 1,68919394480277E-05 | -4,772320484 |
| 1662 | Case4 | M121RH | STPr | V1c | 1 | 0 | 0 | 5,63064648267591E-06 | -5,249441739 |
| 1663 | Case4 | M121RH | SUBI | V1c | 0 | 0 | NA | 0 | -Inf |
| 1664 | Case4 | M121RH | TEa/ma | V1c | 5 | 1 | 0,2 | 2,81532324133795E-05 | -4,550471734 |
| 1665 | Case4 | M121RH | TEa/mp | V1c | 263 | 21 | 0,079847909 | 0,00148086 | -2,82948599 |
| 1666 | Case4 | M121RH | TEad | V1c | 3 | 0 | 0 | 1,68919394480277E-05 | -4,772320484 |
| 1667 | Case4 | M121RH | TEav | V1c | 27 | 0 | 0 | 0,000152027 | -3,818077975 |
| 1668 | Case4 | M121RH | TEO | V1c | 326 | 13 | 0,039877301 | 0,001835591 | -2,736224139 |
| 1669 | Case4 | M121RH | TEOm | V1c | 866 | 67 | 0,077367206 | 0,00487614 | -2,311923847 |
| 1670 | Case4 | M121RH | TEpd | V1c | 137 | 10 | 0,072992701 | 0,000771399 | -3,112721172 |
| 1671 | Case4 | M121RH | TEpv | V1c | 245 | 1 | 0,004081633 | 0,001379508 | -2,860275654 |
| 1672 | Case4 | M121RH | TH_TF | V1c | 142 | 0 | 0 | 0,000799552 | -3,097153394 |
| 1673 | Case4 | M121RH | TPt | V1c | 0 | 0 | NA | 0 | -Inf |
| 1674 | Case4 | M121RH | V1c | V1c | 0 | 0 | NA | 0 | -Inf |
| 1675 | Case4 | M121RH | V1fpLF | V1c | 0 | 0 | NA | 0 | -Inf |
| 1676 | Case4 | M121RH | V1fpUF | V1c | 0 | 0 | NA | 0 | -Inf |
| 1677 | Case4 | M121RH | V1pcLF | V1c | 0 | 0 | NA | 0 | -Inf |
| 1678 | Case4 | M121RH | V1pcUF | V1c | 0 | 0 | NA | 0 | -Inf |
| 1679 | Case4 | M121RH | V2c | V1c | 57600 | 28152 | 0,48875 | 0,324325237 | -0,489019255 |
| 1680 | Case4 | M121RH | V2fpLF | V1c | 0 | 0 | NA | 0 | -Inf |
| 1681 | Case4 | M121RH | V2fpUF | V1c | 0 | 0 | NA | 0 | -Inf |
| 1682 | Case4 | M121RH | V2pcLF | V1c | 0 | 0 | NA | 0 | -Inf |
| 1683 | Case4 | M121RH | V2pcUF | V1c | 0 | 0 | NA | 0 | -Inf |
| 1684 | Case4 | M121RH | V3A | V1c | 62 | 2 | 0,032258065 | 0,0003491 | -3,457050049 |
| 1685 | Case4 | M121RH | V3c | V1c | 449 | 196 | 0,436525612 | 0,00252816 | -2,597195398 |
| 1686 | Case4 | M121RH | V3fpLF | V1c | 0 | 0 | NA | 0 | -Inf |
| 1687 | Case4 | M121RH | V3fpUF | V1c | 0 | 0 | NA | 0 | -Inf |
| 1688 | Case4 | M121RH | V3pcLF | V1c | 162 | 35 | 0,216049383 | 0,000912165 | -3,039926724 |
| 1689 | Case4 | M121RH | V3pcUF | V1c | 7 | 0 | 0 | 3,94145253787314E-05 | -4,404343699 |
| 1690 | Case4 | M121RH | V4c | V1c | 15763 | 6045 | 0,38349299 | 0,088755881 | -1,051802863 |
| 1691 | Case4 | M121RH | V4LF | V1c | 1147 | 352 | 0,306887533 | 0,006458352 | -2,189878321 |
| 1692 | Case4 | M121RH | V4t | V1c | 14 | 4 | 0,285714286 | 7,88290507574627E-05 | -4,103313703 |
| 1693 | Case4 | M121RH | V4UF | V1c | 430 | 27 | 0,062790698 | 0,002421178 | -2,615973283 |
| 1694 | Case4 | M121RH | V6 | V1c | 0 | 0 | NA | 0 | -Inf |
| 1695 | Case4 | M121RH | V6A | V1c | 0 | 0 | NA | 0 | -Inf |
| 1696 | Case4 | M121RH | VIP | V1c | 2 | 0 | 0 | 1,12612929653518E-05 | -4,948411743 |
| 1061 | Case5 | C062 | 1 | V1pcLF | 0 | 0 | NA | 0 | -Inf |
| 1062 | Case5 | C062 | 10 | V1pcLF | 0 | 0 | NA | 0 | -Inf |
| 1063 | Case5 | C062 | 11 | V1pcLF | 0 | 0 | NA | 0 | -Inf |
| 1064 | Case5 | C062 | 12 | V1pcLF | 0 | 0 | NA | 0 | -Inf |
| 1065 | Case5 | C062 | 13 | V1pcLF | 0 | 0 | NA | 0 | -Inf |
| 1066 | Case5 | C062 | 14 | V1pcLF | 0 | 0 | NA | 0 | -Inf |
| 1067 | Case5 | C062 | 2 | V1pcLF | 0 | 0 | NA | 0 | -Inf |
| 1068 | Case5 | C062 | 23 | V1pcLF | 0 | 0 | NA | 0 | -Inf |
| 1069 | Case5 | C062 | 24a | V1pcLF | 0 | 0 | NA | 0 | -Inf |
| 1070 | Case5 | C062 | 24b | V1pcLF | 0 | 0 | NA | 0 | -Inf |
| 1071 | Case5 | C062 | 24c | V1pcLF | 0 | 0 | NA | 0 | -Inf |
| 1072 | Case5 | C062 | 24d | V1pcLF | 0 | 0 | NA | 0 | -Inf |
| 1073 | Case5 | C062 | 25 | V1pcLF | 0 | 0 | NA | 0 | -Inf |
| 1074 | Case5 | C062 | 29/30 | V1pcLF | 0 | 0 | NA | 0 | -Inf |
| 1075 | Case5 | C062 | 3 | V1pcLF | 0 | 0 | NA | 0 | -Inf |
| 1076 | Case5 | C062 | 31 | V1pcLF | 0 | 0 | NA | 0 | -Inf |
| 1077 | Case5 | C062 | 32 | V1pcLF | 0 | 0 | NA | 0 | -Inf |
| 1078 | Case5 | C062 | 44 | V1pcLF | 0 | 0 | NA | 0 | -Inf |
| 1079 | Case5 | C062 | 45A | V1pcLF | 0 | 0 | NA | 0 | -Inf |
| 1080 | Case5 | C062 | 45B | V1pcLF | 0 | 0 | NA | 0 | -Inf |
| 1081 | Case5 | C062 | 46d | V1pcLF | 0 | 0 | NA | 0 | -Inf |
| 1082 | Case5 | C062 | 46v | V1pcLF | 0 | 0 | NA | 0 | -Inf |
| 1083 | Case5 | C062 | 5 | V1pcLF | 0 | 0 | NA | 0 | -Inf |
| 1084 | Case5 | C062 | 7A | V1pcLF | 0 | 0 | NA | 0 | -Inf |

| | Case | MONKEY | SOURCE | TARGET | TOT | SUP | SLN | FLN | lgFLN |
|---|---|---|---|---|---|---|---|---|---|
| 1085 | Case5 | C062 | 7B | V1pcLF | 0 | 0 | NA | 0 | -Inf |
| 1086 | Case5 | C062 | 7m | V1pcLF | 0 | 0 | NA | 0 | -Inf |
| 1087 | Case5 | C062 | 7op | V1pcLF | 0 | 0 | NA | 0 | -Inf |
| 1088 | Case5 | C062 | 8B | V1pcLF | 0 | 0 | NA | 0 | -Inf |
| 1089 | Case5 | C062 | 8l | V1pcLF | 0 | 0 | NA | 0 | -Inf |
| 1090 | Case5 | C062 | 8m | V1pcLF | 0 | 0 | NA | 0 | -Inf |
| 1091 | Case5 | C062 | 8r | V1pcLF | 0 | 0 | NA | 0 | -Inf |
| 1092 | Case5 | C062 | 9 | V1pcLF | 0 | 0 | NA | 0 | -Inf |
| 1093 | Case5 | C062 | 9/46d | V1pcLF | 0 | 0 | NA | 0 | -Inf |
| 1094 | Case5 | C062 | 9/46v | V1pcLF | 0 | 0 | NA | 0 | -Inf |
| 1095 | Case5 | C062 | AIP | V1pcLF | 0 | 0 | NA | 0 | -Inf |
| 1096 | Case5 | C062 | CORE | V1pcLF | 0 | 0 | NA | 0 | -Inf |
| 1097 | Case5 | C062 | DP | V1pcLF | 0 | 0 | NA | 0 | -Inf |
| 1098 | Case5 | C062 | ENTO | V1pcLF | 0 | 0 | NA | 0 | -Inf |
| 1099 | Case5 | C062 | F1 | V1pcLF | 0 | 0 | NA | 0 | -Inf |
| 1100 | Case5 | C062 | F2 | V1pcLF | 0 | 0 | NA | 0 | -Inf |
| 1101 | Case5 | C062 | F3 | V1pcLF | 0 | 0 | NA | 0 | -Inf |
| 1102 | Case5 | C062 | F4 | V1pcLF | 0 | 0 | NA | 0 | -Inf |
| 1103 | Case5 | C062 | F5 | V1pcLF | 0 | 0 | NA | 0 | -Inf |
| 1104 | Case5 | C062 | F6 | V1pcLF | 0 | 0 | NA | 0 | -Inf |
| 1105 | Case5 | C062 | F7 | V1pcLF | 0 | 0 | NA | 0 | -Inf |
| 1106 | Case5 | C062 | FST | V1pcLF | 62 | 0 | 0 | 0,000234668 | -3,629545233 |
| 1107 | Case5 | C062 | Gu | V1pcLF | 0 | 0 | NA | 0 | -Inf |
| 1108 | Case5 | C062 | INSULA | V1pcLF | 0 | 0 | NA | 0 | -Inf |
| 1109 | Case5 | C062 | IPa | V1pcLF | 18 | 0 | 0 | 6,81295597127204E-05 | -4,166664418 |
| 1110 | Case5 | C062 | LB | V1pcLF | 0 | 0 | NA | 0 | -Inf |
| 1111 | Case5 | C062 | LIP | V1pcLF | 0 | 0 | NA | 0 | -Inf |
| 1112 | Case5 | C062 | MB | V1pcLF | 0 | 0 | NA | 0 | -Inf |
| 1113 | Case5 | C062 | MIP | V1pcLF | 0 | 0 | NA | 0 | -Inf |
| 1114 | Case5 | C062 | MST | V1pcLF | 76 | 0 | 0 | 0,000287658 | -3,541123331 |
| 1115 | Case5 | C062 | MTc | V1pcLF | 336 | 4 | 0,011904762 | 0,001271752 | -2,895597645 |
| 1116 | Case5 | C062 | MTp | V1pcLF | 1206 | 26 | 0,021558872 | 0,004564681 | -2,340589615 |
| 1117 | Case5 | C062 | OPAl | V1pcLF | 0 | 0 | NA | 0 | -Inf |
| 1118 | Case5 | C062 | OPro | V1pcLF | 0 | 0 | NA | 0 | -Inf |
| 1119 | Case5 | C062 | PBc | V1pcLF | 2 | 0 | 0 | 7,56995107919115E-06 | -5,120906927 |
| 1120 | Case5 | C062 | PBr | V1pcLF | 0 | 0 | NA | 0 | -Inf |
| 1121 | Case5 | C062 | PERI | V1pcLF | 18 | 0 | 0 | 6,81295597127204E-05 | -4,166664418 |
| 1122 | Case5 | C062 | PGa | V1pcLF | 38 | 0 | 0 | 0,000143829 | -3,842153326 |
| 1123 | Case5 | C062 | Pi | V1pcLF | 0 | 0 | NA | 0 | -Inf |
| 1124 | Case5 | C062 | PIP | V1pcLF | 6 | 0 | 0 | 2,27098532375735E-05 | -4,643785672 |
| 1125 | Case5 | C062 | Pir | V1pcLF | 0 | 0 | NA | 0 | -Inf |
| 1126 | Case5 | C062 | POLE | V1pcLF | 0 | 0 | NA | 0 | -Inf |
| 1127 | Case5 | C062 | Pro.St | V1pcLF | 0 | 0 | NA | 0 | -Inf |
| 1128 | Case5 | C062 | ProM | V1pcLF | 0 | 0 | NA | 0 | -Inf |
| 1129 | Case5 | C062 | SII | V1pcLF | 0 | 0 | NA | 0 | -Inf |
| 1130 | Case5 | C062 | STPc | V1pcLF | 6 | 0 | 0 | 2,27098532375735E-05 | -4,643785672 |
| 1131 | Case5 | C062 | STPi | V1pcLF | 4 | 0 | 0 | 1,51399021583823E-05 | -4,819876931 |
| 1132 | Case5 | C062 | STPr | V1pcLF | 0 | 0 | NA | 0 | -Inf |
| 1133 | Case5 | C062 | SUBI | V1pcLF | 0 | 0 | NA | 0 | -Inf |
| 1134 | Case5 | C062 | TEa/ma | V1pcLF | 0 | 0 | NA | 0 | -Inf |
| 1135 | Case5 | C062 | TEa/mp | V1pcLF | 4 | 0 | 0 | 1,51399021583823E-05 | -4,819876931 |
| 1136 | Case5 | C062 | TEad | V1pcLF | 0 | 0 | NA | 0 | -Inf |
| 1137 | Case5 | C062 | TEav | V1pcLF | 18 | 0 | 0 | 6,81295597127204E-05 | -4,166664418 |
| 1138 | Case5 | C062 | TEO | V1pcLF | 154 | 0 | 0 | 0,000582886 | -3,234416202 |
| 1139 | Case5 | C062 | TEOm | V1pcLF | 20 | 0 | 0 | 7,56995107919115E-05 | -4,120906927 |
| 1140 | Case5 | C062 | TEpd | V1pcLF | 46 | 4 | 0,086956522 | 0,000174109 | -3,759179091 |
| 1141 | Case5 | C062 | TEpv | V1pcLF | 338 | 8 | 0,023668639 | 0,001279322 | -2,893020223 |
| 1142 | Case5 | C062 | TH_TF | V1pcLF | 136 | 2 | 0,014705882 | 0,000514757 | -3,288398014 |
| 1143 | Case5 | C062 | TPt | V1pcLF | 0 | 0 | NA | 0 | -Inf |
| 1144 | Case5 | C062 | V1c | V1pcLF | 666 | 24 | 0,036036036 | 0,002520794 | -2,598462694 |
| 1145 | Case5 | C062 | V1fpLF | V1pcLF | 0 | 0 | NA | 0 | -Inf |
| 1146 | Case5 | C062 | V1fpUF | V1pcLF | 0 | 0 | NA | 0 | -Inf |
| 1147 | Case5 | C062 | V1pcLF | V1pcLF | 0 | 0 | NA | 0 | -Inf |
| 1148 | Case5 | C062 | V1pcUF | V1pcLF | 0 | 0 | NA | 0 | -Inf |
| 1149 | Case5 | C062 | V2c | V1pcLF | 2690 | 1982 | 0,736802974 | 0,010181584 | -1,992184643 |
| 1150 | Case5 | C062 | V2fpLF | V1pcLF | 0 | 0 | NA | 0 | -Inf |
| 1151 | Case5 | C062 | V2fpUF | V1pcLF | 0 | 0 | NA | 0 | -Inf |
| 1152 | Case5 | C062 | V2pcLF | V1pcLF | 19608 | 10222 | 0,521317829 | 0,0742158 | -1,129503625 |
| 1153 | Case5 | C062 | V2pcUF | V1pcLF | 0 | 0 | NA | 0 | -Inf |
| 1154 | Case5 | C062 | V3A | V1pcLF | 12 | 0 | 0 | 4,54197064751469E-05 | -4,342755677 |
| 1155 | Case5 | C062 | V3c | V1pcLF | 62 | 0 | 0 | 0,000234668 | -3,629545233 |
| 1156 | Case5 | C062 | V3fpLF | V1pcLF | 0 | 0 | NA | 0 | -Inf |
| 1157 | Case5 | C062 | V3fpUF | V1pcLF | 0 | 0 | NA | 0 | -Inf |
| 1158 | Case5 | C062 | V3pcLF | V1pcLF | 2914 | 752 | 0,258064516 | 0,011029419 | -1,957447375 |
| 1159 | Case5 | C062 | V3pcUF | V1pcLF | 0 | 0 | NA | 0 | -Inf |

| | Case | MONKEY | SOURCE | TARGET | TOT | SUP | SLN | FLN | lgFLN |
|---|---|---|---|---|---|---|---|---|---|
| 1160 | Case5 | C062 | V4c | V1pcLF | 86 | 2 | 0,023255814 | 0,000325508 | -3,487438472 |
| 1161 | Case5 | C062 | V4LF | V1pcLF | 3246 | 158 | 0,048675293 | 0,012286031 | -1,910588407 |
| 1162 | Case5 | C062 | V4t | V1pcLF | 150 | 14 | 0,093333333 | 0,000567746 | -3,245845664 |
| 1163 | Case5 | C062 | V4UF | V1pcLF | 30 | 0 | 0 | 0,000113549 | -3,944815668 |
| 1164 | Case5 | C062 | V6 | V1pcLF | 0 | 0 | NA | 0 | -Inf |
| 1165 | Case5 | C062 | V6A | V1pcLF | 0 | 0 | NA | 0 | -Inf |
| 1166 | Case5 | C062 | VIP | V1pcLF | 0 | 0 | NA | 0 | -Inf |
| 2015 | Case6 | M122RH | 1 | V1pcLF | 0 | 0 | NA | 0 | -Inf |
| 2016 | Case6 | M122RH | 10 | V1pcLF | 0 | 0 | NA | 0 | -Inf |
| 2017 | Case6 | M122RH | 11 | V1pcLF | 0 | 0 | NA | 0 | -Inf |
| 2018 | Case6 | M122RH | 12 | V1pcLF | 0 | 0 | NA | 0 | -Inf |
| 2019 | Case6 | M122RH | 13 | V1pcLF | 0 | 0 | NA | 0 | -Inf |
| 2020 | Case6 | M122RH | 14 | V1pcLF | 0 | 0 | NA | 0 | -Inf |
| 2021 | Case6 | M122RH | 2 | V1pcLF | 0 | 0 | NA | 0 | -Inf |
| 2022 | Case6 | M122RH | 23 | V1pcLF | 0 | 0 | NA | 0 | -Inf |
| 2023 | Case6 | M122RH | 24a | V1pcLF | 0 | 0 | NA | 0 | -Inf |
| 2024 | Case6 | M122RH | 24b | V1pcLF | 0 | 0 | NA | 0 | -Inf |
| 2025 | Case6 | M122RH | 24c | V1pcLF | 0 | 0 | NA | 0 | -Inf |
| 2026 | Case6 | M122RH | 24d | V1pcLF | 0 | 0 | NA | 0 | -Inf |
| 2027 | Case6 | M122RH | 25 | V1pcLF | 0 | 0 | NA | 0 | -Inf |
| 2028 | Case6 | M122RH | 29/30 | V1pcLF | 0 | 0 | NA | 0 | -Inf |
| 2029 | Case6 | M122RH | 3 | V1pcLF | 0 | 0 | NA | 0 | -Inf |
| 2030 | Case6 | M122RH | 31 | V1pcLF | 0 | 0 | NA | 0 | -Inf |
| 2031 | Case6 | M122RH | 32 | V1pcLF | 0 | 0 | NA | 0 | -Inf |
| 2032 | Case6 | M122RH | 44 | V1pcLF | 0 | 0 | NA | 0 | -Inf |
| 2033 | Case6 | M122RH | 45A | V1pcLF | 0 | 0 | NA | 0 | -Inf |
| 2034 | Case6 | M122RH | 45B | V1pcLF | 0 | 0 | NA | 0 | -Inf |
| 2035 | Case6 | M122RH | 46d | V1pcLF | 0 | 0 | NA | 0 | -Inf |
| 2036 | Case6 | M122RH | 46v | V1pcLF | 0 | 0 | NA | 0 | -Inf |
| 2037 | Case6 | M122RH | 5 | V1pcLF | 0 | 0 | NA | 0 | -Inf |
| 2038 | Case6 | M122RH | 7A | V1pcLF | 4 | 0 | 0 | 1,51399021583823E-05 | -4,819876931 |
| 2039 | Case6 | M122RH | 7B | V1pcLF | 0 | 0 | NA | 0 | -Inf |
| 2040 | Case6 | M122RH | 7m | V1pcLF | 0 | 0 | NA | 0 | -Inf |
| 2041 | Case6 | M122RH | 7op | V1pcLF | 0 | 0 | NA | 0 | -Inf |
| 2042 | Case6 | M122RH | 8B | V1pcLF | 0 | 0 | NA | 0 | -Inf |
| 2043 | Case6 | M122RH | 8l | V1pcLF | 0 | 0 | NA | 0 | -Inf |
| 2044 | Case6 | M122RH | 8m | V1pcLF | 0 | 0 | NA | 0 | -Inf |
| 2045 | Case6 | M122RH | 8r | V1pcLF | 0 | 0 | NA | 0 | -Inf |
| 2046 | Case6 | M122RH | 9 | V1pcLF | 0 | 0 | NA | 0 | -Inf |
| 2047 | Case6 | M122RH | 9/46d | V1pcLF | 0 | 0 | NA | 0 | -Inf |
| 2048 | Case6 | M122RH | 9/46v | V1pcLF | 0 | 0 | NA | 0 | -Inf |
| 2049 | Case6 | M122RH | AIP | V1pcLF | 0 | 0 | NA | 0 | -Inf |
| 2050 | Case6 | M122RH | CORE | V1pcLF | 4 | 0 | 0 | 1,51399021583823E-05 | -4,819876931 |
| 2051 | Case6 | M122RH | DP | V1pcLF | 233 | 26 | 0,111587983 | 0,000881899 | -3,054581002 |
| 2052 | Case6 | M122RH | ENTO | V1pcLF | 0 | 0 | NA | 0 | -Inf |
| 2053 | Case6 | M122RH | F1 | V1pcLF | 0 | 0 | NA | 0 | -Inf |
| 2054 | Case6 | M122RH | F2 | V1pcLF | 2 | 0 | 0 | 7,56995107919115E-06 | -5,120906927 |
| 2055 | Case6 | M122RH | F3 | V1pcLF | 0 | 0 | NA | 0 | -Inf |
| 2056 | Case6 | M122RH | F4 | V1pcLF | 0 | 0 | NA | 0 | -Inf |
| 2057 | Case6 | M122RH | F5 | V1pcLF | 0 | 0 | NA | 0 | -Inf |
| 2058 | Case6 | M122RH | F6 | V1pcLF | 0 | 0 | NA | 0 | -Inf |
| 2059 | Case6 | M122RH | F7 | V1pcLF | 0 | 0 | NA | 0 | -Inf |
| 2060 | Case6 | M122RH | FST | V1pcLF | 212 | 9 | 0,04245283 | 0,000802415 | -3,095601062 |
| 2061 | Case6 | M122RH | Gu | V1pcLF | 0 | 0 | NA | 0 | -Inf |
| 2062 | Case6 | M122RH | INSULA | V1pcLF | 0 | 0 | NA | 0 | -Inf |
| 2063 | Case6 | M122RH | IPa | V1pcLF | 16 | 0 | 0 | 6,05596086335292E-05 | -4,21781694 |
| 2064 | Case6 | M122RH | LB | V1pcLF | 23 | 0 | 0 | 8,70544374106982E-05 | -4,060209087 |
| 2065 | Case6 | M122RH | LIP | V1pcLF | 168 | 7 | 0,041666667 | 0,000635876 | -3,196627641 |
| 2066 | Case6 | M122RH | MB | V1pcLF | 12 | 0 | 0 | 4,54197064751469E-05 | -4,342755677 |
| 2067 | Case6 | M122RH | MIP | V1pcLF | 3 | 0 | 0 | 1,13549266187867E-05 | -4,944815668 |
| 2068 | Case6 | M122RH | MST | V1pcLF | 512 | 0 | 0 | 0,001937907 | -2,712666962 |
| 2069 | Case6 | M122RH | MTc | V1pcLF | 198 | 0 | 0 | 0,000749425 | -3,125271733 |
| 2070 | Case6 | M122RH | MTp | V1pcLF | 6347 | 345 | 0,054356389 | 0,02402324 | -1,619368424 |
| 2071 | Case6 | M122RH | OPAl | V1pcLF | 0 | 0 | NA | 0 | -Inf |
| 2072 | Case6 | M122RH | OPro | V1pcLF | 0 | 0 | NA | 0 | -Inf |
| 2073 | Case6 | M122RH | PBc | V1pcLF | 63 | 0 | 0 | 0,000238453 | -3,622596373 |
| 2074 | Case6 | M122RH | PBr | V1pcLF | 0 | 0 | NA | 0 | -Inf |
| 2075 | Case6 | M122RH | PERI | V1pcLF | 62 | 0 | 0 | 0,000234668 | -3,629545233 |
| 2076 | Case6 | M122RH | PGa | V1pcLF | 100 | 15 | 0,15 | 0,000378498 | -3,421936923 |
| 2077 | Case6 | M122RH | Pi | V1pcLF | 2 | 0 | 0 | 7,56995107919115E-06 | -5,120906927 |
| 2078 | Case6 | M122RH | PIP | V1pcLF | 340 | 15 | 0,044117647 | 0,001286892 | -2,890458006 |
| 2079 | Case6 | M122RH | Pir | V1pcLF | 0 | 0 | NA | 0 | -Inf |
| 2080 | Case6 | M122RH | POLE | V1pcLF | 6 | 0 | 0 | 2,27098532375735E-05 | -4,643785672 |
| 2081 | Case6 | M122RH | Pro.St | V1pcLF | 0 | 0 | NA | 0 | -Inf |
| 2082 | Case6 | M122RH | ProM | V1pcLF | 0 | 0 | NA | 0 | -Inf |

| | Case | MONKEY | SOURCE | TARGET | TOT | SUP | SLN | FLN | lgFLN |
|---|---|---|---|---|---|---|---|---|---|
| 2083 | Case6 | M122RH | SII | V1pcLF | 0 | 0 | NA | 0 | -Inf |
| 2084 | Case6 | M122RH | STPc | V1pcLF | 247 | 0 | 0 | 0,000934889 | -3,02923997 |
| 2085 | Case6 | M122RH | STPi | V1pcLF | 104 | 4 | 0,038461538 | 0,000393637 | -3,404903583 |
| 2086 | Case6 | M122RH | STPr | V1pcLF | 0 | 0 | NA | 0 | -Inf |
| 2087 | Case6 | M122RH | SUBI | V1pcLF | 0 | 0 | NA | 0 | -Inf |
| 2088 | Case6 | M122RH | TEa/ma | V1pcLF | 0 | 0 | NA | 0 | -Inf |
| 2089 | Case6 | M122RH | TEa/mp | V1pcLF | 48 | 0 | 0 | 0,000181679 | -3,740695685 |
| 2090 | Case6 | M122RH | TEad | V1pcLF | 47 | 0 | 0 | 0,000177894 | -3,749839065 |
| 2091 | Case6 | M122RH | TEav | V1pcLF | 68 | 0 | 0 | 0,000257378 | -3,58942801 |
| 2092 | Case6 | M122RH | TEO | V1pcLF | 826 | 4 | 0,004842615 | 0,00312639 | -2,504956875 |
| 2093 | Case6 | M122RH | TEOm | V1pcLF | 82 | 4 | 0,048780488 | 0,000310368 | -3,50812307 |
| 2094 | Case6 | M122RH | TEpd | V1pcLF | 244 | 0 | 0 | 0,000923534 | -3,034547096 |
| 2095 | Case6 | M122RH | TEpv | V1pcLF | 2514 | 30 | 0,011933174 | 0,009515429 | -2,021571649 |
| 2096 | Case6 | M122RH | TH_TF | V1pcLF | 759 | 4 | 0,005270092 | 0,002872796 | -2,541695147 |
| 2097 | Case6 | M122RH | TPt | V1pcLF | 12 | 0 | 0 | 4,54197064751469E-05 | -4,342755677 |
| 2098 | Case6 | M122RH | V1c | V1pcLF | 0 | 0 | NA | 0 | -Inf |
| 2099 | Case6 | M122RH | V1fpLF | V1pcLF | 0 | 0 | NA | 0 | -Inf |
| 2100 | Case6 | M122RH | V1fpUF | V1pcLF | 0 | 0 | NA | 0 | -Inf |
| 2101 | Case6 | M122RH | V1pcLF | V1pcLF | 0 | 0 | NA | 0 | -Inf |
| 2102 | Case6 | M122RH | V1pcUF | V1pcLF | 0 | 0 | NA | 0 | -Inf |
| 2103 | Case6 | M122RH | V2c | V1pcLF | 154 | 6 | 0,038961039 | 0,000582886 | -3,234416202 |
| 2104 | Case6 | M122RH | V2fpLF | V1pcLF | 2281 | 180 | 0,078912758 | 0,008633529 | -2,063811638 |
| 2105 | Case6 | M122RH | V2fpUF | V1pcLF | 35 | 0 | 0 | 0,000132474 | -3,877868878 |
| 2106 | Case6 | M122RH | V2pcLF | V1pcLF | 84222 | 37706 | 0,447697751 | 0,31877821 | -0,496511372 |
| 2107 | Case6 | M122RH | V2pcUF | V1pcLF | 75 | 4 | 0,053333333 | 0,000283873 | -3,546875659 |
| 2108 | Case6 | M122RH | V3A | V1pcLF | 404 | 14 | 0,034653465 | 0,00152913 | -2,815555558 |
| 2109 | Case6 | M122RH | V3c | V1pcLF | 172 | 25 | 0,145348837 | 0,000651016 | -3,186408476 |
| 2110 | Case6 | M122RH | V3fpLF | V1pcLF | 972 | 122 | 0,125514403 | 0,003678996 | -2,434270658 |
| 2111 | Case6 | M122RH | V3fpUF | V1pcLF | 16 | 0 | 0 | 6,05596086335292E-05 | -4,21781694 |
| 2112 | Case6 | M122RH | V3pcLF | V1pcLF | 2327 | 384 | 0,165019338 | 0,008807638 | -2,055140539 |
| 2113 | Case6 | M122RH | V3pcUF | V1pcLF | 16 | 2 | 0,125 | 6,05596086335292E-05 | -4,21781694 |
| 2114 | Case6 | M122RH | V4c | V1pcLF | 12 | 0 | 0 | 4,54197064751469E-05 | -4,342755677 |
| 2115 | Case6 | M122RH | V4LF | V1pcLF | 15350 | 4756 | 0,309837134 | 0,058099375 | -1,235828543 |
| 2116 | Case6 | M122RH | V4t | V1pcLF | 378 | 8 | 0,021164021 | 0,001430721 | -2,844445123 |
| 2117 | Case6 | M122RH | V4UF | V1pcLF | 853 | 45 | 0,052754982 | 0,003228584 | -2,490987892 |
| 2118 | Case6 | M122RH | V6 | V1pcLF | 6 | 0 | 0 | 2,27098532375735E-05 | -4,643785672 |
| 2119 | Case6 | M122RH | V6A | V1pcLF | 68 | 2 | 0,029411765 | 0,000257378 | -3,58942801 |
| 2120 | Case6 | M122RH | VIP | V1pcLF | 25 | 0 | 0 | 9,46243884898894E-05 | -4,023996914 |
| 955 | Case7 | C060 | 1 | V1pcLF | 0 | 0 | NA | 0 | -Inf |
| 956 | Case7 | C060 | 10 | V1pcLF | 0 | 0 | NA | 0 | -Inf |
| 957 | Case7 | C060 | 11 | V1pcLF | 0 | 0 | NA | 0 | -Inf |
| 958 | Case7 | C060 | 12 | V1pcLF | 0 | 0 | NA | 0 | -Inf |
| 959 | Case7 | C060 | 13 | V1pcLF | 0 | 0 | NA | 0 | -Inf |
| 960 | Case7 | C060 | 14 | V1pcLF | 0 | 0 | NA | 0 | -Inf |
| 961 | Case7 | C060 | 2 | V1pcLF | 0 | 0 | NA | 0 | -Inf |
| 962 | Case7 | C060 | 23 | V1pcLF | 0 | 0 | NA | 0 | -Inf |
| 963 | Case7 | C060 | 24a | V1pcLF | 0 | 0 | NA | 0 | -Inf |
| 964 | Case7 | C060 | 24b | V1pcLF | 0 | 0 | NA | 0 | -Inf |
| 965 | Case7 | C060 | 24c | V1pcLF | 0 | 0 | NA | 0 | -Inf |
| 966 | Case7 | C060 | 24d | V1pcLF | 0 | 0 | NA | 0 | -Inf |
| 967 | Case7 | C060 | 25 | V1pcLF | 0 | 0 | NA | 0 | -Inf |
| 968 | Case7 | C060 | 29/30 | V1pcLF | 0 | 0 | NA | 0 | -Inf |
| 969 | Case7 | C060 | 3 | V1pcLF | 0 | 0 | NA | 0 | -Inf |
| 970 | Case7 | C060 | 31 | V1pcLF | 0 | 0 | NA | 0 | -Inf |
| 971 | Case7 | C060 | 32 | V1pcLF | 0 | 0 | NA | 0 | -Inf |
| 972 | Case7 | C060 | 44 | V1pcLF | 0 | 0 | NA | 0 | -Inf |
| 973 | Case7 | C060 | 45A | V1pcLF | 0 | 0 | NA | 0 | -Inf |
| 974 | Case7 | C060 | 45B | V1pcLF | 0 | 0 | NA | 0 | -Inf |
| 975 | Case7 | C060 | 46d | V1pcLF | 0 | 0 | NA | 0 | -Inf |
| 976 | Case7 | C060 | 46v | V1pcLF | 0 | 0 | NA | 0 | -Inf |
| 977 | Case7 | C060 | 5 | V1pcLF | 0 | 0 | NA | 0 | -Inf |
| 978 | Case7 | C060 | 7A | V1pcLF | 0 | 0 | NA | 0 | -Inf |
| 979 | Case7 | C060 | 7B | V1pcLF | 0 | 0 | NA | 0 | -Inf |
| 980 | Case7 | C060 | 7m | V1pcLF | 0 | 0 | NA | 0 | -Inf |
| 981 | Case7 | C060 | 7op | V1pcLF | 0 | 0 | NA | 0 | -Inf |
| 982 | Case7 | C060 | 8B | V1pcLF | 0 | 0 | NA | 0 | -Inf |
| 983 | Case7 | C060 | 8l | V1pcLF | 0 | 0 | NA | 0 | -Inf |
| 984 | Case7 | C060 | 8m | V1pcLF | 0 | 0 | NA | 0 | -Inf |
| 985 | Case7 | C060 | 8r | V1pcLF | 0 | 0 | NA | 0 | -Inf |
| 986 | Case7 | C060 | 9 | V1pcLF | 0 | 0 | NA | 0 | -Inf |
| 987 | Case7 | C060 | 9/46d | V1pcLF | 0 | 0 | NA | 0 | -Inf |
| 988 | Case7 | C060 | 9/46v | V1pcLF | 0 | 0 | NA | 0 | -Inf |
| 989 | Case7 | C060 | AIP | V1pcLF | 0 | 0 | NA | 0 | -Inf |
| 990 | Case7 | C060 | CORE | V1pcLF | 0 | 0 | NA | 0 | -Inf |
| 991 | Case7 | C060 | DP | V1pcLF | 0 | 0 | NA | 0 | -Inf |

| | Case | MONKEY | SOURCE | TARGET | TOT | SUP | SLN | FLN | lgFLN |
|---|---|---|---|---|---|---|---|---|---|
| 992 | Case7 | C060 | ENTO | V1pcLF | 0 | 0 | NA | 0 | -Inf |
| 993 | Case7 | C060 | F1 | V1pcLF | 0 | 0 | NA | 0 | -Inf |
| 994 | Case7 | C060 | F2 | V1pcLF | 0 | 0 | NA | 0 | -Inf |
| 995 | Case7 | C060 | F3 | V1pcLF | 0 | 0 | NA | 0 | -Inf |
| 996 | Case7 | C060 | F4 | V1pcLF | 0 | 0 | NA | 0 | -Inf |
| 997 | Case7 | C060 | F5 | V1pcLF | 0 | 0 | NA | 0 | -Inf |
| 998 | Case7 | C060 | F6 | V1pcLF | 0 | 0 | NA | 0 | -Inf |
| 999 | Case7 | C060 | F7 | V1pcLF | 0 | 0 | NA | 0 | -Inf |
| 1000 | Case7 | C060 | FST | V1pcLF | 102 | 2 | 0,019607843 | 0,000386068 | -3,413336751 |
| 1001 | Case7 | C060 | Gu | V1pcLF | 0 | 0 | NA | 0 | -Inf |
| 1002 | Case7 | C060 | INSULA | V1pcLF | 5 | 0 | 0 | 1,89248776979779E-05 | -4,722966918 |
| 1003 | Case7 | C060 | IPa | V1pcLF | 3 | 0 | 0 | 1,13549266187867E-05 | -4,944815668 |
| 1004 | Case7 | C060 | LB | V1pcLF | 8 | 0 | 0 | 3,02798043167646E-05 | -4,518846936 |
| 1005 | Case7 | C060 | LIP | V1pcLF | 2 | 0 | 0 | 7,56995107919115E-06 | -5,120906927 |
| 1006 | Case7 | C060 | MB | V1pcLF | 2 | 0 | 0 | 7,56995107919115E-06 | -5,120906927 |
| 1007 | Case7 | C060 | MIP | V1pcLF | 0 | 0 | NA | 0 | -Inf |
| 1008 | Case7 | C060 | MST | V1pcLF | 136 | 0 | 0 | 0,000514757 | -3,288398014 |
| 1009 | Case7 | C060 | MTc | V1pcLF | 226 | 0 | 0 | 0,000855404 | -3,067828484 |
| 1010 | Case7 | C060 | MTp | V1pcLF | 1340 | 40 | 0,029850746 | 0,005071867 | -2,294832124 |
| 1011 | Case7 | C060 | OPAl | V1pcLF | 0 | 0 | NA | 0 | -Inf |
| 1012 | Case7 | C060 | OPro | V1pcLF | 0 | 0 | NA | 0 | -Inf |
| 1013 | Case7 | C060 | PBc | V1pcLF | 51,5 | 0 | 0 | 0,000194926 | -3,710129694 |
| 1014 | Case7 | C060 | PBr | V1pcLF | 2 | 0 | 0 | 7,56995107919115E-06 | -5,120906927 |
| 1015 | Case7 | C060 | PERI | V1pcLF | 40 | 0 | 0 | 0,000151399 | -3,819876931 |
| 1016 | Case7 | C060 | PGa | V1pcLF | 12 | 0 | 0 | 4,54197064751469E-05 | -4,342755677 |
| 1017 | Case7 | C060 | Pi | V1pcLF | 0 | 0 | NA | 0 | -Inf |
| 1018 | Case7 | C060 | PIP | V1pcLF | 34 | 0 | 0 | 0,000128689 | -3,890458006 |
| 1019 | Case7 | C060 | Pir | V1pcLF | 0 | 0 | NA | 0 | -Inf |
| 1020 | Case7 | C060 | POLE | V1pcLF | 0 | 0 | NA | 0 | -Inf |
| 1021 | Case7 | C060 | Pro.St | V1pcLF | 0 | 0 | NA | 0 | -Inf |
| 1022 | Case7 | C060 | ProM | V1pcLF | 0 | 0 | NA | 0 | -Inf |
| 1023 | Case7 | C060 | SII | V1pcLF | 0 | 0 | NA | 0 | -Inf |
| 1024 | Case7 | C060 | STPc | V1pcLF | 102 | 0 | 0 | 0,000386068 | -3,413336751 |
| 1025 | Case7 | C060 | STPi | V1pcLF | 19,5 | 0 | 0 | 7,38070230221137E-05 | -4,131902311 |
| 1026 | Case7 | C060 | STPr | V1pcLF | 0 | 0 | NA | 0 | -Inf |
| 1027 | Case7 | C060 | SUBI | V1pcLF | 0 | 0 | NA | 0 | -Inf |
| 1028 | Case7 | C060 | TEa/ma | V1pcLF | 0 | 0 | NA | 0 | -Inf |
| 1029 | Case7 | C060 | TEa/mp | V1pcLF | 5,5 | 0 | 0 | 2,08173654677757E-05 | -4,681574233 |
| 1030 | Case7 | C060 | TEad | V1pcLF | 0 | 0 | NA | 0 | -Inf |
| 1031 | Case7 | C060 | TEav | V1pcLF | 2 | 0 | 0 | 7,56995107919115E-06 | -5,120906927 |
| 1032 | Case7 | C060 | TEO | V1pcLF | 410 | 2 | 0,004878049 | 0,00155184 | -2,809153066 |
| 1033 | Case7 | C060 | TEOm | V1pcLF | 64 | 0 | 0 | 0,000242238 | -3,615756949 |
| 1034 | Case7 | C060 | TEpd | V1pcLF | 8 | 0 | 0 | 3,02798043167646E-05 | -4,518846936 |
| 1035 | Case7 | C060 | TEpv | V1pcLF | 388,5 | 2 | 0,005148005 | 0,001470463 | -2,8325459 |
| 1036 | Case7 | C060 | TH_TF | V1pcLF | 332 | 0 | 0 | 0,001256612 | -2,900798839 |
| 1037 | Case7 | C060 | TPt | V1pcLF | 8 | 0 | 0 | 3,02798043167646E-05 | -4,518846936 |
| 1038 | Case7 | C060 | V1c | V1pcLF | 0 | 0 | NA | 0 | -Inf |
| 1039 | Case7 | C060 | V1fpLF | V1pcLF | 5848 | 3950 | 0,675444596 | 0,022134537 | -1,654929559 |
| 1040 | Case7 | C060 | V1fpUF | V1pcLF | 12394 | 12318 | 0,993868001 | 0,046910987 | -1,328725431 |
| 1041 | Case7 | C060 | V1pcLF | V1pcLF | 0 | 0 | NA | 0 | -Inf |
| 1042 | Case7 | C060 | V1pcUF | V1pcLF | 1282 | 1214 | 0,946957878 | 0,004852339 | -2,314048898 |
| 1043 | Case7 | C060 | V2c | V1pcLF | 25204 | 11142 | 0,442072687 | 0,095396523 | -1,020467452 |
| 1044 | Case7 | C060 | V2fpLF | V1pcLF | 14 | 0 | 0 | 5,29896575543381E-05 | -4,275808887 |
| 1045 | Case7 | C060 | V2fpUF | V1pcLF | 32 | 4 | 0,125 | 0,000121119 | -3,916786944 |
| 1046 | Case7 | C060 | V2pcLF | V1pcLF | 8598 | 3622 | 0,421260758 | 0,03254322 | -1,487539482 |
| 1047 | Case7 | C060 | V2pcUF | V1pcLF | 1418 | 338 | 0,238363893 | 0,005367095 | -2,270260692 |
| 1048 | Case7 | C060 | V3A | V1pcLF | 0 | 0 | NA | 0 | -Inf |
| 1049 | Case7 | C060 | V3c | V1pcLF | 82 | 0 | 0 | 0,000310368 | -3,50812307 |
| 1050 | Case7 | C060 | V3fpLF | V1pcLF | 80 | 6 | 0,075 | 0,000302798 | -3,518846936 |
| 1051 | Case7 | C060 | V3fpUF | V1pcLF | 44 | 0 | 0 | 0,000166539 | -3,778484246 |
| 1052 | Case7 | C060 | V3pcLF | V1pcLF | 542 | 6 | 0,011070111 | 0,002051457 | -2,687937636 |
| 1053 | Case7 | C060 | V3pcUF | V1pcLF | 314 | 44 | 0,140127389 | 0,001188482 | -2,925007275 |
| 1054 | Case7 | C060 | V4c | V1pcLF | 2 | 0 | 0 | 7,56995107919115E-06 | -5,120906927 |
| 1055 | Case7 | C060 | V4LF | V1pcLF | 408 | 20 | 0,049019608 | 0,00154427 | -2,81127676 |
| 1056 | Case7 | C060 | V4t | V1pcLF | 148 | 22 | 0,148648649 | 0,000560176 | -3,251675207 |
| 1057 | Case7 | C060 | V4UF | V1pcLF | 1030 | 100 | 0,097087379 | 0,003898525 | -2,409099698 |
| 1058 | Case7 | C060 | V6 | V1pcLF | 0 | 0 | NA | 0 | -Inf |
| 1059 | Case7 | C060 | V6A | V1pcLF | 2 | 0 | 0 | 7,56995107919115E-06 | -5,120906927 |
| 1060 | Case7 | C060 | VIP | V1pcLF | 0 | 0 | NA | 0 | -Inf |
| 1167 | Case8 | C081 | 1 | V1pcLF | 0 | 0 | NA | 0 | -Inf |
| 1168 | Case8 | C081 | 10 | V1pcLF | 0 | 0 | NA | 0 | -Inf |
| 1169 | Case8 | C081 | 11 | V1pcLF | 0 | 0 | NA | 0 | -Inf |
| 1170 | Case8 | C081 | 12 | V1pcLF | 0 | 0 | NA | 0 | -Inf |
| 1171 | Case8 | C081 | 13 | V1pcLF | 0 | 0 | NA | 0 | -Inf |
| 1172 | Case8 | C081 | 14 | V1pcLF | 0 | 0 | NA | 0 | -Inf |

| | Case | MONKEY | SOURCE | TARGET | TOT | SUP | SLN | FLN | lgFLN |
|---|---|---|---|---|---|---|---|---|---|
| 1173 | Case8 | C081 | 2 | V1pcLF | 0 | 0 | NA | 0 | -Inf |
| 1174 | Case8 | C081 | 23 | V1pcLF | 0 | 0 | NA | 0 | -Inf |
| 1175 | Case8 | C081 | 24a | V1pcLF | 0 | 0 | NA | 0 | -Inf |
| 1176 | Case8 | C081 | 24b | V1pcLF | 0 | 0 | NA | 0 | -Inf |
| 1177 | Case8 | C081 | 24c | V1pcLF | 0 | 0 | NA | 0 | -Inf |
| 1178 | Case8 | C081 | 24d | V1pcLF | 0 | 0 | NA | 0 | -Inf |
| 1179 | Case8 | C081 | 25 | V1pcLF | 0 | 0 | NA | 0 | -Inf |
| 1180 | Case8 | C081 | 29/30 | V1pcLF | 0 | 0 | NA | 0 | -Inf |
| 1181 | Case8 | C081 | 3 | V1pcLF | 0 | 0 | NA | 0 | -Inf |
| 1182 | Case8 | C081 | 31 | V1pcLF | 0 | 0 | NA | 0 | -Inf |
| 1183 | Case8 | C081 | 32 | V1pcLF | 0 | 0 | NA | 0 | -Inf |
| 1184 | Case8 | C081 | 44 | V1pcLF | 0 | 0 | NA | 0 | -Inf |
| 1185 | Case8 | C081 | 45A | V1pcLF | 0 | 0 | NA | 0 | -Inf |
| 1186 | Case8 | C081 | 45B | V1pcLF | 0 | 0 | NA | 0 | -Inf |
| 1187 | Case8 | C081 | 46d | V1pcLF | 0 | 0 | NA | 0 | -Inf |
| 1188 | Case8 | C081 | 46v | V1pcLF | 0 | 0 | NA | 0 | -Inf |
| 1189 | Case8 | C081 | 5 | V1pcLF | 0 | 0 | NA | 0 | -Inf |
| 1190 | Case8 | C081 | 7A | V1pcLF | 2 | 2 | 1 | 7,56995107919115E-06 | -5,120906927 |
| 1191 | Case8 | C081 | 7B | V1pcLF | 0 | 0 | NA | 0 | -Inf |
| 1192 | Case8 | C081 | 7m | V1pcLF | 0 | 0 | NA | 0 | -Inf |
| 1193 | Case8 | C081 | 7op | V1pcLF | 0 | 0 | NA | 0 | -Inf |
| 1194 | Case8 | C081 | 8B | V1pcLF | 0 | 0 | NA | 0 | -Inf |
| 1195 | Case8 | C081 | 8l | V1pcLF | 0 | 0 | NA | 0 | -Inf |
| 1196 | Case8 | C081 | 8m | V1pcLF | 0 | 0 | NA | 0 | -Inf |
| 1197 | Case8 | C081 | 8r | V1pcLF | 0 | 0 | NA | 0 | -Inf |
| 1198 | Case8 | C081 | 9 | V1pcLF | 0 | 0 | NA | 0 | -Inf |
| 1199 | Case8 | C081 | 9/46d | V1pcLF | 0 | 0 | NA | 0 | -Inf |
| 1200 | Case8 | C081 | 9/46v | V1pcLF | 0 | 0 | NA | 0 | -Inf |
| 1201 | Case8 | C081 | AIP | V1pcLF | 0 | 0 | NA | 0 | -Inf |
| 1202 | Case8 | C081 | CORE | V1pcLF | 4 | 4 | 1 | 1,51399021583823E-05 | -4,819876931 |
| 1203 | Case8 | C081 | DP | V1pcLF | 34 | 34 | 1 | 0,000128689 | -3,890458006 |
| 1204 | Case8 | C081 | ENTO | V1pcLF | 0 | 0 | NA | 0 | -Inf |
| 1205 | Case8 | C081 | F1 | V1pcLF | 0 | 0 | NA | 0 | -Inf |
| 1206 | Case8 | C081 | F2 | V1pcLF | 0 | 0 | NA | 0 | -Inf |
| 1207 | Case8 | C081 | F3 | V1pcLF | 0 | 0 | NA | 0 | -Inf |
| 1208 | Case8 | C081 | F4 | V1pcLF | 0 | 0 | NA | 0 | -Inf |
| 1209 | Case8 | C081 | F5 | V1pcLF | 0 | 0 | NA | 0 | -Inf |
| 1210 | Case8 | C081 | F6 | V1pcLF | 0 | 0 | NA | 0 | -Inf |
| 1211 | Case8 | C081 | F7 | V1pcLF | 0 | 0 | NA | 0 | -Inf |
| 1212 | Case8 | C081 | FST | V1pcLF | 74 | 74 | 1 | 0,000280088 | -3,552705203 |
| 1213 | Case8 | C081 | Gu | V1pcLF | 0 | 0 | NA | 0 | -Inf |
| 1214 | Case8 | C081 | INSULA | V1pcLF | 4 | 4 | 1 | 1,51399021583823E-05 | -4,819876931 |
| 1215 | Case8 | C081 | IPa | V1pcLF | 18 | 18 | 1 | 6,81295597127204E-05 | -4,166664418 |
| 1216 | Case8 | C081 | LB | V1pcLF | 8 | 8 | 1 | 3,02798043167646E-05 | -4,518846936 |
| 1217 | Case8 | C081 | LIP | V1pcLF | 28 | 28 | 1 | 0,000105979 | -3,974778891 |
| 1218 | Case8 | C081 | MB | V1pcLF | 1 | 1 | 1 | 3,78497553959558E-06 | -5,421936923 |
| 1219 | Case8 | C081 | MIP | V1pcLF | 0 | 0 | NA | 0 | -Inf |
| 1220 | Case8 | C081 | MST | V1pcLF | 196,5 | 196,5 | 1 | 0,000743748 | -3,128574368 |
| 1221 | Case8 | C081 | MTc | V1pcLF | 211 | 211 | 1 | 0,00079863 | -3,097654467 |
| 1222 | Case8 | C081 | MTp | V1pcLF | 3111 | 3111 | 1 | 0,011775059 | -1,929036912 |
| 1223 | Case8 | C081 | OPAl | V1pcLF | 0 | 0 | NA | 0 | -Inf |
| 1224 | Case8 | C081 | OPro | V1pcLF | 0 | 0 | NA | 0 | -Inf |
| 1225 | Case8 | C081 | PBc | V1pcLF | 14 | 14 | 1 | 5,29896575543381E-05 | -4,275808887 |
| 1226 | Case8 | C081 | PBr | V1pcLF | 1 | 1 | 1 | 3,78497553959558E-06 | -5,421936923 |
| 1227 | Case8 | C081 | PERI | V1pcLF | 90 | 90 | 1 | 0,000340648 | -3,467694413 |
| 1228 | Case8 | C081 | PGa | V1pcLF | 41 | 41 | 1 | 0,000155184 | -3,809153066 |
| 1229 | Case8 | C081 | Pi | V1pcLF | 0 | 0 | NA | 0 | -Inf |
| 1230 | Case8 | C081 | PIP | V1pcLF | 104 | 104 | 1 | 0,000393637 | -3,404903583 |
| 1231 | Case8 | C081 | Pir | V1pcLF | 0 | 0 | NA | 0 | -Inf |
| 1232 | Case8 | C081 | POLE | V1pcLF | 0 | 0 | NA | 0 | -Inf |
| 1233 | Case8 | C081 | Pro.St | V1pcLF | 0 | 0 | NA | 0 | -Inf |
| 1234 | Case8 | C081 | ProM | V1pcLF | 0 | 0 | NA | 0 | -Inf |
| 1235 | Case8 | C081 | SII | V1pcLF | 0 | 0 | NA | 0 | -Inf |
| 1236 | Case8 | C081 | STPc | V1pcLF | 36 | 36 | 1 | 0,000136259 | -3,865634422 |
| 1237 | Case8 | C081 | STPi | V1pcLF | 9 | 9 | 1 | 3,40647798563602E-05 | -4,467694413 |
| 1238 | Case8 | C081 | STPr | V1pcLF | 4 | 4 | 1 | 1,51399021583823E-05 | -4,819876931 |
| 1239 | Case8 | C081 | SUBI | V1pcLF | 0 | 0 | NA | 0 | -Inf |
| 1240 | Case8 | C081 | TEa/ma | V1pcLF | 1 | 1 | 1 | 3,78497553959558E-06 | -5,421936923 |
| 1241 | Case8 | C081 | TEa/mp | V1pcLF | 48 | 48 | 1 | 0,000181679 | -3,740695685 |
| 1242 | Case8 | C081 | TEad | V1pcLF | 29 | 29 | 1 | 0,000109764 | -3,959538925 |
| 1243 | Case8 | C081 | TEav | V1pcLF | 47 | 47 | 1 | 0,000177894 | -3,749839065 |
| 1244 | Case8 | C081 | TEO | V1pcLF | 327 | 327 | 1 | 0,001237687 | -2,90738917 |
| 1245 | Case8 | C081 | TEOm | V1pcLF | 70 | 70 | 1 | 0,000264948 | -3,576838883 |
| 1246 | Case8 | C081 | TEpd | V1pcLF | 325 | 325 | 1 | 0,001230117 | -2,910053562 |
| 1247 | Case8 | C081 | TEpv | V1pcLF | 1221 | 1221 | 1 | 0,004621455 | -2,335221259 |

| | Case | MONKEY | SOURCE | TARGET | TOT | SUP | SLN | FLN | lgFLN |
|---|---|---|---|---|---|---|---|---|---|
| 1248 | Case8 | C081 | TH_TF | V1pcLF | 465 | 465 | 1 | 0,001760014 | -2,75448397 |
| 1249 | Case8 | C081 | TPt | V1pcLF | 3 | 3 | 1 | 1,13549266187867E-05 | -4,944815668 |
| 1250 | Case8 | C081 | V1c | V1pcLF | 2318 | 2318 | 1 | 0,008773573 | -2,056823491 |
| 1251 | Case8 | C081 | V1fpLF | V1pcLF | 112 | 112 | 1 | 0,000423917 | -3,3727189 |
| 1252 | Case8 | C081 | V1fpUF | V1pcLF | 28 | 28 | 1 | 0,000105979 | -3,974778891 |
| 1253 | Case8 | C081 | V1pcLF | V1pcLF | 0 | 0 | NA | 0 | -Inf |
| 1254 | Case8 | C081 | V1pcUF | V1pcLF | 182 | 182 | 1 | 0,000688866 | -3,161865535 |
| 1255 | Case8 | C081 | V2c | V1pcLF | 5800 | 5800 | 1 | 0,021952858 | -1,658508929 |
| 1256 | Case8 | C081 | V2fpLF | V1pcLF | 16 | 16 | 1 | 6,05596086335292E-05 | -4,21781694 |
| 1257 | Case8 | C081 | V2fpUF | V1pcLF | 0 | 0 | NA | 0 | -Inf |
| 1258 | Case8 | C081 | V2pcLF | V1pcLF | 30468 | 30468 | 1 | 0,115320635 | -0,938092976 |
| 1259 | Case8 | C081 | V2pcUF | V1pcLF | 20 | 20 | 1 | 7,56995107919115E-05 | -4,120906927 |
| 1260 | Case8 | C081 | V3A | V1pcLF | 34 | 34 | 1 | 0,000128689 | -3,890458006 |
| 1261 | Case8 | C081 | V3c | V1pcLF | 284 | 284 | 1 | 0,001074933 | -2,968618583 |
| 1262 | Case8 | C081 | V3fpLF | V1pcLF | 88 | 88 | 1 | 0,000333078 | -3,477454251 |
| 1263 | Case8 | C081 | V3fpUF | V1pcLF | 11 | 11 | 1 | 4,16347309355513E-05 | -4,380544238 |
| 1264 | Case8 | C081 | V3pcLF | V1pcLF | 1770 | 1770 | 1 | 0,006699407 | -2,173963656 |
| 1265 | Case8 | C081 | V3pcUF | V1pcLF | 20 | 20 | 1 | 7,56995107919115E-05 | -4,120906927 |
| 1266 | Case8 | C081 | V4c | V1pcLF | 114,5 | 114,5 | 1 | 0,00043338 | -3,363131436 |
| 1267 | Case8 | C081 | V4LF | V1pcLF | 1973 | 1973 | 1 | 0,007467757 | -2,126809838 |
| 1268 | Case8 | C081 | V4t | V1pcLF | 267,5 | 267,5 | 1 | 0,001012481 | -2,994613136 |
| 1269 | Case8 | C081 | V4UF | V1pcLF | 789 | 789 | 1 | 0,002986346 | -2,52485992 |
| 1270 | Case8 | C081 | V6 | V1pcLF | 6 | 6 | 1 | 2,27098532375735E-05 | -4,643785672 |
| 1271 | Case8 | C081 | V6A | V1pcLF | 52 | 52 | 1 | 0,000196819 | -3,705933579 |
| 1272 | Case8 | C081 | VIP | V1pcLF | 2 | 2 | 1 | 7,56995107919115E-06 | -5,120906927 |
| 319 | Case9 | C027 | 1 | V1fpLF | 0 | 0 | NA | 0 | -Inf |
| 320 | Case9 | C027 | 10 | V1fpLF | 0 | 0 | NA | 0 | -Inf |
| 321 | Case9 | C027 | 11 | V1fpLF | 0 | 0 | NA | 0 | -Inf |
| 322 | Case9 | C027 | 12 | V1fpLF | 0 | 0 | NA | 0 | -Inf |
| 323 | Case9 | C027 | 13 | V1fpLF | 0 | 0 | NA | 0 | -Inf |
| 324 | Case9 | C027 | 14 | V1fpLF | 0 | 0 | NA | 0 | -Inf |
| 325 | Case9 | C027 | 2 | V1fpLF | 0 | 0 | NA | 0 | -Inf |
| 326 | Case9 | C027 | 23 | V1fpLF | 0 | 0 | NA | 0 | -Inf |
| 327 | Case9 | C027 | 24a | V1fpLF | 0 | 0 | NA | 0 | -Inf |
| 328 | Case9 | C027 | 24b | V1fpLF | 0 | 0 | NA | 0 | -Inf |
| 329 | Case9 | C027 | 24c | V1fpLF | 0 | 0 | NA | 0 | -Inf |
| 330 | Case9 | C027 | 24d | V1fpLF | 0 | 0 | NA | 0 | -Inf |
| 331 | Case9 | C027 | 25 | V1fpLF | 0 | 0 | NA | 0 | -Inf |
| 332 | Case9 | C027 | 29/30 | V1fpLF | 0 | 0 | NA | 0 | -Inf |
| 333 | Case9 | C027 | 3 | V1fpLF | 0 | 0 | NA | 0 | -Inf |
| 334 | Case9 | C027 | 31 | V1fpLF | 0 | 0 | NA | 0 | -Inf |
| 335 | Case9 | C027 | 32 | V1fpLF | 0 | 0 | NA | 0 | -Inf |
| 336 | Case9 | C027 | 44 | V1fpLF | 0 | 0 | NA | 0 | -Inf |
| 337 | Case9 | C027 | 45A | V1fpLF | 0 | 0 | NA | 0 | -Inf |
| 338 | Case9 | C027 | 45B | V1fpLF | 0 | 0 | NA | 0 | -Inf |
| 339 | Case9 | C027 | 46d | V1fpLF | 0 | 0 | NA | 0 | -Inf |
| 340 | Case9 | C027 | 46v | V1fpLF | 0 | 0 | NA | 0 | -Inf |
| 341 | Case9 | C027 | 5 | V1fpLF | 0 | 0 | NA | 0 | -Inf |
| 342 | Case9 | C027 | 7A | V1fpLF | 2 | 2 | 1 | 1,07449471885846E-05 | -4,968795715 |
| 343 | Case9 | C027 | 7B | V1fpLF | 0 | 0 | NA | 0 | -Inf |
| 344 | Case9 | C027 | 7m | V1fpLF | 0 | 0 | NA | 0 | -Inf |
| 345 | Case9 | C027 | 7op | V1fpLF | 0 | 0 | NA | 0 | -Inf |
| 346 | Case9 | C027 | 8B | V1fpLF | 0 | 0 | NA | 0 | -Inf |
| 347 | Case9 | C027 | 8l | V1fpLF | 1 | 0 | 0 | 5,37247359429228E-06 | -5,26982571 |
| 348 | Case9 | C027 | 8m | V1fpLF | 0 | 0 | NA | 0 | -Inf |
| 349 | Case9 | C027 | 8r | V1fpLF | 0 | 0 | NA | 0 | -Inf |
| 350 | Case9 | C027 | 9 | V1fpLF | 0 | 0 | NA | 0 | -Inf |
| 351 | Case9 | C027 | 9/46d | V1fpLF | 0 | 0 | NA | 0 | -Inf |
| 352 | Case9 | C027 | 9/46v | V1fpLF | 0 | 0 | NA | 0 | -Inf |
| 353 | Case9 | C027 | AIP | V1fpLF | 0 | 0 | NA | 0 | -Inf |
| 354 | Case9 | C027 | CORE | V1fpLF | 1 | 0 | 0 | 5,37247359429228E-06 | -5,26982571 |
| 355 | Case9 | C027 | DP | V1fpLF | 384 | 24 | 0,0625 | 0,00206303 | -2,685494486 |
| 356 | Case9 | C027 | ENTO | V1fpLF | 0 | 0 | NA | 0 | -Inf |
| 357 | Case9 | C027 | F1 | V1fpLF | 0 | 0 | NA | 0 | -Inf |
| 358 | Case9 | C027 | F2 | V1fpLF | 0 | 0 | NA | 0 | -Inf |
| 359 | Case9 | C027 | F3 | V1fpLF | 0 | 0 | NA | 0 | -Inf |
| 360 | Case9 | C027 | F4 | V1fpLF | 0 | 0 | NA | 0 | -Inf |
| 361 | Case9 | C027 | F5 | V1fpLF | 0 | 0 | NA | 0 | -Inf |
| 362 | Case9 | C027 | F6 | V1fpLF | 0 | 0 | NA | 0 | -Inf |
| 363 | Case9 | C027 | F7 | V1fpLF | 0 | 0 | NA | 0 | -Inf |
| 364 | Case9 | C027 | FST | V1fpLF | 29 | 2 | 0,068965517 | 0,000155802 | -3,807427712 |
| 365 | Case9 | C027 | Gu | V1fpLF | 0 | 0 | NA | 0 | -Inf |
| 366 | Case9 | C027 | INSULA | V1fpLF | 6 | 2 | 0,333333333 | 3,22348415657537E-05 | -4,49167446 |
| 367 | Case9 | C027 | IPa | V1fpLF | 1 | 0 | 0 | 5,37247359429228E-06 | -5,26982571 |
| 368 | Case9 | C027 | LB | V1fpLF | 1 | 0 | 0 | 5,37247359429228E-06 | -5,26982571 |

| | Case | MONKEY | SOURCE | TARGET | TOT | SUP | SLN | FLN | lgFLN |
|---|---|---|---|---|---|---|---|---|---|
| 369 | Case9 | C027 | LIP | V1fpLF | 48 | 2 | 0,041666667 | 0,000257879 | -3,588584473 |
| 370 | Case9 | C027 | MB | V1fpLF | 4 | 2 | 0,5 | 2,14898943771691E-05 | -4,667765719 |
| 371 | Case9 | C027 | MIP | V1fpLF | 4 | 0 | 0 | 2,14898943771691E-05 | -4,667765719 |
| 372 | Case9 | C027 | MST | V1fpLF | 106 | 0 | 0 | 0,000569482 | -3,244519845 |
| 373 | Case9 | C027 | MTc | V1fpLF | 44 | 0 | 0 | 0,000236389 | -3,626373034 |
| 374 | Case9 | C027 | MTp | V1fpLF | 1132 | 42 | 0,037102473 | 0,00608164 | -2,215979284 |
| 375 | Case9 | C027 | OPAl | V1fpLF | 0 | 0 | NA | 0 | -Inf |
| 376 | Case9 | C027 | OPro | V1fpLF | 0 | 0 | NA | 0 | -Inf |
| 377 | Case9 | C027 | PBc | V1fpLF | 6 | 0 | 0 | 3,22348415657537E-05 | -4,49167446 |
| 378 | Case9 | C027 | PBr | V1fpLF | 0 | 0 | NA | 0 | -Inf |
| 379 | Case9 | C027 | PERI | V1fpLF | 2 | 0 | 0 | 1,07449471885846E-05 | -4,968795715 |
| 380 | Case9 | C027 | PGa | V1fpLF | 4,5 | 0 | 0 | 2,41761311743153E-05 | -4,616613197 |
| 381 | Case9 | C027 | Pi | V1fpLF | 0 | 0 | NA | 0 | -Inf |
| 382 | Case9 | C027 | PIP | V1fpLF | 82 | 2 | 0,024390244 | 0,000440543 | -3,356011858 |
| 383 | Case9 | C027 | Pir | V1fpLF | 0 | 0 | NA | 0 | -Inf |
| 384 | Case9 | C027 | POLE | V1fpLF | 0 | 0 | NA | 0 | -Inf |
| 385 | Case9 | C027 | Pro.St | V1fpLF | 2 | 0 | 0 | 1,07449471885846E-05 | -4,968795715 |
| 386 | Case9 | C027 | ProM | V1fpLF | 0 | 0 | NA | 0 | -Inf |
| 387 | Case9 | C027 | SII | V1fpLF | 2 | 0 | 0 | 1,07449471885846E-05 | -4,968795715 |
| 388 | Case9 | C027 | STPc | V1fpLF | 8 | 0 | 0 | 4,29797887543383E-05 | -4,366735723 |
| 389 | Case9 | C027 | STPi | V1fpLF | 3 | 0 | 0 | 1,61174207828769E-05 | -4,792704456 |
| 390 | Case9 | C027 | STPr | V1fpLF | 1 | 0 | 0 | 5,37247359429228E-06 | -5,26982571 |
| 391 | Case9 | C027 | SUBI | V1fpLF | 0 | 0 | NA | 0 | -Inf |
| 392 | Case9 | C027 | TEa/ma | V1fpLF | 0 | 0 | NA | 0 | -Inf |
| 393 | Case9 | C027 | TEa/mp | V1fpLF | 1,5 | 0 | 0 | 8,05871039143843E-06 | -5,093734451 |
| 394 | Case9 | C027 | TEad | V1fpLF | 2 | 0 | 0 | 1,07449471885846E-05 | -4,968795715 |
| 395 | Case9 | C027 | TEav | V1fpLF | 2 | 0 | 0 | 1,07449471885846E-05 | -4,968795715 |
| 396 | Case9 | C027 | TEO | V1fpLF | 23,5 | 0 | 0 | 0,000126253 | -3,898757848 |
| 397 | Case9 | C027 | TEOm | V1fpLF | 0 | 0 | NA | 0 | -Inf |
| 398 | Case9 | C027 | TEpd | V1fpLF | 10 | 0 | 0 | 5,37247359429228E-05 | -4,26982571 |
| 399 | Case9 | C027 | TEpv | V1fpLF | 201,5 | 7 | 0,034739454 | 0,001082553 | -2,96555066 |
| 400 | Case9 | C027 | TH_TF | V1fpLF | 126 | 0 | 0 | 0,000676932 | -3,169455165 |
| 401 | Case9 | C027 | TPt | V1fpLF | 10 | 0 | 0 | 5,37247359429228E-05 | -4,26982571 |
| 402 | Case9 | C027 | V1c | V1fpLF | 0 | 0 | NA | 0 | -Inf |
| 403 | Case9 | C027 | V1fpLF | V1fpLF | 0 | 0 | NA | 0 | -Inf |
| 404 | Case9 | C027 | V1fpUF | V1fpLF | 2 | 0 | 0 | 1,07449471885846E-05 | -4,968795715 |
| 405 | Case9 | C027 | V1pcLF | V1fpLF | 532 | 18 | 0,033834586 | 0,002858156 | -2,543914078 |
| 406 | Case9 | C027 | V1pcUF | V1fpLF | 0 | 0 | NA | 0 | -Inf |
| 407 | Case9 | C027 | V2c | V1fpLF | 0 | 0 | NA | 0 | -Inf |
| 408 | Case9 | C027 | V2fpLF | V1fpLF | 26860,5 | 12012,5 | 0,447218034 | 0,144307327 | -0,840711618 |
| 409 | Case9 | C027 | V2fpUF | V1fpLF | 2 | 0 | 0 | 1,07449471885846E-05 | -4,968795715 |
| 410 | Case9 | C027 | V2pcLF | V1fpLF | 274 | 24 | 0,087591241 | 0,001472058 | -2,832075148 |
| 411 | Case9 | C027 | V2pcUF | V1fpLF | 0 | 0 | NA | 0 | -Inf |
| 412 | Case9 | C027 | V3A | V1fpLF | 8644 | 3156 | 0,365108746 | 0,046439662 | -1,333110952 |
| 413 | Case9 | C027 | V3c | V1fpLF | 2 | 0 | 0 | 1,07449471885846E-05 | -4,968795715 |
| 414 | Case9 | C027 | V3fpLF | V1fpLF | 6828 | 496 | 0,072642062 | 0,03668325 | -1,435532198 |
| 415 | Case9 | C027 | V3fpUF | V1fpLF | 2 | 0 | 0 | 1,07449471885846E-05 | -4,968795715 |
| 416 | Case9 | C027 | V3pcLF | V1fpLF | 184 | 8 | 0,043478261 | 0,000988535 | -3,005007887 |
| 417 | Case9 | C027 | V3pcUF | V1fpLF | 0 | 0 | NA | 0 | -Inf |
| 418 | Case9 | C027 | V4c | V1fpLF | 0 | 0 | NA | 0 | -Inf |
| 419 | Case9 | C027 | V4LF | V1fpLF | 2570 | 486 | 0,189105058 | 0,013807257 | -1,859892587 |
| 420 | Case9 | C027 | V4t | V1fpLF | 18 | 0 | 0 | 9,67045246972611E-05 | -4,014553205 |
| 421 | Case9 | C027 | V4UF | V1fpLF | 0 | 0 | NA | 0 | -Inf |
| 422 | Case9 | C027 | V6 | V1fpLF | 8 | 0 | 0 | 4,29797887543383E-05 | -4,366735723 |
| 423 | Case9 | C027 | V6A | V1fpLF | 130 | 0 | 0 | 0,000698422 | -3,155882358 |
| 424 | Case9 | C027 | VIP | V1fpLF | 2 | 0 | 0 | 1,07449471885846E-05 | -4,968795715 |
| 425 | Case10 | C054 | 1 | V1fpLF | 0 | 0 | NA | 0 | -Inf |
| 426 | Case10 | C054 | 10 | V1fpLF | 0 | 0 | NA | 0 | -Inf |
| 427 | Case10 | C054 | 11 | V1fpLF | 0 | 0 | NA | 0 | -Inf |
| 428 | Case10 | C054 | 12 | V1fpLF | 0 | 0 | NA | 0 | -Inf |
| 429 | Case10 | C054 | 13 | V1fpLF | 0 | 0 | NA | 0 | -Inf |
| 430 | Case10 | C054 | 14 | V1fpLF | 0 | 0 | NA | 0 | -Inf |
| 431 | Case10 | C054 | 2 | V1fpLF | 0 | 0 | NA | 0 | -Inf |
| 432 | Case10 | C054 | 23 | V1fpLF | 0 | 0 | NA | 0 | -Inf |
| 433 | Case10 | C054 | 24a | V1fpLF | 0 | 0 | NA | 0 | -Inf |
| 434 | Case10 | C054 | 24b | V1fpLF | 0 | 0 | NA | 0 | -Inf |
| 435 | Case10 | C054 | 24c | V1fpLF | 0 | 0 | NA | 0 | -Inf |
| 436 | Case10 | C054 | 24d | V1fpLF | 2 | 0 | 0 | 1,07449471885846E-05 | -4,968795715 |
| 437 | Case10 | C054 | 25 | V1fpLF | 0 | 0 | NA | 0 | -Inf |
| 438 | Case10 | C054 | 29/30 | V1fpLF | 0 | 0 | NA | 0 | -Inf |
| 439 | Case10 | C054 | 3 | V1fpLF | 0 | 0 | NA | 0 | -Inf |
| 440 | Case10 | C054 | 31 | V1fpLF | 0 | 0 | NA | 0 | -Inf |
| 441 | Case10 | C054 | 32 | V1fpLF | 0 | 0 | NA | 0 | -Inf |
| 442 | Case10 | C054 | 44 | V1fpLF | 0 | 0 | NA | 0 | -Inf |
| 443 | Case10 | C054 | 45A | V1fpLF | 0 | 0 | NA | 0 | -Inf |

| | Case | MONKEY | SOURCE | TARGET | TOT | SUP | SLN | FLN | lgFLN |
|---|---|---|---|---|---|---|---|---|---|
| 444 | Case10 | C054 | 45B | V1fpLF | 0 | 0 | NA | 0 | -Inf |
| 445 | Case10 | C054 | 46d | V1fpLF | 0 | 0 | NA | 0 | -Inf |
| 446 | Case10 | C054 | 46v | V1fpLF | 0 | 0 | NA | 0 | -Inf |
| 447 | Case10 | C054 | 5 | V1fpLF | 0 | 0 | NA | 0 | -Inf |
| 448 | Case10 | C054 | 7A | V1fpLF | 0 | 0 | NA | 0 | -Inf |
| 449 | Case10 | C054 | 7B | V1fpLF | 0 | 0 | NA | 0 | -Inf |
| 450 | Case10 | C054 | 7m | V1fpLF | 0 | 0 | NA | 0 | -Inf |
| 451 | Case10 | C054 | 7op | V1fpLF | 0 | 0 | NA | 0 | -Inf |
| 452 | Case10 | C054 | 8B | V1fpLF | 0 | 0 | NA | 0 | -Inf |
| 453 | Case10 | C054 | 8l | V1fpLF | 10 | 6 | 0,6 | 5,37247359429228E-05 | -4,26982571 |
| 454 | Case10 | C054 | 8m | V1fpLF | 0 | 0 | NA | 0 | -Inf |
| 455 | Case10 | C054 | 8r | V1fpLF | 0 | 0 | NA | 0 | -Inf |
| 456 | Case10 | C054 | 9 | V1fpLF | 0 | 0 | NA | 0 | -Inf |
| 457 | Case10 | C054 | 9/46d | V1fpLF | 0 | 0 | NA | 0 | -Inf |
| 458 | Case10 | C054 | 9/46v | V1fpLF | 0 | 0 | NA | 0 | -Inf |
| 459 | Case10 | C054 | AIP | V1fpLF | 0 | 0 | NA | 0 | -Inf |
| 460 | Case10 | C054 | CORE | V1fpLF | 37 | 0 | 0 | 0,000198782 | -3,701623986 |
| 461 | Case10 | C054 | DP | V1fpLF | 944 | 98 | 0,103813559 | 0,005071615 | -2,294853716 |
| 462 | Case10 | C054 | ENTO | V1fpLF | 0 | 0 | NA | 0 | -Inf |
| 463 | Case10 | C054 | F1 | V1fpLF | 0 | 0 | NA | 0 | -Inf |
| 464 | Case10 | C054 | F2 | V1fpLF | 0 | 0 | NA | 0 | -Inf |
| 465 | Case10 | C054 | F3 | V1fpLF | 0 | 0 | NA | 0 | -Inf |
| 466 | Case10 | C054 | F4 | V1fpLF | 0 | 0 | NA | 0 | -Inf |
| 467 | Case10 | C054 | F5 | V1fpLF | 0 | 0 | NA | 0 | -Inf |
| 468 | Case10 | C054 | F6 | V1fpLF | 0 | 0 | NA | 0 | -Inf |
| 469 | Case10 | C054 | F7 | V1fpLF | 0 | 0 | NA | 0 | -Inf |
| 470 | Case10 | C054 | FST | V1fpLF | 112 | 0 | 0 | 0,000601717 | -3,220607688 |
| 471 | Case10 | C054 | Gu | V1fpLF | 0 | 0 | NA | 0 | -Inf |
| 472 | Case10 | C054 | INSULA | V1fpLF | 41 | 0 | 0 | 0,000220271 | -3,657041854 |
| 473 | Case10 | C054 | IPa | V1fpLF | 10 | 0 | 0 | 5,37247359429228E-05 | -4,26982571 |
| 474 | Case10 | C054 | LB | V1fpLF | 64 | 0 | 0 | 0,000343838 | -3,463645736 |
| 475 | Case10 | C054 | LIP | V1fpLF | 258 | 0 | 0 | 0,001386098 | -2,858206004 |
| 476 | Case10 | C054 | MB | V1fpLF | 48 | 2 | 0,041666667 | 0,000257879 | -3,588584473 |
| 477 | Case10 | C054 | MIP | V1fpLF | 6 | 0 | 0 | 3,22348415657537E-05 | -4,49167446 |
| 478 | Case10 | C054 | MST | V1fpLF | 884 | 2 | 0,002262443 | 0,004749267 | -2,323373445 |
| 479 | Case10 | C054 | MTc | V1fpLF | 270 | 2 | 0,007407407 | 0,001450568 | -2,838461946 |
| 480 | Case10 | C054 | MTp | V1fpLF | 2224 | 106 | 0,047661871 | 0,011948381 | -1,922690927 |
| 481 | Case10 | C054 | OPAl | V1fpLF | 0 | 0 | NA | 0 | -Inf |
| 482 | Case10 | C054 | OPro | V1fpLF | 0 | 0 | NA | 0 | -Inf |
| 483 | Case10 | C054 | PBc | V1fpLF | 103 | 0 | 0 | 0,000553365 | -3,256988486 |
| 484 | Case10 | C054 | PBr | V1fpLF | 2 | 0 | 0 | 1,07449471885846E-05 | -4,968795715 |
| 485 | Case10 | C054 | PERI | V1fpLF | 36 | 0 | 0 | 0,000193409 | -3,71352321 |
| 486 | Case10 | C054 | PGa | V1fpLF | 46 | 2 | 0,043478261 | 0,000247134 | -3,607067879 |
| 487 | Case10 | C054 | Pi | V1fpLF | 0 | 0 | NA | 0 | -Inf |
| 488 | Case10 | C054 | PIP | V1fpLF | 596 | 8 | 0,013422819 | 0,003201994 | -2,494579451 |
| 489 | Case10 | C054 | Pir | V1fpLF | 0 | 0 | NA | 0 | -Inf |
| 490 | Case10 | C054 | POLE | V1fpLF | 0 | 0 | NA | 0 | -Inf |
| 491 | Case10 | C054 | Pro.St | V1fpLF | 48 | 2 | 0,041666667 | 0,000257879 | -3,588584473 |
| 492 | Case10 | C054 | ProM | V1fpLF | 0 | 0 | NA | 0 | -Inf |
| 493 | Case10 | C054 | SII | V1fpLF | 2 | 0 | 0 | 1,07449471885846E-05 | -4,968795715 |
| 494 | Case10 | C054 | STPc | V1fpLF | 142 | 0 | 0 | 0,000762891 | -3,117537366 |
| 495 | Case10 | C054 | STPi | V1fpLF | 13 | 0 | 0 | 6,98421567257997E-05 | -4,155882358 |
| 496 | Case10 | C054 | STPr | V1fpLF | 6 | 0 | 0 | 3,22348415657537E-05 | -4,49167446 |
| 497 | Case10 | C054 | SUBI | V1fpLF | 0 | 0 | NA | 0 | -Inf |
| 498 | Case10 | C054 | TEa/ma | V1fpLF | 0 | 0 | NA | 0 | -Inf |
| 499 | Case10 | C054 | TEa/mp | V1fpLF | 16 | 0 | 0 | 8,59595775086765E-05 | -4,065705728 |
| 500 | Case10 | C054 | TEad | V1fpLF | 4 | 0 | 0 | 2,14898943771691E-05 | -4,667765719 |
| 501 | Case10 | C054 | TEav | V1fpLF | 6 | 0 | 0 | 3,22348415657537E-05 | -4,49167446 |
| 502 | Case10 | C054 | TEO | V1fpLF | 106 | 0 | 0 | 0,000569482 | -3,244519845 |
| 503 | Case10 | C054 | TEOm | V1fpLF | 6 | 0 | 0 | 3,22348415657537E-05 | -4,49167446 |
| 504 | Case10 | C054 | TEpd | V1fpLF | 32 | 8 | 0,25 | 0,000171919 | -3,764675732 |
| 505 | Case10 | C054 | TEpv | V1fpLF | 397 | 4 | 0,010075567 | 0,002132872 | -2,671035204 |
| 506 | Case10 | C054 | TH_TF | V1fpLF | 335 | 4 | 0,011940299 | 0,001799779 | -2,744780903 |
| 507 | Case10 | C054 | TPt | V1fpLF | 2 | 0 | 0 | 1,07449471885846E-05 | -4,968795715 |
| 508 | Case10 | C054 | V1c | V1fpLF | 60 | 22 | 0,366666667 | 0,000322348 | -3,49167446 |
| 509 | Case10 | C054 | V1fpLF | V1fpLF | 0 | 0 | NA | 0 | -Inf |
| 510 | Case10 | C054 | V1fpUF | V1fpLF | 2 | 0 | 0 | 1,07449471885846E-05 | -4,968795715 |
| 511 | Case10 | C054 | V1pcLF | V1fpLF | 656 | 92 | 0,140243902 | 0,003524343 | -2,452921871 |
| 512 | Case10 | C054 | V1pcUF | V1fpLF | 8 | 0 | 0 | 4,29797887543383E-05 | -4,366735723 |
| 513 | Case10 | C054 | V2c | V1fpLF | 12 | 0 | 0 | 6,44696831315074E-05 | -4,190644464 |
| 514 | Case10 | C054 | V2fpLF | V1fpLF | 26568 | 7092 | 0,266937669 | 0,142735878 | -0,845466848 |
| 515 | Case10 | C054 | V2fpUF | V1fpLF | 8 | 0 | 0 | 4,29797887543383E-05 | -4,366735723 |
| 516 | Case10 | C054 | V2pcLF | V1fpLF | 790 | 132 | 0,167088608 | 0,004244254 | -2,372198619 |
| 517 | Case10 | C054 | V2pcUF | V1fpLF | 6 | 0 | 0 | 3,22348415657537E-05 | -4,49167446 |
| 518 | Case10 | C054 | V3A | V1fpLF | 6004 | 1518 | 0,252831446 | 0,032256331 | -1,491385027 |

| | Case | MONKEY | SOURCE | TARGET | TOT | SUP | SLN | FLN | lgFLN |
|---|---|---|---|---|---|---|---|---|---|
| 519 | Case10 | C054 | V3c | V1fpLF | 34 | 0 | 0 | 0,000182664 | -3,738346793 |
| 520 | Case10 | C054 | V3fpLF | V1fpLF | 6084 | 882 | 0,144970414 | 0,032686129 | -1,485636505 |
| 521 | Case10 | C054 | V3fpUF | V1fpLF | 22 | 2 | 0,090909091 | 0,000118194 | -3,92740303 |
| 522 | Case10 | C054 | V3pcLF | V1fpLF | 70 | 0 | 0 | 0,000376073 | -3,42472767 |
| 523 | Case10 | C054 | V3pcUF | V1fpLF | 2 | 0 | 0 | 1,07449471885846E-05 | -4,968795715 |
| 524 | Case10 | C054 | V4c | V1fpLF | 4 | 0 | 0 | 2,14898943771691E-05 | -4,667765719 |
| 525 | Case10 | C054 | V4LF | V1fpLF | 2254 | 144 | 0,063886424 | 0,012109555 | -1,916871799 |
| 526 | Case10 | C054 | V4t | V1fpLF | 24 | 2 | 0,083333333 | 0,000128939 | -3,889614469 |
| 527 | Case10 | C054 | V4UF | V1fpLF | 34 | 0 | 0 | 0,000182664 | -3,738346793 |
| 528 | Case10 | C054 | V6 | V1fpLF | 80 | 2 | 0,025 | 0,000429798 | -3,366735723 |
| 529 | Case10 | C054 | V6A | V1fpLF | 324 | 12 | 0,037037037 | 0,001740681 | -2,7592807 |
| 530 | Case10 | C054 | VIP | V1fpLF | 140 | 0 | 0 | 0,000752146 | -3,123697675 |
| 531 | Case11 | C081 | 1 | V1fpLF | 0 | 0 | NA | 0 | -Inf |
| 532 | Case11 | C081 | 10 | V1fpLF | 0 | 0 | NA | 0 | -Inf |
| 533 | Case11 | C081 | 11 | V1fpLF | 0 | 0 | NA | 0 | -Inf |
| 534 | Case11 | C081 | 12 | V1fpLF | 0 | 0 | NA | 0 | -Inf |
| 535 | Case11 | C081 | 13 | V1fpLF | 0 | 0 | NA | 0 | -Inf |
| 536 | Case11 | C081 | 14 | V1fpLF | 0 | 0 | NA | 0 | -Inf |
| 537 | Case11 | C081 | 2 | V1fpLF | 0 | 0 | NA | 0 | -Inf |
| 538 | Case11 | C081 | 23 | V1fpLF | 4 | 4 | 1 | 2,14898943771691E-05 | -4,667765719 |
| 539 | Case11 | C081 | 24a | V1fpLF | 0 | 0 | NA | 0 | -Inf |
| 540 | Case11 | C081 | 24b | V1fpLF | 0 | 0 | NA | 0 | -Inf |
| 541 | Case11 | C081 | 24c | V1fpLF | 0 | 0 | NA | 0 | -Inf |
| 542 | Case11 | C081 | 24d | V1fpLF | 0 | 0 | NA | 0 | -Inf |
| 543 | Case11 | C081 | 25 | V1fpLF | 0 | 0 | NA | 0 | -Inf |
| 544 | Case11 | C081 | 29/30 | V1fpLF | 0 | 0 | NA | 0 | -Inf |
| 545 | Case11 | C081 | 3 | V1fpLF | 0 | 0 | NA | 0 | -Inf |
| 546 | Case11 | C081 | 31 | V1fpLF | 0 | 0 | NA | 0 | -Inf |
| 547 | Case11 | C081 | 32 | V1fpLF | 0 | 0 | NA | 0 | -Inf |
| 548 | Case11 | C081 | 44 | V1fpLF | 0 | 0 | NA | 0 | -Inf |
| 549 | Case11 | C081 | 45A | V1fpLF | 0 | 0 | NA | 0 | -Inf |
| 550 | Case11 | C081 | 45B | V1fpLF | 0 | 0 | NA | 0 | -Inf |
| 551 | Case11 | C081 | 46d | V1fpLF | 0 | 0 | NA | 0 | -Inf |
| 552 | Case11 | C081 | 46v | V1fpLF | 0 | 0 | NA | 0 | -Inf |
| 553 | Case11 | C081 | 5 | V1fpLF | 0 | 0 | NA | 0 | -Inf |
| 554 | Case11 | C081 | 7A | V1fpLF | 9 | 9 | 1 | 4,83522623486306E-05 | -4,315583201 |
| 555 | Case11 | C081 | 7B | V1fpLF | 0 | 0 | NA | 0 | -Inf |
| 556 | Case11 | C081 | 7m | V1fpLF | 0 | 0 | NA | 0 | -Inf |
| 557 | Case11 | C081 | 7op | V1fpLF | 1 | 1 | 1 | 5,37247359429228E-06 | -5,26982571 |
| 558 | Case11 | C081 | 8B | V1fpLF | 0 | 0 | NA | 0 | -Inf |
| 559 | Case11 | C081 | 8l | V1fpLF | 0 | 0 | NA | 0 | -Inf |
| 560 | Case11 | C081 | 8m | V1fpLF | 0 | 0 | NA | 0 | -Inf |
| 561 | Case11 | C081 | 8r | V1fpLF | 0 | 0 | NA | 0 | -Inf |
| 562 | Case11 | C081 | 9 | V1fpLF | 0 | 0 | NA | 0 | -Inf |
| 563 | Case11 | C081 | 9/46d | V1fpLF | 0 | 0 | NA | 0 | -Inf |
| 564 | Case11 | C081 | 9/46v | V1fpLF | 0 | 0 | NA | 0 | -Inf |
| 565 | Case11 | C081 | AIP | V1fpLF | 1 | 0 | 0 | 5,37247359429228E-06 | -5,26982571 |
| 566 | Case11 | C081 | CORE | V1fpLF | 4 | 4 | 1 | 2,14898943771691E-05 | -4,667765719 |
| 567 | Case11 | C081 | DP | V1fpLF | 820 | 820 | 1 | 0,004405428 | -2,356011858 |
| 568 | Case11 | C081 | ENTO | V1fpLF | 0 | 0 | NA | 0 | -Inf |
| 569 | Case11 | C081 | F1 | V1fpLF | 0 | 0 | NA | 0 | -Inf |
| 570 | Case11 | C081 | F2 | V1fpLF | 0 | 0 | NA | 0 | -Inf |
| 571 | Case11 | C081 | F3 | V1fpLF | 0 | 0 | NA | 0 | -Inf |
| 572 | Case11 | C081 | F4 | V1fpLF | 0 | 0 | NA | 0 | -Inf |
| 573 | Case11 | C081 | F5 | V1fpLF | 0 | 0 | NA | 0 | -Inf |
| 574 | Case11 | C081 | F6 | V1fpLF | 0 | 0 | NA | 0 | -Inf |
| 575 | Case11 | C081 | F7 | V1fpLF | 0 | 0 | NA | 0 | -Inf |
| 576 | Case11 | C081 | FST | V1fpLF | 92 | 92 | 1 | 0,000494268 | -3,306037883 |
| 577 | Case11 | C081 | Gu | V1fpLF | 0 | 0 | NA | 0 | -Inf |
| 578 | Case11 | C081 | INSULA | V1fpLF | 18 | 18 | 1 | 9,67045246972611E-05 | -4,014553205 |
| 579 | Case11 | C081 | IPa | V1fpLF | 11 | 11 | 1 | 5,90972095372151E-05 | -4,228433025 |
| 580 | Case11 | C081 | LB | V1fpLF | 28 | 28 | 1 | 0,000150429 | -3,822667679 |
| 581 | Case11 | C081 | LIP | V1fpLF | 246 | 246 | 1 | 0,001321629 | -2,878890603 |
| 582 | Case11 | C081 | MB | V1fpLF | 15 | 15 | 1 | 8,05871039143843E-05 | -4,093734451 |
| 583 | Case11 | C081 | MIP | V1fpLF | 7 | 7 | 1 | 3,7607315160046E-05 | -4,42472767 |
| 584 | Case11 | C081 | MST | V1fpLF | 925 | 925 | 1 | 0,004969538 | -2,303683978 |
| 585 | Case11 | C081 | MTc | V1fpLF | 107 | 107 | 1 | 0,000574855 | -3,240441933 |
| 586 | Case11 | C081 | MTp | V1fpLF | 3953,5 | 3953,5 | 1 | 0,021240074 | -1,672843967 |
| 587 | Case11 | C081 | OPAl | V1fpLF | 0 | 0 | NA | 0 | -Inf |
| 588 | Case11 | C081 | OPro | V1fpLF | 0 | 0 | NA | 0 | -Inf |
| 589 | Case11 | C081 | PBc | V1fpLF | 49 | 49 | 1 | 0,000263251 | -3,57962963 |
| 590 | Case11 | C081 | PBr | V1fpLF | 1 | 1 | 1 | 5,37247359429228E-06 | -5,26982571 |
| 591 | Case11 | C081 | PERI | V1fpLF | 68 | 68 | 1 | 0,000365328 | -3,437316798 |
| 592 | Case11 | C081 | PGa | V1fpLF | 66 | 66 | 1 | 0,000354583 | -3,450281775 |
| 593 | Case11 | C081 | Pi | V1fpLF | 0 | 0 | NA | 0 | -Inf |

| | Case | MONKEY | SOURCE | TARGET | TOT | SUP | SLN | FLN | lgFLN |
|---|---|---|---|---|---|---|---|---|---|
| 594 | Case11 | C081 | PIP | V1fpLF | 318 | 318 | 1 | 0,001708447 | -2,76739859 |
| 595 | Case11 | C081 | Pir | V1fpLF | 0 | 0 | NA | 0 | -Inf |
| 596 | Case11 | C081 | POLE | V1fpLF | 0 | 0 | NA | 0 | -Inf |
| 597 | Case11 | C081 | Pro.St | V1fpLF | 10 | 10 | 1 | 5,37247359429228E-05 | -4,26982571 |
| 598 | Case11 | C081 | ProM | V1fpLF | 0 | 0 | NA | 0 | -Inf |
| 599 | Case11 | C081 | SII | V1fpLF | 0 | 0 | NA | 0 | -Inf |
| 600 | Case11 | C081 | STPc | V1fpLF | 161,5 | 161,5 | 1 | 0,000867654 | -3,061653184 |
| 601 | Case11 | C081 | STPi | V1fpLF | 23 | 23 | 1 | 0,000123567 | -3,908097874 |
| 602 | Case11 | C081 | STPr | V1fpLF | 1 | 1 | 1 | 5,37247359429228E-06 | -5,26982571 |
| 603 | Case11 | C081 | SUBI | V1fpLF | 0 | 0 | NA | 0 | -Inf |
| 604 | Case11 | C081 | TEa/ma | V1fpLF | 1 | 1 | 1 | 5,37247359429228E-06 | -5,26982571 |
| 605 | Case11 | C081 | TEa/mp | V1fpLF | 9 | 9 | 1 | 4,83522623486306E-05 | -4,315583201 |
| 606 | Case11 | C081 | TEad | V1fpLF | 6 | 6 | 1 | 3,22348415657537E-05 | -4,49167446 |
| 607 | Case11 | C081 | TEav | V1fpLF | 13 | 13 | 1 | 6,98421567257997E-05 | -4,155882358 |
| 608 | Case11 | C081 | TEO | V1fpLF | 127,5 | 127,5 | 1 | 0,00068499 | -3,164315526 |
| 609 | Case11 | C081 | TEOm | V1fpLF | 35 | 35 | 1 | 0,000188037 | -3,725757666 |
| 610 | Case11 | C081 | TEpd | V1fpLF | 54 | 54 | 1 | 0,000290114 | -3,537431951 |
| 611 | Case11 | C081 | TEpv | V1fpLF | 1051 | 1051 | 1 | 0,00564647 | -2,248222994 |
| 612 | Case11 | C081 | TH_TF | V1fpLF | 881 | 881 | 1 | 0,004733149 | -2,324849802 |
| 613 | Case11 | C081 | TPt | V1fpLF | 29 | 29 | 1 | 0,000155802 | -3,807427712 |
| 614 | Case11 | C081 | V1c | V1fpLF | 6 | 6 | 1 | 3,22348415657537E-05 | -4,49167446 |
| 615 | Case11 | C081 | V1fpLF | V1fpLF | 0 | 0 | NA | 0 | -Inf |
| 616 | Case11 | C081 | V1fpUF | V1fpLF | 6252 | 6252 | 1 | 0,033588705 | -1,473806741 |
| 617 | Case11 | C081 | V1pcLF | V1fpLF | 606 | 606 | 1 | 0,003255719 | -2,487353086 |
| 618 | Case11 | C081 | V1pcUF | V1fpLF | 0 | 0 | NA | 0 | -Inf |
| 619 | Case11 | C081 | V2c | V1fpLF | 122 | 122 | 1 | 0,000655442 | -3,18346588 |
| 620 | Case11 | C081 | V2fpLF | V1fpLF | 41186 | 41186 | 1 | 0,221270697 | -0,655076095 |
| 621 | Case11 | C081 | V2fpUF | V1fpLF | 529 | 529 | 1 | 0,002842039 | -2,546370038 |
| 622 | Case11 | C081 | V2pcLF | V1fpLF | 2236 | 2236 | 1 | 0,012012851 | -1,920353911 |
| 623 | Case11 | C081 | V2pcUF | V1fpLF | 698 | 698 | 1 | 0,003749987 | -2,425970288 |
| 624 | Case11 | C081 | V3A | V1fpLF | 578 | 578 | 1 | 0,00310529 | -2,507897872 |
| 625 | Case11 | C081 | V3c | V1fpLF | 4 | 4 | 1 | 2,14898943771691E-05 | -4,667765719 |
| 626 | Case11 | C081 | V3fpLF | V1fpLF | 16488 | 16488 | 1 | 0,088581345 | -1,052657732 |
| 627 | Case11 | C081 | V3fpUF | V1fpLF | 635 | 635 | 1 | 0,003411521 | -2,467051985 |
| 628 | Case11 | C081 | V3pcLF | V1fpLF | 812 | 812 | 1 | 0,004362449 | -2,360269681 |
| 629 | Case11 | C081 | V3pcUF | V1fpLF | 190 | 190 | 1 | 0,00102077 | -2,991072109 |
| 630 | Case11 | C081 | V4c | V1fpLF | 4 | 4 | 1 | 2,14898943771691E-05 | -4,667765719 |
| 631 | Case11 | C081 | V4LF | V1fpLF | 1868 | 1868 | 1 | 0,010035781 | -1,998448838 |
| 632 | Case11 | C081 | V4t | V1fpLF | 539,5 | 539,5 | 1 | 0,00289845 | -2,537834261 |
| 633 | Case11 | C081 | V4UF | V1fpLF | 922 | 922 | 1 | 0,004953421 | -2,305094789 |
| 634 | Case11 | C081 | V6 | V1fpLF | 3610 | 3610 | 1 | 0,01939463 | -1,712318508 |
| 635 | Case11 | C081 | V6A | V1fpLF | 1352 | 1352 | 1 | 0,007263584 | -2,138849019 |
| 636 | Case11 | C081 | VIP | V1fpLF | 48,5 | 48,5 | 1 | 0,000260565 | -3,584083972 |
| 637 | Case12 | C057 | 1 | V1fpUF | 0 | 0 | NA | 0 | -Inf |
| 638 | Case12 | C057 | 10 | V1fpUF | 0 | 0 | NA | 0 | -Inf |
| 639 | Case12 | C057 | 11 | V1fpUF | 0 | 0 | NA | 0 | -Inf |
| 640 | Case12 | C057 | 12 | V1fpUF | 0 | 0 | NA | 0 | -Inf |
| 641 | Case12 | C057 | 13 | V1fpUF | 0 | 0 | NA | 0 | -Inf |
| 642 | Case12 | C057 | 14 | V1fpUF | 0 | 0 | NA | 0 | -Inf |
| 643 | Case12 | C057 | 2 | V1fpUF | 0 | 0 | NA | 0 | -Inf |
| 644 | Case12 | C057 | 23 | V1fpUF | 4 | 0 | 0 | 5,04044960810504E-05 | -4,297530723 |
| 645 | Case12 | C057 | 24a | V1fpUF | 0 | 0 | NA | 0 | -Inf |
| 646 | Case12 | C057 | 24b | V1fpUF | 0 | 0 | NA | 0 | -Inf |
| 647 | Case12 | C057 | 24c | V1fpUF | 0 | 0 | NA | 0 | -Inf |
| 648 | Case12 | C057 | 24d | V1fpUF | 0 | 0 | NA | 0 | -Inf |
| 649 | Case12 | C057 | 25 | V1fpUF | 0 | 0 | NA | 0 | -Inf |
| 650 | Case12 | C057 | 29/30 | V1fpUF | 0 | 0 | NA | 0 | -Inf |
| 651 | Case12 | C057 | 3 | V1fpUF | 0 | 0 | NA | 0 | -Inf |
| 652 | Case12 | C057 | 31 | V1fpUF | 0 | 0 | NA | 0 | -Inf |
| 653 | Case12 | C057 | 32 | V1fpUF | 0 | 0 | NA | 0 | -Inf |
| 654 | Case12 | C057 | 44 | V1fpUF | 0 | 0 | NA | 0 | -Inf |
| 655 | Case12 | C057 | 45A | V1fpUF | 0 | 0 | NA | 0 | -Inf |
| 656 | Case12 | C057 | 45B | V1fpUF | 0 | 0 | NA | 0 | -Inf |
| 657 | Case12 | C057 | 46d | V1fpUF | 0 | 0 | NA | 0 | -Inf |
| 658 | Case12 | C057 | 46v | V1fpUF | 0 | 0 | NA | 0 | -Inf |
| 659 | Case12 | C057 | 5 | V1fpUF | 0 | 0 | NA | 0 | -Inf |
| 660 | Case12 | C057 | 7A | V1fpUF | 4 | 0 | 0 | 5,04044960810504E-05 | -4,297530723 |
| 661 | Case12 | C057 | 7B | V1fpUF | 0 | 0 | NA | 0 | -Inf |
| 662 | Case12 | C057 | 7m | V1fpUF | 0 | 0 | NA | 0 | -Inf |
| 663 | Case12 | C057 | 7op | V1fpUF | 0 | 0 | NA | 0 | -Inf |
| 664 | Case12 | C057 | 8B | V1fpUF | 0 | 0 | NA | 0 | -Inf |
| 665 | Case12 | C057 | 8l | V1fpUF | 0 | 0 | NA | 0 | -Inf |
| 666 | Case12 | C057 | 8m | V1fpUF | 0 | 0 | NA | 0 | -Inf |
| 667 | Case12 | C057 | 8r | V1fpUF | 0 | 0 | NA | 0 | -Inf |
| 668 | Case12 | C057 | 9 | V1fpUF | 0 | 0 | NA | 0 | -Inf |

| | Case | MONKEY | SOURCE | TARGET | TOT | SUP | SLN | FLN | lgFLN |
|---|---|---|---|---|---|---|---|---|---|
| 669 | Case12 | C057 | 9/46d | V1fpUF | 0 | 0 | NA | 0 | -Inf |
| 670 | Case12 | C057 | 9/46v | V1fpUF | 0 | 0 | NA | 0 | -Inf |
| 671 | Case12 | C057 | AIP | V1fpUF | 0 | 0 | NA | 0 | -Inf |
| 672 | Case12 | C057 | CORE | V1fpUF | 1 | 1 | 1 | 1,26011240202626E-05 | -4,899590714 |
| 673 | Case12 | C057 | DP | V1fpUF | 0 | 0 | NA | 0 | -Inf |
| 674 | Case12 | C057 | ENTO | V1fpUF | 0 | 0 | NA | 0 | -Inf |
| 675 | Case12 | C057 | F1 | V1fpUF | 0 | 0 | NA | 0 | -Inf |
| 676 | Case12 | C057 | F2 | V1fpUF | 0 | 0 | NA | 0 | -Inf |
| 677 | Case12 | C057 | F3 | V1fpUF | 0 | 0 | NA | 0 | -Inf |
| 678 | Case12 | C057 | F4 | V1fpUF | 0 | 0 | NA | 0 | -Inf |
| 679 | Case12 | C057 | F5 | V1fpUF | 0 | 0 | NA | 0 | -Inf |
| 680 | Case12 | C057 | F6 | V1fpUF | 0 | 0 | NA | 0 | -Inf |
| 681 | Case12 | C057 | F7 | V1fpUF | 0 | 0 | NA | 0 | -Inf |
| 682 | Case12 | C057 | FST | V1fpUF | 20,5 | 0 | 0 | 0,000258323 | -3,587836853 |
| 683 | Case12 | C057 | Gu | V1fpUF | 0 | 0 | NA | 0 | -Inf |
| 684 | Case12 | C057 | INSULA | V1fpUF | 2 | 0 | 0 | 2,52022480405252E-05 | -4,598560718 |
| 685 | Case12 | C057 | IPa | V1fpUF | 8,5 | 0 | 0 | 0,00010711 | -3,970171788 |
| 686 | Case12 | C057 | LB | V1fpUF | 3 | 0 | 0 | 3,78033720607878E-05 | -4,422469459 |
| 687 | Case12 | C057 | LIP | V1fpUF | 5 | 0 | 0 | 6,3005620101313E-05 | -4,20062071 |
| 688 | Case12 | C057 | MB | V1fpUF | 0 | 0 | NA | 0 | -Inf |
| 689 | Case12 | C057 | MIP | V1fpUF | 0 | 0 | NA | 0 | -Inf |
| 690 | Case12 | C057 | MST | V1fpUF | 362 | 1 | 0,002762431 | 0,004561607 | -2,340882144 |
| 691 | Case12 | C057 | MTc | V1fpUF | 8 | 0 | 0 | 0,000100809 | -3,996500727 |
| 692 | Case12 | C057 | MTp | V1fpUF | 20 | 0 | 0 | 0,000252022 | -3,598560718 |
| 693 | Case12 | C057 | OPAl | V1fpUF | 0 | 0 | NA | 0 | -Inf |
| 694 | Case12 | C057 | OPro | V1fpUF | 0 | 0 | NA | 0 | -Inf |
| 695 | Case12 | C057 | PBc | V1fpUF | 14 | 0 | 0 | 0,000176416 | -3,753462678 |
| 696 | Case12 | C057 | PBr | V1fpUF | 5 | 0 | 0 | 6,3005620101313E-05 | -4,20062071 |
| 697 | Case12 | C057 | PERI | V1fpUF | 14 | 0 | 0 | 0,000176416 | -3,753462678 |
| 698 | Case12 | C057 | PGa | V1fpUF | 52 | 0 | 0 | 0,000655258 | -3,18358737 |
| 699 | Case12 | C057 | Pi | V1fpUF | 0 | 0 | NA | 0 | -Inf |
| 700 | Case12 | C057 | PIP | V1fpUF | 6 | 0 | 0 | 7,56067441215756E-05 | -4,121439464 |
| 701 | Case12 | C057 | Pir | V1fpUF | 0 | 0 | NA | 0 | -Inf |
| 702 | Case12 | C057 | POLE | V1fpUF | 0 | 0 | NA | 0 | -Inf |
| 703 | Case12 | C057 | Pro.St | V1fpUF | 2,5 | 1 | 0,4 | 3,15028100506565E-05 | -4,501650705 |
| 704 | Case12 | C057 | ProM | V1fpUF | 0 | 0 | NA | 0 | -Inf |
| 705 | Case12 | C057 | SII | V1fpUF | 0 | 0 | NA | 0 | -Inf |
| 706 | Case12 | C057 | STPc | V1fpUF | 68 | 0 | 0 | 0,000856876 | -3,067081801 |
| 707 | Case12 | C057 | STPi | V1fpUF | 15,5 | 0 | 0 | 0,000195317 | -3,709259016 |
| 708 | Case12 | C057 | STPr | V1fpUF | 11,5 | 2,5 | 0,217391304 | 0,000144913 | -3,838892874 |
| 709 | Case12 | C057 | SUBI | V1fpUF | 0 | 0 | NA | 0 | -Inf |
| 710 | Case12 | C057 | TEa/ma | V1fpUF | 3,5 | 0 | 0 | 4,41039340709191E-05 | -4,35552267 |
| 711 | Case12 | C057 | TEa/mp | V1fpUF | 3 | 0 | 0 | 3,78033720607878E-05 | -4,422469459 |
| 712 | Case12 | C057 | TEad | V1fpUF | 1,5 | 0 | 0 | 1,89016860303939E-05 | -4,723499455 |
| 713 | Case12 | C057 | TEav | V1fpUF | 1 | 0 | 0 | 1,26011240202626E-05 | -4,899590714 |
| 714 | Case12 | C057 | TEO | V1fpUF | 1 | 0 | 0 | 1,26011240202626E-05 | -4,899590714 |
| 715 | Case12 | C057 | TEOm | V1fpUF | 0 | 0 | NA | 0 | -Inf |
| 716 | Case12 | C057 | TEpd | V1fpUF | 1 | 0 | 0 | 1,26011240202626E-05 | -4,899590714 |
| 717 | Case12 | C057 | TEpv | V1fpUF | 284,5 | 3 | 0,010544815 | 0,00358502 | -2,445508443 |
| 718 | Case12 | C057 | TH_TF | V1fpUF | 241 | 1 | 0,004149378 | 0,003036871 | -2,517573672 |
| 719 | Case12 | C057 | TPt | V1fpUF | 10 | 0 | 0 | 0,000126011 | -3,899590714 |
| 720 | Case12 | C057 | V1c | V1fpUF | 0 | 0 | NA | 0 | -Inf |
| 721 | Case12 | C057 | V1fpLF | V1fpUF | 4 | 2 | 0,5 | 5,04044960810504E-05 | -4,297530723 |
| 722 | Case12 | C057 | V1fpUF | V1fpUF | 0 | 0 | NA | 0 | -Inf |
| 723 | Case12 | C057 | V1pcLF | V1fpUF | 0 | 0 | NA | 0 | -Inf |
| 724 | Case12 | C057 | V1pcUF | V1fpUF | 0 | 0 | NA | 0 | -Inf |
| 725 | Case12 | C057 | V2c | V1fpUF | 0 | 0 | NA | 0 | -Inf |
| 726 | Case12 | C057 | V2fpLF | V1fpUF | 26 | 12 | 0,461538462 | 0,000327629 | -3,484617366 |
| 727 | Case12 | C057 | V2fpUF | V1fpUF | 14591 | 7220 | 0,494825577 | 0,183863001 | -0,735505657 |
| 728 | Case12 | C057 | V2pcLF | V1fpUF | 0 | 0 | NA | 0 | -Inf |
| 729 | Case12 | C057 | V2pcUF | V1fpUF | 0 | 0 | NA | 0 | -Inf |
| 730 | Case12 | C057 | V3A | V1fpUF | 0 | 0 | NA | 0 | -Inf |
| 731 | Case12 | C057 | V3c | V1fpUF | 0 | 0 | NA | 0 | -Inf |
| 732 | Case12 | C057 | V3fpLF | V1fpUF | 0 | 0 | NA | 0 | -Inf |
| 733 | Case12 | C057 | V3fpUF | V1fpUF | 845,5 | 229,5 | 0,27143702 | 0,01065425 | -1,972477102 |
| 734 | Case12 | C057 | V3pcLF | V1fpUF | 0 | 0 | NA | 0 | -Inf |
| 735 | Case12 | C057 | V3pcUF | V1fpUF | 0 | 0 | NA | 0 | -Inf |
| 736 | Case12 | C057 | V4c | V1fpUF | 0 | 0 | NA | 0 | -Inf |
| 737 | Case12 | C057 | V4LF | V1fpUF | 0 | 0 | NA | 0 | -Inf |
| 738 | Case12 | C057 | V4t | V1fpUF | 0 | 0 | NA | 0 | -Inf |
| 739 | Case12 | C057 | V4UF | V1fpUF | 1195,5 | 214 | 0,179004601 | 0,015064644 | -1,822041134 |
| 740 | Case12 | C057 | V6 | V1fpUF | 0 | 0 | NA | 0 | -Inf |
| 741 | Case12 | C057 | V6A | V1fpUF | 150 | 58 | 0,386666667 | 0,001890169 | -2,723499455 |
| 742 | Case12 | C057 | VIP | V1fpUF | 4 | 0 | 0 | 5,04044960810504E-05 | -4,297530723 |
| 849 | Case13 | C083 | 1 | V1fpUF | 0 | 0 | NA | 0 | -Inf |

| | Case | MONKEY | SOURCE | TARGET | TOT | SUP | SLN | FLN | lgFLN |
|---|---|---|---|---|---|---|---|---|---|
| 850 | Case13 | C083 | 10 | V1fpUF | 0 | 0 | NA | 0 | -Inf |
| 851 | Case13 | C083 | 11 | V1fpUF | 0 | 0 | NA | 0 | -Inf |
| 852 | Case13 | C083 | 12 | V1fpUF | 0 | 0 | NA | 0 | -Inf |
| 853 | Case13 | C083 | 13 | V1fpUF | 0 | 0 | NA | 0 | -Inf |
| 854 | Case13 | C083 | 14 | V1fpUF | 0 | 0 | NA | 0 | -Inf |
| 855 | Case13 | C083 | 2 | V1fpUF | 0 | 0 | NA | 0 | -Inf |
| 856 | Case13 | C083 | 23 | V1fpUF | 2 | 2 | 1 | 2,52022480405252E-05 | -4,598560718 |
| 857 | Case13 | C083 | 24a | V1fpUF | 0 | 0 | NA | 0 | -Inf |
| 858 | Case13 | C083 | 24b | V1fpUF | 0 | 0 | NA | 0 | -Inf |
| 859 | Case13 | C083 | 24c | V1fpUF | 0 | 0 | NA | 0 | -Inf |
| 860 | Case13 | C083 | 24d | V1fpUF | 0 | 0 | NA | 0 | -Inf |
| 861 | Case13 | C083 | 25 | V1fpUF | 0 | 0 | NA | 0 | -Inf |
| 862 | Case13 | C083 | 29/30 | V1fpUF | 0 | 0 | NA | 0 | -Inf |
| 863 | Case13 | C083 | 3 | V1fpUF | 0 | 0 | NA | 0 | -Inf |
| 864 | Case13 | C083 | 31 | V1fpUF | 0 | 0 | NA | 0 | -Inf |
| 865 | Case13 | C083 | 32 | V1fpUF | 0 | 0 | NA | 0 | -Inf |
| 866 | Case13 | C083 | 44 | V1fpUF | 0 | 0 | NA | 0 | -Inf |
| 867 | Case13 | C083 | 45A | V1fpUF | 0 | 0 | NA | 0 | -Inf |
| 868 | Case13 | C083 | 45B | V1fpUF | 0 | 0 | NA | 0 | -Inf |
| 869 | Case13 | C083 | 46d | V1fpUF | 0 | 0 | NA | 0 | -Inf |
| 870 | Case13 | C083 | 46v | V1fpUF | 0 | 0 | NA | 0 | -Inf |
| 871 | Case13 | C083 | 5 | V1fpUF | 0 | 0 | NA | 0 | -Inf |
| 872 | Case13 | C083 | 7A | V1fpUF | 8 | 8 | 1 | 0,000100809 | -3,996500727 |
| 873 | Case13 | C083 | 7B | V1fpUF | 0 | 0 | NA | 0 | -Inf |
| 874 | Case13 | C083 | 7m | V1fpUF | 0 | 0 | NA | 0 | -Inf |
| 875 | Case13 | C083 | 7op | V1fpUF | 0 | 0 | NA | 0 | -Inf |
| 876 | Case13 | C083 | 8B | V1fpUF | 0 | 0 | NA | 0 | -Inf |
| 877 | Case13 | C083 | 8l | V1fpUF | 0 | 0 | NA | 0 | -Inf |
| 878 | Case13 | C083 | 8m | V1fpUF | 0 | 0 | NA | 0 | -Inf |
| 879 | Case13 | C083 | 8r | V1fpUF | 0 | 0 | NA | 0 | -Inf |
| 880 | Case13 | C083 | 9 | V1fpUF | 0 | 0 | NA | 0 | -Inf |
| 881 | Case13 | C083 | 9/46d | V1fpUF | 0 | 0 | NA | 0 | -Inf |
| 882 | Case13 | C083 | 9/46v | V1fpUF | 0 | 0 | NA | 0 | -Inf |
| 883 | Case13 | C083 | AIP | V1fpUF | 0 | 0 | NA | 0 | -Inf |
| 884 | Case13 | C083 | CORE | V1fpUF | 5 | 5 | 1 | 6,3005620101313E-05 | -4,20062071 |
| 885 | Case13 | C083 | DP | V1fpUF | 0 | 0 | NA | 0 | -Inf |
| 886 | Case13 | C083 | ENTO | V1fpUF | 0 | 0 | NA | 0 | -Inf |
| 887 | Case13 | C083 | F1 | V1fpUF | 0 | 0 | NA | 0 | -Inf |
| 888 | Case13 | C083 | F2 | V1fpUF | 0 | 0 | NA | 0 | -Inf |
| 889 | Case13 | C083 | F3 | V1fpUF | 0 | 0 | NA | 0 | -Inf |
| 890 | Case13 | C083 | F4 | V1fpUF | 0 | 0 | NA | 0 | -Inf |
| 891 | Case13 | C083 | F5 | V1fpUF | 0 | 0 | NA | 0 | -Inf |
| 892 | Case13 | C083 | F6 | V1fpUF | 0 | 0 | NA | 0 | -Inf |
| 893 | Case13 | C083 | F7 | V1fpUF | 0 | 0 | NA | 0 | -Inf |
| 894 | Case13 | C083 | FST | V1fpUF | 35,5 | 35,5 | 1 | 0,00044734 | -3,349362361 |
| 895 | Case13 | C083 | Gu | V1fpUF | 0 | 0 | NA | 0 | -Inf |
| 896 | Case13 | C083 | INSULA | V1fpUF | 11,5 | 11,5 | 1 | 0,000144913 | -3,838892874 |
| 897 | Case13 | C083 | IPa | V1fpUF | 2 | 2 | 1 | 2,52022480405252E-05 | -4,598560718 |
| 898 | Case13 | C083 | LB | V1fpUF | 22,5 | 22,5 | 1 | 0,000283525 | -3,547408196 |
| 899 | Case13 | C083 | LIP | V1fpUF | 10 | 10 | 1 | 0,000126011 | -3,899590714 |
| 900 | Case13 | C083 | MB | V1fpUF | 14,5 | 14,5 | 1 | 0,000182716 | -3,738222712 |
| 901 | Case13 | C083 | MIP | V1fpUF | 0 | 0 | NA | 0 | -Inf |
| 902 | Case13 | C083 | MST | V1fpUF | 890,5 | 890,5 | 1 | 0,011221301 | -1,94995679 |
| 903 | Case13 | C083 | MTc | V1fpUF | 48 | 48 | 1 | 0,000604854 | -3,218349477 |
| 904 | Case13 | C083 | MTp | V1fpUF | 482 | 482 | 1 | 0,006073742 | -2,216543676 |
| 905 | Case13 | C083 | OPAl | V1fpUF | 0 | 0 | NA | 0 | -Inf |
| 906 | Case13 | C083 | OPro | V1fpUF | 0 | 0 | NA | 0 | -Inf |
| 907 | Case13 | C083 | PBc | V1fpUF | 30 | 30 | 1 | 0,000378034 | -3,422469459 |
| 908 | Case13 | C083 | PBr | V1fpUF | 0 | 0 | NA | 0 | -Inf |
| 909 | Case13 | C083 | PERI | V1fpUF | 9 | 9 | 1 | 0,00011341 | -3,945348205 |
| 910 | Case13 | C083 | PGa | V1fpUF | 64,5 | 64,5 | 1 | 0,000812772 | -3,090030999 |
| 911 | Case13 | C083 | Pi | V1fpUF | 0 | 0 | NA | 0 | -Inf |
| 912 | Case13 | C083 | PIP | V1fpUF | 26 | 26 | 1 | 0,000327629 | -3,484617366 |
| 913 | Case13 | C083 | Pir | V1fpUF | 0 | 0 | NA | 0 | -Inf |
| 914 | Case13 | C083 | POLE | V1fpUF | 0 | 0 | NA | 0 | -Inf |
| 915 | Case13 | C083 | Pro.St | V1fpUF | 50,5 | 50,5 | 1 | 0,000636357 | -3,196299336 |
| 916 | Case13 | C083 | ProM | V1fpUF | 0 | 0 | NA | 0 | -Inf |
| 917 | Case13 | C083 | SII | V1fpUF | 0 | 0 | NA | 0 | -Inf |
| 918 | Case13 | C083 | STPc | V1fpUF | 143,5 | 143,5 | 1 | 0,001808261 | -2,742738813 |
| 919 | Case13 | C083 | STPi | V1fpUF | 12 | 12 | 1 | 0,000151213 | -3,820409468 |
| 920 | Case13 | C083 | STPr | V1fpUF | 0 | 0 | NA | 0 | -Inf |
| 921 | Case13 | C083 | SUBI | V1fpUF | 0 | 0 | NA | 0 | -Inf |
| 922 | Case13 | C083 | TEa/ma | V1fpUF | 1 | 1 | 1 | 1,26011240202626E-05 | -4,899590714 |
| 923 | Case13 | C083 | TEa/mp | V1fpUF | 1 | 1 | 1 | 1,26011240202626E-05 | -4,899590714 |
| 924 | Case13 | C083 | TEad | V1fpUF | 1 | 1 | 1 | 1,26011240202626E-05 | -4,899590714 |

| | Case | MONKEY | SOURCE | TARGET | TOT | SUP | SLN | FLN | lgFLN |
|---|---|---|---|---|---|---|---|---|---|
| 925 | Case13 | C083 | TEav | V1fpUF | 6 | 6 | 1 | 7,56067441215756E-05 | -4,121439464 |
| 926 | Case13 | C083 | TEO | V1fpUF | 8 | 8 | 1 | 0,000100809 | -3,996500727 |
| 927 | Case13 | C083 | TEOm | V1fpUF | 1,5 | 1,5 | 1 | 1,89016860303939E-05 | -4,723499455 |
| 928 | Case13 | C083 | TEpd | V1fpUF | 2 | 2 | 1 | 2,52022480405252E-05 | -4,598560718 |
| 929 | Case13 | C083 | TEpv | V1fpUF | 537,5 | 537,5 | 1 | 0,006773104 | -2,169212245 |
| 930 | Case13 | C083 | TH_TF | V1fpUF | 1258 | 1258 | 1 | 0,015852214 | -1,799910073 |
| 931 | Case13 | C083 | TPt | V1fpUF | 14 | 14 | 1 | 0,000176416 | -3,753462678 |
| 932 | Case13 | C083 | V1c | V1fpUF | 12 | 12 | 1 | 0,000151213 | -3,820409468 |
| 933 | Case13 | C083 | V1fpLF | V1fpUF | 862 | 862 | 1 | 0,010862169 | -1,964083448 |
| 934 | Case13 | C083 | V1fpUF | V1fpUF | 0 | 0 | NA | 0 | -Inf |
| 935 | Case13 | C083 | V1pcLF | V1fpUF | 0 | 0 | NA | 0 | -Inf |
| 936 | Case13 | C083 | V1pcUF | V1fpUF | 2 | 2 | 1 | 2,52022480405252E-05 | -4,598560718 |
| 937 | Case13 | C083 | V2c | V1fpUF | 594 | 594 | 1 | 0,007485068 | -2,125804269 |
| 938 | Case13 | C083 | V2fpLF | V1fpUF | 22 | 22 | 1 | 0,000277225 | -3,557168033 |
| 939 | Case13 | C083 | V2fpUF | V1fpUF | 24581 | 24581 | 1 | 0,30974823 | -0,508991167 |
| 940 | Case13 | C083 | V2pcLF | V1fpUF | 0 | 0 | NA | 0 | -Inf |
| 941 | Case13 | C083 | V2pcUF | V1fpUF | 0 | 0 | NA | 0 | -Inf |
| 942 | Case13 | C083 | V3A | V1fpUF | 0 | 0 | NA | 0 | -Inf |
| 943 | Case13 | C083 | V3c | V1fpUF | 2 | 2 | 1 | 2,52022480405252E-05 | -4,598560718 |
| 944 | Case13 | C083 | V3fpLF | V1fpUF | 0 | 0 | NA | 0 | -Inf |
| 945 | Case13 | C083 | V3fpUF | V1fpUF | 775 | 775 | 1 | 0,009765871 | -2,010289012 |
| 946 | Case13 | C083 | V3pcLF | V1fpUF | 0 | 0 | NA | 0 | -Inf |
| 947 | Case13 | C083 | V3pcUF | V1fpUF | 2 | 2 | 1 | 2,52022480405252E-05 | -4,598560718 |
| 948 | Case13 | C083 | V4c | V1fpUF | 1 | 1 | 1 | 1,26011240202626E-05 | -4,899590714 |
| 949 | Case13 | C083 | V4LF | V1fpUF | 32 | 32 | 1 | 0,000403236 | -3,394440736 |
| 950 | Case13 | C083 | V4t | V1fpUF | 0 | 0 | NA | 0 | -Inf |
| 951 | Case13 | C083 | V4UF | V1fpUF | 816,5 | 816,5 | 1 | 0,010288818 | -1,987634525 |
| 952 | Case13 | C083 | V6 | V1fpUF | 12 | 12 | 1 | 0,000151213 | -3,820409468 |
| 953 | Case13 | C083 | V6A | V1fpUF | 224 | 224 | 1 | 0,002822652 | -2,549342696 |
| 954 | Case13 | C083 | VIP | V1fpUF | 6 | 6 | 1 | 7,56067441215756E-05 | -4,121439464 |
| 743 | Case14 | C060 | 1 | V1fpUF | 0 | 0 | NA | 0 | -Inf |
| 744 | Case14 | C060 | 10 | V1fpUF | 0 | 0 | NA | 0 | -Inf |
| 745 | Case14 | C060 | 11 | V1fpUF | 0 | 0 | NA | 0 | -Inf |
| 746 | Case14 | C060 | 12 | V1fpUF | 0 | 0 | NA | 0 | -Inf |
| 747 | Case14 | C060 | 13 | V1fpUF | 0 | 0 | NA | 0 | -Inf |
| 748 | Case14 | C060 | 14 | V1fpUF | 0 | 0 | NA | 0 | -Inf |
| 749 | Case14 | C060 | 2 | V1fpUF | 0 | 0 | NA | 0 | -Inf |
| 750 | Case14 | C060 | 23 | V1fpUF | 0 | 0 | NA | 0 | -Inf |
| 751 | Case14 | C060 | 24a | V1fpUF | 0 | 0 | NA | 0 | -Inf |
| 752 | Case14 | C060 | 24b | V1fpUF | 0 | 0 | NA | 0 | -Inf |
| 753 | Case14 | C060 | 24c | V1fpUF | 0 | 0 | NA | 0 | -Inf |
| 754 | Case14 | C060 | 24d | V1fpUF | 0 | 0 | NA | 0 | -Inf |
| 755 | Case14 | C060 | 25 | V1fpUF | 0 | 0 | NA | 0 | -Inf |
| 756 | Case14 | C060 | 29/30 | V1fpUF | 0 | 0 | NA | 0 | -Inf |
| 757 | Case14 | C060 | 3 | V1fpUF | 0 | 0 | NA | 0 | -Inf |
| 758 | Case14 | C060 | 31 | V1fpUF | 0 | 0 | NA | 0 | -Inf |
| 759 | Case14 | C060 | 32 | V1fpUF | 0 | 0 | NA | 0 | -Inf |
| 760 | Case14 | C060 | 44 | V1fpUF | 0 | 0 | NA | 0 | -Inf |
| 761 | Case14 | C060 | 45A | V1fpUF | 0 | 0 | NA | 0 | -Inf |
| 762 | Case14 | C060 | 45B | V1fpUF | 0 | 0 | NA | 0 | -Inf |
| 763 | Case14 | C060 | 46d | V1fpUF | 0 | 0 | NA | 0 | -Inf |
| 764 | Case14 | C060 | 46v | V1fpUF | 0 | 0 | NA | 0 | -Inf |
| 765 | Case14 | C060 | 5 | V1fpUF | 0 | 0 | NA | 0 | -Inf |
| 766 | Case14 | C060 | 7A | V1fpUF | 0 | 0 | NA | 0 | -Inf |
| 767 | Case14 | C060 | 7B | V1fpUF | 0 | 0 | NA | 0 | -Inf |
| 768 | Case14 | C060 | 7m | V1fpUF | 0 | 0 | NA | 0 | -Inf |
| 769 | Case14 | C060 | 7op | V1fpUF | 2 | 0 | 0 | 2,52022480405252E-05 | -4,598560718 |
| 770 | Case14 | C060 | 8B | V1fpUF | 0 | 0 | NA | 0 | -Inf |
| 771 | Case14 | C060 | 8l | V1fpUF | 0 | 0 | NA | 0 | -Inf |
| 772 | Case14 | C060 | 8m | V1fpUF | 0 | 0 | NA | 0 | -Inf |
| 773 | Case14 | C060 | 8r | V1fpUF | 0 | 0 | NA | 0 | -Inf |
| 774 | Case14 | C060 | 9 | V1fpUF | 0 | 0 | NA | 0 | -Inf |
| 775 | Case14 | C060 | 9/46d | V1fpUF | 0 | 0 | NA | 0 | -Inf |
| 776 | Case14 | C060 | 9/46v | V1fpUF | 0 | 0 | NA | 0 | -Inf |
| 777 | Case14 | C060 | AIP | V1fpUF | 0 | 0 | NA | 0 | -Inf |
| 778 | Case14 | C060 | CORE | V1fpUF | 2 | 0 | 0 | 2,52022480405252E-05 | -4,598560718 |
| 779 | Case14 | C060 | DP | V1fpUF | 0 | 0 | NA | 0 | -Inf |
| 780 | Case14 | C060 | ENTO | V1fpUF | 0 | 0 | NA | 0 | -Inf |
| 781 | Case14 | C060 | F1 | V1fpUF | 0 | 0 | NA | 0 | -Inf |
| 782 | Case14 | C060 | F2 | V1fpUF | 0 | 0 | NA | 0 | -Inf |
| 783 | Case14 | C060 | F3 | V1fpUF | 0 | 0 | NA | 0 | -Inf |
| 784 | Case14 | C060 | F4 | V1fpUF | 0 | 0 | NA | 0 | -Inf |
| 785 | Case14 | C060 | F5 | V1fpUF | 0 | 0 | NA | 0 | -Inf |
| 786 | Case14 | C060 | F6 | V1fpUF | 0 | 0 | NA | 0 | -Inf |
| 787 | Case14 | C060 | F7 | V1fpUF | 0 | 0 | NA | 0 | -Inf |

| | Case | MONKEY | SOURCE | TARGET | TOT | SUP | SLN | FLN | lgFLN |
|---|---|---|---|---|---|---|---|---|---|
| 788 | Case14 | C060 | FST | V1fpUF | 98 | 2 | 0,020408163 | 0,00123491 | -2,908364638 |
| 789 | Case14 | C060 | Gu | V1fpUF | 0 | 0 | NA | 0 | -Inf |
| 790 | Case14 | C060 | INSULA | V1fpUF | 3,5 | 0 | 0 | 4,41039340709191E-05 | -4,35552267 |
| 791 | Case14 | C060 | IPa | V1fpUF | 5 | 1,5 | 0,3 | 6,3005620101313E-05 | -4,20062071 |
| 792 | Case14 | C060 | LB | V1fpUF | 18 | 0 | 0 | 0,00022682 | -3,644318209 |
| 793 | Case14 | C060 | LIP | V1fpUF | 4 | 0 | 0 | 5,04044960810504E-05 | -4,297530723 |
| 794 | Case14 | C060 | MB | V1fpUF | 8 | 0 | 0 | 0,000100809 | -3,996500727 |
| 795 | Case14 | C060 | MIP | V1fpUF | 0 | 0 | NA | 0 | -Inf |
| 796 | Case14 | C060 | MST | V1fpUF | 224 | 0 | 0 | 0,002822652 | -2,549342696 |
| 797 | Case14 | C060 | MTc | V1fpUF | 224 | 2 | 0,008928571 | 0,002822652 | -2,549342696 |
| 798 | Case14 | C060 | MTp | V1fpUF | 666 | 16 | 0,024024024 | 0,008392349 | -2,076116485 |
| 799 | Case14 | C060 | OPAl | V1fpUF | 0 | 0 | NA | 0 | -Inf |
| 800 | Case14 | C060 | OPro | V1fpUF | 0 | 0 | NA | 0 | -Inf |
| 801 | Case14 | C060 | PBc | V1fpUF | 24 | 0 | 0 | 0,000302427 | -3,519379472 |
| 802 | Case14 | C060 | PBr | V1fpUF | 12,5 | 0 | 0 | 0,000157514 | -3,802680701 |
| 803 | Case14 | C060 | PERI | V1fpUF | 63,5 | 0 | 0 | 0,000800171 | -3,096816989 |
| 804 | Case14 | C060 | PGa | V1fpUF | 4 | 0 | 0 | 5,04044960810504E-05 | -4,297530723 |
| 805 | Case14 | C060 | Pi | V1fpUF | 0 | 0 | NA | 0 | -Inf |
| 806 | Case14 | C060 | PIP | V1fpUF | 88 | 8 | 0,090909091 | 0,001108899 | -2,955108042 |
| 807 | Case14 | C060 | Pir | V1fpUF | 0 | 0 | NA | 0 | -Inf |
| 808 | Case14 | C060 | POLE | V1fpUF | 0 | 0 | NA | 0 | -Inf |
| 809 | Case14 | C060 | Pro.St | V1fpUF | 2 | 0 | 0 | 2,52022480405252E-05 | -4,598560718 |
| 810 | Case14 | C060 | ProM | V1fpUF | 0 | 0 | NA | 0 | -Inf |
| 811 | Case14 | C060 | SII | V1fpUF | 0 | 0 | NA | 0 | -Inf |
| 812 | Case14 | C060 | STPc | V1fpUF | 36 | 0 | 0 | 0,00045364 | -3,343288213 |
| 813 | Case14 | C060 | STPi | V1fpUF | 12 | 0 | 0 | 0,000151213 | -3,820409468 |
| 814 | Case14 | C060 | STPr | V1fpUF | 5 | 0 | 0 | 6,3005620101313E-05 | -4,20062071 |
| 815 | Case14 | C060 | SUBI | V1fpUF | 0 | 0 | NA | 0 | -Inf |
| 816 | Case14 | C060 | TEa/ma | V1fpUF | 1 | 0 | 0 | 1,26011240202626E-05 | -4,899590714 |
| 817 | Case14 | C060 | TEa/mp | V1fpUF | 1 | 0 | 0 | 1,26011240202626E-05 | -4,899590714 |
| 818 | Case14 | C060 | TEad | V1fpUF | 0 | 0 | NA | 0 | -Inf |
| 819 | Case14 | C060 | TEav | V1fpUF | 13,5 | 0 | 0 | 0,000170115 | -3,769256946 |
| 820 | Case14 | C060 | TEO | V1fpUF | 234 | 10 | 0,042735043 | 0,002948663 | -2,530374857 |
| 821 | Case14 | C060 | TEOm | V1fpUF | 26 | 0 | 0 | 0,000327629 | -3,484617366 |
| 822 | Case14 | C060 | TEpd | V1fpUF | 6 | 1 | 0,166666667 | 7,56067441215756E-05 | -4,121439464 |
| 823 | Case14 | C060 | TEpv | V1fpUF | 788 | 76 | 0,096446701 | 0,009929686 | -2,003064497 |
| 824 | Case14 | C060 | TH_TF | V1fpUF | 763,5 | 18 | 0,023575639 | 0,009620958 | -2,016781673 |
| 825 | Case14 | C060 | TPt | V1fpUF | 6 | 0 | 0 | 7,56067441215756E-05 | -4,121439464 |
| 826 | Case14 | C060 | V1c | V1fpUF | 0 | 0 | NA | 0 | -Inf |
| 827 | Case14 | C060 | V1fpLF | V1fpUF | 270 | 94 | 0,348148148 | 0,003402303 | -2,46822695 |
| 828 | Case14 | C060 | V1fpUF | V1fpUF | 0 | 0 | NA | 0 | -Inf |
| 829 | Case14 | C060 | V1pcLF | V1fpUF | 0 | 0 | NA | 0 | -Inf |
| 830 | Case14 | C060 | V1pcUF | V1fpUF | 62 | 8 | 0,129032258 | 0,00078127 | -3,107199025 |
| 831 | Case14 | C060 | V2c | V1fpUF | 88 | 12 | 0,136363636 | 0,001108899 | -2,955108042 |
| 832 | Case14 | C060 | V2fpLF | V1fpUF | 0 | 0 | NA | 0 | -Inf |
| 833 | Case14 | C060 | V2fpUF | V1fpUF | 20838 | 8734 | 0,419138113 | 0,262582222 | -0,58073468 |
| 834 | Case14 | C060 | V2pcLF | V1fpUF | 0 | 0 | NA | 0 | -Inf |
| 835 | Case14 | C060 | V2pcUF | V1fpUF | 626 | 38 | 0,060702875 | 0,007888304 | -2,103016381 |
| 836 | Case14 | C060 | V3A | V1fpUF | 0 | 0 | NA | 0 | -Inf |
| 837 | Case14 | C060 | V3c | V1fpUF | 4 | 0 | 0 | 5,04044960810504E-05 | -4,297530723 |
| 838 | Case14 | C060 | V3fpLF | V1fpUF | 0 | 0 | NA | 0 | -Inf |
| 839 | Case14 | C060 | V3fpUF | V1fpUF | 2464 | 680 | 0,275974026 | 0,03104917 | -1,507950011 |
| 840 | Case14 | C060 | V3pcLF | V1fpUF | 0 | 0 | NA | 0 | -Inf |
| 841 | Case14 | C060 | V3pcUF | V1fpUF | 518 | 76 | 0,146718147 | 0,006527382 | -2,185260954 |
| 842 | Case14 | C060 | V4c | V1fpUF | 0 | 0 | NA | 0 | -Inf |
| 843 | Case14 | C060 | V4LF | V1fpUF | 2 | 0 | 0 | 2,52022480405252E-05 | -4,598560718 |
| 844 | Case14 | C060 | V4t | V1fpUF | 0 | 0 | NA | 0 | -Inf |
| 845 | Case14 | C060 | V4UF | V1fpUF | 1516 | 326 | 0,215039578 | 0,019103304 | -1,718891513 |
| 846 | Case14 | C060 | V6 | V1fpUF | 0 | 0 | NA | 0 | -Inf |
| 847 | Case14 | C060 | V6A | V1fpUF | 0 | 0 | NA | 0 | -Inf |
| 848 | Case14 | C060 | VIP | V1fpUF | 0 | 0 | NA | 0 | -Inf |
| 1273 | Case15 | M101LH | 1 | V2c | 0 | 0 | NA | 0 | -Inf |
| 1274 | Case15 | M101LH | 10 | V2c | 0 | 0 | NA | 0 | -Inf |
| 1275 | Case15 | M101LH | 11 | V2c | 0 | 0 | NA | 0 | -Inf |
| 1276 | Case15 | M101LH | 12 | V2c | 0 | 0 | NA | 0 | -Inf |
| 1277 | Case15 | M101LH | 13 | V2c | 0 | 0 | NA | 0 | -Inf |
| 1278 | Case15 | M101LH | 14 | V2c | 0 | 0 | NA | 0 | -Inf |
| 1279 | Case15 | M101LH | 2 | V2c | 0 | 0 | NA | 0 | -Inf |
| 1280 | Case15 | M101LH | 23 | V2c | 0 | 0 | NA | 0 | -Inf |
| 1281 | Case15 | M101LH | 24a | V2c | 0 | 0 | NA | 0 | -Inf |
| 1282 | Case15 | M101LH | 24b | V2c | 0 | 0 | NA | 0 | -Inf |
| 1283 | Case15 | M101LH | 24c | V2c | 0 | 0 | NA | 0 | -Inf |
| 1284 | Case15 | M101LH | 24d | V2c | 0 | 0 | NA | 0 | -Inf |
| 1285 | Case15 | M101LH | 25 | V2c | 0 | 0 | NA | 0 | -Inf |
| 1286 | Case15 | M101LH | 29/30 | V2c | 0 | 0 | NA | 0 | -Inf |

| | Case | MONKEY | SOURCE | TARGET | TOT | SUP | SLN | FLN | lgFLN |
|---|---|---|---|---|---|---|---|---|---|
| 1287 | Case15 | M101LH | 3 | V2c | 0 | 0 | NA | 0 | -Inf |
| 1288 | Case15 | M101LH | 31 | V2c | 0 | 0 | NA | 0 | -Inf |
| 1289 | Case15 | M101LH | 32 | V2c | 0 | 0 | NA | 0 | -Inf |
| 1290 | Case15 | M101LH | 44 | V2c | 0 | 0 | NA | 0 | -Inf |
| 1291 | Case15 | M101LH | 45A | V2c | 0 | 0 | NA | 0 | -Inf |
| 1292 | Case15 | M101LH | 45B | V2c | 0 | 0 | NA | 0 | -Inf |
| 1293 | Case15 | M101LH | 46d | V2c | 0 | 0 | NA | 0 | -Inf |
| 1294 | Case15 | M101LH | 46v | V2c | 0 | 0 | NA | 0 | -Inf |
| 1295 | Case15 | M101LH | 5 | V2c | 0 | 0 | NA | 0 | -Inf |
| 1296 | Case15 | M101LH | 7A | V2c | 0 | 0 | NA | 0 | -Inf |
| 1297 | Case15 | M101LH | 7B | V2c | 0 | 0 | NA | 0 | -Inf |
| 1298 | Case15 | M101LH | 7m | V2c | 0 | 0 | NA | 0 | -Inf |
| 1299 | Case15 | M101LH | 7op | V2c | 0 | 0 | NA | 0 | -Inf |
| 1300 | Case15 | M101LH | 8B | V2c | 0 | 0 | NA | 0 | -Inf |
| 1301 | Case15 | M101LH | 8l | V2c | 32 | 4 | 0,125 | 4,0196106755835E-05 | -4,395816009 |
| 1302 | Case15 | M101LH | 8m | V2c | 6 | 0 | 0 | 7,53677001671907E-06 | -5,122814737 |
| 1303 | Case15 | M101LH | 8r | V2c | 0 | 0 | NA | 0 | -Inf |
| 1304 | Case15 | M101LH | 9 | V2c | 0 | 0 | NA | 0 | -Inf |
| 1305 | Case15 | M101LH | 9/46d | V2c | 0 | 0 | NA | 0 | -Inf |
| 1306 | Case15 | M101LH | 9/46v | V2c | 0 | 0 | NA | 0 | -Inf |
| 1307 | Case15 | M101LH | AIP | V2c | 0 | 0 | NA | 0 | -Inf |
| 1308 | Case15 | M101LH | CORE | V2c | 0 | 0 | NA | 0 | -Inf |
| 1309 | Case15 | M101LH | DP | V2c | 2 | 0 | 0 | 2,51225667223969E-06 | -5,599935992 |
| 1310 | Case15 | M101LH | ENTO | V2c | 0 | 0 | NA | 0 | -Inf |
| 1311 | Case15 | M101LH | F1 | V2c | 0 | 0 | NA | 0 | -Inf |
| 1312 | Case15 | M101LH | F2 | V2c | 0 | 0 | NA | 0 | -Inf |
| 1313 | Case15 | M101LH | F3 | V2c | 0 | 0 | NA | 0 | -Inf |
| 1314 | Case15 | M101LH | F4 | V2c | 0 | 0 | NA | 0 | -Inf |
| 1315 | Case15 | M101LH | F5 | V2c | 0 | 0 | NA | 0 | -Inf |
| 1316 | Case15 | M101LH | F6 | V2c | 0 | 0 | NA | 0 | -Inf |
| 1317 | Case15 | M101LH | F7 | V2c | 0 | 0 | NA | 0 | -Inf |
| 1318 | Case15 | M101LH | FST | V2c | 847 | 40 | 0,047225502 | 0,001063941 | -2,973082577 |
| 1319 | Case15 | M101LH | Gu | V2c | 0 | 0 | NA | 0 | -Inf |
| 1320 | Case15 | M101LH | INSULA | V2c | 0 | 0 | NA | 0 | -Inf |
| 1321 | Case15 | M101LH | IPa | V2c | 27 | 3 | 0,111111111 | 3,39154650752358E-05 | -4,469602223 |
| 1322 | Case15 | M101LH | LB | V2c | 0 | 0 | NA | 0 | -Inf |
| 1323 | Case15 | M101LH | LIP | V2c | 198 | 9 | 0,045454545 | 0,000248713 | -3,604300797 |
| 1324 | Case15 | M101LH | MB | V2c | 3 | 0 | 0 | 3,76838500835953E-06 | -5,423844733 |
| 1325 | Case15 | M101LH | MIP | V2c | 0 | 0 | NA | 0 | -Inf |
| 1326 | Case15 | M101LH | MST | V2c | 250 | 12 | 0,048 | 0,000314032 | -3,503025979 |
| 1327 | Case15 | M101LH | MTc | V2c | 9292 | 3511 | 0,377851916 | 0,011671944 | -1,932856786 |
| 1328 | Case15 | M101LH | MTp | V2c | 3240 | 631 | 0,194753086 | 0,004069856 | -2,390420977 |
| 1329 | Case15 | M101LH | OPAl | V2c | 0 | 0 | NA | 0 | -Inf |
| 1330 | Case15 | M101LH | OPro | V2c | 0 | 0 | NA | 0 | -Inf |
| 1331 | Case15 | M101LH | PBc | V2c | 0 | 0 | NA | 0 | -Inf |
| 1332 | Case15 | M101LH | PBr | V2c | 0 | 0 | NA | 0 | -Inf |
| 1333 | Case15 | M101LH | PERI | V2c | 30 | 0 | 0 | 3,76838500835953E-05 | -4,423844733 |
| 1334 | Case15 | M101LH | PGa | V2c | 38 | 2 | 0,052631579 | 4,77328767725541E-05 | -4,321182391 |
| 1335 | Case15 | M101LH | Pi | V2c | 0 | 0 | NA | 0 | -Inf |
| 1336 | Case15 | M101LH | PIP | V2c | 320 | 2 | 0,00625 | 0,000401961 | -3,395816009 |
| 1337 | Case15 | M101LH | Pir | V2c | 0 | 0 | NA | 0 | -Inf |
| 1338 | Case15 | M101LH | POLE | V2c | 0 | 0 | NA | 0 | -Inf |
| 1339 | Case15 | M101LH | Pro.St | V2c | 0 | 0 | NA | 0 | -Inf |
| 1340 | Case15 | M101LH | ProM | V2c | 0 | 0 | NA | 0 | -Inf |
| 1341 | Case15 | M101LH | SII | V2c | 0 | 0 | NA | 0 | -Inf |
| 1342 | Case15 | M101LH | STPc | V2c | 3 | 0 | 0 | 3,76838500835953E-06 | -5,423844733 |
| 1343 | Case15 | M101LH | STPi | V2c | 9 | 2 | 0,222222222 | 1,13051550250786E-05 | -4,946723478 |
| 1344 | Case15 | M101LH | STPr | V2c | 0 | 0 | NA | 0 | -Inf |
| 1345 | Case15 | M101LH | SUBI | V2c | 0 | 0 | NA | 0 | -Inf |
| 1346 | Case15 | M101LH | TEa/ma | V2c | 1 | 0 | 0 | 1,25612833611984E-06 | -5,900965987 |
| 1347 | Case15 | M101LH | TEa/mp | V2c | 22 | 0 | 0 | 2,76348233946366E-05 | -4,558543307 |
| 1348 | Case15 | M101LH | TEad | V2c | 19 | 0 | 0 | 2,3866438386277E-05 | -4,622212386 |
| 1349 | Case15 | M101LH | TEav | V2c | 17 | 0 | 0 | 2,13541817140374E-05 | -4,670517066 |
| 1350 | Case15 | M101LH | TEO | V2c | 1930 | 87 | 0,04507772 | 0,002424328 | -2,615408678 |
| 1351 | Case15 | M101LH | TEOm | V2c | 1989 | 70 | 0,035193565 | 0,002498439 | -2,602331204 |
| 1352 | Case15 | M101LH | TEpd | V2c | 285 | 4 | 0,014035088 | 0,000357997 | -3,446121127 |
| 1353 | Case15 | M101LH | TEpv | V2c | 298 | 8 | 0,026845638 | 0,000374326 | -3,426749723 |
| 1354 | Case15 | M101LH | TH_TF | V2c | 723 | 5 | 0,006915629 | 0,000908181 | -3,04182769 |
| 1355 | Case15 | M101LH | TPt | V2c | 2 | 0 | 0 | 2,51225667223969E-06 | -5,599935992 |
| 1356 | Case15 | M101LH | V1c | V2c | 206680 | 167366 | 0,80978324 | 0,259616605 | -0,585667534 |
| 1357 | Case15 | M101LH | V1fpLF | V2c | 0 | 0 | NA | 0 | -Inf |
| 1358 | Case15 | M101LH | V1fpUF | V2c | 16 | 12 | 0,75 | 2,00980533779175E-05 | -4,696846005 |
| 1359 | Case15 | M101LH | V1pcLF | V2c | 0 | 0 | NA | 0 | -Inf |
| 1360 | Case15 | M101LH | V1pcUF | V2c | 0 | 0 | NA | 0 | -Inf |
| 1361 | Case15 | M101LH | V2c | V2c | 0 | 0 | NA | 0 | -Inf |

| | Case | MONKEY | SOURCE | TARGET | TOT | SUP | SLN | FLN | lgFLN |
|---|---|---|---|---|---|---|---|---|---|
| 1362 | Case15 | M101LH | V2fpLF | V2c | 0 | 0 | NA | 0 | -Inf |
| 1363 | Case15 | M101LH | V2fpUF | V2c | 4 | 2 | 0,5 | 5,02451334447938E-06 | -5,298905996 |
| 1364 | Case15 | M101LH | V2pcLF | V2c | 13 | 4 | 0,307692308 | 1,6329668369558E-05 | -4,787022635 |
| 1365 | Case15 | M101LH | V2pcUF | V2c | 0 | 0 | NA | 0 | -Inf |
| 1366 | Case15 | M101LH | V3A | V2c | 3 | 0 | 0 | 3,76838500835953E-06 | -5,423844733 |
| 1367 | Case15 | M101LH | V3c | V2c | 14951 | 5239 | 0,350411344 | 0,018780375 | -1,726295746 |
| 1368 | Case15 | M101LH | V3fpLF | V2c | 0 | 0 | NA | 0 | -Inf |
| 1369 | Case15 | M101LH | V3fpUF | V2c | 6 | 0 | 0 | 7,53677001671907E-06 | -5,122814737 |
| 1370 | Case15 | M101LH | V3pcLF | V2c | 82 | 4 | 0,048780488 | 0,000103003 | -3,987152135 |
| 1371 | Case15 | M101LH | V3pcUF | V2c | 6 | 2 | 0,333333333 | 7,53677001671907E-06 | -5,122814737 |
| 1372 | Case15 | M101LH | V4c | V2c | 52479 | 11445 | 0,218087235 | 0,065920359 | -1,180980436 |
| 1373 | Case15 | M101LH | V4LF | V2c | 4139 | 2466 | 0,595796086 | 0,005199115 | -2,284070561 |
| 1374 | Case15 | M101LH | V4t | V2c | 44 | 5 | 0,113636364 | 5,52696467892732E-05 | -4,257513311 |
| 1375 | Case15 | M101LH | V4UF | V2c | 246 | 10 | 0,040650407 | 0,000309008 | -3,51003088 |
| 1376 | Case15 | M101LH | V6 | V2c | 0 | 0 | NA | 0 | -Inf |
| 1377 | Case15 | M101LH | V6A | V2c | 15 | 9 | 0,6 | 1,88419250417977E-05 | -4,724874728 |
| 1378 | Case15 | M101LH | VIP | V2c | 2 | 0 | 0 | 2,51225667223969E-06 | -5,599935992 |
| 1379 | Case16 | M101RH | 1 | V2c | 0 | 0 | NA | 0 | -Inf |
| 1380 | Case16 | M101RH | 10 | V2c | 0 | 0 | NA | 0 | -Inf |
| 1381 | Case16 | M101RH | 11 | V2c | 0 | 0 | NA | 0 | -Inf |
| 1382 | Case16 | M101RH | 12 | V2c | 0 | 0 | NA | 0 | -Inf |
| 1383 | Case16 | M101RH | 13 | V2c | 0 | 0 | NA | 0 | -Inf |
| 1384 | Case16 | M101RH | 14 | V2c | 0 | 0 | NA | 0 | -Inf |
| 1385 | Case16 | M101RH | 2 | V2c | 0 | 0 | NA | 0 | -Inf |
| 1386 | Case16 | M101RH | 23 | V2c | 0 | 0 | NA | 0 | -Inf |
| 1387 | Case16 | M101RH | 24a | V2c | 0 | 0 | NA | 0 | -Inf |
| 1388 | Case16 | M101RH | 24b | V2c | 0 | 0 | NA | 0 | -Inf |
| 1389 | Case16 | M101RH | 24c | V2c | 0 | 0 | NA | 0 | -Inf |
| 1390 | Case16 | M101RH | 24d | V2c | 0 | 0 | NA | 0 | -Inf |
| 1391 | Case16 | M101RH | 25 | V2c | 0 | 0 | NA | 0 | -Inf |
| 1392 | Case16 | M101RH | 29/30 | V2c | 0 | 0 | NA | 0 | -Inf |
| 1393 | Case16 | M101RH | 3 | V2c | 0 | 0 | NA | 0 | -Inf |
| 1394 | Case16 | M101RH | 31 | V2c | 0 | 0 | NA | 0 | -Inf |
| 1395 | Case16 | M101RH | 32 | V2c | 0 | 0 | NA | 0 | -Inf |
| 1396 | Case16 | M101RH | 44 | V2c | 0 | 0 | NA | 0 | -Inf |
| 1397 | Case16 | M101RH | 45A | V2c | 0 | 0 | NA | 0 | -Inf |
| 1398 | Case16 | M101RH | 45B | V2c | 0 | 0 | NA | 0 | -Inf |
| 1399 | Case16 | M101RH | 46d | V2c | 0 | 0 | NA | 0 | -Inf |
| 1400 | Case16 | M101RH | 46v | V2c | 0 | 0 | NA | 0 | -Inf |
| 1401 | Case16 | M101RH | 5 | V2c | 0 | 0 | NA | 0 | -Inf |
| 1402 | Case16 | M101RH | 7A | V2c | 0 | 0 | NA | 0 | -Inf |
| 1403 | Case16 | M101RH | 7B | V2c | 0 | 0 | NA | 0 | -Inf |
| 1404 | Case16 | M101RH | 7m | V2c | 0 | 0 | NA | 0 | -Inf |
| 1405 | Case16 | M101RH | 7op | V2c | 0 | 0 | NA | 0 | -Inf |
| 1406 | Case16 | M101RH | 8B | V2c | 0 | 0 | NA | 0 | -Inf |
| 1407 | Case16 | M101RH | 8l | V2c | 28 | 12 | 0,428571429 | 3,51715934113557E-05 | -4,453807956 |
| 1408 | Case16 | M101RH | 8m | V2c | 14 | 0 | 0 | 1,75857967056778E-05 | -4,754837952 |
| 1409 | Case16 | M101RH | 8r | V2c | 0 | 0 | NA | 0 | -Inf |
| 1410 | Case16 | M101RH | 9 | V2c | 0 | 0 | NA | 0 | -Inf |
| 1411 | Case16 | M101RH | 9/46d | V2c | 0 | 0 | NA | 0 | -Inf |
| 1412 | Case16 | M101RH | 9/46v | V2c | 0 | 0 | NA | 0 | -Inf |
| 1413 | Case16 | M101RH | AIP | V2c | 0 | 0 | NA | 0 | -Inf |
| 1414 | Case16 | M101RH | CORE | V2c | 0 | 0 | NA | 0 | -Inf |
| 1415 | Case16 | M101RH | DP | V2c | 14 | 14 | 1 | 1,75857967056778E-05 | -4,754837952 |
| 1416 | Case16 | M101RH | ENTO | V2c | 0 | 0 | NA | 0 | -Inf |
| 1417 | Case16 | M101RH | F1 | V2c | 0 | 0 | NA | 0 | -Inf |
| 1418 | Case16 | M101RH | F2 | V2c | 0 | 0 | NA | 0 | -Inf |
| 1419 | Case16 | M101RH | F3 | V2c | 0 | 0 | NA | 0 | -Inf |
| 1420 | Case16 | M101RH | F4 | V2c | 0 | 0 | NA | 0 | -Inf |
| 1421 | Case16 | M101RH | F5 | V2c | 0 | 0 | NA | 0 | -Inf |
| 1422 | Case16 | M101RH | F6 | V2c | 0 | 0 | NA | 0 | -Inf |
| 1423 | Case16 | M101RH | F7 | V2c | 0 | 0 | NA | 0 | -Inf |
| 1424 | Case16 | M101RH | FST | V2c | 600 | 47 | 0,078333333 | 0,000753677 | -3,122814737 |
| 1425 | Case16 | M101RH | Gu | V2c | 0 | 0 | NA | 0 | -Inf |
| 1426 | Case16 | M101RH | INSULA | V2c | 0 | 0 | NA | 0 | -Inf |
| 1427 | Case16 | M101RH | IPa | V2c | 2 | 2 | 1 | 2,51225667223969E-06 | -5,599935992 |
| 1428 | Case16 | M101RH | LB | V2c | 0 | 0 | NA | 0 | -Inf |
| 1429 | Case16 | M101RH | LIP | V2c | 0 | 0 | NA | 0 | -Inf |
| 1430 | Case16 | M101RH | MB | V2c | 1 | 0 | 0 | 1,25612833611984E-06 | -5,900965987 |
| 1431 | Case16 | M101RH | MIP | V2c | 0 | 0 | NA | 0 | -Inf |
| 1432 | Case16 | M101RH | MST | V2c | 321 | 4 | 0,012461059 | 0,000403217 | -3,394460955 |
| 1433 | Case16 | M101RH | MTc | V2c | 8117 | 2656 | 0,327214488 | 0,010195994 | -1,991570441 |
| 1434 | Case16 | M101RH | MTp | V2c | 1662 | 54 | 0,032490975 | 0,002087685 | -2,680334968 |
| 1435 | Case16 | M101RH | OPAl | V2c | 0 | 0 | NA | 0 | -Inf |
| 1436 | Case16 | M101RH | OPro | V2c | 0 | 0 | NA | 0 | -Inf |

| | Case | MONKEY | SOURCE | TARGET | TOT | SUP | SLN | FLN | lgFLN |
|---|---|---|---|---|---|---|---|---|---|
| 1437 | Case16 | M101RH | PBc | V2c | 0 | 0 | NA | 0 | -Inf |
| 1438 | Case16 | M101RH | PBr | V2c | 0 | 0 | NA | 0 | -Inf |
| 1439 | Case16 | M101RH | PERI | V2c | 25 | 2 | 0,08 | 3,14032084029961E-05 | -4,503025979 |
| 1440 | Case16 | M101RH | PGa | V2c | 13 | 0 | 0 | 1,6329668369558E-05 | -4,787022635 |
| 1441 | Case16 | M101RH | Pi | V2c | 0 | 0 | NA | 0 | -Inf |
| 1442 | Case16 | M101RH | PIP | V2c | 252 | 2 | 0,007936508 | 0,000316544 | -3,499565447 |
| 1443 | Case16 | M101RH | Pir | V2c | 0 | 0 | NA | 0 | -Inf |
| 1444 | Case16 | M101RH | POLE | V2c | 0 | 0 | NA | 0 | -Inf |
| 1445 | Case16 | M101RH | Pro.St | V2c | 0 | 0 | NA | 0 | -Inf |
| 1446 | Case16 | M101RH | ProM | V2c | 0 | 0 | NA | 0 | -Inf |
| 1447 | Case16 | M101RH | SII | V2c | 0 | 0 | NA | 0 | -Inf |
| 1448 | Case16 | M101RH | STPc | V2c | 6 | 2 | 0,333333333 | 7,53677001671907E-06 | -5,122814737 |
| 1449 | Case16 | M101RH | STPi | V2c | 20 | 0 | 0 | 2,51225667223969E-05 | -4,599935992 |
| 1450 | Case16 | M101RH | STPr | V2c | 0 | 0 | NA | 0 | -Inf |
| 1451 | Case16 | M101RH | SUBI | V2c | 0 | 0 | NA | 0 | -Inf |
| 1452 | Case16 | M101RH | TEa/ma | V2c | 4 | 0 | 0 | 5,02451334447938E-06 | -5,298905996 |
| 1453 | Case16 | M101RH | TEa/mp | V2c | 38 | 0 | 0 | 4,77328767725541E-05 | -4,321182391 |
| 1454 | Case16 | M101RH | TEad | V2c | 7 | 0 | 0 | 8,79289835283891E-06 | -5,055867947 |
| 1455 | Case16 | M101RH | TEav | V2c | 10 | 0 | 0 | 1,25612833611984E-05 | -4,900965987 |
| 1456 | Case16 | M101RH | TEO | V2c | 1029 | 77 | 0,074829932 | 0,001292556 | -2,888550613 |
| 1457 | Case16 | M101RH | TEOm | V2c | 1013 | 100 | 0,098716683 | 0,001272458 | -2,895356542 |
| 1458 | Case16 | M101RH | TEpd | V2c | 149 | 6 | 0,040268456 | 0,000187163 | -3,727779719 |
| 1459 | Case16 | M101RH | TEpv | V2c | 392 | 1 | 0,00255102 | 0,000492402 | -3,30767992 |
| 1460 | Case16 | M101RH | TH_TF | V2c | 567 | 5 | 0,008818342 | 0,000712225 | -3,147382928 |
| 1461 | Case16 | M101RH | TPt | V2c | 0 | 0 | NA | 0 | -Inf |
| 1462 | Case16 | M101RH | V1c | V2c | 155780 | 121082 | 0,777262807 | 0,195679672 | -0,708454288 |
| 1463 | Case16 | M101RH | V1fpLF | V2c | 0 | 0 | NA | 0 | -Inf |
| 1464 | Case16 | M101RH | V1fpUF | V2c | 0 | 0 | NA | 0 | -Inf |
| 1465 | Case16 | M101RH | V1pcLF | V2c | 0 | 0 | NA | 0 | -Inf |
| 1466 | Case16 | M101RH | V1pcUF | V2c | 0 | 0 | NA | 0 | -Inf |
| 1467 | Case16 | M101RH | V2c | V2c | 0 | 0 | NA | 0 | -Inf |
| 1468 | Case16 | M101RH | V2fpLF | V2c | 7 | 7 | 1 | 8,79289835283891E-06 | -5,055867947 |
| 1469 | Case16 | M101RH | V2fpUF | V2c | 4 | 2 | 0,5 | 5,02451334447938E-06 | -5,298905996 |
| 1470 | Case16 | M101RH | V2pcLF | V2c | 45 | 12 | 0,266666667 | 5,6525775125393E-05 | -4,247753474 |
| 1471 | Case16 | M101RH | V2pcUF | V2c | 9 | 0 | 0 | 1,13051550250786E-05 | -4,946723478 |
| 1472 | Case16 | M101RH | V3A | V2c | 17 | 0 | 0 | 2,13541817140374E-05 | -4,670517066 |
| 1473 | Case16 | M101RH | V3c | V2c | 14950 | 7122 | 0,47638796 | 0,018779119 | -1,726324795 |
| 1474 | Case16 | M101RH | V3fpLF | V2c | 0 | 0 | NA | 0 | -Inf |
| 1475 | Case16 | M101RH | V3fpUF | V2c | 0 | 0 | NA | 0 | -Inf |
| 1476 | Case16 | M101RH | V3pcLF | V2c | 391 | 38 | 0,097186701 | 0,000491146 | -3,30878923 |
| 1477 | Case16 | M101RH | V3pcUF | V2c | 4 | 2 | 0,5 | 5,02451334447938E-06 | -5,298905996 |
| 1478 | Case16 | M101RH | V4c | V2c | 26720 | 4164 | 0,155838323 | 0,033563749 | -1,474129534 |
| 1479 | Case16 | M101RH | V4LF | V2c | 913 | 39 | 0,04271632 | 0,001146845 | -2,94049521 |
| 1480 | Case16 | M101RH | V4t | V2c | 21 | 6 | 0,285714286 | 2,63786950585167E-05 | -4,578746693 |
| 1481 | Case16 | M101RH | V4UF | V2c | 156 | 0 | 0 | 0,000195956 | -3,707841389 |
| 1482 | Case16 | M101RH | V6 | V2c | 0 | 0 | NA | 0 | -Inf |
| 1483 | Case16 | M101RH | V6A | V2c | 8 | 0 | 0 | 1,00490266889588E-05 | -4,997876 |
| 1484 | Case16 | M101RH | VIP | V2c | 0 | 0 | NA | 0 | -Inf |
| 1485 | Case17 | M103LH | 1 | V2c | 0 | 0 | NA | 0 | -Inf |
| 1486 | Case17 | M103LH | 10 | V2c | 0 | 0 | NA | 0 | -Inf |
| 1487 | Case17 | M103LH | 11 | V2c | 0 | 0 | NA | 0 | -Inf |
| 1488 | Case17 | M103LH | 12 | V2c | 0 | 0 | NA | 0 | -Inf |
| 1489 | Case17 | M103LH | 13 | V2c | 0 | 0 | NA | 0 | -Inf |
| 1490 | Case17 | M103LH | 14 | V2c | 0 | 0 | NA | 0 | -Inf |
| 1491 | Case17 | M103LH | 2 | V2c | 0 | 0 | NA | 0 | -Inf |
| 1492 | Case17 | M103LH | 23 | V2c | 0 | 0 | NA | 0 | -Inf |
| 1493 | Case17 | M103LH | 24a | V2c | 0 | 0 | NA | 0 | -Inf |
| 1494 | Case17 | M103LH | 24b | V2c | 0 | 0 | NA | 0 | -Inf |
| 1495 | Case17 | M103LH | 24c | V2c | 0 | 0 | NA | 0 | -Inf |
| 1496 | Case17 | M103LH | 24d | V2c | 0 | 0 | NA | 0 | -Inf |
| 1497 | Case17 | M103LH | 25 | V2c | 0 | 0 | NA | 0 | -Inf |
| 1498 | Case17 | M103LH | 29/30 | V2c | 0 | 0 | NA | 0 | -Inf |
| 1499 | Case17 | M103LH | 3 | V2c | 0 | 0 | NA | 0 | -Inf |
| 1500 | Case17 | M103LH | 31 | V2c | 0 | 0 | NA | 0 | -Inf |
| 1501 | Case17 | M103LH | 32 | V2c | 0 | 0 | NA | 0 | -Inf |
| 1502 | Case17 | M103LH | 44 | V2c | 0 | 0 | NA | 0 | -Inf |
| 1503 | Case17 | M103LH | 45A | V2c | 0 | 0 | NA | 0 | -Inf |
| 1504 | Case17 | M103LH | 45B | V2c | 0 | 0 | NA | 0 | -Inf |
| 1505 | Case17 | M103LH | 46d | V2c | 0 | 0 | NA | 0 | -Inf |
| 1506 | Case17 | M103LH | 46v | V2c | 0 | 0 | NA | 0 | -Inf |
| 1507 | Case17 | M103LH | 5 | V2c | 0 | 0 | NA | 0 | -Inf |
| 1508 | Case17 | M103LH | 7A | V2c | 0 | 0 | NA | 0 | -Inf |
| 1509 | Case17 | M103LH | 7B | V2c | 0 | 0 | NA | 0 | -Inf |
| 1510 | Case17 | M103LH | 7m | V2c | 0 | 0 | NA | 0 | -Inf |
| 1511 | Case17 | M103LH | 7op | V2c | 0 | 0 | NA | 0 | -Inf |

| | Case | MONKEY | SOURCE | TARGET | TOT | SUP | SLN | FLN | lgFLN |
|---|---|---|---|---|---|---|---|---|---|
| 1512 | Case17 | M103LH | 8B | V2c | 0 | 0 | NA | 0 | -Inf |
| 1513 | Case17 | M103LH | 8l | V2c | 23 | 4 | 0,173913043 | 2,88909517307564E-05 | -4,539238151 |
| 1514 | Case17 | M103LH | 8m | V2c | 0 | 0 | NA | 0 | -Inf |
| 1515 | Case17 | M103LH | 8r | V2c | 0 | 0 | NA | 0 | -Inf |
| 1516 | Case17 | M103LH | 9 | V2c | 0 | 0 | NA | 0 | -Inf |
| 1517 | Case17 | M103LH | 9/46d | V2c | 0 | 0 | NA | 0 | -Inf |
| 1518 | Case17 | M103LH | 9/46v | V2c | 0 | 0 | NA | 0 | -Inf |
| 1519 | Case17 | M103LH | AIP | V2c | 0 | 0 | NA | 0 | -Inf |
| 1520 | Case17 | M103LH | CORE | V2c | 0 | 0 | NA | 0 | -Inf |
| 1521 | Case17 | M103LH | DP | V2c | 7 | 1 | 0,142857143 | 8,79289835283891E-06 | -5,055867947 |
| 1522 | Case17 | M103LH | ENTO | V2c | 0 | 0 | NA | 0 | -Inf |
| 1523 | Case17 | M103LH | F1 | V2c | 0 | 0 | NA | 0 | -Inf |
| 1524 | Case17 | M103LH | F2 | V2c | 0 | 0 | NA | 0 | -Inf |
| 1525 | Case17 | M103LH | F3 | V2c | 0 | 0 | NA | 0 | -Inf |
| 1526 | Case17 | M103LH | F4 | V2c | 0 | 0 | NA | 0 | -Inf |
| 1527 | Case17 | M103LH | F5 | V2c | 0 | 0 | NA | 0 | -Inf |
| 1528 | Case17 | M103LH | F6 | V2c | 0 | 0 | NA | 0 | -Inf |
| 1529 | Case17 | M103LH | F7 | V2c | 0 | 0 | NA | 0 | -Inf |
| 1530 | Case17 | M103LH | FST | V2c | 220 | 19 | 0,086363636 | 0,000276348 | -3,558543307 |
| 1531 | Case17 | M103LH | Gu | V2c | 0 | 0 | NA | 0 | -Inf |
| 1532 | Case17 | M103LH | INSULA | V2c | 0 | 0 | NA | 0 | -Inf |
| 1533 | Case17 | M103LH | IPa | V2c | 20 | 0 | 0 | 2,51225667223969E-05 | -4,599935992 |
| 1534 | Case17 | M103LH | LB | V2c | 0 | 0 | NA | 0 | -Inf |
| 1535 | Case17 | M103LH | LIP | V2c | 93 | 7 | 0,075268817 | 0,00011682 | -3,932483039 |
| 1536 | Case17 | M103LH | MB | V2c | 3 | 0 | 0 | 3,76838500835953E-06 | -5,423844733 |
| 1537 | Case17 | M103LH | MIP | V2c | 0 | 0 | NA | 0 | -Inf |
| 1538 | Case17 | M103LH | MST | V2c | 44 | 1 | 0,022727273 | 5,52696467892732E-05 | -4,257513311 |
| 1539 | Case17 | M103LH | MTc | V2c | 4242 | 1167 | 0,275106082 | 0,005328496 | -2,273395323 |
| 1540 | Case17 | M103LH | MTp | V2c | 929 | 42 | 0,045209903 | 0,001166943 | -2,932950273 |
| 1541 | Case17 | M103LH | OPAl | V2c | 0 | 0 | NA | 0 | -Inf |
| 1542 | Case17 | M103LH | OPro | V2c | 0 | 0 | NA | 0 | -Inf |
| 1543 | Case17 | M103LH | PBc | V2c | 1 | 0 | 0 | 1,25612833611984E-06 | -5,900965987 |
| 1544 | Case17 | M103LH | PBr | V2c | 0 | 0 | NA | 0 | -Inf |
| 1545 | Case17 | M103LH | PERI | V2c | 80 | 2 | 0,025 | 0,00010049 | -3,997876 |
| 1546 | Case17 | M103LH | PGa | V2c | 58 | 1 | 0,017241379 | 7,2855443494951E-05 | -4,137537994 |
| 1547 | Case17 | M103LH | Pi | V2c | 0 | 0 | NA | 0 | -Inf |
| 1548 | Case17 | M103LH | PIP | V2c | 75 | 0 | 0 | 9,42096252089883E-05 | -4,025904724 |
| 1549 | Case17 | M103LH | Pir | V2c | 0 | 0 | NA | 0 | -Inf |
| 1550 | Case17 | M103LH | POLE | V2c | 0 | 0 | NA | 0 | -Inf |
| 1551 | Case17 | M103LH | Pro.St | V2c | 0 | 0 | NA | 0 | -Inf |
| 1552 | Case17 | M103LH | ProM | V2c | 0 | 0 | NA | 0 | -Inf |
| 1553 | Case17 | M103LH | SII | V2c | 0 | 0 | NA | 0 | -Inf |
| 1554 | Case17 | M103LH | STPc | V2c | 16 | 0 | 0 | 2,00980533779175E-05 | -4,696846005 |
| 1555 | Case17 | M103LH | STPi | V2c | 19 | 0 | 0 | 2,3866438386277E-05 | -4,622212386 |
| 1556 | Case17 | M103LH | STPr | V2c | 5 | 2 | 0,4 | 6,28064168059922E-06 | -5,201995983 |
| 1557 | Case17 | M103LH | SUBI | V2c | 0 | 0 | NA | 0 | -Inf |
| 1558 | Case17 | M103LH | TEa/ma | V2c | 29 | 0 | 0 | 3,64277217474755E-05 | -4,438567989 |
| 1559 | Case17 | M103LH | TEa/mp | V2c | 19 | 3 | 0,157894737 | 2,3866438386277E-05 | -4,622212386 |
| 1560 | Case17 | M103LH | TEad | V2c | 80 | 4 | 0,05 | 0,00010049 | -3,997876 |
| 1561 | Case17 | M103LH | TEav | V2c | 117 | 5 | 0,042735043 | 0,000146967 | -3,832780126 |
| 1562 | Case17 | M103LH | TEO | V2c | 910 | 218 | 0,23956044 | 0,001143077 | -2,941924595 |
| 1563 | Case17 | M103LH | TEOm | V2c | 1367 | 333 | 0,243599122 | 0,001717127 | -2,765197473 |
| 1564 | Case17 | M103LH | TEpd | V2c | 329 | 14 | 0,042553191 | 0,000413266 | -3,383770089 |
| 1565 | Case17 | M103LH | TEpv | V2c | 253 | 6 | 0,023715415 | 0,0003178 | -3,497845466 |
| 1566 | Case17 | M103LH | TH_TF | V2c | 230 | 3 | 0,013043478 | 0,00028891 | -3,539238151 |
| 1567 | Case17 | M103LH | TPt | V2c | 0 | 0 | NA | 0 | -Inf |
| 1568 | Case17 | M103LH | V1c | V2c | 148105 | 109014 | 0,736058877 | 0,186038887 | -0,730396267 |
| 1569 | Case17 | M103LH | V1fpLF | V2c | 0 | 0 | NA | 0 | -Inf |
| 1570 | Case17 | M103LH | V1fpUF | V2c | 0 | 0 | NA | 0 | -Inf |
| 1571 | Case17 | M103LH | V1pcLF | V2c | 0 | 0 | NA | 0 | -Inf |
| 1572 | Case17 | M103LH | V1pcUF | V2c | 0 | 0 | NA | 0 | -Inf |
| 1573 | Case17 | M103LH | V2c | V2c | 0 | 0 | NA | 0 | -Inf |
| 1574 | Case17 | M103LH | V2fpLF | V2c | 0 | 0 | NA | 0 | -Inf |
| 1575 | Case17 | M103LH | V2fpUF | V2c | 0 | 0 | NA | 0 | -Inf |
| 1576 | Case17 | M103LH | V2pcLF | V2c | 0 | 0 | NA | 0 | -Inf |
| 1577 | Case17 | M103LH | V2pcUF | V2c | 0 | 0 | NA | 0 | -Inf |
| 1578 | Case17 | M103LH | V3A | V2c | 1 | 0 | 0 | 1,25612833611984E-06 | -5,900965987 |
| 1579 | Case17 | M103LH | V3c | V2c | 2982 | 1034 | 0,34674715 | 0,003745775 | -2,426458348 |
| 1580 | Case17 | M103LH | V3fpLF | V2c | 0 | 0 | NA | 0 | -Inf |
| 1581 | Case17 | M103LH | V3fpUF | V2c | 0 | 0 | NA | 0 | -Inf |
| 1582 | Case17 | M103LH | V3pcLF | V2c | 124 | 5 | 0,040322581 | 0,00015576 | -3,807544302 |
| 1583 | Case17 | M103LH | V3pcUF | V2c | 3 | 0 | 0 | 3,76838500835953E-06 | -5,423844733 |
| 1584 | Case17 | M103LH | V4c | V2c | 28758 | 9162 | 0,31858961 | 0,036123739 | -1,442207308 |
| 1585 | Case17 | M103LH | V4LF | V2c | 1424 | 1118 | 0,78511236 | 0,001788727 | -2,747455998 |
| 1586 | Case17 | M103LH | V4t | V2c | 20 | 2 | 0,1 | 2,51225667223969E-05 | -4,599935992 |

| | Case | MONKEY | SOURCE | TARGET | TOT | SUP | SLN | FLN | lgFLN |
|---|---|---|---|---|---|---|---|---|---|
| 1587 | Case17 | M103LH | V4UF | V2c | 31 | 12 | 0,387096774 | 3,89399784197152E-05 | -4,409604293 |
| 1588 | Case17 | M103LH | V6 | V2c | 5 | 0 | 0 | 6,28064168059922E-06 | -5,201995983 |
| 1589 | Case17 | M103LH | V6A | V2c | 6 | 2 | 0,333333333 | 7,53677001671907E-06 | -5,122814737 |
| 1590 | Case17 | M103LH | VIP | V2c | 0 | 0 | NA | 0 | -Inf |
| 1697 | Case18 | M146LH | 1 | V2c | 0 | 0 | NA | 0 | -Inf |
| 1698 | Case18 | M146LH | 10 | V2c | 0 | 0 | NA | 0 | -Inf |
| 1699 | Case18 | M146LH | 11 | V2c | 0 | 0 | NA | 0 | -Inf |
| 1700 | Case18 | M146LH | 12 | V2c | 0 | 0 | NA | 0 | -Inf |
| 1701 | Case18 | M146LH | 13 | V2c | 0 | 0 | NA | 0 | -Inf |
| 1702 | Case18 | M146LH | 14 | V2c | 0 | 0 | NA | 0 | -Inf |
| 1703 | Case18 | M146LH | 2 | V2c | 0 | 0 | NA | 0 | -Inf |
| 1704 | Case18 | M146LH | 23 | V2c | 0 | 0 | NA | 0 | -Inf |
| 1705 | Case18 | M146LH | 24a | V2c | 2 | 0 | 0 | 2,51225667223969E-06 | -5,599935992 |
| 1706 | Case18 | M146LH | 24b | V2c | 2 | 0 | 0 | 2,51225667223969E-06 | -5,599935992 |
| 1707 | Case18 | M146LH | 24c | V2c | 0 | 0 | NA | 0 | -Inf |
| 1708 | Case18 | M146LH | 24d | V2c | 0 | 0 | NA | 0 | -Inf |
| 1709 | Case18 | M146LH | 25 | V2c | 0 | 0 | NA | 0 | -Inf |
| 1710 | Case18 | M146LH | 29/30 | V2c | 0 | 0 | NA | 0 | -Inf |
| 1711 | Case18 | M146LH | 3 | V2c | 0 | 0 | NA | 0 | -Inf |
| 1712 | Case18 | M146LH | 31 | V2c | 0 | 0 | NA | 0 | -Inf |
| 1713 | Case18 | M146LH | 32 | V2c | 0 | 0 | NA | 0 | -Inf |
| 1714 | Case18 | M146LH | 44 | V2c | 0 | 0 | NA | 0 | -Inf |
| 1715 | Case18 | M146LH | 45A | V2c | 0 | 0 | NA | 0 | -Inf |
| 1716 | Case18 | M146LH | 45B | V2c | 2 | 2 | 1 | 2,51225667223969E-06 | -5,599935992 |
| 1717 | Case18 | M146LH | 46d | V2c | 0 | 0 | NA | 0 | -Inf |
| 1718 | Case18 | M146LH | 46v | V2c | 0 | 0 | NA | 0 | -Inf |
| 1719 | Case18 | M146LH | 5 | V2c | 0 | 0 | NA | 0 | -Inf |
| 1720 | Case18 | M146LH | 7A | V2c | 0 | 0 | NA | 0 | -Inf |
| 1721 | Case18 | M146LH | 7B | V2c | 0 | 0 | NA | 0 | -Inf |
| 1722 | Case18 | M146LH | 7m | V2c | 0 | 0 | NA | 0 | -Inf |
| 1723 | Case18 | M146LH | 7op | V2c | 0 | 0 | NA | 0 | -Inf |
| 1724 | Case18 | M146LH | 8B | V2c | 0 | 0 | NA | 0 | -Inf |
| 1725 | Case18 | M146LH | 8l | V2c | 0 | 0 | NA | 0 | -Inf |
| 1726 | Case18 | M146LH | 8m | V2c | 0 | 0 | NA | 0 | -Inf |
| 1727 | Case18 | M146LH | 8r | V2c | 0 | 0 | NA | 0 | -Inf |
| 1728 | Case18 | M146LH | 9 | V2c | 0 | 0 | NA | 0 | -Inf |
| 1729 | Case18 | M146LH | 9/46d | V2c | 0 | 0 | NA | 0 | -Inf |
| 1730 | Case18 | M146LH | 9/46v | V2c | 0 | 0 | NA | 0 | -Inf |
| 1731 | Case18 | M146LH | AIP | V2c | 0 | 0 | NA | 0 | -Inf |
| 1732 | Case18 | M146LH | CORE | V2c | 0 | 0 | NA | 0 | -Inf |
| 1733 | Case18 | M146LH | DP | V2c | 4 | 0 | 0 | 5,02451334447938E-06 | -5,298905996 |
| 1734 | Case18 | M146LH | ENTO | V2c | 0 | 0 | NA | 0 | -Inf |
| 1735 | Case18 | M146LH | F1 | V2c | 0 | 0 | NA | 0 | -Inf |
| 1736 | Case18 | M146LH | F2 | V2c | 0 | 0 | NA | 0 | -Inf |
| 1737 | Case18 | M146LH | F3 | V2c | 0 | 0 | NA | 0 | -Inf |
| 1738 | Case18 | M146LH | F4 | V2c | 0 | 0 | NA | 0 | -Inf |
| 1739 | Case18 | M146LH | F5 | V2c | 0 | 0 | NA | 0 | -Inf |
| 1740 | Case18 | M146LH | F6 | V2c | 0 | 0 | NA | 0 | -Inf |
| 1741 | Case18 | M146LH | F7 | V2c | 0 | 0 | NA | 0 | -Inf |
| 1742 | Case18 | M146LH | FST | V2c | 74 | 4 | 0,054054054 | 9,29534968728685E-05 | -4,031734268 |
| 1743 | Case18 | M146LH | Gu | V2c | 0 | 0 | NA | 0 | -Inf |
| 1744 | Case18 | M146LH | INSULA | V2c | 0 | 0 | NA | 0 | -Inf |
| 1745 | Case18 | M146LH | IPa | V2c | 108 | 2 | 0,018518519 | 0,000135662 | -3,867542232 |
| 1746 | Case18 | M146LH | LB | V2c | 0 | 0 | NA | 0 | -Inf |
| 1747 | Case18 | M146LH | LIP | V2c | 18 | 0 | 0 | 2,26103100501572E-05 | -4,645693482 |
| 1748 | Case18 | M146LH | MB | V2c | 0 | 0 | NA | 0 | -Inf |
| 1749 | Case18 | M146LH | MIP | V2c | 0 | 0 | NA | 0 | -Inf |
| 1750 | Case18 | M146LH | MST | V2c | 12 | 0 | 0 | 1,50735400334381E-05 | -4,821784741 |
| 1751 | Case18 | M146LH | MTc | V2c | 5252 | 1250 | 0,23800457 | 0,006597186 | -2,18064127 |
| 1752 | Case18 | M146LH | MTp | V2c | 192 | 12 | 0,0625 | 0,000241177 | -3,617664759 |
| 1753 | Case18 | M146LH | OPAl | V2c | 0 | 0 | NA | 0 | -Inf |
| 1754 | Case18 | M146LH | OPro | V2c | 0 | 0 | NA | 0 | -Inf |
| 1755 | Case18 | M146LH | PBc | V2c | 2 | 0 | 0 | 2,51225667223969E-06 | -5,599935992 |
| 1756 | Case18 | M146LH | PBr | V2c | 0 | 0 | NA | 0 | -Inf |
| 1757 | Case18 | M146LH | PERI | V2c | 26 | 0 | 0 | 3,2659336739116E-05 | -4,485992639 |
| 1758 | Case18 | M146LH | PGa | V2c | 37 | 2 | 0,054054054 | 4,64767484364343E-05 | -4,332764263 |
| 1759 | Case18 | M146LH | Pi | V2c | 0 | 0 | NA | 0 | -Inf |
| 1760 | Case18 | M146LH | PIP | V2c | 82 | 0 | 0 | 0,000103003 | -3,987152135 |
| 1761 | Case18 | M146LH | Pir | V2c | 0 | 0 | NA | 0 | -Inf |
| 1762 | Case18 | M146LH | POLE | V2c | 0 | 0 | NA | 0 | -Inf |
| 1763 | Case18 | M146LH | Pro.St | V2c | 0 | 0 | NA | 0 | -Inf |
| 1764 | Case18 | M146LH | ProM | V2c | 0 | 0 | NA | 0 | -Inf |
| 1765 | Case18 | M146LH | SII | V2c | 0 | 0 | NA | 0 | -Inf |
| 1766 | Case18 | M146LH | STPc | V2c | 6 | 0 | 0 | 7,53677001671907E-06 | -5,122814737 |
| 1767 | Case18 | M146LH | STPi | V2c | 8 | 2 | 0,25 | 1,00490266889588E-05 | -4,997876 |

| | Case | MONKEY | SOURCE | TARGET | TOT | SUP | SLN | FLN | lgFLN |
|---|---|---|---|---|---|---|---|---|---|
| 1768 | Case18 | M146LH | STPr | V2c | 0 | 0 | NA | 0 | -Inf |
| 1769 | Case18 | M146LH | SUBI | V2c | 0 | 0 | NA | 0 | -Inf |
| 1770 | Case18 | M146LH | TEa/ma | V2c | 30 | 0 | 0 | 3,76838500835953E-05 | -4,423844733 |
| 1771 | Case18 | M146LH | TEa/mp | V2c | 145 | 0 | 0 | 0,000182139 | -3,739597985 |
| 1772 | Case18 | M146LH | TEad | V2c | 32 | 4 | 0,125 | 4,0196106755835E-05 | -4,395816009 |
| 1773 | Case18 | M146LH | TEav | V2c | 14 | 4 | 0,285714286 | 1,75857967056778E-05 | -4,754837952 |
| 1774 | Case18 | M146LH | TEO | V2c | 744 | 156 | 0,209677419 | 0,000934559 | -3,029393052 |
| 1775 | Case18 | M146LH | TEOm | V2c | 978 | 70 | 0,071574642 | 0,001228494 | -2,910627133 |
| 1776 | Case18 | M146LH | TEpd | V2c | 247 | 115 | 0,465587045 | 0,000310264 | -3,508269034 |
| 1777 | Case18 | M146LH | TEpv | V2c | 180 | 22 | 0,122222222 | 0,000226103 | -3,645693482 |
| 1778 | Case18 | M146LH | TH_TF | V2c | 132 | 0 | 0 | 0,000165809 | -3,780392056 |
| 1779 | Case18 | M146LH | TPt | V2c | 0 | 0 | NA | 0 | -Inf |
| 1780 | Case18 | M146LH | V1c | V2c | 48808 | 42334 | 0,86735781 | 0,061309112 | -1,212474975 |
| 1781 | Case18 | M146LH | V1fpLF | V2c | 0 | 0 | NA | 0 | -Inf |
| 1782 | Case18 | M146LH | V1fpUF | V2c | 0 | 0 | NA | 0 | -Inf |
| 1783 | Case18 | M146LH | V1pcLF | V2c | 0 | 0 | NA | 0 | -Inf |
| 1784 | Case18 | M146LH | V1pcUF | V2c | 0 | 0 | NA | 0 | -Inf |
| 1785 | Case18 | M146LH | V2c | V2c | 0 | 0 | NA | 0 | -Inf |
| 1786 | Case18 | M146LH | V2fpLF | V2c | 0 | 0 | NA | 0 | -Inf |
| 1787 | Case18 | M146LH | V2fpUF | V2c | 0 | 0 | NA | 0 | -Inf |
| 1788 | Case18 | M146LH | V2pcLF | V2c | 0 | 0 | NA | 0 | -Inf |
| 1789 | Case18 | M146LH | V2pcUF | V2c | 2 | 0 | 0 | 2,51225667223969E-06 | -5,599935992 |
| 1790 | Case18 | M146LH | V3A | V2c | 22 | 2 | 0,090909091 | 2,76348233946366E-05 | -4,558543307 |
| 1791 | Case18 | M146LH | V3c | V2c | 7710 | 1776 | 0,230350195 | 0,009684749 | -2,013911609 |
| 1792 | Case18 | M146LH | V3fpLF | V2c | 0 | 0 | NA | 0 | -Inf |
| 1793 | Case18 | M146LH | V3fpUF | V2c | 0 | 0 | NA | 0 | -Inf |
| 1794 | Case18 | M146LH | V3pcLF | V2c | 122 | 4 | 0,032786885 | 0,000153248 | -3,814606157 |
| 1795 | Case18 | M146LH | V3pcUF | V2c | 4 | 2 | 0,5 | 5,02451334447938E-06 | -5,298905996 |
| 1796 | Case18 | M146LH | V4c | V2c | 26122 | 4264 | 0,163234056 | 0,032812584 | -1,483959562 |
| 1797 | Case18 | M146LH | V4LF | V2c | 2750 | 1794 | 0,652363636 | 0,003454353 | -2,461633294 |
| 1798 | Case18 | M146LH | V4t | V2c | 8 | 2 | 0,25 | 1,00490266889588E-05 | -4,997876 |
| 1799 | Case18 | M146LH | V4UF | V2c | 14 | 2 | 0,142857143 | 1,75857967056778E-05 | -4,754837952 |
| 1800 | Case18 | M146LH | V6 | V2c | 0 | 0 | NA | 0 | -Inf |
| 1801 | Case18 | M146LH | V6A | V2c | 0 | 0 | NA | 0 | -Inf |
| 1802 | Case18 | M146LH | VIP | V2c | 0 | 0 | NA | 0 | -Inf |
| 2121 | Case19 | M148RH | 1 | V2pcUF | 0 | 0 | NA | 0 | -Inf |
| 2122 | Case19 | M148RH | 10 | V2pcUF | 0 | 0 | NA | 0 | -Inf |
| 2123 | Case19 | M148RH | 11 | V2pcUF | 0 | 0 | NA | 0 | -Inf |
| 2124 | Case19 | M148RH | 12 | V2pcUF | 0 | 0 | NA | 0 | -Inf |
| 2125 | Case19 | M148RH | 13 | V2pcUF | 0 | 0 | NA | 0 | -Inf |
| 2126 | Case19 | M148RH | 14 | V2pcUF | 0 | 0 | NA | 0 | -Inf |
| 2127 | Case19 | M148RH | 2 | V2pcUF | 0 | 0 | NA | 0 | -Inf |
| 2128 | Case19 | M148RH | 23 | V2pcUF | 0 | 0 | NA | 0 | -Inf |
| 2129 | Case19 | M148RH | 24a | V2pcUF | 0 | 0 | NA | 0 | -Inf |
| 2130 | Case19 | M148RH | 24b | V2pcUF | 0 | 0 | NA | 0 | -Inf |
| 2131 | Case19 | M148RH | 24c | V2pcUF | 0 | 0 | NA | 0 | -Inf |
| 2132 | Case19 | M148RH | 24d | V2pcUF | 0 | 0 | NA | 0 | -Inf |
| 2133 | Case19 | M148RH | 25 | V2pcUF | 0 | 0 | NA | 0 | -Inf |
| 2134 | Case19 | M148RH | 29/30 | V2pcUF | 0 | 0 | NA | 0 | -Inf |
| 2135 | Case19 | M148RH | 3 | V2pcUF | 0 | 0 | NA | 0 | -Inf |
| 2136 | Case19 | M148RH | 31 | V2pcUF | 0 | 0 | NA | 0 | -Inf |
| 2137 | Case19 | M148RH | 32 | V2pcUF | 0 | 0 | NA | 0 | -Inf |
| 2138 | Case19 | M148RH | 44 | V2pcUF | 0 | 0 | NA | 0 | -Inf |
| 2139 | Case19 | M148RH | 45A | V2pcUF | 0 | 0 | NA | 0 | -Inf |
| 2140 | Case19 | M148RH | 45B | V2pcUF | 2 | 0 | 0 | 8,60707415424738E-06 | -5,065144455 |
| 2141 | Case19 | M148RH | 46d | V2pcUF | 0 | 0 | NA | 0 | -Inf |
| 2142 | Case19 | M148RH | 46v | V2pcUF | 0 | 0 | NA | 0 | -Inf |
| 2143 | Case19 | M148RH | 5 | V2pcUF | 2 | 0 | 0 | 8,60707415424738E-06 | -5,065144455 |
| 2144 | Case19 | M148RH | 7A | V2pcUF | 8 | 0 | 0 | 3,44282966169895E-05 | -4,463084464 |
| 2145 | Case19 | M148RH | 7B | V2pcUF | 4 | 2 | 0,5 | 1,72141483084948E-05 | -4,76411446 |
| 2146 | Case19 | M148RH | 7m | V2pcUF | 0 | 0 | NA | 0 | -Inf |
| 2147 | Case19 | M148RH | 7op | V2pcUF | 0 | 0 | NA | 0 | -Inf |
| 2148 | Case19 | M148RH | 8B | V2pcUF | 0 | 0 | NA | 0 | -Inf |
| 2149 | Case19 | M148RH | 8l | V2pcUF | 56 | 28 | 0,5 | 0,000240998 | -3,617986424 |
| 2150 | Case19 | M148RH | 8m | V2pcUF | 0 | 0 | NA | 0 | -Inf |
| 2151 | Case19 | M148RH | 8r | V2pcUF | 0 | 0 | NA | 0 | -Inf |
| 2152 | Case19 | M148RH | 9 | V2pcUF | 2 | 0 | 0 | 8,60707415424738E-06 | -5,065144455 |
| 2153 | Case19 | M148RH | 9/46d | V2pcUF | 0 | 0 | NA | 0 | -Inf |
| 2154 | Case19 | M148RH | 9/46v | V2pcUF | 0 | 0 | NA | 0 | -Inf |
| 2155 | Case19 | M148RH | AIP | V2pcUF | 0 | 0 | NA | 0 | -Inf |
| 2156 | Case19 | M148RH | CORE | V2pcUF | 0 | 0 | NA | 0 | -Inf |
| 2157 | Case19 | M148RH | DP | V2pcUF | 6 | 0 | 0 | 2,58212224627421E-05 | -4,588023201 |
| 2158 | Case19 | M148RH | ENTO | V2pcUF | 6 | 0 | 0 | 2,58212224627421E-05 | -4,588023201 |
| 2159 | Case19 | M148RH | F1 | V2pcUF | 0 | 0 | NA | 0 | -Inf |
| 2160 | Case19 | M148RH | F2 | V2pcUF | 0 | 0 | NA | 0 | -Inf |

| | Case | MONKEY | SOURCE | TARGET | TOT | SUP | SLN | FLN | lgFLN |
|---|---|---|---|---|---|---|---|---|---|
| 2161 | Case19 | M148RH | F3 | V2pcUF | 0 | 0 | NA | 0 | -Inf |
| 2162 | Case19 | M148RH | F4 | V2pcUF | 0 | 0 | NA | 0 | -Inf |
| 2163 | Case19 | M148RH | F5 | V2pcUF | 2 | 0 | 0 | 8,60707415424738E-06 | -5,065144455 |
| 2164 | Case19 | M148RH | F6 | V2pcUF | 0 | 0 | NA | 0 | -Inf |
| 2165 | Case19 | M148RH | F7 | V2pcUF | 0 | 0 | NA | 0 | -Inf |
| 2166 | Case19 | M148RH | FST | V2pcUF | 218 | 0 | 0 | 0,000938171 | -3,027717957 |
| 2167 | Case19 | M148RH | Gu | V2pcUF | 0 | 0 | NA | 0 | -Inf |
| 2168 | Case19 | M148RH | INSULA | V2pcUF | 0 | 0 | NA | 0 | -Inf |
| 2169 | Case19 | M148RH | IPa | V2pcUF | 390 | 8 | 0,020512821 | 0,001678379 | -2,775109844 |
| 2170 | Case19 | M148RH | LB | V2pcUF | 0 | 0 | NA | 0 | -Inf |
| 2171 | Case19 | M148RH | LIP | V2pcUF | 582 | 76 | 0,130584192 | 0,002504659 | -2,601251466 |
| 2172 | Case19 | M148RH | MB | V2pcUF | 4 | 0 | 0 | 1,72141483084948E-05 | -4,76411446 |
| 2173 | Case19 | M148RH | MIP | V2pcUF | 0 | 0 | NA | 0 | -Inf |
| 2174 | Case19 | M148RH | MST | V2pcUF | 374 | 12 | 0,032085561 | 0,001609523 | -2,793302849 |
| 2175 | Case19 | M148RH | MTc | V2pcUF | 2242 | 340 | 0,151650312 | 0,00964853 | -2,015538843 |
| 2176 | Case19 | M148RH | MTp | V2pcUF | 3884 | 860 | 0,221421215 | 0,016714938 | -1,77689523 |
| 2177 | Case19 | M148RH | OPAl | V2pcUF | 0 | 0 | NA | 0 | -Inf |
| 2178 | Case19 | M148RH | OPro | V2pcUF | 0 | 0 | NA | 0 | -Inf |
| 2179 | Case19 | M148RH | PBc | V2pcUF | 16 | 0 | 0 | 6,8856593233979E-05 | -4,162054468 |
| 2180 | Case19 | M148RH | PBr | V2pcUF | 0 | 0 | NA | 0 | -Inf |
| 2181 | Case19 | M148RH | PERI | V2pcUF | 182 | 2 | 0,010989011 | 0,000783244 | -3,106103063 |
| 2182 | Case19 | M148RH | PGa | V2pcUF | 146 | 2 | 0,01369863 | 0,000628316 | -3,201821595 |
| 2183 | Case19 | M148RH | Pi | V2pcUF | 0 | 0 | NA | 0 | -Inf |
| 2184 | Case19 | M148RH | PIP | V2pcUF | 2276 | 400 | 0,175746924 | 0,00979485 | -2,009002193 |
| 2185 | Case19 | M148RH | Pir | V2pcUF | 0 | 0 | NA | 0 | -Inf |
| 2186 | Case19 | M148RH | POLE | V2pcUF | 2 | 0 | 0 | 8,60707415424738E-06 | -5,065144455 |
| 2187 | Case19 | M148RH | Pro.St | V2pcUF | 2 | 0 | 0 | 8,60707415424738E-06 | -5,065144455 |
| 2188 | Case19 | M148RH | ProM | V2pcUF | 0 | 0 | NA | 0 | -Inf |
| 2189 | Case19 | M148RH | SII | V2pcUF | 0 | 0 | NA | 0 | -Inf |
| 2190 | Case19 | M148RH | STPc | V2pcUF | 80 | 0 | 0 | 0,000344283 | -3,463084464 |
| 2191 | Case19 | M148RH | STPi | V2pcUF | 26 | 0 | 0 | 0,000111892 | -3,951201103 |
| 2192 | Case19 | M148RH | STPr | V2pcUF | 4 | 0 | 0 | 1,72141483084948E-05 | -4,76411446 |
| 2193 | Case19 | M148RH | SUBI | V2pcUF | 0 | 0 | NA | 0 | -Inf |
| 2194 | Case19 | M148RH | TEa/ma | V2pcUF | 16 | 0 | 0 | 6,8856593233979E-05 | -4,162054468 |
| 2195 | Case19 | M148RH | TEa/mp | V2pcUF | 40 | 0 | 0 | 0,000172141 | -3,76411446 |
| 2196 | Case19 | M148RH | TEad | V2pcUF | 2 | 0 | 0 | 8,60707415424738E-06 | -5,065144455 |
| 2197 | Case19 | M148RH | TEav | V2pcUF | 6 | 2 | 0,333333333 | 2,58212224627421E-05 | -4,588023201 |
| 2198 | Case19 | M148RH | TEO | V2pcUF | 927 | 62 | 0,066882416 | 0,003989379 | -2,399094717 |
| 2199 | Case19 | M148RH | TEOm | V2pcUF | 92 | 8 | 0,086956522 | 0,000395925 | -3,402386624 |
| 2200 | Case19 | M148RH | TEpd | V2pcUF | 341 | 2 | 0,005865103 | 0,001467506 | -2,833420072 |
| 2201 | Case19 | M148RH | TEpv | V2pcUF | 2960 | 46 | 0,015540541 | 0,01273847 | -1,89488274 |
| 2202 | Case19 | M148RH | TH_TF | V2pcUF | 2072 | 2 | 0,000965251 | 0,008916929 | -2,0497847 |
| 2203 | Case19 | M148RH | TPt | V2pcUF | 0 | 0 | NA | 0 | -Inf |
| 2204 | Case19 | M148RH | V1c | V2pcUF | 4 | 0 | 0 | 1,72141483084948E-05 | -4,76411446 |
| 2205 | Case19 | M148RH | V1fpLF | V2pcUF | 3392 | 3379 | 0,996167453 | 0,014597598 | -1,835718607 |
| 2206 | Case19 | M148RH | V1fpUF | V2pcUF | 78440 | 77806 | 0,991917389 | 0,337569448 | -0,471636866 |
| 2207 | Case19 | M148RH | V1pcLF | V2pcUF | 198 | 186 | 0,939393939 | 0,0008521 | -3,069509261 |
| 2208 | Case19 | M148RH | V1pcUF | V2pcUF | 11803 | 10269 | 0,870033042 | 0,050794648 | -1,294182044 |
| 2209 | Case19 | M148RH | V2c | V2pcUF | 71600 | 39224 | 0,547821229 | 0,308133255 | -0,511261429 |
| 2210 | Case19 | M148RH | V2fpLF | V2pcUF | 8 | 6 | 0,75 | 3,44282966169895E-05 | -4,463084464 |
| 2211 | Case19 | M148RH | V2fpUF | V2pcUF | 1916 | 1550 | 0,808977035 | 0,008245577 | -2,083778946 |
| 2212 | Case19 | M148RH | V2pcLF | V2pcUF | 90 | 68 | 0,755555556 | 0,000387318 | -3,411931942 |
| 2213 | Case19 | M148RH | V2pcUF | V2pcUF | 0 | 0 | NA | 0 | -Inf |
| 2214 | Case19 | M148RH | V3A | V2pcUF | 34 | 14 | 0,411764706 | 0,00014632 | -3,834695534 |
| 2215 | Case19 | M148RH | V3c | V2pcUF | 492 | 74 | 0,150406504 | 0,00211734 | -2,674209348 |
| 2216 | Case19 | M148RH | V3fpLF | V2pcUF | 12 | 12 | 1 | 5,16424449254843E-05 | -4,286993205 |
| 2217 | Case19 | M148RH | V3fpUF | V2pcUF | 750 | 150 | 0,2 | 0,003227653 | -2,491113188 |
| 2218 | Case19 | M148RH | V3pcLF | V2pcUF | 382 | 58 | 0,151832461 | 0,001643951 | -2,784111088 |
| 2219 | Case19 | M148RH | V3pcUF | V2pcUF | 17506 | 6910 | 0,39472181 | 0,07533772 | -1,122987527 |
| 2220 | Case19 | M148RH | V4c | V2pcUF | 4 | 0 | 0 | 1,72141483084948E-05 | -4,76411446 |
| 2221 | Case19 | M148RH | V4LF | V2pcUF | 324 | 24 | 0,074074074 | 0,001394346 | -2,855629441 |
| 2222 | Case19 | M148RH | V4t | V2pcUF | 108 | 2 | 0,018518519 | 0,000464782 | -3,332750696 |
| 2223 | Case19 | M148RH | V4UF | V2pcUF | 28324 | 7436 | 0,26253354 | 0,121893384 | -0,914019865 |
| 2224 | Case19 | M148RH | V6 | V2pcUF | 2 | 2 | 1 | 8,60707415424738E-06 | -5,065144455 |
| 2225 | Case19 | M148RH | V6A | V2pcUF | 4 | 4 | 1 | 1,72141483084948E-05 | -4,76411446 |
| 2226 | Case19 | M148RH | VIP | V2pcUF | 2 | 0 | 0 | 8,60707415424738E-06 | -5,065144455 |
| 1909 | Case20 | M097LH | 1 | V2fpLF | 0 | 0 | NA | 0 | -Inf |
| 1910 | Case20 | M097LH | 10 | V2fpLF | 0 | 0 | NA | 0 | -Inf |
| 1911 | Case20 | M097LH | 11 | V2fpLF | 0 | 0 | NA | 0 | -Inf |
| 1912 | Case20 | M097LH | 12 | V2fpLF | 0 | 0 | NA | 0 | -Inf |
| 1913 | Case20 | M097LH | 13 | V2fpLF | 0 | 0 | NA | 0 | -Inf |
| 1914 | Case20 | M097LH | 14 | V2fpLF | 0 | 0 | NA | 0 | -Inf |
| 1915 | Case20 | M097LH | 2 | V2fpLF | 0 | 0 | NA | 0 | -Inf |
| 1916 | Case20 | M097LH | 23 | V2fpLF | 35 | 0 | 0 | 0,000253187 | -3,596559398 |
| 1917 | Case20 | M097LH | 24a | V2fpLF | 0 | 0 | NA | 0 | -Inf |

| | Case | MONKEY | SOURCE | TARGET | TOT | SUP | SLN | FLN | lgFLN |
|---|---|---|---|---|---|---|---|---|---|
| 1918 | Case20 | M097LH | 24b | V2fpLF | 0 | 0 | NA | 0 | -Inf |
| 1919 | Case20 | M097LH | 24c | V2fpLF | 0 | 0 | NA | 0 | -Inf |
| 1920 | Case20 | M097LH | 24d | V2fpLF | 0 | 0 | NA | 0 | -Inf |
| 1921 | Case20 | M097LH | 25 | V2fpLF | 0 | 0 | NA | 0 | -Inf |
| 1922 | Case20 | M097LH | 29/30 | V2fpLF | 0 | 0 | NA | 0 | -Inf |
| 1923 | Case20 | M097LH | 3 | V2fpLF | 0 | 0 | NA | 0 | -Inf |
| 1924 | Case20 | M097LH | 31 | V2fpLF | 0 | 0 | NA | 0 | -Inf |
| 1925 | Case20 | M097LH | 32 | V2fpLF | 0 | 0 | NA | 0 | -Inf |
| 1926 | Case20 | M097LH | 44 | V2fpLF | 0 | 0 | NA | 0 | -Inf |
| 1927 | Case20 | M097LH | 45A | V2fpLF | 0 | 0 | NA | 0 | -Inf |
| 1928 | Case20 | M097LH | 45B | V2fpLF | 2 | 0 | 0 | 1,44678019068563E-05 | -4,839597446 |
| 1929 | Case20 | M097LH | 46d | V2fpLF | 0 | 0 | NA | 0 | -Inf |
| 1930 | Case20 | M097LH | 46v | V2fpLF | 0 | 0 | NA | 0 | -Inf |
| 1931 | Case20 | M097LH | 5 | V2fpLF | 0 | 0 | NA | 0 | -Inf |
| 1932 | Case20 | M097LH | 7A | V2fpLF | 22 | 0 | 0 | 0,000159146 | -3,798204761 |
| 1933 | Case20 | M097LH | 7B | V2fpLF | 0 | 0 | NA | 0 | -Inf |
| 1934 | Case20 | M097LH | 7m | V2fpLF | 34 | 0 | 0 | 0,000245953 | -3,609148525 |
| 1935 | Case20 | M097LH | 7op | V2fpLF | 38 | 0 | 0 | 0,000274888 | -3,560843845 |
| 1936 | Case20 | M097LH | 8B | V2fpLF | 1 | 0 | 0 | 7,23390095342815E-06 | -5,140627442 |
| 1937 | Case20 | M097LH | 8l | V2fpLF | 0 | 0 | NA | 0 | -Inf |
| 1938 | Case20 | M097LH | 8m | V2fpLF | 78 | 2 | 0,025641026 | 0,000564244 | -3,248532839 |
| 1939 | Case20 | M097LH | 8r | V2fpLF | 5 | 0 | 0 | 3,61695047671407E-05 | -4,441657438 |
| 1940 | Case20 | M097LH | 9 | V2fpLF | 0 | 0 | NA | 0 | -Inf |
| 1941 | Case20 | M097LH | 9/46d | V2fpLF | 7 | 1 | 0,142857143 | 5,0637306673997E-05 | -4,295529402 |
| 1942 | Case20 | M097LH | 9/46v | V2fpLF | 5 | 1 | 0,2 | 3,61695047671407E-05 | -4,441657438 |
| 1943 | Case20 | M097LH | AIP | V2fpLF | 0 | 0 | NA | 0 | -Inf |
| 1944 | Case20 | M097LH | CORE | V2fpLF | 0 | 0 | NA | 0 | -Inf |
| 1945 | Case20 | M097LH | DP | V2fpLF | 1576 | 139 | 0,08819797 | 0,011400628 | -1,943071229 |
| 1946 | Case20 | M097LH | ENTO | V2fpLF | 0 | 0 | NA | 0 | -Inf |
| 1947 | Case20 | M097LH | F1 | V2fpLF | 0 | 0 | NA | 0 | -Inf |
| 1948 | Case20 | M097LH | F2 | V2fpLF | 0 | 0 | NA | 0 | -Inf |
| 1949 | Case20 | M097LH | F3 | V2fpLF | 0 | 0 | NA | 0 | -Inf |
| 1950 | Case20 | M097LH | F4 | V2fpLF | 0 | 0 | NA | 0 | -Inf |
| 1951 | Case20 | M097LH | F5 | V2fpLF | 0 | 0 | NA | 0 | -Inf |
| 1952 | Case20 | M097LH | F6 | V2fpLF | 0 | 0 | NA | 0 | -Inf |
| 1953 | Case20 | M097LH | F7 | V2fpLF | 0 | 0 | NA | 0 | -Inf |
| 1954 | Case20 | M097LH | FST | V2fpLF | 122 | 4 | 0,032786885 | 0,000882536 | -3,054267611 |
| 1955 | Case20 | M097LH | Gu | V2fpLF | 0 | 0 | NA | 0 | -Inf |
| 1956 | Case20 | M097LH | INSULA | V2fpLF | 0 | 0 | NA | 0 | -Inf |
| 1957 | Case20 | M097LH | IPa | V2fpLF | 0 | 0 | NA | 0 | -Inf |
| 1958 | Case20 | M097LH | LB | V2fpLF | 24 | 0 | 0 | 0,000173614 | -3,7604162 |
| 1959 | Case20 | M097LH | LIP | V2fpLF | 119 | 25 | 0,210084034 | 0,000860834 | -3,065080481 |
| 1960 | Case20 | M097LH | MB | V2fpLF | 4 | 0 | 0 | 2,89356038137126E-05 | -4,538567451 |
| 1961 | Case20 | M097LH | MIP | V2fpLF | 503 | 20 | 0,039761431 | 0,003638652 | -2,439059457 |
| 1962 | Case20 | M097LH | MST | V2fpLF | 11520 | 2393 | 0,207725694 | 0,083334539 | -1,079174963 |
| 1963 | Case20 | M097LH | MTc | V2fpLF | 93 | 0 | 0 | 0,000672753 | -3,172144493 |
| 1964 | Case20 | M097LH | MTp | V2fpLF | 7596 | 1371 | 0,180489731 | 0,054948712 | -1,260042486 |
| 1965 | Case20 | M097LH | OPAl | V2fpLF | 0 | 0 | NA | 0 | -Inf |
| 1966 | Case20 | M097LH | OPro | V2fpLF | 0 | 0 | NA | 0 | -Inf |
| 1967 | Case20 | M097LH | PBc | V2fpLF | 30 | 0 | 0 | 0,000217017 | -3,663506187 |
| 1968 | Case20 | M097LH | PBr | V2fpLF | 0 | 0 | NA | 0 | -Inf |
| 1969 | Case20 | M097LH | PERI | V2fpLF | 81 | 3 | 0,037037037 | 0,000585946 | -3,232142423 |
| 1970 | Case20 | M097LH | PGa | V2fpLF | 30 | 0 | 0 | 0,000217017 | -3,663506187 |
| 1971 | Case20 | M097LH | Pi | V2fpLF | 0 | 0 | NA | 0 | -Inf |
| 1972 | Case20 | M097LH | PIP | V2fpLF | 465 | 24 | 0,051612903 | 0,003363764 | -2,473174489 |
| 1973 | Case20 | M097LH | Pir | V2fpLF | 0 | 0 | NA | 0 | -Inf |
| 1974 | Case20 | M097LH | POLE | V2fpLF | 0 | 0 | NA | 0 | -Inf |
| 1975 | Case20 | M097LH | Pro.St | V2fpLF | 706 | 48 | 0,067988669 | 0,005107134 | -2,291822741 |
| 1976 | Case20 | M097LH | ProM | V2fpLF | 0 | 0 | NA | 0 | -Inf |
| 1977 | Case20 | M097LH | SII | V2fpLF | 6 | 0 | 0 | 4,34034057205689E-05 | -4,362476192 |
| 1978 | Case20 | M097LH | STPc | V2fpLF | 790 | 120 | 0,151898734 | 0,005714782 | -2,243000351 |
| 1979 | Case20 | M097LH | STPi | V2fpLF | 13 | 0 | 0 | 9,40407123945659E-05 | -4,02668409 |
| 1980 | Case20 | M097LH | STPr | V2fpLF | 1 | 0 | 0 | 7,23390095342815E-06 | -5,140627442 |
| 1981 | Case20 | M097LH | SUBI | V2fpLF | 0 | 0 | NA | 0 | -Inf |
| 1982 | Case20 | M097LH | TEa/ma | V2fpLF | 0 | 0 | NA | 0 | -Inf |
| 1983 | Case20 | M097LH | TEa/mp | V2fpLF | 42 | 0 | 0 | 0,000303824 | -3,517378151 |
| 1984 | Case20 | M097LH | TEad | V2fpLF | 0 | 0 | NA | 0 | -Inf |
| 1985 | Case20 | M097LH | TEav | V2fpLF | 10 | 0 | 0 | 7,23390095342815E-05 | -4,140627442 |
| 1986 | Case20 | M097LH | TEO | V2fpLF | 4 | 4 | 1 | 2,89356038137126E-05 | -4,538567451 |
| 1987 | Case20 | M097LH | TEOm | V2fpLF | 0 | 0 | NA | 0 | -Inf |
| 1988 | Case20 | M097LH | TEpd | V2fpLF | 2 | 0 | 0 | 1,44678019068563E-05 | -4,839597446 |
| 1989 | Case20 | M097LH | TEpv | V2fpLF | 472 | 53 | 0,112288136 | 0,003414401 | -2,466685443 |
| 1990 | Case20 | M097LH | TH_TF | V2fpLF | 1901 | 299 | 0,157285639 | 0,013751646 | -1,861645325 |
| 1991 | Case20 | M097LH | TPt | V2fpLF | 82 | 0 | 0 | 0,00059318 | -3,22681359 |
| 1992 | Case20 | M097LH | V1c | V2fpLF | 28 | 28 | 1 | 0,000202549 | -3,693469411 |

| | Case | MONKEY | SOURCE | TARGET | TOT | SUP | SLN | FLN | lgFLN |
|---|---|---|---|---|---|---|---|---|---|
| 1993 | Case20 | M097LH | V1fpLF | V2fpLF | 93078 | 83169 | 0,893540901 | 0,673317033 | -0,171780399 |
| 1994 | Case20 | M097LH | V1fpUF | V2fpLF | 2622 | 2576 | 0,98245614 | 0,018967288 | -1,721994755 |
| 1995 | Case20 | M097LH | V1pcLF | V2fpLF | 0 | 0 | NA | 0 | -Inf |
| 1996 | Case20 | M097LH | V1pcUF | V2fpLF | 0 | 0 | NA | 0 | -Inf |
| 1997 | Case20 | M097LH | V2c | V2fpLF | 0 | 0 | NA | 0 | -Inf |
| 1998 | Case20 | M097LH | V2fpLF | V2fpLF | 0 | 0 | NA | 0 | -Inf |
| 1999 | Case20 | M097LH | V2fpUF | V2fpLF | 0 | 0 | NA | 0 | -Inf |
| 2000 | Case20 | M097LH | V2pcLF | V2fpLF | 9 | 0 | 0 | 6,51051085808533E-05 | -4,186384932 |
| 2001 | Case20 | M097LH | V2pcUF | V2fpLF | 22 | 11 | 0,5 | 0,000159146 | -3,798204761 |
| 2002 | Case20 | M097LH | V3A | V2fpLF | 215 | 211 | 0,981395349 | 0,001555289 | -2,808188982 |
| 2003 | Case20 | M097LH | V3c | V2fpLF | 0 | 0 | NA | 0 | -Inf |
| 2004 | Case20 | M097LH | V3fpLF | V2fpLF | 13 | 11 | 0,846153846 | 9,40407123945659E-05 | -4,02668409 |
| 2005 | Case20 | M097LH | V3fpUF | V2fpLF | 337 | 141 | 0,418397626 | 0,002437825 | -2,612997541 |
| 2006 | Case20 | M097LH | V3pcLF | V2fpLF | 0 | 0 | NA | 0 | -Inf |
| 2007 | Case20 | M097LH | V3pcUF | V2fpLF | 4 | 0 | 0 | 2,89356038137126E-05 | -4,538567451 |
| 2008 | Case20 | M097LH | V4c | V2fpLF | 3 | 3 | 1 | 2,17017028602844E-05 | -4,663506187 |
| 2009 | Case20 | M097LH | V4LF | V2fpLF | 756 | 140 | 0,185185185 | 0,005468829 | -2,262105646 |
| 2010 | Case20 | M097LH | V4t | V2fpLF | 0 | 0 | NA | 0 | -Inf |
| 2011 | Case20 | M097LH | V4UF | V2fpLF | 1942 | 725 | 0,373326468 | 0,014048236 | -1,852378216 |
| 2012 | Case20 | M097LH | V6 | V2fpLF | 8659 | 6080 | 0,702159603 | 0,062638348 | -1,203159702 |
| 2013 | Case20 | M097LH | V6A | V2fpLF | 3559 | 499 | 0,140207924 | 0,025745453 | -1,589299454 |
| 2014 | Case20 | M097LH | VIP | V2fpLF | 572 | 54 | 0,094405594 | 0,004137791 | -2,383231413 |
| 1803 | Case21 | M096LH | 1 | V2fpUF | 0 | 0 | NA | 0 | -Inf |
| 1804 | Case21 | M096LH | 10 | V2fpUF | 0 | 0 | NA | 0 | -Inf |
| 1805 | Case21 | M096LH | 11 | V2fpUF | 0 | 0 | NA | 0 | -Inf |
| 1806 | Case21 | M096LH | 12 | V2fpUF | 0 | 0 | NA | 0 | -Inf |
| 1807 | Case21 | M096LH | 13 | V2fpUF | 0 | 0 | NA | 0 | -Inf |
| 1808 | Case21 | M096LH | 14 | V2fpUF | 0 | 0 | NA | 0 | -Inf |
| 1809 | Case21 | M096LH | 2 | V2fpUF | 0 | 0 | NA | 0 | -Inf |
| 1810 | Case21 | M096LH | 23 | V2fpUF | 23 | 9 | 0,391304348 | 9,92260369120857E-05 | -4,003374354 |
| 1811 | Case21 | M096LH | 24a | V2fpUF | 2 | 0 | 0 | 8,62835103583354E-06 | -5,064072194 |
| 1812 | Case21 | M096LH | 24b | V2fpUF | 0 | 0 | NA | 0 | -Inf |
| 1813 | Case21 | M096LH | 24c | V2fpUF | 0 | 0 | NA | 0 | -Inf |
| 1814 | Case21 | M096LH | 24d | V2fpUF | 3 | 0 | 0 | 1,29425265537503E-05 | -4,887980935 |
| 1815 | Case21 | M096LH | 25 | V2fpUF | 0 | 0 | NA | 0 | -Inf |
| 1816 | Case21 | M096LH | 29/30 | V2fpUF | 0 | 0 | NA | 0 | -Inf |
| 1817 | Case21 | M096LH | 3 | V2fpUF | 0 | 0 | NA | 0 | -Inf |
| 1818 | Case21 | M096LH | 31 | V2fpUF | 0 | 0 | NA | 0 | -Inf |
| 1819 | Case21 | M096LH | 32 | V2fpUF | 0 | 0 | NA | 0 | -Inf |
| 1820 | Case21 | M096LH | 44 | V2fpUF | 0 | 0 | NA | 0 | -Inf |
| 1821 | Case21 | M096LH | 45A | V2fpUF | 0 | 0 | NA | 0 | -Inf |
| 1822 | Case21 | M096LH | 45B | V2fpUF | 0 | 0 | NA | 0 | -Inf |
| 1823 | Case21 | M096LH | 46d | V2fpUF | 0 | 0 | NA | 0 | -Inf |
| 1824 | Case21 | M096LH | 46v | V2fpUF | 0 | 0 | NA | 0 | -Inf |
| 1825 | Case21 | M096LH | 5 | V2fpUF | 0 | 0 | NA | 0 | -Inf |
| 1826 | Case21 | M096LH | 7A | V2fpUF | 63 | 2 | 0,031746032 | 0,000271793 | -3,565761641 |
| 1827 | Case21 | M096LH | 7B | V2fpUF | 0 | 0 | NA | 0 | -Inf |
| 1828 | Case21 | M096LH | 7m | V2fpUF | 3 | 0 | 0 | 1,29425265537503E-05 | -4,887980935 |
| 1829 | Case21 | M096LH | 7op | V2fpUF | 0 | 0 | NA | 0 | -Inf |
| 1830 | Case21 | M096LH | 8B | V2fpUF | 5 | 1 | 0,2 | 2,15708775895839E-05 | -4,666132186 |
| 1831 | Case21 | M096LH | 8l | V2fpUF | 0 | 0 | NA | 0 | -Inf |
| 1832 | Case21 | M096LH | 8m | V2fpUF | 206 | 74 | 0,359223301 | 0,00088872 | -3,05123497 |
| 1833 | Case21 | M096LH | 8r | V2fpUF | 8 | 3 | 0,375 | 3,45134041433342E-05 | -4,462012203 |
| 1834 | Case21 | M096LH | 9 | V2fpUF | 0 | 0 | NA | 0 | -Inf |
| 1835 | Case21 | M096LH | 9/46d | V2fpUF | 5 | 1 | 0,2 | 2,15708775895839E-05 | -4,666132186 |
| 1836 | Case21 | M096LH | 9/46v | V2fpUF | 1 | 1 | 1 | 4,31417551791677E-06 | -5,36510219 |
| 1837 | Case21 | M096LH | AIP | V2fpUF | 0 | 0 | NA | 0 | -Inf |
| 1838 | Case21 | M096LH | CORE | V2fpUF | 0 | 0 | NA | 0 | -Inf |
| 1839 | Case21 | M096LH | DP | V2fpUF | 6984 | 1450 | 0,207617411 | 0,030130202 | -1,520997959 |
| 1840 | Case21 | M096LH | ENTO | V2fpUF | 27 | 2 | 0,074074074 | 0,000116483 | -3,933738426 |
| 1841 | Case21 | M096LH | F1 | V2fpUF | 0 | 0 | NA | 0 | -Inf |
| 1842 | Case21 | M096LH | F2 | V2fpUF | 0 | 0 | NA | 0 | -Inf |
| 1843 | Case21 | M096LH | F3 | V2fpUF | 0 | 0 | NA | 0 | -Inf |
| 1844 | Case21 | M096LH | F4 | V2fpUF | 0 | 0 | NA | 0 | -Inf |
| 1845 | Case21 | M096LH | F5 | V2fpUF | 0 | 0 | NA | 0 | -Inf |
| 1846 | Case21 | M096LH | F6 | V2fpUF | 0 | 0 | NA | 0 | -Inf |
| 1847 | Case21 | M096LH | F7 | V2fpUF | 2 | 0 | 0 | 8,62835103583354E-06 | -5,064072194 |
| 1848 | Case21 | M096LH | FST | V2fpUF | 395 | 14 | 0,035443038 | 0,001704099 | -2,768505094 |
| 1849 | Case21 | M096LH | Gu | V2fpUF | 0 | 0 | NA | 0 | -Inf |
| 1850 | Case21 | M096LH | INSULA | V2fpUF | 1 | 0 | 0 | 4,31417551791677E-06 | -5,36510219 |
| 1851 | Case21 | M096LH | IPa | V2fpUF | 3 | 0 | 0 | 1,29425265537503E-05 | -4,887980935 |
| 1852 | Case21 | M096LH | LB | V2fpUF | 10 | 0 | 0 | 4,31417551791677E-05 | -4,36510219 |
| 1853 | Case21 | M096LH | LIP | V2fpUF | 1010 | 380 | 0,376237624 | 0,004357317 | -2,360780816 |
| 1854 | Case21 | M096LH | MB | V2fpUF | 8 | 0 | 0 | 3,45134041433342E-05 | -4,462012203 |
| 1855 | Case21 | M096LH | MIP | V2fpUF | 643 | 38 | 0,059097978 | 0,002774015 | -2,556891217 |

| | Case | MONKEY | SOURCE | TARGET | TOT | SUP | SLN | FLN | lgFLN |
|---|---|---|---|---|---|---|---|---|---|
| 1856 | Case21 | M096LH | MST | V2fpUF | 3023 | 188 | 0,062189878 | 0,013041753 | -1,884664043 |
| 1857 | Case21 | M096LH | MTc | V2fpUF | 556 | 144 | 0,258992806 | 0,002398682 | -2,620027398 |
| 1858 | Case21 | M096LH | MTp | V2fpUF | 14298 | 5585 | 0,390614072 | 0,061684082 | -1,209826897 |
| 1859 | Case21 | M096LH | OPAl | V2fpUF | 0 | 0 | NA | 0 | -Inf |
| 1860 | Case21 | M096LH | OPro | V2fpUF | 0 | 0 | NA | 0 | -Inf |
| 1861 | Case21 | M096LH | PBc | V2fpUF | 17 | 0 | 0 | 7,33409838045851E-05 | -4,134653269 |
| 1862 | Case21 | M096LH | PBr | V2fpUF | 0 | 0 | NA | 0 | -Inf |
| 1863 | Case21 | M096LH | PERI | V2fpUF | 135 | 0 | 0 | 0,000582414 | -3,234768422 |
| 1864 | Case21 | M096LH | PGa | V2fpUF | 194 | 6 | 0,030927835 | 0,00083695 | -3,07730046 |
| 1865 | Case21 | M096LH | Pi | V2fpUF | 0 | 0 | NA | 0 | -Inf |
| 1866 | Case21 | M096LH | PIP | V2fpUF | 4043 | 1597 | 0,39500371 | 0,017442212 | -1,758398449 |
| 1867 | Case21 | M096LH | Pir | V2fpUF | 0 | 0 | NA | 0 | -Inf |
| 1868 | Case21 | M096LH | POLE | V2fpUF | 0 | 0 | NA | 0 | -Inf |
| 1869 | Case21 | M096LH | Pro.St | V2fpUF | 1 | 0 | 0 | 4,31417551791677E-06 | -5,36510219 |
| 1870 | Case21 | M096LH | ProM | V2fpUF | 0 | 0 | NA | 0 | -Inf |
| 1871 | Case21 | M096LH | SII | V2fpUF | 0 | 0 | NA | 0 | -Inf |
| 1872 | Case21 | M096LH | STPc | V2fpUF | 242 | 7 | 0,02892562 | 0,00104403 | -2,981286824 |
| 1873 | Case21 | M096LH | STPi | V2fpUF | 13 | 0 | 0 | 5,6084281732918E-05 | -4,251158838 |
| 1874 | Case21 | M096LH | STPr | V2fpUF | 0 | 0 | NA | 0 | -Inf |
| 1875 | Case21 | M096LH | SUBI | V2fpUF | 0 | 0 | NA | 0 | -Inf |
| 1876 | Case21 | M096LH | TEa/ma | V2fpUF | 2 | 0 | 0 | 8,62835103583354E-06 | -5,064072194 |
| 1877 | Case21 | M096LH | TEa/mp | V2fpUF | 4 | 0 | 0 | 1,72567020716671E-05 | -4,763042199 |
| 1878 | Case21 | M096LH | TEad | V2fpUF | 4 | 0 | 0 | 1,72567020716671E-05 | -4,763042199 |
| 1879 | Case21 | M096LH | TEav | V2fpUF | 14 | 0 | 0 | 6,03984572508348E-05 | -4,218974154 |
| 1880 | Case21 | M096LH | TEO | V2fpUF | 1142 | 40 | 0,03502627 | 0,004926788 | -2,307436086 |
| 1881 | Case21 | M096LH | TEOm | V2fpUF | 7 | 2 | 0,285714286 | 3,01992286254174E-05 | -4,52000415 |
| 1882 | Case21 | M096LH | TEpd | V2fpUF | 301 | 2 | 0,006644518 | 0,001298567 | -2,886535694 |
| 1883 | Case21 | M096LH | TEpv | V2fpUF | 1883 | 18 | 0,009559214 | 0,008123593 | -2,09025187 |
| 1884 | Case21 | M096LH | TH_TF | V2fpUF | 4789 | 221 | 0,046147421 | 0,020660587 | -1,684857353 |
| 1885 | Case21 | M096LH | TPt | V2fpUF | 9 | 0 | 0 | 3,88275796612509E-05 | -4,410859681 |
| 1886 | Case21 | M096LH | V1c | V2fpUF | 2 | 2 | 1 | 8,62835103583354E-06 | -5,064072194 |
| 1887 | Case21 | M096LH | V1fpLF | V2fpUF | 39296 | 33771 | 0,859400448 | 0,169529841 | -0,770753845 |
| 1888 | Case21 | M096LH | V1fpUF | V2fpUF | 65731 | 62357 | 0,948669578 | 0,283575071 | -0,547331951 |
| 1889 | Case21 | M096LH | V1pcLF | V2fpUF | 0 | 0 | NA | 0 | -Inf |
| 1890 | Case21 | M096LH | V1pcUF | V2fpUF | 0 | 0 | NA | 0 | -Inf |
| 1891 | Case21 | M096LH | V2c | V2fpUF | 0 | 0 | NA | 0 | -Inf |
| 1892 | Case21 | M096LH | V2fpLF | V2fpUF | 0 | 0 | NA | 0 | -Inf |
| 1893 | Case21 | M096LH | V2fpUF | V2fpUF | 0 | 0 | NA | 0 | -Inf |
| 1894 | Case21 | M096LH | V2pcLF | V2fpUF | 0 | 0 | NA | 0 | -Inf |
| 1895 | Case21 | M096LH | V2pcUF | V2fpUF | 0 | 0 | NA | 0 | -Inf |
| 1896 | Case21 | M096LH | V3A | V2fpUF | 656 | 279 | 0,425304878 | 0,002830099 | -2,548198351 |
| 1897 | Case21 | M096LH | V3c | V2fpUF | 0 | 0 | NA | 0 | -Inf |
| 1898 | Case21 | M096LH | V3fpLF | V2fpUF | 564 | 182 | 0,322695035 | 0,002433195 | -2,613823086 |
| 1899 | Case21 | M096LH | V3fpUF | V2fpUF | 876 | 311 | 0,355022831 | 0,003779218 | -2,422598084 |
| 1900 | Case21 | M096LH | V3pcLF | V2fpUF | 0 | 0 | NA | 0 | -Inf |
| 1901 | Case21 | M096LH | V3pcUF | V2fpUF | 2285 | 428 | 0,187308534 | 0,009857891 | -2,006215986 |
| 1902 | Case21 | M096LH | V4c | V2fpUF | 0 | 0 | NA | 0 | -Inf |
| 1903 | Case21 | M096LH | V4LF | V2fpUF | 7640 | 2656 | 0,347643979 | 0,032960301 | -1,482008831 |
| 1904 | Case21 | M096LH | V4t | V2fpUF | 0 | 0 | NA | 0 | -Inf |
| 1905 | Case21 | M096LH | V4UF | V2fpUF | 45989 | 15749 | 0,342451456 | 0,198404618 | -0,702448224 |
| 1906 | Case21 | M096LH | V6 | V2fpUF | 24708 | 12546 | 0,507770763 | 0,106594649 | -0,972264597 |
| 1907 | Case21 | M096LH | V6A | V2fpUF | 3654 | 734 | 0,200875753 | 0,015763997 | -1,802333647 |
| 1908 | Case21 | M096LH | VIP | V2fpUF | 314 | 76 | 0,242038217 | 0,001354651 | -2,868172542 |
