## Supplementary material for "Retinotopic organization of feedback projections in primate early visual cortex: implications for active vision": Table S3

Tractography Distance of Visual Area V1 and V2

| | V1c | V1pcLF | V1pcUF | V1fpLF | V1fpUF | V2c | V2pcLF | V2pcUF | V2fpLF | V2fpUF |
|---|---|---|---|---|---|---|---|---|---|---|
| 1 | 34,13915 | 37,84365 | 34,04545 | 35,5716 | 34,44775 | 37,54155 | 35,58335 | 31,15345 | 27,2533 | 30,2044831 |
| 10 | 37,19505 | 40,74705 | 40,2027 | 30,0361 | 30,31775 | 37,6271 | 40,6564 | 40,66695 | 35,61255 | 32,3087 |
| 11 | 37,5221 | 37,5472 | 36,5203 | 29,18705 | 28,1926 | 35,65335 | 39,8028 | 35,995 | 35,4368 | 33,4655 |
| 12 | 39,0362 | 40,4979 | 40,86875 | 31,3934 | 31,43485 | 39,13 | 41,04845 | 41,9206 | 35,247 | 34,374 |
| 13 | 31,0883 | 36,67955 | 35,19 | 28,74725 | 30,3301 | 31,3949 | 37,605 | 33,84494652 | 31,69685 | 28,9943 |
| 14 | 28,88645 | 35,5499 | 33,1346 | 29,00205 | 29,3605 | 28,94295 | 37,26785 | 28,30345 | 33,56375 | 26,5009 |
| 2 | 34,88335 | 37,4675 | 35,6138 | 34,50035 | 34,04505 | 35,1025 | 38,48105 | 40,18705 | 33,2286 | 38,92755 |
| 23 | 22,969 | 24,41938333 | 28,66523333 | 13,00426833 | 21,49476667 | 22,96276667 | 22,42616667 | 28,16535 | 10,18061167 | 19,07566667 |
| 24a | 28,71055 | 32,8775 | 33,53765 | 23,1066 | 27,8453 | 30,5333 | 33,8278 | 28,9315 | 27,93265 | 24,0606 |
| 24b | 30,41745 | 37,08755 | 34,7683 | 16,96265 | 20,6496 | 28,17803932 | 33,92505 | 31,95597745 | 18,20505 | 22,25 |
| 24c | 33,6581 | 39,2936 | 38,01675 | 21,9403 | 27,07555 | 35,5391 | 39,61755 | 35,65546643 | 24,89925 | 30,3735 |
| 24d | 31,878 | 33,5712 | 33,36785 | 18,29375 | 25,7277 | 29,65355 | 24,8987 | 29,21 | 18,6871 | 21,3384 |
| 25 | 28,4094 | 33,26345 | 32,65545 | 29,05665 | 28,71285 | 29,39805 | 34,0041 | 29,3145 | 30,84705 | 26,3737 |
| 29/30 | 20,1452 | 21,6015 | 25,54545 | 12,6543 | 14,4862 | 17,97705 | 18,451825 | 23,6001 | 14,1625 | 21,52405 |
| 3 | 35,44115 | 38,55135 | 35,0785 | 36,29755 | 35,66325 | 36,06725 | 39,5743 | 35,1756 | 32,511 | 29,53695 |
| 31 | 24,454 | 23,4093 | 31,32985 | 15,24315 | 24,6416 | 22,9764 | 23,6088 | 30,17705 | 14,1737 | 16,99365 |
| 32 | 33,63465 | 37,5435 | 37,2178 | 29,0404 | 29,3704 | 35,52965 | 38,73655 | 34,5575 | 37,349 | 30,37075 |
| 44 | 40,05625 | 40,0147 | 40,72825 | 32,28205 | 31,56055 | 39,50975 | 38,2921 | 48,1852 | 34,214 | 33,09125 |
| 45A | 39,2139 | 39,22835 | 41,89145 | 32,54605 | 31,8696 | 37,6218 | 38,18005 | 37,49 | 34,39455 | 32,623 |
| 45B | 37,5015 | 39,89935 | 38,73145 | 30,3423 | 30,04075 | 35,60895 | 38,35435 | 38,35142015 | 35,83205 | 28,93685 |
| 46d | 36,1018 | 39,1144 | 40,36795 | 29,85875 | 29,98275 | 36,5355663 | 37,3606 | 37,87312688 | 33,31165 | 32,90145 |
| 46v | 35,70815 | 38,78445 | 39,0056 | 29,19155 | 29,5915 | 36,70145 | 38,8538 | 37,0115 | 34,47205 | 30,5028 |
| 5 | 30,56425 | 32,8647 | 33,05635 | 30,65595 | 32,9048 | 31,6137 | 32,5713 | 33,97955 | 21,20145 | 28,3475 |
| 7A | 25,4315 | 23,9324 | 30,2124 | 16,68435 | 26,99675 | 23,23375 | 21,1968 | 27,57055 | 10,84515 | 25,8261 |
| 7B | 39,13135 | 38,9454 | 35,6521 | 35,8493 | 34,3996 | 36,57175 | 37,49005 | 34,75050839 | 27,1568 | 35,6321 |
| 7m | 26,56075 | 26,86285 | 33,05195 | 19,23865 | 31,2063 | 27,126 | 27,20035 | 28,55535 | 15,279 | 23,2853 |
| 7op | 27,2087 | 28,70475 | 30,7789 | 25,7931 | 30,71155 | 24,2545 | 27,0846 | 29,8204 | 18,74725 | 28,74035 |
| 8B | 37,30075 | 42,5699 | 39,8867 | 27,37585 | 30,4885 | 37,80625 | 43,1826 | 37,57482888 | 29,9958 | 31,26395 |
| 8l | 35,9584 | 40,3322 | 41,0455 | 31,27175 | 29,2665 | 34,96 | 37,7201 | 37,98181846 | 34,0678 | 25,8558 |
| 8m | 34,73 | 39,3301 | 40,29315 | 24,9536 | 28,0357 | 32,68875 | 43,1251 | 36,57370899 | 36,1265 | 33,8401 |
| 8r | 36,6873 | 37,61325 | 41,03865 | 28,5973 | 30,94365 | 41,5151 | 41,2851 | 40,4226 | 33,005 | 28,31295 |
| 9 | 37,08465 | 39,50695 | 41,881 | 30,0022 | 30,88655 | 38,5737 | 38,18 | 38,9851 | 37,4774 | 28,34345 |
| 9/46d | 36,7617 | 37,2025 | 39,2725 | 28,9152 | 29,8832 | 37,881 | 38,3216306 | 35,42 | 33,8675 | 27,0537 |
| 9/46v | 35,9271 | 39,0181 | 40,79835 | 31,26395 | 29,82575 | 37,85145 | 39,9769 | 39,65335405 | 34,4865 | 31,42475 |
| AIP | 34,8764 | 38,8495 | 33,52245 | 31,93105 | 31,0118 | 47,3802 | 46,6902 | 34,2412 | 31,29665 | 24,4949 |
| CORE | 19,22955 | 21,7388 | 27,8846 | 20,91275 | 27,063 | 19,26305 | 22,47625 | 29,03095 | 20,3659 | 28,60765 |
| DP | 21,4292 | 16,98965 | 27,9179 | 13,12385 | 24,7418 | 15,27155 | 11,62445 | 20,4295 | 12,4588 | 24,91045 |
| ENTO | 27,765 | 27,8338 | 30,75845 | 24,01605 | 20,55495 | 20,73845 | 26,1898 | 20,94565 | 24,8284 | 16,53335 |
| F1 | 33,23615 | 35,7994 | 35,0271 | 33,56445 | 33,37885 | 32,961 | 36,27385 | 34,57355 | 31,16775 | 29,802 |
| F2 | 34,9326 | 34,77775 | 36,2301 | 33,01005 | 33,6553 | 34,8425 | 36,26915 | 43,31675 | 30,0023 | 31,7133 |
| F3 | 37,82035 | 38,8583 | 38,86075 | 27,84785 | 33,99825 | 36,61605 | 39,7414 | 39,3301 | 25,0705 | 21,488 |
| F4 | 38,73165 | 39,3561 | 41,3267 | 33,31025 | 34,14405 | 31,75435 | 34,6708 | 43,0676 | 34,9741 | 33,86745 |
| F5 | 35,80095 | 40,0022 | 44,81525 | 35,13985 | 33,11355 | 44,0399 | 37,49575 | 43,7823 | 37,9132 | 34,11665 |
| F6 | 30,35995 | 44,5051 | 41,7451 | 24,49965 | 29,8272 | 32,9935 | 38,7551 | 39,27303416 | 28,5701 | 36,25035 |
| F7 | 36,45862675 | 37,3894 | 42,4926 | 34,1911 | 37,1167 | 32,614 | 39,1001 | 48,7602 | 27,6144 | 31,41825606 |
| FST | 19,3722 | 16,21015 | 25,7063 | 14,88905 | 19,03425 | 14,68795 | 16,12325 | 19,5954 | 13,9178 | 18,35605 |
| Gu | 40,32675 | 35,6911 | 42,09005 | 29,8546 | 35,0001 | 37,82361183 | 42,2051 | 37,76423402 | 37,605 | 36,57 |
| INSULA | 21,49895 | 23,16085 | 26,94635 | 21,9853 | 25,43165 | 19,25105 | 23,0025 | 23,19065 | 21,10275 | 21,85735 |
| IPa | 24,77105 | 23,345 | 28,0383 | 21,35585 | 26,70365 | 17,2023 | 24,86295 | 22,05385 | 22,42295 | 20,3572 |
| LB | 23,16265 | 24,13225 | 29,6427 | 19,7806 | 28,7297 | 21,77135 | 22,8893 | 30,4061 | 19,3746 | 27,1675 |
| LIP | 23,74925 | 22,576925 | 27,305525 | 15,194475 | 22,7233 | 20,8032 | 17,469375 | 22,8272 | 10,1315 | 23,674525 |
| MB | 18,35105 | 22,2185 | 24,05045 | 21,17965 | 25,43205 | 19,254 | 22,7953 | 26,0822 | 20,4945 | 26,576 |
| MIP | 26,57605 | 27,2693 | 32,29605 | 20,9976 | 32,4761 | 27,1374 | 28,88485 | 25,75325 | 15,56705 | 21,89535 |
| MST | 18,91965 | 16,1441 | 25,9262 | 12,3941 | 23,68635 | 15,2169 | 14,61885 | 22,03885 | 11,50435 | 20,65375 |
| MTc | 14,2153 | 13,9152 | 22,16905 | 12,6995 | 21,85965 | 13,47825 | 13,54835 | 18,62205 | 12,16405 | 19,97145 |
| MTp | 18,33885 | 14,9997 | 23,9119 | 13,2985 | 23,70215 | 16,2176 | 13,5967 | 20,1517 | 12,43995 | 20,942 |
| OPAI | 24,48495 | 32,78075 | 30,44875 | 30,44315 | 29,9685 | 24,79685 | 30,29765 | 23,17815 | 27,51575 | 27,3699 |
| OPro | 21,97305 | 26,9599 | 24,0619 | 24,3791 | 28,23745 | 19,8601 | 23,7799 | 23,38395 | 23,43555 | 18,8052 |
| PBc | 24,085 | 24,4889 | 30,0332 | 21,2502 | 28,49995 | 21,5181 | 22,1664 | 29,9439 | 21,4376 | 31,5867 |
| PBr | 24,30875 | 26,86615 | 26,1898 | 23,9271 | 24,88155 | 20,6357 | 24,36785 | 23,35215 | 22,27495 | 23,70575 |
| PERI | 28,10205 | 31,43995 | 34,16925 | 26,964 | 26,81365 | 23,8073 | 31,5336 | 24,55625 | 29,8956 | 20,70705 |
| PGa | 18,6094 | 17,0932 | 26,02755 | 17,1186 | 23,2141 | 16,255 | 17,34845 | 21,122 | 17,0172 | 18,5598 |
| Pi | 19,40205 | 26,1343 | 22,77505 | 25,0225 | 24,95195 | 19,3495 | 24,9187 | 23,6162 | 24,18185 | 24,97335 |
| PIP | 17,0902 | 14,2648 | 21,4899 | 8,958965 | 20,81925 | 15,343 | 9,771825 | 17,4843 | 8,84411 | 18,7961 |
| Pir | 22,06428896 | 25,78144032 | 25,90422155 | 24,83476686 | 26,45774348 | 21,70424811 | 24,81317951 | 23,90746667 | 22,68347872 | 20,71500378 |
| POLE | 25,27651667 | 28,86455 | 29,4748 | 27,52296667 | 26,0374 | 23,70153333 | 28,55773333 | 23,9245 | 27,5448 | 21,37001667 |
| Pro.St | 15,38955 | 17,96985 | 19,13325 | 3,911975 | 3,26359 | 14,6714 | 13,32385 | 18,84775 | 8,62587 | 3,04787 |
| ProM | 38,35955 | 41,4383 | 44,83815 | 34,87755 | 34,1575 | 37,14505 | 45,0802 | 37,145 | 40,0351 | 38,2663 |
| SII | 34,30025 | 36,5331 | 32,95125 | 34,3982 | 33,5639 | 39,0713 | 33,4284 | 37,96259615 | 29,8542 | 35,8608 |
| STPc | 19,72645 | 20,33065 | 27,14375 | 17,47895 | 25,4393 | 17,5591 | 19,8219 | 25,2932 | 17,09035 | 25,62195 |
| STPi | 22,65855 | 22,06475 | 28,79405 | 21,1654 | 23,56345 | 17,4873 | 22,14865 | 20,84475 | 20,2905 | 21,58885 |
| STPr | 27,05725 | 26,45915 | 31,57975 | 24,709 | 26,3253 | 20,56165 | 25,64255 | 22,5558 | 24,576 | 20,87065 |
| SUBI | 17,5276 | 22,8207 | 19,54285 | 8,86674 | 11,5512 | 16,1886 | 23,3463 | 17,06815 | 12,47605 | 4,750455 |
| TEa/ma | 26,70015 | 27,55575 | 30,71025 | 25,5515 | 29,7208 | 21,2267 | 28,42805 | 22,88745 | 26,67505 | 21,1371 |
| TEa/mp | 20,89455 | 21,58025 | 24,67655 | 20,91325 | 23,3976 | 15,76595 | 22,7002 | 19,65945 | 20,83905 | 19,35085 |

| | V1c | V1pcLF | V1pcUF | V1fpLF | V1fpUF | V2c | V2pcLF | V2pcUF | V2fpLF | V2fpUF |
|---|---|---|---|---|---|---|---|---|---|---|
| TEad | 28,66055 | 28,96285 | 36,10375 | 25,0534 | 27,89575 | 21,297 | 28,5293 | 23,2789 | 27,05885 | 19,4292 |
| TEav | 26,8517 | 29,6286 | 29,5082 | 26,87625 | 26,69485 | 23,38635 | 29,93945 | 24,2604 | 28,24275 | 20,6165 |
| TEO | 16,5645 | 20,80845 | 19,6503 | 17,60975 | 18,7511 | 11,32945 | 19,46155 | 14,18885 | 19,5728 | 17,2477 |
| TEOm | 15,67325 | 17,9565 | 20,22705 | 16,987 | 20,0118 | 13,28075 | 18,12245 | 17,7745 | 16,70765 | 20,049 |
| TEpd | 21,25625 | 23,9713 | 26,2402 | 21,14645 | 23,8848 | 16,1311 | 23,71115 | 19,48425 | 22,50715 | 18,68025 |
| TEpv | 17,44405 | 22,17825 | 19,23135 | 19,8982 | 17,6295 | 15,73145 | 22,73345 | 16,0682 | 21,6169 | 12,6189 |
| TH/TF | 18,463775 | 21,831325 | 20,1289 | 14,376625 | 13,713025 | 15,431575 | 23,421375 | 16,426125 | 19,333125 | 8,09902 |
| TPt | 26,6769 | 26,2625 | 30,95375 | 18,7028 | 29,8936 | 23,7827 | 23,36275 | 31,3221 | 16,4753 | 31,25035 |
| V1c | 0 | 7,256835 | 6,873385 | 8,55797 | 8,526375 | 3,591785 | 9,537195 | 5,4547 | 14,4859 | 13,66835 |
| V1fpLF | 8,55797 | 4,93912 | 14,4893 | 0 | 5,363955 | 9,092225 | 5,33578 | 17,7693 | 4,0818 | 10,046735 |
| V1fpUF | 8,526375 | 5,9522 | 5,224035 | 5,363955 | 0 | 10,07716 | 10,84655 | 5,640085 | 14,43095 | 5,303345 |
| V1pcLF | 7,256835 | 0 | 5,323915 | 4,93912 | 5,9522 | 7,64493 | 3,97704 | 19,81135 | 9,629475 | 21,92425 |
| V1pcUF | 6,873385 | 5,323915 | 0 | 14,4893 | 5,224035 | 11,25615 | 19,8721 | 5,300695 | 24,31155 | 9,261585 |
| V2c | 3,591785 | 7,64493 | 11,25615 | 9,092225 | 10,07716 | 0 | 4,6481 | 3,365095 | 12,2412 | 12,2763 |
| V2fpLF | 14,4859 | 9,629475 | 24,31155 | 4,0818 | 14,43095 | 12,2412 | 2,875845 | 21,8488 | 0 | 18,7366 |
| V2fpUF | 13,66835 | 21,92425 | 9,261585 | 10,046735 | 5,303345 | 12,2763 | 23,5191 | 3,642715 | 18,7366 | 0 |
| V2pcLF | 9,537195 | 3,97704 | 19,8721 | 5,33578 | 10,84655 | 4,6481 | 0 | 20,21345 | 2,875845 | 23,5191 |
| V2pcUF | 5,4547 | 19,81135 | 5,300695 | 17,7693 | 5,640085 | 3,365095 | 20,21345 | 0 | 21,8488 | 3,642715 |
| V3A | 15,94715 | 10,654675 | 22,0246 | 7,231115 | 18,47755 | 8,01093 | 7,366415 | 16,93035 | 6,71341 | 17,44755 |
| V3c | 6,113065 | 10,394505 | 12,60815 | 10,699115 | 10,66235 | 3,472995 | 13,51515 | 10,37276 | 13,8082 | 13,67885 |
| V3fpLF | 12,8216 | 7,80343 | 22,80575 | 5,63029 | 14,21465 | 11,4286 | 2,65526 | 19,0572 | 3,608145 | 19,9177 |
| V3fpUF | 13,6725 | 19,79865 | 13,2498 | 17,9593 | 9,8991 | 11,7947 | 19,80495 | 8,10327 | 20,74905 | 3,995545 |
| V3pcLF | 10,1217 | 8,05069 | 17,92415 | 7,031655 | 12,91335 | 4,22989 | 5,083705 | 15,67805 | 9,34155 | 18,67095 |
| V3pcUF | 9,869485 | 18,6406 | 10,21316 | 16,0064 | 12,44315 | 4,375125 | 18,07475 | 5,3402 | 19,09415 | 5,75036 |
| V4c | 10,3329 | 17,616 | 16,3685 | 13,54125 | 12,414 | 5,85574 | 17,067 | 13,85215 | 17,68495 | 21,34475 |
| V4LF | 10,45002 | 15,17785 | 19,2378 | 12,40855 | 15,99845 | 7,89634 | 13,9235 | 16,7162 | 13,67385 | 25,06085 |
| V4t | 14,6671 | 15,058 | 21,7575 | 13,9482 | 21,3795 | 12,2873 | 14,45955 | 17,85125 | 14,5846 | 21,49525 |
| V4UF | 13,66 | 19,21795 | 13,94905 | 16,9812 | 13,0762 | 9,311455 | 18,68975 | 10,29069 | 18,9721 | 7,67959 |
| V6 | 15,43075 | 16,5535 | 24,43505 | 6,25332 | 19,74845 | 15,4697 | 11,7039 | 20,9555 | 3,634475 | 19,9537 |
| V6A | 20,49 | 20,4993 | 26,8482 | 8,24409 | 24,1839 | 19,8406 | 17,65645 | 21,82125 | 6,744795 | 19,43965 |
| VIP | 25,9426 | 26,08005 | 30,0052 | 16,15575 | 30,0808 | 25,4781 | 26,83355 | 27,81845 | 12,507535 | 22,130275 |
