## Supplementary material for "Retinotopic organization of feedback projections in primate early visual cortex: implications for active vision": Table S4

**Table S4: Index of abbreviations**

| | |
|---|---|
| 1 | Somatosensory Area 1 |
| 2 | Somatosensory Area 2 |
| 3 | Somatosensory Area 3 |
| 5 | Somatosensory Area 5 |
| 9 | Area 9 |
| 10 | Area 10 |
| 11 | Area 11 |
| 12 | Area 12 |
| 13 | Area 13 |
| 14 | Area 14 |
| 23 | Area 23 |
| 25 | Area 25 |
| 31 | Area 31 |
| 32 | Area 32 |
| 44 | Area 44 |
| 24a | Area 24a |
| 24b | Area 24b |
| 24c | Area 24c |
| 24d | Area 24d |
| 29/30 | Area 29/30 |
| 45A | Area 45a |
| 45B | Area 45b |
| 46d | Area 46, Dorsal Part |
| 46v | Area 46, Ventral Part |
| 7A | Area 7A |
| 7B | Area 7B |
| 7m | Area 7m |
| 7op | Area 7op |
| 8B | Area 8B |
| 8L | Area 8, Lateral |
| 8m | Area 8, Medial |
| 8r | 8r |
| 9/46d | Area 9/46, Dorsal Part |
| 9/46v | Area 9/46, Ventral Part |
| AIP | Anterior Intraparietal Area |
| CORE | Core Region of The Auditory Cortex |
| DP | Dorsal Prelunate Area |
| ENTO | Entorhinal |
| F1 | Frontal Area F1 |
| F2 | Frontal Area F2 |
| F3 | Frontal Area F3 |

| | |
|---|---|
| F4 | Fronta Area F4 |
| F5 | Frontal Area F5 |
| F6 | Frontal Area F6 |
| F7 | Frontal Area F7 |
| FST | Fundus of Superior Temporal Area |
| Gu | Gustatory Cortex |
| INS | Insula |
| IPa | Area IPa |
| LB | Lateral Belt |
| LIP | Lateral Intraparietal Area |
| MB | Medial Belt |
| MIP | Medial Intraparietal Area |
| MST | Medial Superior Temporal Area |
| MTc | Middle Temporal Area Central |
| MTp | Middle Temporal Area Peripheral |
| OPAl | Orbital Periallocortex |
| OPro | Orbital Proisocortex |
| PBc | Parabelt, Caudal Part |
| PBr | Parabelt, Rostral Part |
| PERI | Perirhinal |
| PGa | Area PGa |
| Pi | Area Parainsula |
| PIP | Posterior Intraparietal Area |
| Pir | Piriform |
| POLE | Temporal Pole |
| Pro.St | Prostriata |
| ProM | Area ProM |
| SII | Secondary Somatosensory Area |
| STPc | Superior Temporal Polysensory, Caudal Part |
| STPi | Superior Temporal Polysensory, Intermediate Part |
| STPr | Superior Temporal Polysensory, Rostral Part |
| SIBI | Subiculum |
| TEa/ma | Anterior Area TEa/m |
| TEa/mp | Posterior Area TEa/m |
| TEad | Anterior Dorsal TE |
| TEav | Anterior Ventral TE |
| TEO | Area TEO |
| TEOm | Area TEOm |
| TEpd | Posterior Dorsal TE |
| TEpv | Posterior Ventral TE |
| TH/TF | Area TH/TF |
| TPt | Temporo-Parietal Area |
| V1c | Visual Area 1, Central |

| | |
|---|---|
| V1fpLF | Visual Area 1, Far Periphery Lower Field |
| V1fpUF | Visual Area 1, Far Periphery Upper Field |
| V1pcLF | Visual Area 1, Paracentral Lower Field |
| V1pcUF | Visual Area 1, Paracentral Upper Field |
| V2c | Visual Area 2, Central |
| V2fpLF | Visual Area 2, Far Periphery Lower Field |
| V2fpUF | Visual Area 2, Far Periphery Upper Field |
| V2pcLF | Visual Area 2, Paracentral Lower Field |
| V2pcUF | Visual Area 2, Paracentral Upper Field |
| V3A | Visual Area 3A |
| V3c | Visual Area 3, Central |
| V3fpLF | Visual Area 3, Far Periphery Lower Field |
| V3fpUF | Visual Area 3, Far Periphery Upper Field |
| V3pcLF | Visual Area 3, Paracentral Lower Field |
| V3pcUF | Visual Area 3, Paracentral Upper Field |
| V4c | Visual Area 4, Central |
| V4LF | Visual Area 4, Peripheral Lower Field |
| V4t | Transitional Visual Area 4 |
| V4UF | Visual Area 4, Peripheral Upper Field |
| V6 | Area V6 |
| V6A | Area V6a |
| VIP | Ventral Intraparietal Area |
